# Epithelial tumor cells utilize mast cell-derived histamine to regulate perineural invasion

**DOI:** 10.1101/2025.06.23.661147

**Authors:** Ankit Srivastava, Tomas Bencomo, Chaw-Ning Lee, Angela Mah, Jasmine Garcia, Lek Wei Seow, Isoline M. Donohue, Aiko J. Tan, Audrey Nguyen, Tiffany Jiang, Saurabh Gombar, Lilian Phu, Pankaj Dwivedi, Christopher M. Rose, Ryanne Brown, Carolyn S. Lee

## Abstract

Cancer dissemination by perineural invasion (PNI) is associated with poor outcomes in epithelial malignancies, yet the cellular interactions critical to this process are not fully defined. Single-cell transcriptomic analysis of cutaneous squamous cell carcinoma (cSCC) with PNI highlighted *KITLG*-*KIT* signaling activation between keratinocytes and mast cells. Additionally, we observed increased expression of *HRH1* on tumor cells, which encodes the H1 histamine receptor. We confirmed these changes in other cancers, including PNI-positive hepatobiliary and oral mucosal tumors. Histamine enhanced tumor invasion into nerve in organoids that recapitulate key features of PNI, an effect rescued by H1-antihistamines. Mechanistically, broad upregulation of matrix metalloproteinases (MMPs) through P38 activation underscored histamine-mediated PNI by enabling collagen degradation in the nerve sheath. Examination of multiple cancer types with PNI showed co-expression of *HRH1* and *MMP2* on tumor cells as well as type IV collagen loss in involved nerves. Retrospective analysis of >15,000 patient health records in two independent databases demonstrated that H1-antihistamine use improved overall survival and immunotherapy response in cancers that spread by PNI. Together, these data reveal a critical role for mast cells in PNI and present a new therapeutic strategy for targeting this process in epithelial cancers.

## Introduction

Perineural invasion (PNI) is a form of cancer dissemination distinct from lymphovascular metastasis in which tumor cells spread in, around, and through nerves^1^. The presence of PNI is frequently associated with neuropathic pain, tumor recurrence, distant metastasis, and poorer overall survival^2–4^. The incidence of PNI varies across different cancers and affects up to 80-100% of pancreatic cancers, head and neck squamous cell carcinomas (HNSC), and cholangiocarcinomas (CHOL)^2,4,5^. PNI is also observed in up to 14% of cutaneous squamous cell carcinoma (cSCC), where it is associated with worse outcomes and more common in poorly differentiated tumors^3,6,7^.

The mechanisms underlying tumor cell invasion into nerves have not been fully defined. Some studies suggest nerve and tumor cells communicate actively to achieve mutual benefit during PNI^2,8^. Several signaling molecules, including neurotrophins, chemokines, matrix metalloproteinase (MMPs), cytokines, surface receptors, and their corresponding ligands, are known to facilitate PNI^2,8^. Despite this understanding, there are currently no targeted treatments for PNI, and existing cancer therapies do not benefit PNI. Additionally, little is known about the influence of the tumor microenvironment on PNI promotion in cSCC. Key bottlenecks preventing a deeper understanding of the intercellular communication that enables PNI are a lack of single-cell transcriptomic data and limitations of current models used to study this phenomenon.

This study (Fig. 1a) analyzed the single-cell transcriptomes of five cSCC patients exhibiting extensive PNI. By also integrating data from low-risk, PNI-negative cSCC cases and normal skin, we characterized 92,371 distinct single-cell transcriptomes. Our analysis of ligand-receptor relationships revealed the enrichment of keratinocyte-mast cell interactions in PNI-positive tumors. We spatially validated this across various epithelial cancer types, including cSCC, HNSC, and CHOL. To characterize this interaction functionally, we developed a method for generating cancer organoids that recapitulate the features of PNI. Finally, we interrogated two independent electronic health record (EHR) databases to highlight the therapeutic potential of targeting mast cell-mediated orchestration of PNI in epithelial cancers.

**Fig. 1.**
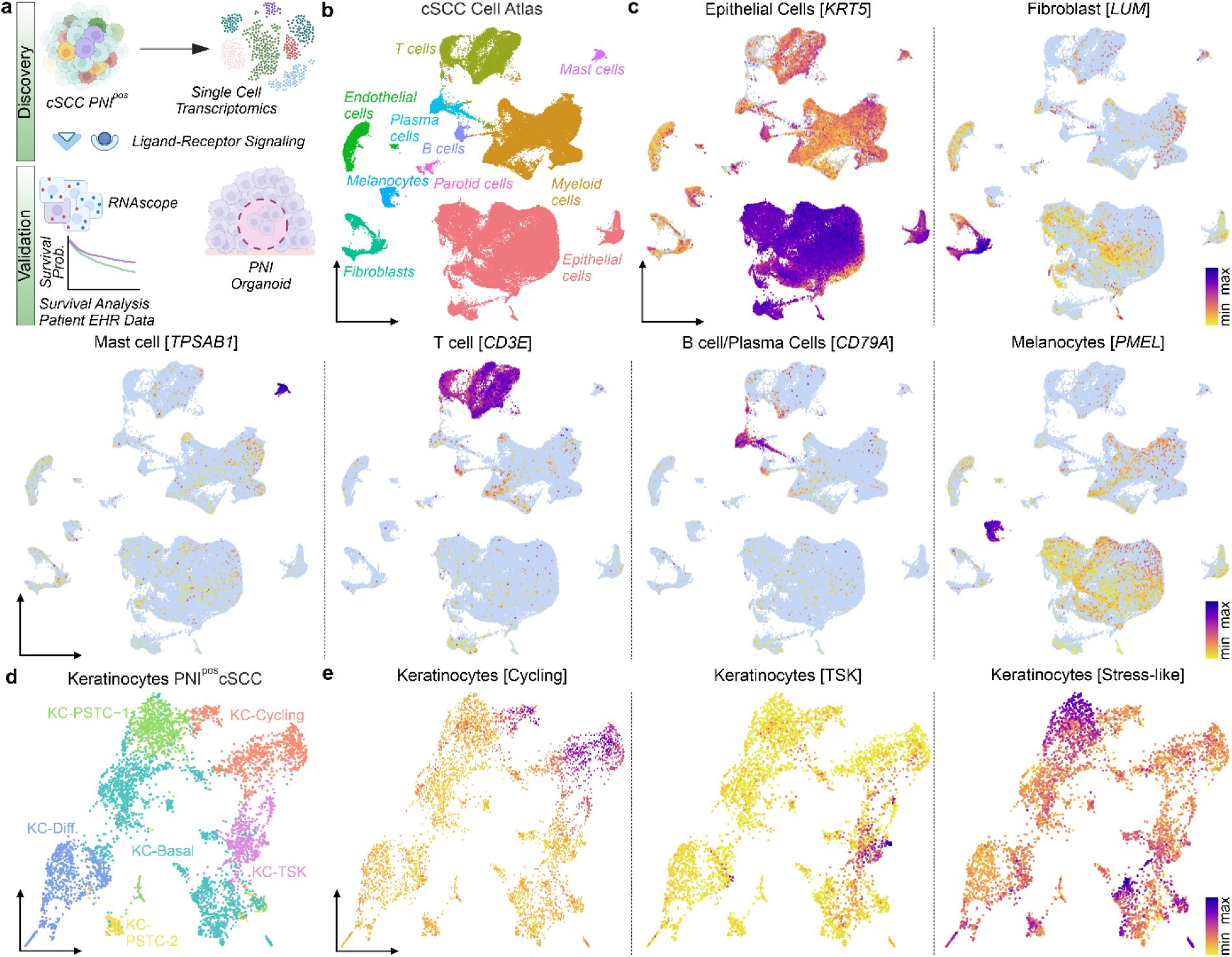
Epithelial cell milieu in PNI^pos^ cSCC. **a**, Study schema depicting the discovery and validation phase (created with BioRender.com). We applied scRNA-seq, RNAscope, novel PNI organoids, and survival analysis to define the tumor cellular niche and therapeutically actionable intercellular pathways of PNI^pos^ cancers. **b**, Uniform manifold approximation and projection (UMAP) of the cSCC cellular atlas, including PNI-positive and PNI-negative tumors compared to normal skin (n=92,371). **c**, UMAP view of epithelial cells, fibroblasts, mast cells, T cells, B/Plasma cells, and melanocytes. **d**, UMAP plot for keratinocytes in PNI^pos^ cSCC (n=4830). **e**, UMAP of different keratinocyte subpopulations involved in PNI^pos^ cSCC.

## Results

### PNI-specific tumor populations

We performed single-cell RNA-Seq (scRNA-Seq) on five cSCC with histopathologically confirmed PNI (PNI^pos^) (Supplementary Table 1). We then combined these single-cell transcriptomes with those obtained from PNI-negative (PNI^neg^) cSCC and clinically normal skin^9^ to create an expanded cellular atlas of cSCC. After evaluating quality control metrics, we retained 92,371 high-quality single-cell transcriptomes (Extended Data Fig. 1a, b) and subjected these to unsupervised clustering. This approach identified ten cell populations in cSCC, including epithelial cells, myeloid cells, endothelial cells, T cells, mast cells, and fibroblasts (Fig. 1b, c, and Extended Data Fig. 2a). Next, we applied Louvain clustering to highlight PNI-specific keratinocytes (Fig. 1d and Supplementary Table 2). A Differentiation-Progenitor-like (DvP) signature^10^ capable of scoring cells along this axis was used to characterize keratinocytes in PNI^pos^ cSCC. Consistent with prior reports, this analysis indicated a marked loss of differentiation programming in keratinocytes from PNI^pos^ cSCC compared to those in PNI^neg^ cSCC or normal skin (Extended Data Fig. 2b). We also detected keratinocyte subpopulations previously reported in PNI^neg^ cSCC^9^, including basal, cycling, and tumor-specific keratinocyte (TSK) subpopulations in PNI^pos^ cSCC (Fig. 1e and Extended Data Fig. 2c), with 70% more TSKs present in these tumors compared to PNI^neg^ cases. Additionally, we identified two distinct keratinocyte subpopulations without an equivalent in PNI^neg^ cSCC or normal skin, which we termed PNI-specific tumor cells (PSTC-1, 2) (Fig. 1d). PSTCs expressed genes associated with neural functions, and stress response activation (Fig. 1e and Supplementary Table 2), and resemble a stress signature found in HNSC^11^. Gene set enrichment analysis of 619 tumors from The Cancer Genome Atlas (TCGA) with annotated PNI status demonstrated enrichment of this stress-like signature in PNI^pos^ tumors originating in tissues other than skin compared to their PNI^neg^ counterparts. These findings revealed stress pathway activation in PNI^pos^ cSCC, a phenomenon also observed in other PNI^pos^ cancers, such as HNSC, colorectal cancer, and CHOL (Extended Data Fig. 2d).

### Cellular interactions in PNI^pos^ epithelial cancers

Next, we sought to identify cellular crosstalk in PNI^pos^ cSCC by analyzing cell-specific ligand-receptor expression patterns. We used CellChat^12^ to nominate distinct ligand-receptor pairs that may regulate PNI (Supplementary Table 3). Signaling pathways known to control neural functions, such as PTN, AGRN, and semaphorins, featured prominently among the activated pathways in PNI^pos^ cSCC (Supplementary Table 3). Among the top enriched pathways not previously associated with PNI in cSCC was KIT (Supplementary Table 3), with TSKs, cycling, and basal keratinocytes as well as fibroblasts were observed to express *KITLG* (stem cell factor, SCF), thereby activating KIT-positive mast cells (Fig. 2a). We detected minimal expression of *KITLG* in the corresponding keratinocyte populations or fibroblasts in PNI^neg^ cSCC (Fig. 2b). Analysis of TCGA tumors with known PNI status further confirmed increased *KITLG* expression and mast cell infiltration in PNI^pos^ tumors (Fig. 2c).

**Fig. 2.**
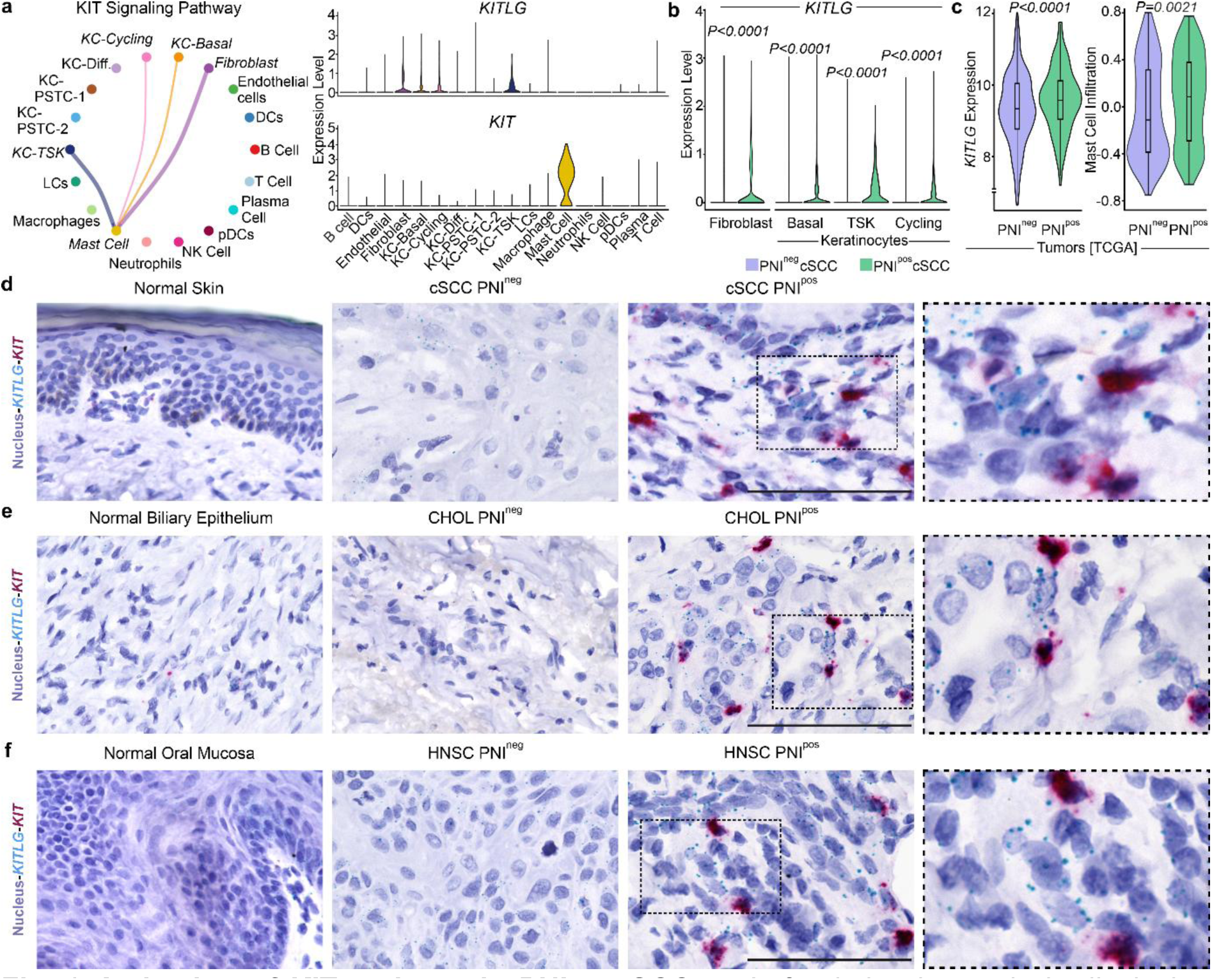
Activation of KIT pathway in PNI^pos^ cSCC. **a**, Left, circle plot analysis displaying cellular crosstalk via ligand (*KITLG*) expressed by keratinocytes/fibroblasts and receptor (*KIT*) present on mast cells in PNI^pos^ cSCC. Right, scRNA-seq data for *KITLG* and *KIT* expression in different cellular populations of PNI^pos^ cSCC. **b**, Expression of *KITLG* in keratinocytes and fibroblasts in cSCC patients. Wilcoxon rank sum test calculated the *P-value*. **c**, Enrichment of *KITLG* and mast cell infiltration gene signature in TCGA tumors with known PNI status. *P-value* calculated by linear regression after adjusting for cancer types. Error lines represent 1.5X interquartile range, and the box represents the interquartile range. **d**, RNAscope duplex *in situ* hybridization of *KITLG* (teal), *KIT* (red), and counterstained cell nuclei (blue) in normal skin, PNI^neg^, and PNI^pos^ cSCC. Scale bar=100μm, images are representative of n=5/group. **e**, RNAscope duplex *in situ* hybridization of *KITLG* (teal), *KIT* (red), and counterstained cell nuclei (blue) in normal biliary epithelium, PNI^neg^, and PNI^pos^ CHOL. Scale bar=100μm, images are representative of n=5/group. **f**, RNAscope duplex *in situ* hybridization of *KITLG* (teal), *KIT* (red), and counterstained cell nuclei (blue) in normal oral mucosa, PNI^neg^, and PNI^pos^ HNSC. Scale bar=100μm, images are representative of n=5/group.

To gain spatial context for KIT signaling in PNI, we employed an RNAscope duplex *in situ* hybridization system to visualize *KITLG*-*KIT* interactions. Both normal skin and PNI^neg^ cSCC displayed minimal expression of *KITLG* and *KIT*, while significant enrichment of *KITLG* and *KIT*-positive cells was detected in PNI^pos^ cSCC (Fig. 2d). Spatially, these cells were present at the invasion front, confirming the interaction between *KITLG*-expressing TSKs and *KIT*-positive mast cells (Fig. 2d). RNAscope analysis further demonstrated marked enrichment of *KITLG* and *KIT* signals in PNI^pos^ HNSC and CHOL compared to their PNI^neg^ counterparts and normal tissues (Fig. 2e, f). Consistent with PNI^pos^ cSCC, *KITLG* and *KIT*-expressing cells were found in close proximity in these PNI^pos^ tumors (Fig. 2e, f). Across all studied cancers, we observed *KITLG*-*KIT* interaction in and around nerves involved by PNI, suggesting that communication between tumor cells and mast cells is maintained throughout the dissemination process (Extended Data Fig. 3a-c). These data indicate that mast cells play a key role in orchestrating PNI in cSCC and other epithelial malignancies.

To investigate the regulation of *KITLG* in PNI, we performed copy number analysis in PNI^neg^ and PNI^pos^ cSCC using scRNA-Seq data. While the gene copy number of *KITLG* was unchanged in PNI^neg^ cSCC patients, 60% of PNI^pos^ cSCC patients displayed *KITLG* amplification (Extended Data Fig. 4a). We then utilized SCENIC to identify potential upstream regulators of *KITLG* in the remaining 40% of PNI^pos^ tumors. This analysis predicted E2F1, E2F7, IRF1, IRF2, and STAT1 as *KITLG* regulators within keratinocytes involved in PNI (Extended Data Fig.4b). Of these, the pathway activities of IRF1/2 and STAT1 were higher in PNI^pos^ cSCC lacking *KITLG* amplification, suggesting these transcription factors might control *KITLG* expression in these cases (Extended Data Fig. 4c). Examination of publicly available ChIP-seq data for keratinocytes using the Integrative Genome Viewer (IGV) subsequently revealed a STAT1 binding peak located downstream of the POL II binding site within the *KITLG* gene body; however, no binding peaks for IRF1/2 were present in this region (Extended Data Fig. 4d). To validate this experimentally, we inhibited STAT1 phosphorylation with fludarabine in primary keratinocytes and assessed *KITLG* expression. Inhibition of STAT1 phosphorylation reduced the expression of *KITLG*, along with *CXCL10*, a known STAT1-regulated gene (Extended Data Fig. 4e). These results suggest that keratinocytes in PNI^pos^ cSCC upregulate *KITLG* either through copy number amplification or transcriptional induction mediated by STAT1.

### Histamine-H1 axis facilitates PNI

To identify mast cell-derived factors that promote PNI in epithelial cancers, we began by examining the expression of various mast cell mediator receptors in keratinocyte populations in PNI^pos^ cSCC. Among these, the expression of *HRH1*, which encodes the H1 histamine receptor, was significantly upregulated in keratinocyte populations in PNI^pos^ cSCC compared to those from PNI^neg^ tumors, with other receptors either lowly expressed or unchanged in this context (Fig. 3a). Analysis of PNI-annotated TCGA tumors further validated higher expression of *HRH1* in PNI^pos^ tumors (Fig. 3b). In addition, we assessed histamine release in normal skin, biliary epithelium, and oral mucosa alongside PNI^neg^ and PNI^pos^ cSCC, CHOL, and HNSC using toluidine blue. We found no evidence of histamine staining in normal tissues and only a few histamine-positive cells in PNI^neg^ tumors (Fig. 3c-e and Extended Data Fig. 5a). By comparison, PNI^pos^ cSCC, CHOL, and HNSC exhibited a three to five-fold enrichment in histamine-positive mast cells (Fig. 3c-e and Extended Data Fig. 5a).

**Fig. 3.**
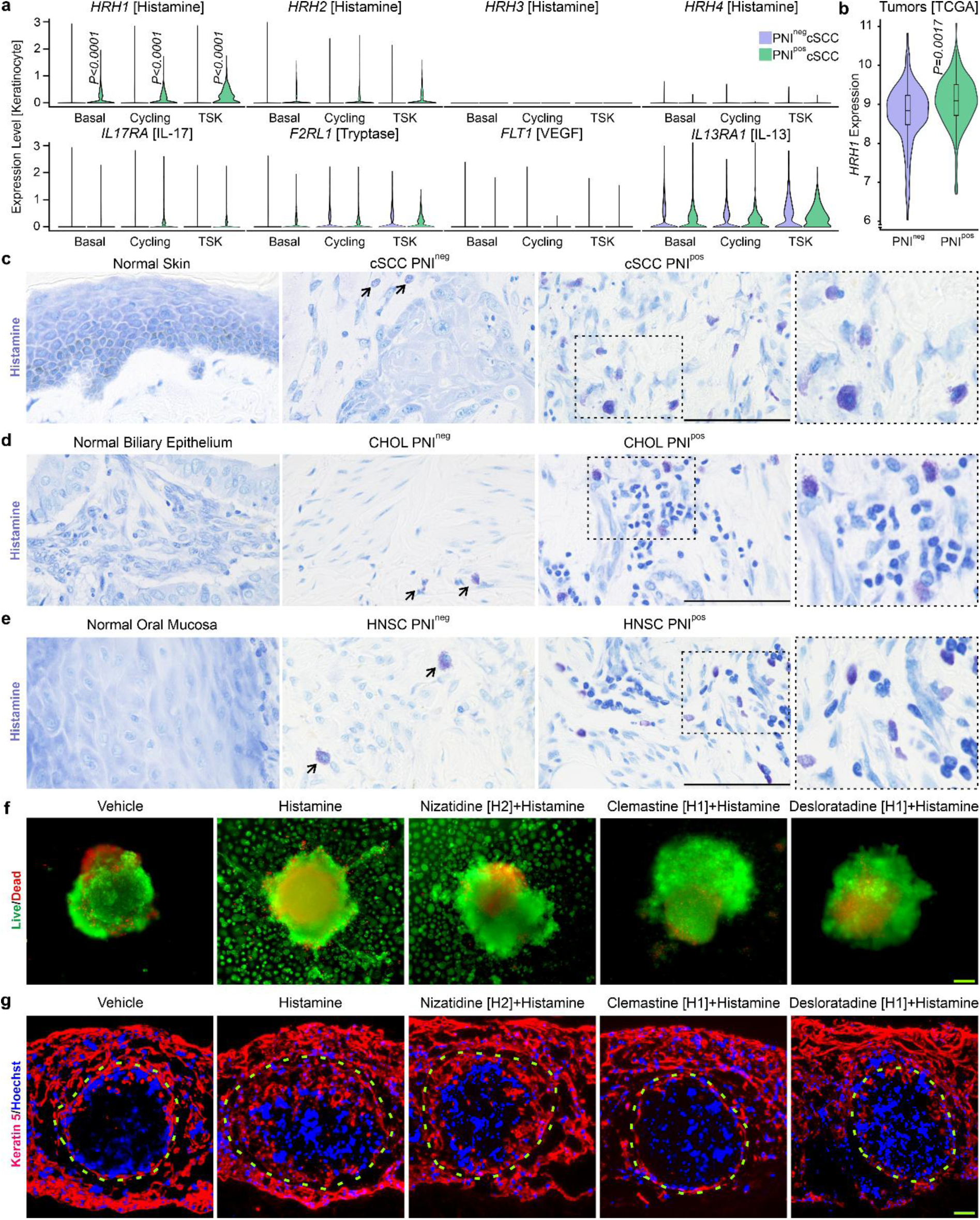
Histamine enhances PNI in an H1-dependent manner. **a**, Expression of various mast cell mediator receptors in keratinocyte populations of PNI^neg^ and PNI^pos^ cSCC assessed by scRNA-seq. Wilcoxon rank sum test calculated the *P-value*. **b**, Expression analysis of *HRH1* in PNI-annotated TCGA tumors. *P-value* calculated by linear regression after adjusting for cancer types. Error lines represent 1.5X interquartile range, and the box represents the interquartile range. **c**, Release of histamine by mast cells investigated with toluidine blue at low pH in normal skin, PNI^neg,^ and PNI^pos^ cSCC. Deep purple-histamine-positive cells, light blue-background, scale bar=100μm, images are representative of n=5/group. **d**, Release of histamine by mast cells in normal biliary epithelium, PNI^neg,^ and PNI^pos^ CHOL. Deep purple-histamine-positive cells, light blue-background, scale bar=100μm, images are representative of n=5/group. **e**, Release of histamine by mast cells in normal oral mucosa, PNI^neg,^ and PNI^pos^ HNSC. Deep purple-positive cells, light blue-background, scale bar=100μm, images are representative of n=5/group. **f**, *HRAS* and *CDK4*-transformed tumor spheroids were treated with either vehicle control, histamine, or a combination of histamine and H1-specific or H2-specific antihistamine. Tumor spheroid invasion was evaluated with live (green) and dead (red) staining and fluorescence microscopy. Scale bar=100μm. Results are representative of three independent experiments conducted in replicates. **g**, Immunofluorescence analysis of PNI organoids treated with either vehicle, histamine, or a combination of histamine and H1-specific or H2-specific antihistamine. The dotted line outlines the nerve sheath. Scale bar=100μm. Results are representative of three independent experiments conducted in replicates.

Next, we sought to define the oncogenic impacts of histamine-H1 interaction. We first determined a dose of histamine that upregulates H1 – a known consequence of histamine pathway activation^13^ – and leads to phosphorylation of JNK, a MAP kinase regulated by histamine^14^ (Extended Data Fig. 5b-d). We then evaluated the efficacy of two H1-antihistamines, clemastine fumarate and desloratadine, identifying doses that rescue histamine-mediated effects (Extended Data Fig. 5e). The inability of nizatidine, an H2-antihistamine, to counteract histamine-mediated *HRH1* induction provided further evidence of the importance of histamine-H1 interaction in keratinocytes (Extended Data Fig. 5e). Furthermore, we transformed human primary keratinocytes with oncogenic *HRAS* and cyclin-dependent kinase 4 (*CDK4*), a combination known to induce malignant conversion^15^. We generated tumor spheroids and cSCC organoids from these transformed primary keratinocytes, and visualized invasion using live/dead staining or immunofluorescence. Histamine treatment significantly enhanced invasion into the hydrogel matrix in spheroids and into the dermis in cSCC organoids (Fig. 3f and Extended Data Fig. 6). While treatment with an H2-antihistamine did not rescue these effects, two different H1-antihistamines effectively suppressed histamine-induced keratinocyte invasion in both of these models (Fig. 3f and Extended Data Fig. 6).

Organoid systems capable of studying PNI, particularly those that recapitulate its histologic features, are currently limited. To address this gap, we developed a novel PNI organoid system incorporating explanted nerves with *HRAS* and *CDK4*-transformed skin tissues. We assessed tumor infiltration into the nerve by treating these organoids with histamine in both the presence and absence of antihistamines. Histamine-treated PNI organoids demonstrated enhanced tumor cell invasion into nerves compared to those treated with vehicle control (Fig. 3g). Treatment with an H2-antihistamine did not alter this effect; however, both, clemastine fumarate and desloratadine effectively reduced histamine-mediated nerve invasion by keratinocytes to near baseline levels (Fig. 3g). These findings implicate histamine as a regulator of tumor dissemination along nerves and suggest that H1-antihistamines may have therapeutic potential in cancers that spread through PNI.

### Histamine activates P38 and MMP signaling in PNI

We used a phosphoproteomic approach to identify the downstream pathways activated by histamine-H1 interaction in keratinocytes. Protein phosphorylation was examined in lysates from human primary keratinocytes treated with either vehicle or histamine for 10 minutes to identify histamine-induced phosphorylated proteins using an FDR cutoff of ≤5% (Fig. 4a and Supplementary Table 4). Pathway enrichment analysis of this group consistently highlighted activation of the MAP kinase P38 and matrix metalloproteinase (MMP) pathways (Fig. 4b). Analysis of TCGA tumors with PNI status confirmed P38 and MMP pathway activation in PNI^pos^ tumors, with increased expression of multiple MMPs detected in PNI^pos^ compared to PNI^neg^ tumors (Fig. 4c). We then validated our phosphoproteomic findings by demonstrating phosphorylation of P38 following 10 minutes of histamine treatment in primary keratinocytes (Fig. 4d). Both H1-antihistamine treatment and pharmacologic inhibition of the P38 pathway were able to rescue this effect (Fig. 4d). Furthermore, we showed that histamine upregulated the expression of several MMPs in keratinocytes and both antihistamine treatment as well as P38 inhibition reversed this effect (Fig. 4e). Among the tested MMPs, the secreted levels of MMP2 and MMP9, quantified using ELISA, showed similar effects (Fig. 4e). *MMP2* and *MMP9* were also upregulated in cSCC organoids treated with histamine, and clemastine fumarate mitigated this effect (Extended Data Fig. 7a). Finally, we evaluated the impact of histamine-induced P38 phosphorylation on invasion using neoplastic spheroids, cSCC organoids, and PNI organoids. In all models, histamine significantly enhanced tumor invasion (Fig. 4f, g and Extended Data Fig. 7b). Inhibition of the P38 pathway in each context suppressed histamine-mediated tumor cell invasion into hydrogel, nerve, or dermis (Fig. 4f, g, and Extended Data Fig. 7b). Taken together, these results indicate histamine activates MMPs through the P38 pathway, and this activation can be pharmacologically blocked using either H1-antihistamines or inhibitors of P38.

**Fig. 4.**
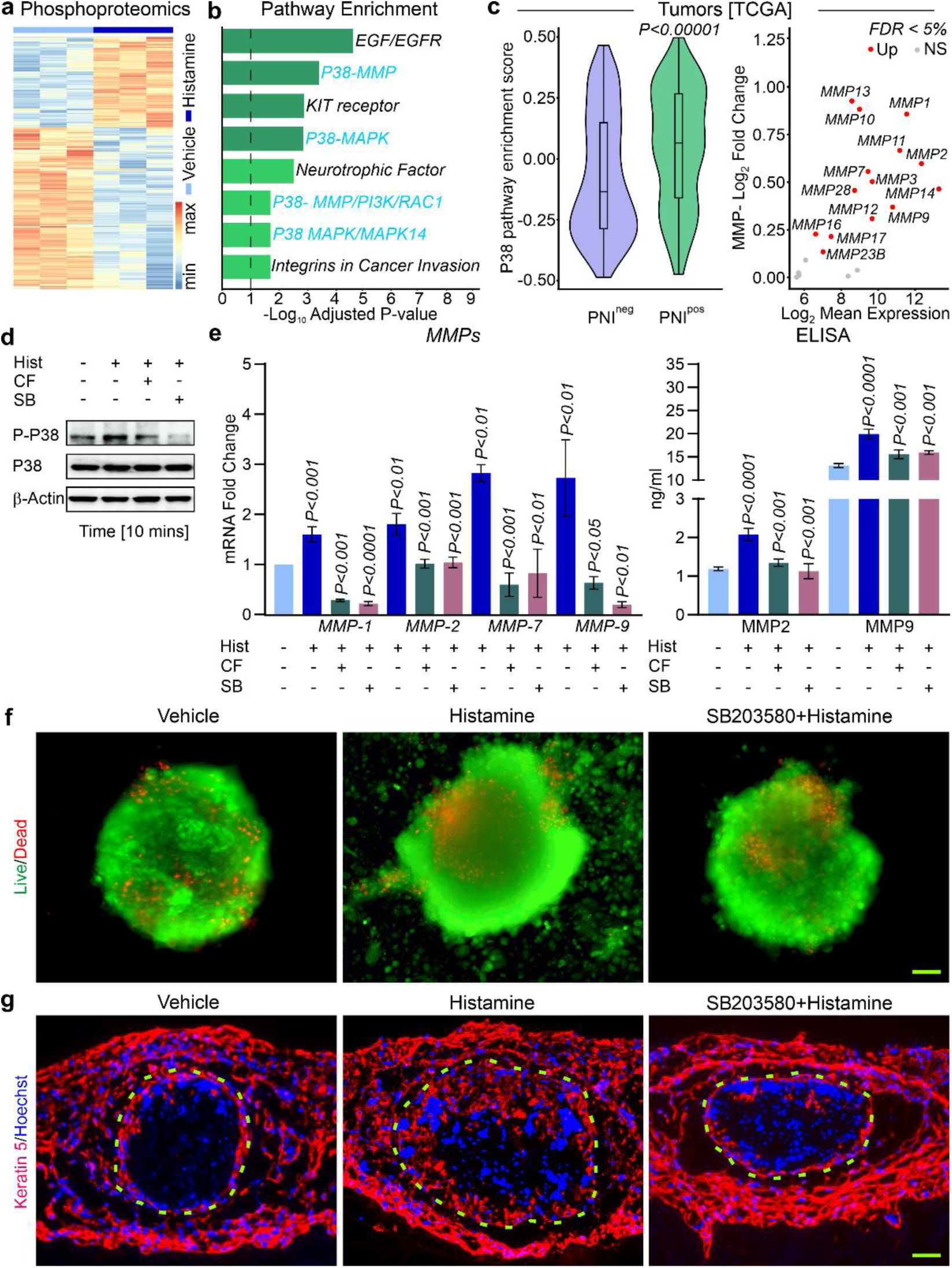
Histamine triggers MMP signaling by P38 activation. **a**, Phosphoproteomic analysis of keratinocytes treated with either vehicle control or histamine for 10 minutes by mass spectrometry. The heatmap highlights the phosphorylation sites altered by histamine compared to the control. **b**, Pathway enrichment analysis by EnrichR of histamine phosphorylated sites in primary keratinocytes. **c**, Left, P38 gene signature enrichment in PNI-annotated TCGA tumors. Right, induction of various MMPs in PNI-positive TCGA tumors. *P-value* calculated by linear regression after adjusting for cancer types. Error lines represent 1.5X interquartile range, and the box represents the interquartile range. **d**, Immunoblotting of P-P38, P38, and β-Actin in primary keratinocytes treated with vehicle control, hist-histamine, SB-SB203580 (P38 inhibitor) with histamine, or CF-clemastine fumarate (H1-antihistamine) with histamine. Results are representative of at least three independent experiments. **e**, Left, expression analysis of *MMPs* in primary keratinocytes treated with control, hist-histamine, SB-SB203580 (P38 inhibitor) with histamine, or CF-clemastine fumarate (H1-antihistamine) with histamine. Results are representative of at least three independent experiments conducted in replicates. Two-tailed Student’s t-tests calculated the *P-value* by comparing Hist to control, then CF and SB to the Hist group. Error bar-standard deviation. Right, ELISA for MMP2 and MMP9 in culture supernatants of primary keratinocytes in different treatment conditions. hist-histamine, SB-SB203580, CF-clemastine fumarate. Results are representative of multiple replicates in an experiment. Two-tailed Student’s t-tests calculated the *P-value* by comparing Hist to control, then CF and SB to the Hist group. Error bar-standard deviation. **f**, Live (green) and dead (red) staining in spheroids depicted tumor spheroid invasion upon the vehicle, histamine, or P38 inhibitor (SB203580) and histamine treatments. Scale bar=100μm, Results are representative of three independent experiments conducted in replicates. **g**, Immunofluorescence of Keratin 5 (red) displaying nerve invading oncogenic *HRAS* and *CDK4*-transformed primary keratinocytes in PNI organoids treated with histamine, or P38 inhibitor (SB203580) and histamine. The dotted line outlines the nerve sheath. Scale bar=100μm, Results are representative of three independent experiments conducted in replicates.

### Epithelial tumor cells induce MMPs to target collagen in the nerve sheath in PNI

To investigate the consequences of histamine-induced upregulation of *MMP2*/*9*, we examined the expression of these *MMPs* and *HRH1* via RNAscope in PNI^neg^ and PNI^pos^ cSCC, CHOL, and HNSC. We occasionally observed *HRH1* or *MMP2*-expressing cells in normal skin, biliary epithelium, oral mucosa, and PNI^neg^ tumors (Fig. 5a-c). However, in PNI^pos^ tumors, we detected numerous epithelial cells with strong co-expression of both *HRH1* and *MMP2* (Fig. 5a-c). These cells were present at the tumor invasion front and areas of PNI and were seen invading deep into the nerves (Fig. 5a-c). In addition, we identified an enrichment of *MMP9*-expressing tumor cells in PNI^pos^ tumors compared to their PNI^neg^ counterparts (Extended Data Fig. 8a-c). However, no marked enrichment of *MMP9*-positive cells was observed at the involved nerves by PNI. These data suggest that histamine-mediated activation of MMPs through *HRH1* facilitates tumor cell invasion into the nerve.

**Fig. 5.**
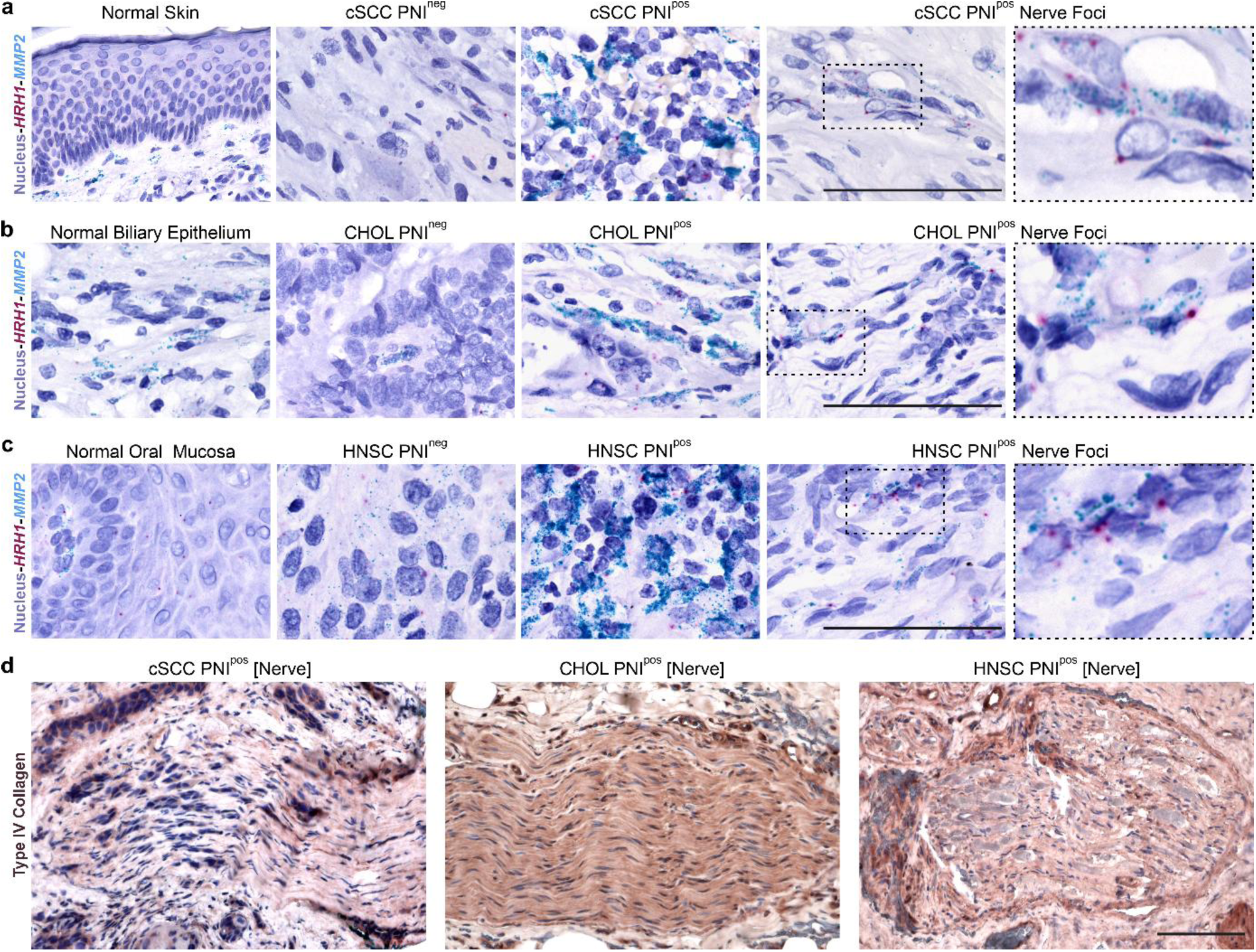
PNI-positive tumor cells co-express *HRH1*-*MMP2* to target type IV collagen. **a**, RNAscope duplex *in situ* hybridization of *MMP2* (teal), *HRH1* (red), and counterstained cell nuclei (blue) in normal skin, PNI^neg^, and PNI^pos^ cSCC. Scale bar=100μm, images are representative of n=5/group. **b**, RNAscope duplex *in situ* hybridization of *MMP2* (teal), *HRH1* (red), and counterstained cell nuclei (blue) in normal biliary epithelium, PNI^neg^, and PNI^pos^ CHOL. Scale bar=100μm, images are representative of n=5/group. **c**, RNAscope duplex *in situ* hybridization of *MMP2* (teal), *HRH1* (red), and counterstained cell nuclei (blue) in normal oral mucosa, PNI^neg^, and PNI^pos^ HNSC. Scale bar=100μm, images are representative of n=5/group. **d**, Immunohistochemistry for type IV collagen in PNI-involved nerves of PNI^pos^ cSCC, CHOL, and HNSC. Scale bar=100μm, images are representative of n=5/group.

Type IV collagen, a principal component of the basement membrane and nerve sheath, is a known substrate for MMP2/9^16^. Therefore, we performed immunohistochemistry (IHC) for Type IV collagen in PNI^pos^ cases of cSCC, CHOL, and HNSC, and examined both nerves with tumor cell infiltration and those without PNI involvement. Uninvolved nerves in these tumors displayed a strong expression of Type IV collagen around an intact nerve sheath (Extended Data Fig. 9a). By contrast, nerves affected by PNI demonstrated a near complete loss of Type IV collagen, accompanied by nerve sheath degradation and tumor cell invasion across all studied cancers (Fig. 5d). We also detected a similar loss of Type IV collagen staining consistent with nerve sheath degradation in histamine-treated PNI organoids, with H1-antihistamine treatment protecting against this degradation (Extended Data Fig. 9b). Altogether, these results support a model wherein *MMP*s are upregulated in tumor cells upon binding of mast cell-derived histamine to the H1 receptor, which confers their ability for collagenolysis and facilitates tumor invasion into the nerves.

### H1-antihistamines improve epithelial cancer patient survival

Given the longstanding accessibility and availability of H1-antihistamines, we next leveraged EHR data to determine whether their use might improve outcomes in cancers that spread by PNI. To explore this possibility, we performed a retrospective analysis using the global EHR database TriNetX to compare survival probabilities in patients with cSCC, HNSC, and CHOL who took H1-antihistamines or H2-blockers. Our study included 1594 patients who took H1-antihistamines, and 5472 patients who took H2-antihistamines for unrelated conditions or secondary symptoms of their malignancy (Supplementary Table 5). We then employed propensity score matching (PSM) to balance treatment groups, resulting in a final cohort of cSCC, CHOL, and HNSC patients for Kaplan-Meier survival analysis (Supplementary Table 5). Individuals with these cancers who took H1-antihistamines demonstrated improved survival compared to those who received H2-antihistamines [*P<0.0001*, adjusted HR 0.66(0.55, 0.78)] (Fig. 6a). Given the increasing use of immunotherapy to treat these malignancies, we next used TriNetX data to examine the effect of H1-antihistamines on immunotherapy response. PSM was performed to define a cohort of cSCC, HNSC, and CHOL patients on immunotherapy who received either H1-antihistamines (n=267) or H2-blockers (n=287) (Supplementary Table 6). The use of H1-antihistamines again extended overall survival compared to H2-blockers in immunotherapy patients, [*P=0.019*, adjusted HR 0.70(0.51, 0.94)] (Extended Data Fig. 10a).

**Fig. 6.**
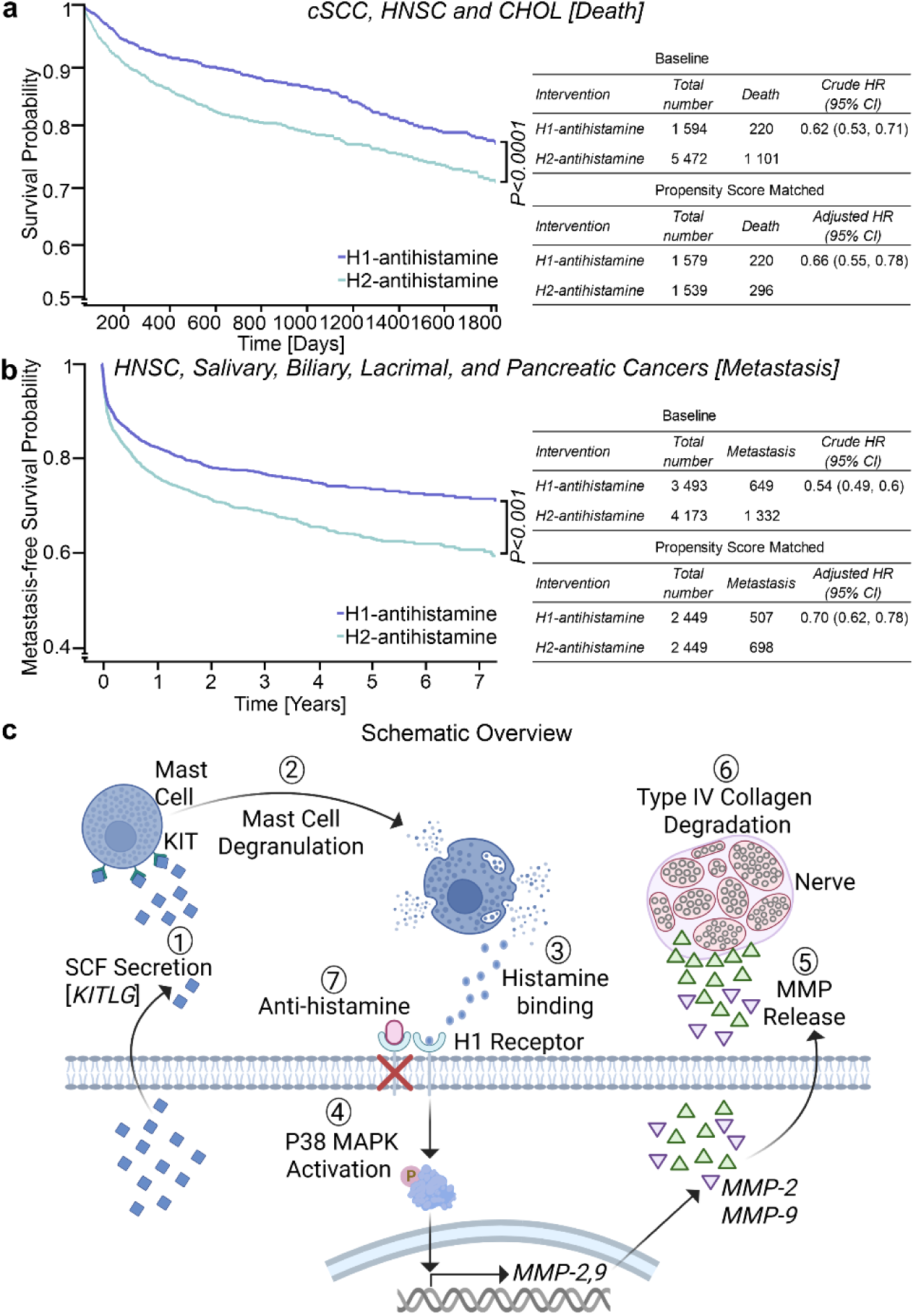
H1-antihistamines enhance patient survival. **a**, Left, Kaplan-Meier survival curve for cSCC, HNSC, and CHOL patients in the global EHR database TriNetX comparing H1-or H2-antihistamine use. Two-tailed Student’s t-tests calculated the *P-value*. Right, crude (baseline)-and propensity-score-matched analysis for hazard ratios. The Cox proportional hazard model calculated hazard ratios. **b**, Left, Kaplan-Meier metastasis-free survival curve for HNSC, Salivary, Lacrimal, and pancreatic cancer patients in the US national EHR database-Atropos comparing H1- or H2-antihistamine use. Two-tailed Student’s t-tests calculated the *P-value*. Right, crude (baseline)- and propensity-score-matched analysis for hazard ratios. The Cox proportional hazard model calculated hazard ratios. **c**, Mechanistic summary of the key conclusions (created with BioRender).

To determine whether the protective effect of H1-antihistamines in these three cancer types prone to PNI extends to a broader range of malignancies, we examined a second independent cohort of HNSC, salivary, biliary, lacrimal, and pancreatic cancers. Using EHR data from Atropos Health, a U.S. national database, we obtained a cohort of 2449 patients treated with H1- or H2-antihistamines after applying PSM (Supplementary Table 7). Patients who used H1-antihistamine exhibited higher metastasis-free survival than those who received H2-antihistamines [*P<0.0001*, adjusted HR 0.70(0.62, 0.78)] (Fig. 6b). Furthermore, treatment with H1-antihistamines was associated with improved overall survival [*P=0.05*, adjusted HR 0.86(0.74, 1)] (Extended Data Fig.10b). Collectively, these analyses suggest H1-antihistamines enhance survival in the examined malignancies by reducing cancer dissemination and are consistent with our observed experimental results demonstrating PNI suppression with H1 receptor blockade.

## Discussion

There is an unmet need for therapeutic strategies that address cancer metastasis by PNI. Our study revealed a previously unrecognized interaction between mast cells and keratinocytes in cSCC with PNI that is also present in other epithelial cancers, including HNSC and CHOL. We present a targetable disease model wherein epithelial tumor cells co-opt mast cells to acquire the ability to invade into nerves. Following SCF (*KITLG*)-mediated degranulation of mast cells, histamine-H1 receptor (*HRH1*) interaction promotes MMP secretion by tumor cells via P38 activation and ultimately enables PNI by enhancing collagenolysis in the nerve sheath (Fig. 6c).

An understanding of PNI at the single-cell level is in its early stages^17–20^, and this process remains unexplored in cSCC. Here, we provide a high-resolution view of the cell states and intercellular interactions in cSCC-associated PNI that confirms previously reported cell populations in cSCC^9^ and highlights differences in cellular composition compared to PNI^neg^ cSCC. Our results also identify new tumor cell populations in PNI^pos^ cSCC that resemble those implicated in the tumor cell dissemination of other cancers via stress response regulation. Future studies are needed to elucidate the functional roles of these stress-related keratinocyte populations in PNI.

By focusing on ligand-receptor relationships enriched in PNI^pos^ cSCC, we identified the recruitment of KIT-positive mast cells through the SCF secretion by keratinocytes in PNI^pos^ tumors, which was upregulated by *KITLG* gene amplification or transcriptional activation by STAT1. While early mast cell recruitment can enhance anti-tumor immunity in certain malignancies, mast cell presence has also been linked to tumor progression and resistance to immunotherapy^21–24^. We noted a significant influx of mast cells in PNI^pos^ cSCC, HNSC, and CHOL compared to their PNI^neg^ counterparts; notably, previous studies reporting increased mast cells in these malignancies did not examine their downstream signaling consequences or functional impacts^25–28^. Although mast cells typically mediate allergic response in an IgE-dependent manner, SCF can also potentiate and facilitate IgE-independent mast cell degranulation^29^. Our results confirm increased histamine release in PNI^pos^ tumors and demonstrate upregulation of the histamine receptor H1 in PNI^pos^ keratinocyte populations compared to other mast cell mediator receptors. Both tumorigenic and tumor suppressive roles have been ascribed to histamine, and these effects appear to be regulated by binding to specific histamine receptors^30^. Our data support a mechanism wherein high *HRH1* (H1) expression enables tumor cells to respond to mast cell-derived histamine, which is critical for PNI. Interestingly, increased *HRH1* expression on macrophages has been associated with T-cell dysfunction and immunotherapy resistance in melanoma and lung cancer^31^. We did not detect *HRH1* expression in cSCC-associated macrophages, suggesting PNI promotion by histamine-H1 receptor interaction is distinct from its effects on macrophages.

Several studies have used various co-culture approaches and a limited number of in vivo models to investigate PNI^32^. The three-dimensional architecture of our PNI organoids closely mimics the human condition, providing an advantage over two-dimensional in vitro and ex vivo models. While sciatic nerve injection has been widely used to study PNI in vivo, it is cell line-dependent and bypasses early interactions that may be necessary for PNI progression^33^. By comparison, tumor inoculation into the mouse whisker pad allows PNI to unfold spontaneously but occurs at low frequencies^34^. We developed a novel PNI model that robustly and reproducibly recapitulates the features of PNI using human primary cells. This approach faithfully captures early features of PNI, including invasion into the extracellular matrix (ECM) and degradation of the nerve sheath.

Using PNI organoids and complementary approaches, we showed that histamine enhanced cancer cell invasion, including into explanted nerves. At the signaling level, histamine upregulated MMPs in keratinocytes via P38 activation. Treatment with commercially available H1-antihistamines or a pharmacologic inhibitor of P38 prevented these effects, while targeting the lowly expressed H2 receptor on keratinocytes did not affect neoplastic invasion or nerve infiltration. These studies suggest that PNI is regulated specifically by histamine binding to H1 receptors and demonstrate the pharmacological modulation of this process. Although previous research has indicated histamine may mediate MMP expression and P38 activation in various cell types^35,36^. To our knowledge, the role of histamine-H1-P38-MMP2 signaling has not been fully characterized in keratinocytes or PNI settings. Our findings show enrichment of *HRH1*-*MMP2* co-expressing tumor cells and loss of its target type IV collagen in the nerve sheath in PNI^pos^ cSCC, HNSC, and CHOL. Notably, these cells were present at the invasion front and deep within the nerves involved by PNI, suggesting their critical role in PNI promotion through nerve sheath degradation.

In patients with melanoma and lung cancer, histamine has been shown to facilitate resistance to anti-PD-1/PDL-1 treatment, and the use of H1-antihistamines is associated with improved survival in these contexts^31^. We leveraged two independent EHR databases and retrospectively analyzed data from patients with cancers that frequently disseminate by PNI, including cSCC, CHOL, and HNSC, as well as salivary, biliary, lacrimal, and pancreatic cancers. Use of H1-antihistamines improved overall survival and metastasis-free survival in these patients compared to H2-antihistamines. While mast cell reduction with imatinib^37^ or prevention of mast cell degranulation with stabilizing agents such as cromolyn^38^ may have therapeutic potential in PNI, these strategies may also impede early immune cell recruitment, as other mast cell mediators are required for anti-tumor immunity. Our results highlight the strong therapeutic and protective impacts of H1-antihistamines across PNI^pos^ cancers. Given their well-established safety profile and ready availability, H1-antihistamines should be explored as adjunct therapies to standard cancer treatments.

Our study is not without limitations. The dissociation protocol utilized was optimized for skin and differs from those used for peripheral nerves^39^, which is likely to account for the absence of nerve and nerve-resident cells in our single-cell transcriptome dataset. We used toluidine blue to demonstrate histamine release, and we recognize that it also co-stains heparin, another mast cell mediator. Lastly, due to a lack of PNI annotation in EHR data, our survival analyses may include a mix of both PNI^pos^ and PNI^neg^ tumors. However, except cSCC, the other cancers in this analysis, which comprise most of the cohort, exhibit high frequencies of PNI ranging from 70%-100%^2,4,5^.

In conclusion, we introduce a therapeutic framework in which mast cells orchestrate tumor spread by PNI in epithelial cancers. By engaging mast cells and co-opting histamine-H1 allergy response pathways, tumor cells gain the capacity to infiltrate and invade nerves. Our analysis of patient data underscored the promising role of H1-antihistamines in treating cancers disseminated by PNI. Furthermore, we confirmed our findings concerning PNI in cSCC across other epithelial cancer types, suggesting that H1-antihistamines may have broad therapeutic applicability. Finally, we developed an organoid model of human PNI. In light of the FDA’s recent guidance on the expanded use of organoids in preclinical testing, we anticipate that adapting this model to other cancer types may streamline drug screening for PNI-associated malignancies.

**Extended Fig. 1.**
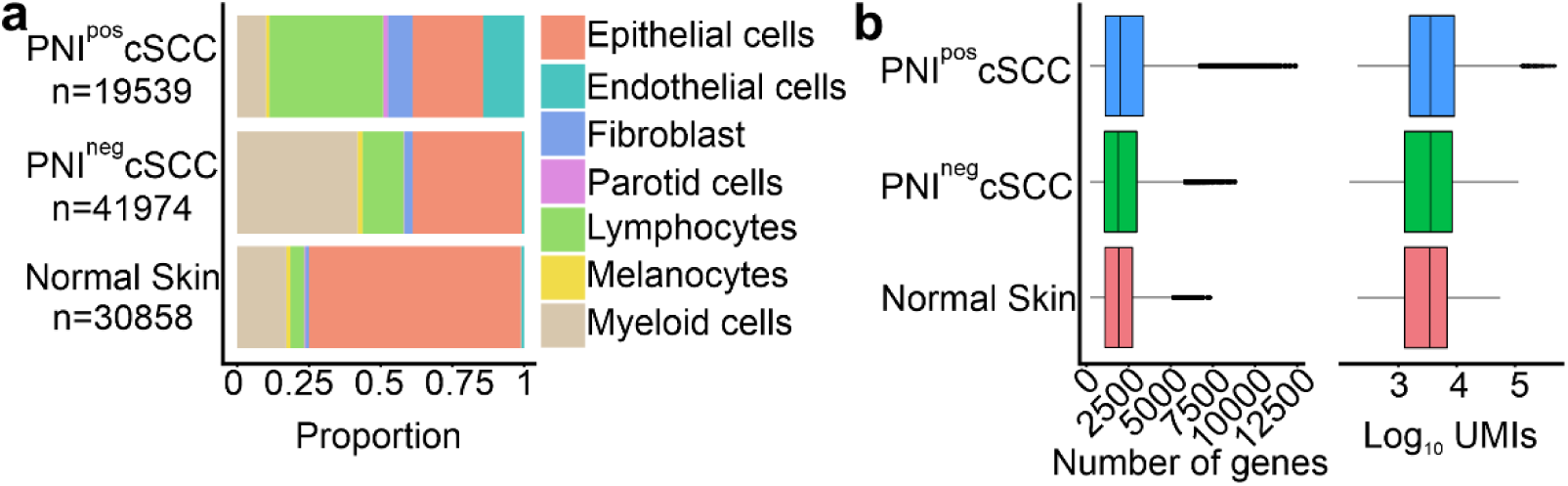
Cellular proportion and gene reads. **a**, Cellular composition of normal skin, PNI^neg,^ and PNI^pos^ cSCC. **b**, Gene read counts and unique molecular identifiers (UMIs) in normal skin, PNI^neg,^ and PNI^pos^ cSCC.

**Extended Fig. 2.**
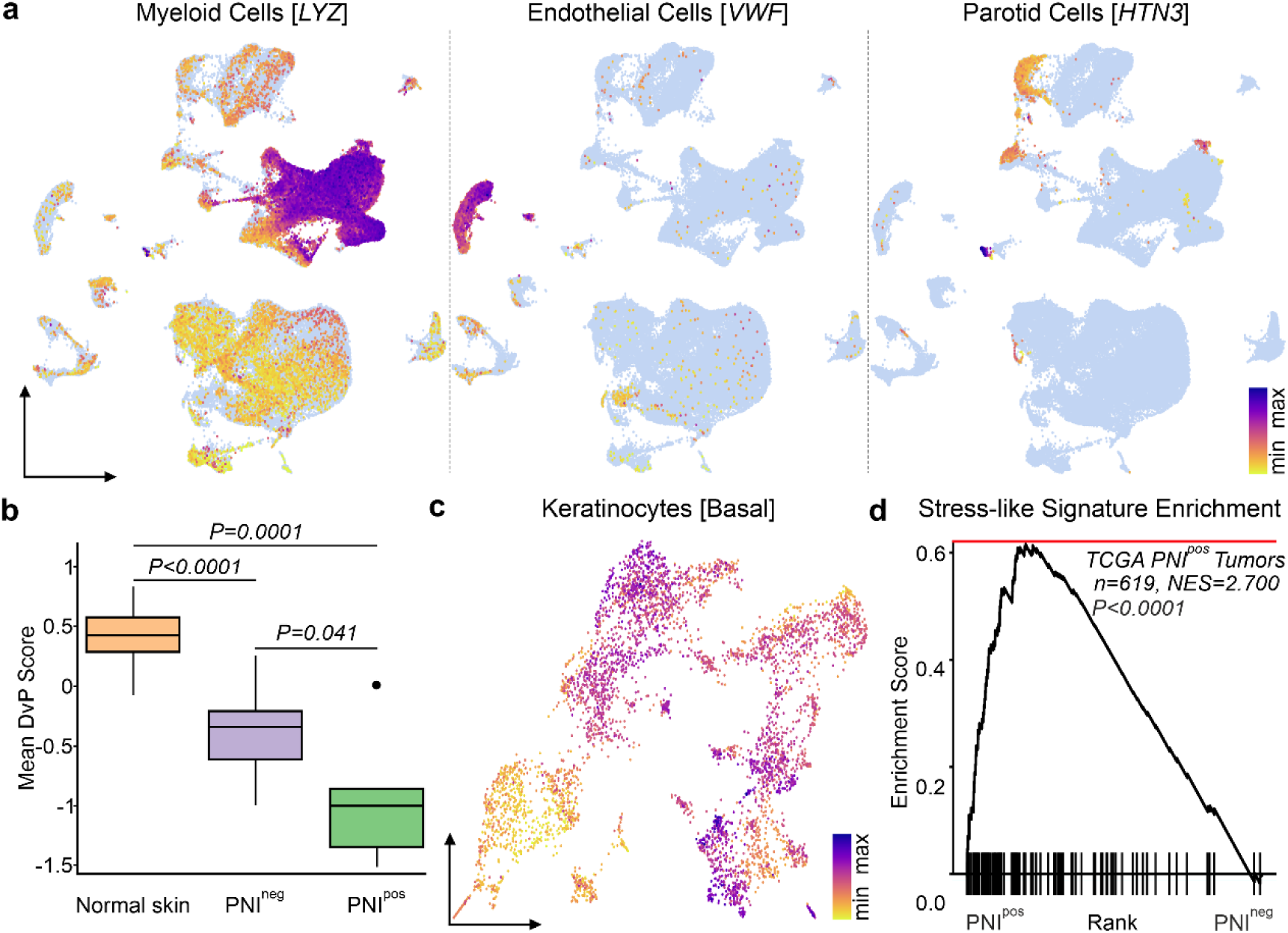
Cell populations in cSCC and their characteristics. **a**, UMAP view of myeloid, endothelial, and parotid cells in cSCC. **b**, DvP score depicting the loss of differentiation programming in keratinocytes in PNI^pos^ cSCC. *P-value* was calculated by mixed effect model adjusting for patient ID with the Satterthwaite method. **c**, UMAP of basal keratinocytes in PNI^pos^ cSCC. **d**, Gene set enrichment analysis (GSEA) for a stress-like signature in TCGA tumors with known PNI status. *P-value* was calculated by permutation test.

**Extended Fig. 3.**
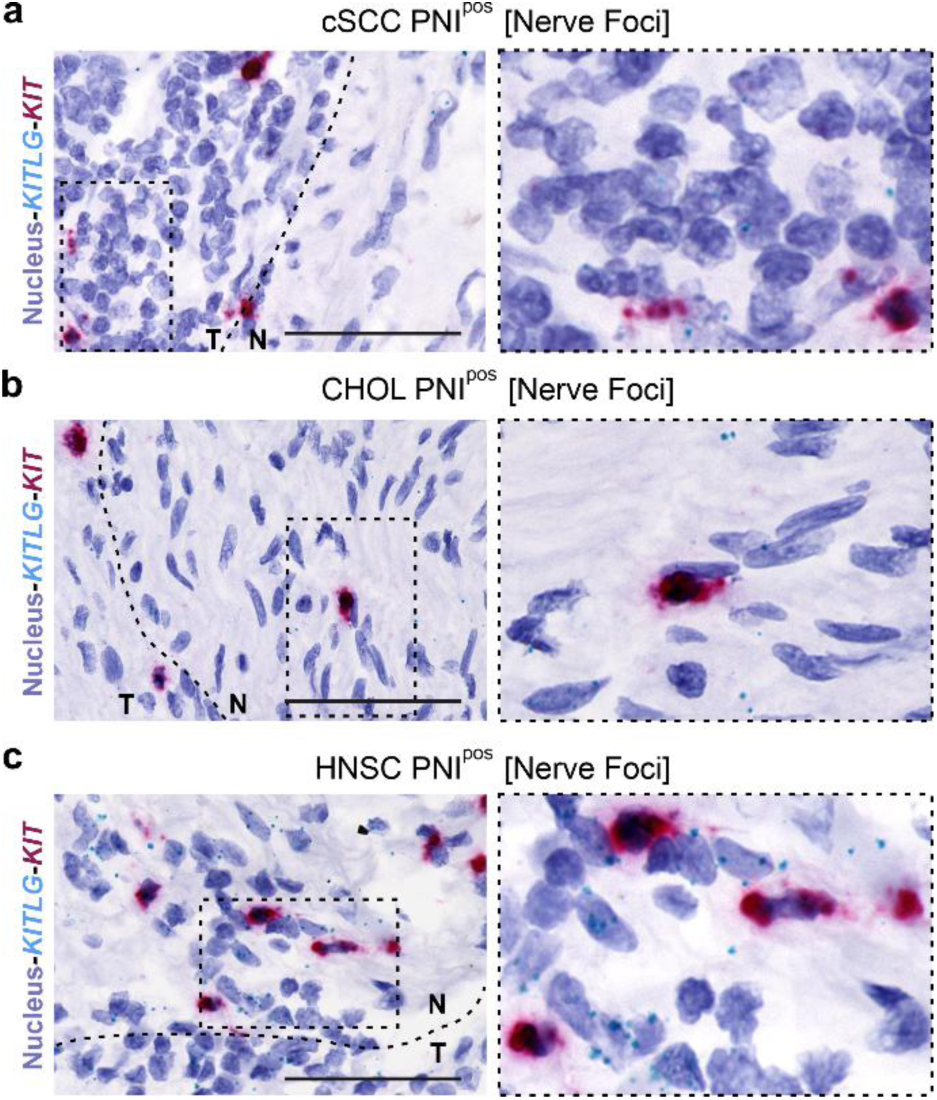
Activation of KIT signaling at tumor-nerve foci. **a-c**, RNAscope duplex ISH of *KITLG* (teal), *KIT* (red), and counterstained cell nuclei (blue) at the tumor-nerve intersection in PNI^pos^ cSCC, CHOL, and HNSC. N=nerve, T=tumor, scale bar=100μm, images are representative of n=5/group.

**Extended Fig. 4.**
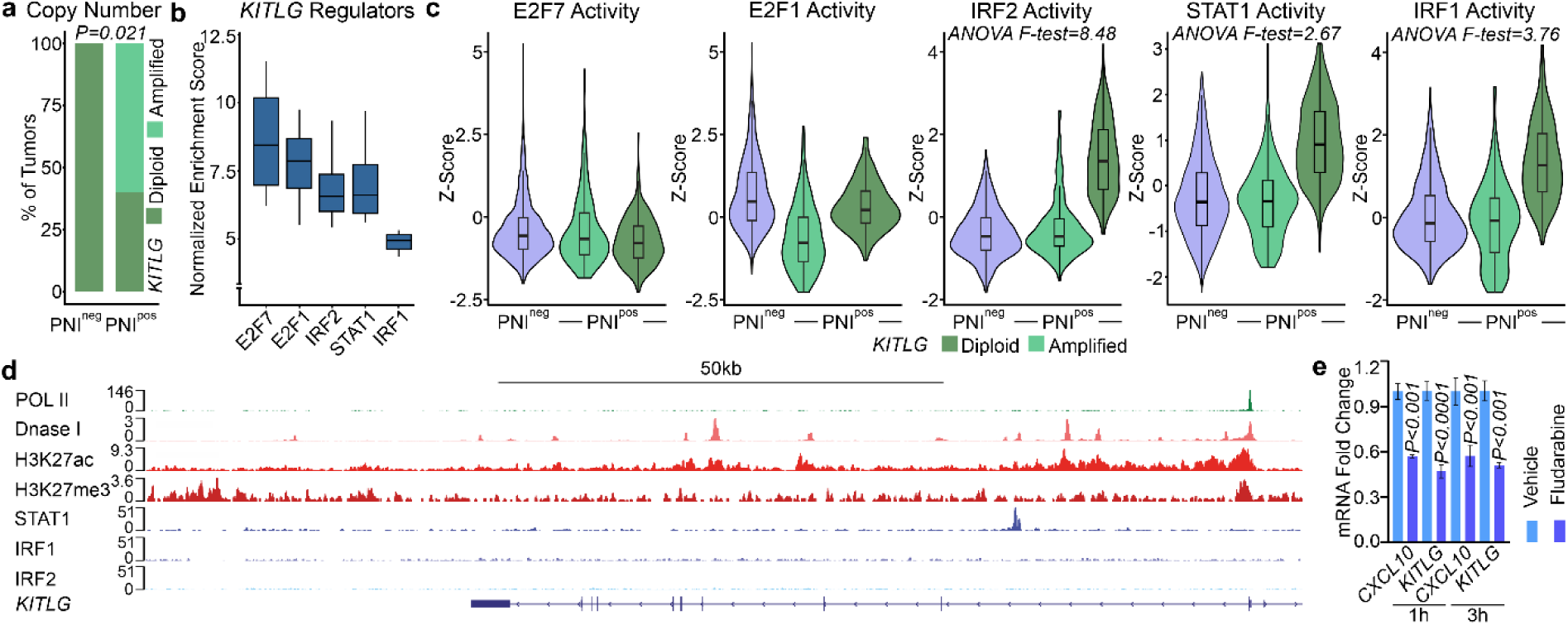
STAT1 regulates *KITLG* in keratinocytes. **a**, Copy number analysis of *KITLG* in primary human cSCC. Fisher’s exact test calculated *P-value*. **b**, Single-cell regulatory network inference and clustering (SCENIC)-based prediction of upstream regulators of *KITLG* in keratinocytes. Error lines represent 1.5X interquartile range, and the box represents the interquartile range. **c**, Pathway activity scores of predicted *KITLG* regulators in PNI^neg^ cSCC and *KITLG*-amplified and *KITLG*-diploid PNI^pos^ tumors. Error lines represent 1.5X interquartile range, and the box represents the interquartile range. **d**, IGV view of *KITLG* gene body with open chromatin, transcription regulators, and transcription factor binding analyzed using publicly available ChIP-seq data. **e**, Expression analysis of *CXCL10* and *KITLG* upon STAT1 phosphorylation inhibition with fludarabine in keratinocytes. Conducted with replicates and repeated four independent times. Two-tailed Student’s t-tests calculated the *P-value*, error bar-standard deviation.

**Extended Fig. 5.**
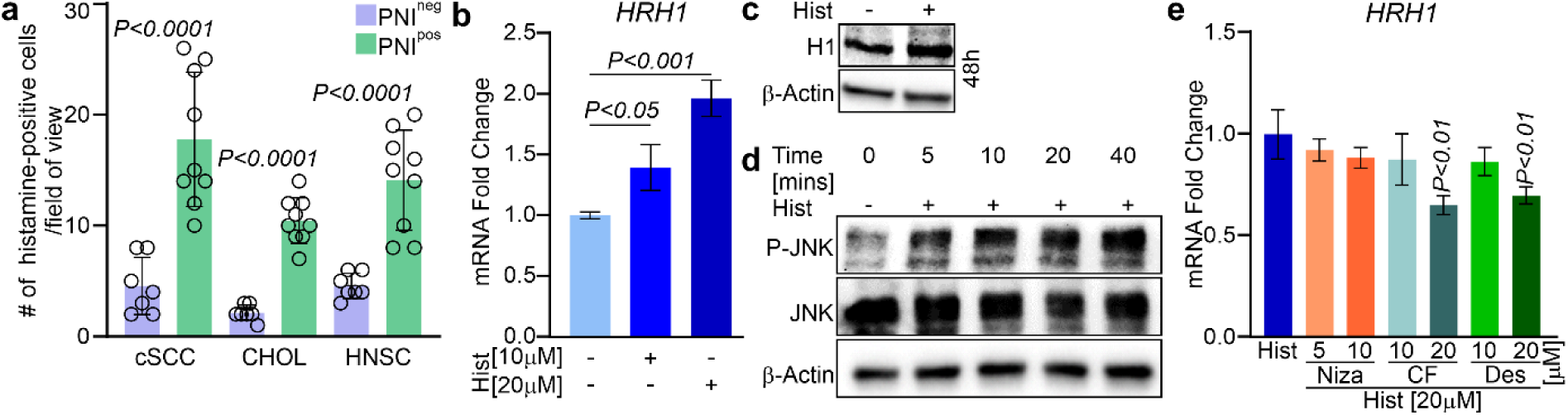
Histamine presence in cancer and dose analysis of histamine and antihistamine response in keratinocytes. **a**, Quantification of histamine-positive mast cells using toluidine blue in PNI^neg^ and PNI^pos^ cSCC, CHOL, and HNSC. A three-five field of view was used to quantify histamine-positive cells from images representing n=5/group. **b-c**, The Dose response of histamine treatments in keratinocytes was evaluated with *HRH1* expression via qRT-PCR and immunoblotting. Conducted with replicates and repeated three independent times. Two-tailed Student’s t-tests calculated the *P-value*, error bar-standard deviation, Hist-histamine. **d**, Immunoblotting for P-JNK, JNK, and β-Actin upon histamine treatment in primary keratinocytes. Repeated three times. **e**, qRT-PCR analysis of *HRH1* in different doses of H1-or H2-antihistamines following histamine treatment in keratinocytes compared to histamine group. Conducted with replicates and repeated three independent times. Two-tailed Student’s t-tests calculated the *P-value*, error bar-standard deviation, Hist-histamine, Niza-Nizatidine, CF-clemastine fumarate, Des-Desloratadine.

**Extended Fig. 6.**
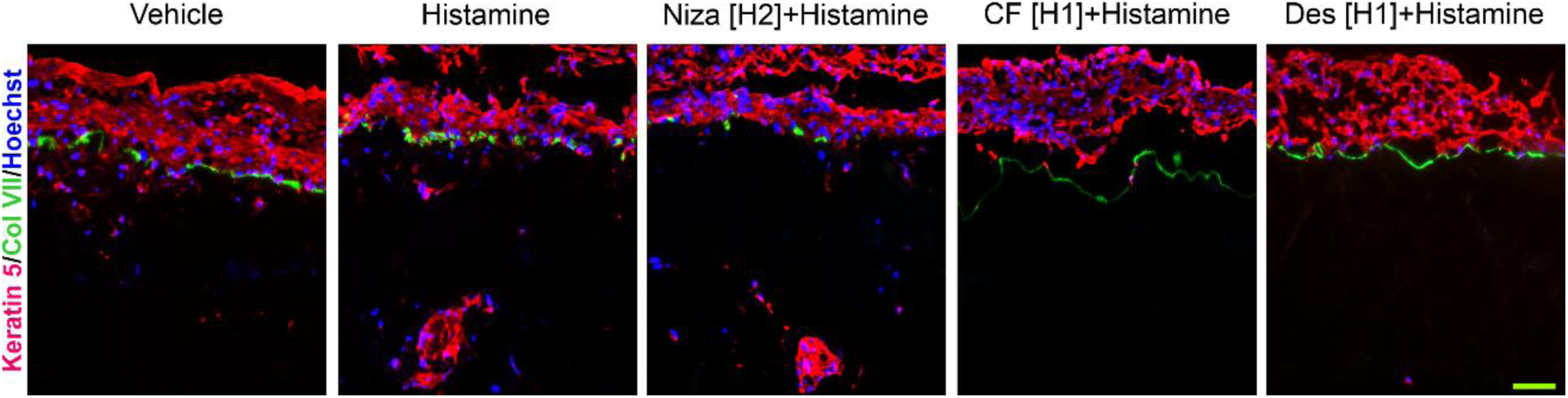
Histamine enhances invasion in an H1-specific manner. Oncogenic *HRAS* and *CDK4*-transformed primary keratinocytes were grown on human dermis at the air-liquid interface for 14 days and subjected to the drug treatments as shown. Immunofluorescence staining of keratin 5 (red), col VII (green), and nuclei (blue) was performed on the resulting cSCC organoids. Niza-Nizatidine, CF-clemastine fumarate, Des-Desloratadine. Scale bar=100μm, conducted with replicates and repeated three independent times.

**Extended Fig. 7.**
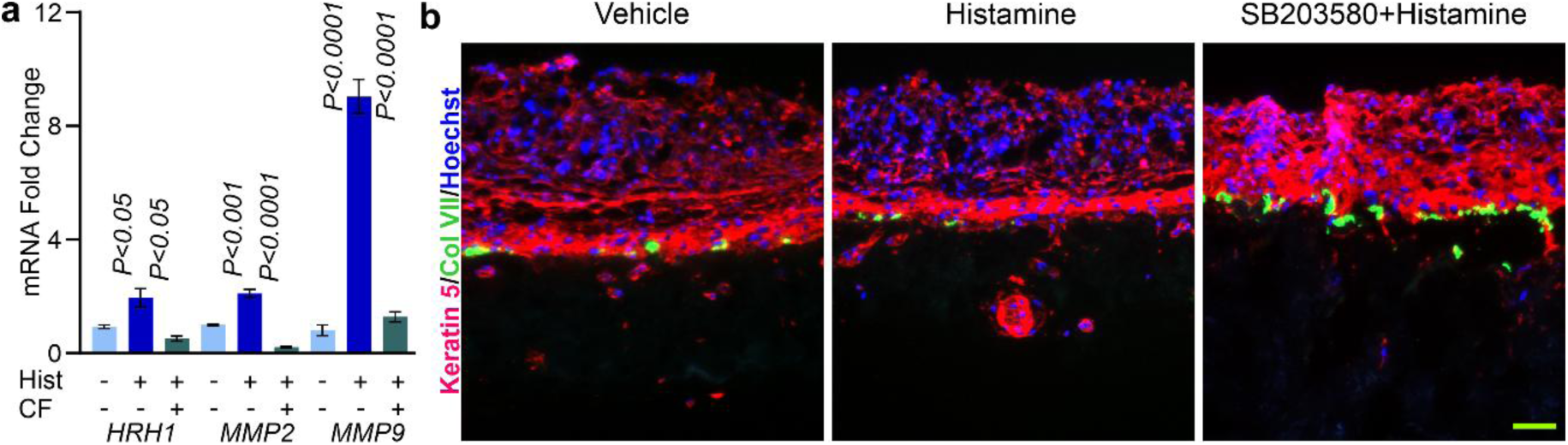
Histamine enhances invasion via P38. **a**, qRT-PCR analysis of *HRH1*, *MMP2*, and *MMP9* in skin organoids at day 7, upon histamine or antihistamine and histamine treatment. Hist-histamine, CF-clemastine fumarate. Results are representative of independent experiments conducted in replicates. Two-tailed Student’s t-tests calculated the *P-value* by comparing Hist to control, then CF to the Hist group for each gene. Error bar-standard deviation. **b**, Keratin 5 (red) and type VII collagen (green) immunofluorescence of oncogenic *HRAS* and *CDK4*-transformed human skin organoids following treatment with histamine or the P38 inhibitor SB203580. Scale bar=100μm, conducted with replicates and repeated three independent times.

**Extended Fig. 8.**
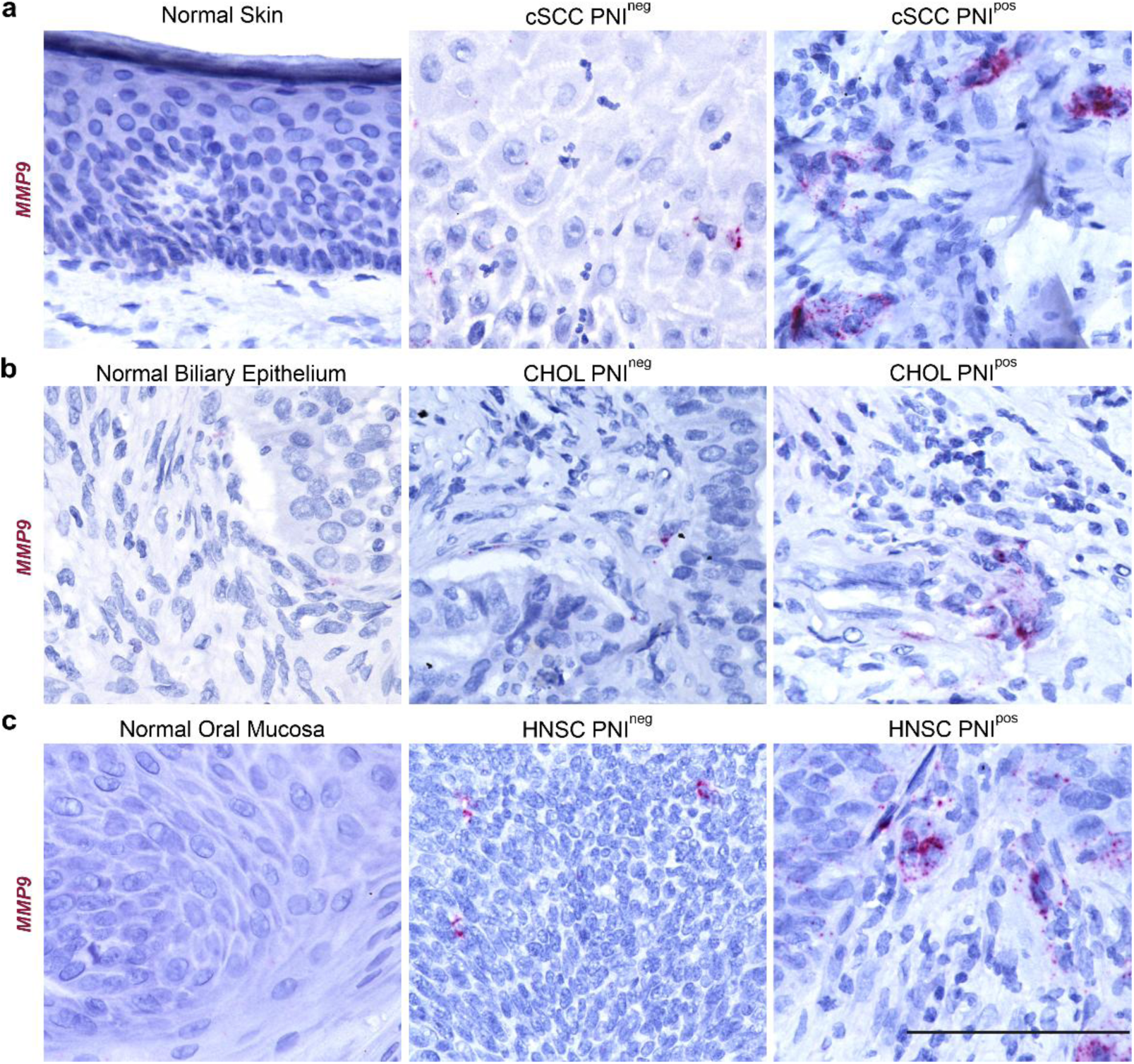
Enrichment of *MMP9* in PNI^pos^ tumors. **a**-**c**, Examination of *MMP9* (red) expression with RNAscope in normal skin, normal biliary epithelium, and oral mucosa compared to PNI^neg^ and PNI^pos^ cSCC, CHOL, and HNSC. Scale bar=100μM, images are representative of n=5/group.

**Extended Fig. 9.**
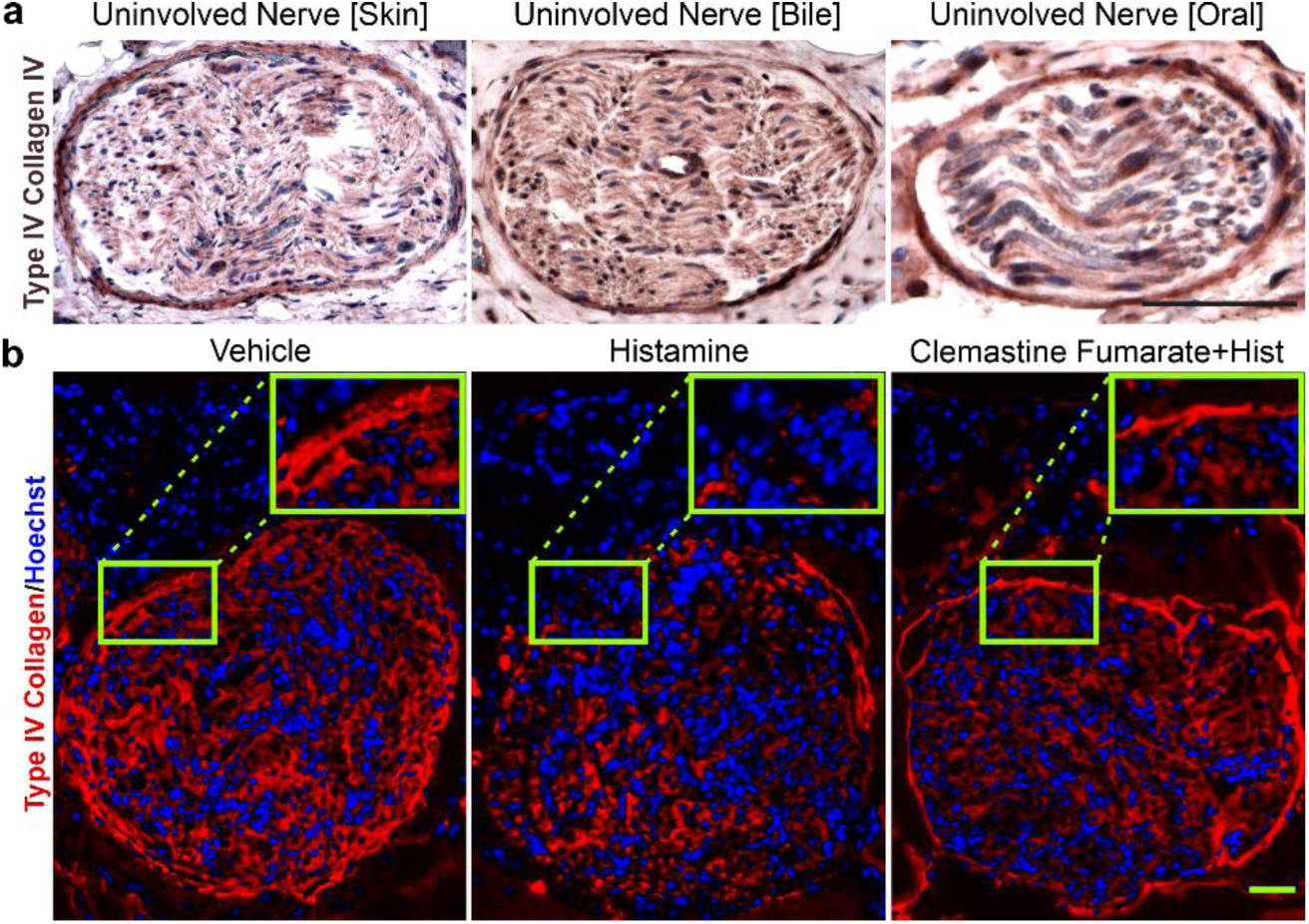
Histamine suppresses type IV collagen in PNI organoids. **a**, Immunohistochemistry for type IV collagen in uninvolved nerves of PNI^pos^ cSCC, CHOL, and HNSC. Scale bar=100μm, images are representative of n=5/group. **b**, Immunofluorescence analysis of type IV collagen (red) in PNI organoids treated with vehicle, histamine, or H1-antihistamine (clemastine fumarate) and histamine. Hist-histamine, Scale bar=100μm, conducted with replicates and repeated three independent times.

**Extended Fig. 10.**
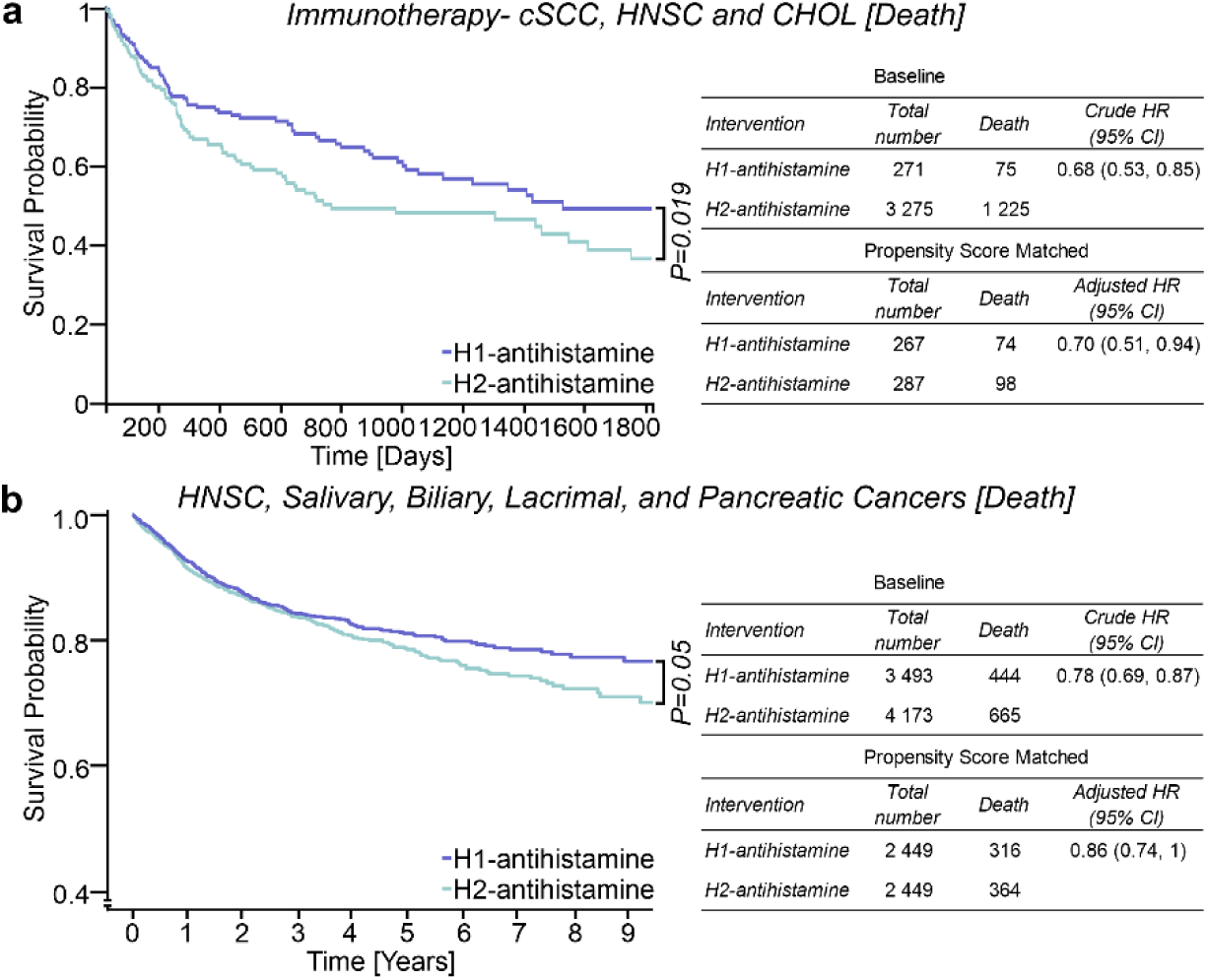
H1-antihistamines improve survival of patients receiving immunotherapy. **a**, Left, Kaplan-Meier survival analysis from TriNetX for cSCC, HNSC, and CHOL patients on immunotherapy and H1-or H2-antihistamines.Two-tailed Student’s t-tests calculated the *P-value*. Right, crude (baseline)-and propensity-score-matched analysis for hazard ratios. The Cox proportional hazard model calculated hazard ratios. **b**, Left, Kaplan-Meier survival evaluation from Atropos for HNSC, salivary, lacrimal, and pancreatic cancer patients with H1-or H2-antihistamines. Two-tailed Student’s t-tests calculated the *P-value*. Right, crude (baseline)-and propensity-score-matched analysis for hazard ratios. The Cox proportional hazard model calculated hazard ratios.

## Methods

### Human patient samples

Tumor samples from cSCC patients with PNI were collected at the time of surgical removal. We collected five samples from both male and female patients aged 70-98. All participating individuals provided informed written consent. A histological review by a board-certified pathologist confirmed PNI involvement and defined tumor differentiation status. The Institutional Review Board (IRB) at Stanford University approved this protocol (#21750). All collected samples were subjected to tissue dissociation at the time of collection.

Archival formalin-fixed paraffin-embedded (FFPE) normal skin, biliary epithelium, oral mucosa, PNI^pos^ and PNI^neg^ cSCC, CHOL, and HNSC tumors were obtained from the Stanford University pathology archives under an approved IRB protocol (#23838). Tumor grade and PNI status were obtained from pathology reports and confirmed by a board-certified pathologist. These samples were used for RNAscope, IHC, and toluidine staining.

### Single-cell preparation and scRNA-seq

Fresh tumor samples from PNI^pos^ cSCC were minced in serum-free Dulbecco’s Modified Eagle Medium (DMEM, Thermo Fisher Scientific) with 0.2 mg/ml DNase I (Worthington Inc.). Tissue samples were incubated in 0.25% trypsin-EDTA (Thermo Fisher Scientific) at 37°C for 30 minutes with trituration every 10 minutes. Trypsin-EDTA activity was quenched with DMEM supplemented with 10% fetal bovine serum (FBS) (Thermo Fisher Scientific) after 30 minutes, and the cell suspension was filtered through a 70µ filter (BD Falcon) to remove tissue debris. After washing two times, cells were resuspended in serum-free DMEM with DNase I, and cell viability was measured using trypan blue staining (Invitrogen). Overall, we maintained >90% cell viability for all tumors in this study. Following the manufacturer’s protocols for Chromium single-cell 3’v3 (10X Genomics), single-cell suspensions were subjected to GEM generation, barcoding, post-GEM RT cleanup, cDNA amplification, and library construction. Captured tumor cells ranged between 2000 and 10,000. Quality checks for constructed sequencing libraries were performed at Novogene Sacramento, California, followed by sequencing using an Illumina sequencer.

### Computational methods

#### scRNA-Seq data processing

scRNA sequencing files (FASTQ) were obtained from Novogene, and publicly available PNI-negative cSCC scRNA-seq data (BAM) was downloaded from NCBI GEO (*GSE144236*)^9^. BAM files were converted to FASTQ files with *bamtofastq* (*v1.3.2*). Raw sequencing data was aligned to the hg38 human reference genome and quantified using Cellranger (*v5.0.1*). Doublets were identified with scrublet (*v0.2.3*) and excluded from the analysis. Cells with >200 non-zero genes and >20% mitochondrial reads were retained for analysis. The Seurat pipeline (*v4.0.6*) was applied to normalize counts, reduce dimensionality, and define cell clusters (Louvain algorithm). We obtained a median of 1908 genes/cell and a median of 5755 UMI/cell. UMAP dimensionality reduction was performed with *RunUMAP* in Seurat.

#### Cell type annotation

The standard Seurat pipeline (*v4.0.6*) was used to identify broad cell types. Key marker genes defining keratinocytes (*KRT5*, *KRT14*, *KRT1*, and *KRT10*), salivary glands (*HTN3*), lymphocytes (*CD3E*, *GNLY*, *CD79A*), myeloid cells (LYZ, *HLA-DRB1*, *HLA-DRA*, and *HLA-DQB2*), fibroblasts (*LUM*, *COL1A1*, and *COL1A1*), endothelial cells (*VWF*, *TFF3*, and *CLDN5*), and melanocytes (*PMEL*, *DCT*, and *MLANA*) were obtained from Human Protein Atlas (https://www.proteinatlas.org/) and compared to marker genes identified by *FindMarkers* as well as previously published markers^9^.

PNI-negative/positive tumor epithelial cell samples were analyzed separately to highlight detailed cellular states. Differentially expressed genes were identified for each group, gene expression was scaled, and principal component dimensionality reduction was performed. The top 50 principal components were input to Harmony (*v0.1.0*) to remove patient-specific batch effects, followed by Louvain clustering. cSCC tumor subpopulation marker genes^9^, were used to define basal, cycling, and tumor-specific keratinocytes (TSK). For PSTC-1/2 gene enrichment analysis, each cluster’s top 100 marker genes were input into EnrichR (https://maayanlab.cloud/Enrichr/)^40,41^. *AddModuleScore*^42^ was used to score tumor cells for gene signatures. The stress-like signature enriched in PSTC-1/2 was obtained from HNSC scRNA-seq^11^. DvP scores were calculated based on gene signatures^10,43^. The Wilcoxon rank sum test with *FindMarkers* calculated P-values for differential gene expression.

#### Cell-cell interaction via ligand-receptor signals

To identify cell-cell interactions, we used CellChat (*v1.1.3*)^12^. Normal skin and PNI-negative/positive cSCC were analyzed separately to define interactions in each group. PNI-negative/positive cSCC datasets were then integrated, and differential signaling analysis was performed with *rankNet*(*mode=’comparison’*) to define signaling pathways with differential activity. Signaling pathways involving tumor cells as senders or receivers were prioritized for further investigation. Pathways identified as PNI-specific by CellChat but possessing known roles in cSCC development were excluded from the final list of PNI-specific signaling pathway candidates to focus on novel interactions.

#### Copy number alteration

*InferCNV* (*v1.7.1*) was used to estimate copy number profiles for *KITLG* on PNI-negative/positive TSK populations. Fibroblasts and endothelial cells were used as controls.

#### SCENIC and ChiP analysis

Gene regulatory networks and transcription factor-target interactions were inferred with *pySCENIC* (*v0.9.18*). Cells from the PNI-positive tumors were used as input. Reference files were downloaded from the SCENIC website (https://scenic.aertslab.org/)^44^. *hs_hgnc_curated_tfs.txt* command identified a list of transcription factors regulating *KITLG*. *Hg38 refseq-r80 10kb_up_and_down_tss.mc9nr.feather* was used for the ranking database and *motifs-v9-nr.hgnc-m0.001-o0.0.tbl* for the motif database. The E2F1, IRF1, IRF2, and STAT1 regulon signatures were created by aggregating all the target genes for a specific transcription factor from the “*pyscenic ctx*” command and then scoring each cell with the *AddModuleScore* function in Seurat. As shown before^45^, publicly available ChIP datasets on ENCODE were analyzed using Integrated Genomics Viewer^46,47^, to visualize transcription factor binding at the KITLG promoter.

#### TCGA data

Pan-cancer cohort RSEM counts were downloaded from UCSC Xena (https://tcga.xenahubs.net:443) along with clinical metadata, including PNI status. Counts were normalized using *DESeq2,* and the variance stabilized transform (*vst function*) was applied to normalized counts for downstream analysis. VST expression data was input for gene signature scoring with GSVA (*v1.52.3*). Differential expression of *KITLG*, *HRH1*, mast cell signature genes, P38 pathway genes, and *MMPs* between PNI-negative/positive tumors was identified using linear regression tests with *RegParallel*. T-statistics from the linear regressions were used as input to *fgsea* (*v1.18.0*) for gene set enrichment analysis. The mast cell infiltration signature consisted of the top 20 marker genes of mast cells from scRNA-Seq data in this article. The P38 pathway signature was obtained from *BIOCARTA_P38MAPK_PATHWAY in MSigDB* (https://www.gsea-msigdb.org/gsea/msigdb/index.jsp).

### RNAscope ISH

Duplex RNAscope ISH (RNAscope® 2.5 HD Duplex Detection kit, Bio-Techne-Advanced Cell Diagnostics) was performed on normal skin, normal biliary epithelium, normal oral mucosa, PNI^neg^ and PNI^pos^ cSCC, CHOL, and HNSC FFPE samples following the manufacturer’s protocol. Briefly, 6µ thick tissue sections were deparaffinized and treated with hydrogen peroxide (RNAscope pretreatment kit, Bio-Techne-Advanced Cell Diagnostics) to quench endogenous peroxidase activity. Next, we performed target retrieval and protease treatment (RNAscope pretreatment kit, Bio-Techne-Advanced Cell Diagnostics). A duplex combination of *KITLG* (RNAscope® Probe-Hs-*KITLG*-C1, Bio-Techne-Advanced Cell Diagnostics)-*KIT* (RNAscope® Probe-Hs-*KIT*-C2, Bio-Techne-Advanced Cell Diagnostics) or *HRH1* (RNAscope® Probe-Hs-*HRH1*-C2, Bio-Techne-Advanced Cell Diagnostics)-*MMP2* (RNAscope® Probe-Hs-*MMP2*-C1, Bio-Techne-Advanced Cell Diagnostics) was hybridized at 40°C for 2 hours using hybridization oven (Bio-Techne-Advanced Cell Diagnostics). *KIT*, *HRH1* [(alkaline phosphatase) AP-red], *KITLG*, and *MMP2* [(horseradish peroxidase) HRP-teal] signals were detected at the end of six or ten pre-amplification and amplification hybridization steps. Samples were counterstained with Gill’s hematoxylin (Stat lab medical products) for 30 seconds at room temperature, and slides were washed and mounted using a Vectamount permanent mounting medium (Vector Laboratories). For *MMP9* (RNAscope® Probe-Hs-*MMP9*-C2, Bio-Techne-Advanced Cell Diagnostics), RNAscope ISH was performed as described above, and the signal (AP-red) was detected at the end of six hybridization cycles, followed by counterstaining and mounting. Both positive (RNAscope® Probe-Hs-*PPIB*-C1/*POLR2A*-C2) and negative controls (RNAscope® Probe, targeting bacterial gene *DapB*) were included in each RNAscope run.

### Cell culture and treatments

Neonatal human primary keratinocytes were isolated from freshly discarded tissue specimens (IRB #76094) following protocols as described earlier^15^. Cells were maintained at 37°C with 5% CO2 in medium 154 (Thermo Fisher Scientific) supplemented with human keratinocyte growth supplement (HKGS) (Thermo Fisher Scientific). To avoid baseline proliferation induced by HKGS in keratinocytes, we changed the culture media to growth supplement-free medium 154 16-24h before any treatments. Human primary keratinocytes were treated with fludarabine (200µM, Selleckchem) for 1-3 hours. Histamine (10 or 20µM, Sigma-Aldrich) alone or with pre-treatment (30-60 minutes) with clemastine fumarate or desloratadine (both at 10 or 20µM, Selleckchem) or with nizatidine (5 or 10µM, Selleckchem) were added to keratinocyte cultures. Human primary keratinocytes were pre-treated with adezmapimod (SB203580, 10µM, Selleckchem) for 1 hour before histamine treatment (20µM).

### Spheroids, cSCC neoplastic skin organoids, and PNI organoids

Oncogenic *CDK4* or *HRAS* were transfected in Phoenix cells to generate a retroviral medium. We then transduced human primary keratinocytes with this medium at low centrifugation in the presence of 5mg/mL polybrene (Sigma-Aldrich). Cells were maintained in medium-154 supplemented with HKGS at 37°C with 5% CO2.

Oncogenic *CDK4*-*HRAS* co-expressing keratinocytes (10,000-20,000 cells) were seeded in ultra-low attachment plates (Nunclon Sphera, Thermo Fisher Scientific) to establish spheroids over 24-36 hours. Spheroids were transferred with a wide-bore pipette tip to a 96-flat bottom plate containing synthetic Vitrogel hydrogel matrix (TheWell Bioscience) mimicking skin extracellular matrix with high HKGS concentration. Spheres were treated with histamine (20µM) alone or pre-treated with either clemastine fumarate (20µM), desloratadine (20µM), nizatidine (5µM), or adezmapimod (SB203580, 10µM) followed by histamine (20µM) treatment. Spheroids were observed over 4-7 days, and invasion in the hydrogel matrix was assessed using a Cyto3D live-dead assay kit (TheWell Bioscience) and fluorescence microscopy.

Human cSCC neoplastic skin organoids were generated as described^15^. Briefly, 0.5-0.75×10^6^ oncogenic *CDK4*-*HRAS* co-expressing keratinocytes were populated over a devitalized acellular dermis (New York Firefighters Skin Bank). The organoids were treated daily with histamine or drugs, and these treatments were placed alternately on the epidermis or in the culture medium. Organoids were maintained at the air-liquid interface to achieve 3D tissue growth over 7 days for mRNA expression analysis or 14 days for invasion assessment in keratinocyte growth medium (KGM)^15^.

For PNI organoids, we isolated vagus nerves from rats marked for euthanasia due to aging or excess breeding (#19807). All isolated nerves were cleaned and washed in Dulbecco’s phosphate-buffered saline (DPBS, Thermo Fisher Scientific) containing a 2X Pen-Strep cocktail. Nerves were briefly submerged in Matrigel (Corning) before placing them on the human devitalized acellular dermis. Oncogenic *CDK4*-*HRAS* co-expressing keratinocytes (0.5-0.75×10^6^) were then seeded on top of the nerves and treated daily with histamine or combined with other inhibitors as described above. PNI organoids were cultured for 14 days in KGM at the air-liquid interface, followed by evaluation of PNI with immunofluorescence upon harvest.

### Gene expression

Total RNA was isolated from cultured keratinocytes treated with histamine or indicated inhibitors following the manufacturer’s instructions using the RNeasy plus mini kit (Qiagen). Cells were lysed with RLT buffer and Qiashredder columns (Qiagen), and gDNA was removed using gDNA eliminator columns (Qiagen). Total RNA was eluted in DNase-, RNase-free ultra-pure water (Qiagen). cSCC neoplastic skin organoids were homogenized with metal beads using a TissueLyser LT (Qiagen) at 50Hz for 4 minutes. Total RNA was isolated from the homogenized samples as described above. Total RNA (0.1-1μg) was reverse transcribed, and cDNA synthesis was performed using the iSCRIPT 20X cDNA kit (Bio-Rad). Gene expression of *KITLG*, *CXCL10*, *HRH1*, *MMP1*, *2*, *7*, and *9* (predesigned PrimeTime® qPCR primer assays; Integrated DNA Technologies) were analyzed with maxima SYBR Green/ROX qPCR master mix (Fisher Scientific) using a LightCycler 480 II (Roche). Gene expression was calculated by normalizing to housekeeping genes *L32* and *18S* using the ΔCt-method.

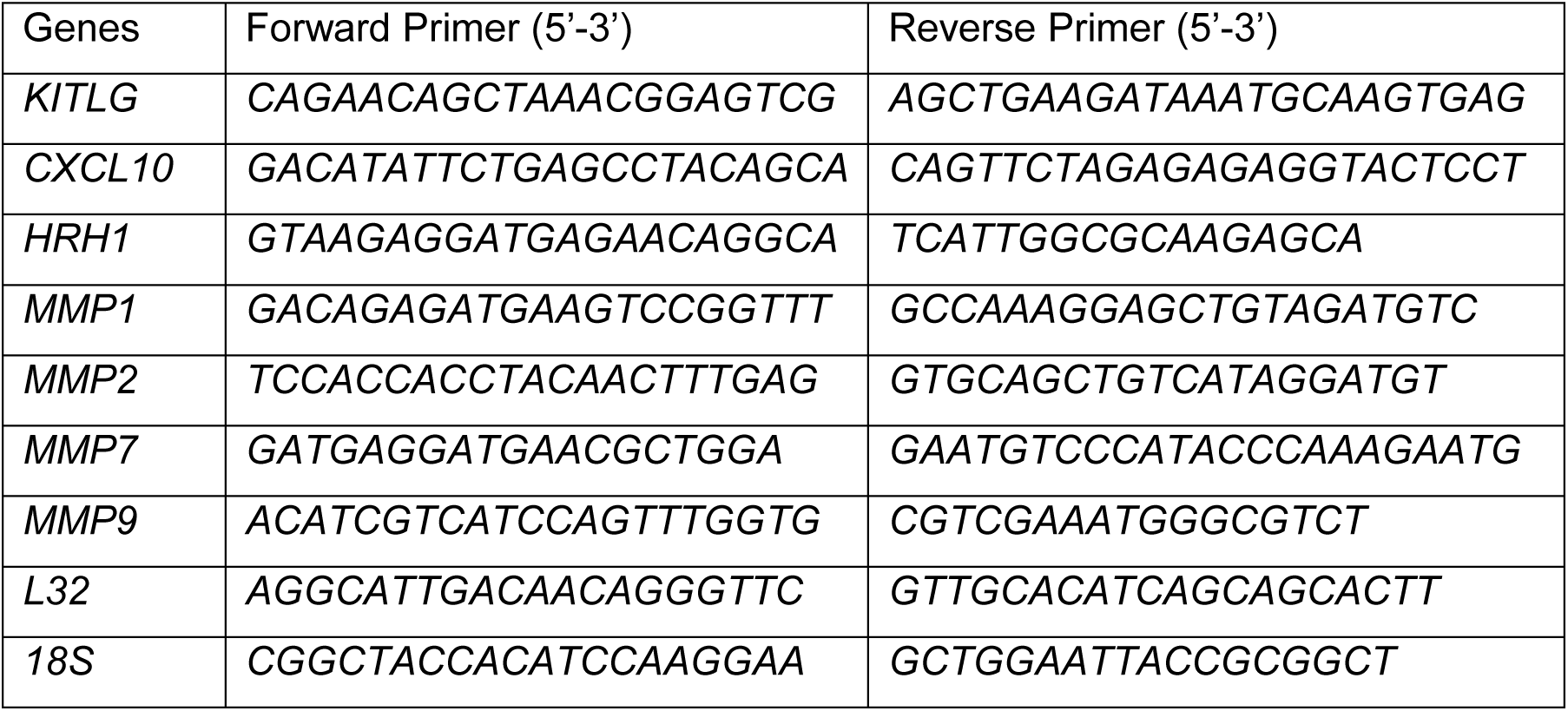

### Protein Expression

#### Immunoblotting

Histamine, vehicle control, or inhibitor-treated human primary keratinocytes were lysed in RIPA buffer (Fisher Scientific) supplemented with phosphatase inhibitor (Fisher Scientific) and protease inhibitor cocktails (Sigma-Aldrich) at the given time points. Protein quantity was measured by bicinchoninic acid (BCA) (Pierce, Thermo Fisher Scientific). Total proteins were loaded on the 4-20% Mini-PROTEAN TGX gel (Bio-Rad) and transferred to the nitrocellulose membrane (Bio-Rad) using the wet transfer method. The membranes were blocked with protein-free Tris-buffered saline (TBS) blocking buffer (Pierce, Thermo Fisher Scientific). Primary antibodies used for immunoblotting were H1 (H1R, rabbit, 1:1000, Abcam), phospho-JNK (T183/y185) (rabbit, 1:1000, Cell Signaling Technology), JNK (rabbit, 1:1000, Cell Signaling Technology), phospho-p38 MAPK (Thr180/Tyr182) (rabbit, 1:1000, Cell Signaling Technology), p38 MAPK (rabbit, 1:1000, Cell Signaling Technology). Horseradish peroxidase (HRP)-conjugated β-Actin (rabbit/mouse, 1:1000, Cell Signaling Technology) was used as a housekeeping marker. Rabbit IgG HRP secondary antibody (donkey, 1:5000, Millipore Sigma) was used prior to chemiluminescence signal detection. Proteins were visualized by SuperSignal West Femto maximum sensitivity substrate (Thermo Fisher Scientific) using a gel documentation system (Bio-Rad).

#### ELISA

Cell culture supernatants were collected at the indicated times following various treatments in human primary keratinocytes. Secreted protein levels of MMP2 and MMP9 were detected by enzyme-linked immunosorbent assay (ELISA) following the manufacturer’s instructions with the human quantikine ELISA kit (Bio-Techne-R&D Systems). Optical density was measured at 450nm with wavelength correction at 540nm using a plate reader (SpectraMax M5, Molecular Devices). The final concentration of MMP2 and MMP9 was calculated based on the standard curve equations obtained by positive standard control readings.

#### Phosphoproteomics

Human primary keratinocytes were treated with histamine (20µM) or vehicle control for 10 minutes. Treated keratinocytes were washed and collected by scraping in ice-cold PBS and centrifuged to collect cells. Cells were lysed by sonicating (3 cycles of 5s) in lysis buffer containing 9M urea, 1mM sodium orthovanadate, 20mM HEPES (Sigma-Aldrich), 2.5mM sodium pyrophosphate, 1mM β-glycerophosphate (Fisher Scientific) at pH 8.0, followed by centrifugation at 16,000g for 10 min. The supernatant was collected, and protein quantitation was performed using BCA assay (Pierce, Thermo Fisher Scientific). Approximately 500μg of total protein was used for the reduction, alkylation, and digestion at 37 °C for 16 h with trypsin (Promega) at a 1:50 protein: protease ratio. Sep-Pak Vac 1 cc (50 mg) tC18 cartridges were used to clean up the digested samples. The trypsin-digested lysates were acidified with 1/20 volume of 20% trifluoroacetic acid (TFA) and centrifuged at 16,000g for 15 min. The peptide solutions were loaded onto the washed columns. The columns were washed, and peptides were eluted with 1 ml of the elution buffer (40% acetonitrile in 0.1% TFA elution buffer). The peptide amount per sample was quantified with peptide assay (Pierce, Thermo Fisher Scientific). The resuspended peptides were volume adjusted by adding 1M EPPS, pH 8.0, in a 3:1 ratio. 50 μg of peptide from each sample was labeled with 200μg of tandem mass tag (TMT) pro reagents resuspended in 5μl, 100% acetonitrile for 3h at 25°C, followed by quenching with hydroxylamine (0.5% final) and acidified with trifluoroacetic acid (2% final). The samples were desalted with 50 mg tC18 Sep-Paks and dried by vacuum.

The dried peptides were resuspended in 0.1% TFA. Approximately 120 μg of peptide mix was subjected to orthogonal pH reverse phase fractionation on a 3×150 mm column packed with 1.9 μm Poroshell C18 material (Agilent) equilibrated with buffer A (5% acetonitrile in 10mM ammonium bicarbonate, pH 8). Peptides were fractionated using a linear gradient from 12% to 45% buffer B (90% acetonitrile in 10mM ammonium bicarbonate, pH 8) at a 0.8 ml/minute flow rate for 45 minutes. The samples were resuspended in 0.1% TFA, desalted on StageTips, and vacuum dried. Peptides were reconstituted in 5% formic acid + 5% acetonitrile for LC-MS3 analysis. The concatenated fractions were cleaned with C18 spin tips (Pierce, Thermo Fisher Scientific) and dried in speed-vac before liquid chromatography-mass spectrometry (LC-MS) analysis. LC-MS was performed as described earlier^48^, using a Dionex Ultimate 3000 RSLCnano system (Thermo Fisher Scientific) and an Orbitrap Eclipse Tribrid MS (Thermo Fisher Scientific).

Peptide-spectrum match (PSM) scores from decoy proteins and redundant PSMs were filtered out from the final data analysis. Quantification and statistical analysis were performed by MSstatsTMT v2.2.7, an open-source R/Bioconductor package. MSstatsTMT generated a normalized quantification report across all the samples at the protein level from the processed PSM report. Global median normalization equalized the median of the reporter ion intensities across all the channels and TMT mixtures. The normalized reporter ion intensities of all the peptide ions mapped to a protein were summarized into a single protein level intensity in each channel and TMT mixture. MSstatsTMT performed differential abundance analysis for the normalized protein intensities. To test the two-sided null hypothesis of no changes in abundance, the model-based test statistics were compared with the Student’s t-test distribution with the degrees of freedom appropriate for each protein and each dataset. The resulting P values were adjusted to control the FDR using Benjamini-Hochberg’s method. The phosphorylated proteins were used to analyze pathway enrichment via EnrichR.

#### Immunofluorescence, Immunohistochemistry, and Toluidine Blue staining

Immunofluorescence (IF) was performed as reported earlier^49^. Briefly, 7μ thick frozen tissue sections for cSCC neoplastic skin organoids and PNI organoids were used to assess invasion, PNI, or nerve sheath degradation by IF. Primary antibodies used were keratin 5 (rabbit, 1:500, BioLegend), type VII collagen (mouse, 1:250, Millipore Sigma), and type IV collagen (rabbit, 1:100, Abcam). Secondary antibodies used were anti-mouse IgG Alexa Fluor 488 (goat, 1:250, Thermo Fisher Scientific) and anti-rabbit IgG Alexa Fluor 594 (goat, 1:250, Thermo Fisher Scientific). Cell nuclei were stained with Hoechst 33342, trihydrochloride, trihydrate (1:4000, Thermo Fisher Scientific). Sections were mounted with Prolong Gold Antifade reagent (Thermo Fisher Scientific).

Immunohistochemistry (IHC) was performed using the VECTASTAIN elite ABC-HRP Kit (Vector Laboratories)^50^ on nerve tissue sections. FFPE sections were deparaffinized and subjected to antigen retrieval. Antigen unmasking solution (Tris-based, pH 9, Vector Laboratories) was used at 98°C in a water bath for antigen retrieval, followed by quenching of endogenous peroxidase activity with Bloxall blocking solution (Vector Laboratories). Primary and secondary antibodies were type IV collagen (rabbit, 1:100, Abcam) and biotinylated Horse anti-rabbit IgG antibody (H+L, horse, 1:200, Vector Laboratories). ABC-HRP Kit and ImmPACT AMEC red peroxidase substrate (Vector Laboratories) were used to develop IHC signals, and slides were counter-stained with hematoxylin (Vector Laboratories). Sections were mounted with VectaMount aqueous mounting medium (Vector Laboratories).

Normal skin, normal biliary epithelium, normal oral mucosa, PNI^neg,^ and PNI^pos^ cSCC, CHOL, and HNSC FFPE samples were deparaffinized and stained with a toluidine blue kit (VitroVivo Biotech, Fisher Scientific) as per the manufacturer’s instructions. Slides were washed, dehydrated, cleared, and mounted using a VectaMount permanent mounting medium (Vector Laboratories).

### Microscopy, image processing, and cell counting

RNAScope ISH, IHC, and toluidine blue-stained images were obtained using a brightfield microscope (BZ-X810, Keyence) at 20- or 40X magnifications. Zeiss Axio Observer. A Z1 fluorescent microscope was used to capture IF images from cSCC neoplastic skin organoids, PNI organoids, and live-dead staining images of the spheroid model at 5X magnification. All images were processed using CorelDraw Suite 2019, and brightness, contrast, intensity, red, blue, green channels, and gamma were adjusted on the control samples. The same adjustment settings were applied to all the samples in an experiment. The number of toluidine blue-positive (histamine) cells/field of view was quantified using the cell counter function in Fiji ImageJ software.

### Electronic Health Record Survival Analysis

We employed two independent US national and global EHR databases, Atropos (https://www.atroposhealth.com/) and TriNetX (https://trinetx.com/), for the retrospective patient survival analysis. We included patients aged 20 and above, diagnosed between 2008 and 2024, with cSCC, HNSC, CHOL, salivary, biliary, lacrimal, and pancreatic cancers. Patients with at least three prescriptions of antihistamines one month before/after diagnosis (TriNetX) or with the use of antihistamines for more than 6 months before the diagnosis (Atropos) were included. For immunotherapy (TriNetX), patients with at least three prescriptions of antihistamines one month before/after treatment with anti-PD-1/PD-L1 inhibitors were included. The TriNetX database comprises de-identified patient records and contains no identifiable information. For this study, we received ethical approval from the IRB of National Cheng Kung University Hospital, Taiwan (#AER-113-157).

H1-antihistamines used in this study were loratadine, desloratadine, cetirizine, levocetirizine, ketotifen, azelastine, fexofenadine, and clemastine; H2-antihistamines used were famotidine, nizatidine, ranitidine, and roxatidine. We included cSCC, HNSC, and CHOL patients treated with anti-PD-1/PD-L1 inhibitors to analyze the effects of antihistamine treatment on immunotherapy response. Overall survival or time to first metastasis (metastasis-free survival) was analyzed. Propensity score matching (PSM, 1:1 matching) was conducted to balance the two-arm study groups (H1-vs. H2-antihistamines). Matching with covariates was performed based on age, sex, race, and specific confounding comorbidities and medication. All statistics, including PSM, were analyzed using Atropos or the TriNetX platform. Descriptive analysis utilized counts and proportions for categorical variables and means ± standard deviation (SD) for continuous variables. Multivariable Cox proportional-hazards models were adopted to calculate hazard ratios (HR) and 95% confidence intervals to compare patients’ overall/metastasis-free survival. The absolute standardized mean difference (SMD) was used to examine the heterogeneity levels between the two treatment groups, with a 10% threshold.

## Statistical analysis

In addition to the scRNA-Seq, bioinformatics, and phosphoproteomics data analysis, all the other standard deviations and two-tailed Student’s t-tests were performed using GraphPad Prism version 9/10 for Windows (GraphPad Software). P values <0.5 were considered significant. Survival and metastasis-free survival analysis was performed using Kaplan-Meier methods and the Cox proportional hazard model calculated hazard ratios.

## Code and data availability

The code to reproduce figures and statistical tests for scRNA-Seq, CellChat, SCENIC, TCGA, and other data analyses is available at https://github.com/tjbencomo/pni-histamine. scRNA-Seq data will be provided upon request. Phosphoproteomics data are included as a supplementary table (supplementary table 4).

## Acknowledgment

We are deeply grateful to the patients who participated in this study. We acknowledge Elias Godoy and veterinary pathologist Dr. Kerriann M. Casey of Stanford University, RAF-1 (pathology/necropsy lab), for their support in obtaining the animals used in this study for nerve isolation. We thank Stanford Research Computing for the computational and data storage resources related to our scRNA-seq data analysis. We also thank Vanessa Lopez-Pajares, Zurab Siprashvili, Shiying Tao, Sumaira Aasi, Fred Baik, Gretchen Ehrenkaufer, Daria Mochly-Rosen, and Kevin Grimes for their insightful discussions. C.S. Lee is a recipient of the McCormick and Gabilan Faculty Awards. Both C.S. Lee and A. Srivastava are recipients of the SPARK research grant and are mentored by Stanford’s SPARK Translational Research Program.

## Author Contribution

A.S. led the study, hypothesized, designed and performed experiments, supervised assisting researchers, raised funding, analyzed and interpreted data, and wrote the manuscript with input from all authors. T.B. led scRNA-seq data analysis and computational methods, assisted with scRNA-seq study design, and provided valuable input in the manuscript draft. C.-N.L. performed survival analysis using TriNetX datasets. A.M. assisted with organoid generation. J.G. and T.J. helped with tumor collection and processing. L.W.S., I.M.D., A.J.T., and A.N. assisted with in vitro experiments and immunofluorescence assays. S.G. performed survival analysis using Atropos datasets. L.P. assisted with phosphoproteomics. P.D. performed phosphoproteomics, analyzed data, and provided inputs in the manuscript draft. C.M.R. advised on the phosphoproteomics study design and provided expertise. R.B. provided expertise on disease pathology and retrieved archival tumor samples. C.S.L. initiated the study, conceptualized, managed, and provided overall supervision, raised funding for this work, and provided final inputs and edits for the manuscript.

## References

1. Batsakis, J.G. Nerves and neurotropic carcinomas. Ann Otol Rhinol Laryngol 94, 426–427 (1985).

2. Bapat, A.A., Hostetter, G., Von Hoff, D.D. & Han, H. Perineural invasion and associated pain in pancreatic cancer. Nat Rev Cancer 11, 695–707 (2011).

3. Perez Garcia, M.P., Mateu Puchades, A. & Sanmartin Jimenez, O. Perineural Invasion in Cutaneous Squamous Cell Carcinoma. Actas Dermosifiliogr (Engl Ed) 110, 426–433 (2019).

4. Bakst, R.L., et al. Perineural Invasion and Perineural Tumor Spread in Head and Neck Cancer. Int J Radiat Oncol Biol Phys 103, 1109–1124 (2019).

5. Shirai, K., et al. Perineural invasion is a prognostic factor in intrahepatic cholangiocarcinoma. World J Surg 32, 2395–2402 (2008).

6. Massey, P.R., et al. Extensive Perineural Invasion vs Nerve Caliber to Assess Cutaneous Squamous Cell Carcinoma Prognosis. JAMA Dermatol 159, 1332–1338 (2023).

7. Karia, P.S., Morgan, F.C., Ruiz, E.S. & Schmults, C.D. Clinical and Incidental Perineural Invasion of Cutaneous Squamous Cell Carcinoma: A Systematic Review and Pooled Analysis of Outcomes Data. JAMA Dermatol 153, 781–788 (2017).

8. Marchesi, F., Piemonti, L., Mantovani, A. & Allavena, P. Molecular mechanisms of perineural invasion, a forgotten pathway of dissemination and metastasis. Cytokine Growth Factor Rev 21, 77–82 (2010).

9. Ji, A.L., et al. Multimodal Analysis of Composition and Spatial Architecture in Human Squamous Cell Carcinoma. Cell 182, 497–514 e422 (2020).

10. Bailey, P., et al. Driver gene combinations dictate cutaneous squamous cell carcinoma disease continuum progression. Nat Commun 14, 5211 (2023).

11. Puram, S.V., et al. Single-Cell Transcriptomic Analysis of Primary and Metastatic Tumor Ecosystems in Head and Neck Cancer. Cell 171, 1611–1624 e1624 (2017).

12. Jin, S., et al. Inference and analysis of cell-cell communication using CellChat. Nat Commun 12, 1088 (2021).

13. Das, A.K., et al. Stimulation of histamine H1 receptor up-regulates histamine H1 receptor itself through activation of receptor gene transcription. J Pharmacol Sci 103, 374–382 (2007).

14. Michinaga, S., Nagata, A., Ogami, R., Ogawa, Y. & Hishinuma, S. Histamine H(1) Receptor-Mediated JNK Phosphorylation Is Regulated by G(q) Protein-Dependent but Arrestin-Independent Pathways. Int J Mol Sci 25(2024).

15. Ridky, T.W., Chow, J.M., Wong, D.J. & Khavari, P.A. Invasive three-dimensional organotypic neoplasia from multiple normal human epithelia. Nat Med 16, 1450–1455 (2010).

16. Roach, D.M., et al. Up-regulation of MMP-2 and MMP-9 leads to degradation of type IV collagen during skeletal muscle reperfusion injury; protection by the MMP inhibitor, doxycycline. Eur J Vasc Endovasc Surg 23, 260–269 (2002).

17. Hwang, W.L., et al. Single-nucleus and spatial transcriptome profiling of pancreatic cancer identifies multicellular dynamics associated with neoadjuvant treatment. Nat Genet 54, 1178–1191 (2022).

18. Schmitd, L.B., et al. Spatial and Transcriptomic Analysis of Perineural Invasion in Oral Cancer. Clin Cancer Res 28, 3557–3572 (2022).

19. Zhang, B., et al. Single-cell RNA sequencing reveals intratumoral heterogeneity and potential mechanisms of malignant progression in prostate cancer with perineural invasion. Front Genet 13, 1073232 (2022).

20. Yuan, X., Dong, Z., Zhang, B., Li, Q. & Jiang, W. Combining single-cell spatial transcriptomics and molecular simulation to develop in vivo probes targeting the perineural invasion region of adenoid cystic carcinoma. Heliyon 10, e34628 (2024).

21. Varricchi, G., et al. Controversial role of mast cells in skin cancers. Exp Dermatol 26, 11–17 (2017).

22. Somasundaram, R., et al. Tumor-infiltrating mast cells are associated with resistance to anti-PD-1 therapy. Nat Commun 12, 346 (2021).

23. Derakhshani, A., et al. Mast cells: A double-edged sword in cancer. Immunol Lett 209, 28–35 (2019).

24. Ribatti, D. Mast Cells and Resistance to Immunotherapy in Cancer. Arch Immunol Ther Exp (Warsz) 71, 11 (2023).

25. Johnson, C., et al. Inhibition of Mast Cell-Derived Histamine Decreases Human Cholangiocarcinoma Growth and Differentiation via c-Kit/Stem Cell Factor-Dependent Signaling. Am J Pathol 186, 123–133 (2016).

26. Mino, M., Pilch, B.Z. & Faquin, W.C. Expression of KIT (CD117) in neoplasms of the head and neck: an ancillary marker for adenoid cystic carcinoma. Mod Pathol 16, 1224–1231 (2003).

27. Hang, D., et al. KIT polymorphisms were associated with the risk for head and neck squamous carcinoma in Chinese population. Mol Carcinog 56, 232–237 (2017).

28. Artuc, M., et al. Mast cell-derived TNF-alpha and histamine modify IL-6 and IL-8 expression and release from cutaneous tumor cells. Exp Dermatol 20, 1020–1022 (2011).

29. Tsai, M., Valent, P. & Galli, S.J. KIT as a master regulator of the mast cell lineage. J Allergy Clin Immunol 149, 1845–1854 (2022).

30. Sarasola, M.P., Taquez Delgado, M.A., Nicoud, M.B. & Medina, V.A. Histamine in cancer immunology and immunotherapy. Current status and new perspectives. Pharmacol Res Perspect 9, e00778 (2021).

31. Li, H., et al. The allergy mediator histamine confers resistance to immunotherapy in cancer patients via activation of the macrophage histamine receptor H1. Cancer Cell 40, 36–52 e39 (2022).

32. Jiang, S.H., Zhang, S., Wang, H., Xue, J.L. & Zhang, Z.G. Emerging experimental models for assessing perineural invasion in human cancers. Cancer Lett 535, 215610 (2022).

33. Deborde, S., et al. An In Vivo Murine Sciatic Nerve Model of Perineural Invasion. J Vis Exp (2018).

34. de Lima, P.O., et al. Development of an in vivo murine model of perineural invasion and spread of cutaneous squamous cell carcinoma of the head and neck. Front Oncol 13, 1231104 (2023).

35. Gschwandtner, M., et al. Histamine upregulates keratinocyte MMP-9 production via the histamine H1 receptor. J Invest Dermatol 128, 2783–2791 (2008).

36. Ancha, H.R., et al. Histamine stimulation of MMP-1(collagenase-1) secretion and gene expression in gastric epithelial cells: role of EGFR transactivation and the MAP kinase pathway. Int J Biochem Cell Biol 39, 2143–2152 (2007).

37. Cahill, K.N., et al. KIT Inhibition by Imatinib in Patients with Severe Refractory Asthma. N Engl J Med 376, 1911–1920 (2017).

38. Puzzovio, P.G., et al. Cromolyn Sodium differentially regulates human mast cell and mouse leukocyte responses to control allergic inflammation. Pharmacol Res 178, 106172 (2022).

39. Zhang, X., Gopalan, V., Syed, N., Hannenhalli, S. & Shern, J.F. Protocol for using single-cell sequencing to study the heterogeneity of NF1 nerve sheath tumors from clinical biospecimens. STAR Protoc 4, 102297 (2023).

40. Kuleshov, M.V., et al. Enrichr: a comprehensive gene set enrichment analysis web server 2016 update. Nucleic Acids Res 44, W90–97 (2016).

41. Chen, E.Y., et al. Enrichr: interactive and collaborative HTML5 gene list enrichment analysis tool. BMC Bioinformatics 14, 128 (2013).

42. Tirosh, I., et al. Dissecting the multicellular ecosystem of metastatic melanoma by single-cell RNA-seq. Science 352, 189–196 (2016).

43. Bencomo, T. & Lee, C.S. Gene expression landscape of cutaneous squamous cell carcinoma progression. Br J Dermatol 191, 760–774 (2024).

44. Aibar, S., et al. SCENIC: single-cell regulatory network inference and clustering. Nat Methods 14, 1083–1086 (2017).

45. Srivastava, A., et al. Cross-talk between IFN-gamma and TWEAK through miR-149 amplifies skin inflammation in psoriasis. J Allergy Clin Immunol 147, 2225–2235 (2021).

46. Robinson, J.T., Thorvaldsdottir, H., Wenger, A.M., Zehir, A. & Mesirov, J.P. Variant Review with the Integrative Genomics Viewer. Cancer Res 77, e31–e34 (2017).

47. Robinson, J.T., et al. Integrative genomics viewer. Nat Biotechnol 29, 24–26 (2011).

48. Marei, H., et al. Antibody targeting of E3 ubiquitin ligases for receptor degradation. Nature 610, 182–189 (2022).

49. Srivastava, A., et al. MAB21L4 Deficiency Drives Squamous Cell Carcinoma via Activation of RET. Cancer Res 82, 3143–3157 (2022).

50. Srivastava, A., et al. MicroRNA-146a suppresses IL-17-mediated skin inflammation and is genetically associated with psoriasis. J Allergy Clin Immunol 139, 550–561 (2017).

