## Supplemental Tables 1-7 for "Epithelial tumor cells utilize mast cell-derived histamine to regulate perineural invasion"

**Supplementary Table 1. Patient characteristics**

| Patient Code | Sex | Age | Anatomical Site | Tumor Differentiation Status | Immune Status |
|---|---|---|---|---|---|
| *LEE01* | Male | 74 | Left Neck | Poor | Immunocompromised |
| *LEE02* | Male | 87 | Left Cheek | Moderate | Immunocompetent |
| *LEE03* | Male | 70 | Left Lateral Forehead | Poor | Immunocompromised |
| *LEE04* | Male | 70 | Right Cheek | Poor | Immunocompetent |
| *LEE05* | Female | 98 | Left Temple | Moderate | Immunocompetent |

**Supplementary Table 2. Diverse keratinocyte cell subtype markers**

| Gene Symbol | Cell Cluster | Fold Change [$Log_2$] | P-Value | Adjusted P-Value |
|---|---|---|---|---|
| *SFRP1* | KC_Basal | 1.94 | *1.66E-31* | *6.09E-27* |
| *TIMP1* | KC_Basal | 1.87 | *5.35E-19* | *1.96E-14* |
| *SLC47A2* | KC_Basal | 1.68 | *6.25E-49* | *2.29E-44* |
| *WNT3* | KC_Basal | 1.62 | *1.74E-49* | *6.36E-45* |
| *CXCL14* | KC_Basal | 1.52 | *8.04E-92* | *2.94E-87* |
| *BGN* | KC_Basal | 1.46 | *1.38E-47* | *5.05E-43* |
| *AKR1C1* | KC_Basal | 1.43 | *3.00E-19* | *1.10E-14* |
| *BOC* | KC_Basal | 1.37 | *1.11E-27* | *4.08E-23* |
| *FOXP2* | KC_Basal | 1.37 | *1.23E-35* | *4.51E-31* |
| *SOX6* | KC_Basal | 1.36 | *1.11E-56* | *4.07E-52* |
| *IL20* | KC_PSTC1 | 2.34 | *1.25E-43* | *4.58E-39* |
| *CCL2* | KC_PSTC1 | 2.31 | *1.66E-47* | *6.08E-43* |
| *ZBTB16* | KC_PSTC1 | 2.01 | *1.63E-53* | *5.98E-49* |
| *SEMA5A* | KC_PSTC1 | 1.96 | *1.55E-50* | *5.68E-46* |
| *NFIL3* | KC_PSTC1 | 1.94 | *2.41E-24* | *8.82E-20* |
| *DUSP2* | KC_PSTC1 | 1.86 | *3.11E-29* | *1.14E-24* |
| *HES1* | KC_PSTC1 | 1.85 | *7.92E-30* | *2.90E-25* |
| *EGR2* | KC_PSTC1 | 1.81 | *4.75E-54* | *1.74E-49* |
| *MAFF* | KC_PSTC1 | 1.81 | *2.84E-33* | *1.04E-28* |
| *SOCS3* | KC_PSTC1 | 1.80 | *3.21E-27* | *1.18E-22* |
| *CENPA* | KC_Cycling | 5.99 | *0.00E+00* | *0.00E+00* |
| *CDC25C* | KC_Cycling | 5.85 | *9.77E-241* | *3.58E-236* |
| *ASF1B* | KC_Cycling | 5.64 | *0.00E+00* | *0.00E+00* |
| *CCNA2* | KC_Cycling | 5.63 | *0.00E+00* | *0.00E+00* |
| *GTSE1* | KC_Cycling | 5.56 | *0.00E+00* | *0.00E+00* |
| *DEPDC1* | KC_Cycling | 5.55 | *0.00E+00* | *0.00E+00* |
| *ERCC6L* | KC_Cycling | 5.51 | *9.37E-272* | *3.43E-267* |

| | | | | |
|---|---|---|---|---|
| *NEK2* | KC_Cycling | 5.50 | *0.00E+00* | *0.00E+00* |
| *CDCA2* | KC_Cycling | 5.46 | *0.00E+00* | *0.00E+00* |
| *UBE2C* | KC_Cycling | 5.45 | *0.00E+00* | *0.00E+00* |
| *CLCA4* | KC_PSTC2 | 3.33 | *3.71E-40* | *1.36E-35* |
| *MMP7* | KC_PSTC2 | 3.02 | *5.58E-13* | *2.04E-08* |
| *SOSTDC1* | KC_PSTC2 | 2.98 | *7.55E-88* | *2.77E-83* |
| *EN1* | KC_PSTC2 | 2.88 | *8.56E-19* | *3.13E-14* |
| *NES* | KC_PSTC2 | 2.69 | *4.88E-32* | *1.78E-27* |
| *PRICKLE1* | KC_PSTC2 | 2.64 | *4.91E-43* | *1.80E-38* |
| *BBOX1* | KC_PSTC2 | 2.57 | *2.13E-47* | *7.79E-43* |
| *CYP1B1* | KC_PSTC2 | 2.50 | *3.46E-54* | *1.27E-49* |
| *AGR2* | KC_PSTC2 | 2.46 | *5.17E-14* | *1.89E-09* |
| *RHOV* | KC_PSTC2 | 2.44 | *1.84E-89* | *6.73E-85* |
| *MMP1* | KC_TSK | 4.76 | *1.60E-126* | *5.85E-122* |
| *MMP10* | KC_TSK | 4.75 | *5.04E-114* | *1.84E-109* |
| *PTGS2* | KC_TSK | 3.96 | *1.06E-27* | *3.89E-23* |
| *LAMC2* | KC_TSK | 3.60 | *1.48E-119* | *5.43E-115* |
| *IGFBP2* | KC_TSK | 3.28 | *4.50E-40* | *1.65E-35* |
| *NT5E* | KC_TSK | 3.10 | *3.99E-101* | *1.46E-96* |
| *IGFL1* | KC_TSK | 3.10 | *1.48E-49* | *5.43E-45* |
| *INHBA* | KC_TSK | 3.01 | *2.40E-160* | *8.77E-156* |
| *SERPINE1* | KC_TSK | 2.96 | *4.23E-80* | *1.55E-75* |
| *LAMA3* | KC_TSK | 2.90 | *1.12E-89* | *4.11E-85* |
| *CST6* | KC_Diff | 8.85 | *5.12E-235* | *1.87E-230* |
| *DEFB4A* | KC_Diff | 8.80 | *4.69E-197* | *1.72E-192* |
| *SPRR2B* | KC_Diff | 8.47 | *1.75E-113* | *6.40E-109* |
| *DEFB4B* | KC_Diff | 8.38 | *2.99E-147* | *1.09E-142* |
| *SLURP2* | KC_Diff | 8.21 | *6.41E-212* | *2.35E-207* |
| *KRT1* | KC_Diff | 7.61 | *3.36E-239* | *1.23E-234* |
| *KLK6* | KC_Diff | 7.45 | *1.51E-138* | *5.53E-134* |
| *SDR9C7* | KC_Diff | 7.28 | *5.43E-201* | *1.99E-196* |

| | | | | |
|---|---|---|---|---|
| *SPRR2E* | KC_Diff | 7.22 | *3.07E-245* | *1.12E-240* |
| *CNFN* | KC_Diff | 7.15 | *9.78E-166* | *3.58E-161* |

**Supplementary Table 3. Top activated pathways in PNI$^{pos}$ cSCC**

| Pathway | Contribution | Scaled Contribution | P-Value | Top Ligand-Receptor Pair |
|---|---|---|---|---|
| PTN | 0.3053 | 0.843 | *P<0.0001* | *PTN-NCL* |
| SAA | 0.1951 | 0.612 | *P<0.0001* | *SAA1-FPR2* |
| ***KIT*** | ***0.0204*** | ***0.257*** | ***P<0.0001*** | ***KITLG-KIT*** |
| AGRN | 0.0180 | 0.249 | *P<0.0001* | *AGRN-DAG1* |
| CD226 | 0.0124 | 0.228 | *P<0.0001* | *CD226-NECTIN2* |
| LIGHT | 0.0112 | 0.222 | *P<0.0001* | *TNFSF14-TNFRSF14* |
| PVR | 0.0084 | 0.209 | *P<0.0001* | *PVR-TIGIT* |
| SEMA5 | 0.0062 | 0.196 | *P<0.0001* | *SEMA5A-PLXNA1* |
| CSF | 0.0028 | 0.171 | *P<0.0001* | *CSF1-CSF1R* |
| CHEMERIN | 0.0024 | 0.165 | *P<0.0001* | *RARRES2-CMKLR1* |
| ACTIVIN | 0.0022 | 0.163 | *P<0.0001* | *INHBA - (ACVR1B and ACVR2A)* |
| NT | 0.0018 | 0.155 | *P<0.0001* | *NTF5-NTRK2* |
| SLURP | 0.0008 | 0.141 | *P<0.0001* | *SLURP21-CHRNB1* |
| CD137 | 0.0007 | 0.137 | *P<0.0001* | *TNFSF9-TNFRSF9* |
| NCAM | 0.0005 | 0.130 | *P<0.0001* | *NCAM1-FGFR1* |
| SEMA6 | 0.0003 | 0.125 | *P<0.0001* | *SEMA6B-PLXNA4* |
| FLT3 | 0.0001 | 0.102 | *P<0.0001* | *FLT3L-FLT3* |

**Supplementary Table 4. Phosphorylated proteins by histamine at 10 minutes**

| Proteins | Fold Change | P-value | Adjusted P-value |
|---|---|---|---|
| ROA1_HUMAN\|P09651_S364_S365 | 12.51409463 | 9.51E-04 | 1.16E-02 |
| ROA1_HUMAN\|P09651_S364_S365 | 8.76149993 | 3.22E-03 | 3.22E-02 |
| ZC3HD_HUMAN\|Q5T200_S1014_S1017 | 7.867639309 | 2.85E-15 | 7.73E-12 |
| ROA1_HUMAN\|P09651_S360_Y366 | 7.499270642 | 1.38E-08 | 4.97E-06 |
| SNUT1_HUMAN\|O43290_T430 | 6.541147197 | 5.40E-11 | 1.46E-07 |
| IF4B_HUMAN\|P23588_S348 | 6.088989326 | 2.91E-07 | 4.14E-05 |
| RL34_HUMAN\|P49207_S12 | 5.802045451 | 1.69E-08 | 3.53E-06 |
| SNUT1_HUMAN\|O43290_T430 | 5.770007536 | 1.85E-10 | 3.34E-07 |
| AMPD2_HUMAN\|Q01433_S100 | 5.135847568 | 2.58E-08 | 4.82E-06 |
| DC1L1_HUMAN\|Q9Y6G9_S412 | 5.103812424 | 8.39E-07 | 5.08E-05 |
| SUN2_HUMAN\|Q9UH99_S112_S113_T107 | 5.080480383 | 2.65E-07 | 8.69E-06 |
| PLEC_HUMAN\|Q15149_S4681 | 4.985978573 | 7.89E-06 | 3.37E-04 |
| SRC8_HUMAN\|Q14247_Y446 | 4.790850482 | 3.60E-07 | 4.16E-05 |
| HIRP3_HUMAN\|Q9BW71_S555 | 4.706407972 | 1.76E-06 | 4.04E-05 |
| KC1A_HUMAN\|P48729-3_T323 | 4.701643059 | 1.13E-05 | 4.32E-04 |
| DC1L1_HUMAN\|Q9Y6G9_S415 | 4.399176241 | 3.31E-07 | 2.60E-05 |
| PLEC_HUMAN\|Q15149_S4681 | 4.389257157 | 2.16E-05 | 5.34E-04 |
| ROA1_HUMAN\|P09651_S365 | 4.34015541 | 4.13E-04 | 6.15E-03 |
| MYPT1_HUMAN\|O14974_T696 | 4.331163693 | 5.38E-05 | 8.22E-04 |
| RL34_HUMAN\|P49207_S12 | 4.313691842 | 3.75E-07 | 5.29E-05 |
| SUN2_HUMAN\|Q9UH99_S112_S113_T107 | 4.309099235 | 1.46E-06 | 1.61E-04 |
| RS3A_HUMAN\|P61247_S154 | 4.287244022 | 3.70E-06 | 2.75E-04 |
| RL29_HUMAN\|P47914_S142 | 4.263740073 | 1.43E-04 | 2.81E-03 |
| MK01_HUMAN\|P28482_T185_Y187 | 4.254516699 | 4.90E-11 | 1.77E-08 |
| IF4B_HUMAN\|P23588_S348 | 4.238006922 | 6.83E-06 | 2.32E-04 |
| IWS1_HUMAN\|Q96ST2_S398 | 4.129149014 | 1.41E-08 | 1.07E-06 |
| HIRP3_HUMAN\|Q9BW71_S555 | 4.113078131 | 5.80E-06 | 9.79E-05 |
| RAI14_HUMAN\|Q9P0K7-4_S261 | 4.088593452 | 6.46E-14 | 3.50E-10 |

| | | | |
|---|---|---|---|
| RS3A_HUMAN\|P61247_S154 | 4.031066044 | 6.42E-06 | 1.07E-04 |
| TRI29_HUMAN\|Q14134_S166 | 4.00759812 | 6.34E-05 | 1.19E-03 |
| UBXN7_HUMAN\|O94888_S288 | 3.994069857 | 2.12E-07 | 3.59E-05 |
| T22D4_HUMAN\|Q9Y3Q8_T167 | 3.949224162 | 8.35E-13 | 6.47E-10 |
| EGFR_HUMAN\|P00533_Y1197 | 3.919193103 | 2.41E-07 | 2.08E-05 |
| RS3A_HUMAN\|P61247_S154 | 3.847079393 | 9.82E-06 | 3.07E-04 |
| ROA1_HUMAN\|P09651_S365 | 3.846483789 | 9.08E-04 | 1.36E-02 |
| MK14_HUMAN\|Q16539_T180_Y182 | 3.829016102 | 9.68E-06 | 3.07E-04 |
| DDX3X_HUMAN\|O00571_S612 | 3.80992326 | 1.65E-07 | 3.09E-05 |
| AHNK2_HUMAN\|Q8IVF2_S5175 | 3.789626914 | 5.42E-12 | 2.10E-09 |
| RSF1_HUMAN\|Q96T23_S1277_T1278 | 3.787207111 | 3.76E-05 | 4.25E-04 |
| ROA1_HUMAN\|P09651_S360_Y366 | 3.702374179 | 7.68E-06 | 5.07E-04 |
| TIA1_HUMAN\|P31483_T385 | 3.67225255 | 6.11E-06 | 2.83E-04 |
| MK03_HUMAN\|P27361_T198_T202 | 3.656077203 | 3.22E-11 | 1.34E-08 |
| CR025_HUMAN\|Q96B23_S66_S67_T74 | 3.571242841 | 1.69E-05 | 2.21E-04 |
| UBP5_HUMAN\|P45974_S785 | 3.558521173 | 2.05E-07 | 3.47E-05 |
| IF4B_HUMAN\|P23588_S348 | 3.550388656 | 3.54E-05 | 1.48E-03 |
| SRRM2_HUMAN\|Q9UQ35_S1152 | 3.527327627 | 1.08E-09 | 1.17E-06 |
| KC1A_HUMAN\|P48729-3_T323 | 3.507561807 | 1.48E-04 | 2.23E-03 |
| ROA1_HUMAN\|P09651_S365 | 3.503759296 | 1.67E-03 | 1.36E-02 |
| PLEC_HUMAN\|Q15149_S4681 | 3.495921709 | 1.40E-04 | 3.75E-03 |
| ITA2_HUMAN\|P17301_S1180 | 3.493705323 | 3.16E-11 | 2.86E-08 |
| AHNK_HUMAN\|Q09666_S5739 | 3.472553728 | 4.90E-10 | 1.21E-07 |
| MAP4_HUMAN\|P27816_S1145 | 3.437924233 | 2.36E-03 | 2.32E-02 |
| SRRM2_HUMAN\|Q9UQ35_S1152 | 3.430256994 | 1.61E-09 | 1.59E-06 |
| TM230_HUMAN\|Q96A57_T25 | 3.424389869 | 4.43E-10 | 6.15E-08 |
| MTAP2_HUMAN\|P11137_T1602_T1605 | 3.416755219 | 1.94E-08 | 1.24E-06 |
| SF3B2_HUMAN\|Q13435_S307_T311 | 3.398261811 | 3.87E-05 | 8.17E-04 |
| CRBG1_HUMAN\|Q9Y4K1_S410 | 3.374027784 | 1.38E-08 | 2.27E-06 |
| ROA1_HUMAN\|P09651_S360_Y366 | 3.356536996 | 2.06E-05 | 7.98E-04 |
| KC1A_HUMAN\|P48729-3_T323 | 3.347452556 | 2.24E-04 | 5.13E-03 |

| | | | |
|---|---|---|---|
| UTP18_HUMAN\|Q9Y5J1_S121 | 3.342573673 | 8.36E-06 | 1.29E-04 |
| ROA1_HUMAN\|P09651_S363 | 3.342226709 | 1.25E-06 | 9.65E-05 |
| TIA1_HUMAN\|P31483_T385 | 3.315750317 | 1.70E-05 | 4.57E-04 |
| TRI22_HUMAN\|Q8IYM9_S384 | 3.307446101 | 6.52E-11 | 4.42E-08 |
| CCDC9_HUMAN\|Q9Y3X0_S376_S386_T381 | 3.304758561 | 8.30E-12 | 5.00E-09 |
| ROA1_HUMAN\|P09651_S364 | 3.304298646 | 1.22E-07 | 2.64E-05 |
| G3BP1_HUMAN\|Q13283_S250 | 3.275318067 | 2.64E-06 | 1.54E-04 |
| HIRP3_HUMAN\|Q9BW71_S555 | 3.274734444 | 4.84E-05 | 9.65E-04 |
| LMNA_HUMAN\|P02545_S636 | 3.271062792 | 3.17E-04 | 5.02E-03 |
| SF3B2_HUMAN\|Q13435_S307_S309 | 3.270817821 | 1.55E-07 | 1.56E-05 |
| UBXN7_HUMAN\|O94888_S288 | 3.268572639 | 2.21E-06 | 1.36E-04 |
| ITA2_HUMAN\|P17301_S1180 | 3.266908593 | 1.18E-10 | 1.28E-07 |
| AHNK_HUMAN\|Q09666_S5746_S5752 | 3.262793185 | 3.80E-13 | 5.15E-10 |
| NOLC1_HUMAN\|Q14978_S688_S698 | 3.257359709 | 1.66E-05 | 5.41E-04 |
| K2C5_HUMAN\|P13647_S26_T19 | 3.24416315 | 1.57E-05 | 5.20E-04 |
| CR025_HUMAN\|Q96B23_S143_S145 | 3.227311667 | 2.04E-13 | 2.47E-10 |
| CE170_HUMAN\|Q5SW79_S633_S636 | 3.198284558 | 8.11E-08 | 1.91E-05 |
| CR025_HUMAN\|Q96B23_S66_S67_T74 | 3.198250349 | 4.99E-05 | 5.33E-04 |
| UTP18_HUMAN\|Q9Y5J1_S121 | 3.198210007 | 1.33E-05 | 1.85E-04 |
| SRC8_HUMAN\|Q14247_S405_S418_T401 | 3.197893466 | 3.86E-08 | 8.72E-06 |
| AHNK2_HUMAN\|Q8IVF2_S5175 | 3.187897126 | 1.09E-10 | 2.95E-07 |
| FOSL2_HUMAN\|P15408_S19 | 3.160774219 | 2.83E-08 | 1.67E-06 |
| LPP_HUMAN\|Q93052_S609 | 3.159461582 | 7.71E-09 | 1.49E-06 |
| FUBP2_HUMAN\|Q92945_T195 | 3.1410848 | 8.36E-07 | 5.08E-05 |
| TRI22_HUMAN\|Q8IYM9_S383 | 3.127832473 | 6.60E-08 | 6.51E-06 |
| MET_HUMAN\|P08581_S990_S997 | 3.114058736 | 3.56E-05 | 3.96E-04 |
| AHNK_HUMAN\|Q09666_S5745_S5749 | 3.103003581 | 1.70E-13 | 3.08E-10 |
| SRC8_HUMAN\|Q14247_Y446 | 3.099403774 | 2.93E-05 | 6.54E-04 |
| PHF14_HUMAN\|O94880_T287 | 3.098624909 | 3.50E-10 | 4.74E-07 |
| MDC1_HUMAN\|Q14676_S453 | 3.093199661 | 3.91E-04 | 2.80E-03 |
| REQU_HUMAN\|Q92785_T149 | 3.087062593 | 1.54E-10 | 2.78E-07 |

| | | | |
|---|---|---|---|
| MYPT1_HUMAN|O14974_T696 | 3.084522451 | 7.49E-04 | 2.45E-02 |
| HEAT6_HUMAN|Q6AI08_S395_S397 | 3.082215723 | 3.98E-06 | 7.36E-05 |
| PKP3_HUMAN|Q9Y446_S180_S183 | 3.077145221 | 2.06E-09 | 1.59E-06 |
| ANXA2_HUMAN|P07355_S26 | 3.067230092 | 6.55E-05 | 1.58E-03 |
| 4EBP1_HUMAN|Q13541_S112 | 3.054502743 | 5.68E-06 | 2.78E-04 |
| ROA1_HUMAN|P09651_S362_S365 | 3.051365331 | 3.25E-05 | 9.41E-04 |
| EF2K_HUMAN|O00418_S470 | 3.045839288 | 1.08E-10 | 1.15E-07 |
| IF4B_HUMAN|P23588_S348 | 3.044571225 | 1.54E-04 | 2.66E-02 |
| ROA1_HUMAN|P09651_S360_Y366 | 3.022933797 | 6.05E-05 | 1.15E-03 |
| RL29_HUMAN|P47914_S142 | 3.022783041 | 1.76E-03 | 1.41E-02 |
| IF4B_HUMAN|P23588_S348 | 3.021627764 | 1.66E-04 | 4.46E-03 |
| COHA1_HUMAN|Q9UMD9_S76_S79 | 3.02161947 | 7.99E-07 | 3.61E-05 |
| MAP4_HUMAN|P27816_S1145 | 3.01932658 | 5.34E-03 | 3.23E-02 |
| RL34_HUMAN|P49207_S12 | 3.019325676 | 2.18E-05 | 2.73E-04 |
| DBNL_HUMAN|Q9UJU6_S269 | 3.01884898 | 9.07E-09 | 3.78E-06 |
| MDC1_HUMAN|Q14676_S453 | 3.014667844 | 4.89E-04 | 3.11E-03 |
| PLEC_HUMAN|Q15149_S4681 | 3.013058129 | 4.94E-04 | 9.66E-03 |
| EPHA2_HUMAN|P29317_Y772 | 3.011896021 | 5.59E-06 | 2.08E-04 |
| ZC3HD_HUMAN|Q5T200_S1014_S1017 | 3.010837226 | 1.03E-09 | 1.87E-06 |
| PABP1_HUMAN|P11940_S629 | 3.010587054 | 2.18E-06 | 1.36E-04 |
| K2C6B_HUMAN|P04259_S37 | 2.986577644 | 1.19E-08 | 5.40E-06 |
| ZO1_HUMAN|Q07157_T868_Y843 | 2.971505178 | 2.48E-07 | 8.63E-06 |
| HIRP3_HUMAN|Q9BW71_S551 | 2.970319278 | 6.73E-07 | 1.80E-05 |
| ROA1_HUMAN|P09651_S361_S364 | 2.96754819 | 1.05E-08 | 4.06E-06 |
| LPP_HUMAN|Q93052_S609 | 2.967389667 | 1.98E-08 | 7.15E-06 |
| NIBL1_HUMAN|Q96TA1_S641_S646 | 2.966275825 | 6.58E-08 | 1.37E-05 |
| TV23B_HUMAN|Q9NYZ1_T199 | 2.965722713 | 3.22E-06 | 6.26E-05 |
| SRC8_HUMAN|Q14247_Y446 | 2.952956786 | 4.94E-05 | 1.91E-03 |
| ROA1_HUMAN|P09651_S360_Y366 | 2.951417872 | 7.77E-05 | 2.79E-03 |
| EIF3H_HUMAN|O15372_S183 | 2.942010456 | 4.80E-08 | 1.25E-05 |
| EIF3H_HUMAN|O15372_S183 | 2.939767146 | 4.86E-08 | 7.11E-06 |

| | | | |
|---|---|---|---|
| IWS1_HUMAN\|Q96ST2_S398 | 2.939351054 | 8.69E-07 | 2.19E-05 |
| RL29_HUMAN\|P47914_S142 | 2.929587403 | 2.21E-03 | 2.50E-02 |
| ERBB2_HUMAN\|P04626_S998 | 2.927586746 | 4.47E-08 | 4.85E-06 |
| KC1A_HUMAN\|P48729_S312 | 2.925393576 | 1.97E-06 | 1.35E-04 |
| KPCD2_HUMAN\|Q9BZL6_S710 | 2.924830999 | 1.59E-06 | 1.39E-04 |
| IF4B_HUMAN\|P23588_S352 | 2.905694387 | 2.03E-04 | 3.71E-03 |
| RSF1_HUMAN\|Q96T23_S1277_T1278 | 2.896354465 | 3.98E-04 | 2.65E-03 |
| HSPB1_HUMAN\|P04792_S78_S82 | 2.892022353 | 6.47E-08 | 8.25E-06 |
| KMT2A_HUMAN\|Q03164_T3028 | 2.885144148 | 1.20E-06 | 4.66E-05 |
| PLEC_HUMAN\|Q15149_S4406 | 2.883790471 | 1.10E-05 | 6.16E-04 |
| EGFR_HUMAN\|P00533_Y1197 | 2.878190874 | 1.08E-05 | 4.29E-04 |
| HIRP3_HUMAN\|Q9BW71_S551 | 2.876831375 | 1.04E-06 | 2.65E-05 |
| EF2K_HUMAN\|O00418_S470 | 2.875333123 | 3.07E-10 | 2.38E-07 |
| FOXO3_HUMAN\|O43524_S413 | 2.865588099 | 5.15E-07 | 3.58E-05 |
| RAI14_HUMAN\|Q9P0K7_S290 | 2.863405896 | 4.92E-08 | 4.44E-05 |
| HIRP3_HUMAN\|Q9BW71_S555 | 2.861893552 | 1.79E-04 | 4.65E-03 |
| K2C5_HUMAN\|P13647_S21_T19 | 2.85816279 | 1.96E-07 | 3.44E-05 |
| DPYL3_HUMAN\|Q14195_S568 | 2.847277547 | 3.54E-07 | 4.16E-05 |
| SMCR8_HUMAN\|Q8TEV9_S417 | 2.842732452 | 4.55E-05 | 1.35E-03 |
| EIF3H_HUMAN\|O15372_S183 | 2.841011986 | 8.45E-08 | 1.76E-05 |
| NIPA4_HUMAN\|Q0D2K0_S427 | 2.830770483 | 3.33E-05 | 3.79E-04 |
| TIA1_HUMAN\|P31483_T385 | 2.826560091 | 8.92E-05 | 2.73E-03 |
| SP100_HUMAN\|P23497_S407_S410 | 2.823341913 | 7.50E-05 | 7.52E-04 |
| COBL1_HUMAN\|Q53SF7_T322 | 2.821272928 | 2.93E-11 | 1.46E-07 |
| CAVN1_HUMAN\|Q6NZI2_Y308 | 2.820125328 | 1.73E-06 | 8.84E-05 |
| GGYF2_HUMAN\|Q6Y7W6_S201 | 2.813479482 | 1.06E-09 | 5.25E-07 |
| IF4B_HUMAN\|P23588_S352 | 2.811984587 | 2.80E-04 | 3.65E-03 |
| REQU_HUMAN\|Q92785_T149 | 2.80982394 | 8.95E-10 | 1.21E-06 |
| DBNL_HUMAN\|Q9UJU6_T270 | 2.808101653 | 3.92E-08 | 1.12E-05 |
| NEB2_HUMAN\|Q96SB3_S100 | 2.806354962 | 1.32E-07 | 1.35E-05 |
| ANXA2_HUMAN\|P07355_S26 | 2.800335161 | 1.62E-04 | 2.39E-03 |

| | | | |
|---|---|---|---|
| PAWR_HUMAN\|Q96IZ0_S231 | 2.798937929 | 2.43E-11 | 6.58E-08 |
| ROA1_HUMAN\|P09651_S362_S365 | 2.78663352 | 8.73E-05 | 2.69E-03 |
| HMGA2_HUMAN\|P52926_S105 | 2.784811056 | 2.30E-06 | 4.95E-05 |
| AHNK_HUMAN\|Q09666_S5397 | 2.78456811 | 1.37E-05 | 6.10E-04 |
| LMNA_HUMAN\|P02545_S22_T19 | 2.77931192 | 7.35E-06 | 3.07E-03 |
| TP53B_HUMAN\|Q12888_S356 | 2.774662579 | 1.64E-06 | 1.18E-04 |
| NIBL1_HUMAN\|Q96TA1_S641_S646 | 2.770623496 | 1.76E-07 | 6.58E-06 |
| NIPA_HUMAN\|Q86WB0_S408_S409 | 2.769176872 | 2.05E-12 | 1.39E-09 |
| SF3B2_HUMAN\|Q13435_S303_S309 | 2.764450898 | 3.88E-06 | 1.64E-04 |
| EGFR_HUMAN\|P00533_Y1197 | 2.759832476 | 1.85E-05 | 9.20E-04 |
| JAM1_HUMAN\|Q9Y624_S296 | 2.742276762 | 2.77E-05 | 8.24E-04 |
| RS3A_HUMAN\|P61247_S154 | 2.737843016 | 2.53E-04 | 1.28E-02 |
| TB10A_HUMAN\|Q9BXI6_S40_S45 | 2.735865378 | 7.28E-06 | 1.17E-04 |
| EPHA2_HUMAN\|P29317_Y772 | 2.735535568 | 2.00E-05 | 6.40E-04 |
| SP100_HUMAN\|P23497_S407_S410 | 2.726945568 | 1.11E-04 | 9.88E-04 |
| STMN1_HUMAN\|P16949_S25 | 2.725017274 | 2.30E-11 | 2.49E-08 |
| PKP3_HUMAN\|Q9Y446_S180_S183 | 2.723750209 | 1.49E-08 | 6.20E-06 |
| I2BPL_HUMAN\|Q9H1B7_S659 | 2.71455656 | 1.23E-05 | 2.71E-04 |
| K1C18_HUMAN\|P05783_S7 | 2.713719091 | 6.77E-08 | 5.76E-06 |
| CCAR2_HUMAN\|Q8N163_S678 | 2.713178472 | 1.14E-05 | 1.62E-04 |
| B4DKV4_HUMAN\|B4DKV4_S19_S22 | 2.711476288 | 1.77E-06 | 1.47E-04 |
| ZO1_HUMAN\|Q07157_T868_Y843 | 2.709613168 | 9.54E-07 | 2.39E-05 |
| HIRP3_HUMAN\|Q9BW71_S551_S555 | 2.708799654 | 9.51E-06 | 1.45E-04 |
| NGAP_HUMAN\|Q9UJF2_S946_S954 | 2.707449172 | 2.00E-04 | 1.56E-03 |
| IASPP_HUMAN\|Q8WUF5_T341 | 2.697036832 | 9.25E-06 | 3.78E-04 |
| CBPC1_HUMAN\|Q9UPW5_T1170 | 2.696186803 | 3.94E-05 | 4.29E-04 |
| COHA1_HUMAN\|Q9UMD9_S76_S79 | 2.693945447 | 3.85E-06 | 2.04E-04 |
| TP53B_HUMAN\|Q12888_S356 | 2.692812247 | 2.45E-06 | 1.87E-04 |
| RL1D1_HUMAN\|O76021_T465 | 2.692604842 | 1.95E-07 | 1.51E-04 |
| HDGF_HUMAN\|P51858_S103 | 2.689962333 | 1.22E-05 | 4.48E-04 |
| ITA2_HUMAN\|P17301_S1181 | 2.689182252 | 1.60E-08 | 2.47E-06 |

| | | | |
|---|---|---|---|
| EF2K_HUMAN\|O00418_S470 | 2.684164756 | 1.12E-09 | 6.09E-06 |
| MTAP2_HUMAN\|P11137_T1602_T1605 | 2.677488848 | 7.09E-07 | 1.99E-05 |
| EGFR_HUMAN\|P00533_S1039 | 2.668139526 | 1.67E-04 | 1.35E-03 |
| MK01_HUMAN\|P28482_T185_Y187 | 2.661591197 | 4.99E-08 | 1.23E-05 |
| DC1L1_HUMAN\|Q9Y6G9_S415 | 2.660172766 | 6.34E-05 | 6.55E-04 |
| DDX3X_HUMAN\|O00571_S612 | 2.654327326 | 1.22E-05 | 6.90E-04 |
| PKP3_HUMAN\|Q9Y446_S223 | 2.648484798 | 1.24E-04 | 2.78E-03 |
| TV23B_HUMAN\|Q9NYZ1_T199 | 2.639440244 | 1.61E-05 | 4.28E-04 |
| ITA2_HUMAN\|P17301_S1180 | 2.639175688 | 1.00E-08 | 3.38E-06 |
| K2C6A_HUMAN\|P02538_S37 | 2.63799199 | 7.57E-05 | 1.71E-03 |
| RS3A_HUMAN\|P61247_S154 | 2.636906528 | 3.67E-04 | 4.45E-03 |
| PHF14_HUMAN\|O94880_T287 | 2.625983416 | 5.88E-09 | 3.14E-06 |
| MEF2D_HUMAN\|Q14814_S231 | 2.621318204 | 6.40E-07 | 3.04E-05 |
| K2C5_HUMAN\|P13647_S18_S21 | 2.61859927 | 2.68E-06 | 1.54E-04 |
| AHNK_HUMAN\|Q09666_S5749_S5752 | 2.617218994 | 9.12E-11 | 3.09E-08 |
| MK14_HUMAN\|Q16539_Y182 | 2.616977971 | 5.77E-09 | 1.56E-06 |
| LMNA_HUMAN\|P02545_S616_S619 | 2.612812286 | 3.20E-04 | 3.33E-03 |
| AHNK2_HUMAN\|Q8IVF2_S457 | 2.603719685 | 2.10E-06 | 1.35E-04 |
| PKP3_HUMAN\|Q9Y446_S223 | 2.602848102 | 1.50E-04 | 2.90E-03 |
| CE170_HUMAN\|Q5SW79_S633_S636 | 2.602061585 | 1.56E-06 | 1.39E-04 |
| FKBP5_HUMAN\|Q13451_S445 | 2.593735814 | 2.86E-05 | 1.28E-03 |
| FOSL2_HUMAN\|P15408_S19 | 2.592665524 | 9.24E-07 | 2.43E-05 |
| AMPD2_HUMAN\|Q01433_S100 | 2.591107031 | 3.70E-05 | 4.35E-03 |
| DFFA_HUMAN\|O00273-2_S257 | 2.585554715 | 1.44E-05 | 4.92E-04 |
| LAD1_HUMAN\|O00515_S38_S39 | 2.581657311 | 8.44E-07 | 4.53E-05 |
| LMNA_HUMAN\|P02545_S618 | 2.5774348 | 5.37E-05 | 1.35E-03 |
| LPP_HUMAN\|Q93052_S609 | 2.575581422 | 1.87E-07 | 3.27E-05 |
| ARHG5_HUMAN\|Q12774_S445_S450 | 2.572707911 | 5.65E-12 | 1.53E-08 |
| SNX6_HUMAN\|Q9UNH7_S316 | 2.570509767 | 2.47E-04 | 1.94E-03 |
| COBL1_HUMAN\|Q53SF7_S294_T298 | 2.562605121 | 1.11E-08 | 5.40E-06 |
| DC1L1_HUMAN\|Q9Y6G9_S412 | 2.561799977 | 4.81E-04 | 3.35E-03 |

| | | | |
|---|---|---|---|
| TV23B_HUMAN\|Q9NYZ1_T199 | 2.556675494 | 2.51E-05 | 3.00E-04 |
| DKC1_HUMAN\|O60832_S485 | 2.555751285 | 1.19E-09 | 5.37E-07 |
| RSF1_HUMAN\|Q96T23_S1277_T1278 | 2.554932531 | 1.23E-03 | 1.16E-02 |
| PR38B_HUMAN\|Q5VTL8_S529 | 2.554773626 | 3.49E-10 | 9.46E-08 |
| B4DKV4_HUMAN\|B4DKV4_S22 | 2.552874342 | 7.13E-06 | 3.87E-04 |
| MET_HUMAN\|P08581_S990_S997 | 2.551606285 | 3.05E-04 | 6.74E-03 |
| SARG_HUMAN\|Q9BW04_T151 | 2.551137927 | 8.45E-06 | 4.30E-04 |
| PAK2_HUMAN\|Q13177_S197 | 2.549726815 | 4.32E-05 | 1.15E-03 |
| IASPP_HUMAN\|Q8WUF5_T341 | 2.549469843 | 1.91E-05 | 3.73E-04 |
| AHNK2_HUMAN\|Q8IVF2_S1710 | 2.54390845 | 1.11E-08 | 8.15E-07 |
| CR025_HUMAN\|Q96B23_S143_S145 | 2.537094426 | 5.89E-11 | 9.86E-08 |
| AKTS1_HUMAN\|Q96B36_S203 | 2.5345411 | 7.60E-06 | 1.21E-04 |
| EF2K_HUMAN\|O00418_S470 | 2.533905143 | 3.47E-09 | 2.35E-06 |
| TTP_HUMAN\|P26651_S60 | 2.533096329 | 8.99E-10 | 5.25E-07 |
| TM230_HUMAN\|Q96A57_T25 | 2.52596226 | 1.69E-07 | 1.67E-05 |
| AMPD2_HUMAN\|Q01433_S100 | 2.522574391 | 5.11E-05 | 1.78E-03 |
| TIA1_HUMAN\|P31483_T385 | 2.522065392 | 3.01E-04 | 6.63E-03 |
| MAGD2_HUMAN\|Q9UNF1_S190_S191 | 2.520337486 | 2.09E-05 | 2.65E-04 |
| SUN2_HUMAN\|Q9UH99_S112_S113_T107 | 2.518992619 | 5.10E-04 | 3.51E-03 |
| MTAP2_HUMAN\|P11137_T1602_T1605 | 2.517965598 | 1.86E-06 | 2.09E-04 |
| DC1I2_HUMAN\|Q13409-2_S81 | 2.51384975 | 1.03E-06 | 1.10E-04 |
| EF2_HUMAN\|P13639_T57 | 2.51299329 | 2.69E-05 | 1.24E-03 |
| MPZL1_HUMAN\|O95297_S221 | 2.512763674 | 3.45E-05 | 3.95E-04 |
| RYBP_HUMAN\|Q8N488_T186 | 2.512730061 | 2.49E-03 | 1.15E-02 |
| COTL1_HUMAN\|Q14019_T141 | 2.511213944 | 4.66E-05 | 1.22E-03 |
| E7ETA6_HUMAN\|E7ETA6_T1557 | 2.511148955 | 1.33E-06 | 4.96E-05 |
| HIRP3_HUMAN\|Q9BW71_S551_S555 | 2.510822371 | 2.56E-05 | 3.09E-04 |
| EPHA2_HUMAN\|P29317_T771 | 2.506672753 | 3.77E-05 | 1.06E-03 |
| K2C6A_HUMAN\|P02538_S37 | 2.503632347 | 1.50E-04 | 3.23E-03 |
| ROA1_HUMAN\|P09651_S362_S365 | 2.502462153 | 2.87E-04 | 3.72E-03 |
| SNUT1_HUMAN\|O43290_T430 | 2.501384917 | 5.12E-06 | 1.38E-04 |

| | | | |
|---|---|---|---|
| RTKN_HUMAN\|Q9BST9_S520_S529 | 2.501288974 | 2.78E-08 | 3.36E-06 |
| RU17_HUMAN\|P08621_S320 | 2.498869627 | 6.06E-06 | 3.53E-04 |
| 1433Z_HUMAN\|P63104_T232 | 2.498824943 | 5.41E-07 | 5.43E-05 |
| K2C5_HUMAN\|P13647_S18_S21 | 2.495646161 | 5.64E-06 | 1.48E-04 |
| RL23_HUMAN\|P62829_S138 | 2.492548882 | 9.50E-05 | 1.65E-03 |
| ENAH_HUMAN\|Q8N8S7_T489 | 2.49219042 | 7.04E-05 | 9.92E-04 |
| IASPP_HUMAN\|Q8WUF5_S526 | 2.490296599 | 2.08E-05 | 5.20E-04 |
| TBC9B_HUMAN\|Q66K14_S411 | 2.488143198 | 2.71E-07 | 1.47E-04 |
| AKTS1_HUMAN\|Q96B36_S203 | 2.485550453 | 1.01E-05 | 6.75E-04 |
| RL34_HUMAN\|P49207_S12 | 2.481168207 | 2.26E-04 | 1.22E-02 |
| K2C5_HUMAN\|P13647_S18_S21 | 2.474409446 | 6.44E-06 | 3.60E-04 |
| UBP5_HUMAN\|P45974_S779 | 2.472824974 | 3.48E-07 | 4.16E-05 |
| RAI14_HUMAN\|Q9P0K7_T297 | 2.470769947 | 2.07E-07 | 1.37E-05 |
| K2C75_HUMAN\|O95678_S61 | 2.469681224 | 2.12E-06 | 1.71E-04 |
| CAVN1_HUMAN\|Q6NZI2_Y308 | 2.468960706 | 1.08E-05 | 6.48E-04 |
| RAI14_HUMAN\|Q9P0K7_S281_S290 | 2.468892547 | 3.82E-06 | 1.48E-04 |
| ROA1_HUMAN\|P09651_S363 | 2.467615768 | 5.78E-05 | 2.07E-03 |
| LARP1_HUMAN\|Q6PKG0_S766 | 2.46465902 | 3.65E-05 | 1.03E-03 |
| CT2NL_HUMAN\|Q9P2B4_S443 | 2.464458209 | 3.38E-04 | 6.78E-03 |
| K2C5_HUMAN\|P13647_S16_S26 | 2.462052009 | 5.85E-06 | 2.78E-04 |
| AMPD2_HUMAN\|Q01433_S100 | 2.460544564 | 6.93E-05 | 7.12E-04 |
| OSBL1_HUMAN\|Q9BXW6_S472_S481 | 2.457238834 | 2.34E-04 | 1.77E-03 |
| K1C14_HUMAN\|P02533_S32 | 2.446664794 | 7.68E-10 | 1.74E-07 |
| RBM7_HUMAN\|Q9Y580_S136 | 2.445284035 | 7.67E-08 | 3.58E-06 |
| MTAP2_HUMAN\|P11137_S1782_S1790 | 2.444278011 | 9.12E-06 | 1.35E-04 |
| LAD1_HUMAN\|O00515_S38_T40_T41 | 2.444130167 | 6.98E-08 | 6.76E-06 |
| TMF1_HUMAN\|P82094_S72_S77 | 2.443745045 | 8.75E-07 | 4.65E-05 |
| RTKN_HUMAN\|Q9BST9_S529_T518 | 2.443538563 | 2.17E-11 | 9.80E-09 |
| JAM1_HUMAN\|Q9Y624_S296 | 2.443463517 | 1.27E-04 | 3.46E-03 |
| DC1L1_HUMAN\|Q9Y6G9_S412 | 2.442732021 | 7.74E-04 | 2.47E-02 |
| NGAP_HUMAN\|Q9UJF2_S946_S954 | 2.442346357 | 5.80E-04 | 1.06E-02 |

| | | | |
|---|---|---|---|
| SEPT7_HUMAN|Q16181_S423 | 2.436305802 | 1.16E-06 | 9.22E-05 |
| AHNK2_HUMAN|Q8IVF2_S5175 | 2.435046678 | 2.10E-08 | 1.46E-06 |
| MK14_HUMAN|Q16539_T180_Y182 | 2.43464105 | 8.12E-04 | 5.18E-03 |
| SRRM1_HUMAN|Q8IYB3_S738_S743 | 2.433101813 | 1.59E-05 | 6.61E-04 |
| CR025_HUMAN|Q96B23_S66_S67_T74 | 2.431334906 | 8.05E-04 | 8.28E-03 |
| CCDC9_HUMAN|Q9Y3X0_S386_S390 | 2.429406263 | 3.72E-06 | 1.60E-04 |
| PARD3_HUMAN|Q8TEW0_S220 | 2.428492692 | 2.05E-05 | 6.54E-04 |
| K2C5_HUMAN|P13647_S21_T19 | 2.427565176 | 2.91E-06 | 1.62E-04 |
| AHNK2_HUMAN|Q8IVF2_S5175 | 2.425951296 | 2.27E-08 | 5.87E-06 |
| AHNK2_HUMAN|Q8IVF2_S1710 | 2.425134082 | 2.96E-08 | 6.98E-06 |
| MA7D1_HUMAN|Q3KQU3_S254 | 2.424853735 | 1.00E-10 | 3.20E-08 |
| IASPP_HUMAN|Q8WUF5_T341 | 2.423341845 | 3.72E-05 | 1.17E-03 |
| NELFE_HUMAN|P18615_S251 | 2.423232277 | 5.47E-07 | 3.71E-05 |
| CPSF4_HUMAN|O95639_S202 | 2.42291527 | 2.82E-07 | 1.99E-05 |
| SPAS2_HUMAN|Q86XZ4_S538 | 2.421763326 | 1.91E-05 | 4.94E-04 |
| EPHA2_HUMAN|P29317_T771 | 2.420847535 | 6.16E-05 | 1.17E-03 |
| PABP1_HUMAN|P11940_S629 | 2.418200895 | 4.00E-05 | 1.82E-03 |
| MK14_HUMAN|Q16539_Y182 | 2.417332592 | 2.76E-08 | 1.74E-06 |
| RL34_HUMAN|P49207_S12 | 2.417027094 | 3.09E-04 | 3.90E-03 |
| UFD1_HUMAN|Q92890_S299 | 2.416561517 | 4.13E-06 | 2.15E-04 |
| HEAT6_HUMAN|Q6AI08_S395_S397 | 2.412559044 | 8.32E-05 | 7.79E-04 |
| COBL1_HUMAN|Q53SF7_S294_T298 | 2.411580189 | 3.44E-08 | 3.84E-06 |
| SF3B2_HUMAN|Q13435_S307_T311 | 2.406814449 | 1.08E-03 | 1.63E-02 |
| GRSF1_HUMAN|Q12849_S118 | 2.404233197 | 3.31E-09 | 3.22E-07 |
| PAWR_HUMAN|Q96IZ0_S231 | 2.401651074 | 1.01E-09 | 5.25E-07 |
| MFF_HUMAN|Q9GZY8_S157 | 2.401512032 | 6.31E-07 | 7.24E-05 |
| RL34_HUMAN|P49207_S12 | 2.400488379 | 3.36E-04 | 6.94E-03 |
| NOLC1_HUMAN|Q14978_S688_S698 | 2.399738777 | 4.30E-04 | 8.38E-03 |
| K1C14_HUMAN|P02533_S44 | 2.397822484 | 1.50E-07 | 1.07E-05 |
| CAMP2_HUMAN|Q08AD1_S691 | 2.395324448 | 5.32E-07 | 5.43E-05 |
| ARM10_HUMAN|Q8N2F6-2_S45 | 2.393113254 | 3.02E-03 | 4.45E-02 |

| | | | |
|---|---|---|---|
| RU17_HUMAN\|P08621_S320 | 2.392962262 | 1.13E-05 | 4.32E-04 |
| SF3B2_HUMAN\|Q13435_S307_S309 | 2.392465747 | 1.12E-05 | 1.59E-04 |
| RENT1_HUMAN\|Q92900_S1107_T1111 | 2.386580762 | 8.49E-06 | 4.30E-04 |
| KC1A_HUMAN\|P48729_S312 | 2.386233156 | 4.38E-05 | 1.76E-03 |
| HOME3_HUMAN\|Q9NSC5_S159 | 2.38542718 | 1.55E-05 | 4.23E-04 |
| K2C6B_HUMAN\|P04259_S37 | 2.383983431 | 8.35E-07 | 7.24E-05 |
| ACAP2_HUMAN\|Q15057_S381_S384 | 2.382056761 | 1.73E-08 | 6.70E-06 |
| RHG29_HUMAN\|Q52LW3_S176 | 2.380273911 | 1.68E-06 | 7.71E-05 |
| S20A2_HUMAN\|Q08357_Y423 | 2.379636239 | 2.16E-06 | 1.72E-04 |
| ZC3HD_HUMAN\|Q5T200_S1014_S1017 | 2.379068479 | 7.58E-08 | 3.16E-05 |
| IWS1_HUMAN\|Q96ST2_S398 | 2.377776263 | 1.60E-05 | 4.35E-04 |
| DC1L1_HUMAN\|Q9Y6G9_S412 | 2.37673311 | 1.02E-03 | 1.46E-02 |
| MK14_HUMAN\|Q16539_T180_Y182 | 2.374720763 | 1.04E-03 | 1.48E-02 |
| NIBL1_HUMAN\|Q96TA1_S692_S696 | 2.374181244 | 4.18E-08 | 9.06E-06 |
| IWS1_HUMAN\|Q96ST2_S398 | 2.372993796 | 1.64E-05 | 4.32E-04 |
| EF2_HUMAN\|P13639_T57 | 2.370159542 | 5.86E-05 | 2.50E-03 |
| GRAP1_HUMAN\|Q4V328_S666_S669 | 2.368734548 | 1.14E-04 | 2.59E-03 |
| JAM1_HUMAN\|Q9Y624_S296 | 2.368135033 | 1.92E-04 | 2.69E-03 |
| PKP3_HUMAN\|Q9Y446_S314 | 2.36523662 | 1.71E-09 | 1.54E-06 |
| SF3A3_HUMAN\|Q12874_S365_S367_S369 | 2.361949283 | 1.17E-04 | 2.63E-03 |
| RAI14_HUMAN\|Q9P0K7_S281_S290 | 2.360613612 | 7.65E-06 | 1.21E-04 |
| RRBP1_HUMAN\|Q9P2E9_T225 | 2.358362218 | 1.32E-06 | 6.51E-04 |
| UBXN7_HUMAN\|O94888_S288 | 2.355969436 | 1.36E-04 | 1.69E-03 |
| RBM7_HUMAN\|Q9Y580_S136 | 2.355525036 | 1.58E-07 | 6.06E-06 |
| TRI29_HUMAN\|Q14134_S166 | 2.355248787 | 5.04E-03 | 2.29E-02 |
| GYS1_HUMAN\|P13807_S727_S731 | 2.355245096 | 2.90E-07 | 6.29E-05 |
| K1C14_HUMAN\|P02533_S54_Y46 | 2.35486499 | 1.57E-07 | 3.04E-05 |
| K2C75_HUMAN\|O95678_S61 | 2.35356105 | 4.57E-06 | 2.31E-04 |
| DPYL2_HUMAN\|Q16555_S570 | 2.353514162 | 7.85E-06 | 3.37E-04 |
| SF3B2_HUMAN\|Q13435_S307_S309 | 2.35302066 | 1.43E-05 | 7.44E-04 |
| PHLB2_HUMAN\|Q86SQ0_S334_T328 | 2.350818929 | 2.60E-06 | 1.93E-04 |

| | | | |
|---|---|---|---|
| K1C14_HUMAN\|P02533_S44 | 2.35028897 | 2.29E-07 | 3.76E-05 |
| K2C6A_HUMAN\|P02538_S37 | 2.350121304 | 3.42E-04 | 3.52E-03 |
| K2C5_HUMAN\|P13647_S21_S26 | 2.349348238 | 1.39E-07 | 2.79E-05 |
| ZC3HD_HUMAN\|Q5T200_S1014_S1017 | 2.349080079 | 9.71E-08 | 1.93E-05 |
| COTL1_HUMAN\|Q14019_T141 | 2.348427077 | 1.10E-04 | 1.81E-03 |
| K2C5_HUMAN\|P13647_S21_T19 | 2.348071383 | 5.12E-06 | 1.38E-04 |
| SCAM2_HUMAN\|O15127_S319 | 2.347521511 | 8.44E-07 | 5.08E-05 |
| AHNK_HUMAN\|Q09666_T5824 | 2.343799719 | 1.51E-04 | 2.91E-03 |
| TRI29_HUMAN\|Q14134_S166 | 2.342558381 | 5.27E-03 | 4.50E-02 |
| SPTB2_HUMAN\|Q01082-3_S14 | 2.341398847 | 1.08E-07 | 9.17E-06 |
| NDRG1_HUMAN\|Q92597_S333_T328 | 2.340046987 | 6.80E-05 | 1.63E-03 |
| SF3B2_HUMAN\|Q13435_S307_T311 | 2.338625938 | 1.44E-03 | 1.98E-02 |
| RL34_HUMAN\|P49207_S12 | 2.338432934 | 4.59E-04 | 9.28E-03 |
| FOXO3_HUMAN\|O43524_S413 | 2.337329036 | 1.10E-05 | 1.57E-04 |
| TB182_HUMAN\|Q9C0C2_S691 | 2.337023355 | 9.26E-09 | 6.87E-07 |
| LMNA_HUMAN\|P02545_S636 | 2.335685146 | 5.28E-03 | 4.50E-02 |
| AHNK_HUMAN\|Q09666_S5784 | 2.33546626 | 2.70E-04 | 2.93E-03 |
| COHA1_HUMAN\|Q9UMD9_S93_T97 | 2.335049993 | 1.52E-06 | 5.51E-05 |
| HMGA2_HUMAN\|P52926_S105 | 2.332608762 | 3.35E-05 | 7.28E-04 |
| DC1L1_HUMAN\|Q9Y6G9_S415 | 2.331955137 | 2.89E-04 | 1.35E-02 |
| ELMO3_HUMAN\|Q96BJ8-3_S5 | 2.331362537 | 9.37E-03 | 3.29E-02 |
| CRBG1_HUMAN\|Q9Y4K1_S410 | 2.329101069 | 2.98E-06 | 2.42E-04 |
| ROA1_HUMAN\|P09651_S361_S364 | 2.325191923 | 8.86E-07 | 9.69E-05 |
| HIRP3_HUMAN\|Q9BW71_S555 | 2.324499148 | 1.42E-03 | 1.27E-02 |
| K1C17_HUMAN\|Q04695_S11_S13 | 2.32314157 | 5.02E-08 | 1.25E-05 |
| GGYF2_HUMAN\|Q6Y7W6_S201 | 2.321499247 | 3.73E-08 | 2.22E-06 |
| CCDC9_HUMAN\|Q9Y3X0_S376_S386_T381 | 2.320550199 | 1.01E-08 | 5.40E-06 |
| PABP1_HUMAN\|P11940_S629 | 2.318528806 | 7.13E-05 | 1.31E-03 |
| K1C14_HUMAN\|P02533_S44 | 2.312303517 | 3.23E-07 | 4.16E-05 |
| AHNK_HUMAN\|Q09666_S5397 | 2.310485936 | 1.44E-04 | 2.81E-03 |
| EF2_HUMAN\|P13639_T57 | 2.309867096 | 8.28E-05 | 1.13E-03 |

| | | | |
|---|---|---|---|
| XPC_HUMAN\|Q01831_S883_S884_S891 | 2.308647379 | 8.24E-07 | 7.24E-05 |
| ETV3_HUMAN\|P41162_S139 | 2.304887151 | 3.46E-05 | 5.90E-04 |
| AMPD2_HUMAN\|Q01433_S100 | 2.303499801 | 1.57E-04 | 3.97E-03 |
| PHLB1_HUMAN\|Q86UU1_S430 | 2.303360033 | 6.98E-08 | 1.80E-05 |
| REQU_HUMAN\|Q92785_S142 | 2.302504425 | 4.60E-08 | 1.25E-05 |
| LAP2A_HUMAN\|P42166_S156_T160 | 2.30226459 | 3.53E-06 | 2.42E-04 |
| RANG_HUMAN\|P43487_S60 | 2.300614173 | 2.67E-04 | 2.06E-03 |
| UBP5_HUMAN\|P45974_S785 | 2.299428772 | 5.02E-05 | 9.85E-04 |
| EIF3H_HUMAN\|O15372_S183 | 2.299016952 | 3.12E-06 | 2.20E-04 |
| K2C8_HUMAN\|P05787_Y25 | 2.298257626 | 5.69E-04 | 7.85E-03 |
| K1522_HUMAN\|Q9P206_S612 | 2.297975107 | 4.23E-07 | 2.52E-05 |
| EIF3H_HUMAN\|O15372_S183 | 2.297263929 | 3.16E-06 | 2.48E-04 |
| TTP_HUMAN\|P26651_S186 | 2.297259512 | 1.18E-05 | 3.46E-04 |
| CT2NL_HUMAN\|Q9P2B4_S443 | 2.296491983 | 7.25E-04 | 7.60E-03 |
| K2C7_HUMAN\|P08729_S38 | 2.294314324 | 2.65E-05 | 9.44E-04 |
| COHA1_HUMAN\|Q9UMD9_S93_S96 | 2.29424182 | 3.09E-06 | 9.41E-05 |
| LARP1_HUMAN\|Q6PKG0_S766 | 2.294103769 | 9.28E-05 | 2.29E-03 |
| CCDC9_HUMAN\|Q9Y3X0_S376 | 2.293036246 | 5.64E-09 | 9.71E-07 |
| CR025_HUMAN\|Q96B23_S66_S69 | 2.292994348 | 1.27E-05 | 5.85E-04 |
| LMNA_HUMAN\|P02545_S618 | 2.292828879 | 2.33E-04 | 2.60E-03 |
| LMNA_HUMAN\|P02545_S636 | 2.290248684 | 6.20E-03 | 3.63E-02 |
| ROA1_HUMAN\|P09651_S364 | 2.285720069 | 2.71E-05 | 1.24E-03 |
| COHA1_HUMAN\|Q9UMD9_S93_T97 | 2.285697023 | 2.22E-06 | 1.74E-04 |
| CT2NL_HUMAN\|Q9P2B4_S443 | 2.285192516 | 7.65E-04 | 2.47E-02 |
| K1C14_HUMAN\|P02533_S32 | 2.285078892 | 4.43E-09 | 8.00E-06 |
| NEB2_HUMAN\|Q96SB3_S99 | 2.284890209 | 2.93E-08 | 5.29E-06 |
| SRRM1_HUMAN\|Q8IYB3_S452 | 2.283460863 | 1.69E-03 | 2.15E-02 |
| RU17_HUMAN\|P08621_S320 | 2.280167363 | 2.30E-05 | 4.32E-04 |
| AKTS1_HUMAN\|Q96B36_S187 | 2.278797505 | 2.78E-07 | 4.07E-05 |
| TBCD5_HUMAN\|Q92609_S522 | 2.277142155 | 1.00E-07 | 1.13E-05 |
| RAI14_HUMAN\|Q9P0K7_S296 | 2.275639826 | 3.48E-07 | 1.93E-05 |

| | | | |
|---|---|---|---|
| NUP98_HUMAN\|P52948_S623 | 2.275366851 | 1.71E-07 | 6.49E-06 |
| TM230_HUMAN\|Q96A57_T25 | 2.275198073 | 1.46E-06 | 6.79E-05 |
| G3BP1_HUMAN\|Q13283_S250 | 2.274912398 | 2.01E-04 | 2.80E-03 |
| K2C5_HUMAN\|P13647_S64 | 2.274562941 | 1.29E-06 | 1.25E-04 |
| ACAP2_HUMAN\|Q15057_S384_T382 | 2.274184331 | 1.95E-12 | 1.06E-08 |
| K2C6A_HUMAN\|P02538_S85 | 2.273118085 | 3.45E-04 | 3.54E-03 |
| KPCD2_HUMAN\|Q9BZL6_S212 | 2.272316862 | 1.84E-07 | 1.38E-05 |
| HDGF_HUMAN\|P51858_S103 | 2.271540629 | 1.10E-04 | 3.13E-03 |
| HSPB1_HUMAN\|P04792_S78_S82 | 2.270720743 | 3.20E-06 | 6.26E-05 |
| UBP5_HUMAN\|P45974_S785 | 2.268139915 | 6.05E-05 | 2.13E-03 |
| PHLB2_HUMAN\|Q86SQ0_S330_S334 | 2.266232936 | 5.14E-06 | 3.10E-04 |
| TB182_HUMAN\|Q9C0C2_S1029 | 2.265181123 | 2.01E-09 | 5.18E-07 |
| CRBG1_HUMAN\|Q9Y4K1_S141 | 2.264546806 | 2.55E-07 | 1.62E-05 |
| LAD1_HUMAN\|O00515_S512 | 2.26420454 | 6.32E-09 | 1.27E-06 |
| SRP09_HUMAN\|P49458_T85 | 2.257878768 | 3.61E-08 | 1.09E-05 |
| ARHG5_HUMAN\|Q12774_S450 | 2.256007212 | 1.49E-06 | 1.52E-04 |
| NOLC1_HUMAN\|Q14978_S688_S698 | 2.253797866 | 8.55E-04 | 1.14E-02 |
| MPZL1_HUMAN\|O95297_S221 | 2.253361192 | 1.54E-04 | 9.73E-03 |
| ACAP2_HUMAN\|Q15057_S381_S384 | 2.253115552 | 5.06E-08 | 1.25E-05 |
| PEA15_HUMAN\|Q15121_S116 | 2.251104315 | 1.32E-06 | 6.27E-05 |
| FKBP5_HUMAN\|Q13451_T443 | 2.247672126 | 1.32E-05 | 4.62E-04 |
| GGYF2_HUMAN\|Q6Y7W6_S30 | 2.244946909 | 6.79E-05 | 9.64E-04 |
| LYRIC_HUMAN\|Q86UE4_S298 | 2.24473545 | 1.53E-05 | 2.06E-04 |
| ROA1_HUMAN\|P09651_S368 | 2.243866502 | 5.94E-05 | 2.35E-03 |
| MK14_HUMAN\|Q16539_T180_Y182 | 2.243349805 | 1.86E-03 | 4.17E-02 |
| JAM1_HUMAN\|Q9Y624_S284 | 2.24100732 | 2.88E-04 | 4.72E-03 |
| DC1L1_HUMAN\|Q9Y6G9_S415 | 2.238391959 | 4.67E-04 | 8.90E-03 |
| MISP_HUMAN\|Q8IVT2_S155 | 2.237693961 | 2.95E-06 | 2.12E-04 |
| FGD6_HUMAN\|Q6ZV73_T693 | 2.234943342 | 1.18E-07 | 9.81E-06 |
| LAP2A_HUMAN\|P42166_S156_T160 | 2.230906921 | 5.91E-06 | 2.78E-04 |
| IF4B_HUMAN\|P23588_S352 | 2.230850666 | 2.78E-03 | 2.92E-02 |

| | | | |
|---|---|---|---|
| AMPD2_HUMAN\|Q01433_S100 | 2.229584553 | 2.36E-04 | 3.20E-03 |
| KPCD2_HUMAN\|Q9BZL6_S212 | 2.227633806 | 2.73E-07 | 2.24E-05 |
| AHNK_HUMAN\|Q09666_T4430 | 2.227479175 | 1.58E-07 | 3.13E-05 |
| PKP3_HUMAN\|Q9Y446_S180 | 2.226837924 | 2.62E-07 | 3.95E-05 |
| K1C14_HUMAN\|P02533_S32 | 2.226833335 | 8.70E-09 | 4.29E-06 |
| K2C8_HUMAN\|P05787_Y25 | 2.226760778 | 8.30E-04 | 6.89E-03 |
| EGFR_HUMAN\|P00533_Y1197 | 2.224635529 | 3.10E-04 | 6.74E-03 |
| LMNA_HUMAN\|P02545_S618 | 2.222238848 | 3.45E-04 | 5.92E-03 |
| HSPB1_HUMAN\|P04792_S78_S82 | 2.22071519 | 4.68E-06 | 1.68E-04 |
| EIF3H_HUMAN\|O15372_S183 | 2.220092285 | 5.81E-06 | 9.73E-04 |
| RAI14_HUMAN\|Q9P0K7_S281 | 2.219250444 | 2.47E-09 | 2.73E-07 |
| AHNK_HUMAN\|Q09666_S572 | 2.217236164 | 6.54E-07 | 6.24E-05 |
| ANXA2_HUMAN\|P07355_S26 | 2.217089887 | 1.73E-03 | 2.11E-02 |
| K2C8_HUMAN\|P05787_S24 | 2.216745494 | 6.10E-07 | 2.95E-05 |
| K2C5_HUMAN\|P13647_S64 | 2.215038823 | 2.17E-06 | 7.16E-05 |
| CT2NL_HUMAN\|Q9P2B4_S443 | 2.214590978 | 1.08E-03 | 9.98E-03 |
| SNX6_HUMAN\|Q9UNH7_S316 | 2.213689523 | 1.22E-03 | 1.09E-02 |
| UIMC1_HUMAN\|Q96RL1_S101 | 2.213575078 | 4.02E-03 | 1.69E-02 |
| GRSF1_HUMAN\|Q12849_S118 | 2.213377466 | 2.23E-08 | 1.52E-06 |
| EGFR_HUMAN\|P00533_Y1197 | 2.212437361 | 3.33E-04 | 3.84E-02 |
| K2C5_HUMAN\|P13647_S21_S26 | 2.211859828 | 5.06E-07 | 2.58E-05 |
| EPHA2_HUMAN\|P29317_Y772 | 2.211313767 | 3.44E-04 | 7.07E-03 |
| K1C14_HUMAN\|P02533_Y46 | 2.210563981 | 1.65E-07 | 1.16E-05 |
| EDC4_HUMAN\|Q6P2E9_S723 | 2.209589987 | 4.66E-04 | 1.82E-02 |
| HOME3_HUMAN\|Q9NSC5_S159 | 2.208633489 | 4.69E-05 | 5.05E-04 |
| PLEC_HUMAN\|Q15149_S4384_S4386_S4389 | 2.207338151 | 3.34E-04 | 6.94E-03 |
| GYS1_HUMAN\|P13807_S727_S731 | 2.206935456 | 9.78E-07 | 1.08E-04 |
| SLK_HUMAN\|Q9H2G2_S655 | 2.205118189 | 2.07E-06 | 1.35E-04 |
| IASPP_HUMAN\|Q8WUF5_S526 | 2.204730708 | 1.09E-04 | 7.75E-03 |
| MRCKG_HUMAN\|Q6DT37_S1544 | 2.204106223 | 2.85E-06 | 5.76E-05 |
| K1C14_HUMAN\|P02533_Y46 | 2.201595014 | 1.81E-07 | 3.30E-05 |

| | | | |
|---|---|---|---|
| FYN_HUMAN\|P06241_Y420 | 2.201421161 | 9.36E-05 | 1.64E-03 |
| PABP1_HUMAN\|P11940_S629 | 2.20097229 | 1.47E-04 | 3.85E-03 |
| AFTIN_HUMAN\|Q6ULP2_S518 | 2.197935972 | 2.77E-08 | 7.52E-06 |
| MICA3_HUMAN\|Q7RTP6_S1636 | 2.197319605 | 1.43E-05 | 4.92E-04 |
| DBNL_HUMAN\|Q9UJU6_S269 | 2.196053648 | 1.80E-06 | 1.48E-04 |
| SRC8_HUMAN\|Q14247_Y446 | 2.195411245 | 1.29E-03 | 1.84E-02 |
| RAI14_HUMAN\|Q9P0K7_S290 | 2.19533557 | 4.43E-06 | 1.80E-04 |
| K2C5_HUMAN\|P13647_S26_T19 | 2.195316 | 1.21E-03 | 1.48E-02 |
| K1C16_HUMAN\|P08779_S449_S452 | 2.194967975 | 2.46E-05 | 7.56E-04 |
| K1C17_HUMAN\|Q04695_S24 | 2.194159574 | 4.42E-08 | 4.52E-06 |
| 1433Z_HUMAN\|P63104_T232 | 2.192542795 | 4.68E-06 | 3.57E-04 |
| PSF2_HUMAN\|Q9Y248_S182_T180 | 2.191568136 | 1.49E-06 | 1.14E-04 |
| CRBG1_HUMAN\|Q9Y4K1_S410 | 2.190228628 | 8.05E-06 | 2.63E-04 |
| NFIB_HUMAN\|O00712_S276_S300_T286 | 2.189446023 | 2.08E-03 | 1.11E-02 |
| HSPB1_HUMAN\|P04792_S78_S82 | 2.187198898 | 6.09E-06 | 5.00E-04 |
| GYS1_HUMAN\|P13807_S727_S731 | 2.186601786 | 1.17E-06 | 2.82E-05 |
| B4DKV4_HUMAN\|B4DKV4_S22 | 2.185873124 | 9.25E-05 | 5.97E-03 |
| DDX3X_HUMAN\|O00571_S612 | 2.181939776 | 1.61E-04 | 2.38E-03 |
| R3HD1_HUMAN\|Q15032_S973 | 2.181665939 | 9.04E-10 | 2.58E-07 |
| SRRM1_HUMAN\|Q8IYB3_S756 | 2.181291454 | 3.20E-05 | 3.73E-04 |
| SCRIB_HUMAN\|Q14160_S1285 | 2.18103309 | 2.30E-06 | 1.78E-04 |
| GTR1_HUMAN\|P11166_S490 | 2.179049668 | 7.96E-04 | 1.09E-02 |
| SF3B2_HUMAN\|Q13435_S307_S309 | 2.177989159 | 4.51E-05 | 1.95E-03 |
| RENT1_HUMAN\|Q92900_S1127 | 2.177719285 | 6.04E-04 | 8.93E-03 |
| R3HD2_HUMAN\|Q9Y2K5_S349 | 2.177466447 | 2.36E-04 | 3.20E-03 |
| MPZL1_HUMAN\|O95297_S221 | 2.176250082 | 2.50E-04 | 5.48E-03 |
| ROA1_HUMAN\|P09651_S364 | 2.171582291 | 6.00E-05 | 1.66E-03 |
| JAM1_HUMAN\|Q9Y624_S284 | 2.170255202 | 4.45E-04 | 5.16E-03 |
| FA83B_HUMAN\|Q5T0W9_S802 | 2.169600372 | 1.02E-07 | 4.41E-06 |
| DBNL_HUMAN\|Q9UJU6_T270 | 2.169518738 | 2.97E-06 | 2.12E-04 |
| NU214_HUMAN\|P35658_S1023 | 2.169016374 | 2.77E-07 | 1.98E-05 |

| | | | |
|---|---|---|---|
| K2C6A_HUMAN\|P02538_S60 | 2.168402445 | 4.98E-04 | 7.10E-03 |
| ACINU_HUMAN\|Q9UKV3_S328 | 2.166062804 | 3.09E-04 | 3.89E-03 |
| K1C18_HUMAN\|P05783_S7 | 2.164956916 | 4.55E-06 | 3.52E-04 |
| FKBP5_HUMAN\|Q13451_S445 | 2.164889116 | 2.89E-04 | 3.09E-03 |
| AHNK_HUMAN\|Q09666_T5824 | 2.164813728 | 4.12E-04 | 4.04E-03 |
| TTP_HUMAN\|P26651_S60 | 2.164770813 | 4.17E-08 | 2.35E-06 |
| PHF14_HUMAN\|O94880_T287 | 2.163733558 | 2.38E-07 | 8.61E-05 |
| TM1L1_HUMAN\|O75674_S321 | 2.162742433 | 6.11E-06 | 3.99E-04 |
| NIBL1_HUMAN\|Q96TA1_S691_S696 | 2.161475737 | 9.10E-08 | 1.59E-05 |
| SF3B2_HUMAN\|Q13435_S307_T311 | 2.15953884 | 3.14E-03 | 1.56E-02 |
| K2C6A_HUMAN\|P02538_S85 | 2.158027292 | 6.81E-04 | 9.09E-03 |
| HIRP3_HUMAN\|Q9BW71_S555 | 2.157228554 | 2.98E-03 | 3.21E-02 |
| K1C19_HUMAN\|P08727_S13 | 2.155190569 | 5.26E-06 | 1.40E-04 |
| COHA1_HUMAN\|Q9UMD9_S93_S96 | 2.154609345 | 9.36E-06 | 4.62E-04 |
| T22D4_HUMAN\|Q9Y3Q8_T167 | 2.154480854 | 1.13E-07 | 4.37E-05 |
| SRRM1_HUMAN\|Q8IYB3_S754 | 2.152428093 | 8.99E-06 | 1.34E-04 |
| K2C8_HUMAN\|P05787_S24 | 2.151909401 | 1.19E-06 | 9.33E-05 |
| TOP2B_HUMAN\|Q02880_S1550_S1552 | 2.151677123 | 1.15E-05 | 1.68E-04 |
| K2C7_HUMAN\|P08729_S38 | 2.150680898 | 7.33E-05 | 1.71E-03 |
| CPSF4_HUMAN\|O95639_S202 | 2.149247308 | 2.31E-06 | 1.07E-04 |
| DC1L1_HUMAN\|Q9Y6G9_S412 | 2.147406626 | 2.81E-03 | 2.09E-02 |
| PARD3_HUMAN\|Q8TEW0_S220 | 2.146816894 | 1.29E-04 | 2.85E-03 |
| COTL1_HUMAN\|Q14019_T141 | 2.146725052 | 3.52E-04 | 7.18E-03 |
| CAVN1_HUMAN\|Q6NZI2_Y308 | 2.145900832 | 8.15E-05 | 2.85E-03 |
| PDLI2_HUMAN\|Q96JY6_S137 | 2.145864723 | 1.88E-08 | 5.40E-06 |
| K1C17_HUMAN\|Q04695_S11_S13 | 2.145159179 | 3.34E-07 | 4.41E-05 |
| B4DET1_HUMAN\|B4DET1_S720_T711 | 2.143740934 | 1.46E-03 | 8.25E-03 |
| ROA1_HUMAN\|P09651_S362_S365 | 2.143634503 | 1.61E-03 | 2.14E-02 |
| PKP3_HUMAN\|Q9Y446_S180_S183 | 2.143572089 | 1.12E-06 | 1.19E-04 |
| K2C5_HUMAN\|P13647_S26_T19 | 2.142926542 | 1.59E-03 | 1.15E-02 |
| IWS1_HUMAN\|Q96ST2_S300_S313_S315 | 2.142674781 | 1.23E-04 | 1.13E-03 |

| | | | |
|---|---|---|---|
| COHA1_HUMAN\|Q9UMD9_S76_S79 | 2.140700038 | 1.07E-04 | 2.48E-03 |
| HNRPU_HUMAN\|Q00839_S59 | 2.139662353 | 4.69E-03 | 3.77E-02 |
| K2C5_HUMAN\|P13647_S64 | 2.139658195 | 4.30E-06 | 2.22E-04 |
| IASPP_HUMAN\|Q8WUF5_T341 | 2.139165695 | 1.99E-04 | 4.67E-03 |
| MICA3_HUMAN\|Q7RTP6_S685_S687 | 2.139005368 | 1.59E-05 | 8.29E-04 |
| DBNL_HUMAN\|Q9UJU6_S269 | 2.138852954 | 2.92E-06 | 9.09E-05 |
| NCK2_HUMAN\|O43639_S90_T95 | 2.138162229 | 2.23E-03 | 1.49E-02 |
| IASPP_HUMAN\|Q8WUF5_S526 | 2.137520234 | 1.67E-04 | 4.13E-03 |
| NELFE_HUMAN\|P18615_S251 | 2.136705797 | 4.78E-06 | 8.56E-05 |
| K1C17_HUMAN\|Q04695_S11_S12 | 2.135986407 | 8.53E-07 | 7.24E-05 |
| B4DKV4_HUMAN\|B4DKV4_S19_S22 | 2.134526416 | 9.99E-05 | 6.23E-03 |
| K2C8_HUMAN\|P05787_Y25 | 2.13412851 | 1.37E-03 | 1.61E-02 |
| GGYF2_HUMAN\|Q6Y7W6_S201 | 2.133453245 | 2.07E-07 | 1.24E-04 |
| NUP50_HUMAN\|Q9UKX7_S221 | 2.131465468 | 7.96E-09 | 2.06E-06 |
| SPTB2_HUMAN\|Q01082-3_S14_S8 | 2.129892653 | 3.43E-08 | 4.04E-06 |
| SF3B2_HUMAN\|Q13435_S303 | 2.127716764 | 1.27E-03 | 1.12E-02 |
| DC1L1_HUMAN\|Q9Y6G9_S412 | 2.127609773 | 3.08E-03 | 3.30E-02 |
| MET_HUMAN\|P08581_S997_T993 | 2.127009261 | 2.97E-06 | 5.79E-05 |
| RS6_HUMAN\|P62753_S236 | 2.126660247 | 3.16E-04 | 3.97E-03 |
| GYS1_HUMAN\|P13807_S710 | 2.126618289 | 1.58E-03 | 1.74E-02 |
| HEAT6_HUMAN\|Q6AI08_S395_S397 | 2.126254448 | 4.26E-04 | 8.18E-03 |
| SERF2_HUMAN\|P84101_S21 | 2.124432744 | 3.81E-07 | 5.29E-05 |
| RERE_HUMAN\|Q9P2R6_S679 | 2.124406009 | 2.43E-05 | 2.96E-04 |
| K2C75_HUMAN\|O95678_S61 | 2.123570849 | 2.47E-05 | 4.58E-04 |
| GIT1_HUMAN\|Q9Y2X7_S388 | 2.123472476 | 4.07E-05 | 4.51E-04 |
| BL1S3_HUMAN\|Q6QNY0_S65 | 2.123034402 | 7.10E-03 | 3.05E-02 |
| UBP5_HUMAN\|P45974_S785 | 2.118508946 | 1.56E-04 | 2.66E-02 |
| K2C6B_HUMAN\|P04259_S37 | 2.118315559 | 8.54E-06 | 2.03E-04 |
| COHA1_HUMAN\|Q9UMD9_S76_S79 | 2.11807386 | 1.25E-04 | 3.45E-03 |
| WFS1_HUMAN\|O76024_S235 | 2.117510555 | 6.67E-05 | 1.20E-03 |
| IPP2M_HUMAN\|Q6NXS1_S121 | 2.115757789 | 2.19E-06 | 4.75E-05 |

| | | | |
|---|---|---|---|
| TBD2A_HUMAN|Q9BYX2_S920 | 2.115015303 | 4.30E-08 | 4.48E-06 |
| PKP3_HUMAN|Q9Y446_S180 | 2.11488545 | 7.31E-07 | 6.60E-05 |
| ACAP2_HUMAN|Q15057_S384_T382 | 2.114211497 | 1.47E-11 | 6.58E-08 |
| K1C14_HUMAN|P02533_S54_Y46 | 2.114073878 | 1.65E-06 | 1.39E-04 |
| IWS1_HUMAN|Q96ST2_T725 | 2.113586272 | 5.21E-06 | 1.39E-04 |
| PPM1G_HUMAN|O15355_S201 | 2.112157285 | 5.69E-05 | 2.07E-03 |
| SLTM_HUMAN|Q9NWH9_S371 | 2.111357275 | 1.94E-03 | 2.01E-02 |
| ITA2_HUMAN|P17301_S1180 | 2.110850036 | 1.46E-06 | 1.33E-04 |
| ARM10_HUMAN|Q8N2F6-2_S45 | 2.109412019 | 9.44E-03 | 3.85E-02 |
| NIBL1_HUMAN|Q96TA1_S691 | 2.107112628 | 7.99E-10 | 6.18E-07 |
| NIBL1_HUMAN|Q96TA1_S692 | 2.105449076 | 7.04E-08 | 1.41E-05 |
| K1522_HUMAN|Q9P206_S339 | 2.104921156 | 4.80E-04 | 6.88E-03 |
| NIBAN_HUMAN|Q9BZQ8_S926 | 2.103541 | 2.43E-08 | 4.70E-06 |
| CAVN1_HUMAN|Q6NZI2_Y308 | 2.103164103 | 1.10E-04 | 2.24E-02 |
| K1C16_HUMAN|P08779_S448_S451 | 2.101922684 | 3.11E-07 | 4.16E-05 |
| AHNK_HUMAN|Q09666_S5745_S5749 | 2.100109006 | 2.75E-09 | 1.86E-06 |
| PAR10_HUMAN|Q53GL7_S431 | 2.099782835 | 1.54E-08 | 4.93E-06 |
| CR025_HUMAN|Q96B23_S66_S67_T74 | 2.098948772 | 3.61E-03 | 3.79E-02 |
| GYS1_HUMAN|P13807_S710 | 2.098850938 | 1.82E-03 | 1.92E-02 |
| JUN_HUMAN|P05412_S63 | 2.097848207 | 1.28E-06 | 3.00E-05 |
| TB182_HUMAN|Q9C0C2_S691 | 2.097561809 | 1.15E-07 | 1.89E-05 |
| SERF2_HUMAN|P84101_S21 | 2.096010939 | 5.25E-07 | 3.60E-05 |
| DC1L1_HUMAN|Q9Y6G9_S415 | 2.09589789 | 1.01E-03 | 1.55E-02 |
| NKTR_HUMAN|P30414_S463 | 2.09527086 | 5.82E-07 | 3.89E-05 |
| ROA1_HUMAN|P09651_S368 | 2.09351731 | 1.67E-04 | 2.45E-03 |
| CCD86_HUMAN|Q9H6F5_S217 | 2.093382587 | 4.03E-06 | 1.52E-04 |
| NGAP_HUMAN|Q9UJF2_S954 | 2.093152614 | 3.93E-07 | 2.43E-05 |
| K2C5_HUMAN|P13647_T19 | 2.092447158 | 3.17E-05 | 1.08E-03 |
| MRCKG_HUMAN|Q6DT37_S1544 | 2.092059428 | 7.60E-06 | 1.20E-04 |
| DC1I2_HUMAN|Q13409-2_S84 | 2.091313202 | 8.92E-08 | 1.59E-05 |
| PAK2_HUMAN|Q13177_S192 | 2.09018891 | 7.32E-04 | 9.63E-03 |

| | | | |
|---|---|---|---|
| DC1L1_HUMAN\|Q9Y6G9_S412 | 2.089386957 | 3.69E-03 | 3.84E-02 |
| NCOA7_HUMAN\|Q8NI08_S208_T210 | 2.089051861 | 4.43E-08 | 4.85E-06 |
| PININ_HUMAN\|Q9H307_S66 | 2.088630719 | 1.38E-06 | 3.33E-05 |
| ROA1_HUMAN\|P09651_S363 | 2.088239312 | 5.27E-04 | 5.83E-03 |
| CCAR2_HUMAN\|Q8N163_S678 | 2.087934261 | 3.32E-04 | 4.11E-03 |
| AHNK2_HUMAN\|Q8IVF2_S1583 | 2.087588599 | 6.98E-06 | 3.15E-04 |
| ITA2_HUMAN\|P17301_S1181 | 2.087563897 | 3.72E-06 | 2.84E-04 |
| NELFE_HUMAN\|P18615_S51 | 2.087408054 | 1.08E-07 | 1.20E-05 |
| AMPD2_HUMAN\|Q01433_S100 | 2.087280859 | 5.41E-04 | 7.54E-03 |
| DBNL_HUMAN\|Q9UJU6_T270 | 2.086990583 | 6.00E-06 | 1.53E-04 |
| PKP3_HUMAN\|Q9Y446_S313_S314 | 2.086694104 | 3.03E-08 | 9.67E-06 |
| TB182_HUMAN\|Q9C0C2_T1282 | 2.086522979 | 6.80E-08 | 5.76E-06 |
| STX5_HUMAN\|Q13190_S9 | 2.085955875 | 4.38E-05 | 1.32E-03 |
| TBCD5_HUMAN\|Q92609_S522 | 2.085143061 | 5.76E-07 | 5.68E-05 |
| DPYL3_HUMAN\|Q14195_S568 | 2.083377821 | 4.09E-05 | 1.26E-03 |
| TB10A_HUMAN\|Q9BXI6_S40_S43 | 2.083152695 | 3.44E-05 | 3.95E-04 |
| AHNK_HUMAN\|Q09666_S210_S216_S220 | 2.082740094 | 2.28E-08 | 7.72E-06 |
| CLIP4_HUMAN\|Q8N3C7_T582 | 2.082408433 | 5.26E-08 | 7.31E-06 |
| GRAP1_HUMAN\|Q4V328_S664 | 2.081789633 | 5.39E-07 | 6.49E-05 |
| NUP53_HUMAN\|Q8NFH5_S105 | 2.080821809 | 1.95E-06 | 6.75E-05 |
| ATRX_HUMAN\|P46100_S784 | 2.080426173 | 5.47E-06 | 9.32E-05 |
| FIL1L_HUMAN\|Q4L180_S1049 | 2.080232774 | 6.30E-06 | 2.25E-04 |
| MICA3_HUMAN\|Q7RTP6_S685_S687 | 2.079158931 | 2.54E-05 | 1.28E-03 |
| T22D4_HUMAN\|Q9Y3Q8_T167 | 2.079019666 | 2.57E-07 | 3.95E-05 |
| CNKR1_HUMAN\|Q969H4_S305_S314 | 2.078582563 | 1.76E-06 | 8.91E-05 |
| GOGA4_HUMAN\|Q13439_S89_S92 | 2.078142946 | 1.88E-07 | 3.29E-05 |
| PLEC_HUMAN\|Q15149_S4406 | 2.075887667 | 6.73E-04 | 3.99E-03 |
| FA83B_HUMAN\|Q5T0W9_S802 | 2.075527268 | 2.60E-07 | 2.20E-05 |
| KPCD2_HUMAN\|Q9BZL6_S212 | 2.074605627 | 1.16E-06 | 4.86E-04 |
| RFIP3_HUMAN\|O75154_S52 | 2.074427799 | 2.88E-07 | 4.33E-05 |
| AHNK2_HUMAN\|Q8IVF2_S1583 | 2.074191196 | 7.90E-06 | 4.12E-04 |

| | | | |
|---|---|---|---|
| MAP7_HUMAN\|Q14244_T184 | 2.073868304 | 5.43E-08 | 4.99E-06 |
| K2C75_HUMAN\|O95678_S40 | 2.07365092 | 1.02E-04 | 2.10E-03 |
| E7ETA6_HUMAN\|E7ETA6_T1557 | 2.073108884 | 4.59E-05 | 1.35E-03 |
| BCR_HUMAN\|P11274_T385 | 2.072287707 | 6.30E-03 | 4.77E-02 |
| VIP2_HUMAN\|O43314_S1006 | 2.071476516 | 7.61E-09 | 1.25E-06 |
| T22D4_HUMAN\|Q9Y3Q8_T167 | 2.070284433 | 2.84E-07 | 4.40E-05 |
| MTAP2_HUMAN\|P11137_T1602_T1605 | 2.069433277 | 4.66E-05 | 9.29E-04 |
| REPS1_HUMAN\|Q96D71_S162_S166_T173 | 2.06937346 | 1.21E-08 | 1.72E-06 |
| SF3B2_HUMAN\|Q13435_S303_S309 | 2.068517679 | 2.09E-04 | 1.72E-03 |
| AKTS1_HUMAN\|Q96B36_S187 | 2.068109583 | 2.07E-06 | 1.35E-04 |
| PKP3_HUMAN\|Q9Y446_S223 | 2.066913403 | 1.93E-03 | 1.33E-02 |
| NDRG1_HUMAN\|Q92597_S330_S336_T335 | 2.064026262 | 7.36E-03 | 3.56E-02 |
| GRAP1_HUMAN\|Q4V328_S666_S669 | 2.063862472 | 6.50E-04 | 5.80E-03 |
| SRC8_HUMAN\|Q14247_Y446 | 2.062919669 | 2.56E-03 | 2.54E-02 |
| FLVC1_HUMAN\|Q9Y5Y0_S536 | 2.062125118 | 1.43E-06 | 1.47E-04 |
| HMGN1_HUMAN\|P05114_S7 | 2.061368936 | 1.54E-06 | 3.43E-05 |
| SF3B2_HUMAN\|Q13435_S303_S309 | 2.060938955 | 2.20E-04 | 5.30E-03 |
| LAP2A_HUMAN\|P42166_S156_T160 | 2.060757142 | 2.23E-05 | 4.23E-04 |
| DKC1_HUMAN\|O60832_S485 | 2.060401794 | 1.39E-07 | 2.79E-05 |
| NDRG1_HUMAN\|Q92597_S326_S330_S333 | 2.058809098 | 6.41E-07 | 7.24E-05 |
| K1143_HUMAN\|Q96AT1_S146 | 2.058031613 | 8.80E-06 | 2.07E-04 |
| WAC2A_HUMAN\|Q641Q2_S284 | 2.057444106 | 6.71E-06 | 1.10E-04 |
| DKC1_HUMAN\|O60832_S485 | 2.055583076 | 1.47E-07 | 2.57E-05 |
| AHNK_HUMAN\|Q09666_S5864 | 2.055370402 | 1.57E-04 | 3.97E-03 |
| HMGA2_HUMAN\|P52926_S105 | 2.054264701 | 2.36E-04 | 1.78E-03 |
| ARM10_HUMAN\|Q8N2F6-2_S45 | 2.053666194 | 1.19E-02 | 3.95E-02 |
| K2C5_HUMAN\|P13647_S16_S21 | 2.053551383 | 6.75E-05 | 1.63E-03 |
| NGAP_HUMAN\|Q9UJF2_T620 | 2.052434484 | 2.37E-04 | 2.64E-03 |
| NUP50_HUMAN\|Q9UKX7_S221 | 2.052402258 | 2.09E-08 | 3.15E-06 |
| SLAI2_HUMAN\|Q9P270_S349_S353_S357 | 2.052400275 | 3.47E-08 | 3.84E-06 |
| TTP_HUMAN\|P26651_S60 | 2.052155642 | 1.63E-07 | 1.25E-05 |

| | | | |
|---|---|---|---|
| HMGA1_HUMAN\|P17096_S49 | 2.052081775 | 2.34E-04 | 4.12E-03 |
| TT39B_HUMAN\|Q5VTQ0_S120 | 2.051971327 | 1.18E-05 | 4.40E-04 |
| BORG4_HUMAN\|Q9H3Q1_S64 | 2.051284959 | 8.87E-08 | 2.19E-05 |
| GGYF2_HUMAN\|Q6Y7W6_S30 | 2.05049458 | 2.41E-04 | 1.26E-02 |
| COBL1_HUMAN\|Q53SF7_T322 | 2.049642976 | 3.32E-08 | 1.09E-05 |
| AHNK2_HUMAN\|Q8IVF2_S5175 | 2.048401725 | 9.13E-07 | 5.38E-05 |
| K2C6A_HUMAN\|P02538_S58 | 2.04833791 | 8.44E-04 | 6.97E-03 |
| K2C5_HUMAN\|P13647_S16_S21 | 2.047353436 | 7.11E-05 | 1.85E-03 |
| EFNB1_HUMAN\|P98172_T284 | 2.047154183 | 6.71E-05 | 1.78E-03 |
| TAGL2_HUMAN\|P37802_S163 | 2.046490786 | 1.34E-05 | 8.74E-04 |
| SRRM1_HUMAN\|Q8IYB3_S754 | 2.046429187 | 2.27E-05 | 2.77E-04 |
| CPSF4_HUMAN\|O95639_S202 | 2.046158258 | 5.68E-06 | 1.28E-03 |
| FA83B_HUMAN\|Q5T0W9_T687 | 2.045810466 | 2.57E-03 | 2.55E-02 |
| NIPA4_HUMAN\|Q0D2K0_S427 | 2.045718647 | 1.64E-03 | 9.08E-03 |
| DCAF6_HUMAN\|Q58WW2_S657 | 2.044968078 | 2.05E-06 | 4.51E-05 |
| Q659E9_HUMAN\|Q659E9_S168 | 2.044909492 | 2.30E-09 | 1.38E-06 |
| K1C17_HUMAN\|Q04695_S28 | 2.043196936 | 2.32E-06 | 7.45E-05 |
| HEAT6_HUMAN\|Q6AI08_S395_S397 | 2.040836264 | 7.26E-04 | 7.60E-03 |
| SRRM2_HUMAN\|Q9UQ35_S2692 | 2.040709025 | 9.16E-05 | 2.27E-03 |
| GGYF2_HUMAN\|Q6Y7W6_S201 | 2.040444871 | 5.29E-07 | 3.05E-05 |
| TBCD5_HUMAN\|Q92609_S522 | 2.039911543 | 9.04E-07 | 4.71E-05 |
| PININ_HUMAN\|Q9H307_S66 | 2.039659116 | 2.24E-06 | 1.05E-04 |
| CCAR2_HUMAN\|Q8N163_S678 | 2.039338966 | 4.54E-04 | 2.92E-03 |
| TOM34_HUMAN\|Q15785_S44 | 2.038622598 | 1.04E-02 | 3.57E-02 |
| TANC2_HUMAN\|Q9HCD6_S1827 | 2.037372796 | 2.93E-08 | 3.69E-06 |
| DBNL_HUMAN\|Q9UJU6_S269 | 2.037241187 | 7.19E-06 | 4.87E-04 |
| SRRM1_HUMAN\|Q8IYB3_S752_S754_S756 | 2.036410327 | 5.33E-09 | 9.63E-07 |
| GRAP1_HUMAN\|Q4V328_S666_S669 | 2.036253899 | 7.72E-04 | 9.98E-03 |
| HNRPM_HUMAN\|P52272_S575 | 2.036172718 | 1.10E-05 | 3.34E-04 |
| PAWR_HUMAN\|Q96IZ0_S231 | 2.035134348 | 7.83E-08 | 1.75E-05 |
| TV23B_HUMAN\|Q9NYZ1_T199 | 2.034448556 | 6.26E-04 | 6.58E-03 |

| | | | |
|---|---|---|---|
| SCAM2_HUMAN\|O15127_S319 | 2.032551481 | 1.43E-05 | 8.97E-04 |
| MTAP2_HUMAN\|P11137_T1602_T1605 | 2.031724287 | 6.36E-05 | 2.24E-03 |
| MFAP1_HUMAN\|P55081_S116 | 2.031104745 | 1.16E-04 | 1.08E-03 |
| SRRM2_HUMAN\|Q9UQ35_S2692 | 2.030926744 | 9.88E-05 | 2.05E-03 |
| NEB2_HUMAN\|Q96SB3_S100 | 2.030579914 | 2.76E-05 | 6.25E-04 |
| K2C5_HUMAN\|P13647_S77_S82 | 2.029664591 | 1.34E-04 | 2.93E-03 |
| SQSTM_HUMAN\|Q13501_S355_S365 | 2.028678782 | 1.03E-03 | 1.47E-02 |
| IRS1_HUMAN\|P35568_S330 | 2.028652758 | 2.60E-05 | 1.17E-03 |
| RBM7_HUMAN\|Q9Y580_S136 | 2.026517275 | 3.26E-06 | 1.31E-04 |
| FKBP5_HUMAN\|Q13451_S445 | 2.024573631 | 6.92E-04 | 2.26E-02 |
| VIME_HUMAN\|P08670_S34 | 2.023881457 | 5.72E-07 | 2.83E-05 |
| ITA2_HUMAN\|P17301_S1181 | 2.023441715 | 7.39E-06 | 4.50E-04 |
| K2C8_HUMAN\|P05787_S477 | 2.023170662 | 6.03E-06 | 2.82E-04 |
| AHNK_HUMAN\|Q09666_S212_S216 | 2.023165259 | 2.75E-08 | 3.36E-06 |
| CALD1_HUMAN\|Q05682_S789 | 2.023112812 | 1.94E-06 | 1.84E-04 |
| EFNB1_HUMAN\|P98172_T284 | 2.023049294 | 8.21E-05 | 1.82E-03 |
| M3K3_HUMAN\|Q99759_S337 | 2.022914041 | 3.32E-06 | 6.38E-05 |
| PKP3_HUMAN\|Q9Y446_S180_S183 | 2.022181819 | 3.51E-06 | 1.03E-04 |
| ETV3_HUMAN\|P41162_S139 | 2.020118923 | 2.71E-04 | 4.53E-03 |
| CING_HUMAN\|Q9P2M7_S129_S134 | 2.019707479 | 1.90E-06 | 1.33E-04 |
| UBP32_HUMAN\|Q8NFA0_S1393 | 2.019252312 | 2.98E-08 | 1.70E-06 |
| PRC2A_HUMAN\|P48634_S1325_S1328_T1323 | 2.018108056 | 7.35E-04 | 1.03E-02 |
| UBXN7_HUMAN\|O94888_S280_S288 | 2.01800615 | 5.94E-05 | 1.47E-03 |
| CAVN1_HUMAN\|Q6NZI2_T302 | 2.017347662 | 1.68E-08 | 3.04E-05 |
| TPC12_HUMAN\|Q8WVT3_S184 | 2.016129845 | 9.95E-06 | 1.49E-04 |
| PRC2A_HUMAN\|P48634_S761_S764 | 2.015347499 | 6.79E-09 | 4.60E-06 |
| ZO1_HUMAN\|Q07157_T868_Y843 | 2.015183299 | 9.88E-05 | 1.57E-03 |
| TB182_HUMAN\|Q9C0C2_S429 | 2.01450794 | 6.66E-05 | 6.02E-03 |
| TOP2B_HUMAN\|Q02880_S1550_S1552 | 2.013140732 | 3.86E-05 | 1.60E-03 |
| RENT1_HUMAN\|Q92900_S1107_T1111 | 2.012992868 | 1.13E-04 | 2.30E-03 |
| ROA1_HUMAN\|P09651_S364 | 2.012940693 | 1.97E-04 | 4.96E-03 |

| | | | |
|---|---|---|---|
| UTP18_HUMAN\|Q9Y5J1_S121 | 2.012426911 | 2.33E-03 | 1.83E-02 |
| COBL1_HUMAN\|Q53SF7_T322 | 2.011785109 | 5.25E-08 | 4.99E-06 |
| LASP1_HUMAN\|Q14847_T104 | 2.010088539 | 7.78E-08 | 6.39E-06 |
| AHNK_HUMAN\|Q09666_S5746_S5752 | 2.009617717 | 3.43E-08 | 1.09E-05 |
| CAVN1_HUMAN\|Q6NZI2_Y308 | 2.008760961 | 2.17E-04 | 3.92E-03 |
| PSF2_HUMAN\|Q9Y248_S182_T180 | 2.007591755 | 9.97E-06 | 2.28E-04 |
| COHA1_HUMAN\|Q9UMD9_S61_S62_S70 | 2.007544254 | 2.92E-04 | 5.25E-03 |
| SF3B2_HUMAN\|Q13435_S303_S309 | 2.006740802 | 3.22E-04 | 6.52E-03 |
| SPTB2_HUMAN\|Q01082_T2328 | 2.006569889 | 3.79E-04 | 5.80E-03 |
| CR025_HUMAN\|Q96B23_S69 | 2.006377816 | 2.67E-04 | 1.94E-03 |
| MARCS_HUMAN\|P29966_S118 | 2.006327637 | 3.15E-05 | 3.62E-04 |
| GRSF1_HUMAN\|Q12849_S118 | 2.005669336 | 2.43E-07 | 6.12E-05 |
| TAGL2_HUMAN\|P37802_S163 | 2.005618178 | 1.97E-05 | 2.47E-04 |
| LMNB1_HUMAN\|P20700_S23_T20 | 2.005365033 | 1.05E-03 | 5.72E-03 |
| RL23_HUMAN\|P62829_S138 | 2.003610668 | 1.37E-03 | 1.87E-02 |
| ARG35_HUMAN\|A5YM69_S450 | 2.003599563 | 1.49E-05 | 2.03E-04 |
| TB10A_HUMAN\|Q9BXI6_S40_S45 | 2.002647861 | 4.90E-04 | 5.48E-03 |
| FOSL1_HUMAN\|P15407_S252_S265 | 2.002585799 | 2.27E-04 | 5.51E-03 |
| G3BP1_HUMAN\|Q13283_S250 | 2.001264724 | 1.01E-03 | 1.29E-02 |
| K1C17_HUMAN\|Q04695_S11_S12 | 2.000998077 | 3.89E-06 | 2.60E-04 |
| K2C6A_HUMAN\|P02538_T548 | 2.00043697 | 1.29E-03 | 9.70E-03 |
| PLEC_HUMAN\|Q15149_S4675_Y4668 | 1.999471312 | 6.49E-05 | 1.22E-03 |
| ZDHC5_HUMAN\|Q9C0B5_S704 | 1.999307495 | 5.92E-05 | 1.65E-03 |
| HOME3_HUMAN\|Q9NSC5_S159 | 1.998962656 | 2.05E-04 | 5.15E-03 |
| T22D4_HUMAN\|Q9Y3Q8_T167 | 1.997772243 | 6.56E-07 | 6.24E-05 |
| IWS1_HUMAN\|Q96ST2_T53 | 1.996823869 | 1.90E-04 | 1.60E-03 |
| WAC2A_HUMAN\|Q641Q2_S284 | 1.996641013 | 1.23E-05 | 3.51E-04 |
| MK01_HUMAN\|P28482_T185_Y187 | 1.995795616 | 9.14E-06 | 3.78E-04 |
| OTU7B_HUMAN\|Q6GQQ9_S449 | 1.99374368 | 1.63E-03 | 1.40E-02 |
| K2C5_HUMAN\|P13647_S21_S26 | 1.993233671 | 4.96E-06 | 3.06E-04 |
| SLTM_HUMAN\|Q9NWH9_S371 | 1.992630867 | 3.64E-03 | 2.12E-02 |

| | | | |
|---|---|---|---|
| AHNK_HUMAN\|Q09666_S2397 | 1.992610648 | 6.78E-06 | 2.24E-04 |
| DC1L1_HUMAN\|Q9Y6G9_S412 | 1.992275928 | 5.92E-03 | 4.49E-02 |
| AHNK_HUMAN\|Q09666_S5746_S5752 | 1.991936678 | 4.37E-08 | 1.25E-05 |
| ATRX_HUMAN\|P46100_S784 | 1.991108975 | 1.39E-05 | 1.89E-04 |
| SPD2A_HUMAN\|Q5TCZ1_S651 | 1.990245606 | 9.08E-06 | 4.52E-04 |
| LAP2A_HUMAN\|P42166_S66_S67_T74 | 1.989999297 | 2.73E-06 | 1.56E-04 |
| K1C17_HUMAN\|Q04695_S11_S13 | 1.989753329 | 2.06E-06 | 6.96E-05 |
| RD23B_HUMAN\|P54727_T162 | 1.988733428 | 1.34E-04 | 4.01E-03 |
| NEB2_HUMAN\|Q96SB3_S100 | 1.987977027 | 4.01E-05 | 1.52E-03 |
| MK01_HUMAN\|P28482_T185_Y187 | 1.987917501 | 9.87E-06 | 1.86E-03 |
| MAP7_HUMAN\|Q14244_S183 | 1.987829312 | 2.79E-08 | 3.36E-06 |
| B4DKV4_HUMAN\|B4DKV4_S19_S22 | 1.987409698 | 3.26E-04 | 5.13E-03 |
| MKL2_HUMAN\|Q9ULH7_S921 | 1.9860525 | 5.40E-07 | 6.65E-05 |
| TB10A_HUMAN\|Q9BXI6_S40_S45 | 1.985454598 | 5.53E-04 | 3.43E-03 |
| NDRG1_HUMAN\|Q92597_S333_S336 | 1.982605026 | 1.56E-06 | 1.17E-04 |
| AMPD2_HUMAN\|Q01433_S100 | 1.982105528 | 1.04E-03 | 1.59E-02 |
| AAK1_HUMAN\|Q2M2I8_T620 | 1.981652027 | 4.37E-07 | 3.20E-05 |
| AKTS1_HUMAN\|Q96B36_T246 | 1.981387226 | 1.10E-05 | 4.30E-04 |
| MB12A_HUMAN\|Q96EY5_S188 | 1.981056153 | 2.11E-05 | 8.07E-04 |
| MTCL1_HUMAN\|Q9Y4B5_S1679_T1675 | 1.980276643 | 4.95E-08 | 4.85E-06 |
| JUND_HUMAN\|P17535_S100 | 1.980047486 | 2.74E-06 | 2.85E-04 |
| SRC8_HUMAN\|Q14247_S405_S418_T401 | 1.979548181 | 9.58E-05 | 2.78E-03 |
| ZO2_HUMAN\|Q9UDY2_S398 | 1.979514275 | 3.75E-05 | 4.24E-04 |
| K2C6A_HUMAN\|P02538_S60 | 1.979232681 | 1.64E-03 | 1.79E-02 |
| VIP2_HUMAN\|O43314_S1006 | 1.977964685 | 2.67E-08 | 8.12E-06 |
| PAWR_HUMAN\|Q96IZ0_S231 | 1.977880802 | 1.71E-07 | 1.16E-04 |
| IASPP_HUMAN\|Q8WUF5_S526 | 1.977591048 | 4.94E-04 | 5.50E-03 |
| LARP1_HUMAN\|Q6PKG0_S766_S774 | 1.977563733 | 3.05E-05 | 1.06E-03 |
| LPP_HUMAN\|Q93052_S609 | 1.977281552 | 1.95E-05 | 6.31E-04 |
| TB182_HUMAN\|Q9C0C2_T875 | 1.977110363 | 1.42E-07 | 3.09E-05 |
| STAT1_HUMAN\|P42224_S727 | 1.975709242 | 2.38E-07 | 3.68E-05 |

| | | | |
|---|---|---|---|
| K2C5_HUMAN\|P13647_S16_S21 | 1.975243018 | 1.31E-04 | 1.64E-03 |
| LARP1_HUMAN\|Q6PKG0_S766 | 1.974271305 | 6.89E-04 | 6.01E-03 |
| MTAP2_HUMAN\|P11137_S1782 | 1.974010375 | 1.92E-04 | 1.61E-03 |
| DESP_HUMAN\|P15924_T13 | 1.973989036 | 2.28E-03 | 2.28E-02 |
| K1C14_HUMAN\|P02533_S44_S53_S54 | 1.973900224 | 2.16E-07 | 3.54E-05 |
| K1C17_HUMAN\|Q04695_S11_S12 | 1.973833476 | 5.34E-06 | 1.41E-04 |
| PDS5A_HUMAN\|Q29RF7_S1276 | 1.973202479 | 4.06E-06 | 1.52E-04 |
| MTAP2_HUMAN\|P11137_T1602_T1605 | 1.973165876 | 1.04E-04 | 1.76E-03 |
| ACAP2_HUMAN\|Q15057_S381_S384 | 1.973115635 | 7.78E-07 | 4.68E-04 |
| SARG_HUMAN\|Q9BW04_S131 | 1.972235182 | 1.27E-04 | 2.53E-03 |
| COHA1_HUMAN\|Q9UMD9_T182 | 1.971964526 | 1.44E-05 | 6.29E-04 |
| PDLI2_HUMAN\|Q96JY6_S127_S129 | 1.971752178 | 2.39E-05 | 5.69E-04 |
| SMAG2_HUMAN\|Q5PRF9_S271 | 1.971518476 | 2.65E-05 | 6.06E-04 |
| PKP4_HUMAN\|Q99569_S233 | 1.971102228 | 1.62E-07 | 3.13E-05 |
| ACSA_HUMAN\|Q9NR19_S267 | 1.97101848 | 8.21E-07 | 9.89E-05 |
| HS90A_HUMAN\|P07900_S726 | 1.971009182 | 2.39E-03 | 2.34E-02 |
| PR38B_HUMAN\|Q5VTL8_S529 | 1.970993453 | 2.84E-07 | 4.40E-05 |
| AAK1_HUMAN\|Q2M2I8_T620 | 1.969400913 | 5.02E-07 | 1.56E-05 |
| TB10A_HUMAN\|Q9BXI6_S40_S45 | 1.969272125 | 6.19E-04 | 1.09E-02 |
| TB182_HUMAN\|Q9C0C2_S1138 | 1.968584428 | 2.04E-03 | 2.58E-02 |
| EDC4_HUMAN\|Q6P2E9_S723 | 1.968106775 | 1.86E-03 | 3.27E-02 |
| JUND_HUMAN\|P17535_S100 | 1.967695145 | 3.16E-06 | 2.49E-04 |
| PLEC_HUMAN\|Q15149_S4384_S4386_S4389 | 1.966745149 | 1.44E-03 | 1.23E-02 |
| MACF1_HUMAN\|Q9UPN3_T7287 | 1.966626594 | 1.15E-04 | 2.33E-03 |
| DESP_HUMAN\|P15924_T13 | 1.966320334 | 2.39E-03 | 2.42E-02 |
| SCAM2_HUMAN\|O15127_S319 | 1.965502067 | 2.79E-05 | 3.31E-04 |
| DC1L1_HUMAN\|Q9Y6G9_S415 | 1.965328825 | 2.16E-03 | 1.73E-02 |
| DDX3X_HUMAN\|O00571_S612 | 1.965132802 | 6.68E-04 | 1.18E-02 |
| CR025_HUMAN\|Q96B23_S143_S145_S147 | 1.965004019 | 5.23E-06 | 3.81E-04 |
| PAXI_HUMAN\|P49023_S103_S106 | 1.964996315 | 3.53E-05 | 1.53E-03 |
| K1C14_HUMAN\|P02533_S51_S54 | 1.964496593 | 1.07E-05 | 5.10E-04 |

| | | | |
|---|---|---|---|
| STMN1_HUMAN\|P16949_S25 | 1.963781942 | 7.74E-08 | 1.46E-05 |
| TTP_HUMAN\|P26651_S60 | 1.963568341 | 5.12E-07 | 2.31E-04 |
| AHNK_HUMAN\|Q09666_S5745_S5749 | 1.963422481 | 1.89E-08 | 7.15E-06 |
| STAT1_HUMAN\|P42224_S727 | 1.96338415 | 2.78E-07 | 9.01E-06 |
| ZO1_HUMAN\|Q07157_S841_Y853 | 1.963263497 | 3.29E-05 | 3.81E-04 |
| HEAT6_HUMAN\|Q6AI08_S395_S397 | 1.96310806 | 1.20E-03 | 1.08E-02 |
| HSPB1_HUMAN\|P04792_S78_S82 | 1.963078221 | 4.10E-05 | 1.53E-03 |
| K1C16_HUMAN\|P08779_S448_S451 | 1.962432542 | 1.70E-06 | 1.51E-04 |
| DBNL_HUMAN\|Q9UJU6_T270 | 1.960649958 | 1.90E-05 | 9.31E-04 |
| MISP_HUMAN\|Q8IVT2_S155 | 1.959553189 | 3.03E-05 | 8.87E-04 |
| TM230_HUMAN\|Q96A57_T25 | 1.957639441 | 3.34E-05 | 3.79E-04 |
| CNKR1_HUMAN\|Q969H4_S305_S314 | 1.957280181 | 6.18E-06 | 1.34E-03 |
| K2C5_HUMAN\|P13647_S77_S82 | 1.956288313 | 2.44E-04 | 2.70E-03 |
| PHLB2_HUMAN\|Q86SQ0_S334_T328 | 1.955854695 | 5.67E-05 | 1.41E-03 |
| CAVN1_HUMAN\|Q6NZI2_T302 | 1.955434141 | 3.91E-08 | 6.06E-06 |
| NDRG1_HUMAN\|Q92597_S333_S336_T328 | 1.955183395 | 3.53E-08 | 1.06E-05 |
| TBC15_HUMAN\|Q8TC07-2_S205 | 1.954840407 | 1.18E-05 | 6.90E-04 |
| K2C5_HUMAN\|P13647_S18 | 1.954273936 | 8.55E-07 | 7.24E-05 |
| NUP50_HUMAN\|Q9UKX7_S221 | 1.953866998 | 7.64E-08 | 5.91E-05 |
| CT2NL_HUMAN\|Q9P2B4_S443 | 1.953731035 | 4.18E-03 | 3.58E-02 |
| K2C8_HUMAN\|P05787_S24 | 1.953546421 | 1.06E-05 | 5.07E-04 |
| DPYL2_HUMAN\|Q16555_S570 | 1.953108716 | 1.40E-04 | 3.02E-03 |
| PKP1_HUMAN\|Q13835_S185 | 1.953094694 | 4.56E-04 | 7.19E-03 |
| ARG28_HUMAN\|Q8N1W1_S476_S486 | 1.951850575 | 2.64E-06 | 2.24E-04 |
| TISB_HUMAN\|Q07352_S336 | 1.9514007 | 1.33E-05 | 2.85E-04 |
| TB10A_HUMAN\|Q9BXI6_S40_S45 | 1.949952262 | 7.10E-04 | 7.28E-03 |
| GGYF2_HUMAN\|Q6Y7W6_S30 | 1.948889479 | 4.96E-04 | 9.21E-03 |
| ACINU_HUMAN\|Q9UKV3_S328 | 1.948033927 | 1.22E-03 | 1.66E-02 |
| CLIP2_HUMAN\|Q9UDT6_S207_T198 | 1.947510537 | 2.31E-05 | 2.84E-04 |
| FUBP2_HUMAN\|Q92945_T195 | 1.947483152 | 5.66E-04 | 1.00E-02 |
| STXB5_HUMAN\|Q5T5C0-2_S712 | 1.946853597 | 3.50E-08 | 9.03E-06 |

| | | | |
|---|---|---|---|
| SPD2A_HUMAN\|Q5TCZ1_S1043 | 1.946663792 | 1.43E-05 | 2.95E-04 |
| PHRF1_HUMAN\|Q9P1Y6_S915_T917 | 1.946524189 | 4.14E-04 | 6.76E-03 |
| DESP_HUMAN\|P15924_S166 | 1.94631304 | 8.35E-06 | 2.69E-04 |
| RBM7_HUMAN\|Q9Y580_S136 | 1.946288616 | 7.58E-06 | 2.52E-04 |
| STXB5_HUMAN\|Q5T5C0-2_S712 | 1.946203322 | 3.53E-08 | 5.80E-06 |
| ROA1_HUMAN\|P09651_S361_S364 | 1.945835823 | 2.87E-05 | 6.51E-04 |
| HNRH3_HUMAN\|P31942_S298 | 1.945787688 | 1.04E-06 | 8.52E-05 |
| BD1L1_HUMAN\|Q8NFC6_S2986 | 1.945694882 | 5.94E-05 | 8.77E-04 |
| K1C14_HUMAN\|P02533_S32 | 1.945683807 | 3.32E-07 | 4.41E-05 |
| K1C14_HUMAN\|P02533_S51_S54 | 1.945065816 | 1.33E-05 | 4.63E-04 |
| K1C19_HUMAN\|P08727_S13 | 1.943656814 | 4.15E-05 | 1.12E-03 |
| MPLKI_HUMAN\|Q8TAP9_S80 | 1.943570282 | 3.51E-04 | 5.97E-03 |
| MAP4_HUMAN\|P27816_S928 | 1.943480936 | 8.42E-07 | 3.65E-05 |
| KMT2A_HUMAN\|Q03164_T3028 | 1.943479023 | 4.95E-04 | 1.81E-02 |
| K2C5_HUMAN\|P13647_S82_T78 | 1.943425892 | 4.18E-07 | 2.22E-05 |
| LYRIC_HUMAN\|Q86UE4_S298 | 1.943110399 | 2.09E-04 | 1.72E-03 |
| K2C6A_HUMAN\|P02538_T548 | 1.942998923 | 1.88E-03 | 1.97E-02 |
| ENSA_HUMAN\|O43768_S109 | 1.942954284 | 4.01E-06 | 1.04E-03 |
| COHA1_HUMAN\|Q9UMD9_S93_T97 | 1.942663846 | 4.42E-05 | 3.70E-03 |
| SNX6_HUMAN\|Q9UNH7_S316 | 1.942592852 | 4.91E-03 | 4.59E-02 |
| MILK1_HUMAN\|Q8N3F8_T318 | 1.942418975 | 1.64E-04 | 1.96E-03 |
| MK01_HUMAN\|P28482_Y187 | 1.942093603 | 7.56E-06 | 1.58E-03 |
| EIF3C_HUMAN\|Q99613_S909 | 1.941111121 | 4.75E-03 | 2.99E-02 |
| PKP3_HUMAN\|Q9Y446_S183 | 1.940859775 | 6.42E-06 | 2.95E-04 |
| OTU7B_HUMAN\|Q6GQQ9_S449 | 1.940586873 | 2.26E-03 | 1.71E-02 |
| HS90A_HUMAN\|P07900_S726 | 1.940347653 | 2.86E-03 | 2.76E-02 |
| CCDC6_HUMAN\|Q16204_S244 | 1.939637776 | 9.89E-06 | 1.49E-04 |
| COHA1_HUMAN\|Q9UMD9_S93_T97 | 1.939426836 | 4.56E-05 | 1.20E-03 |
| VIME_HUMAN\|P08670_S34 | 1.939231221 | 1.59E-06 | 1.17E-04 |
| ARI4B_HUMAN\|Q4LE39_S790 | 1.938625626 | 3.74E-03 | 1.79E-02 |
| PININ_HUMAN\|Q9H307_S690 | 1.938431126 | 2.79E-05 | 3.31E-04 |

| | | | |
|---|---|---|---|
| JAM1_HUMAN\|Q9Y624_S284 | 1.938279807 | 2.01E-03 | 4.36E-02 |
| K1C14_HUMAN\|P02533_S462 | 1.937643828 | 1.44E-05 | 7.80E-04 |
| NBEL2_HUMAN\|Q6ZNJ1_T1867 | 1.935664167 | 2.39E-05 | 1.13E-03 |
| CCDC6_HUMAN\|Q16204_S244 | 1.935641197 | 1.03E-05 | 1.49E-04 |
| CCD86_HUMAN\|Q9H6F5_S217 | 1.935328188 | 1.79E-05 | 4.74E-04 |
| ACAP2_HUMAN\|Q15057_S384_T382 | 1.935267279 | 2.05E-10 | 1.11E-06 |
| DDX3X_HUMAN\|O00571_S612 | 1.93474269 | 8.27E-04 | 1.12E-02 |
| KLC3_HUMAN\|Q6P597_S497_S502 | 1.933992265 | 6.37E-09 | 3.14E-06 |
| CHSP1_HUMAN\|Q9Y2V2_S52 | 1.933408172 | 1.27E-05 | 5.85E-04 |
| PAWR_HUMAN\|Q96IZ0_S231 | 1.933137591 | 3.21E-07 | 4.41E-05 |
| ARG35_HUMAN\|A5YM69_S450 | 1.932770099 | 3.02E-05 | 1.34E-03 |
| SOS1_HUMAN\|Q07889_S1082 | 1.932670485 | 1.81E-03 | 1.27E-02 |
| AKAP2_HUMAN\|Q9Y2D5_S152_T157 | 1.932569714 | 4.53E-07 | 5.12E-05 |
| UFD1_HUMAN\|Q92890_S299 | 1.932158204 | 1.47E-04 | 3.85E-03 |
| COTL1_HUMAN\|Q14019_T141 | 1.931577831 | 1.40E-03 | 1.94E-02 |
| PPM1G_HUMAN\|O15355_S183 | 1.931542282 | 1.30E-06 | 1.30E-04 |
| AHNK_HUMAN\|Q09666_S5864 | 1.931295281 | 4.02E-04 | 4.79E-03 |
| NAV1_HUMAN\|Q8NEY1_T1006 | 1.930396113 | 4.49E-05 | 1.19E-03 |
| PAXI_HUMAN\|P49023_S103_S106 | 1.930009349 | 4.99E-05 | 2.08E-03 |
| K2C8_HUMAN\|P05787_S21_S24_S27 | 1.929762752 | 3.21E-06 | 9.67E-05 |
| AMOL2_HUMAN\|Q9Y2J4_S759_S762 | 1.929528909 | 3.68E-04 | 3.72E-03 |
| R3HD2_HUMAN\|Q9Y2K5_T856 | 1.929182037 | 9.23E-05 | 1.51E-03 |
| NIBL1_HUMAN\|Q96TA1_S692_S696 | 1.92914738 | 2.73E-06 | 5.40E-05 |
| COF1_HUMAN\|P23528_S24 | 1.929098085 | 5.50E-05 | 1.84E-03 |
| K2C5_HUMAN\|P13647_S21_S26 | 1.928792795 | 1.03E-05 | 6.28E-04 |
| MAP7_HUMAN\|Q14244_T184 | 1.928486912 | 3.63E-07 | 4.69E-05 |
| AHNK_HUMAN\|Q09666_S5864 | 1.928048642 | 4.12E-04 | 4.04E-03 |
| BAIP2_HUMAN\|Q9UQB8_T340 | 1.928011923 | 1.69E-04 | 2.01E-03 |
| RHG12_HUMAN\|Q8IWW6_S215 | 1.926838992 | 8.20E-07 | 9.25E-05 |
| MPZL1_HUMAN\|O95297_S221 | 1.926718092 | 1.33E-03 | 1.23E-02 |
| RELB_HUMAN\|Q01201_S573 | 1.926582134 | 3.95E-07 | 2.43E-05 |

| | | | |
|---|---|---|---|
| NGAP_HUMAN\|Q9UJF2_S954 | 1.926426317 | 2.44E-06 | 2.17E-04 |
| B4DKV4_HUMAN\|B4DKV4_S22 | 1.926096927 | 7.09E-04 | 6.13E-03 |
| ARHG5_HUMAN\|Q12774_S450 | 1.925986289 | 2.73E-05 | 3.19E-04 |
| ROA1_HUMAN\|P09651_S363 | 1.925341225 | 1.54E-03 | 2.07E-02 |
| VIME_HUMAN\|P08670_S51 | 1.924995736 | 2.93E-06 | 1.62E-04 |
| P63_HUMAN\|Q9H3D4_S463_T451 | 1.924762465 | 4.30E-04 | 5.02E-03 |
| IPP2M_HUMAN\|Q6NXS1_S122 | 1.924467761 | 2.65E-06 | 5.51E-05 |
| LAP2A_HUMAN\|P42166_S156_T160 | 1.924406614 | 7.24E-05 | 5.16E-03 |
| ROA1_HUMAN\|P09651_S364 | 1.923815998 | 4.00E-04 | 4.77E-03 |
| CALD1_HUMAN\|Q05682_S789 | 1.923735763 | 5.97E-06 | 3.86E-04 |
| DC1L1_HUMAN\|Q9Y6G9_S516 | 1.92217431 | 2.63E-07 | 8.69E-06 |
| IASPP_HUMAN\|Q8WUF5_T341 | 1.922083407 | 8.65E-04 | 2.67E-02 |
| AHNK_HUMAN\|Q09666_T5824 | 1.922029293 | 1.86E-03 | 2.20E-02 |
| RL34_HUMAN\|P49207_S12 | 1.921636177 | 4.60E-03 | 3.74E-02 |
| MED24_HUMAN\|O75448_S862_S873 | 1.921578914 | 2.08E-04 | 2.91E-03 |
| NUP98_HUMAN\|P52948_S612 | 1.92148636 | 3.57E-07 | 1.12E-05 |
| CND1_HUMAN\|Q15021_S1330_S1333 | 1.921176315 | 4.38E-04 | 7.07E-03 |
| MISP_HUMAN\|Q8IVT2_S395_S397 | 1.920868365 | 7.08E-06 | 3.15E-04 |
| COHA1_HUMAN\|Q9UMD9_S93_S96 | 1.919830603 | 7.54E-05 | 1.71E-03 |
| K2C5_HUMAN\|P13647_S82_T78 | 1.919760853 | 5.83E-07 | 6.87E-05 |
| HIRP3_HUMAN\|Q9BW71_S551 | 1.919480416 | 3.84E-04 | 8.07E-03 |
| NEB2_HUMAN\|Q96SB3_S100 | 1.919258372 | 7.48E-05 | 7.52E-04 |
| K2C6A_HUMAN\|P02538_S60 | 1.919220596 | 2.44E-03 | 1.59E-02 |
| SRC8_HUMAN\|Q14247_S405_S418_T401 | 1.919092964 | 1.64E-04 | 4.41E-03 |
| EPN1_HUMAN\|Q9Y6I3_S454 | 1.918754203 | 1.54E-07 | 1.21E-05 |
| NIBL1_HUMAN\|Q96TA1_S641_S646 | 1.918584206 | 6.63E-05 | 1.19E-03 |
| CE170_HUMAN\|Q5SW79_S633_S636 | 1.917660602 | 1.95E-04 | 4.60E-03 |
| NAV1_HUMAN\|Q8NEY1_S648 | 1.91755521 | 1.02E-03 | 2.86E-02 |
| SENP3_HUMAN\|Q9H4L4_S169_S181_T176 | 1.91741022 | 1.66E-04 | 1.99E-03 |
| UBXN7_HUMAN\|O94888_S285 | 1.917352305 | 9.49E-05 | 2.32E-03 |
| SF3B2_HUMAN\|Q13435_S307_S309 | 1.917246527 | 3.17E-04 | 3.97E-03 |

| | | | |
|---|---|---|---|
| MAGD2_HUMAN|Q9UNF1_S190_S191 | 1.917072917 | 8.31E-04 | 4.75E-03 |
| CF132_HUMAN|Q5T0Z8_T297 | 1.916878115 | 6.30E-05 | 1.71E-03 |
| ENAH_HUMAN|Q8N8S7_T489 | 1.916284026 | 1.98E-03 | 4.49E-02 |
| TERA_HUMAN|P55072_S794 | 1.916146911 | 6.22E-05 | 1.52E-03 |
| JUNB_HUMAN|P17275_T255 | 1.915996373 | 9.67E-09 | 1.45E-06 |
| DBNL_HUMAN|Q9UJU6_T271 | 1.91576919 | 1.59E-06 | 1.17E-04 |
| ACINU_HUMAN|Q9UKV3_S328 | 1.915659532 | 1.52E-03 | 1.66E-02 |
| CCDC9_HUMAN|Q9Y3X0_S386_S390 | 1.915620524 | 1.74E-04 | 1.50E-03 |
| COHA1_HUMAN|Q9UMD9_T182 | 1.915409271 | 2.58E-05 | 4.70E-04 |
| TM230_HUMAN|Q96A57_T25 | 1.915349486 | 5.25E-05 | 1.80E-03 |
| COF1_HUMAN|P23528_S24 | 1.914896131 | 6.30E-05 | 2.23E-03 |
| PACN3_HUMAN|Q9UKS6_S344_S354 | 1.9143786 | 5.63E-07 | 1.57E-05 |
| PDLI2_HUMAN|Q96JY6_S137_S143 | 1.913732042 | 1.26E-06 | 4.81E-05 |
| AHNK2_HUMAN|Q8IVF2_S1710 | 1.913690251 | 5.25E-06 | 4.43E-04 |
| K2C6B_HUMAN|P04259_S34 | 1.912996723 | 2.27E-04 | 4.02E-03 |
| ACAP2_HUMAN|Q15057_S384 | 1.912632442 | 3.03E-08 | 4.11E-05 |
| K2C6A_HUMAN|P02538_S541 | 1.912421521 | 2.08E-03 | 2.13E-02 |
| DDX10_HUMAN|Q13206_S831 | 1.911437377 | 9.68E-07 | 4.91E-05 |
| SRRM1_HUMAN|Q8IYB3_S452 | 1.911086616 | 9.94E-03 | 4.51E-02 |
| SARG_HUMAN|Q9BW04_T151 | 1.910849695 | 5.46E-04 | 5.08E-03 |
| SEPT7_HUMAN|Q16181_S423 | 1.910750169 | 6.92E-05 | 1.81E-03 |
| EIF3E_HUMAN|P60228_S442 | 1.91047761 | 2.70E-04 | 6.18E-03 |
| UBXN7_HUMAN|O94888_S280_S288 | 1.909483728 | 1.58E-04 | 3.34E-03 |
| MISP_HUMAN|Q8IVT2_S395_S400 | 1.908540263 | 1.71E-04 | 3.50E-03 |
| ADCY6_HUMAN|O43306_S54 | 1.907402773 | 1.25E-06 | 1.23E-04 |
| EF2_HUMAN|P13639_T57 | 1.906259061 | 1.14E-03 | 1.03E-02 |
| K1C14_HUMAN|P02533_S28 | 1.905994821 | 9.00E-07 | 3.78E-05 |
| RAI14_HUMAN|Q9P0K7_S290 | 1.905767908 | 5.91E-05 | 8.77E-04 |
| MAGD2_HUMAN|Q9UNF1_S190_S191 | 1.905629529 | 9.02E-04 | 8.83E-03 |
| CD97_HUMAN|P48960_S831 | 1.905444209 | 5.08E-07 | 5.41E-05 |
| FOXK1_HUMAN|P85037_T436 | 1.905417212 | 6.44E-03 | 2.82E-02 |

| | | | |
|---|---|---|---|
| CAMP2_HUMAN\|Q08AD1_S980 | 1.905052464 | 1.53E-05 | 5.15E-04 |
| CAVN1_HUMAN\|Q6NZI2_T302 | 1.904720917 | 8.06E-08 | 1.75E-05 |
| PKP3_HUMAN\|Q9Y446_S280 | 1.904522901 | 8.35E-05 | 2.11E-03 |
| K1C14_HUMAN\|P02533_S39 | 1.903895578 | 6.31E-04 | 5.68E-03 |
| NUP98_HUMAN\|P52948_S623 | 1.903819666 | 8.88E-06 | 1.34E-04 |
| UFD1_HUMAN\|Q92890_S299 | 1.903559576 | 1.88E-04 | 3.80E-03 |
| K1C14_HUMAN\|P02533_Y46 | 1.903520559 | 5.89E-06 | 2.78E-04 |
| TBC15_HUMAN\|Q8TC07_S70 | 1.903469589 | 5.77E-04 | 8.66E-03 |
| K1C14_HUMAN\|P02533_S54_Y46 | 1.903341898 | 1.70E-05 | 3.38E-04 |
| PAR10_HUMAN\|Q53GL7_S431 | 1.902830659 | 1.68E-07 | 6.41E-06 |
| ARHG5_HUMAN\|Q12774_S15 | 1.902141467 | 8.18E-03 | 3.89E-02 |
| PSF2_HUMAN\|Q9Y248_S182_T180 | 1.901365059 | 3.24E-05 | 1.38E-03 |
| STXB5_HUMAN\|Q5T5C0-2_S712 | 1.901256282 | 6.61E-08 | 3.16E-05 |
| NBEL2_HUMAN\|Q6ZNJ1_S2739 | 1.901121933 | 9.66E-04 | 9.57E-03 |
| ROA1_HUMAN\|P09651_S361_S364 | 1.901023858 | 4.56E-05 | 1.95E-03 |
| SEC62_HUMAN\|Q99442_T375 | 1.900720922 | 4.29E-04 | 2.81E-03 |
| STXB5_HUMAN\|Q5T5C0-2_S712 | 1.900621237 | 6.67E-08 | 1.72E-05 |
| NDRG1_HUMAN\|Q92597_S333_S336 | 1.900345991 | 4.22E-06 | 2.76E-04 |
| MAP7_HUMAN\|Q14244_S183 | 1.900332518 | 1.03E-07 | 2.42E-05 |
| WFS1_HUMAN\|O76024_S235 | 1.900290433 | 4.25E-04 | 3.00E-03 |
| FOSL2_HUMAN\|P15408_S19 | 1.900284451 | 2.57E-04 | 5.96E-03 |
| LAR4B_HUMAN\|Q92615_S731_S736 | 1.90026599 | 4.27E-06 | 2.76E-04 |
| K1522_HUMAN\|Q9P206_S404_S409 | 1.899303453 | 2.40E-07 | 3.72E-05 |
| CR025_HUMAN\|Q96B23_S143_S145_S147 | 1.899289357 | 1.11E-05 | 1.64E-04 |
| CCDC9_HUMAN\|Q9Y3X0_S376_S386_T381 | 1.898866916 | 1.15E-06 | 2.49E-04 |
| K2C5_HUMAN\|P13647_S16_S26 | 1.898483203 | 3.90E-04 | 3.89E-03 |
| RAI14_HUMAN\|Q9P0K7_S281 | 1.898216209 | 1.64E-07 | 2.70E-05 |
| DNJC1_HUMAN\|Q96KC8_S381 | 1.897967003 | 4.90E-04 | 3.11E-03 |
| MINK1_HUMAN\|Q8N4C8_S701 | 1.897846379 | 2.89E-04 | 4.72E-03 |
| PKP3_HUMAN\|Q9Y446_S180_S183 | 1.897393365 | 1.27E-05 | 1.63E-03 |
| NOLC1_HUMAN\|Q14978_S688_S698 | 1.896614442 | 5.64E-03 | 2.91E-02 |

| | | | |
|---|---|---|---|
| PKP4_HUMAN\|Q99569_S236 | 1.896356421 | 1.35E-06 | 1.29E-04 |
| RAI14_HUMAN\|Q9P0K7_S281_S290 | 1.8961148 | 2.58E-04 | 3.43E-03 |
| K1C14_HUMAN\|P02533_S32 | 1.896089267 | 6.79E-07 | 6.24E-05 |
| K1522_HUMAN\|Q9P206_S404_S409 | 1.895975234 | 2.53E-07 | 3.95E-05 |
| KC1A_HUMAN\|P48729-3_T321 | 1.895472669 | 1.44E-03 | 1.66E-02 |
| AHNK_HUMAN\|Q09666_S210_S216_S220 | 1.89523411 | 2.81E-07 | 4.14E-05 |
| IWS1_HUMAN\|Q96ST2_S313 | 1.895181272 | 1.02E-03 | 6.16E-03 |
| WDR44_HUMAN\|Q5JSH3_S561 | 1.894615783 | 5.63E-07 | 3.81E-04 |
| SRC8_HUMAN\|Q14247_S405_T401 | 1.894546998 | 2.72E-04 | 6.28E-03 |
| PDLI2_HUMAN\|Q96JY6_S137 | 1.893964772 | 4.19E-07 | 2.52E-05 |
| K2C5_HUMAN\|P13647_S36 | 1.893804934 | 4.61E-06 | 2.94E-04 |
| FOSL2_HUMAN\|P15408_S320 | 1.892879742 | 9.82E-04 | 5.97E-03 |
| SRC8_HUMAN\|Q14247_S405_S418_T401 | 1.89256965 | 2.08E-04 | 1.61E-03 |
| AKTS1_HUMAN\|Q96B36_S203 | 1.892550026 | 5.92E-04 | 3.60E-03 |
| PEA15_HUMAN\|Q15121_S104 | 1.892409805 | 5.40E-05 | 1.03E-03 |
| FUBP2_HUMAN\|Q92945_T195 | 1.891456682 | 8.60E-04 | 1.37E-02 |
| HOME3_HUMAN\|Q9NSC5_S159 | 1.891028553 | 4.72E-04 | 5.35E-03 |
| SMCR8_HUMAN\|Q8TEV9_S417 | 1.889143899 | 4.60E-03 | 3.74E-02 |
| ROA1_HUMAN\|P09651_S368 | 1.888991838 | 7.78E-04 | 1.25E-02 |
| PR38B_HUMAN\|Q5VTL8_S529 | 1.888591357 | 9.10E-07 | 2.06E-04 |
| TM237_HUMAN\|Q96Q45_S62 | 1.888161865 | 1.33E-03 | 3.36E-02 |
| AKTS1_HUMAN\|Q96B36_S203 | 1.888156574 | 6.14E-04 | 6.53E-03 |
| MILK1_HUMAN\|Q8N3F8_S640_S644 | 1.887803748 | 2.23E-06 | 7.32E-05 |
| TBC15_HUMAN\|Q8TC07_S70 | 1.887736351 | 6.50E-04 | 8.79E-03 |
| K1C18_HUMAN\|P05783_S7 | 1.887699718 | 6.59E-05 | 5.10E-03 |
| ITA2_HUMAN\|P17301_S1181 | 1.887212392 | 3.42E-05 | 1.12E-03 |
| ASPP2_HUMAN\|Q13625_S556 | 1.887069307 | 3.68E-06 | 1.44E-04 |
| GTR1_HUMAN\|P11166_S490 | 1.886909759 | 4.71E-03 | 3.79E-02 |
| FA83G_HUMAN\|A6ND36_S726 | 1.886560559 | 1.20E-05 | 4.45E-04 |
| DC1L1_HUMAN\|Q9Y6G9_S415 | 1.886475502 | 3.49E-03 | 3.69E-02 |
| LAD1_HUMAN\|O00515_S38_T40_T41 | 1.885980916 | 1.62E-05 | 6.69E-04 |

| | | | |
|---|---|---|---|
| GRAP1_HUMAN\|Q4V328_S666 | 1.885438637 | 1.21E-05 | 4.45E-04 |
| IFIT3_HUMAN\|O14879_S484 | 1.88496225 | 1.37E-03 | 1.80E-02 |
| FUBP2_HUMAN\|Q92945_T195 | 1.884731346 | 9.05E-04 | 5.61E-03 |
| LAR4B_HUMAN\|Q92615_T732 | 1.884679206 | 9.90E-09 | 1.45E-06 |
| ROA1_HUMAN\|P09651_S363 | 1.883664219 | 2.06E-03 | 2.14E-02 |
| TPR_HUMAN\|P12270_S1847_T1839 | 1.883197224 | 2.59E-04 | 4.83E-03 |
| TBC9B_HUMAN\|Q66K14_S411 | 1.883160269 | 4.01E-05 | 6.48E-04 |
| SRPK2_HUMAN\|P78362_S494_T498 | 1.883160267 | 1.62E-04 | 1.31E-03 |
| FAT1_HUMAN\|Q14517_S4473 | 1.882992161 | 1.36E-04 | 2.96E-03 |
| NKTR_HUMAN\|P30414_S463 | 1.882756324 | 6.03E-06 | 1.02E-04 |
| TBC15_HUMAN\|Q8TC07_S70 | 1.882589966 | 6.77E-04 | 2.32E-02 |
| AFAD_HUMAN\|P55196_T1232 | 1.882522498 | 1.90E-05 | 3.72E-04 |
| RAI14_HUMAN\|Q9P0K7-4_S261 | 1.882440868 | 2.47E-06 | 5.23E-05 |
| IWS1_HUMAN\|Q96ST2_S398 | 1.882035568 | 5.08E-04 | 9.34E-03 |
| PAIRB_HUMAN\|Q8NC51_S392 | 1.881603706 | 7.00E-06 | 4.44E-04 |
| FLVC1_HUMAN\|Q9Y5Y0_S536 | 1.881289874 | 1.25E-05 | 1.63E-03 |
| AHNK2_HUMAN\|Q8IVF2_S457 | 1.881098069 | 3.60E-04 | 7.28E-03 |
| K2C5_HUMAN\|P13647_S16_S26 | 1.880987606 | 4.53E-04 | 8.71E-03 |
| HDGF_HUMAN\|P51858_S103 | 1.880795119 | 1.40E-03 | 1.04E-02 |
| SEPT9_HUMAN\|Q9UHD8_T38_T42 | 1.880611295 | 3.57E-07 | 1.93E-05 |
| CE170_HUMAN\|Q5SW79_S633_S636 | 1.879053047 | 2.73E-04 | 2.94E-03 |
| PHLB2_HUMAN\|Q86SQ0_S565 | 1.87905149 | 3.47E-07 | 4.16E-05 |
| OSBP1_HUMAN\|P22059_S379_S382_S386 | 1.87887996 | 8.88E-05 | 8.26E-04 |
| COHA1_HUMAN\|Q9UMD9_S85_S93_T97 | 1.878713975 | 1.12E-04 | 1.44E-03 |
| SP100_HUMAN\|P23497_S407_S410 | 1.87746978 | 7.14E-03 | 4.19E-02 |
| KPCD2_HUMAN\|Q9BZL6_S214 | 1.877365244 | 1.40E-05 | 4.00E-04 |
| CE170_HUMAN\|Q5SW79_S630_S633 | 1.87733533 | 2.83E-07 | 4.14E-05 |
| MAP1B_HUMAN\|P46821_S1792 | 1.877253028 | 1.41E-05 | 2.94E-04 |
| UTP18_HUMAN\|Q9Y5J1_S121 | 1.876548569 | 5.11E-03 | 3.13E-02 |
| MRCKG_HUMAN\|Q6DT37_S1544 | 1.876191452 | 6.16E-05 | 2.00E-03 |
| SF3B2_HUMAN\|Q13435_S307 | 1.875652856 | 1.19E-04 | 1.11E-03 |

| | | | |
|---|---|---|---|
| MTAP2_HUMAN\|P11137_S1782_S1790 | 1.875588672 | 5.37E-04 | 5.93E-03 |
| RBM7_HUMAN\|Q9Y580_S136 | 1.874846232 | 1.67E-05 | 9.66E-04 |
| K2C5_HUMAN\|P13647_T19 | 1.874755388 | 2.22E-04 | 2.51E-03 |
| HIRP3_HUMAN\|Q9BW71_S551 | 1.874511115 | 5.52E-04 | 1.42E-02 |
| TOP2B_HUMAN\|Q02880_S1550_S1552 | 1.874192324 | 1.44E-04 | 4.05E-03 |
| K2C6A_HUMAN\|P02538_S31 | 1.873727579 | 3.16E-03 | 2.86E-02 |
| LRRF2_HUMAN\|Q9Y608_S320_T317 | 1.872482756 | 1.35E-04 | 1.22E-03 |
| SRRM2_HUMAN\|Q9UQ35_S1383_S1387 | 1.872465736 | 3.50E-05 | 7.53E-04 |
| TTP_HUMAN\|P26651_S60 | 1.872190711 | 1.79E-06 | 1.72E-04 |
| K2C8_HUMAN\|P05787_S477 | 1.871209985 | 3.43E-05 | 1.12E-03 |
| TANK_HUMAN\|Q92844_S129 | 1.871085012 | 2.75E-05 | 4.90E-04 |
| ARG28_HUMAN\|Q8N1W1_S476_S486 | 1.870759549 | 7.10E-06 | 3.15E-04 |
| I2BPL_HUMAN\|Q9H1B7_S659 | 1.870576156 | 1.76E-03 | 4.06E-02 |
| MICA3_HUMAN\|Q7RTP6_S977 | 1.870292847 | 2.04E-09 | 1.59E-06 |
| HOME3_HUMAN\|Q9NSC5_S159 | 1.87017256 | 5.58E-04 | 1.01E-02 |
| SNX6_HUMAN\|Q9UNH7_S316 | 1.869914417 | 7.34E-03 | 2.68E-02 |
| WDR44_HUMAN\|Q5JSH3_S561 | 1.86936588 | 7.89E-07 | 9.10E-05 |
| CSCL2_HUMAN\|Q5T3F8_S111_S114 | 1.869215937 | 1.73E-04 | 2.48E-03 |
| K2C5_HUMAN\|P13647_S16_S26 | 1.868782722 | 5.03E-04 | 7.73E-03 |
| PHLB2_HUMAN\|Q86SQ0_S565 | 1.867944371 | 4.04E-07 | 5.09E-05 |
| M3K3_HUMAN\|Q99759_S337 | 1.866910517 | 1.93E-05 | 2.44E-04 |
| SHOT1_HUMAN\|A0MZ66_T537 | 1.86663764 | 1.68E-04 | 2.45E-03 |
| K2C5_HUMAN\|P13647_S571_S574 | 1.866239033 | 5.95E-06 | 1.53E-04 |
| K1C14_HUMAN\|P02533_S39 | 1.866015107 | 8.60E-04 | 1.07E-02 |
| PPM1G_HUMAN\|O15355_S201 | 1.864906983 | 3.87E-04 | 5.89E-03 |
| LAP2A_HUMAN\|P42166_S156_T160 | 1.864760485 | 1.26E-04 | 3.45E-03 |
| TTP_HUMAN\|P26651_S186 | 1.864241318 | 3.83E-04 | 1.57E-02 |
| ACINU_HUMAN\|Q9UKV3_S240 | 1.864115031 | 6.28E-06 | 1.04E-04 |
| K1C15_HUMAN\|P19012_S48 | 1.864092299 | 7.77E-04 | 1.25E-02 |
| MICA3_HUMAN\|Q7RTP6_S685_S687 | 1.864057975 | 1.60E-04 | 1.30E-03 |
| RAI14_HUMAN\|Q9P0K7_T251 | 1.863967523 | 1.31E-05 | 3.67E-04 |

| | | | |
|---|---|---|---|
| K1C14_HUMAN\|P02533_T9 | 1.863587028 | 4.54E-05 | 7.05E-04 |
| EGFR_HUMAN\|P00533_S1204 | 1.863554167 | 8.43E-05 | 1.84E-03 |
| DHX16_HUMAN\|O60231_S103 | 1.863248939 | 2.59E-03 | 1.33E-02 |
| IRS2_HUMAN\|Q9Y4H2_S1185_S1203 | 1.862995262 | 1.03E-05 | 3.05E-04 |
| IRS2_HUMAN\|Q9Y4H2_S608 | 1.862795332 | 4.57E-07 | 2.36E-05 |
| KPCD2_HUMAN\|Q9BZL6_S212 | 1.861786143 | 1.16E-05 | 6.49E-04 |
| SFR19_HUMAN\|Q9H7N4_T989 | 1.86001192 | 8.19E-04 | 8.21E-03 |
| DBNL_HUMAN\|Q9UJU6_T271 | 1.859524884 | 3.16E-06 | 9.57E-05 |
| HIRP3_HUMAN\|Q9BW71_S551 | 1.85906664 | 6.26E-04 | 6.58E-03 |
| SF3B2_HUMAN\|Q13435_S307 | 1.858997344 | 1.40E-04 | 2.15E-03 |
| CAMP1_HUMAN\|Q5T5Y3_S827 | 1.85887397 | 3.31E-05 | 1.37E-03 |
| LARP1_HUMAN\|Q6PKG0_S631_Y633 | 1.858798022 | 1.08E-03 | 8.46E-03 |
| JAM1_HUMAN\|Q9Y624_S296 | 1.858382445 | 4.35E-03 | 3.67E-02 |
| I2BPL_HUMAN\|Q9H1B7_S659 | 1.857819581 | 1.93E-03 | 2.26E-02 |
| CNKR1_HUMAN\|Q969H4_S305_S314 | 1.85738716 | 1.89E-05 | 6.39E-03 |
| TB182_HUMAN\|Q9C0C2_S691 | 1.857165433 | 2.31E-06 | 2.45E-04 |
| REPS1_HUMAN\|Q96D71_S162_S170_S174 | 1.85714818 | 1.60E-06 | 5.78E-05 |
| MK03_HUMAN\|P27361_T198_T202 | 1.856782412 | 1.47E-05 | 7.89E-04 |
| GGYF2_HUMAN\|Q6Y7W6_S201 | 1.856724849 | 4.21E-06 | 3.04E-04 |
| MEF2D_HUMAN\|Q14814_S231 | 1.856604783 | 2.32E-04 | 5.24E-03 |
| S6A15_HUMAN\|Q9H2J7_S699 | 1.85623648 | 2.15E-04 | 1.22E-02 |
| ZCCHV_HUMAN\|Q7Z2W4_T380 | 1.855547285 | 6.13E-06 | 3.54E-04 |
| MK01_HUMAN\|P28482_T185_Y187 | 1.855479632 | 3.82E-05 | 3.51E-03 |
| PPM1G_HUMAN\|O15355_S201 | 1.855400798 | 4.19E-04 | 4.92E-03 |
| IRS2_HUMAN\|Q9Y4H2_S608 | 1.855102789 | 5.09E-07 | 5.41E-05 |
| C2CD5_HUMAN\|Q86YS7_S307 | 1.854768101 | 2.07E-06 | 4.55E-05 |
| K2C6B_HUMAN\|P04259_S541 | 1.854706567 | 3.41E-06 | 1.01E-04 |
| GYS1_HUMAN\|P13807_S727 | 1.85445868 | 4.71E-07 | 6.23E-05 |
| B4DKV4_HUMAN\|B4DKV4_S19_S22 | 1.853944175 | 1.01E-03 | 8.00E-03 |
| WFS1_HUMAN\|O76024_S235 | 1.853774733 | 6.45E-04 | 3.87E-03 |
| PLEC_HUMAN\|Q15149_S4384_S4386_S4389 | 1.853529258 | 3.04E-03 | 4.46E-02 |

| | | | |
|---|---|---|---|
| EIF3C_HUMAN|Q99613_S909 | 1.853478211 | 7.77E-03 | 3.29E-02 |
| EPHA2_HUMAN|P29317_T771 | 1.853069487 | 2.54E-03 | 2.75E-02 |
| KLC3_HUMAN|Q6P597_S495_S502 | 1.852945357 | 1.93E-04 | 3.88E-03 |
| AAK1_HUMAN|Q2M2I8_T620 | 1.852828634 | 2.04E-06 | 1.88E-04 |
| GAB1_HUMAN|Q13480_S683 | 1.852793355 | 2.97E-07 | 2.04E-05 |
| NELFE_HUMAN|P18615_S51 | 1.851833004 | 1.76E-06 | 7.93E-05 |
| CR025_HUMAN|Q96B23_S143_S145 | 1.851831227 | 3.32E-07 | 6.66E-05 |
| AKTS1_HUMAN|Q96B36_T246 | 1.851748638 | 4.35E-05 | 1.32E-03 |
| AKTS1_HUMAN|Q96B36_S203 | 1.851660021 | 8.24E-04 | 1.27E-02 |
| GGYF1_HUMAN|O75420_S157 | 1.851548603 | 6.14E-04 | 6.60E-03 |
| HIRP3_HUMAN|Q9BW71_S551_S555 | 1.851372101 | 1.61E-03 | 2.14E-02 |
| XPC_HUMAN|Q01831_S883_S884_S891 | 1.850963751 | 7.04E-05 | 9.92E-04 |
| HOME3_HUMAN|Q9NSC5_S159 | 1.850811419 | 6.53E-04 | 3.90E-03 |
| SRRM1_HUMAN|Q8IYB3_S756 | 1.850689533 | 4.71E-04 | 5.39E-03 |
| PKP3_HUMAN|Q9Y446_S183 | 1.849559956 | 1.74E-05 | 7.10E-04 |
| HDGF_HUMAN|P51858_S103 | 1.849184772 | 1.76E-03 | 1.41E-02 |
| K1C13_HUMAN|P13646_S11 | 1.848434168 | 2.83E-03 | 2.74E-02 |
| PHF14_HUMAN|O94880_T287 | 1.847962833 | 6.81E-06 | 1.71E-04 |
| EF2K_HUMAN|O00418_S470 | 1.84735584 | 3.56E-06 | 1.01E-03 |
| CPSF4_HUMAN|O95639_S202 | 1.846776296 | 3.99E-05 | 1.52E-03 |
| GRSF1_HUMAN|Q12849_S118 | 1.846452881 | 1.96E-06 | 9.64E-05 |
| AHNK_HUMAN|Q09666_S572 | 1.84642849 | 2.77E-05 | 1.25E-03 |
| UBP5_HUMAN|P45974_S779 | 1.846013604 | 6.14E-05 | 2.13E-03 |
| ZO2_HUMAN|Q9UDY2_S400 | 1.845944207 | 2.37E-04 | 4.52E-03 |
| NIBAN_HUMAN|Q9BZQ8_S926 | 1.845757131 | 7.15E-07 | 1.25E-04 |
| TCF20_HUMAN|Q9UGU0_S538 | 1.844163569 | 4.34E-06 | 3.10E-04 |
| GGYF1_HUMAN|O75420_S157 | 1.844091542 | 6.52E-04 | 6.79E-03 |
| K1C14_HUMAN|P02533_S44 | 1.843565748 | 4.37E-05 | 3.70E-03 |
| SLK_HUMAN|Q9H2G2_S655 | 1.843208015 | 5.71E-05 | 2.07E-03 |
| NAV1_HUMAN|Q8NEY1_T1006 | 1.842502932 | 1.09E-04 | 2.50E-03 |
| SPD2A_HUMAN|Q5TCZ1_S651 | 1.841985227 | 4.07E-05 | 1.12E-03 |

| | | | |
|---|---|---|---|
| KPCD2_HUMAN\|Q9BZL6_S710 | 1.841965752 | 1.03E-03 | 1.57E-02 |
| SNUT1_HUMAN\|O43290_T430 | 1.841806887 | 5.57E-04 | 2.02E-02 |
| K1C14_HUMAN\|P02533_S44_S53_S54 | 1.84177276 | 1.48E-06 | 1.33E-04 |
| K1C18_HUMAN\|P05783_S7 | 1.841758194 | 1.07E-04 | 2.19E-03 |
| GRAP1_HUMAN\|Q4V328_S666 | 1.841404492 | 1.98E-05 | 7.80E-04 |
| MISP_HUMAN\|Q8IVT2_S395 | 1.841166164 | 9.79E-05 | 1.56E-03 |
| PKP3_HUMAN\|Q9Y446_S123 | 1.840937731 | 3.38E-05 | 1.12E-03 |
| TBD2A_HUMAN\|Q9BYX2_S920 | 1.840865648 | 1.28E-06 | 1.30E-04 |
| MAP7_HUMAN\|Q14244_S209 | 1.840711974 | 4.38E-05 | 6.85E-04 |
| CTNA1_HUMAN\|P35221_S652_S655_T658 | 1.840708862 | 1.41E-04 | 2.77E-03 |
| CCDC9_HUMAN\|Q9Y3X0_S386_S390 | 1.840538281 | 3.40E-04 | 4.18E-03 |
| CRBG1_HUMAN\|Q9Y4K1_S410 | 1.840246068 | 1.55E-04 | 3.30E-03 |
| COHA1_HUMAN\|Q9UMD9_S61_S62_S70 | 1.84017774 | 1.06E-03 | 8.34E-03 |
| SRRM1_HUMAN\|Q8IYB3_S738_S743 | 1.840086367 | 1.04E-03 | 8.21E-03 |
| HCFC1_HUMAN\|P51610_S1507 | 1.839958551 | 1.81E-04 | 4.82E-03 |
| NUFP2_HUMAN\|Q7Z417_S212_S214 | 1.839738786 | 6.87E-06 | 1.71E-04 |
| MK01_HUMAN\|P28482_Y187 | 1.839497828 | 2.33E-05 | 5.62E-04 |
| LAP2A_HUMAN\|P42166_S66_S67_T74 | 1.839427739 | 1.42E-05 | 7.78E-04 |
| RANG_HUMAN\|P43487_S60 | 1.839014487 | 4.05E-03 | 1.70E-02 |
| 4EBP1_HUMAN\|Q13541_S112 | 1.838453136 | 3.13E-03 | 2.95E-02 |
| K1C17_HUMAN\|Q04695_S11 | 1.838189395 | 5.03E-06 | 1.38E-04 |
| AHNK_HUMAN\|Q09666_T4430 | 1.837516881 | 1.10E-05 | 4.31E-04 |
| K2C6A_HUMAN\|P02538_Y62 | 1.837302541 | 4.27E-03 | 2.40E-02 |
| RD23B_HUMAN\|P54727_S160 | 1.837278109 | 1.17E-04 | 3.61E-03 |
| SQSTM_HUMAN\|Q13501_S355_S365 | 1.837183299 | 3.58E-03 | 2.43E-02 |
| MISP_HUMAN\|Q8IVT2_S400 | 1.837075333 | 6.78E-05 | 1.79E-03 |
| NAV1_HUMAN\|Q8NEY1_S648 | 1.836571445 | 1.85E-03 | 1.97E-02 |
| PLEC_HUMAN\|Q15149_S4673_S4675 | 1.836428131 | 2.56E-05 | 1.19E-03 |
| ARP19_HUMAN\|P56211_S62 | 1.836410561 | 1.61E-06 | 1.39E-04 |
| PPR18_HUMAN\|Q6NYC8_S209_S213 | 1.836347536 | 6.68E-07 | 7.39E-05 |
| K1522_HUMAN\|Q9P206_S339 | 1.836332807 | 2.84E-03 | 2.96E-02 |

| | | | |
|---|---|---|---|
| HELZ_HUMAN\|P42694_S1740_S1741 | 1.836197183 | 8.79E-03 | 4.83E-02 |
| MISP_HUMAN\|Q8IVT2_S395_S400 | 1.835678638 | 3.23E-04 | 4.03E-03 |
| K2C5_HUMAN\|P13647_S36 | 1.83506381 | 9.98E-06 | 4.04E-04 |
| IWS1_HUMAN\|Q96ST2_T53 | 1.835037539 | 6.88E-04 | 4.06E-03 |
| NOP2_HUMAN\|P46087_T605 | 1.834396769 | 2.83E-06 | 5.56E-05 |
| SF3B2_HUMAN\|Q13435_S303 | 1.83377482 | 6.74E-03 | 2.91E-02 |
| PHF14_HUMAN\|O94880_T287 | 1.833693527 | 8.09E-06 | 5.28E-04 |
| K2C6B_HUMAN\|P04259_S541 | 1.83362098 | 4.61E-06 | 2.32E-04 |
| PKP1_HUMAN\|Q13835_S142 | 1.833326517 | 9.34E-04 | 1.15E-02 |
| FKBP5_HUMAN\|Q13451_S445 | 1.83230421 | 2.55E-03 | 2.46E-02 |
| GOGA4_HUMAN\|Q13439_S89_S92 | 1.832065553 | 4.19E-06 | 3.39E-04 |
| GIT1_HUMAN\|Q9Y2X7_S388 | 1.832003277 | 4.12E-04 | 4.86E-03 |
| SOS1_HUMAN\|Q07889_S1082 | 1.831463905 | 3.58E-03 | 4.87E-02 |
| DESP_HUMAN\|P15924_S176 | 1.830941987 | 1.04E-03 | 8.21E-03 |
| K2C6A_HUMAN\|P02538_S58 | 1.830430976 | 3.61E-03 | 3.15E-02 |
| TM237_HUMAN\|Q96Q45_S62 | 1.830318914 | 2.08E-03 | 1.61E-02 |
| KPCD2_HUMAN\|Q9BZL6_S214 | 1.83018852 | 2.39E-05 | 5.71E-04 |
| ROA1_HUMAN\|P09651_S361_S364 | 1.830143215 | 9.74E-05 | 2.12E-02 |
| B4DKV4_HUMAN\|B4DKV4_S19 | 1.829995864 | 5.53E-04 | 8.35E-03 |
| PDLI2_HUMAN\|Q96JY6_S124_S127_S129 | 1.828744477 | 8.33E-05 | 1.49E-03 |
| MAP4_HUMAN\|P27816_S1073 | 1.828170254 | 2.35E-06 | 7.48E-05 |
| PLEC_HUMAN\|Q15149_S4675 | 1.827905708 | 1.21E-04 | 1.94E-03 |
| FAT1_HUMAN\|Q14517_S4473 | 1.82772394 | 2.38E-04 | 2.64E-03 |
| IWS1_HUMAN\|Q96ST2_S69_S80_T67 | 1.827572238 | 7.66E-04 | 4.94E-03 |
| LIMA1_HUMAN\|Q9UHB6_S507 | 1.827051553 | 7.45E-05 | 1.71E-03 |
| K2C6A_HUMAN\|P02538_S541 | 1.826573424 | 3.74E-03 | 2.17E-02 |
| FLNB_HUMAN\|O75369-8_S1474 | 1.826551873 | 2.04E-03 | 2.58E-02 |
| OXR1_HUMAN\|Q8N573_S201_S202 | 1.826209261 | 6.44E-07 | 3.60E-05 |
| PININ_HUMAN\|Q9H307_S66 | 1.825696099 | 2.31E-05 | 5.69E-04 |
| HNRPM_HUMAN\|P52272_S575 | 1.825645194 | 8.49E-05 | 1.85E-03 |
| SMC4_HUMAN\|Q9NTJ3_S22 | 1.823987353 | 2.48E-03 | 1.91E-02 |

| | | | |
|---|---|---|---|
| ZO2_HUMAN|Q9UDY2_S170 | 1.823875514 | 5.39E-07 | 1.64E-05 |
| RAI14_HUMAN|Q9P0K7-4_S261 | 1.823753112 | 5.63E-06 | 3.68E-04 |
| NFIB_HUMAN|O00712_S276_S295 | 1.823573363 | 3.46E-05 | 7.47E-04 |
| IRS2_HUMAN|Q9Y4H2_S1185_S1203 | 1.82335459 | 1.64E-05 | 8.45E-04 |
| ACAP2_HUMAN|Q15057_S381_S384 | 1.823340245 | 4.45E-06 | 1.26E-04 |
| MRCKG_HUMAN|Q6DT37_S1544 | 1.823064515 | 1.08E-04 | 1.80E-03 |
| PHLB2_HUMAN|Q86SQ0_S330_S334 | 1.822827542 | 2.07E-04 | 3.76E-03 |
| JUND_HUMAN|P17535_S100 | 1.822147254 | 1.91E-05 | 2.42E-04 |
| I2BP2_HUMAN|Q7Z5L9_S460 | 1.82214367 | 4.41E-10 | 1.14E-07 |
| K1C14_HUMAN|P02533_S435 | 1.821869217 | 3.81E-05 | 1.06E-03 |
| LIMA1_HUMAN|Q9UHB6_S617 | 1.821775116 | 8.33E-05 | 1.84E-03 |
| MACF1_HUMAN|Q9UPN3_T7287 | 1.821574978 | 4.09E-04 | 4.85E-03 |
| SRRM2_HUMAN|Q9UQ35_S1152 | 1.8209734 | 8.54E-05 | 1.15E-03 |
| ARAF_HUMAN|P10398_S257_T253 | 1.820387607 | 1.39E-05 | 2.92E-04 |
| TRAD1_HUMAN|O14545_S409_S415 | 1.820210699 | 2.04E-04 | 2.83E-03 |
| NOLC1_HUMAN|Q14978_S688_S698 | 1.819962795 | 8.78E-03 | 4.71E-02 |
| XPC_HUMAN|Q01831_S883_S884_S891 | 1.819875202 | 9.94E-05 | 2.40E-03 |
| AHNK2_HUMAN|Q8IVF2_S294_T298 | 1.819836735 | 1.46E-05 | 6.29E-04 |
| ARH40_HUMAN|Q8TER5_S262 | 1.81917916 | 6.73E-07 | 3.15E-05 |
| OSB11_HUMAN|Q9BXB4_S177_S181 | 1.81915272 | 1.74E-03 | 3.10E-02 |
| FUBP2_HUMAN|Q92945_T195 | 1.818933419 | 1.50E-03 | 2.04E-02 |
| ZBT21_HUMAN|Q9ULJ3_S422 | 1.818573459 | 6.71E-04 | 3.99E-03 |
| K1671_HUMAN|Q9BY89_S1441 | 1.818499577 | 7.56E-05 | 1.71E-03 |
| OSBL8_HUMAN|Q9BZF1_S807_S808_S810 | 1.818433878 | 4.51E-05 | 4.84E-04 |
| NUP98_HUMAN|P52948_T1000 | 1.818402764 | 6.89E-05 | 1.22E-03 |
| LPP_HUMAN|Q93052_S609 | 1.817840164 | 9.53E-05 | 7.17E-03 |
| INT12_HUMAN|Q96CB8_S389 | 1.817559413 | 2.13E-04 | 2.95E-03 |
| COBL1_HUMAN|Q53SF7_S364 | 1.817408093 | 2.08E-06 | 9.96E-05 |
| CING_HUMAN|Q9P2M7_S129_S134 | 1.817338198 | 2.03E-05 | 9.65E-04 |
| RL23_HUMAN|P62829_S138 | 1.816612805 | 4.53E-03 | 2.10E-02 |
| PHRF1_HUMAN|Q9P1Y6_S915_T917 | 1.81658368 | 1.14E-03 | 8.79E-03 |

| | | | |
|---|---|---|---|
| AHNK_HUMAN\|Q09666_S5739 | 1.816581311 | 5.80E-05 | 1.62E-03 |
| DESP_HUMAN\|P15924_T13 | 1.81632338 | 6.07E-03 | 3.08E-02 |
| GA2L1_HUMAN\|Q99501_S352 | 1.815866884 | 4.08E-04 | 7.90E-03 |
| TBC15_HUMAN\|Q8TC07-2_S205 | 1.815771947 | 5.93E-05 | 8.77E-04 |
| COBL1_HUMAN\|Q53SF7_S294_T298 | 1.815532264 | 1.21E-05 | 1.63E-03 |
| K2C5_HUMAN\|P13647_S50 | 1.81542727 | 7.26E-07 | 7.87E-05 |
| MISP_HUMAN\|Q8IVT2_S394_S395 | 1.815410384 | 1.75E-06 | 3.75E-05 |
| PLEC_HUMAN\|Q15149_S4622_Y4612 | 1.814863671 | 1.20E-05 | 2.68E-04 |
| GIT1_HUMAN\|Q9Y2X7_S388_T392 | 1.814495152 | 1.04E-04 | 9.96E-04 |
| K1C14_HUMAN\|P02533_S44 | 1.813769974 | 6.24E-05 | 2.15E-03 |
| ZO2_HUMAN\|Q9UDY2_S394 | 1.813720335 | 5.34E-03 | 2.09E-02 |
| K2C6A_HUMAN\|P02538_S546 | 1.813090062 | 6.35E-03 | 3.19E-02 |
| PLEC_HUMAN\|Q15149_S4389_S4391 | 1.81274999 | 4.37E-05 | 4.76E-04 |
| K1C14_HUMAN\|P02533_S51_S54 | 1.812497335 | 6.24E-05 | 9.07E-04 |
| TIF1B_HUMAN\|Q13263_S596_S598_S601 | 1.812359526 | 4.07E-03 | 3.43E-02 |
| WAC2A_HUMAN\|Q641Q2_S284 | 1.81218218 | 8.91E-05 | 2.91E-03 |
| TCAL3_HUMAN\|Q969E4_S125 | 1.81197172 | 7.30E-05 | 2.43E-03 |
| TV23B_HUMAN\|Q9NYZ1_T199 | 1.811878146 | 3.09E-03 | 3.30E-02 |
| AHNK_HUMAN\|Q09666_S5864 | 1.811659466 | 1.06E-03 | 2.92E-02 |
| TMF1_HUMAN\|P82094_S338_S344 | 1.811185766 | 3.23E-07 | 4.16E-05 |
| ICE1_HUMAN\|Q9Y2F5_S1712 | 1.811184661 | 5.93E-06 | 1.53E-04 |
| HEXI1_HUMAN\|O94992_S237_S252 | 1.810773404 | 5.35E-04 | 3.65E-03 |
| DC1L1_HUMAN\|Q9Y6G9_S516_T515 | 1.810287026 | 2.94E-06 | 2.34E-04 |
| FOXK2_HUMAN\|Q01167_S398 | 1.810139931 | 2.28E-04 | 1.83E-03 |
| RENT1_HUMAN\|Q92900_S1127 | 1.810077868 | 5.41E-03 | 4.17E-02 |
| NBEL2_HUMAN\|Q6ZNJ1_S2739 | 1.810068714 | 1.94E-03 | 1.05E-02 |
| SPTB2_HUMAN\|Q01082-3_S14_S8 | 1.809890729 | 2.07E-06 | 1.35E-04 |
| DTX3L_HUMAN\|Q8TDB6_S202 | 1.809821708 | 1.12E-03 | 1.71E-02 |
| NIBL1_HUMAN\|Q96TA1_S691_S696 | 1.809706035 | 4.68E-06 | 8.19E-05 |
| ACAP2_HUMAN\|Q15057_S381_S384 | 1.809463315 | 5.29E-06 | 2.00E-04 |
| PARD3_HUMAN\|Q8TEW0_S225 | 1.808464722 | 8.57E-06 | 3.60E-04 |

| | | | |
|---|---|---|---|
| NELFE_HUMAN\|P18615_S51 | 1.808443034 | 3.12E-06 | 6.15E-05 |
| MILK1_HUMAN\|Q8N3F8_T318 | 1.808247961 | 5.14E-04 | 7.26E-03 |
| K2C6A_HUMAN\|P02538_S85 | 1.808118324 | 6.39E-03 | 4.82E-02 |
| MED24_HUMAN\|O75448_S862 | 1.807749559 | 2.18E-03 | 1.03E-02 |
| AMOL2_HUMAN\|Q9Y2J4_S759_S762 | 1.807218987 | 9.97E-04 | 1.45E-02 |
| PHLB1_HUMAN\|Q86UU1_S430 | 1.806868071 | 1.46E-05 | 4.01E-04 |
| AHNK_HUMAN\|Q09666_T158 | 1.806476058 | 8.84E-04 | 7.21E-03 |
| DOCK9_HUMAN\|Q9BZ29-5_S37 | 1.806110716 | 5.35E-05 | 1.99E-03 |
| CE170_HUMAN\|Q5SW79_S630_S633 | 1.805708252 | 8.32E-07 | 8.84E-05 |
| K2C8_HUMAN\|P05787_S21_S24_S27 | 1.805503033 | 1.67E-05 | 5.42E-04 |
| K2C5_HUMAN\|P13647_T19 | 1.805484702 | 4.32E-04 | 6.32E-03 |
| EIF3F_HUMAN\|O00303_S258 | 1.805445251 | 1.07E-03 | 2.92E-02 |
| ZO2_HUMAN\|Q9UDY2_T1027 | 1.804570109 | 2.33E-06 | 1.42E-04 |
| TANC2_HUMAN\|Q9HCD6_S1827 | 1.803728995 | 6.20E-07 | 4.10E-05 |
| FOXO3_HUMAN\|O43524_S413 | 1.803310405 | 6.97E-04 | 1.18E-02 |
| KPCD2_HUMAN\|Q9BZL6_S710 | 1.803307421 | 1.40E-03 | 1.26E-02 |
| K2C6B_HUMAN\|P04259_S37 | 1.80318239 | 2.08E-04 | 1.06E-02 |
| TB10A_HUMAN\|Q9BXI6_S40_S43 | 1.803155444 | 3.93E-04 | 2.62E-03 |
| GGYF2_HUMAN\|Q6Y7W6_S30 | 1.802679614 | 1.51E-03 | 1.65E-02 |
| CLAP1_HUMAN\|Q7Z460_S1091 | 1.802264888 | 8.31E-07 | 3.65E-05 |
| HMGA2_HUMAN\|P52926_S105 | 1.801767359 | 1.73E-03 | 2.17E-02 |
| UTP18_HUMAN\|Q9Y5J1_S205 | 1.801322223 | 1.45E-03 | 8.19E-03 |
| K2C6A_HUMAN\|P02538_S22 | 1.801071277 | 4.30E-03 | 3.64E-02 |
| CRBG1_HUMAN\|Q9Y4K1_S22 | 1.800876866 | 3.40E-07 | 1.93E-05 |
| ITB4_HUMAN\|P16144-2_S1364_T1369 | 1.800867112 | 5.53E-03 | 2.14E-02 |
| KC1A_HUMAN\|P48729_S312 | 1.800729618 | 2.91E-03 | 2.07E-02 |
| SRPK1_HUMAN\|Q96SB4_S39_S51 | 1.800662229 | 9.99E-05 | 1.71E-03 |
| FLVC1_HUMAN\|Q9Y5Y0_S536 | 1.800459749 | 3.51E-05 | 1.53E-03 |
| PDLI2_HUMAN\|Q96JY6_S124_S127_S129 | 1.800081375 | 1.13E-04 | 1.06E-03 |
| K2C6A_HUMAN\|P02538_S22 | 1.799719989 | 4.34E-03 | 2.42E-02 |
| ARHG5_HUMAN\|Q12774_S445_S450 | 1.799605358 | 9.47E-08 | 2.12E-05 |

| | | | |
|---|---|---|---|
| AAK1_HUMAN\|Q2M2I8_S678 | 1.799470301 | 7.11E-06 | 4.87E-04 |
| ARG28_HUMAN\|Q8N1W1_S476_S486 | 1.799223726 | 1.77E-05 | 3.50E-04 |
| RERE_HUMAN\|Q9P2R6_S679 | 1.798886098 | 4.12E-04 | 4.86E-03 |
| TM1L1_HUMAN\|O75674_S321 | 1.798836089 | 1.85E-04 | 1.45E-03 |
| DESP_HUMAN\|P15924_S166 | 1.798727447 | 4.58E-05 | 1.95E-03 |
| MTAP2_HUMAN\|P11137_S1782 | 1.798684822 | 8.46E-04 | 4.81E-03 |
| FOXO3_HUMAN\|O43524_S413 | 1.798608938 | 7.27E-04 | 1.24E-02 |
| 2A5E_HUMAN\|Q16537_S32_S34 | 1.798484976 | 3.08E-04 | 3.24E-03 |
| PAK2_HUMAN\|Q13177_S197 | 1.798477203 | 3.62E-03 | 3.26E-02 |
| K1C13_HUMAN\|P13646_S11 | 1.798428442 | 4.12E-03 | 3.46E-02 |
| DESP_HUMAN\|P15924_S53 | 1.7983423 | 1.61E-03 | 1.16E-02 |
| EF2_HUMAN\|P13639_T57 | 1.797910928 | 2.52E-03 | 1.30E-02 |
| SRPK2_HUMAN\|P78362_S494_T498 | 1.797706021 | 3.53E-04 | 4.28E-03 |
| RTKN_HUMAN\|Q9BST9_S529 | 1.797608811 | 1.67E-05 | 3.35E-04 |
| CDK9_HUMAN\|P50750_S347 | 1.797527576 | 1.58E-03 | 1.31E-02 |
| DKC1_HUMAN\|O60832_S485_S494 | 1.797238917 | 3.43E-07 | 4.16E-05 |
| AKAP2_HUMAN\|Q9Y2D5_S152_S155 | 1.796744019 | 2.68E-06 | 1.54E-04 |
| MAP4_HUMAN\|P27816_S941 | 1.796398809 | 1.28E-05 | 2.78E-04 |
| PRC2A_HUMAN\|P48634_S761_S764 | 1.796228298 | 1.86E-07 | 3.29E-05 |
| AKAP8_HUMAN\|O43823_S323 | 1.795740242 | 4.01E-04 | 2.67E-03 |
| PRC2C_HUMAN\|Q9Y520_S2694_T2691 | 1.795499501 | 8.60E-05 | 1.15E-03 |
| MRP_HUMAN\|P49006_T178 | 1.795401823 | 5.06E-07 | 6.23E-05 |
| RB3GP_HUMAN\|Q15042_S537 | 1.795002057 | 5.12E-07 | 6.60E-05 |
| TPPP_HUMAN\|O94811_S160 | 1.794525949 | 1.54E-04 | 1.86E-03 |
| NSD3_HUMAN\|Q9BZ95_S457 | 1.794109649 | 2.04E-06 | 4.26E-05 |
| NAV1_HUMAN\|Q8NEY1_T1006 | 1.793812496 | 1.81E-04 | 2.11E-03 |
| TIF1B_HUMAN\|Q13263_S594_S601 | 1.793728305 | 2.55E-03 | 1.64E-02 |
| LMNB2_HUMAN\|Q03252_S420 | 1.793577439 | 2.54E-06 | 1.91E-04 |
| K1C17_HUMAN\|Q04695_S13 | 1.793541513 | 7.39E-06 | 3.26E-04 |
| 2A5E_HUMAN\|Q16537_S32_S34 | 1.793371999 | 3.24E-04 | 5.11E-03 |
| EIF3A_HUMAN\|Q14152_S882 | 1.792390265 | 3.59E-04 | 6.04E-03 |

| | | | |
|---|---|---|---|
| RPB1_HUMAN\|P24928_S1847_T1856 | 1.791924899 | 1.58E-04 | 1.28E-03 |
| SLK_HUMAN\|Q9H2G2_S779 | 1.791898943 | 8.92E-04 | 1.18E-02 |
| LAP2A_HUMAN\|P42166_S167_T160 | 1.791790611 | 1.26E-05 | 4.51E-04 |
| MA7D1_HUMAN\|Q3KQU3_S366 | 1.791460448 | 6.57E-08 | 5.75E-06 |
| DC1L1_HUMAN\|Q9Y6G9_S516_T515 | 1.790687323 | 3.82E-06 | 6.97E-05 |
| TMF1_HUMAN\|P82094_S77 | 1.790630844 | 4.34E-06 | 1.61E-04 |
| ETV3_HUMAN\|P41162_S139 | 1.790300242 | 1.73E-03 | 1.87E-02 |
| ACSA_HUMAN\|Q9NR19_S267 | 1.790182638 | 8.43E-06 | 2.70E-04 |
| NIBL1_HUMAN\|Q96TA1_S692_S696 | 1.790023195 | 1.40E-05 | 8.91E-04 |
| PLEC_HUMAN\|Q15149_S4673_S4675 | 1.789830677 | 4.46E-05 | 9.13E-04 |
| OPTN_HUMAN\|Q96CV9_T204 | 1.789682079 | 2.22E-07 | 3.54E-05 |
| COBL1_HUMAN\|Q53SF7_S1146 | 1.789669686 | 6.66E-06 | 2.29E-04 |
| K1C16_HUMAN\|P08779_S449_S452 | 1.789440514 | 8.61E-04 | 1.31E-02 |
| PDLI2_HUMAN\|Q96JY6_S137 | 1.789008221 | 1.86E-06 | 3.95E-05 |
| SGTA_HUMAN\|O43765_S301_S305 | 1.788598652 | 6.65E-04 | 9.52E-03 |
| AHNK2_HUMAN\|Q8IVF2_S294_T298 | 1.788589169 | 2.19E-05 | 6.94E-04 |
| RAI14_HUMAN\|Q9P0K7_T297 | 1.788515283 | 1.01E-04 | 7.30E-03 |
| IWS1_HUMAN\|Q96ST2_T725 | 1.788048271 | 1.19E-04 | 7.06E-03 |
| KLC3_HUMAN\|Q6P597_S167_S173 | 1.788020522 | 3.69E-05 | 1.16E-03 |
| K2C75_HUMAN\|O95678_S40 | 1.787470805 | 1.02E-03 | 1.30E-02 |
| MPZL1_HUMAN\|O95297_S221 | 1.78646841 | 3.69E-03 | 2.49E-02 |
| FOXO3_HUMAN\|O43524_S413 | 1.786003133 | 8.14E-04 | 1.27E-02 |
| TACC1_HUMAN\|O75410_S44 | 1.785514724 | 7.58E-06 | 4.03E-04 |
| LAD1_HUMAN\|O00515_S38_T40_T41 | 1.785507814 | 5.35E-05 | 1.99E-03 |
| K1C19_HUMAN\|P08727_S13 | 1.784663043 | 2.28E-04 | 4.39E-03 |
| CPSF4_HUMAN\|O95639_S202 | 1.784438251 | 7.79E-05 | 7.74E-04 |
| COF1_HUMAN\|P23528_S3 | 1.783830816 | 1.04E-02 | 4.68E-02 |
| WFS1_HUMAN\|O76024_S235 | 1.783624598 | 1.22E-03 | 1.09E-02 |
| BCL10_HUMAN\|O95999_S138 | 1.783620417 | 1.52E-05 | 4.23E-04 |
| SRRM1_HUMAN\|Q8IYB3_S738_S743 | 1.783145653 | 1.66E-03 | 1.79E-02 |
| B4DET1_HUMAN\|B4DET1_S720_T711 | 1.782931176 | 1.24E-02 | 4.08E-02 |

| | | | |
|---|---|---|---|
| TRI47_HUMAN\|Q96LD4_S588 | 1.782696336 | 2.69E-05 | 1.32E-03 |
| MARCS_HUMAN\|P29966_S118 | 1.782466418 | 2.96E-04 | 2.22E-03 |
| MAP1B_HUMAN\|P46821_S1427 | 1.782401687 | 3.40E-07 | 1.93E-05 |
| CCDC9_HUMAN\|Q9Y3X0_S386_T381 | 1.782346921 | 7.71E-06 | 4.06E-04 |
| TBCD5_HUMAN\|Q92609_S541 | 1.782272276 | 6.18E-09 | 3.14E-06 |
| K1522_HUMAN\|Q9P206_S929 | 1.782012068 | 4.62E-07 | 1.35E-05 |
| BNIP2_HUMAN\|Q12982_S114 | 1.781835414 | 1.20E-04 | 1.05E-03 |
| UBXN7_HUMAN\|O94888_S285 | 1.781354846 | 3.68E-04 | 5.66E-03 |
| MRCKG_HUMAN\|Q6DT37_S1544 | 1.780814361 | 1.70E-04 | 4.61E-03 |
| IRS1_HUMAN\|P35568_S330 | 1.780396352 | 2.73E-04 | 3.58E-03 |
| GGYF2_HUMAN\|Q6Y7W6_S30 | 1.78008099 | 1.80E-03 | 1.43E-02 |
| NUFP2_HUMAN\|Q7Z417_S629 | 1.780062172 | 4.66E-04 | 9.15E-03 |
| IASPP_HUMAN\|Q8WUF5_S134 | 1.779894392 | 4.90E-05 | 7.50E-04 |
| CAMP2_HUMAN\|Q08AD1_S691 | 1.779806783 | 1.30E-04 | 3.50E-03 |
| NBEL2_HUMAN\|Q6ZNJ1_S2739 | 1.7797023 | 2.47E-03 | 2.80E-02 |
| CE170_HUMAN\|Q5SW79_S636_T628 | 1.77968382 | 4.52E-05 | 1.35E-03 |
| EF2K_HUMAN\|O00418_S470 | 1.779657154 | 8.71E-06 | 1.33E-04 |
| LMNA_HUMAN\|P02545_S428 | 1.779247784 | 3.57E-07 | 1.93E-05 |
| COBL1_HUMAN\|Q53SF7_T298 | 1.778812234 | 2.54E-05 | 4.67E-04 |
| PDLI2_HUMAN\|Q96JY6_S134 | 1.778576404 | 1.94E-08 | 2.62E-06 |
| DC1L1_HUMAN\|Q9Y6G9_S516 | 1.778525478 | 1.85E-06 | 1.80E-04 |
| MICA3_HUMAN\|Q7RTP6_S1636 | 1.778404458 | 4.61E-04 | 7.23E-03 |
| PFKAL_HUMAN\|P17858_S775 | 1.778387689 | 3.00E-04 | 3.83E-03 |
| PFKAL_HUMAN\|P17858_S775 | 1.778182844 | 3.00E-04 | 6.63E-03 |
| AHNK_HUMAN\|Q09666_S5752 | 1.77801321 | 1.61E-07 | 3.13E-05 |
| FXR2_HUMAN\|P51116_S453 | 1.777790339 | 8.32E-06 | 3.52E-04 |
| PKP3_HUMAN\|Q9Y446_S314 | 1.777501813 | 1.74E-06 | 1.24E-04 |
| SSFA2_HUMAN\|P28290_S92 | 1.776742618 | 2.48E-03 | 1.61E-02 |
| NIBL1_HUMAN\|Q96TA1_S641_S646 | 1.77631787 | 2.58E-04 | 5.96E-03 |
| CLIP4_HUMAN\|Q8N3C7_S610 | 1.775683919 | 3.00E-05 | 6.71E-04 |
| LAP2A_HUMAN\|P42166_S167_T160 | 1.775300942 | 1.55E-05 | 6.57E-04 |

| | | | |
|---|---|---|---|
| EIF3H_HUMAN\|O15372_S183 | 1.775282055 | 3.44E-04 | 1.47E-02 |
| ZO2_HUMAN\|Q9UDY2_S170 | 1.774976922 | 1.13E-06 | 1.28E-04 |
| FOSL2_HUMAN\|P15408_S320 | 1.774904188 | 2.57E-03 | 4.90E-02 |
| CAVN3_HUMAN\|Q969G5_T205 | 1.77487197 | 7.04E-05 | 1.66E-03 |
| LAP2A_HUMAN\|P42166_T74 | 1.774435996 | 1.35E-04 | 1.67E-03 |
| 3BP1_HUMAN\|Q9Y3L3_S596 | 1.774161678 | 8.88E-06 | 2.07E-04 |
| AHNK2_HUMAN\|Q8IVF2_S294_T298 | 1.773881782 | 2.66E-05 | 4.77E-04 |
| PKP4_HUMAN\|Q99569_S447 | 1.773810412 | 6.26E-04 | 9.16E-03 |
| SRRM2_HUMAN\|Q9UQ35_S1463 | 1.773797978 | 1.39E-04 | 1.25E-03 |
| ACSA_HUMAN\|Q9NR19_S267 | 1.773763081 | 1.06E-05 | 1.51E-03 |
| TB182_HUMAN\|Q9C0C2_T1282 | 1.773558681 | 4.46E-06 | 2.28E-04 |
| NELFE_HUMAN\|P18615_S251 | 1.773341791 | 1.49E-04 | 2.20E-03 |
| AHNK_HUMAN\|Q09666_S210 | 1.77309103 | 1.94E-07 | 1.44E-05 |
| NIBAN_HUMAN\|Q9BZQ8_S926 | 1.773005545 | 2.12E-06 | 1.35E-04 |
| PAXI_HUMAN\|P49023_S106 | 1.772903935 | 1.84E-05 | 9.08E-04 |
| K1C14_HUMAN\|P02533_S39 | 1.772751189 | 1.88E-03 | 1.99E-02 |
| K2C5_HUMAN\|P13647_S21 | 1.772458981 | 8.30E-03 | 3.94E-02 |
| RABE1_HUMAN\|Q15276_S407 | 1.772077129 | 2.53E-06 | 5.31E-05 |
| ZO1_HUMAN\|Q07157_T868_Y843 | 1.772067995 | 8.10E-04 | 1.33E-02 |
| K1671_HUMAN\|Q9BY89_S1441 | 1.772064475 | 1.25E-04 | 8.46E-03 |
| K1C14_HUMAN\|P02533_S433_S438 | 1.771859615 | 8.23E-05 | 1.12E-03 |
| K1C14_HUMAN\|P02533_S13 | 1.771780808 | 1.06E-06 | 4.27E-05 |
| MK14_HUMAN\|Q16539_Y182 | 1.771451992 | 2.38E-05 | 5.70E-04 |
| KPCD2_HUMAN\|Q9BZL6_S710 | 1.771338756 | 1.81E-03 | 1.92E-02 |
| DC1I2_HUMAN\|Q13409-2_S81 | 1.771162475 | 4.73E-04 | 7.35E-03 |
| JUN_HUMAN\|P05412_S63 | 1.771095475 | 6.43E-05 | 2.49E-03 |
| SEPT7_HUMAN\|Q16181_T426 | 1.770994167 | 5.97E-04 | 8.90E-03 |
| SRPK2_HUMAN\|P78362_S494_S497 | 1.770774549 | 5.89E-06 | 3.94E-04 |
| AKAP2_HUMAN\|Q9Y2D5_S152_T157 | 1.770350395 | 4.54E-06 | 3.52E-04 |
| AHNK2_HUMAN\|Q8IVF2_S457 | 1.770190111 | 9.63E-04 | 9.19E-03 |
| CR025_HUMAN\|Q96B23_S69 | 1.769934617 | 1.77E-03 | 1.49E-02 |

| | | | |
|---|---|---|---|
| B4DET1_HUMAN\|B4DET1_S725_Y731 | 1.769659252 | 1.36E-02 | 4.41E-02 |
| PHC2_HUMAN\|Q8IXK0_S751 | 1.769653759 | 9.46E-04 | 9.07E-03 |
| NAV1_HUMAN\|Q8NEY1_S648 | 1.769512009 | 3.08E-03 | 2.23E-02 |
| COMT_HUMAN\|P21964_S267 | 1.769203271 | 3.01E-06 | 1.65E-04 |
| SMRCD_HUMAN\|Q9H4L7_S96 | 1.76889196 | 5.87E-06 | 9.73E-04 |
| GRSF1_HUMAN\|Q12849_S118 | 1.768693035 | 5.95E-06 | 2.03E-04 |
| DP13A_HUMAN\|Q9UKG1_S401 | 1.768419376 | 3.96E-05 | 6.44E-04 |
| ZO1_HUMAN\|Q07157_S841_Y853 | 1.768090054 | 2.37E-04 | 3.20E-03 |
| ARG28_HUMAN\|Q8N1W1_S513 | 1.767658443 | 8.17E-06 | 1.96E-04 |
| ZCRB1_HUMAN\|Q8TBF4_S216 | 1.767269447 | 1.31E-06 | 3.04E-05 |
| EGFR_HUMAN\|P00533_S991 | 1.765798996 | 7.01E-06 | 1.13E-04 |
| ZO1_HUMAN\|Q07157_T868_Y843 | 1.765161156 | 8.63E-04 | 8.57E-03 |
| K2C6B_HUMAN\|P04259_S541 | 1.765100325 | 1.26E-05 | 7.04E-04 |
| L7UQD0_HUMAN\|L7UQD0_S268 | 1.764679792 | 3.15E-05 | 1.08E-03 |
| MAP7_HUMAN\|Q14244_S315 | 1.764413601 | 3.98E-06 | 1.15E-04 |
| PAK1_HUMAN\|Q13153_S223 | 1.764397405 | 6.05E-05 | 6.30E-04 |
| PAXI_HUMAN\|P49023_S250 | 1.76401091 | 7.50E-04 | 1.05E-02 |
| ZO1_HUMAN\|Q07157_S1617 | 1.764000653 | 2.56E-06 | 2.21E-04 |
| E7ETA6_HUMAN\|E7ETA6_T1557 | 1.763881547 | 8.62E-04 | 2.67E-02 |
| UBP32_HUMAN\|Q8NFA0_S1393 | 1.763538382 | 1.56E-06 | 3.69E-05 |
| DESP_HUMAN\|P15924_S166 | 1.763408892 | 7.05E-05 | 1.25E-03 |
| RERE_HUMAN\|Q9P2R6_S679 | 1.763180734 | 5.81E-04 | 6.28E-03 |
| MAP7_HUMAN\|Q14244_S202 | 1.763170678 | 9.01E-07 | 9.39E-05 |
| NBN_HUMAN\|O60934_S343 | 1.763156952 | 1.08E-05 | 4.29E-04 |
| K1C16_HUMAN\|P08779_S446_S449 | 1.762813694 | 2.87E-05 | 8.46E-04 |
| DESP_HUMAN\|P15924_S166 | 1.762798942 | 7.10E-05 | 7.28E-04 |
| COBL1_HUMAN\|Q53SF7_S1146 | 1.762612611 | 9.59E-06 | 5.65E-04 |
| K1C19_HUMAN\|P08727_S22 | 1.762542675 | 4.09E-05 | 1.12E-03 |
| UBP5_HUMAN\|P45974_S779 | 1.762487942 | 1.48E-04 | 2.66E-02 |
| K1C14_HUMAN\|P02533_S44_S53_S54 | 1.762021247 | 5.10E-06 | 1.38E-04 |
| PDLI2_HUMAN\|Q96JY6_S134_S137_S143 | 1.761408154 | 4.53E-05 | 1.77E-03 |

| | | | |
|---|---|---|---|
| MFAP1_HUMAN\|P55081_S116 | 1.761398279 | 1.13E-03 | 6.06E-03 |
| PSF2_HUMAN\|Q9Y248_S182_T180 | 1.761242667 | 1.68E-04 | 3.46E-03 |
| CTNB1_HUMAN\|P35222_T551 | 1.761115333 | 6.40E-04 | 9.28E-03 |
| K2C5_HUMAN\|P13647_S571_S574 | 1.76067943 | 2.51E-05 | 7.69E-04 |
| PKP3_HUMAN\|Q9Y446_S123 | 1.76057283 | 8.34E-05 | 1.84E-03 |
| DC1L1_HUMAN\|Q9Y6G9_S510_S516 | 1.760517356 | 5.31E-06 | 3.81E-04 |
| SGTA_HUMAN\|O43765_S305 | 1.7600753 | 7.50E-06 | 5.81E-04 |
| MK14_HUMAN\|Q16539_Y182 | 1.759475352 | 2.78E-05 | 3.50E-03 |
| RRP1B_HUMAN\|Q14684_S245 | 1.759415588 | 4.10E-07 | 3.08E-05 |
| PININ_HUMAN\|Q9H307_S690 | 1.759249231 | 1.78E-04 | 1.41E-03 |
| MED24_HUMAN\|O75448_S862_S873 | 1.759007193 | 9.80E-04 | 1.52E-02 |
| BRAF_HUMAN\|P15056_S446 | 1.7585368 | 9.02E-05 | 8.38E-04 |
| K2C5_HUMAN\|P13647_S77_S82 | 1.758320629 | 1.36E-03 | 1.54E-02 |
| K1C14_HUMAN\|P02533_S433_S438 | 1.757709623 | 9.84E-05 | 2.05E-03 |
| SF3B2_HUMAN\|Q13435_S307 | 1.757602963 | 3.77E-04 | 7.41E-03 |
| MED24_HUMAN\|O75448_S862_S873 | 1.757556399 | 9.94E-04 | 1.56E-02 |
| SLK_HUMAN\|Q9H2G2_S655 | 1.757074074 | 1.44E-04 | 2.19E-03 |
| SCAM2_HUMAN\|O15127_S320 | 1.756786872 | 2.42E-04 | 1.92E-03 |
| K2C5_HUMAN\|P13647_S82_T78 | 1.756616726 | 6.67E-06 | 3.04E-04 |
| PRAG1_HUMAN\|Q86YV5_S745 | 1.756302666 | 2.73E-05 | 1.23E-03 |
| VIP2_HUMAN\|O43314_S1006 | 1.756152027 | 7.89E-07 | 1.94E-04 |
| NEB2_HUMAN\|Q96SB3_S100 | 1.756072184 | 3.68E-04 | 1.55E-02 |
| SPF30_HUMAN\|O75940_S201 | 1.755860634 | 1.11E-04 | 2.24E-02 |
| TAGL2_HUMAN\|P37802_S163 | 1.755819227 | 2.55E-04 | 6.01E-03 |
| K1C14_HUMAN\|P02533_Y46 | 1.755488343 | 4.18E-05 | 3.70E-03 |
| MA7D1_HUMAN\|Q3KQU3_S452 | 1.755205348 | 4.82E-05 | 1.24E-03 |
| MAP7_HUMAN\|Q14244_T184 | 1.755102942 | 4.61E-06 | 8.34E-04 |
| DEN1A_HUMAN\|Q8TEH3_T532 | 1.754452503 | 2.29E-03 | 1.80E-02 |
| BCL10_HUMAN\|O95999_S138 | 1.754230362 | 2.25E-05 | 2.79E-04 |
| TV23B_HUMAN\|Q9NYZ1_T199 | 1.753847296 | 4.78E-03 | 4.55E-02 |
| REQU_HUMAN\|Q92785_S142 | 1.753800308 | 2.33E-05 | 7.25E-04 |

| | | | |
|---|---|---|---|
| PAK2_HUMAN\|Q13177_S192 | 1.753776375 | 6.32E-03 | 4.77E-02 |
| MILK1_HUMAN\|Q8N3F8_S640_S644 | 1.753735471 | 1.22E-05 | 6.90E-04 |
| ENSA_HUMAN\|O43768_S109 | 1.753652769 | 4.00E-05 | 1.52E-03 |
| KCD12_HUMAN\|Q96CX2_S200_T196 | 1.753575264 | 9.99E-04 | 9.43E-03 |
| AHNK_HUMAN\|Q09666_S3426 | 1.753383475 | 3.95E-06 | 1.50E-04 |
| IRS2_HUMAN\|Q9Y4H2_S1186_S1203 | 1.753279621 | 9.09E-06 | 5.42E-04 |
| F120A_HUMAN\|Q9NZB2_S1023 | 1.752895005 | 4.80E-06 | 1.34E-04 |
| EGFR_HUMAN\|P00533_S991 | 1.75284917 | 8.58E-06 | 5.23E-04 |
| K2C6A_HUMAN\|P02538_S22 | 1.752112114 | 6.08E-03 | 4.58E-02 |
| TMF1_HUMAN\|P82094_S338_S344 | 1.752021134 | 8.94E-07 | 9.69E-05 |
| PTN3_HUMAN\|P26045_S425 | 1.751850176 | 5.56E-05 | 8.46E-04 |
| AHNK_HUMAN\|Q09666_S5746_S5752 | 1.751651322 | 1.63E-06 | 1.51E-04 |
| CY561_HUMAN\|P49447_S250 | 1.751471016 | 1.34E-04 | 4.01E-03 |
| K1C17_HUMAN\|Q04695_S11_S13 | 1.751269254 | 4.61E-05 | 1.82E-03 |
| UBP5_HUMAN\|P45974_S779 | 1.75114984 | 1.67E-04 | 2.45E-03 |
| OSB10_HUMAN\|Q9BXB5_S32 | 1.750288504 | 1.22E-05 | 2.70E-04 |
| TLN1_HUMAN\|Q9Y490_S1227 | 1.750042972 | 3.45E-04 | 7.39E-03 |
| SRPK2_HUMAN\|P78362_S494_S497 | 1.750009488 | 7.85E-06 | 1.24E-04 |
| TM230_HUMAN\|Q96A57_T25 | 1.749244421 | 3.36E-04 | 1.47E-02 |
| M3K3_HUMAN\|Q99759_S337 | 1.748649036 | 8.20E-05 | 1.39E-03 |
| LMNA_HUMAN\|P02545_S414 | 1.748388288 | 1.75E-05 | 3.47E-04 |
| AAK1_HUMAN\|Q2M2I8_S678 | 1.747880897 | 1.39E-05 | 8.53E-04 |
| COHA1_HUMAN\|Q9UMD9_S172_T176_T182 | 1.747829731 | 3.65E-03 | 3.27E-02 |
| JMY_HUMAN\|Q8N9B5_S115 | 1.74762552 | 1.95E-04 | 2.26E-03 |
| HMGA2_HUMAN\|P52926_S105 | 1.747561425 | 2.71E-03 | 2.03E-02 |
| MB12A_HUMAN\|Q96EY5_S188 | 1.7461289 | 2.61E-04 | 4.46E-03 |
| PHF2_HUMAN\|O75151_S1058 | 1.746047335 | 5.22E-05 | 5.37E-04 |
| CBX3_HUMAN\|Q13185_S176 | 1.745878537 | 7.36E-05 | 2.43E-03 |
| AHNK_HUMAN\|Q09666_S5739 | 1.74526133 | 1.32E-04 | 7.67E-03 |
| DESP_HUMAN\|P15924_S166 | 1.745204538 | 8.83E-05 | 2.63E-03 |
| K1C14_HUMAN\|P02533_S12 | 1.744292829 | 4.19E-06 | 1.20E-04 |

| | | | |
|---|---|---|---|
| CLIP4_HUMAN\|Q8N3C7_S610 | 1.74398009 | 4.64E-05 | 9.41E-04 |
| K1C14_HUMAN\|P02533_S437_S438 | 1.743965323 | 7.16E-05 | 1.01E-03 |
| LARP1_HUMAN\|Q6PKG0_S631_Y633 | 1.743712164 | 2.65E-03 | 2.60E-02 |
| PKP3_HUMAN\|Q9Y446_S180 | 1.743690844 | 4.31E-05 | 6.81E-04 |
| SRP09_HUMAN\|P49458_T85 | 1.743660977 | 1.54E-05 | 6.57E-04 |
| CCDC9_HUMAN\|Q9Y3X0_T389 | 1.743262432 | 1.28E-07 | 1.35E-05 |
| PRC2A_HUMAN\|P48634_S1085 | 1.742907069 | 3.93E-05 | 4.38E-04 |
| NAV1_HUMAN\|Q8NEY1_S648 | 1.742825933 | 3.78E-03 | 3.27E-02 |
| HMGA2_HUMAN\|P52926_S101_S105_T100 | 1.742679474 | 4.77E-05 | 1.40E-03 |
| TLN1_HUMAN\|Q9Y490_S1641 | 1.742365971 | 4.95E-07 | 5.41E-05 |
| K2C5_HUMAN\|P13647_S75 | 1.742272133 | 1.43E-04 | 9.32E-03 |
| K1C19_HUMAN\|P08727_S22 | 1.74223804 | 5.40E-05 | 1.53E-03 |
| CLIP4_HUMAN\|Q8N3C7_T582 | 1.742144115 | 7.28E-06 | 1.17E-04 |
| CSCL2_HUMAN\|Q5T3F8_S114_T110 | 1.742140436 | 2.91E-05 | 6.54E-04 |
| CRBG1_HUMAN\|Q9Y4K1_S410 | 1.742080701 | 4.07E-04 | 1.63E-02 |
| TANC2_HUMAN\|Q9HCD6_S1827 | 1.741771675 | 1.56E-06 | 3.69E-05 |
| ZO2_HUMAN\|Q9UDY2_S170 | 1.741563172 | 1.92E-06 | 9.55E-05 |
| RBP10_HUMAN\|Q6VN20_S365 | 1.74144625 | 8.52E-07 | 3.66E-05 |
| K2C5_HUMAN\|P13647_S574 | 1.741443 | 6.43E-06 | 1.63E-04 |
| K1C17_HUMAN\|Q04695_S13 | 1.740611728 | 1.61E-05 | 3.26E-04 |
| K2C5_HUMAN\|P13647_S573 | 1.740587479 | 5.06E-03 | 4.08E-02 |
| TOM1_HUMAN\|O60784_S462 | 1.740493787 | 7.56E-05 | 7.19E-04 |
| MARCS_HUMAN\|P29966_S118 | 1.740263973 | 4.65E-04 | 9.15E-03 |
| MAP7_HUMAN\|Q14244_T184 | 1.740230356 | 5.82E-06 | 2.78E-04 |
| IWS1_HUMAN\|Q96ST2_S398 | 1.740058917 | 1.66E-03 | 3.01E-02 |
| PGAM1_HUMAN\|P18669_S14 | 1.739981326 | 3.97E-05 | 1.23E-03 |
| WFS1_HUMAN\|O76024_S235 | 1.739964667 | 1.83E-03 | 2.40E-02 |
| CD97_HUMAN\|P48960_S831 | 1.739641838 | 6.57E-06 | 3.64E-04 |
| LAP2A_HUMAN\|P42166_S167_T160 | 1.739640735 | 2.46E-05 | 4.58E-04 |
| DC1L2_HUMAN\|O43237_S446 | 1.739427001 | 2.33E-03 | 2.31E-02 |
| YTDC2_HUMAN\|Q9H6S0_S1202 | 1.739111096 | 4.61E-06 | 1.67E-04 |

| | | | |
|---|---|---|---|
| K1C17_HUMAN\|Q04695_S28 | 1.738886375 | 9.91E-05 | 2.40E-03 |
| K1522_HUMAN\|Q9P206_S161 | 1.738345258 | 2.50E-06 | 1.51E-04 |
| INT12_HUMAN\|Q96CB8_S389 | 1.738020783 | 4.67E-04 | 3.26E-03 |
| SMTN_HUMAN\|P53814_S301 | 1.737961845 | 2.41E-03 | 1.57E-02 |
| ZC3HD_HUMAN\|Q5T200_S1014_S1017 | 1.737881892 | 5.52E-05 | 2.41E-03 |
| JUN_HUMAN\|P05412_S63 | 1.737641904 | 9.96E-05 | 3.20E-03 |
| IPP2M_HUMAN\|Q6NXS1_S121 | 1.736645862 | 1.26E-04 | 2.00E-03 |
| AHNK_HUMAN\|Q09666_S4092 | 1.736405018 | 7.68E-04 | 1.06E-02 |
| AHNK_HUMAN\|Q09666_S5746_S5752 | 1.736239927 | 2.10E-06 | 4.21E-04 |
| MA7D1_HUMAN\|Q3KQU3_S313 | 1.73615308 | 2.42E-04 | 2.68E-03 |
| WDR44_HUMAN\|Q5JSH3_S561 | 1.735617689 | 5.31E-06 | 4.43E-04 |
| AHNK_HUMAN\|Q09666_S212 | 1.735092185 | 9.59E-05 | 1.53E-03 |
| HES1_HUMAN\|Q14469_S10 | 1.734889898 | 1.51E-03 | 1.69E-02 |
| HMGN1_HUMAN\|P05114_S7 | 1.734858037 | 8.75E-05 | 3.00E-03 |
| FA83H_HUMAN\|Q6ZRV2_S1025 | 1.734819123 | 1.28E-05 | 4.54E-04 |
| WAC2A_HUMAN\|Q641Q2_S284 | 1.733885417 | 2.20E-04 | 1.79E-03 |
| SRS10_HUMAN\|O75494_S106 | 1.733880284 | 3.65E-04 | 6.12E-03 |
| PKP1_HUMAN\|Q13835_S142 | 1.733592465 | 2.30E-03 | 1.52E-02 |
| PDLI2_HUMAN\|Q96JY6_S127_S129 | 1.733549711 | 2.69E-04 | 2.06E-03 |
| PML_HUMAN\|P29590_S504 | 1.733468926 | 5.21E-06 | 9.14E-05 |
| MK03_HUMAN\|P27361_T198_T202 | 1.733182309 | 6.83E-05 | 1.63E-03 |
| LARP1_HUMAN\|Q6PKG0_S766 | 1.732846126 | 3.98E-03 | 3.73E-02 |
| B4DQA8_HUMAN\|B4DQA8_S49 | 1.73269636 | 1.17E-04 | 1.82E-03 |
| PPM1G_HUMAN\|O15355_S183 | 1.732675788 | 1.53E-05 | 5.15E-04 |
| LAD1_HUMAN\|O00515_T41 | 1.732467744 | 3.63E-04 | 5.60E-03 |
| NHSL1_HUMAN\|Q5SYE7_S722_S723 | 1.732332432 | 1.95E-04 | 1.63E-03 |
| ARP19_HUMAN\|P56211_Y59 | 1.731734679 | 5.58E-06 | 3.33E-04 |
| NUP98_HUMAN\|P52948_T1000 | 1.731695087 | 1.90E-04 | 1.60E-03 |
| DFFA_HUMAN\|O00273-2_S257 | 1.731247655 | 4.60E-03 | 4.11E-02 |
| PKP3_HUMAN\|Q9Y446_S313_S314 | 1.730621335 | 3.53E-06 | 1.90E-04 |
| CRBG1_HUMAN\|Q9Y4K1_S410 | 1.730521563 | 4.57E-04 | 6.66E-03 |

| | | | |
|---|---|---|---|
| RBBP6_HUMAN\|Q7Z6E9_S1644_S1651 | 1.730492485 | 1.01E-02 | 3.49E-02 |
| STXB4_HUMAN\|Q6ZWJ1_S463 | 1.730472371 | 2.92E-03 | 1.81E-02 |
| IWS1_HUMAN\|Q96ST2_S300_S313_S315 | 1.730398531 | 2.63E-03 | 1.20E-02 |
| MRCKG_HUMAN\|Q6DT37_S1544 | 1.730388158 | 2.97E-04 | 6.59E-03 |
| PKP1_HUMAN\|Q13835_S142 | 1.729841344 | 2.38E-03 | 2.41E-02 |
| NBEL2_HUMAN\|Q6ZNJ1_S2739 | 1.729607211 | 3.68E-03 | 2.48E-02 |
| PKP2_HUMAN\|Q99959_S132 | 1.729592082 | 1.32E-04 | 2.90E-03 |
| K1C14_HUMAN\|P02533_S17 | 1.729491832 | 7.01E-04 | 9.27E-03 |
| K2C75_HUMAN\|O95678_S40 | 1.72925985 | 1.70E-03 | 1.21E-02 |
| SEPT9_HUMAN\|Q9UHD8_T42 | 1.729099659 | 5.13E-06 | 1.38E-04 |
| OSB11_HUMAN\|Q9BXB4_S181 | 1.729003501 | 3.38E-06 | 1.01E-04 |
| K1C15_HUMAN\|P19012_S48 | 1.728999329 | 2.60E-03 | 2.49E-02 |
| SRPK2_HUMAN\|P78362_S497_T492 | 1.728885445 | 6.40E-06 | 4.13E-04 |
| DCBD2_HUMAN\|Q96PD2_S724_S727 | 1.727740174 | 6.32E-04 | 5.68E-03 |
| BAIP2_HUMAN\|Q9UQB8_T360 | 1.726951866 | 3.26E-03 | 1.98E-02 |
| TTP_HUMAN\|P26651_S186 | 1.726897469 | 1.36E-03 | 1.54E-02 |
| MAGD2_HUMAN\|Q9UNF1_S190_S191 | 1.726699514 | 3.50E-03 | 2.47E-02 |
| K2C5_HUMAN\|P13647_S6 | 1.726583448 | 2.54E-05 | 7.72E-04 |
| PAXX_HUMAN\|Q9BUH6_S152 | 1.72629138 | 7.01E-04 | 1.15E-02 |
| AHNK_HUMAN\|Q09666_S5749 | 1.726276288 | 1.42E-07 | 3.09E-05 |
| CSCL2_HUMAN\|Q5T3F8_T110 | 1.726206544 | 5.14E-04 | 3.24E-03 |
| PHLB2_HUMAN\|Q86SQ0_Y567 | 1.726085547 | 3.30E-06 | 1.79E-04 |
| NCOA7_HUMAN\|Q8NI08_S208_T210 | 1.725774943 | 6.57E-06 | 2.27E-04 |
| SMRCD_HUMAN\|Q9H4L7_S96 | 1.725710822 | 1.16E-05 | 6.90E-04 |
| CCDC9_HUMAN\|Q9Y3X0_S376_S386_S390 | 1.725594335 | 8.12E-04 | 6.80E-03 |
| ARP19_HUMAN\|P56211_S62 | 1.725553365 | 9.07E-06 | 3.78E-04 |
| EXPH5_HUMAN\|Q8NEV8_S341 | 1.725490341 | 1.29E-05 | 5.88E-04 |
| PARP4_HUMAN\|Q9UKK3_S1722 | 1.725345099 | 1.48E-05 | 7.94E-04 |
| K1C14_HUMAN\|P02533_S462 | 1.725164331 | 1.82E-04 | 3.38E-03 |
| MFAP1_HUMAN\|P55081_S116 | 1.72515805 | 1.58E-03 | 1.38E-02 |
| CBLB_HUMAN\|Q13191_S480_S484 | 1.72449796 | 1.59E-06 | 8.31E-05 |

| | | | |
|---|---|---|---|
| SH24A_HUMAN|Q9H788_S316 | 1.724428024 | 9.25E-05 | 1.51E-03 |
| RIN1_HUMAN|Q13671_S333_S337 | 1.724385075 | 8.91E-06 | 1.34E-04 |
| UBP5_HUMAN|P45974_S785 | 1.72438099 | 2.87E-03 | 3.22E-02 |
| ITB4_HUMAN|P16144_S1454_S1457 | 1.724354446 | 3.18E-03 | 2.97E-02 |
| AHNK2_HUMAN|Q8IVF2_S1712 | 1.723960114 | 1.24E-05 | 1.76E-04 |
| CDK13_HUMAN|Q14004_T1056 | 1.723826293 | 3.51E-05 | 5.97E-04 |
| ARHG5_HUMAN|Q12774_S445_S450 | 1.72380016 | 3.60E-07 | 1.13E-04 |
| RPP25_HUMAN|Q9BUL9_S162 | 1.72355888 | 1.89E-05 | 4.91E-04 |
| ARG35_HUMAN|A5YM69_S450 | 1.723514732 | 2.92E-04 | 3.77E-03 |
| FA83H_HUMAN|Q6ZRV2_S513_S516 | 1.723388702 | 3.12E-04 | 5.49E-03 |
| TPR_HUMAN|P12270_S2155 | 1.7233488 | 2.66E-05 | 3.50E-03 |
| PDZD2_HUMAN|O15018_S920 | 1.723322442 | 1.45E-03 | 1.92E-02 |
| RNF25_HUMAN|Q96BH1_S450 | 1.723279292 | 9.15E-05 | 7.09E-03 |
| ALDOA_HUMAN|P04075_S46 | 1.723217525 | 4.38E-05 | 1.32E-03 |
| K1C17_HUMAN|Q04695_S12 | 1.722798615 | 1.29E-04 | 1.63E-03 |
| SEPT7_HUMAN|Q16181_S423 | 1.722710707 | 4.39E-04 | 5.11E-03 |
| AFAD_HUMAN|P55196_S1172_S1173_S1182 | 1.721924194 | 1.72E-06 | 6.09E-05 |
| FGD6_HUMAN|Q6ZV73_S721 | 1.721711545 | 1.21E-04 | 1.87E-03 |
| CR025_HUMAN|Q96B23_S69 | 1.721585251 | 2.67E-03 | 3.11E-02 |
| K1C17_HUMAN|Q04695_S13 | 1.721579697 | 2.14E-05 | 8.12E-04 |
| RAI14_HUMAN|Q9P0K7-4_S261 | 1.721364608 | 2.54E-05 | 5.97E-04 |
| EIF3E_HUMAN|P60228_S442 | 1.721304423 | 1.44E-03 | 1.23E-02 |
| JUPI2_HUMAN|Q9H910_S97 | 1.720703488 | 7.47E-05 | 1.04E-03 |
| HMGA2_HUMAN|P52926_S105 | 1.72068975 | 3.41E-03 | 3.63E-02 |
| K1C14_HUMAN|P02533_S435_S437 | 1.72061486 | 3.57E-04 | 3.62E-03 |
| PKP1_HUMAN|Q13835_S121 | 1.720372074 | 2.57E-03 | 2.55E-02 |
| UT14A_HUMAN|Q9BVJ6_S437 | 1.720198886 | 1.19E-04 | 1.04E-03 |
| K1C14_HUMAN|P02533_S12 | 1.720165141 | 6.27E-06 | 3.58E-04 |
| PKP1_HUMAN|Q13835_S205 | 1.720103455 | 1.12E-02 | 4.93E-02 |
| CR025_HUMAN|Q96B23_S66_S69 | 1.719812981 | 1.43E-03 | 1.60E-02 |
| IRS2_HUMAN|Q9Y4H2_S1186_S1203 | 1.719572223 | 1.46E-05 | 4.01E-04 |

| | | | |
|---|---|---|---|
| GA2L1_HUMAN\|Q99501_S352 | 1.719196419 | 1.11E-03 | 2.97E-02 |
| CLIP4_HUMAN\|Q8N3C7_T582 | 1.719118955 | 1.06E-05 | 6.16E-04 |
| MINK1_HUMAN\|Q8N4C8_S701 | 1.71889916 | 1.40E-03 | 1.63E-02 |
| K1C17_HUMAN\|Q04695_S10 | 1.718524746 | 6.50E-05 | 9.30E-04 |
| TLN1_HUMAN\|Q9Y490_S1641 | 1.718015658 | 7.53E-07 | 8.87E-05 |
| RTKN_HUMAN\|Q9BST9_S543 | 1.717773046 | 5.41E-06 | 1.42E-04 |
| SERF2_HUMAN\|P84101_S21 | 1.717466023 | 6.50E-05 | 1.74E-03 |
| HSPB1_HUMAN\|P04792_S78_S82 | 1.71731657 | 4.67E-04 | 2.99E-03 |
| TRI29_HUMAN\|Q14134_S152 | 1.717056056 | 2.26E-05 | 8.34E-04 |
| L7UQD0_HUMAN\|L7UQD0_S268 | 1.717009934 | 6.04E-05 | 1.48E-03 |
| SF3B2_HUMAN\|Q13435_S303_S309 | 1.716724269 | 3.03E-03 | 3.12E-02 |
| K1C16_HUMAN\|P08779_S447_S450 | 1.71649642 | 3.87E-05 | 1.07E-03 |
| MILK1_HUMAN\|Q8N3F8_T318 | 1.716487091 | 1.19E-03 | 1.62E-02 |
| ATX2L_HUMAN\|Q8WWM7_S339 | 1.716265647 | 4.06E-08 | 4.31E-06 |
| LIMA1_HUMAN\|Q9UHB6_S507 | 1.716219535 | 2.54E-04 | 3.40E-02 |
| LPP_HUMAN\|Q93052_S609 | 1.71620522 | 2.87E-04 | 3.72E-03 |
| IF4B_HUMAN\|P23588_S497_S504_T500 | 1.71611166 | 6.84E-05 | 2.36E-03 |
| KPCD2_HUMAN\|Q9BZL6_S212 | 1.716107624 | 6.95E-05 | 1.65E-03 |
| HIRP3_HUMAN\|Q9BW71_S551_S555 | 1.716061386 | 4.50E-03 | 2.88E-02 |
| K2C5_HUMAN\|P13647_S50 | 1.716013949 | 3.89E-06 | 2.05E-04 |
| EIF3H_HUMAN\|O15372_S183 | 1.71564527 | 6.43E-04 | 5.76E-03 |
| EFNB1_HUMAN\|P98172_T284 | 1.715138438 | 1.36E-03 | 3.40E-02 |
| TISB_HUMAN\|Q07352_S92 | 1.715078015 | 1.69E-04 | 2.01E-03 |
| K1C14_HUMAN\|P02533_S435_S437 | 1.714809433 | 3.82E-04 | 5.83E-03 |
| PRC2A_HUMAN\|P48634_S1085 | 1.714730289 | 5.66E-05 | 1.09E-03 |
| K2C5_HUMAN\|P13647_S18 | 1.714530818 | 2.58E-05 | 9.28E-04 |
| PCM1_HUMAN\|Q15154_S93 | 1.714309011 | 7.62E-07 | 4.86E-05 |
| NFIB_HUMAN\|O00712_S276_S295 | 1.714287674 | 1.40E-04 | 3.96E-03 |
| MTMR3_HUMAN\|Q13615_T611 | 1.714211631 | 2.78E-04 | 6.28E-03 |
| AHNK2_HUMAN\|Q8IVF2_S5175 | 1.714198734 | 5.68E-05 | 2.44E-03 |
| DKC1_HUMAN\|O60832_S485_S494 | 1.71406024 | 1.39E-06 | 7.47E-05 |

| | | | |
|---|---|---|---|
| PPIG_HUMAN\|Q13427_S696 | 1.713836759 | 4.17E-04 | 8.18E-03 |
| MTAP2_HUMAN\|P11137_S1782_S1790 | 1.713791383 | 2.17E-03 | 1.03E-02 |
| K1C19_HUMAN\|P08727_S13 | 1.713550505 | 5.07E-04 | 9.39E-03 |
| SRSF6_HUMAN\|Q13247_S303 | 1.71343527 | 9.50E-05 | 8.77E-04 |
| TBC9B_HUMAN\|Q66K14_S411_S435 | 1.713420737 | 6.51E-03 | 3.93E-02 |
| DC1I2_HUMAN\|Q13409-2_S81 | 1.713053814 | 8.55E-04 | 1.37E-02 |
| TBC9B_HUMAN\|Q66K14_S411 | 1.712982467 | 2.48E-04 | 5.47E-03 |
| MYPT1_HUMAN\|O14974_S554 | 1.7129626 | 2.72E-03 | 2.65E-02 |
| ABI2_HUMAN\|Q9NYB9_Y371 | 1.712780802 | 3.30E-06 | 9.88E-05 |
| ANS1A_HUMAN\|Q92625_S649 | 1.71274255 | 3.55E-07 | 1.93E-05 |
| DFFA_HUMAN\|O00273-2_S257 | 1.712701937 | 5.33E-03 | 4.23E-02 |
| EF2K_HUMAN\|O00418_S470 | 1.711475315 | 2.25E-05 | 8.34E-04 |
| ARG28_HUMAN\|Q8N1W1_S513 | 1.711118193 | 1.91E-05 | 9.31E-04 |
| SF3B2_HUMAN\|Q13435_S309 | 1.710293772 | 1.31E-03 | 7.57E-03 |
| DDX10_HUMAN\|Q13206_S539 | 1.710134245 | 1.01E-04 | 9.72E-04 |
| MRCKG_HUMAN\|Q6DT37_S1544 | 1.70999889 | 3.74E-04 | 4.50E-03 |
| ITB4_HUMAN\|P16144_S1515_S1517 | 1.709850933 | 4.07E-04 | 2.70E-03 |
| MK14_HUMAN\|Q16539_Y182 | 1.709677987 | 5.40E-05 | 1.83E-03 |
| IFIT3_HUMAN\|O14879_S484 | 1.709674423 | 5.10E-03 | 4.74E-02 |
| CLD4_HUMAN\|O14493_Y208 | 1.709588458 | 1.31E-03 | 1.51E-02 |
| IASPP_HUMAN\|Q8WUF5_S74 | 1.709232793 | 5.24E-07 | 2.63E-05 |
| NIBAN_HUMAN\|Q9BZQ8_S926 | 1.708878711 | 5.80E-06 | 4.25E-04 |
| CSKI2_HUMAN\|Q8WXE0_S403 | 1.708813437 | 1.41E-06 | 5.21E-05 |
| PML_HUMAN\|P29590_S518_S527 | 1.708704853 | 4.10E-04 | 4.85E-03 |
| TB182_HUMAN\|Q9C0C2_S920 | 1.708576291 | 5.51E-05 | 2.03E-03 |
| CLAP2_HUMAN\|O75122_S463 | 1.708455027 | 8.12E-04 | 5.18E-03 |
| GGYF1_HUMAN\|O75420_S157 | 1.70830087 | 2.07E-03 | 4.42E-02 |
| NEB2_HUMAN\|Q96SB3_S99 | 1.708149748 | 2.51E-05 | 1.15E-03 |
| DC1L1_HUMAN\|Q9Y6G9_S516 | 1.707935687 | 5.32E-06 | 4.43E-04 |
| PAR10_HUMAN\|Q53GL7_S431 | 1.707820228 | 2.81E-06 | 2.87E-04 |
| CNKR1_HUMAN\|Q969H4_S314 | 1.707522366 | 4.68E-06 | 1.15E-03 |

| | | | |
|---|---|---|---|
| ACAP2_HUMAN\|Q15057_T382 | 1.707490256 | 2.60E-07 | 4.26E-05 |
| FRM4B_HUMAN\|Q9Y2L6_S628 | 1.707448057 | 4.55E-05 | 4.84E-03 |
| COBL1_HUMAN\|Q53SF7_S294_T298 | 1.707095019 | 4.83E-05 | 1.24E-03 |
| K1522_HUMAN\|Q9P206_S161 | 1.706915779 | 4.26E-06 | 2.76E-04 |
| MAN1_HUMAN\|Q9Y2U8_S144 | 1.706346838 | 6.24E-04 | 3.76E-03 |
| MAP7_HUMAN\|Q14244_S183 | 1.706301433 | 2.55E-06 | 1.52E-04 |
| HMGA1_HUMAN\|P17096_S49 | 1.706098818 | 3.75E-03 | 3.31E-02 |
| SF3B2_HUMAN\|Q13435_S307_S309 | 1.705997521 | 1.93E-03 | 2.26E-02 |
| GYS1_HUMAN\|P13807_S727_S731 | 1.705947746 | 1.68E-04 | 2.45E-03 |
| NFIC_HUMAN\|P08651_S284_S304_T293 | 1.705655426 | 6.73E-05 | 1.20E-03 |
| TRM1_HUMAN\|Q9NXH9_S625 | 1.705425791 | 8.59E-05 | 1.15E-03 |
| WDR13_HUMAN\|Q9H1Z4_S70_S74 | 1.705172602 | 1.90E-05 | 7.65E-04 |
| MACF1_HUMAN\|Q9UPN3_T7287 | 1.704437428 | 1.23E-03 | 1.67E-02 |
| 1433Z_HUMAN\|P63104_T232 | 1.704285704 | 4.23E-04 | 4.95E-03 |
| BAD_HUMAN\|Q92934_Y76 | 1.70421863 | 2.63E-06 | 2.23E-04 |
| K1C14_HUMAN\|P02533_S432_S433_S438 | 1.704206532 | 3.72E-04 | 7.41E-03 |
| NKTR_HUMAN\|P30414_S463 | 1.704150053 | 5.80E-05 | 2.07E-03 |
| CAMP2_HUMAN\|Q08AD1_S691 | 1.703785088 | 3.03E-04 | 5.45E-03 |
| 2A5E_HUMAN\|Q16537_S33_S34 | 1.703717736 | 4.97E-07 | 6.74E-05 |
| TBCD5_HUMAN\|Q92609_S522 | 1.70364281 | 4.51E-05 | 1.62E-03 |
| MY18A_HUMAN\|Q92614_S1998 | 1.703640489 | 6.98E-06 | 1.14E-04 |
| F122A_HUMAN\|Q96E09_S37 | 1.703290871 | 4.70E-03 | 3.87E-02 |
| RPRD2_HUMAN\|Q5VT52_T723 | 1.702166115 | 6.16E-07 | 2.95E-05 |
| RFIP1_HUMAN\|Q6WKZ4_S501 | 1.702140663 | 7.54E-04 | 9.79E-03 |
| SCAM2_HUMAN\|O15127_S319 | 1.701788085 | 4.89E-04 | 3.11E-03 |
| ATN1_HUMAN\|P54259_S77_S79 | 1.701538323 | 2.79E-04 | 5.14E-03 |
| K1C14_HUMAN\|P02533_S435_S438 | 1.701268083 | 2.63E-04 | 2.87E-03 |
| PHLB1_HUMAN\|Q86UU1_S430 | 1.701219047 | 5.91E-05 | 1.94E-03 |
| MTUS1_HUMAN\|Q9ULD2_T1204 | 1.700834216 | 1.70E-05 | 4.46E-04 |
| K2C75_HUMAN\|O95678_S535 | 1.700811626 | 1.00E-06 | 4.09E-05 |
| ZBT21_HUMAN\|Q9ULJ3_S435_T431 | 1.700659246 | 4.84E-06 | 1.34E-04 |

| | | | |
|---|---|---|---|
| COBL1_HUMAN\|Q53SF7_T298 | 1.700637335 | 6.85E-05 | 1.80E-03 |
| PAK1_HUMAN\|Q13153_S223 | 1.700368447 | 1.30E-04 | 1.11E-03 |
| KLC3_HUMAN\|Q6P597_S167_S173 | 1.700355219 | 1.07E-04 | 2.20E-03 |
| TFE3_HUMAN\|P19532_S548 | 1.700225698 | 7.16E-04 | 4.66E-03 |
| ACAP2_HUMAN\|Q15057_T382 | 1.699649178 | 2.99E-07 | 2.38E-05 |
| KLC1_HUMAN\|Q07866-4_S631 | 1.699372631 | 1.11E-06 | 9.01E-05 |
| SOS1_HUMAN\|Q07889_S1078_S1082 | 1.699336295 | 7.27E-05 | 2.99E-03 |
| JUNB_HUMAN\|P17275_T255 | 1.699284173 | 4.27E-07 | 2.10E-04 |
| DTD1_HUMAN\|Q8TEA8_S196_S197 | 1.699271711 | 1.66E-06 | 1.67E-04 |
| TBCD5_HUMAN\|Q92609_S554 | 1.699147267 | 7.94E-05 | 2.66E-03 |
| EMAL3_HUMAN\|Q32P44_S889 | 1.698996988 | 2.79E-03 | 1.26E-02 |
| HNRPC_HUMAN\|P07910_S240 | 1.698714166 | 4.31E-05 | 1.15E-03 |
| FYN_HUMAN\|P06241_Y420 | 1.698542943 | 4.51E-03 | 3.69E-02 |
| K2C5_HUMAN\|P13647_S64 | 1.698108928 | 4.46E-04 | 1.78E-02 |
| HES1_HUMAN\|Q14469_S10 | 1.697378244 | 2.14E-03 | 2.49E-02 |
| PFD4_HUMAN\|Q9NQP4_S125 | 1.697359628 | 2.23E-04 | 3.98E-03 |
| STK10_HUMAN\|O94804_T952 | 1.697224827 | 2.58E-04 | 4.82E-03 |
| P3C2A_HUMAN\|O00443_S259 | 1.696488468 | 1.10E-05 | 6.16E-04 |
| PALLD_HUMAN\|Q8WX93_S1352 | 1.696457323 | 4.15E-06 | 1.73E-04 |
| TAOK3_HUMAN\|Q9H2K8_S324 | 1.695933152 | 5.72E-06 | 1.97E-04 |
| K1C16_HUMAN\|P08779_S448_S451 | 1.695882667 | 6.52E-05 | 9.30E-04 |
| SERF2_HUMAN\|P84101_S21 | 1.695787561 | 8.83E-05 | 7.04E-03 |
| SOS1_HUMAN\|Q07889_S1161 | 1.695288959 | 4.68E-03 | 4.67E-02 |
| K2C5_HUMAN\|P13647_S11_S14 | 1.695210176 | 2.96E-06 | 9.16E-05 |
| MAP7_HUMAN\|Q14244_S183 | 1.694767652 | 3.14E-06 | 6.07E-04 |
| PLEC_HUMAN\|Q15149_S4389_S4391 | 1.694663228 | 1.82E-04 | 6.33E-03 |
| SERF2_HUMAN\|P84101_S21 | 1.694488837 | 9.00E-05 | 1.93E-03 |
| LASP1_HUMAN\|Q14847_T104 | 1.694471131 | 7.72E-06 | 3.35E-04 |
| NRK_HUMAN\|Q7Z2Y5_S852_Y858 | 1.694084272 | 1.44E-04 | 1.29E-03 |
| AHNK_HUMAN\|Q09666_S5745_S5749 | 1.693858205 | 1.64E-06 | 1.51E-04 |
| BAIP2_HUMAN\|Q9UQB8_T340 | 1.693724068 | 1.48E-03 | 1.67E-02 |

| | | | |
|---|---|---|---|
| MRP_HUMAN\|P49006_T178 | 1.693699941 | 2.93E-06 | 1.62E-04 |
| PPIG_HUMAN\|Q13427_S696 | 1.693634826 | 5.19E-04 | 5.80E-03 |
| SPTB2_HUMAN\|Q01082_T2328 | 1.693251547 | 4.01E-03 | 3.48E-02 |
| K2C5_HUMAN\|P13647_S11_S8 | 1.692942851 | 7.23E-06 | 1.78E-04 |
| IWS1_HUMAN\|Q96ST2_S398 | 1.692629435 | 2.52E-03 | 3.00E-02 |
| ENSA_HUMAN\|O43768_S109 | 1.692427956 | 8.97E-05 | 1.58E-03 |
| WAC2A_HUMAN\|Q641Q2_S375 | 1.692330757 | 4.40E-04 | 7.07E-03 |
| AKAP2_HUMAN\|Q9Y2D5_T157 | 1.692174938 | 4.18E-05 | 1.72E-03 |
| SSH2_HUMAN\|Q76I76_T486 | 1.691919219 | 8.78E-06 | 2.07E-04 |
| F122A_HUMAN\|Q96E09_S37 | 1.691590048 | 5.16E-03 | 2.75E-02 |
| ROA3_HUMAN\|P51991_S355 | 1.69132079 | 9.57E-07 | 3.93E-05 |
| CTSL2_HUMAN\|Q05D32_S9 | 1.691027728 | 1.05E-03 | 5.72E-03 |
| K1C14_HUMAN\|P02533_S435 | 1.69028944 | 2.05E-04 | 4.06E-03 |
| SRSF6_HUMAN\|Q13247_S303 | 1.690264949 | 1.27E-04 | 1.16E-03 |
| ML12A_HUMAN\|P19105_T18 | 1.690091111 | 6.43E-05 | 1.73E-03 |
| PGRC1_HUMAN\|O00264_S54 | 1.690088436 | 8.24E-04 | 5.22E-03 |
| SYNE2_HUMAN\|Q8WXH0_T2876 | 1.69002058 | 8.99E-04 | 5.06E-03 |
| SRC8_HUMAN\|Q14247_S405_S418_T401 | 1.68970979 | 1.46E-03 | 1.66E-02 |
| TAGL2_HUMAN\|P37802_S163 | 1.689191301 | 5.37E-04 | 9.87E-03 |
| ZBT21_HUMAN\|Q9ULJ3_S435_T431 | 1.688659362 | 5.92E-06 | 9.73E-04 |
| TERA_HUMAN\|P55072_S794 | 1.688607626 | 6.13E-04 | 9.01E-03 |
| IASPP_HUMAN\|Q8WUF5_Y109 | 1.688522329 | 1.63E-05 | 3.27E-04 |
| AHNK2_HUMAN\|Q8IVF2_S1712 | 1.688439879 | 2.15E-05 | 1.02E-03 |
| PKP3_HUMAN\|Q9Y446_Y176 | 1.688196605 | 6.84E-05 | 5.16E-03 |
| FA83H_HUMAN\|Q6ZRV2_S974 | 1.687753044 | 7.12E-06 | 1.15E-04 |
| PLEC_HUMAN\|Q15149_S4675 | 1.687522233 | 5.50E-04 | 9.96E-03 |
| IWS1_HUMAN\|Q96ST2_S313 | 1.687429834 | 5.01E-03 | 2.00E-02 |
| USO1_HUMAN\|O60763_S952 | 1.687290572 | 6.72E-04 | 3.99E-03 |
| LIMA1_HUMAN\|Q9UHB6_S507 | 1.687265937 | 3.55E-04 | 4.29E-03 |
| IRS2_HUMAN\|Q9Y4H2_S608 | 1.687200582 | 6.36E-06 | 4.47E-04 |
| SEPT9_HUMAN\|Q9UHD8_T38_T42 | 1.687124959 | 6.73E-06 | 3.04E-03 |

| | | | |
|---|---|---|---|
| MTMRA_HUMAN\|Q9NXD2_T583 | 1.686843121 | 1.48E-03 | 1.09E-02 |
| PININ_HUMAN\|Q9H307_S66 | 1.686797356 | 1.28E-04 | 3.34E-03 |
| KLC3_HUMAN\|Q6P597_S497_S502 | 1.686532114 | 4.41E-07 | 6.29E-05 |
| SSFA2_HUMAN\|P28290_S92 | 1.686529148 | 5.04E-03 | 3.99E-02 |
| COHA1_HUMAN\|Q9UMD9_S93_S96 | 1.686469915 | 8.07E-04 | 2.56E-02 |
| WAC2A_HUMAN\|Q641Q2_S352 | 1.686294563 | 2.82E-03 | 3.25E-02 |
| SMC4_HUMAN\|Q9NTJ3_S22 | 1.686209252 | 6.85E-03 | 2.96E-02 |
| ATX2L_HUMAN\|Q8WWM7_S680_S684 | 1.685847599 | 7.10E-07 | 8.75E-05 |
| DBNL_HUMAN\|Q9UJU6_T271 | 1.685590084 | 3.24E-05 | 1.38E-03 |
| ZO2_HUMAN\|Q9UDY2_S398 | 1.685360946 | 6.76E-04 | 4.01E-03 |
| SPTB2_HUMAN\|Q01082-3_S8 | 1.685231875 | 7.51E-05 | 1.71E-03 |
| PAK2_HUMAN\|Q13177_S141 | 1.684520283 | 1.75E-04 | 2.06E-03 |
| NELFE_HUMAN\|P18615_S251 | 1.68449416 | 3.95E-04 | 7.67E-03 |
| NUP98_HUMAN\|P52948_S612 | 1.684424095 | 1.41E-05 | 1.90E-04 |
| RB6I2_HUMAN\|Q8IUD2_S37_S42 | 1.684323603 | 8.42E-04 | 5.30E-03 |
| CCD86_HUMAN\|Q9H6F5_S217 | 1.684249611 | 2.76E-04 | 6.28E-03 |
| SRRM2_HUMAN\|Q9UQ35_S1152 | 1.684035901 | 3.92E-04 | 1.64E-02 |
| K2C5_HUMAN\|P13647_S36 | 1.684023292 | 8.21E-05 | 1.12E-03 |
| IF4G1_HUMAN\|Q04637_S1231 | 1.683798503 | 2.10E-05 | 1.02E-03 |
| 2A5E_HUMAN\|Q16537_S32_S34 | 1.683480666 | 1.01E-03 | 1.45E-02 |
| ARP19_HUMAN\|P56211_Y59 | 1.683289431 | 1.26E-05 | 4.51E-04 |
| PR40B_HUMAN\|Q6NWY9_S852_T842 | 1.682727773 | 3.29E-04 | 2.29E-03 |
| K1C19_HUMAN\|P08727_S22 | 1.682387679 | 1.25E-04 | 1.58E-03 |
| IWS1_HUMAN\|Q96ST2_S69_S80_T67 | 1.682191708 | 2.69E-03 | 1.22E-02 |
| ZFP91_HUMAN\|Q96JP5_S82 | 1.681870602 | 3.71E-05 | 6.16E-04 |
| LYAR_HUMAN\|Q9NX58_S276 | 1.681654795 | 1.97E-06 | 8.75E-05 |
| MICA3_HUMAN\|Q7RTP6_S1636 | 1.681646722 | 1.18E-03 | 9.10E-03 |
| COBL1_HUMAN\|Q53SF7_S1145 | 1.681453344 | 1.27E-04 | 2.00E-03 |
| HMGN1_HUMAN\|P05114_S7 | 1.681288445 | 1.81E-04 | 4.82E-03 |
| SCAM2_HUMAN\|O15127_S320 | 1.681236118 | 5.79E-04 | 3.54E-03 |
| LMNA_HUMAN\|P02545_S619 | 1.680990803 | 2.24E-04 | 4.35E-03 |

| | | | |
|---|---|---|---|
| AKAP2_HUMAN\|Q9Y2D5_S152_S155 | 1.680810177 | 1.62E-05 | 8.42E-04 |
| ABLM1_HUMAN\|O14639_S458 | 1.680613824 | 8.40E-04 | 6.95E-03 |
| ATAT_HUMAN\|Q5SQI0_S315 | 1.680555495 | 8.40E-07 | 3.65E-05 |
| HNRH3_HUMAN\|P31942_S298 | 1.680366851 | 3.17E-05 | 1.38E-03 |
| GA2L1_HUMAN\|Q99501_S352 | 1.680202673 | 1.67E-03 | 3.98E-02 |
| UBP5_HUMAN\|P45974_S785 | 1.679729123 | 4.15E-03 | 3.56E-02 |
| K1C14_HUMAN\|P02533_S11 | 1.679687886 | 7.72E-06 | 1.88E-04 |
| KC1A_HUMAN\|P48729_S312 | 1.679672008 | 7.63E-03 | 3.67E-02 |
| NFIC_HUMAN\|P08651_S473 | 1.679659422 | 1.27E-03 | 3.25E-02 |
| PLEC_HUMAN\|Q15149_S4389_S4391 | 1.679516271 | 2.21E-04 | 1.68E-03 |
| ZCH18_HUMAN\|Q86VM9_S532_S534 | 1.679106162 | 1.83E-07 | 3.30E-05 |
| PKHG3_HUMAN\|A1L390_T1042 | 1.678787262 | 5.95E-04 | 5.43E-03 |
| PGAM1_HUMAN\|P18669_S14 | 1.678378352 | 8.86E-05 | 1.91E-03 |
| NOP2_HUMAN\|P46087_T605 | 1.678360058 | 2.75E-05 | 3.29E-04 |
| HIRP3_HUMAN\|Q9BW71_S551_S555 | 1.678200907 | 6.07E-03 | 3.73E-02 |
| I2BP2_HUMAN\|Q7Z5L9_S455_S460 | 1.678153895 | 6.10E-05 | 2.19E-03 |
| TP53B_HUMAN\|Q12888_S366_T357 | 1.677725591 | 6.02E-05 | 1.66E-03 |
| LAP2A_HUMAN\|P42166_S159 | 1.677378746 | 4.02E-05 | 6.48E-04 |
| CR025_HUMAN\|Q96B23_S145_S147 | 1.677354034 | 3.85E-06 | 2.60E-04 |
| RBM7_HUMAN\|Q9Y580_S136 | 1.677188843 | 1.84E-04 | 4.43E-03 |
| E41L2_HUMAN\|O43491_S87 | 1.677075985 | 2.92E-05 | 5.14E-04 |
| CCD86_HUMAN\|Q9H6F5_S113 | 1.67693868 | 2.84E-03 | 4.29E-02 |
| CCDC9_HUMAN\|Q9Y3X0_S386_S390 | 1.67692687 | 1.60E-03 | 3.79E-02 |
| AHNK_HUMAN\|Q09666_S5752_S5763 | 1.676851172 | 3.93E-04 | 6.52E-03 |
| TRI25_HUMAN\|Q14258_S100 | 1.676685673 | 4.32E-04 | 4.20E-03 |
| PHLB2_HUMAN\|Q86SQ0_Y567 | 1.676350321 | 7.31E-06 | 3.92E-04 |
| PAWR_HUMAN\|Q96IZ0_S231 | 1.67603579 | 1.75E-05 | 3.47E-04 |
| MTAP2_HUMAN\|P11137_S1782 | 1.675974566 | 2.58E-03 | 4.05E-02 |
| Q659E9_HUMAN\|Q659E9_S168 | 1.675793352 | 1.36E-06 | 3.12E-05 |
| VIME_HUMAN\|P08670_S34 | 1.675719625 | 5.74E-05 | 2.07E-03 |
| EGFR_HUMAN\|P00533_S695 | 1.675647876 | 1.60E-03 | 1.40E-02 |

| | | | |
|---|---|---|---|
| ACSA_HUMAN\|Q9NR19_S267 | 1.675498663 | 4.34E-05 | 4.71E-03 |
| PKP3_HUMAN\|Q9Y446_Y176 | 1.675238557 | 8.15E-05 | 2.07E-03 |
| PR40B_HUMAN\|Q6NWY9_S852_T842 | 1.67485962 | 3.62E-04 | 2.63E-03 |
| SRSF6_HUMAN\|Q13247_S301 | 1.674804689 | 1.23E-04 | 1.07E-03 |
| NU214_HUMAN\|P35658_S1023 | 1.674424813 | 1.12E-04 | 3.02E-03 |
| NEK9_HUMAN\|Q8TD19_T333 | 1.674362679 | 7.74E-04 | 9.98E-03 |
| SCRIB_HUMAN\|Q14160_S1475_S1486 | 1.674113956 | 2.11E-03 | 1.00E-02 |
| AAK1_HUMAN\|Q2M2I8_T620 | 1.673824071 | 2.35E-05 | 5.70E-04 |
| K2C5_HUMAN\|P13647_S6 | 1.673783686 | 5.64E-05 | 2.07E-03 |
| TBCD4_HUMAN\|O60343_T752 | 1.673688805 | 2.29E-04 | 1.84E-03 |
| AHNK_HUMAN\|Q09666_S5864 | 1.673537425 | 3.47E-03 | 3.07E-02 |
| S20A2_HUMAN\|Q08357_Y423 | 1.673408659 | 1.34E-03 | 1.53E-02 |
| AHNK_HUMAN\|Q09666_S5745_S5749 | 1.673241233 | 2.40E-06 | 7.23E-04 |
| K1522_HUMAN\|Q9P206_S339 | 1.673096262 | 9.28E-03 | 4.86E-02 |
| PALLD_HUMAN\|Q8WX93_S726 | 1.673000745 | 6.28E-04 | 8.52E-03 |
| P63_HUMAN\|Q9H3D4_S463_T451 | 1.67285053 | 3.72E-03 | 1.79E-02 |
| STMN1_HUMAN\|P16949_S25 | 1.672712019 | 7.12E-06 | 4.43E-04 |
| SOS1_HUMAN\|Q07889_S1082 | 1.672552439 | 1.10E-02 | 4.35E-02 |
| ITB4_HUMAN\|P16144_S1457_T1455 | 1.672514026 | 5.45E-03 | 4.30E-02 |
| BORG4_HUMAN\|Q9H3Q1_S140_S141 | 1.672452923 | 3.28E-03 | 3.05E-02 |
| CING_HUMAN\|Q9P2M7_S129 | 1.672242425 | 7.23E-04 | 1.02E-02 |
| K2C8_HUMAN\|P05787_S21_S24_S27 | 1.672216177 | 1.10E-04 | 2.54E-03 |
| YETS2_HUMAN\|Q9ULM3_S536 | 1.672163142 | 1.75E-05 | 2.64E-03 |
| TFE3_HUMAN\|P19532_S548 | 1.671928176 | 9.65E-04 | 9.57E-03 |
| PLEC_HUMAN\|Q15149_S4613_S4618 | 1.671797907 | 1.92E-05 | 3.73E-04 |
| RAI14_HUMAN\|Q9P0K7_S281_S290 | 1.67148779 | 2.02E-03 | 9.70E-03 |
| AAK1_HUMAN\|Q2M2I8_T620 | 1.671176221 | 2.44E-05 | 3.31E-03 |
| 2A5E_HUMAN\|Q16537_S33_S34 | 1.671025658 | 9.57E-07 | 3.93E-05 |
| SARG_HUMAN\|Q9BW04_S429 | 1.67084367 | 4.39E-04 | 1.69E-02 |
| K2C75_HUMAN\|O95678_S61 | 1.669976766 | 1.46E-03 | 3.77E-02 |
| MISP_HUMAN\|Q8IVT2_S471 | 1.669751232 | 1.15E-04 | 2.60E-03 |

| | | | |
|---|---|---|---|
| KPCD2_HUMAN\|Q9BZL6_S214 | 1.669440203 | 1.72E-04 | 3.24E-03 |
| RAI14_HUMAN\|Q9P0K7_S290 | 1.668960672 | 7.11E-04 | 9.38E-03 |
| MDN1_HUMAN\|Q9NU22_S4538 | 1.668872273 | 7.21E-03 | 3.10E-02 |
| ROCK1_HUMAN\|Q13464_S1333_S1336_S1341 | 1.668549965 | 1.79E-03 | 4.11E-02 |
| COBL1_HUMAN\|Q53SF7_S1145 | 1.668545162 | 1.50E-04 | 3.81E-03 |
| PKP1_HUMAN\|Q13835_S121 | 1.66854499 | 4.14E-03 | 3.47E-02 |
| KDM3B_HUMAN\|Q7LBC6_S293 | 1.668382066 | 7.41E-07 | 1.95E-05 |
| MAP7_HUMAN\|Q14244_S202 | 1.668346139 | 4.82E-06 | 1.34E-04 |
| NIBL1_HUMAN\|Q96TA1_S641 | 1.66824297 | 1.24E-03 | 6.57E-03 |
| SCAM2_HUMAN\|O15127_S319 | 1.66803308 | 7.23E-04 | 1.18E-02 |
| NBEL2_HUMAN\|Q6ZNJ1_T1867 | 1.667964443 | 4.92E-04 | 3.12E-03 |
| CE170_HUMAN\|Q5SW79_S630_S633 | 1.667780682 | 7.90E-06 | 1.90E-04 |
| MISP_HUMAN\|Q8IVT2_S471 | 1.667617492 | 1.18E-04 | 2.38E-03 |
| MED24_HUMAN\|O75448_S873 | 1.667509527 | 1.49E-03 | 1.09E-02 |
| PRC2A_HUMAN\|P48634_S1085_S1089_S1092 | 1.667282323 | 1.13E-05 | 5.34E-04 |
| TB182_HUMAN\|Q9C0C2_T1282 | 1.66697573 | 2.33E-05 | 8.48E-04 |
| MAST4_HUMAN\|O15021_S2555 | 1.666686756 | 1.52E-04 | 2.91E-03 |
| FUBP2_HUMAN\|Q92945_T195 | 1.66659551 | 5.18E-03 | 4.08E-02 |
| DC1I2_HUMAN\|Q13409-2_S84 | 1.66611113 | 3.42E-05 | 3.88E-04 |
| INADL_HUMAN\|Q8NI35_S645 | 1.665800654 | 7.55E-06 | 3.30E-04 |
| ACAP2_HUMAN\|Q15057_S384_T382 | 1.665724533 | 3.07E-08 | 5.36E-06 |
| RAI14_HUMAN\|Q9P0K7_T297 | 1.665613135 | 4.21E-04 | 6.22E-03 |
| CTNA1_HUMAN\|P35221_S641_S652 | 1.665507677 | 6.85E-05 | 1.63E-03 |
| MEF2D_HUMAN\|Q14814_S231 | 1.665442645 | 1.54E-03 | 3.71E-02 |
| RFIP3_HUMAN\|O75154_S52 | 1.665282059 | 7.34E-05 | 2.69E-03 |
| MILK1_HUMAN\|Q8N3F8_S418 | 1.665106167 | 1.40E-02 | 4.53E-02 |
| CCDC9_HUMAN\|Q9Y3X0_S376_S386_T381 | 1.664790213 | 3.21E-05 | 9.36E-04 |
| CND1_HUMAN\|Q15021_S1330_S1333 | 1.664337421 | 3.82E-03 | 3.28E-02 |
| SRPK2_HUMAN\|P78362_S494_T498 | 1.664261989 | 1.31E-03 | 1.85E-02 |
| WDR44_HUMAN\|Q5JSH3_S561 | 1.664149196 | 1.61E-05 | 4.35E-04 |
| MA7D1_HUMAN\|Q3KQU3_S452 | 1.6641204 | 1.61E-04 | 3.38E-03 |

| | | | |
|---|---|---|---|
| PKN2_HUMAN\|Q16513_S362 | 1.664103834 | 6.51E-05 | 9.30E-04 |
| YETS2_HUMAN\|Q9ULM3_S536 | 1.663966258 | 2.00E-05 | 3.85E-04 |
| NEB2_HUMAN\|Q96SB3_S99 | 1.663955421 | 4.79E-05 | 9.50E-04 |
| E41L2_HUMAN\|O43491_S87 | 1.663662023 | 3.58E-05 | 1.14E-03 |
| LMNB2_HUMAN\|Q03252_S421_S424 | 1.663513471 | 9.49E-05 | 1.99E-03 |
| SOS2_HUMAN\|Q07890_S1267 | 1.663095032 | 5.74E-04 | 5.29E-03 |
| OSB11_HUMAN\|Q9BXB4_S174_S181 | 1.662919224 | 2.30E-03 | 1.52E-02 |
| NP1L1_HUMAN\|P55209_T62 | 1.662561685 | 1.40E-03 | 1.63E-02 |
| ATX2L_HUMAN\|Q8WWM7_S594 | 1.662336754 | 2.49E-03 | 3.97E-02 |
| KLC3_HUMAN\|Q6P597_S495_S502 | 1.662274059 | 1.25E-03 | 1.75E-02 |
| PKP2_HUMAN\|Q99959_S151 | 1.662264808 | 1.66E-05 | 8.51E-04 |
| RALY_HUMAN\|Q9UKM9_T286_T298 | 1.662070392 | 3.14E-04 | 2.33E-03 |
| CLIP2_HUMAN\|Q9UDT6_S207_T198 | 1.662034355 | 4.88E-04 | 5.46E-03 |
| TPC12_HUMAN\|Q8WVT3_S184 | 1.662028683 | 4.54E-04 | 9.24E-03 |
| MP2K4_HUMAN\|P45985_S393 | 1.661979485 | 1.75E-03 | 2.34E-02 |
| MARK2_HUMAN\|Q7KZI7-13_S376 | 1.661717384 | 1.83E-03 | 4.16E-02 |
| FA83B_HUMAN\|Q5T0W9_S802 | 1.661568652 | 4.13E-05 | 4.57E-03 |
| K2C5_HUMAN\|P13647_S571_S574 | 1.661564187 | 1.05E-04 | 3.03E-03 |
| AT11C_HUMAN\|Q8NB49_T1124 | 1.661562129 | 3.24E-05 | 1.35E-03 |
| MICA1_HUMAN\|Q8TDZ2_S960 | 1.661556794 | 4.43E-06 | 7.82E-05 |
| DC1L1_HUMAN\|Q9Y6G9_T513_T515 | 1.661416048 | 2.43E-05 | 1.27E-03 |
| PININ_HUMAN\|Q9H307_S690 | 1.661267624 | 5.39E-04 | 5.94E-03 |
| K2C75_HUMAN\|O95678_S535 | 1.660608529 | 2.15E-06 | 1.35E-04 |
| TMF1_HUMAN\|P82094_S72_S77 | 1.660535475 | 1.01E-03 | 6.09E-03 |
| NBN_HUMAN\|O60934_S343 | 1.66049681 | 4.79E-05 | 1.40E-03 |
| GBF1_HUMAN\|Q92538_S1781_S1784 | 1.660381686 | 1.32E-04 | 1.12E-03 |
| K2C5_HUMAN\|P13647_S572 | 1.660361329 | 1.04E-04 | 2.45E-03 |
| GRAP1_HUMAN\|Q4V328_S664 | 1.660352413 | 8.42E-05 | 1.84E-03 |
| KCD12_HUMAN\|Q96CX2_S200_T196 | 1.659694548 | 2.41E-03 | 2.91E-02 |
| PLEC_HUMAN\|Q15149_S4384_S4385 | 1.659535485 | 2.10E-04 | 1.73E-03 |
| COBL1_HUMAN\|Q53SF7_S1145 | 1.659269378 | 1.69E-04 | 2.45E-03 |

| | | | |
|---|---|---|---|
| SLK_HUMAN\|Q9H2G2_S655 | 1.659206965 | 4.38E-04 | 4.56E-02 |
| FIL1L_HUMAN\|Q4L180_S1049 | 1.65902393 | 7.32E-04 | 4.74E-03 |
| MED24_HUMAN\|O75448_S873 | 1.659001898 | 1.63E-03 | 3.93E-02 |
| MKL2_HUMAN\|Q9ULH7_S921 | 1.658895739 | 5.18E-05 | 5.34E-04 |
| RABE1_HUMAN\|Q15276_S407 | 1.65886909 | 1.48E-05 | 4.16E-04 |
| ZCH18_HUMAN\|Q86VM9_S110_S95 | 1.658777043 | 5.72E-04 | 3.88E-03 |
| RAI14_HUMAN\|Q9P0K7_S281 | 1.658754251 | 7.92E-06 | 5.96E-04 |
| PAWR_HUMAN\|Q96IZ0_S231 | 1.65874417 | 2.35E-05 | 1.11E-03 |
| NFKB1_HUMAN\|P19838_S907 | 1.658461602 | 4.34E-07 | 6.04E-05 |
| ACAP2_HUMAN\|Q15057_S384_T382 | 1.65807767 | 3.61E-08 | 3.92E-06 |
| LARP1_HUMAN\|Q6PKG0_S1040 | 1.657881556 | 1.57E-05 | 5.20E-04 |
| LMNB1_HUMAN\|P20700_T20 | 1.657703726 | 8.26E-04 | 1.39E-02 |
| MYO9A_HUMAN\|B2RTY4_S1733 | 1.657402962 | 2.62E-04 | 2.02E-03 |
| FA83H_HUMAN\|Q6ZRV2_S1025 | 1.657233917 | 4.15E-05 | 6.60E-04 |
| AHNK_HUMAN\|Q09666_S5099 | 1.657104459 | 1.15E-03 | 1.09E-02 |
| B4DKV4_HUMAN\|B4DKV4_S22 | 1.656966938 | 6.90E-03 | 4.99E-02 |
| SYTL1_HUMAN\|Q8IYJ3_S216 | 1.656920663 | 2.12E-03 | 2.16E-02 |
| TRI25_HUMAN\|Q14258_S100 | 1.656737494 | 5.43E-04 | 9.93E-03 |
| ASPP2_HUMAN\|Q13625_S556 | 1.656418332 | 8.13E-05 | 2.85E-03 |
| EIF3H_HUMAN\|O15372_S183 | 1.655943706 | 1.22E-03 | 1.77E-02 |
| EGFR_HUMAN\|P00533_S1204 | 1.655939985 | 8.86E-04 | 7.21E-03 |
| BORG2_HUMAN\|Q9UKI2_S100 | 1.655530119 | 2.40E-04 | 1.80E-03 |
| TAGL2_HUMAN\|P37802_S163 | 1.655454695 | 7.91E-04 | 1.81E-02 |
| RENT1_HUMAN\|Q92900_S1107_T1111 | 1.655251077 | 2.47E-03 | 1.61E-02 |
| OPTN_HUMAN\|Q96CV9_T204 | 1.654671119 | 2.73E-06 | 2.00E-04 |
| K1C16_HUMAN\|P08779_S468 | 1.654660651 | 4.65E-05 | 7.17E-04 |
| 1433Z_HUMAN\|P63104_T232 | 1.654584287 | 7.30E-04 | 6.24E-03 |
| AHNK_HUMAN\|Q09666_S3426 | 1.654007458 | 1.95E-05 | 2.85E-03 |
| AKAP2_HUMAN\|Q9Y2D5_S720 | 1.653862364 | 1.16E-05 | 5.43E-04 |
| UN13B_HUMAN\|O14795_S254 | 1.65340877 | 3.07E-04 | 2.17E-03 |
| AFTIN_HUMAN\|Q6ULP2_S518 | 1.653342386 | 2.76E-05 | 6.35E-04 |

| | | | |
|---|---|---|---|
| FOSL1_HUMAN\|P15407_S265 | 1.65289334 | 2.81E-03 | 3.07E-02 |
| BIG1_HUMAN\|Q9Y6D6_S397 | 1.652822767 | 1.98E-05 | 7.80E-04 |
| RGS3_HUMAN\|P49796_S1044 | 1.652728451 | 1.63E-05 | 3.27E-04 |
| SVIL_HUMAN\|O95425_T852 | 1.652632215 | 3.41E-03 | 3.05E-02 |
| IF4B_HUMAN\|P23588_S489_S497_S498 | 1.65257499 | 1.53E-05 | 8.11E-04 |
| MTUS1_HUMAN\|Q9ULD2_T1204 | 1.652558528 | 3.69E-05 | 4.08E-04 |
| FLVC1_HUMAN\|Q9Y5Y0_S536 | 1.652544584 | 2.59E-04 | 3.44E-03 |
| EPIPL_HUMAN\|P58107_S2716 | 1.65236209 | 6.71E-07 | 1.90E-05 |
| K1C14_HUMAN\|P02533_S54_Y46 | 1.652261139 | 3.90E-04 | 7.68E-03 |
| IPP2M_HUMAN\|Q6NXS1_S122 | 1.652213012 | 9.85E-05 | 1.70E-03 |
| AHNK_HUMAN\|Q09666_S5739 | 1.652024582 | 4.06E-04 | 6.08E-03 |
| AHNK_HUMAN\|Q09666_S212 | 1.65188039 | 2.75E-04 | 5.89E-03 |
| PLEC_HUMAN\|Q15149_S4384_S4386_S4389 | 1.651500326 | 1.25E-02 | 4.83E-02 |
| PININ_HUMAN\|Q9H307_S66 | 1.651435628 | 2.03E-04 | 1.57E-03 |
| DKC1_HUMAN\|O60832_S485_S494 | 1.65129159 | 4.27E-06 | 3.40E-04 |
| K1C14_HUMAN\|P02533_S435_S438 | 1.650577624 | 5.01E-04 | 7.12E-03 |
| CNKR1_HUMAN\|Q969H4_S314 | 1.650437708 | 1.24E-05 | 2.71E-04 |
| TM237_HUMAN\|Q96Q45_S62 | 1.650039155 | 8.78E-03 | 4.83E-02 |
| MISP_HUMAN\|Q8IVT2_S395_S397 | 1.649882158 | 1.78E-04 | 4.30E-03 |
| KMT2A_HUMAN\|Q03164_T3028 | 1.649789318 | 5.77E-03 | 4.47E-02 |
| BAIP2_HUMAN\|Q9UQB8-4_T520 | 1.649713487 | 2.10E-04 | 4.83E-03 |
| CSCL2_HUMAN\|Q5T3F8_S111_S114 | 1.64956124 | 1.63E-03 | 9.06E-03 |
| SMRC1_HUMAN\|Q92922_S328_S330 | 1.649338856 | 6.53E-05 | 1.22E-03 |
| K2C5_HUMAN\|P13647_S28 | 1.649269344 | 1.10E-03 | 8.57E-03 |
| RAE1_HUMAN\|P24386_S651 | 1.64917641 | 7.93E-06 | 1.25E-04 |
| EF2K_HUMAN\|O00418_S470 | 1.648756141 | 5.63E-05 | 2.06E-03 |
| COHA1_HUMAN\|Q9UMD9_S93_T97 | 1.648361248 | 9.72E-04 | 1.43E-02 |
| CNKR1_HUMAN\|Q969H4_S305_S314 | 1.648097186 | 2.51E-04 | 5.60E-03 |
| SMRCD_HUMAN\|Q9H4L7_S96 | 1.647856979 | 4.14E-05 | 1.53E-03 |
| AHNK_HUMAN\|Q09666_S572 | 1.647459963 | 3.11E-04 | 5.49E-03 |
| FXR2_HUMAN\|P51116_S453 | 1.647372234 | 6.10E-05 | 2.13E-03 |

| | | | |
|---|---|---|---|
| WDR7_HUMAN\|Q9Y4E6_S1153 | 1.646981977 | 3.58E-03 | 2.10E-02 |
| 3BP1_HUMAN\|Q9Y3L3_T601 | 1.646661517 | 2.14E-03 | 2.18E-02 |
| PR38B_HUMAN\|Q5VTL8_S529 | 1.646620633 | 3.99E-05 | 4.57E-03 |
| YAP1_HUMAN\|P46937_T119 | 1.646506616 | 9.40E-04 | 9.06E-03 |
| TREF1_HUMAN\|Q96PN7_S758 | 1.646288619 | 2.64E-03 | 3.06E-02 |
| FOXO3_HUMAN\|O43524_S294 | 1.646130556 | 4.62E-04 | 5.25E-03 |
| K2C5_HUMAN\|P13647_S11_S8 | 1.646058315 | 1.66E-05 | 5.41E-04 |
| TB10A_HUMAN\|Q9BXI6_S40_S43 | 1.645909377 | 1.85E-03 | 1.54E-02 |
| EHBP1_HUMAN\|Q8NDI1_S428_S436 | 1.645503143 | 1.99E-06 | 4.44E-05 |
| RHG12_HUMAN\|Q8IWW6_S215 | 1.645234632 | 3.64E-05 | 6.09E-04 |
| TOM1_HUMAN\|O60784_S462 | 1.6452236 | 2.45E-04 | 3.30E-03 |
| ACINU_HUMAN\|Q9UKV3_T254 | 1.645219935 | 9.51E-04 | 1.16E-02 |
| ARI1A_HUMAN\|O14497_S696_S698_S702 | 1.644580513 | 2.25E-06 | 2.01E-04 |
| NKTR_HUMAN\|P30414_S463 | 1.644386334 | 1.32E-04 | 3.44E-03 |
| DC1L1_HUMAN\|Q9Y6G9_T512_T513 | 1.644284418 | 1.26E-03 | 1.87E-02 |
| CE170_HUMAN\|Q5SW79_S971 | 1.644015877 | 3.78E-05 | 6.25E-04 |
| CC186_HUMAN\|Q7Z3E2_S726 | 1.643691857 | 3.96E-05 | 4.40E-04 |
| HELZ_HUMAN\|P42694_S1766 | 1.643535559 | 1.19E-03 | 2.35E-02 |
| NKTR_HUMAN\|P30414_S463 | 1.643463044 | 1.34E-04 | 3.88E-03 |
| IWS1_HUMAN\|Q96ST2_S313 | 1.643396025 | 7.15E-03 | 4.02E-02 |
| TARA_HUMAN\|Q9H2D6_S1955 | 1.643279087 | 2.02E-03 | 1.38E-02 |
| SVIL_HUMAN\|O95425_T852 | 1.642774866 | 3.72E-03 | 3.31E-02 |
| ARAF_HUMAN\|P10398_S157 | 1.64252546 | 1.32E-05 | 4.62E-04 |
| AFAD_HUMAN\|P55196_S1696 | 1.642522314 | 7.64E-05 | 1.72E-03 |
| K1C15_HUMAN\|P19012_S34 | 1.642290069 | 3.25E-03 | 2.92E-02 |
| AFTIN_HUMAN\|Q6ULP2_S518 | 1.642198202 | 3.29E-05 | 1.10E-03 |
| ZCH18_HUMAN\|Q86VM9_T796 | 1.642146925 | 5.81E-04 | 8.70E-03 |
| PKHG3_HUMAN\|A1L390_S1079 | 1.64166955 | 1.09E-04 | 1.03E-03 |
| K1C17_HUMAN\|Q04695_S11 | 1.64149458 | 9.22E-05 | 1.95E-03 |
| NGAP_HUMAN\|Q9UJF2_S954 | 1.641446159 | 9.90E-05 | 3.19E-03 |
| MAP1B_HUMAN\|P46821_S1797_Y1796 | 1.640645854 | 4.22E-04 | 8.58E-03 |

| | | | |
|---|---|---|---|
| SPTB2_HUMAN\|Q01082-3_S14_S8 | 1.64036765 | 2.82E-05 | 9.93E-04 |
| AT11C_HUMAN\|Q8NB49_T1124 | 1.640358424 | 4.56E-05 | 9.13E-04 |
| LAP2A_HUMAN\|P42166_T160 | 1.640280047 | 1.56E-05 | 5.20E-04 |
| SRPK2_HUMAN\|P78362_S497_T492 | 1.639944565 | 2.48E-05 | 2.98E-04 |
| SRC8_HUMAN\|Q14247_S405_S418_T401 | 1.639775056 | 2.42E-03 | 1.87E-02 |
| AHNK2_HUMAN\|Q8IVF2_S1583 | 1.639688956 | 7.83E-04 | 2.48E-02 |
| K1C14_HUMAN\|P02533_S432_S433_S437 | 1.63959404 | 7.16E-04 | 6.16E-03 |
| YBOX3_HUMAN\|P16989_S369_T366 | 1.639394166 | 2.11E-05 | 8.07E-04 |
| NIBL1_HUMAN\|Q96TA1_S691 | 1.639279053 | 1.17E-06 | 1.67E-04 |
| NRK_HUMAN\|Q7Z2Y5_S854_Y858 | 1.63910533 | 2.05E-05 | 5.16E-04 |
| SF3B2_HUMAN\|Q13435_S309 | 1.63885422 | 2.61E-03 | 1.91E-02 |
| RB3GP_HUMAN\|Q15042_S537 | 1.638699702 | 6.14E-06 | 1.02E-04 |
| K1C17_HUMAN\|Q04695_S28 | 1.638595835 | 3.84E-04 | 1.61E-02 |
| HNRH3_HUMAN\|P31942_S298 | 1.638536 | 5.83E-05 | 1.12E-03 |
| BCL10_HUMAN\|O95999_S138 | 1.638501967 | 1.14E-04 | 3.49E-03 |
| BAIP2_HUMAN\|Q9UQB8_Y491 | 1.638258146 | 1.09E-02 | 4.83E-02 |
| CE170_HUMAN\|Q5SW79_S630_S636 | 1.638144844 | 1.12E-05 | 4.32E-04 |
| KLC4_HUMAN\|Q9NSK0_T612 | 1.637909823 | 2.83E-03 | 1.76E-02 |
| SRRM2_HUMAN\|Q9UQ35_S1152 | 1.637691913 | 6.76E-04 | 1.13E-02 |
| TCAL3_HUMAN\|Q969E4_S125 | 1.637375623 | 5.77E-04 | 2.08E-02 |
| TPR_HUMAN\|P12270_S2155 | 1.637367337 | 9.90E-05 | 1.29E-03 |
| SRSF6_HUMAN\|Q13247_S301 | 1.636958417 | 2.04E-04 | 1.69E-03 |
| SMTN_HUMAN\|P53814_T791 | 1.636647132 | 1.94E-03 | 2.04E-02 |
| ZN185_HUMAN\|O15231_S153 | 1.636644923 | 1.59E-05 | 2.46E-03 |
| FAT1_HUMAN\|Q14517_S4473 | 1.636467489 | 1.81E-03 | 1.92E-02 |
| K2C5_HUMAN\|P13647_S21_S26 | 1.636426088 | 3.99E-04 | 1.64E-02 |
| LAD1_HUMAN\|O00515_S38_S39 | 1.636157066 | 2.12E-03 | 2.46E-02 |
| TM1L1_HUMAN\|O75674_S321 | 1.635835594 | 1.09E-03 | 1.65E-02 |
| HELZ2_HUMAN\|Q9BYK8_S1006 | 1.635728909 | 8.38E-04 | 1.39E-02 |
| AHNK2_HUMAN\|Q8IVF2_S457 | 1.635669206 | 3.40E-03 | 3.12E-02 |
| CHSP1_HUMAN\|Q9Y2V2_S52 | 1.635569523 | 5.26E-04 | 7.38E-03 |

| | | | |
|---|---|---|---|
| S23A2_HUMAN\|Q9UGH3_S78_S81 | 1.635442283 | 2.12E-03 | 1.01E-02 |
| HPBP1_HUMAN\|Q9NZL4_S354 | 1.635432087 | 1.01E-04 | 9.21E-04 |
| CE170_HUMAN\|Q5SW79_S636_T628 | 1.635155987 | 2.84E-04 | 4.68E-03 |
| K1C15_HUMAN\|P19012_S34 | 1.635136171 | 3.49E-03 | 3.18E-02 |
| TFE3_HUMAN\|P19532_S548_S556 | 1.635072584 | 1.66E-04 | 1.45E-03 |
| K1C14_HUMAN\|P02533_S12 | 1.634786976 | 2.72E-05 | 8.20E-04 |
| FA76A_HUMAN\|Q8TAV0_S306 | 1.634564824 | 1.63E-05 | 6.69E-04 |
| DC1L2_HUMAN\|O43237_S407 | 1.634404982 | 4.32E-05 | 6.81E-04 |
| LAD1_HUMAN\|O00515_S498 | 1.634314607 | 1.49E-03 | 2.03E-02 |
| IF2P_HUMAN\|O60841_S182_S183_S190 | 1.634174362 | 7.26E-05 | 7.37E-04 |
| TFE3_HUMAN\|P19532_S548 | 1.633949527 | 1.45E-03 | 3.54E-02 |
| NUP53_HUMAN\|Q8NFH5_S105 | 1.633926469 | 2.91E-04 | 4.72E-03 |
| EPIPL_HUMAN\|P58107_S2716 | 1.633902664 | 9.79E-07 | 5.70E-05 |
| CTSL2_HUMAN\|Q05D32_S9 | 1.633691249 | 1.88E-03 | 2.47E-02 |
| ASPP2_HUMAN\|Q13625_S556 | 1.633609649 | 1.13E-04 | 1.01E-03 |
| OSBL3_HUMAN\|Q9H4L5_S437 | 1.633516526 | 4.12E-04 | 4.86E-03 |
| LAD1_HUMAN\|O00515_S498 | 1.632842926 | 1.52E-03 | 1.28E-02 |
| ML12A_HUMAN\|P19105_S19 | 1.632678646 | 2.85E-05 | 9.98E-04 |
| KLC4_HUMAN\|Q9NSK0_S611 | 1.632672035 | 9.44E-05 | 1.98E-03 |
| TBC9B_HUMAN\|Q66K14_S411 | 1.632398339 | 6.32E-04 | 9.19E-03 |
| 1433Z_HUMAN\|P63104_T232 | 1.63230785 | 9.37E-04 | 1.22E-02 |
| CE170_HUMAN\|Q5SW79_T628 | 1.632078189 | 9.10E-03 | 3.72E-02 |
| AHNK_HUMAN\|Q09666_S210_S216 | 1.632044078 | 4.17E-05 | 8.51E-04 |
| PRAG1_HUMAN\|Q86YV5_S907 | 1.631986574 | 2.41E-05 | 4.51E-04 |
| FOSL2_HUMAN\|P15408_S320 | 1.631981994 | 8.58E-03 | 4.64E-02 |
| TMF1_HUMAN\|P82094_S72 | 1.631927667 | 1.26E-05 | 3.57E-04 |
| TACC1_HUMAN\|O75410_S44 | 1.631870099 | 7.69E-05 | 1.72E-03 |
| ARH40_HUMAN\|Q8TER5_S1513 | 1.631582495 | 1.67E-06 | 3.66E-05 |
| SARG_HUMAN\|Q9BW04_S429 | 1.631318428 | 7.01E-04 | 9.27E-03 |
| PAK1_HUMAN\|Q13153_S223 | 1.631287777 | 3.09E-04 | 3.90E-03 |
| NELFE_HUMAN\|P18615_S251 | 1.631253424 | 7.29E-04 | 2.42E-02 |

| | | | |
|---|---|---|---|
| TAB2_HUMAN|Q9NYJ8_S450 | 1.630957505 | 4.21E-06 | 1.74E-04 |
| CRTC3_HUMAN|Q6UUV7_S370 | 1.630854954 | 6.73E-04 | 5.94E-03 |
| MICA3_HUMAN|Q7RTP6_S1636 | 1.630811133 | 1.97E-03 | 4.32E-02 |
| CAVN1_HUMAN|Q6NZI2_T302 | 1.630320234 | 7.22E-06 | 5.67E-04 |
| PKP2_HUMAN|Q99959_S151 | 1.630206186 | 2.77E-05 | 1.21E-03 |
| IFIT3_HUMAN|O14879_S484 | 1.629813633 | 9.58E-03 | 4.97E-02 |
| PARD3_HUMAN|Q8TEW0_S187 | 1.629327522 | 9.56E-03 | 4.38E-02 |
| TMF1_HUMAN|P82094_S72_S77 | 1.629186559 | 1.43E-03 | 1.22E-02 |
| TB182_HUMAN|Q9C0C2_S1029 | 1.62915673 | 1.74E-05 | 8.71E-04 |
| TRI47_HUMAN|Q96LD4_S588 | 1.62915182 | 1.87E-04 | 4.89E-03 |
| DESP_HUMAN|P15924_S166 | 1.62912891 | 3.93E-04 | 2.62E-03 |
| PR38B_HUMAN|Q5VTL8_S529 | 1.629054715 | 5.35E-05 | 1.35E-03 |
| TAGL2_HUMAN|P37802_S163 | 1.629023161 | 1.07E-03 | 9.98E-03 |
| NUP50_HUMAN|Q9UKX7_S221 | 1.628700635 | 1.23E-05 | 4.48E-04 |
| SP100_HUMAN|P23497_S476 | 1.628646729 | 2.84E-04 | 6.44E-03 |
| EF2K_HUMAN|O00418_S470 | 1.627993787 | 7.69E-05 | 1.07E-03 |
| AAK1_HUMAN|Q2M2I8_S21 | 1.627890216 | 2.39E-04 | 4.54E-03 |
| K1C16_HUMAN|P08779_S449 | 1.62760253 | 3.60E-04 | 5.59E-03 |
| TT39B_HUMAN|Q5VTQ0_S120 | 1.627518057 | 1.27E-03 | 1.54E-02 |
| SCAM2_HUMAN|O15127_S320 | 1.627506469 | 1.10E-03 | 1.01E-02 |
| WDR44_HUMAN|Q5JSH3_S561 | 1.627331758 | 2.91E-05 | 1.53E-03 |
| AHNK_HUMAN|Q09666_S216_S220 | 1.626846125 | 1.12E-05 | 2.54E-04 |
| AHNK_HUMAN|Q09666_S5746_S5752 | 1.626762972 | 1.41E-05 | 2.38E-03 |
| IASPP_HUMAN|Q8WUF5_S395 | 1.626762082 | 7.85E-04 | 6.66E-03 |
| KPCD2_HUMAN|Q9BZL6_S214 | 1.626476528 | 3.02E-04 | 1.36E-02 |
| PML_HUMAN|P29590_T867 | 1.625998337 | 6.36E-05 | 9.20E-04 |
| WAC2A_HUMAN|Q641Q2_S284 | 1.625899528 | 8.09E-04 | 1.34E-02 |
| MA7D3_HUMAN|Q8IWC1_S524 | 1.625856622 | 6.14E-03 | 4.61E-02 |
| DKC1_HUMAN|O60832_S485 | 1.625851895 | 5.03E-05 | 1.26E-02 |
| CR025_HUMAN|Q96B23_S145_S147 | 1.625839633 | 1.02E-05 | 4.08E-04 |
| PSF2_HUMAN|Q9Y248_S182 | 1.625735851 | 4.06E-04 | 6.08E-03 |

| | | | |
|---|---|---|---|
| EFNB1_HUMAN\|P98172_S281 | 1.625599026 | 1.64E-04 | 3.42E-03 |
| CE170_HUMAN\|Q5SW79_S630_S636 | 1.625482997 | 1.41E-05 | 2.94E-04 |
| LAD1_HUMAN\|O00515_S301 | 1.625282204 | 4.08E-04 | 4.34E-02 |
| SVIL_HUMAN\|O95425_T852 | 1.625199279 | 4.37E-03 | 2.44E-02 |
| PML_HUMAN\|P29590_S518_S527 | 1.625095905 | 1.07E-03 | 2.92E-02 |
| RFIP3_HUMAN\|O75154_S52 | 1.624874099 | 1.36E-04 | 1.15E-03 |
| PLEC_HUMAN\|Q15149_S4389_S4396 | 1.624775967 | 8.99E-04 | 5.58E-03 |
| SNUT1_HUMAN\|O43290_T430 | 1.624675198 | 3.94E-03 | 3.73E-02 |
| CCDC9_HUMAN\|Q9Y3X0_T389 | 1.624368446 | 1.41E-06 | 1.47E-04 |
| IWS1_HUMAN\|Q96ST2_S300_S313_S315 | 1.624182375 | 6.45E-03 | 3.90E-02 |
| RGPS2_HUMAN\|Q86X27_S296 | 1.623932033 | 9.38E-03 | 4.89E-02 |
| SRPK2_HUMAN\|P78362_S494_T498 | 1.623791949 | 1.98E-03 | 2.55E-02 |
| TB10A_HUMAN\|Q9BXI6_S40_S43 | 1.62360796 | 2.32E-03 | 1.74E-02 |
| K1522_HUMAN\|Q9P206_S404_S409 | 1.623397601 | 2.17E-05 | 4.14E-04 |
| NDK8_HUMAN\|O60361_S105 | 1.623328486 | 5.96E-05 | 1.47E-03 |
| ACAP2_HUMAN\|Q15057_T382 | 1.623142781 | 1.24E-06 | 6.51E-04 |
| BORG2_HUMAN\|Q9UKI2_S100 | 1.62287684 | 3.75E-04 | 7.84E-03 |
| AF1L2_HUMAN\|Q8N4X5_S344 | 1.622667388 | 7.37E-05 | 1.91E-03 |
| PACS1_HUMAN\|Q6VY07_T504 | 1.622505103 | 3.90E-04 | 5.90E-03 |
| IP3KB_HUMAN\|P27987_S43 | 1.622465519 | 2.88E-05 | 1.29E-03 |
| MISP_HUMAN\|Q8IVT2_S155 | 1.622367781 | 9.84E-04 | 2.85E-02 |
| KLC1_HUMAN\|Q07866-4_S631 | 1.622327463 | 4.88E-06 | 3.05E-04 |
| INADL_HUMAN\|Q8NI35_S645 | 1.621994541 | 1.58E-05 | 6.61E-04 |
| TOM1_HUMAN\|O60784_S462 | 1.621797127 | 3.31E-04 | 7.25E-03 |
| TGS1_HUMAN\|Q96RS0_S85_S89 | 1.621786671 | 1.53E-03 | 1.70E-02 |
| FKBP5_HUMAN\|Q13451_T443 | 1.621729352 | 2.74E-03 | 2.89E-02 |
| NDK8_HUMAN\|O60361_S105 | 1.621675213 | 6.12E-05 | 1.68E-03 |
| K1C14_HUMAN\|P02533_S13 | 1.621630863 | 1.51E-05 | 5.14E-04 |
| K2C5_HUMAN\|P13647_S28 | 1.621588264 | 1.51E-03 | 3.85E-02 |
| ROA1_HUMAN\|P09651_S361_S364 | 1.621484137 | 1.07E-03 | 1.33E-02 |
| K2C5_HUMAN\|P13647_S62 | 1.621479767 | 1.09E-04 | 2.50E-03 |

| | | | |
|---|---|---|---|
| K1C17_HUMAN\|Q04695_S24 | 1.621398451 | 1.01E-04 | 2.41E-03 |
| COF1_HUMAN\|P23528_S24 | 1.621289604 | 1.39E-03 | 7.91E-03 |
| RBL2_HUMAN\|Q08999_S966 | 1.621173789 | 3.76E-03 | 3.25E-02 |
| CTSL2_HUMAN\|Q05D32_S9 | 1.62071825 | 2.16E-03 | 1.73E-02 |
| NIBL1_HUMAN\|Q96TA1_S692 | 1.620703353 | 3.76E-05 | 4.14E-04 |
| RAI14_HUMAN\|Q9P0K7_S290 | 1.620681103 | 1.23E-03 | 1.73E-02 |
| AKTS1_HUMAN\|Q96B36_T246 | 1.620536111 | 6.74E-04 | 5.94E-03 |
| MYPT1_HUMAN\|O14974_S507 | 1.620434362 | 1.11E-04 | 7.80E-03 |
| CCAR2_HUMAN\|Q8N163_S678 | 1.620423911 | 9.63E-03 | 4.99E-02 |
| RAI14_HUMAN\|Q9P0K7_S281 | 1.620143936 | 1.58E-05 | 2.10E-04 |
| LMNB2_HUMAN\|Q03252_S420_S424 | 1.620068698 | 9.56E-04 | 1.23E-02 |
| ANXA2_HUMAN\|P07355_S127 | 1.619922357 | 5.49E-03 | 4.31E-02 |
| MAP1B_HUMAN\|P46821_S1501 | 1.619856516 | 6.59E-06 | 2.27E-04 |
| NDRG1_HUMAN\|Q92597_S333_S336_T328 | 1.619771374 | 1.10E-05 | 2.50E-04 |
| DCAF6_HUMAN\|Q58WW2_S657 | 1.619479861 | 3.16E-04 | 2.23E-03 |
| SLK_HUMAN\|Q9H2G2_S779 | 1.619398036 | 4.21E-03 | 3.50E-02 |
| K1C14_HUMAN\|P02533_S17 | 1.619377698 | 2.30E-03 | 1.52E-02 |
| PSF2_HUMAN\|Q9Y248_S182 | 1.61937185 | 4.44E-04 | 4.28E-03 |
| FA83B_HUMAN\|Q5T0W9_S731 | 1.618976255 | 4.38E-05 | 1.76E-03 |
| LYAR_HUMAN\|Q9NX58_S276 | 1.618973399 | 6.45E-06 | 2.26E-04 |
| ARHG5_HUMAN\|Q12774_S445_S450 | 1.618936612 | 2.67E-06 | 5.33E-05 |
| MED24_HUMAN\|O75448_S862_S873 | 1.618901326 | 3.98E-03 | 1.88E-02 |
| MA7D1_HUMAN\|Q3KQU3_S452 | 1.618728736 | 3.00E-04 | 3.83E-03 |
| XRCC6_HUMAN\|P12956_S477 | 1.618471565 | 1.31E-03 | 2.55E-02 |
| NDRG1_HUMAN\|Q92597_S326_S330_S333 | 1.61802647 | 1.97E-04 | 2.27E-03 |
| FLVC1_HUMAN\|Q9Y5Y0_S536 | 1.61783065 | 4.22E-04 | 6.86E-03 |
| COHA1_HUMAN\|Q9UMD9_S85_S93_T97 | 1.617818872 | 1.71E-03 | 4.05E-02 |
| COBL1_HUMAN\|Q53SF7_S1146 | 1.6177125 | 7.72E-05 | 1.33E-03 |
| K2C5_HUMAN\|P13647_S26 | 1.617571046 | 9.12E-05 | 1.21E-03 |
| MED24_HUMAN\|O75448_S862_S873 | 1.617566088 | 4.03E-03 | 1.69E-02 |
| IWS1_HUMAN\|Q96ST2_T53 | 1.617530259 | 4.67E-03 | 2.96E-02 |

| | | | |
|---|---|---|---|
| LAP2A_HUMAN\|P42166_T160 | 1.617409234 | 2.28E-05 | 8.34E-04 |
| COHA1_HUMAN\|Q9UMD9_S130 | 1.617297811 | 8.78E-03 | 4.10E-02 |
| S6A15_HUMAN\|Q9H2J7_S699 | 1.617202447 | 2.46E-03 | 2.40E-02 |
| K1C17_HUMAN\|Q04695_S11_S13 | 1.617099604 | 3.12E-04 | 1.40E-02 |
| P3C2A_HUMAN\|O00443_S259 | 1.616787651 | 3.96E-05 | 1.52E-03 |
| DCBD2_HUMAN\|Q96PD2_S726_S727 | 1.616745868 | 2.79E-03 | 2.62E-02 |
| CTSL2_HUMAN\|Q05D32_S9 | 1.616716648 | 2.25E-03 | 2.77E-02 |
| K1C17_HUMAN\|Q04695_S17 | 1.616714488 | 2.11E-03 | 1.43E-02 |
| NOP2_HUMAN\|P46087_T605 | 1.616605675 | 7.24E-05 | 2.67E-03 |
| ASPP2_HUMAN\|Q13625_S556 | 1.616462959 | 1.46E-04 | 2.20E-03 |
| ACINU_HUMAN\|Q9UKV3_S243 | 1.616399274 | 1.53E-05 | 8.07E-04 |
| TM1L1_HUMAN\|O75674_S321_S323 | 1.616349192 | 3.67E-04 | 2.50E-03 |
| SRSF4_HUMAN\|Q08170_S460 | 1.616045136 | 6.00E-04 | 4.02E-03 |
| ZO1_HUMAN\|Q07157_T868_Y843 | 1.615887736 | 3.60E-03 | 3.89E-02 |
| CSKI2_HUMAN\|Q8WXE0_S406 | 1.615820587 | 1.65E-06 | 5.91E-05 |
| ATN1_HUMAN\|P54259_S77_S79 | 1.615786272 | 8.29E-04 | 6.89E-03 |
| EF2K_HUMAN\|O00418_S470 | 1.615666882 | 9.28E-05 | 1.96E-03 |
| RBM7_HUMAN\|Q9Y580_S136 | 1.6156243 | 4.12E-04 | 8.46E-03 |
| TGS1_HUMAN\|Q96RS0_S55 | 1.615368949 | 1.20E-04 | 1.93E-03 |
| MON1A_HUMAN\|Q86VX9_S56 | 1.615319798 | 1.52E-05 | 2.43E-03 |
| CLIP1_HUMAN\|P30622_S200 | 1.615258616 | 1.16E-03 | 1.37E-02 |
| NIBL1_HUMAN\|Q96TA1_S641 | 1.614998794 | 2.15E-03 | 2.67E-02 |
| YAP1_HUMAN\|P46937_T110_T119 | 1.614843743 | 6.83E-04 | 5.98E-03 |
| NU214_HUMAN\|P35658_S986_T987 | 1.614716949 | 4.29E-03 | 2.93E-02 |
| ZFP91_HUMAN\|Q96JP5_S83 | 1.614714079 | 5.85E-06 | 1.53E-04 |
| EPIPL_HUMAN\|P58107_S2716 | 1.614494304 | 1.47E-06 | 6.79E-05 |
| ENSA_HUMAN\|O43768_S109 | 1.614464356 | 2.62E-04 | 4.87E-03 |
| MET_HUMAN\|P08581_S997_T993 | 1.614227808 | 9.17E-04 | 1.47E-02 |
| EMAL3_HUMAN\|Q32P44_S889 | 1.614216827 | 5.90E-03 | 3.67E-02 |
| MISP_HUMAN\|Q8IVT2_S543 | 1.614212286 | 1.12E-04 | 3.14E-03 |
| LIMA1_HUMAN\|Q9UHB6_S507 | 1.614092773 | 8.47E-04 | 1.29E-02 |

| | | | |
|---|---|---|---|
| AHNK_HUMAN\|Q09666_T4564 | 1.613773276 | 2.65E-04 | 4.48E-03 |
| STMN1_HUMAN\|P16949_S16 | 1.613643597 | 1.60E-04 | 4.48E-03 |
| COBL1_HUMAN\|Q53SF7_T298 | 1.613603877 | 2.21E-04 | 3.96E-03 |
| PLEC_HUMAN\|Q15149_Y4615 | 1.613506463 | 9.84E-05 | 1.29E-03 |
| KDM4A_HUMAN\|O75164_S523 | 1.613474503 | 4.22E-03 | 3.89E-02 |
| EXPH5_HUMAN\|Q8NEV8_S341 | 1.613119891 | 7.18E-05 | 1.68E-03 |
| ATN1_HUMAN\|P54259_S77_S79 | 1.613099721 | 8.58E-04 | 1.07E-02 |
| TPC12_HUMAN\|Q8WVT3_S184 | 1.613063374 | 8.22E-04 | 1.86E-02 |
| AHNK_HUMAN\|Q09666_S135 | 1.613046845 | 9.08E-04 | 8.84E-03 |
| MON1A_HUMAN\|Q86VX9_S95 | 1.612974957 | 2.54E-05 | 3.07E-04 |
| FUBP2_HUMAN\|Q92945_T195 | 1.612894466 | 8.14E-03 | 4.60E-02 |
| DESP_HUMAN\|P15924_S166 | 1.61286866 | 4.88E-04 | 9.28E-03 |
| RGL2_HUMAN\|O15211_S619 | 1.612773949 | 1.05E-04 | 3.03E-03 |
| WDR13_HUMAN\|Q9H1Z4_S70_S74 | 1.612760622 | 8.17E-05 | 1.12E-03 |
| PININ_HUMAN\|Q9H307_S66 | 1.61271483 | 3.40E-04 | 7.25E-03 |
| UFD1_HUMAN\|Q92890_S299 | 1.612700704 | 2.92E-03 | 2.08E-02 |
| MPLKI_HUMAN\|Q8TAP9_S80 | 1.612452709 | 6.49E-03 | 4.77E-02 |
| ZO2_HUMAN\|Q9UDY2_T1027 | 1.61237697 | 4.36E-05 | 6.85E-04 |
| PPIG_HUMAN\|Q13427_S696 | 1.61230178 | 1.29E-03 | 9.70E-03 |
| IASPP_HUMAN\|Q8WUF5_S526 | 1.612019123 | 8.47E-03 | 3.52E-02 |
| REQU_HUMAN\|Q92785_S142 | 1.611965665 | 1.76E-04 | 2.06E-03 |
| SGTB_HUMAN\|Q96EQ0_S297 | 1.611920835 | 4.93E-05 | 9.69E-04 |
| CLIP4_HUMAN\|Q8N3C7_S610 | 1.611798907 | 3.06E-04 | 2.28E-03 |
| BORG4_HUMAN\|Q9H3Q1_S73 | 1.611750682 | 2.37E-03 | 2.32E-02 |
| FA83B_HUMAN\|Q5T0W9_S802 | 1.61173579 | 8.61E-05 | 1.43E-03 |
| BCL10_HUMAN\|O95999_S138 | 1.611503138 | 1.69E-04 | 1.36E-03 |
| ACSA_HUMAN\|Q9NR19_S267 | 1.611024911 | 1.15E-04 | 3.22E-03 |
| SCRIB_HUMAN\|Q14160_S1348 | 1.610968456 | 1.55E-03 | 1.72E-02 |
| PARD3_HUMAN\|Q8TEW0_S225 | 1.610916987 | 1.65E-04 | 3.42E-03 |
| DCA10_HUMAN\|Q5QP82_S350 | 1.610779414 | 1.98E-04 | 5.07E-03 |
| DCBD2_HUMAN\|Q96PD2_S724 | 1.61074325 | 3.74E-03 | 3.58E-02 |

| | | | |
|---|---|---|---|
| PKP3_HUMAN\|Q9Y446_S180 | 1.610670384 | 2.49E-04 | 1.20E-02 |
| SNTB2_HUMAN\|Q13425_S222 | 1.610476887 | 4.71E-03 | 3.13E-02 |
| TTP_HUMAN\|P26651_S186 | 1.610447902 | 4.22E-03 | 3.89E-02 |
| SF3B2_HUMAN\|Q13435_S303_S309 | 1.610305713 | 7.47E-03 | 4.33E-02 |
| DCBD2_HUMAN\|Q96PD2_S726_S727 | 1.610216286 | 3.00E-03 | 1.85E-02 |
| DC1I2_HUMAN\|Q13409_S101_S104 | 1.61020371 | 2.46E-03 | 1.81E-02 |
| C2CD5_HUMAN\|Q86YS7-2_S855 | 1.610092691 | 9.98E-05 | 7.30E-03 |
| DPYL2_HUMAN\|Q16555_T509 | 1.609808009 | 1.20E-06 | 4.66E-05 |
| NIPA_HUMAN\|Q86WB0_S408_S409 | 1.609791114 | 2.23E-06 | 7.10E-04 |
| K2C8_HUMAN\|P05787_S24 | 1.609618641 | 7.83E-04 | 1.25E-02 |
| ZO1_HUMAN\|Q07157_T868_Y843 | 1.609589626 | 3.84E-03 | 3.93E-02 |
| COHA1_HUMAN\|Q9UMD9_S79 | 1.609458457 | 3.46E-03 | 3.15E-02 |
| COF1_HUMAN\|P23528_S24 | 1.609353726 | 1.59E-03 | 8.01E-03 |
| MTUS1_HUMAN\|Q9ULD2_T1204 | 1.609304299 | 7.51E-05 | 7.52E-04 |
| HPS5_HUMAN\|Q9UPZ3_S534 | 1.608774835 | 1.25E-04 | 2.79E-03 |
| UBP2L_HUMAN\|Q14157_S116 | 1.608595115 | 6.82E-03 | 3.37E-02 |
| KC1D_HUMAN\|P48730_S328 | 1.608558701 | 1.94E-03 | 2.01E-02 |
| K1C14_HUMAN\|P02533_S44_S53_S54 | 1.608451575 | 6.54E-05 | 2.23E-03 |
| S23A2_HUMAN\|Q9UGH3_S78_S81 | 1.608316672 | 2.84E-03 | 1.44E-02 |
| ZCH18_HUMAN\|Q86VM9_S532_S534 | 1.60812672 | 8.63E-07 | 3.68E-05 |
| LAR4B_HUMAN\|Q92615_S731_S736 | 1.608098375 | 1.89E-04 | 3.52E-03 |
| K1C14_HUMAN\|P02533_S432_S433_S437 | 1.607986561 | 1.06E-03 | 1.50E-02 |
| ATX2L_HUMAN\|Q8WWM7_S684_T681 | 1.607882863 | 2.68E-03 | 2.54E-02 |
| KPCD1_HUMAN\|Q15139_S742 | 1.60783677 | 1.68E-03 | 1.43E-02 |
| SMRCD_HUMAN\|Q9H4L7_S96 | 1.607630474 | 8.15E-05 | 1.46E-03 |
| ARG28_HUMAN\|Q8N1W1_S476_S486 | 1.607429959 | 2.54E-04 | 3.40E-02 |
| LMNA_HUMAN\|P02545_S619 | 1.607388193 | 5.88E-04 | 5.39E-03 |
| TX264_HUMAN\|Q9Y6I9_S244 | 1.607373056 | 2.18E-03 | 1.74E-02 |
| K1C14_HUMAN\|P02533_S435 | 1.607363835 | 6.24E-04 | 5.65E-03 |
| ABI2_HUMAN\|Q9NYB9_Y371 | 1.607337171 | 1.98E-05 | 7.80E-04 |
| FOSL2_HUMAN\|P15408_S19 | 1.606551447 | 4.64E-03 | 3.09E-02 |

| | | | |
|---|---|---|---|
| HMGN1_HUMAN\|P05114_S7 | 1.606385777 | 5.13E-04 | 5.69E-03 |
| K2C5_HUMAN\|P13647_S18 | 1.606362807 | 1.40E-04 | 1.73E-03 |
| PRAG1_HUMAN\|Q86YV5_S745 | 1.606168559 | 2.10E-04 | 1.62E-03 |
| MA7D3_HUMAN\|Q8IWC1_S524 | 1.60589894 | 7.33E-03 | 3.55E-02 |
| K1C18_HUMAN\|P05783_S7 | 1.605891736 | 1.53E-03 | 1.70E-02 |
| SPAS2_HUMAN\|Q86XZ4_S538 | 1.605666235 | 7.59E-03 | 4.20E-02 |
| BRD4_HUMAN\|O60885_S1117 | 1.605589823 | 2.34E-03 | 2.38E-02 |
| SRRM2_HUMAN\|Q9UQ35_S2272 | 1.605406072 | 2.93E-06 | 1.21E-04 |
| KI21A_HUMAN\|Q7Z4S6_S855 | 1.605287974 | 7.92E-05 | 1.35E-03 |
| P63_HUMAN\|Q9H3D4_S452_S463 | 1.605185126 | 1.18E-03 | 1.07E-02 |
| CR025_HUMAN\|Q96B23_S143_S145 | 1.604952719 | 2.37E-05 | 5.70E-04 |
| ENSA_HUMAN\|O43768_S109 | 1.604942468 | 3.00E-04 | 3.83E-03 |
| PARD3_HUMAN\|Q8TEW0_S225 | 1.604861489 | 1.81E-04 | 2.11E-03 |
| MON1A_HUMAN\|Q86VX9_S56 | 1.604602085 | 1.85E-05 | 4.86E-04 |
| FOXO3_HUMAN\|O43524_S413 | 1.604469806 | 4.49E-03 | 3.01E-02 |
| VIME_HUMAN\|P08670_S459 | 1.604433532 | 2.43E-05 | 7.35E-03 |
| SHOT1_HUMAN\|A0MZ66_T537 | 1.604396969 | 2.57E-03 | 3.01E-02 |
| OSBL8_HUMAN\|Q9BZF1_S807_S808_S810 | 1.604396126 | 7.16E-04 | 4.66E-03 |
| ARI1A_HUMAN\|O14497_S696_S698_S702 | 1.604055509 | 5.14E-06 | 1.96E-04 |
| CXA1_HUMAN\|P17302_S306 | 1.603905155 | 1.63E-04 | 5.79E-03 |
| STMN1_HUMAN\|P16949_S25 | 1.603681839 | 2.42E-05 | 2.91E-04 |
| UTP18_HUMAN\|Q9Y5J1_S205 | 1.603443908 | 7.68E-03 | 4.24E-02 |
| COBL1_HUMAN\|Q53SF7_S1146 | 1.603326605 | 9.62E-05 | 9.35E-04 |
| MRE11_HUMAN\|P49959_S649 | 1.603253773 | 1.13E-03 | 1.40E-02 |
| CAAP1_HUMAN\|Q9H8G2_S92 | 1.603090042 | 1.86E-05 | 2.37E-04 |
| VIP2_HUMAN\|O43314_S1006 | 1.602950271 | 1.20E-05 | 7.76E-04 |
| UTP18_HUMAN\|Q9Y5J1_S205 | 1.602896847 | 7.72E-03 | 4.44E-02 |
| CCDC9_HUMAN\|Q9Y3X0_S386 | 1.602839938 | 2.10E-06 | 9.96E-05 |
| NIBAN_HUMAN\|Q9BZQ8_S926 | 1.602810737 | 3.43E-05 | 1.40E-03 |
| ERF_HUMAN\|P50548_T526 | 1.602771548 | 4.44E-04 | 7.09E-03 |
| NDRG2_HUMAN\|Q9UN36_S332_T330 | 1.602274327 | 5.84E-05 | 8.77E-04 |

| | | | |
|---|---|---|---|
| AHNK2_HUMAN\|Q8IVF2_S1583 | 1.602085321 | 1.23E-03 | 1.09E-02 |
| FKBP5_HUMAN\|Q13451_T443 | 1.602012275 | 3.34E-03 | 2.01E-02 |
| ERBB2_HUMAN\|P04626_S998 | 1.601791892 | 3.12E-03 | 3.32E-02 |
| SERF2_HUMAN\|P84101_S21 | 1.60175512 | 3.45E-04 | 1.51E-02 |
| KLC3_HUMAN\|Q6P597_S495_S502 | 1.601684653 | 2.36E-03 | 2.32E-02 |
| PCM1_HUMAN\|Q15154_S872 | 1.601537579 | 2.26E-05 | 4.26E-04 |
| PALLD_HUMAN\|Q8WX93_S1104 | 1.601467706 | 4.39E-05 | 9.02E-04 |
| TNIK_HUMAN\|Q9UKE5_S707 | 1.601325126 | 4.65E-03 | 3.75E-02 |
| PAK1_HUMAN\|Q13153_S223 | 1.601219536 | 4.55E-04 | 8.55E-03 |
| CBX3_HUMAN\|Q13185_S93 | 1.601207211 | 3.56E-03 | 3.86E-02 |
| NIBL1_HUMAN\|Q96TA1_S641_S646 | 1.600612767 | 1.66E-03 | 2.13E-02 |
| CSN1_HUMAN\|Q13098_S483_T479 | 1.60052293 | 4.13E-05 | 1.12E-03 |
| CNKR1_HUMAN\|Q969H4_S305_S314 | 1.600497882 | 4.74E-04 | 6.81E-03 |
| AHNK_HUMAN\|Q09666_S210 | 1.600093982 | 5.45E-06 | 3.65E-04 |
| PKP2_HUMAN\|Q99959_S151 | 1.600002891 | 4.52E-05 | 4.84E-04 |
| CBLB_HUMAN\|Q13191_S480_S484 | 1.59999033 | 1.51E-05 | 2.43E-03 |
| NBN_HUMAN\|O60934_S343 | 1.599943701 | 1.21E-04 | 1.55E-03 |
| TBCD5_HUMAN\|Q92609_S522 | 1.599942597 | 1.88E-04 | 4.82E-03 |
| PKP3_HUMAN\|Q9Y446_S183 | 1.599868661 | 3.85E-04 | 3.86E-03 |
| DKC1_HUMAN\|O60832_S485 | 1.599837463 | 7.64E-05 | 6.27E-03 |
| PHF2_HUMAN\|O75151_S1058 | 1.59970292 | 4.11E-04 | 1.15E-02 |
| K1C15_HUMAN\|P19012_S48 | 1.599631293 | 8.50E-03 | 4.00E-02 |
| RL1D1_HUMAN\|O76021_T465 | 1.599615262 | 2.24E-03 | 1.05E-02 |
| DC1L1_HUMAN\|Q9Y6G9_S516_T515 | 1.599588676 | 6.34E-05 | 2.47E-03 |
| LMNB2_HUMAN\|Q03252_S420_S424 | 1.599347808 | 1.24E-03 | 1.44E-02 |
| COHA1_HUMAN\|Q9UMD9_S85_S93_T97 | 1.599346145 | 2.10E-03 | 2.18E-02 |
| K1C14_HUMAN\|P02533_S33 | 1.599198548 | 1.70E-04 | 2.01E-03 |
| PALLD_HUMAN\|Q8WX93_S726 | 1.598905409 | 1.45E-03 | 1.66E-02 |
| GYS1_HUMAN\|P13807_S727_S731 | 1.598524321 | 6.43E-04 | 1.11E-02 |
| NEB2_HUMAN\|Q96SB3_S100 | 1.598086336 | 2.00E-03 | 2.55E-02 |
| SRPK2_HUMAN\|P78362_S496_T498 | 1.598040371 | 1.45E-04 | 4.19E-03 |

| | | | |
|---|---|---|---|
| AAPK1_HUMAN\|Q13131_S508 | 1.597862122 | 8.98E-07 | 5.35E-05 |
| NUP53_HUMAN\|Q8NFH5_S105 | 1.597665172 | 4.69E-04 | 7.30E-03 |
| PCM1_HUMAN\|Q15154_S872 | 1.597564771 | 2.42E-05 | 2.57E-03 |
| DKC1_HUMAN\|O60832_S485 | 1.597506837 | 7.93E-05 | 2.81E-03 |
| AKAP1_HUMAN\|Q92667_S151 | 1.597446702 | 1.15E-03 | 1.73E-02 |
| K2C5_HUMAN\|P13647_S64 | 1.597393776 | 1.48E-03 | 1.88E-02 |
| AKAP8_HUMAN\|O43823_S323 | 1.597387492 | 2.93E-03 | 4.38E-02 |
| FA83H_HUMAN\|Q6ZRV2_S1098 | 1.597156129 | 5.33E-05 | 1.35E-03 |
| COF1_HUMAN\|P23528_S24 | 1.596659075 | 1.84E-03 | 1.45E-02 |
| TAGL2_HUMAN\|P37802_S163 | 1.596488236 | 1.57E-03 | 8.78E-03 |
| K1C14_HUMAN\|P02533_S17 | 1.596416486 | 2.96E-03 | 2.82E-02 |
| ANS1A_HUMAN\|Q92625_S649 | 1.596371872 | 3.51E-06 | 2.88E-04 |
| GIT1_HUMAN\|Q9Y2X7_S385_T392 | 1.596351727 | 7.91E-03 | 3.34E-02 |
| K2C6B_HUMAN\|P04259_S546 | 1.5962548 | 7.39E-05 | 1.71E-03 |
| HMGA2_HUMAN\|P52926_T100 | 1.59579766 | 3.07E-03 | 1.53E-02 |
| K1C14_HUMAN\|P02533_S429_S432_S438 | 1.595468584 | 5.64E-03 | 4.32E-02 |
| FOXO3_HUMAN\|O43524_S294 | 1.595466441 | 8.75E-04 | 4.94E-03 |
| TFE3_HUMAN\|P19532_S548 | 1.595239597 | 2.21E-03 | 1.68E-02 |
| MMTA2_HUMAN\|Q9BU76_S216_S220 | 1.595203579 | 1.44E-04 | 3.80E-03 |
| PRC2C_HUMAN\|Q9Y520_S2694_T2691 | 1.5950504 | 1.01E-03 | 2.95E-02 |
| DPYL3_HUMAN\|Q14195_S568 | 1.594440249 | 3.35E-03 | 2.33E-02 |
| YTDC2_HUMAN\|Q9H6S0_S1202 | 1.594215597 | 5.00E-05 | 9.85E-04 |
| ASPP2_HUMAN\|Q13625_S556 | 1.594204457 | 2.04E-04 | 1.69E-03 |
| K1C14_HUMAN\|P02533_S437_S438 | 1.594078829 | 5.65E-04 | 7.83E-03 |
| CAMP2_HUMAN\|Q08AD1_S1313 | 1.59398015 | 1.52E-04 | 2.91E-03 |
| LAP2A_HUMAN\|P42166_T160 | 1.593978827 | 3.39E-05 | 5.83E-04 |
| RABE1_HUMAN\|Q15276_S410_T408 | 1.593863634 | 3.78E-05 | 1.06E-03 |
| AKAP8_HUMAN\|O43823_S323 | 1.593855164 | 3.04E-03 | 1.52E-02 |
| MED24_HUMAN\|O75448_S873 | 1.593810171 | 3.34E-03 | 3.08E-02 |
| TB182_HUMAN\|Q9C0C2_S936 | 1.593593576 | 4.24E-05 | 6.72E-04 |
| SPB1_HUMAN\|Q8IY81_T573 | 1.593317155 | 1.14E-04 | 1.87E-03 |

| | | | |
|---|---|---|---|
| PPIG_HUMAN\|Q13427_S696 | 1.593296694 | 1.60E-03 | 3.79E-02 |
| SF3B2_HUMAN\|Q13435_S307_S309 | 1.593107517 | 5.49E-03 | 2.13E-02 |
| MAP1B_HUMAN\|P46821_S1501 | 1.593073992 | 1.11E-05 | 6.37E-04 |
| RHG05_HUMAN\|Q13017_S1173 | 1.592986718 | 7.31E-05 | 7.41E-04 |
| C2CD5_HUMAN\|Q86YS7_S307 | 1.592887108 | 1.17E-04 | 1.90E-03 |
| DEFI6_HUMAN\|Q9H4E7_S590_S597_T595 | 1.592447728 | 3.50E-04 | 5.97E-03 |
| MTAP2_HUMAN\|P11137_T1602_T1605 | 1.592130185 | 3.88E-03 | 3.87E-02 |
| UBP20_HUMAN\|Q9Y2K6_S132_S134 | 1.592077264 | 7.24E-03 | 2.65E-02 |
| FA83H_HUMAN\|Q6ZRV2_S513_S516 | 1.592065517 | 1.43E-03 | 1.60E-02 |
| K2C5_HUMAN\|P13647_S572 | 1.591929906 | 2.91E-04 | 3.11E-03 |
| RABL6_HUMAN\|Q3YEC7_S425_S427 | 1.591864895 | 8.33E-06 | 4.30E-04 |
| AHNK2_HUMAN\|Q8IVF2_S1583 | 1.591803707 | 1.39E-03 | 1.88E-02 |
| ERF_HUMAN\|P50548_S24 | 1.591707829 | 6.15E-05 | 2.19E-03 |
| K2C75_HUMAN\|O95678_S61 | 1.591457323 | 3.28E-03 | 3.27E-02 |
| PRC2C_HUMAN\|Q9Y520_T2691 | 1.59143367 | 6.44E-05 | 9.25E-04 |
| MARCS_HUMAN\|P29966_S118 | 1.591392841 | 2.42E-03 | 1.87E-02 |
| ZFN2B_HUMAN\|Q8WV99_S183 | 1.591142477 | 2.26E-04 | 1.22E-02 |
| VIME_HUMAN\|P08670_S51 | 1.590805415 | 2.77E-04 | 5.10E-03 |
| GA2L1_HUMAN\|Q99501_S352 | 1.5907545 | 4.33E-03 | 2.42E-02 |
| KMT2A_HUMAN\|Q03164_S3036 | 1.590708808 | 5.09E-03 | 2.02E-02 |
| SRC8_HUMAN\|Q14247_S418 | 1.590348697 | 5.94E-04 | 1.08E-02 |
| STR3N_HUMAN\|O95772_S39 | 1.590046093 | 1.10E-03 | 1.65E-02 |
| CLAP1_HUMAN\|Q7Z460_S1091 | 1.589697688 | 3.08E-05 | 1.06E-03 |
| ARHG5_HUMAN\|Q12774_S445_S450 | 1.589134431 | 4.89E-06 | 3.05E-04 |
| TANC2_HUMAN\|Q9HCD6_S1827 | 1.589035714 | 1.94E-05 | 9.55E-04 |
| KI21A_HUMAN\|Q7Z4S6_S855 | 1.588855386 | 1.04E-04 | 1.76E-03 |
| CTNB1_HUMAN\|P35222_T551 | 1.588841168 | 3.84E-03 | 3.29E-02 |
| P66B_HUMAN\|Q8WXI9_S334 | 1.588689038 | 3.04E-06 | 9.31E-05 |
| K1671_HUMAN\|Q9BY89_S1441 | 1.588661004 | 1.05E-03 | 1.32E-02 |
| GA2L1_HUMAN\|Q99501_S352 | 1.58854138 | 4.43E-03 | 2.07E-02 |
| PKHG3_HUMAN\|A1L390_S1079 | 1.58847189 | 2.34E-04 | 3.16E-03 |

| | | | |
|---|---|---|---|
| K1C17_HUMAN|Q04695_T9 | 1.588231907 | 8.30E-04 | 6.89E-03 |
| CLIP1_HUMAN|P30622_S200 | 1.588228781 | 1.59E-03 | 1.75E-02 |
| DESP_HUMAN|P15924_S165 | 1.588167813 | 1.61E-03 | 1.33E-02 |
| PLEC_HUMAN|Q15149_S4670_S4675 | 1.588155371 | 9.08E-03 | 4.80E-02 |
| SRRM2_HUMAN|Q9UQ35_S1463 | 1.588100642 | 1.21E-03 | 6.41E-03 |
| MK01_HUMAN|P28482_Y187 | 1.587943282 | 5.31E-04 | 9.66E-03 |
| TANC2_HUMAN|Q9HCD6_S1827 | 1.587821867 | 1.99E-05 | 1.09E-03 |
| MISP_HUMAN|Q8IVT2_S395 | 1.58729259 | 1.52E-03 | 1.70E-02 |
| AHNK_HUMAN|Q09666_S5739 | 1.587165188 | 9.19E-04 | 1.37E-02 |
| MISP_HUMAN|Q8IVT2_S394_S395 | 1.587049717 | 5.59E-05 | 2.23E-03 |
| ACINU_HUMAN|Q9UKV3_S838 | 1.586823633 | 2.17E-05 | 5.34E-04 |
| PHF8_HUMAN|Q9UPP1_S1021 | 1.586797686 | 1.25E-02 | 4.81E-02 |
| AMOL2_HUMAN|Q9Y2J4_S759 | 1.586628264 | 2.62E-05 | 4.74E-04 |
| SH3K1_HUMAN|Q96B97_S587 | 1.586535583 | 6.07E-05 | 8.89E-04 |
| WRIP1_HUMAN|Q96S55_T116 | 1.586535118 | 8.60E-03 | 4.04E-02 |
| FRMD8_HUMAN|Q9BZ67_S446 | 1.586452991 | 8.47E-04 | 1.36E-02 |
| PAXX_HUMAN|Q9BUH6_S152 | 1.586374982 | 2.95E-03 | 1.83E-02 |
| LMNB2_HUMAN|Q03252_S424 | 1.586316085 | 5.68E-04 | 5.26E-03 |
| TB182_HUMAN|Q9C0C2_S1158 | 1.585892736 | 3.55E-04 | 6.00E-03 |
| K1C17_HUMAN|Q04695_S28 | 1.585721822 | 7.98E-04 | 1.02E-02 |
| COF1_HUMAN|P23528_S24 | 1.584904528 | 2.10E-03 | 2.66E-02 |
| CLIP4_HUMAN|Q8N3C7_S610 | 1.584799158 | 4.56E-04 | 8.55E-03 |
| MAP1B_HUMAN|P46821_S1793_S1797 | 1.584410568 | 3.76E-05 | 1.60E-03 |
| IF4B_HUMAN|P23588_S497_S498 | 1.584359757 | 3.56E-04 | 7.56E-03 |
| RAI14_HUMAN|Q9P0K7_S296 | 1.58425858 | 6.71E-04 | 2.31E-02 |
| HELZ_HUMAN|P42694_S1766 | 1.584208672 | 2.28E-03 | 1.20E-02 |
| K2C5_HUMAN|P13647_S62 | 1.584206265 | 1.98E-04 | 2.27E-03 |
| VIME_HUMAN|P08670_S325 | 1.583951073 | 3.28E-03 | 3.05E-02 |
| SPD2A_HUMAN|Q5TCZ1_S1002 | 1.583920051 | 2.78E-05 | 9.87E-04 |
| K2C5_HUMAN|P13647_S571 | 1.583806456 | 3.66E-05 | 6.10E-04 |
| GYS1_HUMAN|P13807_S727_S731 | 1.583796265 | 7.78E-04 | 5.00E-03 |

| | | | |
|---|---|---|---|
| MAP7_HUMAN\|Q14244_T184 | 1.58377192 | 7.76E-05 | 2.49E-03 |
| K1C14_HUMAN\|P02533_S51 | 1.583721338 | 1.79E-03 | 2.15E-02 |
| PALLD_HUMAN\|Q8WX93_S893 | 1.583664743 | 3.77E-04 | 2.55E-03 |
| AHNK_HUMAN\|Q09666_S5745_S5749 | 1.583612694 | 1.35E-05 | 1.68E-03 |
| ACINU_HUMAN\|Q9UKV3_S240 | 1.583364955 | 2.67E-04 | 6.21E-03 |
| ABLM1_HUMAN\|O14639_S431_S435 | 1.583166253 | 2.06E-03 | 2.59E-02 |
| K1C14_HUMAN\|P02533_S429_S432_S433 | 1.582919425 | 7.99E-04 | 1.25E-02 |
| CALD1_HUMAN\|Q05682_S789 | 1.58241909 | 5.05E-04 | 1.90E-02 |
| AKTS1_HUMAN\|Q96B36_S202_S203 | 1.582232615 | 1.43E-05 | 8.53E-04 |
| UTP18_HUMAN\|Q9Y5J1_S205 | 1.581728518 | 9.30E-03 | 3.27E-02 |
| PDLI5_HUMAN\|Q96HC4_S111 | 1.581682521 | 1.25E-04 | 2.49E-03 |
| P63_HUMAN\|Q9H3D4_S463 | 1.581484128 | 1.34E-05 | 2.86E-04 |
| TCAL3_HUMAN\|Q969E4_S125 | 1.581483005 | 1.16E-03 | 2.34E-02 |
| I2BP2_HUMAN\|Q7Z5L9_S455_S460 | 1.581415746 | 2.42E-04 | 1.81E-03 |
| LIPA1_HUMAN\|Q13136_S666_S668 | 1.581403522 | 5.69E-05 | 1.60E-03 |
| CAMP1_HUMAN\|Q5T5Y3_S827 | 1.581349001 | 8.90E-04 | 2.73E-02 |
| HCFC1_HUMAN\|P51610_S1507 | 1.581268678 | 2.56E-03 | 3.01E-02 |
| MICA2_HUMAN\|O94851_S699 | 1.580790253 | 2.84E-04 | 2.04E-03 |
| K2C5_HUMAN\|P13647_S75 | 1.580729641 | 1.12E-03 | 1.64E-02 |
| STR3N_HUMAN\|O95772_S39 | 1.580713934 | 1.24E-03 | 1.10E-02 |
| SFR19_HUMAN\|Q9H7N4_T989 | 1.580518424 | 8.73E-03 | 3.60E-02 |
| DBNL_HUMAN\|Q9UJU6_T271 | 1.580505604 | 1.55E-04 | 3.30E-03 |
| K2C75_HUMAN\|O95678_S535 | 1.580357915 | 1.05E-05 | 5.07E-04 |
| SERF2_HUMAN\|P84101_S21 | 1.580325977 | 4.75E-04 | 8.93E-03 |
| K2C6B_HUMAN\|P04259_S34 | 1.579786738 | 5.75E-03 | 4.46E-02 |
| IASPP_HUMAN\|Q8WUF5_Y109 | 1.579618595 | 9.09E-05 | 1.93E-03 |
| AHNK2_HUMAN\|Q8IVF2_S1583 | 1.579535669 | 1.61E-03 | 1.16E-02 |
| K1C17_HUMAN\|Q04695_T9 | 1.579459354 | 9.37E-04 | 1.22E-02 |
| PAK6_HUMAN\|Q9NQU5_S308 | 1.57914043 | 4.58E-03 | 3.81E-02 |
| CALD1_HUMAN\|Q05682_S789 | 1.579125523 | 5.29E-04 | 8.06E-03 |
| ABI2_HUMAN\|Q9NYB9_S368 | 1.579040321 | 5.03E-06 | 3.07E-04 |

| | | | |
|---|---|---|---|
| DKC1_HUMAN\|O60832_S485_T496 | 1.578978489 | 4.97E-06 | 1.91E-04 |
| CAMP1_HUMAN\|Q5T5Y3_S827 | 1.578753728 | 9.21E-04 | 5.68E-03 |
| TBC15_HUMAN\|Q8TC07_S70 | 1.578629983 | 8.50E-03 | 4.60E-02 |
| PDLI2_HUMAN\|Q96JY6_S124_S127_S129 | 1.5782572 | 1.42E-03 | 1.89E-02 |
| MYO1E_HUMAN\|Q12965_S980 | 1.578219373 | 4.72E-04 | 6.81E-03 |
| RGL2_HUMAN\|O15211_S619 | 1.578161956 | 1.79E-04 | 2.10E-03 |
| KDM2A_HUMAN\|Q9Y2K7_S558 | 1.577922847 | 6.14E-03 | 3.11E-02 |
| K1C14_HUMAN\|P02533_S13 | 1.577889167 | 3.44E-05 | 1.12E-03 |
| IRS1_HUMAN\|P35568_S1100 | 1.577646389 | 9.07E-05 | 1.21E-03 |
| RLA1_HUMAN\|P05386_S101 | 1.57756892 | 1.18E-03 | 6.94E-03 |
| MISP_HUMAN\|Q8IVT2_S395_S397 | 1.577281709 | 4.71E-04 | 5.39E-03 |
| CCDC9_HUMAN\|Q9Y3X0_T389 | 1.577228018 | 3.92E-06 | 1.50E-04 |
| STK10_HUMAN\|O94804_T952 | 1.577226345 | 1.09E-03 | 1.30E-02 |
| RS28_HUMAN\|P62857_S23 | 1.577048371 | 4.83E-03 | 3.02E-02 |
| BORG4_HUMAN\|Q9H3Q1_S64 | 1.577027112 | 8.64E-05 | 1.87E-03 |
| ADCY6_HUMAN\|O43306_S54 | 1.576919206 | 1.59E-04 | 8.80E-03 |
| RAI14_HUMAN\|Q9P0K7_T251 | 1.576634438 | 5.37E-04 | 8.13E-03 |
| K2C5_HUMAN\|P13647_S26 | 1.576509874 | 1.80E-04 | 3.64E-03 |
| TMF1_HUMAN\|P82094_S72 | 1.576316884 | 3.59E-05 | 7.69E-04 |
| DCAF6_HUMAN\|Q58WW2_S657 | 1.576228935 | 5.72E-04 | 6.23E-03 |
| SGTA_HUMAN\|O43765_S305 | 1.576086729 | 1.35E-04 | 1.14E-03 |
| APBB2_HUMAN\|Q92870_S409_S412 | 1.575384691 | 6.62E-05 | 1.77E-03 |
| BIG1_HUMAN\|Q9Y6D6_S397 | 1.575334111 | 7.71E-05 | 1.07E-03 |
| MAP7_HUMAN\|Q14244_S202 | 1.575233266 | 2.79E-05 | 8.28E-04 |
| SMRC1_HUMAN\|Q92922_S328_S330 | 1.575228222 | 1.90E-04 | 4.51E-03 |
| STMN1_HUMAN\|P16949_S25 | 1.57516316 | 4.08E-05 | 1.84E-03 |
| ARHG5_HUMAN\|Q12774_S450 | 1.575066872 | 1.33E-03 | 1.88E-02 |
| LRCH1_HUMAN\|Q9Y2L9_S323_S327 | 1.5749998 | 6.30E-03 | 3.17E-02 |
| PCM1_HUMAN\|Q15154_S93 | 1.574971162 | 1.01E-05 | 1.86E-03 |
| NFIC_HUMAN\|P08651_S284_S305_Y299 | 1.57461411 | 5.94E-04 | 6.40E-03 |
| PUM2_HUMAN\|Q8TB72_S182 | 1.574574399 | 1.52E-03 | 8.50E-03 |

| | | | |
|---|---|---|---|
| BCAR3_HUMAN\|O75815_S370 | 1.574566089 | 1.25E-03 | 1.52E-02 |
| MARCS_HUMAN\|P29966_T150 | 1.574566071 | 7.96E-04 | 6.72E-03 |
| K1C14_HUMAN\|P02533_S33 | 1.574518386 | 2.53E-04 | 4.77E-03 |
| MP2K4_HUMAN\|P45985_S393 | 1.574454664 | 4.20E-03 | 4.13E-02 |
| LMNA_HUMAN\|P02545_S619 | 1.573881071 | 9.20E-04 | 1.14E-02 |
| PHLB1_HUMAN\|Q86UU1_S489 | 1.573825985 | 2.11E-03 | 1.44E-02 |
| K1468_HUMAN\|Q9P260_S180 | 1.573768339 | 4.35E-05 | 4.75E-04 |
| AKTS1_HUMAN\|Q96B36_S187 | 1.573756208 | 7.64E-04 | 6.50E-03 |
| HSPB1_HUMAN\|P04792_S82 | 1.573570478 | 9.87E-06 | 7.13E-04 |
| SRRM2_HUMAN\|Q9UQ35_S1383_S1387 | 1.5734455 | 1.15E-03 | 1.60E-02 |
| ARHGB_HUMAN\|O15085_S1457_T1462 | 1.57333183 | 8.04E-04 | 4.62E-03 |
| TP53B_HUMAN\|Q12888_S366_T357 | 1.572923387 | 2.67E-04 | 4.49E-03 |
| CAMP1_HUMAN\|Q5T5Y3_S827 | 1.572874115 | 9.93E-04 | 9.42E-03 |
| VIME_HUMAN\|P08670_T458 | 1.572824825 | 4.56E-05 | 1.24E-02 |
| ITB4_HUMAN\|P16144_S1454_T1455 | 1.572297566 | 4.64E-03 | 3.84E-02 |
| MACF1_HUMAN\|Q9UPN3_S7330 | 1.572122982 | 1.69E-03 | 9.27E-03 |
| DKC1_HUMAN\|O60832_S485 | 1.571945941 | 1.20E-04 | 3.18E-03 |
| LDLR_HUMAN\|P01130_S854 | 1.571945033 | 8.17E-03 | 2.94E-02 |
| K1C14_HUMAN\|P02533_S33 | 1.571712665 | 2.65E-04 | 1.24E-02 |
| NDRG1_HUMAN\|Q92597_S336 | 1.571668596 | 9.26E-05 | 1.51E-03 |
| CING_HUMAN\|Q9P2M7_S129 | 1.571591691 | 2.30E-03 | 2.30E-02 |
| FOXK2_HUMAN\|Q01167_S398 | 1.571339165 | 2.73E-03 | 1.24E-02 |
| LRRF2_HUMAN\|Q9Y608_S320_T317 | 1.571084196 | 3.34E-03 | 2.38E-02 |
| TB182_HUMAN\|Q9C0C2_S1533 | 1.570771119 | 2.55E-04 | 4.78E-03 |
| K1C15_HUMAN\|P19012_S34 | 1.570718951 | 6.76E-03 | 3.34E-02 |
| AHNK_HUMAN\|Q09666_S210_S216_S220 | 1.570470074 | 5.53E-05 | 4.40E-03 |
| PLEC_HUMAN\|Q15149_S4384 | 1.569858884 | 1.37E-03 | 1.18E-02 |
| PLEC_HUMAN\|Q15149_S4613_S4618_T4623 | 1.569832119 | 1.17E-05 | 2.63E-04 |
| PDLI2_HUMAN\|Q96JY6_S117 | 1.56968233 | 6.28E-04 | 3.78E-03 |
| AHNK_HUMAN\|Q09666_S216_S220 | 1.569533925 | 3.29E-05 | 9.48E-04 |
| CBX3_HUMAN\|Q13185_S93 | 1.569325878 | 4.93E-03 | 1.98E-02 |

| | | | |
|---|---|---|---|
| TRI22_HUMAN\|Q8IYM9_S384 | 1.569195556 | 5.99E-04 | 1.07E-02 |
| NOG1_HUMAN\|Q9BZE4_S468_S470 | 1.569152001 | 1.15E-02 | 4.51E-02 |
| PKP2_HUMAN\|Q99959_S151 | 1.569145059 | 7.55E-05 | 7.56E-04 |
| IF2B_HUMAN\|P20042_T111 | 1.569036962 | 1.33E-03 | 1.58E-02 |
| EPN1_HUMAN\|Q9Y6I3_S454 | 1.568835842 | 3.21E-05 | 3.68E-04 |
| TGS1_HUMAN\|Q96RS0_S55 | 1.568688868 | 2.40E-04 | 1.91E-03 |
| ALKB5_HUMAN\|Q6P6C2_S374_S384 | 1.568598339 | 5.21E-03 | 4.08E-02 |
| ZO1_HUMAN\|Q07157_Y843 | 1.568526365 | 2.82E-03 | 1.43E-02 |
| TR112_HUMAN\|Q9UI30_S125 | 1.568365009 | 1.10E-02 | 3.73E-02 |
| EF2K_HUMAN\|O00418_S470 | 1.568333901 | 1.94E-04 | 6.62E-03 |
| EFNB1_HUMAN\|P98172_S281 | 1.568174256 | 3.95E-04 | 5.97E-03 |
| DEFI6_HUMAN\|Q9H4E7_S597 | 1.568121037 | 3.74E-06 | 1.09E-04 |
| NGAP_HUMAN\|Q9UJF2_S954 | 1.567687748 | 2.93E-04 | 2.10E-03 |
| SOS2_HUMAN\|Q07890_S1267 | 1.567646748 | 1.87E-03 | 4.33E-02 |
| MTAP2_HUMAN\|P11137_S1782_S1790 | 1.567515378 | 8.34E-03 | 4.54E-02 |
| NDRG1_HUMAN\|Q92597_S326_S330_S333 | 1.567417607 | 4.20E-04 | 6.22E-03 |
| ASPP2_HUMAN\|Q13625_S558 | 1.567240115 | 2.41E-05 | 1.27E-03 |
| DESP_HUMAN\|P15924_S165 | 1.567087456 | 2.07E-03 | 1.10E-02 |
| MA7D1_HUMAN\|Q3KQU3_S452 | 1.56697601 | 6.21E-04 | 2.19E-02 |
| COHA1_HUMAN\|Q9UMD9_S85_S93_T97 | 1.56695573 | 3.03E-03 | 3.12E-02 |
| PKP4_HUMAN\|Q99569_S236 | 1.566948999 | 1.56E-04 | 3.94E-03 |
| KDM3B_HUMAN\|Q7LBC6_S289 | 1.566859552 | 1.34E-04 | 3.88E-03 |
| NIPA_HUMAN\|Q86WB0_S408_S409 | 1.566721312 | 5.25E-06 | 4.19E-04 |
| BORG4_HUMAN\|Q9H3Q1_S64 | 1.566401508 | 1.04E-04 | 1.34E-03 |
| SARG_HUMAN\|Q9BW04_S131 | 1.566378257 | 5.37E-03 | 2.83E-02 |
| MAST4_HUMAN\|O15021_S1387_S1394 | 1.566347269 | 4.12E-05 | 4.55E-04 |
| T200A_HUMAN\|Q86VY9_S350 | 1.566194769 | 2.47E-03 | 2.71E-02 |
| K2C5_HUMAN\|P13647_S574 | 1.566101715 | 1.26E-04 | 2.79E-03 |
| SRRM1_HUMAN\|Q8IYB3_S752_S754_S756 | 1.565763929 | 2.10E-05 | 2.92E-03 |
| TB182_HUMAN\|Q9C0C2_S1652 | 1.565677985 | 2.00E-04 | 2.30E-03 |
| RIPK2_HUMAN\|O43353_S531 | 1.565637785 | 4.68E-05 | 9.43E-04 |

| | | | |
|---|---|---|---|
| LAD1_HUMAN\|O00515_S38_S39 | 1.565631924 | 4.46E-03 | 3.74E-02 |
| GGYF1_HUMAN\|O75420_S157 | 1.565601364 | 7.59E-03 | 3.23E-02 |
| K1C14_HUMAN\|P02533_S51_S54 | 1.565592129 | 1.43E-03 | 3.73E-02 |
| LRRF2_HUMAN\|Q9Y608_S320_T317 | 1.565502699 | 3.55E-03 | 2.43E-02 |
| MAP7_HUMAN\|Q14244_S202 | 1.565372157 | 3.39E-05 | 3.37E-03 |
| CAVN3_HUMAN\|Q969G5_T205 | 1.565301177 | 9.71E-04 | 7.76E-03 |
| CE170_HUMAN\|Q5SW79_S630_S636 | 1.565286409 | 4.25E-05 | 1.30E-03 |
| LIMA1_HUMAN\|Q9UHB6_S507 | 1.565040266 | 1.55E-03 | 2.07E-02 |
| SCRIB_HUMAN\|Q14160_S1285 | 1.564974958 | 1.88E-03 | 2.28E-02 |
| GIT1_HUMAN\|Q9Y2X7_S388_T392 | 1.564799631 | 1.64E-03 | 1.35E-02 |
| K1468_HUMAN\|Q9P260_S180 | 1.564593508 | 5.17E-05 | 1.00E-03 |
| 3BP1_HUMAN\|Q9Y3L3_T601 | 1.564530776 | 4.89E-03 | 4.31E-02 |
| RABE1_HUMAN\|Q15276_S407_S410 | 1.564432632 | 6.75E-07 | 6.24E-05 |
| K1C14_HUMAN\|P02533_S432 | 1.564162383 | 1.82E-03 | 1.27E-02 |
| SFR19_HUMAN\|Q9H7N4_S719_S725 | 1.564071438 | 9.95E-04 | 2.08E-02 |
| SRP09_HUMAN\|P49458_T85 | 1.563887829 | 2.29E-04 | 5.21E-03 |
| ROCK1_HUMAN\|Q13464_S1333_S1336_S1341 | 1.563781832 | 5.10E-03 | 4.03E-02 |
| PAXI_HUMAN\|P49023_S103_S106 | 1.56362009 | 2.73E-03 | 1.24E-02 |
| FXR2_HUMAN\|P51116_S453 | 1.563553845 | 2.36E-04 | 4.52E-03 |
| NEB2_HUMAN\|Q96SB3_S99 | 1.563257288 | 2.24E-04 | 1.81E-03 |
| BZW2_HUMAN\|Q9Y6E2_S412 | 1.563082517 | 2.86E-03 | 2.11E-02 |
| K1C14_HUMAN\|P02533_S51 | 1.563058541 | 2.31E-03 | 1.52E-02 |
| NOG1_HUMAN\|Q9BZE4_S470_Y466 | 1.56281031 | 8.65E-05 | 1.43E-03 |
| CSCL2_HUMAN\|Q5T3F8_T110 | 1.562795232 | 3.21E-03 | 1.59E-02 |
| TP53B_HUMAN\|Q12888_S356 | 1.562689898 | 7.50E-03 | 3.61E-02 |
| AHNK_HUMAN\|Q09666_S5749_S5763 | 1.562666503 | 6.45E-06 | 3.60E-04 |
| GYS1_HUMAN\|P13807_S727 | 1.562586465 | 5.46E-05 | 5.51E-04 |
| ACINU_HUMAN\|Q9UKV3_S243 | 1.562580007 | 4.07E-05 | 6.53E-04 |
| GBF1_HUMAN\|Q92538_S1781_S1784 | 1.562559349 | 5.47E-04 | 1.02E-02 |
| YAP1_HUMAN\|P46937_S109_T119 | 1.562456283 | 6.88E-06 | 1.71E-04 |
| CSCL2_HUMAN\|Q5T3F8_S111_S114 | 1.562118229 | 4.22E-03 | 2.89E-02 |

| | | | |
|---|---|---|---|
| NIBL1_HUMAN\|Q96TA1_S692 | 1.562112001 | 9.70E-05 | 3.19E-03 |
| AAK1_HUMAN\|Q2M2I8_S678_T674 | 1.561787572 | 6.36E-05 | 2.69E-03 |
| MAP7_HUMAN\|Q14244_S315 | 1.561715269 | 1.14E-04 | 6.96E-03 |
| LMO7_HUMAN\|Q8WWI1_S1421 | 1.561621217 | 1.88E-03 | 1.31E-02 |
| F120A_HUMAN\|Q9NZB2_S1023 | 1.561389509 | 9.08E-05 | 1.93E-03 |
| EP15R_HUMAN\|Q9UBC2_S229 | 1.561320204 | 2.25E-06 | 2.11E-04 |
| GSK3A_HUMAN\|P49840_T19 | 1.561258233 | 3.56E-05 | 1.02E-03 |
| TB182_HUMAN\|Q9C0C2_S429 | 1.560999969 | 4.93E-03 | 4.02E-02 |
| K1C14_HUMAN\|P02533_S429_S435 | 1.560954632 | 1.27E-03 | 1.47E-02 |
| PR38B_HUMAN\|Q5VTL8_S529 | 1.560948186 | 1.72E-04 | 3.51E-03 |
| HIRP3_HUMAN\|Q9BW71_S196_S199 | 1.560929236 | 8.08E-05 | 2.07E-03 |
| RHG32_HUMAN\|A7KAX9_S682 | 1.560827538 | 2.49E-05 | 1.15E-03 |
| ARG28_HUMAN\|Q8N1W1_S558 | 1.560745758 | 2.07E-03 | 1.42E-02 |
| NOC2L_HUMAN\|Q9Y3T9_S672 | 1.560589744 | 1.55E-02 | 4.91E-02 |
| MILK1_HUMAN\|Q8N3F8_S391 | 1.560465239 | 1.81E-06 | 6.32E-05 |
| RHG01_HUMAN\|Q07960_S51 | 1.560463634 | 1.74E-04 | 4.62E-03 |
| SVIL_HUMAN\|O95425_T679 | 1.560188082 | 9.07E-06 | 5.85E-04 |
| NFIC_HUMAN\|P08651_S294_S304 | 1.560187125 | 6.51E-04 | 5.80E-03 |
| CLIP2_HUMAN\|Q9UDT6_S207_T198 | 1.560164191 | 1.66E-03 | 1.36E-02 |
| IWS1_HUMAN\|Q96ST2_T725 | 1.560149767 | 1.64E-03 | 1.79E-02 |
| K2C6B_HUMAN\|P04259_S541 | 1.560096915 | 3.15E-04 | 5.51E-03 |
| TBCD5_HUMAN\|Q92609_S522 | 1.55999878 | 3.36E-04 | 5.79E-03 |
| COBL1_HUMAN\|Q53SF7_S364 | 1.559449002 | 1.06E-04 | 3.03E-03 |
| IF4G2_HUMAN\|P78344_T508 | 1.559377696 | 5.95E-06 | 1.53E-04 |
| SP100_HUMAN\|P23497_S180 | 1.559356032 | 5.62E-04 | 3.46E-03 |
| BORG4_HUMAN\|Q9H3Q1_S73 | 1.559300799 | 4.15E-03 | 3.56E-02 |
| HMCS1_HUMAN\|Q01581_S516 | 1.55901378 | 5.67E-04 | 7.84E-03 |
| EPN3_HUMAN\|Q9H201_S188_S191 | 1.558914818 | 5.31E-04 | 8.07E-03 |
| AHNK2_HUMAN\|Q8IVF2_S5175 | 1.558809028 | 5.25E-04 | 9.59E-03 |
| FOSL2_HUMAN\|P15408_S19 | 1.558732651 | 7.55E-03 | 4.18E-02 |
| CLIP4_HUMAN\|Q8N3C7_T582 | 1.558563921 | 1.61E-04 | 2.36E-03 |

| | | | |
|---|---|---|---|
| GIT1_HUMAN\|Q9Y2X7_S388 | 1.558464845 | 5.42E-03 | 3.45E-02 |
| DESP_HUMAN\|P15924_S165 | 1.55845581 | 2.30E-03 | 2.80E-02 |
| ACAP2_HUMAN\|Q15057_S384 | 1.558431462 | 1.05E-05 | 7.27E-04 |
| PEAK1_HUMAN\|Q9H792_T574 | 1.558293219 | 4.15E-05 | 4.49E-04 |
| XPC_HUMAN\|Q01831_S883_S884_S891 | 1.558194707 | 2.21E-03 | 2.50E-02 |
| CPSF4_HUMAN\|O95639_S200 | 1.558099813 | 1.27E-04 | 1.94E-03 |
| ENSA_HUMAN\|O43768_S109 | 1.558098993 | 5.87E-04 | 8.07E-03 |
| TMF1_HUMAN\|P82094_S338_S344 | 1.558096054 | 3.43E-05 | 5.87E-04 |
| NFIA_HUMAN\|Q12857_S287 | 1.557766113 | 1.97E-03 | 1.35E-02 |
| DCBD2_HUMAN\|Q96PD2_S724_S727 | 1.557763825 | 4.30E-03 | 3.95E-02 |
| SAFB1_HUMAN\|Q15424_S247 | 1.557624951 | 8.03E-05 | 1.79E-03 |
| NDRG1_HUMAN\|Q92597_S333_S336_T328 | 1.557574222 | 3.72E-05 | 1.05E-03 |
| SCAM2_HUMAN\|O15127_S320 | 1.557515428 | 2.55E-03 | 3.01E-02 |
| KC1A_HUMAN\|P48729-3_S312 | 1.557465136 | 2.95E-03 | 2.73E-02 |
| ZBT21_HUMAN\|Q9ULJ3_S422 | 1.557245312 | 7.50E-03 | 3.20E-02 |
| K1C14_HUMAN\|P02533_S432_S433_S438 | 1.556667755 | 2.28E-03 | 1.52E-02 |
| RIN1_HUMAN\|Q13671_S349 | 1.556533294 | 2.48E-03 | 3.97E-02 |
| EF2K_HUMAN\|O00418_S470 | 1.556458729 | 2.34E-04 | 5.50E-03 |
| GGYF1_HUMAN\|O75420_S638 | 1.556389912 | 2.44E-04 | 1.93E-03 |
| PRC2A_HUMAN\|P48634_S1147 | 1.556176876 | 2.94E-05 | 5.14E-04 |
| K1C14_HUMAN\|P02533_S11 | 1.556004085 | 7.40E-05 | 1.71E-03 |
| PKHG3_HUMAN\|A1L390_S640_S643 | 1.555910404 | 6.74E-04 | 6.95E-03 |
| AHNK_HUMAN\|Q09666_S210 | 1.555903057 | 1.37E-05 | 6.10E-04 |
| LAD1_HUMAN\|O00515_T41 | 1.555794675 | 2.76E-03 | 1.98E-02 |
| RB3GP_HUMAN\|Q15042_S537 | 1.555481193 | 2.70E-05 | 1.32E-03 |
| NDRG1_HUMAN\|Q92597_S393 | 1.555234325 | 1.29E-03 | 1.73E-02 |
| ADAM9_HUMAN\|Q13443_S758 | 1.555183384 | 3.97E-03 | 1.68E-02 |
| TRI29_HUMAN\|Q14134_S152 | 1.554556866 | 2.79E-04 | 4.61E-03 |
| RALY_HUMAN\|Q9UKM9_T286_T298 | 1.554166935 | 1.25E-03 | 1.11E-02 |
| TBC9B_HUMAN\|Q66K14_S411 | 1.554099213 | 1.64E-03 | 1.35E-02 |
| TBCD5_HUMAN\|Q92609_S541 | 1.554047786 | 8.28E-07 | 5.08E-05 |

| | | | |
|---|---|---|---|
| K1C14_HUMAN\|P02533_S53 | 1.553869434 | 8.33E-05 | 1.13E-03 |
| PACS1_HUMAN\|Q6VY07_T504 | 1.553818843 | 9.41E-04 | 1.22E-02 |
| MAP4_HUMAN\|P27816_S928 | 1.553771845 | 2.62E-04 | 4.46E-03 |
| MYOME_HUMAN\|Q5VU43-2_S195 | 1.553737946 | 1.48E-03 | 1.25E-02 |
| P33MX_HUMAN\|Q96A73_S183_T179 | 1.55369048 | 1.74E-03 | 3.10E-02 |
| UBXN7_HUMAN\|O94888_S278 | 1.553682368 | 1.10E-02 | 3.74E-02 |
| NP1L1_HUMAN\|P55209_T62 | 1.553603639 | 4.42E-03 | 2.46E-02 |
| K2C5_HUMAN\|P13647_S11_S8 | 1.553603277 | 9.04E-05 | 2.25E-03 |
| ZCHC8_HUMAN\|Q6NZY4_T479 | 1.553322071 | 9.87E-05 | 1.29E-03 |
| LAD1_HUMAN\|O00515_S38_T40_T41 | 1.553287693 | 1.11E-03 | 1.63E-02 |
| TBCD5_HUMAN\|Q92609_S554 | 1.552983944 | 5.66E-04 | 3.48E-03 |
| DDX10_HUMAN\|Q13206_S831 | 1.552893034 | 2.05E-04 | 2.83E-03 |
| CR025_HUMAN\|Q96B23_S67 | 1.552793095 | 2.12E-03 | 1.71E-02 |
| NIBL1_HUMAN\|Q96TA1_S691_S696 | 1.552738986 | 1.88E-04 | 4.82E-03 |
| RB6I2_HUMAN\|Q8IUD2_S17_S21 | 1.552555378 | 1.38E-04 | 1.71E-03 |
| NUP98_HUMAN\|P52948_S623 | 1.552361416 | 9.01E-04 | 1.95E-02 |
| SPD2A_HUMAN\|Q5TCZ1_S642 | 1.552326974 | 6.54E-04 | 8.80E-03 |
| K1143_HUMAN\|Q96AT1_S146 | 1.552307314 | 2.20E-03 | 2.26E-02 |
| IRS1_HUMAN\|P35568_S330 | 1.55221655 | 3.23E-03 | 1.60E-02 |
| PKHG3_HUMAN\|A1L390_S1079 | 1.552019503 | 4.02E-04 | 4.76E-03 |
| CNKR1_HUMAN\|Q969H4_S305_S314 | 1.551917165 | 9.21E-04 | 7.46E-03 |
| S23A2_HUMAN\|Q9UGH3_S78_S81 | 1.551763525 | 5.25E-03 | 3.38E-02 |
| CR025_HUMAN\|Q96B23_S143_S145_S147 | 1.551570592 | 9.94E-04 | 1.55E-02 |
| K1C16_HUMAN\|P08779_S450 | 1.551341911 | 3.08E-04 | 4.90E-03 |
| CSCL2_HUMAN\|Q5T3F8_S114_T110 | 1.55125231 | 5.02E-04 | 4.69E-02 |
| GOGA4_HUMAN\|Q13439_S89_S92 | 1.551242761 | 2.85E-04 | 5.20E-03 |
| K2C5_HUMAN\|P13647_S26 | 1.551124715 | 2.75E-04 | 4.58E-03 |
| SRPK2_HUMAN\|P78362_S496_T498 | 1.551046538 | 2.95E-04 | 2.11E-03 |
| TT39B_HUMAN\|Q5VTQ0_S120 | 1.550611089 | 3.22E-03 | 1.96E-02 |
| CE170_HUMAN\|Q5SW79_S636 | 1.550456521 | 9.67E-07 | 8.06E-05 |
| RMP_HUMAN\|O94763_S372 | 1.550347601 | 7.45E-04 | 9.72E-03 |

| | | | |
|---|---|---|---|
| K1C14_HUMAN|P02533_S51_S54 | 1.550106905 | 1.76E-03 | 2.13E-02 |
| AHNK_HUMAN|Q09666_T4430 | 1.549998086 | 5.62E-04 | 5.22E-03 |
| NFIB_HUMAN|O00712_S276_S295 | 1.54995084 | 1.29E-03 | 6.75E-03 |
| RGL2_HUMAN|O15211_S761 | 1.549876033 | 2.59E-04 | 2.85E-03 |
| LARP1_HUMAN|Q6PKG0_S1040 | 1.549743224 | 9.44E-05 | 2.32E-03 |
| LYAR_HUMAN|Q9NX58_S276 | 1.549696295 | 2.55E-05 | 1.28E-03 |
| CRK_HUMAN|P46108_S304 | 1.549528848 | 9.71E-06 | 1.47E-04 |
| ARHGB_HUMAN|O15085_S1519 | 1.549417758 | 2.08E-04 | 5.17E-03 |
| M3K3_HUMAN|Q99759_S337 | 1.549382297 | 1.16E-03 | 1.68E-02 |
| AAK1_HUMAN|Q2M2I8_S678 | 1.549349888 | 2.45E-04 | 1.19E-02 |
| FA83B_HUMAN|Q5T0W9_S664 | 1.549237155 | 5.59E-05 | 1.58E-03 |
| COHA1_HUMAN|Q9UMD9_S85_S93_T97 | 1.549063774 | 3.71E-03 | 3.22E-02 |
| ZCCHV_HUMAN|Q7Z2W4_T380 | 1.548751839 | 3.53E-04 | 5.50E-03 |
| ZO2_HUMAN|Q9UDY2_T1027 | 1.54873036 | 1.27E-04 | 3.46E-03 |
| PPR18_HUMAN|Q6NYC8_S224 | 1.548675352 | 7.26E-04 | 2.36E-02 |
| ARHGB_HUMAN|O15085_S1457_T1462 | 1.548554079 | 1.12E-03 | 1.71E-02 |
| TFE3_HUMAN|P19532_S568_S570 | 1.548431849 | 6.01E-05 | 1.11E-03 |
| IASPP_HUMAN|Q8WUF5_S567 | 1.548340977 | 8.78E-06 | 5.29E-04 |
| LAD1_HUMAN|O00515_S38 | 1.548323351 | 2.58E-04 | 3.44E-03 |
| NDE1_HUMAN|Q9NXR1_S282 | 1.548183385 | 6.49E-03 | 3.24E-02 |
| PKP2_HUMAN|Q99959_S151 | 1.548135738 | 1.08E-04 | 3.42E-03 |
| CRK_HUMAN|P46108_S304 | 1.548066838 | 1.00E-05 | 3.03E-04 |
| DC1L1_HUMAN|Q9Y6G9_T513_T515 | 1.547676179 | 1.47E-04 | 1.22E-03 |
| ITA2_HUMAN|P17301_S1180 | 1.547674414 | 1.59E-03 | 8.04E-03 |
| SRRM1_HUMAN|Q8IYB3_S756 | 1.547553732 | 8.23E-03 | 2.96E-02 |
| SYNEM_HUMAN|O15061_S1044_S1049 | 1.547527232 | 2.56E-04 | 1.21E-02 |
| LAP2A_HUMAN|P42166_S66_S67 | 1.547525043 | 3.30E-05 | 1.39E-03 |
| ASHWN_HUMAN|Q9BVC5_S193 | 1.547258362 | 1.13E-05 | 1.57E-03 |
| RPRD2_HUMAN|Q5VT52_T723 | 1.546845233 | 1.32E-05 | 5.98E-04 |
| ERBB2_HUMAN|P04626_S998 | 1.546801491 | 5.69E-03 | 3.38E-02 |
| YBOX3_HUMAN|P16989_S369_S370 | 1.546754031 | 3.71E-04 | 6.19E-03 |

| | | | |
|---|---|---|---|
| NFKB1_HUMAN\|P19838_S907 | 1.546671306 | 4.54E-06 | 4.11E-04 |
| K2C5_HUMAN\|P13647_S11_S14 | 1.546485572 | 5.13E-05 | 1.31E-03 |
| SF3B2_HUMAN\|Q13435_S307 | 1.546431464 | 3.62E-03 | 1.55E-02 |
| PEPL_HUMAN\|O60437_S1657 | 1.546393816 | 1.51E-04 | 3.24E-03 |
| AHNK_HUMAN\|Q09666_S135 | 1.546245984 | 2.07E-03 | 2.12E-02 |
| MARCS_HUMAN\|P29966_S118 | 1.546089499 | 4.07E-03 | 4.04E-02 |
| STXB5_HUMAN\|Q5T5C0-2_S712 | 1.545927002 | 2.57E-05 | 7.79E-04 |
| MINK1_HUMAN\|Q8N4C8_S701 | 1.545636585 | 7.37E-03 | 3.56E-02 |
| REPS1_HUMAN\|Q96D71_S162_S166_T173 | 1.545434367 | 4.28E-05 | 1.15E-03 |
| PKP3_HUMAN\|Q9Y446_S183 | 1.545425223 | 8.11E-04 | 1.27E-02 |
| KAP2_HUMAN\|P13861_S78_S80 | 1.545260927 | 1.94E-04 | 6.62E-03 |
| UN45A_HUMAN\|Q9H3U1_S147_Y145 | 1.54505814 | 2.84E-04 | 2.15E-03 |
| K1C14_HUMAN\|P02533_S433_S438 | 1.544634666 | 1.63E-03 | 2.02E-02 |
| ROAA_HUMAN\|Q99729-3_S255 | 1.544615922 | 2.21E-03 | 2.72E-02 |
| CT2NL_HUMAN\|Q9P2B4_S568 | 1.544610201 | 1.43E-04 | 1.76E-03 |
| MTUS1_HUMAN\|Q9ULD2_T1204 | 1.544464478 | 2.25E-04 | 5.50E-03 |
| TMF1_HUMAN\|P82094_S77 | 1.544415534 | 2.24E-04 | 3.03E-03 |
| RABL6_HUMAN\|Q3YEC7_S425_S427 | 1.544304394 | 2.26E-05 | 7.08E-04 |
| KPCD2_HUMAN\|Q9BZL6_S214 | 1.544146222 | 9.26E-04 | 1.45E-02 |
| RBM7_HUMAN\|Q9Y580_S204 | 1.544000298 | 5.54E-04 | 3.43E-03 |
| RERE_HUMAN\|Q9P2R6_S679 | 1.543633321 | 5.50E-03 | 2.13E-02 |
| SRRM1_HUMAN\|Q8IYB3_S756 | 1.543572676 | 8.57E-03 | 4.77E-02 |
| CPSF4_HUMAN\|O95639_S200_S212 | 1.543437573 | 2.10E-04 | 2.91E-03 |
| AHNK2_HUMAN\|Q8IVF2_S1583 | 1.543311552 | 2.52E-03 | 2.73E-02 |
| APC_HUMAN\|P25054_S2242_S2244 | 1.543226358 | 1.29E-05 | 6.95E-04 |
| K0930_HUMAN\|Q6ICG6_S267 | 1.543130777 | 2.27E-04 | 2.55E-03 |
| K1C17_HUMAN\|Q04695_T9 | 1.543119336 | 1.55E-03 | 1.69E-02 |
| PAK1_HUMAN\|Q13153_S223 | 1.543112207 | 9.86E-04 | 1.55E-02 |
| KC1A_HUMAN\|P48729-3_S312 | 1.543067382 | 3.51E-03 | 3.44E-02 |
| CSN3_HUMAN\|Q9UNS2_S421 | 1.54294282 | 6.04E-06 | 3.99E-04 |
| PLEC_HUMAN\|Q15149_S4673 | 1.542538642 | 2.73E-04 | 3.52E-02 |

| | | | |
|---|---|---|---|
| EPN2_HUMAN\|O95208_S173 | 1.542247385 | 8.84E-04 | 7.21E-03 |
| UBP20_HUMAN\|Q9Y2K6_S132_S134 | 1.542244292 | 1.14E-02 | 4.50E-02 |
| SF3B2_HUMAN\|Q13435_S303_S309 | 1.542110467 | 1.35E-02 | 4.41E-02 |
| 3BP1_HUMAN\|Q9Y3L3_T601 | 1.542060316 | 6.15E-03 | 3.11E-02 |
| K1C16_HUMAN\|P08779_S468 | 1.542039881 | 2.88E-04 | 4.72E-03 |
| LAD1_HUMAN\|O00515_S498 | 1.541906574 | 4.21E-03 | 3.50E-02 |
| CALD1_HUMAN\|Q05682_S789 | 1.541885952 | 9.03E-04 | 1.46E-02 |
| PALLD_HUMAN\|Q8WX93_S726 | 1.541861526 | 2.82E-03 | 2.95E-02 |
| IRS2_HUMAN\|Q9Y4H2_S608 | 1.541846331 | 7.65E-05 | 1.98E-03 |
| K2C5_HUMAN\|P13647_S16 | 1.541846016 | 1.50E-03 | 1.65E-02 |
| PKP2_HUMAN\|Q99959_S132 | 1.541827794 | 1.34E-03 | 1.53E-02 |
| ATX2L_HUMAN\|Q8WWM7_S680_S684 | 1.541658535 | 1.36E-05 | 1.68E-03 |
| PDLI2_HUMAN\|Q96JY6_S137 | 1.541623271 | 1.06E-04 | 2.92E-03 |
| PKP3_HUMAN\|Q9Y446_S280 | 1.5413115 | 3.56E-03 | 3.13E-02 |
| PSF2_HUMAN\|Q9Y248_S179_S182 | 1.541179241 | 3.92E-04 | 3.90E-03 |
| PEX1_HUMAN\|O43933_S1181 | 1.541162902 | 9.04E-04 | 5.08E-03 |
| MTCL1_HUMAN\|Q9Y4B5_S196 | 1.541149128 | 7.51E-04 | 1.05E-02 |
| ZN185_HUMAN\|O15231_S465 | 1.540952557 | 6.43E-03 | 4.74E-02 |
| MAP1B_HUMAN\|P46821_S1793_S1797 | 1.540613535 | 8.39E-05 | 2.57E-03 |
| SVIL_HUMAN\|O95425_T679 | 1.540566458 | 1.35E-05 | 2.86E-04 |
| OTUL_HUMAN\|Q96BN8_S351 | 1.540477573 | 4.59E-06 | 1.82E-04 |
| PPM1G_HUMAN\|O15355_S201 | 1.540428726 | 7.56E-03 | 3.64E-02 |
| PDS5B_HUMAN\|Q9NTI5_S1165 | 1.540395701 | 1.70E-03 | 1.37E-02 |
| SPR2E_HUMAN\|P22531_S60 | 1.540277687 | 7.16E-03 | 3.50E-02 |
| TRM1_HUMAN\|Q9NXH9_S625 | 1.540256776 | 7.85E-04 | 1.25E-02 |
| TGS1_HUMAN\|Q96RS0_S55_T60 | 1.540068745 | 1.14E-05 | 4.32E-04 |
| PDLI2_HUMAN\|Q96JY6_S137 | 1.539863262 | 1.09E-04 | 3.42E-03 |
| ADRM1_HUMAN\|Q16186_S211 | 1.539842411 | 5.27E-04 | 7.38E-03 |
| MA7D1_HUMAN\|Q3KQU3_S254 | 1.539837078 | 5.11E-05 | 1.48E-03 |
| CPSF4_HUMAN\|O95639_S200 | 1.53981515 | 1.71E-04 | 2.47E-03 |
| SRRM1_HUMAN\|Q8IYB3_S740_S743 | 1.53970133 | 5.44E-04 | 5.08E-03 |

| | | | |
|---|---|---|---|
| DKC1_HUMAN\|O60832_S485_T496 | 1.539638568 | 1.16E-05 | 4.37E-04 |
| K1671_HUMAN\|Q9BY89_S404 | 1.539549601 | 2.64E-05 | 4.76E-04 |
| K2C5_HUMAN\|P13647_S21_T24 | 1.539534634 | 2.49E-03 | 2.42E-02 |
| RHG01_HUMAN\|Q07960_S51 | 1.539518085 | 2.46E-04 | 1.83E-03 |
| ASPP2_HUMAN\|Q13625_S558 | 1.539471414 | 4.17E-05 | 8.51E-04 |
| CLAP1_HUMAN\|Q7Z460_S646_S649 | 1.539250617 | 6.23E-04 | 6.57E-03 |
| K1C14_HUMAN\|P02533_S432_S433_S437 | 1.539203092 | 2.55E-03 | 2.46E-02 |
| KI21A_HUMAN\|Q7Z4S6_S855 | 1.539065766 | 2.40E-04 | 5.31E-03 |
| CCNH_HUMAN\|P51946_T315 | 1.539002426 | 1.17E-03 | 1.45E-02 |
| SRC8_HUMAN\|Q14247_T401 | 1.538982344 | 1.22E-04 | 1.55E-03 |
| SVIL_HUMAN\|O95425-2_S263_S270 | 1.538660496 | 4.78E-04 | 9.21E-03 |
| TBC9B_HUMAN\|Q66K14_S1241 | 1.538547795 | 1.45E-05 | 1.99E-04 |
| K1C17_HUMAN\|Q04695_S422 | 1.538340532 | 1.98E-03 | 2.31E-02 |
| K1C14_HUMAN\|P02533_S432 | 1.538305787 | 2.53E-03 | 2.45E-02 |
| MAP1B_HUMAN\|P46821_S1797_Y1796 | 1.538283343 | 1.58E-03 | 2.05E-02 |
| AHNK2_HUMAN\|Q8IVF2_S457 | 1.538280109 | 8.82E-03 | 4.11E-02 |
| IF4B_HUMAN\|P23588_S497_S504_T500 | 1.538255115 | 9.13E-04 | 9.27E-03 |
| MISP_HUMAN\|Q8IVT2_S575 | 1.53813635 | 3.98E-04 | 8.19E-03 |
| VPS50_HUMAN\|Q96JG6_S559_S561 | 1.538012594 | 2.82E-03 | 1.27E-02 |
| ELAV1_HUMAN\|Q15717_S202 | 1.537847021 | 2.78E-03 | 1.74E-02 |
| K1C14_HUMAN\|P02533_S11 | 1.537823609 | 1.04E-04 | 2.45E-03 |
| DESP_HUMAN\|P15924_S165 | 1.537769833 | 2.96E-03 | 1.32E-02 |
| NEB2_HUMAN\|Q96SB3_S99 | 1.5375649 | 3.37E-04 | 1.47E-02 |
| FA83G_HUMAN\|A6ND36_S726 | 1.537518644 | 1.02E-03 | 1.46E-02 |
| ZCCHV_HUMAN\|Q7Z2W4_S378 | 1.537368481 | 6.80E-06 | 3.73E-04 |
| BAD_HUMAN\|Q92934_S74 | 1.537143137 | 5.40E-04 | 5.94E-03 |
| K1C17_HUMAN\|Q04695_S12 | 1.537065075 | 1.64E-03 | 1.79E-02 |
| ZN827_HUMAN\|Q17R98_S963_S984_T959 | 1.536986495 | 7.66E-04 | 1.06E-02 |
| PSF2_HUMAN\|Q9Y248_S182 | 1.536937401 | 1.42E-03 | 1.65E-02 |
| ZCH18_HUMAN\|Q86VM9_S110_S95 | 1.536925974 | 2.47E-03 | 1.82E-02 |
| LMNA_HUMAN\|P02545_S12 | 1.536899547 | 1.01E-03 | 1.29E-02 |

| | | | |
|---|---|---|---|
| ARG35_HUMAN\|A5YM69_S450 | 1.536873984 | 2.72E-03 | 3.10E-02 |
| K2C5_HUMAN\|P13647_S572 | 1.536546876 | 6.83E-04 | 9.10E-03 |
| MAP4_HUMAN\|P27816_S928 | 1.536451291 | 3.49E-04 | 7.13E-03 |
| DCA10_HUMAN\|Q5QP82_S350 | 1.536211746 | 5.79E-04 | 1.08E-02 |
| NOP56_HUMAN\|O00567_S570 | 1.535968423 | 2.46E-06 | 1.13E-04 |
| MA7D1_HUMAN\|Q3KQU3_S254 | 1.535941011 | 5.52E-05 | 1.38E-03 |
| NU214_HUMAN\|P35658_S1023 | 1.53536725 | 8.45E-04 | 1.13E-02 |
| CAMP2_HUMAN\|Q08AD1_S980 | 1.535349362 | 1.34E-03 | 1.17E-02 |
| GOGA4_HUMAN\|Q13439_S89_S92 | 1.535315983 | 3.70E-04 | 3.73E-03 |
| SCAM2_HUMAN\|O15127_S319 | 1.535314285 | 3.53E-03 | 3.65E-02 |
| E41L2_HUMAN\|O43491_S87 | 1.535239784 | 2.74E-04 | 1.32E-02 |
| DC1L1_HUMAN\|Q9Y6G9_T513_T515 | 1.534488341 | 1.83E-04 | 4.82E-03 |
| AKTS1_HUMAN\|Q96B36_S202_S203 | 1.534387741 | 3.82E-05 | 4.29E-04 |
| OPTN_HUMAN\|Q96CV9_S177 | 1.53438384 | 2.33E-03 | 1.54E-02 |
| BAIP2_HUMAN\|Q9UQB8-4_T520 | 1.53436535 | 1.11E-03 | 1.31E-02 |
| ZN768_HUMAN\|Q9H5H4_S83 | 1.534305945 | 2.11E-03 | 2.19E-02 |
| PHLB2_HUMAN\|Q86SQ0_S334_T328 | 1.533981663 | 4.05E-03 | 2.30E-02 |
| RB3GP_HUMAN\|Q15042_S537 | 1.533814615 | 4.04E-05 | 8.37E-04 |
| COHA1_HUMAN\|Q9UMD9_S78 | 1.53373408 | 5.94E-03 | 4.57E-02 |
| K1C16_HUMAN\|P08779_S449 | 1.533670148 | 1.38E-03 | 1.03E-02 |
| MICA3_HUMAN\|Q7RTP6_S977 | 1.533612071 | 2.05E-06 | 6.96E-05 |
| K1C14_HUMAN\|P02533_S33 | 1.533603853 | 4.94E-04 | 7.07E-03 |
| LIMA1_HUMAN\|Q9UHB6_S15 | 1.533539969 | 1.40E-03 | 1.63E-02 |
| SSFA2_HUMAN\|P28290_S668 | 1.533369828 | 1.82E-03 | 1.53E-02 |
| ZO1_HUMAN\|Q07157_S841_Y853 | 1.533345485 | 3.41E-03 | 3.56E-02 |
| LMNB1_HUMAN\|P20700_T20 | 1.533283204 | 3.56E-03 | 3.67E-02 |
| TANC2_HUMAN\|Q9HCD6_S1827 | 1.533280864 | 5.33E-05 | 5.39E-04 |
| CSCL2_HUMAN\|Q5T3F8_S114_T110 | 1.532974818 | 6.68E-04 | 1.18E-02 |
| MARK2_HUMAN\|Q7KZI7-13_S376 | 1.532846822 | 7.06E-03 | 3.99E-02 |
| SF3B2_HUMAN\|Q13435_S307 | 1.532699386 | 4.22E-03 | 4.21E-02 |
| PKHG3_HUMAN\|A1L390_S1081 | 1.532645385 | 1.94E-05 | 4.94E-04 |

| | | | |
|---|---|---|---|
| RPP25_HUMAN\|Q9BUL9_S162 | 1.532528311 | 4.15E-04 | 6.17E-03 |
| IASPP_HUMAN\|Q8WUF5_S134 | 1.532425173 | 1.16E-03 | 3.17E-02 |
| K1671_HUMAN\|Q9BY89_S127 | 1.532350461 | 1.26E-04 | 1.59E-03 |
| AHNK_HUMAN\|Q09666_T4564 | 1.532339677 | 8.52E-04 | 7.03E-03 |
| K1C14_HUMAN\|P02533_S28 | 1.532086392 | 3.37E-04 | 6.95E-03 |
| COBL1_HUMAN\|Q53SF7_S1146 | 1.531732324 | 2.97E-04 | 1.35E-02 |
| CCD86_HUMAN\|Q9H6F5_S217 | 1.531722492 | 1.83E-03 | 2.24E-02 |
| LMNA_HUMAN\|P02545_S414 | 1.531560923 | 4.90E-04 | 9.12E-03 |
| SRC8_HUMAN\|Q14247_T399_T411 | 1.531534143 | 2.07E-04 | 1.60E-03 |
| RHG01_HUMAN\|Q07960_S51 | 1.531450631 | 2.81E-04 | 6.04E-03 |
| NIBL1_HUMAN\|Q96TA1_S696 | 1.53144426 | 3.02E-04 | 3.84E-03 |
| NKTR_HUMAN\|P30414_S463 | 1.531305261 | 6.80E-04 | 1.63E-02 |
| GGA2_HUMAN\|Q9UJY4_S177 | 1.531226123 | 1.30E-03 | 1.21E-02 |
| TCF20_HUMAN\|Q9UGU0_S538 | 1.531058823 | 4.01E-04 | 2.85E-03 |
| H2AX_HUMAN\|P16104_S140 | 1.531054944 | 4.68E-05 | 1.84E-03 |
| PLEC_HUMAN\|Q15149_S4673_S4675 | 1.530948951 | 1.31E-03 | 1.50E-02 |
| K1C14_HUMAN\|P02533_S33 | 1.53087104 | 5.17E-04 | 9.53E-03 |
| MTCL1_HUMAN\|Q9Y4B5_S196 | 1.530752753 | 8.70E-04 | 7.14E-03 |
| AHNK_HUMAN\|Q09666_S216_S220 | 1.530581922 | 6.98E-05 | 2.35E-03 |
| PDLI2_HUMAN\|Q96JY6_S127_S129 | 1.53030267 | 2.84E-03 | 3.20E-02 |
| PALLD_HUMAN\|Q8WX93_S1104 | 1.529917617 | 1.49E-04 | 9.65E-03 |
| PHF8_HUMAN\|Q9UPP1_S804 | 1.529802311 | 9.35E-05 | 9.12E-04 |
| PKP3_HUMAN\|Q9Y446_S180 | 1.529695235 | 7.84E-04 | 1.25E-02 |
| PACN3_HUMAN\|Q9UKS6_S344_S354 | 1.529442594 | 1.77E-04 | 4.76E-03 |
| LAD1_HUMAN\|O00515_S394 | 1.529392236 | 3.09E-04 | 6.34E-03 |
| VPS50_HUMAN\|Q96JG6_S559_S561 | 1.529284758 | 3.13E-03 | 1.55E-02 |
| EPIPL_HUMAN\|P58107_S2716 | 1.529189081 | 9.39E-06 | 1.44E-04 |
| ZFP91_HUMAN\|Q96JP5_S82 | 1.529180246 | 3.99E-04 | 1.64E-02 |
| SCEL_HUMAN\|O95171_S91 | 1.529163545 | 5.69E-03 | 2.93E-02 |
| S6A15_HUMAN\|Q9H2J7_S699 | 1.528973465 | 6.40E-03 | 3.72E-02 |
| DLGP4_HUMAN\|Q9Y2H0_S973 | 1.528836519 | 2.54E-03 | 1.17E-02 |

| | | | |
|---|---|---|---|
| PALLD_HUMAN\|Q8WX93_S893 | 1.528420329 | 8.30E-04 | 1.39E-02 |
| DESP_HUMAN\|P15924_S22 | 1.527902541 | 3.36E-03 | 2.02E-02 |
| ASPP2_HUMAN\|Q13625_S556 | 1.527778775 | 5.63E-04 | 1.04E-02 |
| EI2BE_HUMAN\|Q13144_S544 | 1.527748904 | 4.07E-04 | 2.89E-03 |
| IASPP_HUMAN\|Q8WUF5_S332 | 1.527723985 | 6.45E-04 | 5.76E-03 |
| FKBP5_HUMAN\|Q13451_T443 | 1.527683585 | 7.18E-03 | 4.03E-02 |
| K1C14_HUMAN\|P02533_S51 | 1.527455562 | 3.58E-03 | 3.14E-02 |
| NUCKS_HUMAN\|Q9H1E3_S229_S240 | 1.527311883 | 5.12E-04 | 3.52E-03 |
| WAC2A_HUMAN\|Q641Q2_S284 | 1.527193981 | 2.79E-03 | 3.07E-02 |
| FNBP4_HUMAN\|Q8N3X1_S464 | 1.527055392 | 1.67E-03 | 2.15E-02 |
| PP12C_HUMAN\|Q9BZL4_S453 | 1.526465291 | 8.85E-06 | 4.44E-04 |
| IRS2_HUMAN\|Q9Y4H2_S560 | 1.526419466 | 3.92E-05 | 1.22E-03 |
| PKP2_HUMAN\|Q99959_S132 | 1.526221226 | 1.65E-03 | 1.18E-02 |
| AHNK_HUMAN\|Q09666_S210_S216 | 1.526220152 | 2.47E-04 | 4.69E-03 |
| SRRM1_HUMAN\|Q8IYB3_S754 | 1.526168158 | 4.97E-03 | 3.06E-02 |
| S6A15_HUMAN\|Q9H2J7_S699 | 1.525689755 | 6.64E-03 | 3.97E-02 |
| TACC2_HUMAN\|O95359_S2512 | 1.525582194 | 2.60E-03 | 1.19E-02 |
| PKP3_HUMAN\|Q9Y446_S123 | 1.525569571 | 1.57E-03 | 1.13E-02 |
| K1C16_HUMAN\|P08779_S466 | 1.525494368 | 9.43E-04 | 1.15E-02 |
| TLN1_HUMAN\|Q9Y490_S1641 | 1.525452264 | 2.89E-05 | 6.52E-04 |
| BCL9_HUMAN\|O00512_T315 | 1.525139047 | 1.35E-03 | 3.53E-02 |
| BIG1_HUMAN\|Q9Y6D6_S394 | 1.52490193 | 2.87E-04 | 3.07E-03 |
| BICD2_HUMAN\|Q8TD16_S615 | 1.524698424 | 1.68E-04 | 4.60E-03 |
| BRAF_HUMAN\|P15056_S729 | 1.524680971 | 5.10E-03 | 4.74E-02 |
| GIT1_HUMAN\|Q9Y2X7_S388 | 1.52447154 | 7.66E-03 | 2.78E-02 |
| AAPK1_HUMAN\|Q13131_S508 | 1.524400451 | 4.72E-06 | 1.68E-04 |
| FIL1L_HUMAN\|Q4L180_T986 | 1.524177538 | 6.81E-04 | 1.14E-02 |
| MKL2_HUMAN\|Q9ULH7_S921 | 1.524117187 | 4.48E-04 | 5.13E-03 |
| K1C14_HUMAN\|P02533_S437 | 1.523890487 | 4.02E-03 | 3.41E-02 |
| KDM4A_HUMAN\|O75164_S523 | 1.523821808 | 1.01E-02 | 4.57E-02 |
| LARP1_HUMAN\|Q6PKG0_S627_Y633 | 1.523696384 | 3.55E-03 | 2.09E-02 |

| | | | |
|---|---|---|---|
| DDX10_HUMAN\|Q13206_S539 | 1.523542161 | 1.28E-03 | 1.19E-02 |
| EGFR_HUMAN\|P00533_S1204 | 1.523389938 | 4.34E-03 | 3.98E-02 |
| AT11C_HUMAN\|Q8NB49_T1124 | 1.523353863 | 3.26E-04 | 7.16E-03 |
| CBX4_HUMAN\|O00257_S182 | 1.523342547 | 1.29E-03 | 9.72E-03 |
| LAD1_HUMAN\|O00515_S394 | 1.523314139 | 3.43E-04 | 1.47E-02 |
| C2CD5_HUMAN\|Q86YS7_S307 | 1.523187919 | 3.79E-04 | 1.57E-02 |
| AHNK_HUMAN\|Q09666_S212 | 1.523154683 | 1.54E-03 | 2.15E-02 |
| KDM3B_HUMAN\|Q7LBC6_S293 | 1.523150258 | 1.54E-05 | 9.19E-04 |
| YAP1_HUMAN\|P46937_T119 | 1.522970915 | 3.95E-03 | 3.73E-02 |
| ADCY6_HUMAN\|O43306_S54 | 1.522793547 | 3.88E-04 | 5.89E-03 |
| HNRL2_HUMAN\|Q1KMD3_S161_T165 | 1.522636712 | 9.61E-03 | 3.90E-02 |
| IPP2M_HUMAN\|Q6NXS1_S121 | 1.522531748 | 1.94E-03 | 2.34E-02 |
| EHBP1_HUMAN\|Q8NDI1_S1035 | 1.522485 | 2.06E-03 | 1.10E-02 |
| PAXI_HUMAN\|P49023_S126_S130 | 1.522477987 | 5.26E-03 | 2.07E-02 |
| PPM1G_HUMAN\|O15355_S183 | 1.522206879 | 3.39E-04 | 3.50E-03 |
| CPNE1_HUMAN\|Q99829_S55 | 1.522193827 | 8.75E-04 | 1.32E-02 |
| SRRM1_HUMAN\|Q8IYB3_S752_S754_S756 | 1.52215455 | 5.21E-05 | 2.00E-03 |
| BRE1A_HUMAN\|Q5VTR2_S136 | 1.522124397 | 2.45E-05 | 5.81E-04 |
| OSTF1_HUMAN\|Q92882_S213 | 1.521992546 | 6.75E-04 | 1.63E-02 |
| MAP7_HUMAN\|Q14244_S183 | 1.521725511 | 8.33E-05 | 2.61E-03 |
| FA76B_HUMAN\|Q5HYJ3_S338 | 1.521609331 | 1.07E-03 | 1.33E-02 |
| E41L2_HUMAN\|O43491_S87 | 1.521419636 | 3.44E-04 | 5.37E-03 |
| MA7D1_HUMAN\|Q3KQU3_S254 | 1.521353677 | 7.40E-05 | 5.16E-03 |
| SIPA1_HUMAN\|Q96FS4_S1036 | 1.521150618 | 4.50E-04 | 8.93E-03 |
| BUD13_HUMAN\|Q9BRD0_T159 | 1.520930214 | 6.49E-04 | 6.77E-03 |
| LMNB2_HUMAN\|Q03252_S424 | 1.520801215 | 1.48E-03 | 1.67E-02 |
| K1C17_HUMAN\|Q04695_S11 | 1.520792676 | 6.32E-04 | 9.19E-03 |
| SMG9_HUMAN\|Q9H0W8_T56 | 1.520678627 | 2.12E-04 | 4.18E-03 |
| TISB_HUMAN\|Q07352_S92 | 1.520592062 | 1.99E-03 | 2.32E-02 |
| ARAF_HUMAN\|P10398_S257_T253 | 1.520520499 | 7.39E-04 | 2.37E-02 |
| KHDR1_HUMAN\|Q07666_S20 | 1.520398652 | 1.60E-04 | 2.36E-03 |

| | | | |
|---|---|---|---|
| ACAP2_HUMAN\|Q15057_T382 | 1.520389914 | 1.02E-05 | 7.16E-04 |
| K1C16_HUMAN\|P08779_S448 | 1.520280025 | 9.92E-04 | 1.20E-02 |
| CY561_HUMAN\|P49447_S250 | 1.520247248 | 2.65E-03 | 3.07E-02 |
| SEPT7_HUMAN\|Q16181_S423 | 1.520238031 | 4.11E-03 | 3.83E-02 |
| NU214_HUMAN\|P35658_S1023 | 1.520154322 | 1.06E-03 | 9.89E-03 |
| P3C2A_HUMAN\|O00443_S259 | 1.520144735 | 2.06E-04 | 2.91E-03 |
| MK01_HUMAN\|P28482_Y187 | 1.519768679 | 1.35E-03 | 1.01E-02 |
| MON1A_HUMAN\|Q86VX9_S95 | 1.51972392 | 1.38E-04 | 2.13E-03 |
| PTN13_HUMAN\|Q12923_S908 | 1.519707628 | 7.96E-04 | 1.02E-02 |
| RAI14_HUMAN\|Q9P0K7_T297 | 1.519649711 | 2.59E-03 | 2.56E-02 |
| KLRG2_HUMAN\|A4D1S0_S215 | 1.519439593 | 1.54E-03 | 7.86E-03 |
| INADL_HUMAN\|Q8NI35_S645 | 1.519439281 | 9.92E-05 | 1.29E-03 |
| IF4B_HUMAN\|P23588_S497_S498_S504 | 1.519346472 | 9.45E-05 | 1.52E-03 |
| CAMP3_HUMAN\|Q9P1Y5_S756 | 1.519100044 | 2.58E-04 | 2.83E-03 |
| ZCHC8_HUMAN\|Q6NZY4_S500_T492_T496 | 1.519039189 | 9.67E-04 | 7.75E-03 |
| HSPB1_HUMAN\|P04792_S82 | 1.518977318 | 3.08E-05 | 6.85E-04 |
| K1C17_HUMAN\|Q04695_S39 | 1.518615728 | 4.56E-03 | 2.51E-02 |
| K1C17_HUMAN\|Q04695_S11 | 1.518558954 | 6.56E-04 | 1.11E-02 |
| KLC2_HUMAN\|Q9H0B6_S608_S611 | 1.518514243 | 3.38E-05 | 1.12E-03 |
| PKP2_HUMAN\|Q99959_S151 | 1.518278221 | 1.81E-04 | 2.56E-03 |
| USO1_HUMAN\|O60763_S942_S952 | 1.51819709 | 3.72E-03 | 3.57E-02 |
| TX264_HUMAN\|Q9Y6I9_S244 | 1.518069985 | 6.14E-03 | 3.59E-02 |
| TACC2_HUMAN\|O95359_S2512 | 1.518044727 | 2.86E-03 | 3.26E-02 |
| ZN185_HUMAN\|O15231_S153 | 1.517974878 | 1.45E-04 | 2.20E-03 |
| K1C14_HUMAN\|P02533_Y46 | 1.517812372 | 1.30E-03 | 1.73E-02 |
| CF132_HUMAN\|Q5T0Z8_T297 | 1.51772018 | 4.43E-03 | 2.47E-02 |
| YAP1_HUMAN\|P46937_T110_T119 | 1.51760267 | 2.39E-03 | 4.72E-02 |
| PRC2A_HUMAN\|P48634_S1147 | 1.517512184 | 6.25E-05 | 1.70E-03 |
| MA7D1_HUMAN\|Q3KQU3_S254 | 1.517504376 | 7.99E-05 | 2.52E-03 |
| PRC2A_HUMAN\|P48634_S1085_S1089_S1092 | 1.517257037 | 1.51E-04 | 1.83E-03 |
| NASP_HUMAN\|P49321_S421 | 1.517250347 | 5.38E-05 | 2.05E-03 |

| | | | |
|---|---|---|---|
| VIME_HUMAN\|P08670_S34 | 1.517196718 | 6.59E-04 | 9.46E-03 |
| PHLB2_HUMAN\|Q86SQ0_S330_S334 | 1.517118366 | 5.11E-03 | 2.74E-02 |
| JHD2C_HUMAN\|Q15652_S1058 | 1.517037379 | 3.93E-03 | 2.24E-02 |
| TP53B_HUMAN\|Q12888_S1462_S1481 | 1.517030008 | 5.22E-03 | 2.06E-02 |
| RFIP1_HUMAN\|Q6WKZ4_S501 | 1.517014486 | 5.55E-03 | 4.35E-02 |
| ZO1_HUMAN\|Q07157_S1617 | 1.516985359 | 1.79E-04 | 4.65E-03 |
| CAAP1_HUMAN\|Q9H8G2_S89 | 1.516847068 | 1.13E-05 | 1.67E-04 |
| TB182_HUMAN\|Q9C0C2_S920 | 1.516715154 | 8.92E-04 | 5.03E-03 |
| GRB10_HUMAN\|Q13322_S104 | 1.516675352 | 3.55E-05 | 6.01E-04 |
| PDS5B_HUMAN\|Q9NTI5_S1165 | 1.516588342 | 2.34E-03 | 2.68E-02 |
| MY18A_HUMAN\|Q92614_S1998 | 1.516387289 | 1.59E-04 | 2.38E-03 |
| SCAF8_HUMAN\|Q9UPN6_S617 | 1.516327284 | 4.68E-04 | 4.47E-03 |
| MMTA2_HUMAN\|Q9BU76_S216_S220 | 1.516213315 | 5.01E-04 | 4.73E-03 |
| K1C17_HUMAN\|Q04695_S24 | 1.516208315 | 5.59E-04 | 7.77E-03 |
| HNRPM_HUMAN\|P52272_S575 | 1.515945403 | 2.96E-03 | 3.20E-02 |
| MAP7_HUMAN\|Q14244_S315 | 1.515803152 | 2.57E-04 | 5.64E-03 |
| UBP47_HUMAN\|Q96K76_S910 | 1.515310117 | 4.78E-03 | 1.93E-02 |
| GGYF2_HUMAN\|Q6Y7W6_S201 | 1.515291554 | 4.90E-04 | 4.69E-02 |
| HNRL2_HUMAN\|Q1KMD3_S161_S168_T165 | 1.51525865 | 3.50E-04 | 3.57E-03 |
| DESP_HUMAN\|P15924_S2815_S2820 | 1.515150983 | 1.26E-04 | 1.93E-03 |
| TBCD5_HUMAN\|Q92609_S541 | 1.515129162 | 2.15E-06 | 9.26E-05 |
| PLEC_HUMAN\|Q15149_S4672 | 1.514780692 | 2.67E-03 | 2.85E-02 |
| IRS1_HUMAN\|P35568_S1101 | 1.514705157 | 1.15E-04 | 2.60E-03 |
| DESP_HUMAN\|P15924_S165 | 1.514634455 | 3.92E-03 | 2.73E-02 |
| K1C14_HUMAN\|P02533_S28 | 1.514324929 | 4.58E-04 | 6.66E-03 |
| EMAL4_HUMAN\|Q9HC35_S981 | 1.514324023 | 9.30E-04 | 1.15E-02 |
| DEN1A_HUMAN\|Q8TEH3_S536 | 1.514220173 | 3.74E-04 | 2.70E-03 |
| VIME_HUMAN\|P08670_S65 | 1.51421028 | 2.30E-04 | 4.06E-03 |
| CCDC6_HUMAN\|Q16204_S244 | 1.513941217 | 1.64E-03 | 1.40E-02 |
| DESP_HUMAN\|P15924_S2815 | 1.513690089 | 1.33E-04 | 1.21E-03 |
| ZCH18_HUMAN\|Q86VM9_S110_S95 | 1.513682652 | 3.29E-03 | 4.73E-02 |

| | | | |
|---|---|---|---|
| HMGA2_HUMAN\|P52926_S101_S105_T100 | 1.513635854 | 1.36E-03 | 1.01E-02 |
| CPNE1_HUMAN\|Q99829_S55 | 1.51360304 | 9.91E-04 | 7.90E-03 |
| AHNK_HUMAN\|Q09666_S135 | 1.513533397 | 3.12E-03 | 2.26E-02 |
| SCAF8_HUMAN\|Q9UPN6_S617 | 1.513236833 | 4.92E-04 | 1.81E-02 |
| PLEC_HUMAN\|Q15149_S4618_S4626_T4628 | 1.513202309 | 5.36E-04 | 3.65E-03 |
| K1671_HUMAN\|Q9BY89_S1441 | 1.513181425 | 2.71E-03 | 1.96E-02 |
| ZFN2B_HUMAN\|Q8WV99_S183 | 1.512990664 | 7.95E-04 | 1.31E-02 |
| M3K20_HUMAN\|Q9NYL2_S633 | 1.512983428 | 5.15E-03 | 2.75E-02 |
| NHSL1_HUMAN\|Q5SYE7_S722_S723 | 1.512975879 | 2.67E-03 | 1.22E-02 |
| IF4B_HUMAN\|P23588_S495_S497 | 1.512903773 | 9.89E-05 | 3.19E-03 |
| HELZ2_HUMAN\|Q9BYK8_S1006 | 1.512895374 | 3.78E-03 | 3.85E-02 |
| FRYL_HUMAN\|O94915_S1914_S1935 | 1.512824303 | 3.89E-03 | 2.23E-02 |
| TREF1_HUMAN\|Q96PN7_S758 | 1.512705802 | 1.07E-02 | 3.65E-02 |
| NIBL1_HUMAN\|Q96TA1_S696 | 1.512624918 | 4.08E-04 | 6.09E-03 |
| CBX3_HUMAN\|Q13185_S176 | 1.512337656 | 1.59E-03 | 1.32E-02 |
| IP3KB_HUMAN\|P27987_S43 | 1.512244167 | 1.95E-04 | 4.93E-03 |
| EPIPL_HUMAN\|P58107_S2716 | 1.512105687 | 1.38E-05 | 8.91E-04 |
| PKHA2_HUMAN\|Q9HB19_S184 | 1.511941095 | 4.13E-05 | 1.12E-03 |
| EGFR_HUMAN\|P00533_S1042 | 1.511807187 | 8.56E-03 | 3.06E-02 |
| NELFE_HUMAN\|P18615_S51 | 1.511754653 | 2.86E-04 | 6.03E-03 |
| LAP2A_HUMAN\|P42166_T74 | 1.511467501 | 2.91E-03 | 2.80E-02 |
| COHA1_HUMAN\|Q9UMD9_S85_S93_S96 | 1.511458857 | 7.17E-04 | 6.16E-03 |
| ADCY6_HUMAN\|O43306_S54 | 1.511425617 | 4.69E-04 | 4.47E-03 |
| LAD1_HUMAN\|O00515_T41 | 1.51105097 | 4.71E-03 | 4.19E-02 |
| WDR44_HUMAN\|Q5JSH3_S561 | 1.510900469 | 2.13E-04 | 1.63E-03 |
| RHG01_HUMAN\|Q07960_S51 | 1.510894513 | 3.96E-04 | 1.12E-02 |
| RGL2_HUMAN\|O15211_S761 | 1.510791203 | 4.89E-04 | 1.89E-02 |
| MTCL1_HUMAN\|Q9Y4B5_S776 | 1.510782529 | 6.89E-05 | 2.33E-03 |
| ACINU_HUMAN\|Q9UKV3_T563 | 1.510757664 | 2.34E-05 | 5.70E-04 |
| K1C14_HUMAN\|P02533_S462 | 1.510725873 | 2.96E-03 | 2.10E-02 |
| SP100_HUMAN\|P23497_S476 | 1.510645346 | 1.58E-03 | 8.00E-03 |

| | | | |
|---|---|---|---|
| TRI29_HUMAN\|Q14134_S152 | 1.510417738 | 5.73E-04 | 2.07E-02 |
| K1C17_HUMAN\|Q04695_S28 | 1.510294524 | 2.30E-03 | 4.61E-02 |
| CND1_HUMAN\|Q15021_S1333 | 1.51027257 | 3.38E-04 | 7.25E-03 |
| CTNA1_HUMAN\|P35221_S641_T645 | 1.510156723 | 9.33E-04 | 1.15E-02 |
| K2C75_HUMAN\|O95678_S535 | 1.509920892 | 4.58E-05 | 1.82E-03 |
| STXB5_HUMAN\|Q5T5C0-2_S712 | 1.5097198 | 5.27E-05 | 1.51E-03 |
| ACINU_HUMAN\|Q9UKV3_T254 | 1.509709882 | 4.51E-03 | 3.77E-02 |
| ITA3_HUMAN\|P26006_T1044 | 1.509706805 | 5.64E-03 | 3.36E-02 |
| SRSF1_HUMAN\|Q07955_S199 | 1.509650595 | 1.05E-02 | 4.20E-02 |
| ATX2_HUMAN\|Q99700_T741 | 1.509460286 | 1.30E-06 | 4.88E-05 |
| FA76B_HUMAN\|Q5HYJ3_S338 | 1.509348365 | 1.28E-03 | 3.41E-02 |
| CAMP2_HUMAN\|Q08AD1_S1313 | 1.509256306 | 5.72E-04 | 5.29E-03 |
| ZYX_HUMAN\|Q15942_S259 | 1.509184498 | 1.81E-04 | 3.38E-03 |
| ZO2_HUMAN\|Q9UDY2_S170 | 1.509139121 | 1.13E-04 | 1.78E-03 |
| FA83H_HUMAN\|Q6ZRV2_S513_S516 | 1.509104808 | 3.91E-03 | 2.24E-02 |
| CLAP1_HUMAN\|Q7Z460_S646_S649 | 1.509078632 | 9.83E-04 | 1.19E-02 |
| DTX3L_HUMAN\|Q8TDB6_S202 | 1.509013306 | 1.54E-02 | 4.87E-02 |
| RRBP1_HUMAN\|Q9P2E9_S583 | 1.508916616 | 2.85E-03 | 1.77E-02 |
| K1C17_HUMAN\|Q04695_S10 | 1.508696176 | 1.44E-03 | 1.60E-02 |
| T200A_HUMAN\|Q86VY9_S350 | 1.508664235 | 5.04E-03 | 3.10E-02 |
| YBOX3_HUMAN\|P16989_S369_S370 | 1.508559678 | 6.76E-04 | 9.05E-03 |
| SRPK1_HUMAN\|Q96SB4_S39_S51 | 1.508287948 | 3.07E-03 | 1.53E-02 |
| HSPB1_HUMAN\|P04792_S82 | 1.508269499 | 3.88E-05 | 8.17E-04 |
| ACSA_HUMAN\|Q9NR19_S267 | 1.507818267 | 5.95E-04 | 5.43E-03 |
| FA83H_HUMAN\|Q6ZRV2_S903 | 1.507628233 | 9.06E-04 | 5.08E-03 |
| FLVC1_HUMAN\|Q9Y5Y0_S536 | 1.507626944 | 2.03E-03 | 2.34E-02 |
| K1C14_HUMAN\|P02533_S432_S435 | 1.507455972 | 5.20E-03 | 4.48E-02 |
| K1C14_HUMAN\|P02533_S432_S433_S438 | 1.507428903 | 4.24E-03 | 3.51E-02 |
| TB182_HUMAN\|Q9C0C2_S691 | 1.507392233 | 4.84E-04 | 5.45E-03 |
| K1C14_HUMAN\|P02533_S429_S432 | 1.507142817 | 3.50E-03 | 2.08E-02 |
| GYS1_HUMAN\|P13807_S727_S731 | 1.507133594 | 2.15E-03 | 1.73E-02 |

| | | | |
|---|---|---|---|
| IF4B_HUMAN\|P23588_S497_S504_T500 | 1.506987558 | 1.46E-03 | 2.01E-02 |
| RAI14_HUMAN\|Q9P0K7_S296 | 1.506822108 | 1.93E-03 | 2.01E-02 |
| EIF3A_HUMAN\|Q14152_S882 | 1.50665841 | 6.89E-03 | 4.99E-02 |
| GGYF1_HUMAN\|O75420_S638 | 1.50657628 | 5.31E-04 | 5.86E-03 |
| FA83H_HUMAN\|Q6ZRV2_T938 | 1.506572604 | 3.79E-03 | 3.33E-02 |
| CLIP4_HUMAN\|Q8N3C7_T582 | 1.506527524 | 4.06E-04 | 1.63E-02 |
| PPR18_HUMAN\|Q6NYC8_S209_S213 | 1.506439879 | 1.56E-04 | 1.89E-03 |
| FA83B_HUMAN\|Q5T0W9_S802 | 1.506427809 | 4.46E-04 | 8.48E-03 |
| OSBP1_HUMAN\|P22059_S379_S382_S386 | 1.506401424 | 4.64E-03 | 2.14E-02 |
| ARI1A_HUMAN\|O14497_S696_S698 | 1.506271531 | 1.00E-04 | 2.41E-03 |
| EGFR_HUMAN\|P00533_T693 | 1.506136184 | 3.22E-03 | 2.31E-02 |
| DOCK4_HUMAN\|Q8N1I0_S1620 | 1.506091572 | 1.24E-02 | 4.79E-02 |
| LMNA_HUMAN\|P02545_S303 | 1.505988732 | 5.27E-03 | 4.11E-02 |
| MACF1_HUMAN\|Q9UPN3_S7330_S7345 | 1.505803262 | 8.54E-04 | 7.03E-03 |
| TB182_HUMAN\|Q9C0C2_S1029 | 1.505660509 | 1.70E-04 | 1.47E-03 |
| RPP25_HUMAN\|Q9BUL9_S162 | 1.505636791 | 6.53E-04 | 1.12E-02 |
| SRRM1_HUMAN\|Q8IYB3_S752_S754_S756 | 1.505575403 | 7.39E-05 | 5.16E-03 |
| FIL1L_HUMAN\|Q4L180_S1049 | 1.505534699 | 4.92E-03 | 4.63E-02 |
| ROAA_HUMAN\|Q99729-3_S255 | 1.505494449 | 3.68E-03 | 1.57E-02 |
| SRSF1_HUMAN\|Q07955_S199 | 1.50501102 | 1.10E-02 | 3.74E-02 |
| CLAP1_HUMAN\|Q7Z460_S1091 | 1.504800832 | 1.52E-04 | 8.50E-03 |
| AAK1_HUMAN\|Q2M2I8_S678 | 1.504770559 | 4.96E-04 | 5.57E-03 |
| KDM3B_HUMAN\|Q7LBC6_S289 | 1.50473979 | 3.78E-04 | 2.55E-03 |
| M3K3_HUMAN\|Q99759_S337_S340 | 1.504675793 | 1.44E-04 | 1.20E-03 |
| SSRP1_HUMAN\|Q08945_Y438 | 1.504493657 | 4.82E-03 | 2.61E-02 |
| ZDHC5_HUMAN\|Q9C0B5_S621 | 1.50446601 | 5.86E-03 | 3.66E-02 |
| ATX2L_HUMAN\|Q8WWM7_S680_S684 | 1.504369358 | 3.09E-05 | 5.36E-04 |
| TPC12_HUMAN\|Q8WVT3_S184 | 1.504345751 | 3.22E-03 | 1.59E-02 |
| FAK1_HUMAN\|Q05397_S910 | 1.504307148 | 6.74E-04 | 1.19E-02 |
| DDX3X_HUMAN\|O00571_S90 | 1.504223634 | 1.04E-03 | 1.59E-02 |
| RABL6_HUMAN\|Q3YEC7_S492 | 1.504212885 | 1.20E-04 | 2.42E-03 |

| | | | |
|---|---|---|---|
| ARHG2_HUMAN\|Q92974_S886 | 1.504211006 | 6.12E-03 | 3.75E-02 |
| K2C5_HUMAN\|P13647_S50 | 1.504094285 | 1.97E-04 | 2.27E-03 |
| MTUS1_HUMAN\|Q9ULD2_T1204 | 1.504039514 | 4.52E-04 | 8.55E-03 |
| AAPK1_HUMAN\|Q13131_S508 | 1.503992484 | 7.63E-06 | 5.82E-04 |
| RPRD2_HUMAN\|Q5VT52_Y778 | 1.503886228 | 9.97E-04 | 2.08E-02 |
| TB10B_HUMAN\|Q4KMP7_S656 | 1.503826406 | 2.36E-03 | 2.40E-02 |
| KLC4_HUMAN\|Q9NSK0_S611 | 1.503820493 | 6.67E-04 | 9.52E-03 |
| SSH2_HUMAN\|Q76I76_S487 | 1.503738812 | 1.36E-05 | 2.89E-04 |
| IPP2M_HUMAN\|Q6NXS1_S121 | 1.503624261 | 2.50E-03 | 4.83E-02 |
| K2C5_HUMAN\|P13647_S16 | 1.503594956 | 2.54E-03 | 1.64E-02 |
| TFE3_HUMAN\|P19532_S556 | 1.503582596 | 1.36E-03 | 7.07E-03 |
| FA83H_HUMAN\|Q6ZRV2_S974 | 1.503571946 | 1.93E-04 | 4.99E-03 |
| PATL1_HUMAN\|Q86TB9_S179_S184 | 1.503563041 | 2.85E-03 | 2.75E-02 |
| CSN3_HUMAN\|Q9UNS2_S421 | 1.503507779 | 1.46E-05 | 4.01E-04 |
| FA83H_HUMAN\|Q6ZRV2_S523 | 1.503446906 | 9.14E-06 | 2.12E-04 |
| ZO1_HUMAN\|Q07157_S841_Y853 | 1.503425732 | 4.87E-03 | 3.03E-02 |
| SYNEM_HUMAN\|O15061_S1044_S1049 | 1.503285221 | 5.45E-04 | 9.94E-03 |
| FIL1L_HUMAN\|Q4L180_S1049 | 1.503237141 | 5.07E-03 | 4.43E-02 |
| NIBL1_HUMAN\|Q96TA1_S691 | 1.503177342 | 1.95E-05 | 2.46E-04 |
| COBL1_HUMAN\|Q53SF7_S364 | 1.503172041 | 2.77E-04 | 1.32E-02 |
| GRIN1_HUMAN\|Q7Z2K8_S615 | 1.502942772 | 3.88E-04 | 3.88E-03 |
| MK01_HUMAN\|P28482_Y187 | 1.502683248 | 1.71E-03 | 9.39E-03 |
| FA83H_HUMAN\|Q6ZRV2_S1009 | 1.502614659 | 2.94E-03 | 2.15E-02 |
| RAI14_HUMAN\|Q9P0K7_S290 | 1.50249455 | 4.93E-03 | 3.22E-02 |
| RABE1_HUMAN\|Q15276_S410_T408 | 1.502406967 | 1.94E-04 | 4.60E-03 |
| T2FA_HUMAN\|P35269_S433 | 1.502344923 | 9.49E-05 | 6.05E-03 |
| TB22B_HUMAN\|Q9NU19_S132 | 1.502270815 | 3.53E-04 | 2.58E-03 |
| K2C5_HUMAN\|P13647_Y66 | 1.502213659 | 8.56E-05 | 1.15E-03 |
| PDLI2_HUMAN\|Q96JY6_S134_S137_S143 | 1.502208253 | 1.46E-03 | 1.66E-02 |
| SPD2A_HUMAN\|Q5TCZ1_S651 | 1.5020798 | 2.39E-03 | 1.57E-02 |
| AKAP2_HUMAN\|Q9Y2D5_S152_T157 | 1.501839041 | 3.73E-04 | 4.18E-02 |

| | | | |
|---|---|---|---|
| K2C6B_HUMAN\|P04259_S541 | 1.501797592 | 8.25E-04 | 2.60E-02 |
| TGS1_HUMAN\|Q96RS0_S55_T60 | 1.501762686 | 2.61E-05 | 4.73E-04 |
| YTDC2_HUMAN\|Q9H6S0_S1202 | 1.501523685 | 2.59E-04 | 2.00E-03 |
| OSBL3_HUMAN\|Q9H4L5_S437 | 1.501422714 | 2.35E-03 | 1.23E-02 |
| CPSF4_HUMAN\|O95639_S200_S212 | 1.501238998 | 4.17E-04 | 1.64E-02 |
| NFIC_HUMAN\|P08651_S294_S305 | 1.501052155 | 8.73E-04 | 7.15E-03 |
| RFIP3_HUMAN\|O75154_S52 | 1.500899484 | 9.69E-04 | 1.51E-02 |
| RL23A_HUMAN\|P62750_S43 | 1.500828295 | 4.58E-03 | 4.10E-02 |
| MP2K4_HUMAN\|P45985_S393 | 1.500825551 | 9.02E-03 | 3.19E-02 |
| K1C14_HUMAN\|P02533_S44_S53_S54 | 1.50078624 | 4.38E-04 | 1.78E-02 |
| K2C5_HUMAN\|P13647_S11_S14 | 1.5007373 | 1.29E-04 | 2.85E-03 |
| COBL1_HUMAN\|Q53SF7_S1145 | 1.500642709 | 1.47E-03 | 3.58E-02 |
| DBNL_HUMAN\|Q9UJU6_S275 | 1.500470462 | 4.24E-03 | 2.39E-02 |
| PDLI2_HUMAN\|Q96JY6_S127_S129 | 1.500438372 | 4.09E-03 | 4.05E-02 |
| PKHG3_HUMAN\|A1L390_S1081 | 1.500402416 | 3.85E-05 | 4.31E-04 |
| AFTIN_HUMAN\|Q6ULP2_S518 | 1.500330701 | 3.44E-04 | 4.21E-03 |
| NCOA7_HUMAN\|Q8NI08_S208_T210 | 1.500303668 | 2.99E-04 | 6.21E-03 |
| SKT_HUMAN\|Q5T5P2_S169 | 1.500232417 | 8.23E-04 | 1.04E-02 |
| GGYF2_HUMAN\|Q6Y7W6_S26 | 1.500040353 | 4.89E-05 | 1.88E-03 |
| MTMR3_HUMAN\|Q13615_T611 | 1.500004926 | 3.42E-03 | 2.37E-02 |
| LAP2A_HUMAN\|P42166_S66_S67_T74 | 1.499728552 | 1.23E-03 | 1.50E-02 |
| IRS2_HUMAN\|Q9Y4H2_S560 | 1.49970793 | 6.72E-05 | 2.13E-03 |
| ANR17_HUMAN\|O75179_S2045_S2047 | 1.499595937 | 5.28E-05 | 5.39E-04 |
| K1522_HUMAN\|Q9P206_S406 | 1.49951013 | 1.28E-03 | 1.54E-02 |
| FA83B_HUMAN\|Q5T0W9_S869 | 1.499344372 | 2.57E-06 | 2.21E-04 |
| PAXI_HUMAN\|P49023_S106 | 1.499343943 | 1.10E-03 | 5.94E-03 |
| MAP1B_HUMAN\|P46821_Y1796 | 1.499078886 | 2.04E-04 | 2.33E-03 |
| AHNK2_HUMAN\|Q8IVF2_S1712 | 1.498995311 | 4.97E-04 | 9.69E-03 |
| TB182_HUMAN\|Q9C0C2_S836 | 1.498886136 | 2.04E-04 | 5.17E-03 |
| DC1L1_HUMAN\|Q9Y6G9_S510_S516 | 1.498616287 | 3.16E-04 | 6.41E-03 |
| SRRM1_HUMAN\|Q8IYB3_S740_S743 | 1.497970013 | 1.05E-03 | 1.32E-02 |

| | | | |
|---|---|---|---|
| 1B07_HUMAN\|P01889_S359 | 1.497954088 | 3.42E-04 | 3.52E-03 |
| NRK_HUMAN\|Q7Z2Y5_S852_S855 | 1.497873665 | 1.45E-03 | 1.23E-02 |
| PHF14_HUMAN\|O94880_T287 | 1.49757811 | 8.38E-04 | 1.36E-02 |
| PKP2_HUMAN\|Q99959_S81 | 1.497517218 | 6.91E-03 | 4.99E-02 |
| GGYF2_HUMAN\|Q6Y7W6_S23 | 1.49750239 | 8.61E-05 | 2.84E-03 |
| RAI14_HUMAN\|Q9P0K7_T251 | 1.497418118 | 1.67E-03 | 1.79E-02 |
| TBCD5_HUMAN\|Q92609_S541 | 1.497401416 | 3.37E-06 | 2.34E-04 |
| PLEC_HUMAN\|Q15149_S4673 | 1.497333766 | 5.81E-04 | 6.28E-03 |
| SPD2A_HUMAN\|Q5TCZ1_S642 | 1.497103775 | 1.43E-03 | 1.65E-02 |
| IRS2_HUMAN\|Q9Y4H2_S606 | 1.497004006 | 1.06E-02 | 4.75E-02 |
| IRS2_HUMAN\|Q9Y4H2_S1185_S1203 | 1.496991201 | 1.27E-03 | 1.82E-02 |
| FXR2_HUMAN\|P51116_S453 | 1.496894962 | 7.18E-04 | 6.16E-03 |
| EGFR_HUMAN\|P00533_T693 | 1.496824011 | 3.65E-03 | 3.92E-02 |
| LMNB1_HUMAN\|P20700_T19 | 1.496757103 | 1.34E-04 | 4.01E-03 |
| IPP2M_HUMAN\|Q6NXS1_S122 | 1.496715728 | 1.03E-03 | 5.68E-03 |
| DOCK5_HUMAN\|Q9H7D0_S1805 | 1.49668476 | 2.25E-04 | 5.50E-03 |
| REEP4_HUMAN\|Q9H6H4_S253 | 1.496621238 | 5.74E-03 | 4.46E-02 |
| MISP_HUMAN\|Q8IVT2_S394_S395 | 1.496613114 | 2.64E-04 | 6.18E-03 |
| NBEL2_HUMAN\|Q6ZNJ1_T1867 | 1.496565624 | 4.17E-03 | 4.20E-02 |
| PLEC_HUMAN\|Q15149_Y4612 | 1.496510292 | 1.08E-03 | 2.19E-02 |
| IRS1_HUMAN\|P35568_S330 | 1.496509099 | 6.14E-03 | 3.76E-02 |
| UBP2L_HUMAN\|Q14157_S609 | 1.496479459 | 1.04E-03 | 5.70E-03 |
| NIPA_HUMAN\|Q86WB0_S408_S409 | 1.49640837 | 2.30E-05 | 1.09E-03 |
| PALLD_HUMAN\|Q8WX93_S1352 | 1.496074386 | 1.47E-04 | 2.87E-03 |
| TOP2B_HUMAN\|Q02880_S1552 | 1.495994873 | 3.87E-03 | 2.22E-02 |
| PLEC_HUMAN\|Q15149-2_S42 | 1.495488562 | 6.23E-04 | 1.09E-02 |
| CNKR1_HUMAN\|Q969H4_S314 | 1.495336294 | 2.09E-04 | 2.85E-03 |
| NCK5L_HUMAN\|Q9HCH0_S489_S494 | 1.495285596 | 4.24E-04 | 4.13E-03 |
| IRS2_HUMAN\|Q9Y4H2_S1186_S1203 | 1.495136052 | 4.79E-04 | 3.06E-03 |
| NIBL1_HUMAN\|Q96TA1_S641_S646 | 1.495038089 | 5.52E-03 | 2.48E-02 |
| NGAP_HUMAN\|Q9UJF2_S954 | 1.494438022 | 9.03E-04 | 1.43E-02 |

| | | | |
|---|---|---|---|
| K1C17_HUMAN\|Q04695_S28 | 1.494265071 | 2.88E-03 | 2.99E-02 |
| MYO9A_HUMAN\|B2RTY4_S1733 | 1.494220046 | 2.56E-03 | 1.88E-02 |
| ARVC_HUMAN\|O00192_S203 | 1.494196438 | 7.37E-04 | 2.37E-02 |
| SI1L3_HUMAN\|O60292_S172 | 1.494133259 | 4.20E-06 | 1.20E-04 |
| PER1_HUMAN\|O15534_S809_S815 | 1.494103522 | 2.19E-03 | 1.47E-02 |
| LAP2A_HUMAN\|P42166_S167_T160 | 1.4940111 | 8.48E-04 | 1.29E-02 |
| SVIL_HUMAN\|O95425_T679 | 1.49379198 | 3.56E-05 | 1.02E-03 |
| SYTL1_HUMAN\|Q8IYJ3_S216 | 1.49357604 | 1.12E-02 | 4.93E-02 |
| ZN185_HUMAN\|O15231_S452 | 1.493539085 | 8.33E-04 | 8.33E-03 |
| K1C14_HUMAN\|P02533_S53 | 1.49349714 | 2.63E-04 | 4.46E-03 |
| MICA3_HUMAN\|Q7RTP6_S977 | 1.493441907 | 5.52E-06 | 1.28E-03 |
| K1C14_HUMAN\|P02533_S12 | 1.493417101 | 3.56E-04 | 1.52E-02 |
| VIME_HUMAN\|P08670_S66 | 1.493371949 | 2.58E-04 | 4.44E-03 |
| HSPB1_HUMAN\|P04792_S82 | 1.493292367 | 5.35E-05 | 1.99E-03 |
| ARHGG_HUMAN\|Q5VV41_S107 | 1.493251276 | 3.43E-05 | 5.87E-04 |
| TISB_HUMAN\|Q07352_S92 | 1.493203074 | 2.87E-03 | 2.69E-02 |
| PDS5B_HUMAN\|Q9NTI5_S1362 | 1.493105414 | 1.61E-04 | 1.41E-03 |
| BCR_HUMAN\|P11274_S301 | 1.493031789 | 1.15E-02 | 4.51E-02 |
| CSCL2_HUMAN\|Q5T3F8_S114_T110 | 1.493016428 | 1.25E-03 | 7.27E-03 |
| K1C14_HUMAN\|P02533_S437_S438 | 1.492894844 | 2.41E-03 | 2.67E-02 |
| PACN2_HUMAN\|Q9UNF0_S399_S425 | 1.492854891 | 4.05E-04 | 6.63E-03 |
| FA83H_HUMAN\|Q6ZRV2_S1025 | 1.492845652 | 6.03E-04 | 8.93E-03 |
| BIG1_HUMAN\|Q9Y6D6_S397 | 1.492727753 | 3.47E-04 | 1.51E-02 |
| DDX20_HUMAN\|Q9UHI6_S677_Y673 | 1.492675632 | 4.94E-04 | 5.57E-03 |
| TB182_HUMAN\|Q9C0C2_S1158 | 1.492625957 | 1.42E-03 | 1.60E-02 |
| NOG1_HUMAN\|Q9BZE4_S470_Y466 | 1.492594782 | 3.22E-04 | 4.01E-03 |
| MKL2_HUMAN\|Q9ULH7_S921 | 1.492390947 | 7.60E-04 | 1.28E-02 |
| DESP_HUMAN\|P15924_S2815_S2820 | 1.492374523 | 1.97E-04 | 1.64E-03 |
| RIN1_HUMAN\|Q13671_S349 | 1.492286903 | 5.39E-03 | 2.10E-02 |
| K2C5_HUMAN\|P13647_S11_S8 | 1.492102012 | 2.91E-04 | 6.17E-03 |
| LMNB1_HUMAN\|P20700_T20 | 1.492067354 | 5.84E-03 | 3.65E-02 |

| | | | |
|---|---|---|---|
| CF132_HUMAN\|Q5T0Z8_T297 | 1.492029852 | 5.99E-03 | 4.53E-02 |
| BCL9_HUMAN\|O00512_T315 | 1.492029228 | 2.18E-03 | 1.47E-02 |
| TLN1_HUMAN\|Q9Y490_S1641 | 1.492006839 | 5.78E-05 | 1.36E-02 |
| PKN2_HUMAN\|Q16513_S360 | 1.491975922 | 2.11E-03 | 1.43E-02 |
| FA83H_HUMAN\|Q6ZRV2_S936 | 1.491497319 | 1.87E-03 | 1.99E-02 |
| CCDC9_HUMAN\|Q9Y3X0_T389 | 1.491489333 | 2.74E-05 | 3.29E-04 |
| MAP1B_HUMAN\|P46821_S1427 | 1.491465028 | 8.58E-05 | 2.66E-03 |
| PDLI2_HUMAN\|Q96JY6_S123_S124 | 1.491373733 | 2.06E-04 | 2.84E-03 |
| MA7D1_HUMAN\|Q3KQU3_S313 | 1.491052765 | 4.61E-03 | 3.74E-02 |
| KLC3_HUMAN\|Q6P597_S167 | 1.49103949 | 1.47E-03 | 2.73E-02 |
| ASPP2_HUMAN\|Q13625_S556 | 1.490926386 | 1.00E-03 | 1.55E-02 |
| RAI14_HUMAN\|Q9P0K7_T297 | 1.490843229 | 3.76E-03 | 3.59E-02 |
| PPR18_HUMAN\|Q6NYC8_S209_S213 | 1.490716909 | 2.10E-04 | 3.80E-03 |
| TANC2_HUMAN\|Q9HCD6_S1476 | 1.490588508 | 6.11E-03 | 2.31E-02 |
| PDLI2_HUMAN\|Q96JY6_S137_S143 | 1.490546279 | 6.00E-04 | 8.23E-03 |
| SPTB2_HUMAN\|Q01082-3_S10 | 1.49054495 | 8.38E-04 | 1.06E-02 |
| M3K3_HUMAN\|Q99759_S340 | 1.490324615 | 4.73E-04 | 1.28E-02 |
| AMOL2_HUMAN\|Q9Y2J4_S759 | 1.490316248 | 1.65E-04 | 3.42E-03 |
| UH1BL_HUMAN\|A0JNW5_S953 | 1.490264045 | 8.00E-05 | 5.42E-03 |
| AF1L2_HUMAN\|Q8N4X5_S651 | 1.490229405 | 3.87E-05 | 1.60E-03 |
| CSN1_HUMAN\|Q13098_S483_T479 | 1.490140575 | 2.97E-04 | 6.28E-03 |
| LIPA1_HUMAN\|Q13136_S1172 | 1.489978813 | 1.32E-04 | 2.62E-03 |
| CCDC9_HUMAN\|Q9Y3X0_S376 | 1.48987071 | 3.76E-04 | 6.25E-03 |
| PKHG3_HUMAN\|A1L390_S1079 | 1.489696816 | 1.04E-03 | 5.70E-03 |
| WAC2A_HUMAN\|Q641Q2_S375 | 1.489670818 | 5.18E-03 | 4.08E-02 |
| AHNK_HUMAN\|Q09666_S5749_S5752 | 1.489645715 | 2.51E-04 | 1.20E-02 |
| GSK3A_HUMAN\|P49840_S21 | 1.489633106 | 3.02E-04 | 4.81E-03 |
| RALY_HUMAN\|Q9UKM9_T286_T298 | 1.489518128 | 2.95E-03 | 1.31E-02 |
| DC1L1_HUMAN\|Q9Y6G9_S510_S516 | 1.488924454 | 3.73E-04 | 2.53E-03 |
| HBS1L_HUMAN\|Q9Y450-2_S246 | 1.488799307 | 4.81E-04 | 5.43E-03 |
| ZFP91_HUMAN\|Q96JP5_S82 | 1.488653418 | 7.75E-04 | 2.47E-02 |

| | | | |
|---|---|---|---|
| EHBP1_HUMAN\|Q8NDI1_S1035 | 1.488584178 | 3.19E-03 | 2.23E-02 |
| SOS1_HUMAN\|Q07889_S1078_S1082 | 1.488509826 | 1.61E-03 | 1.16E-02 |
| LMNA_HUMAN\|P02545_S403 | 1.488445256 | 3.21E-03 | 1.41E-02 |
| RB6I2_HUMAN\|Q8IUD2_S37_S42 | 1.488390516 | 7.61E-03 | 2.76E-02 |
| STXB5_HUMAN\|Q5T5C0-2_S712 | 1.48834113 | 8.13E-05 | 1.12E-03 |
| SRRM2_HUMAN\|Q9UQ35_S1582 | 1.488272351 | 6.36E-05 | 9.20E-04 |
| PCIF1_HUMAN\|Q9H4Z3_T150 | 1.488112344 | 4.67E-03 | 2.56E-02 |
| PDLI2_HUMAN\|Q96JY6_T279 | 1.488031612 | 9.29E-03 | 4.28E-02 |
| PDLI2_HUMAN\|Q96JY6_S123_S124 | 1.487881688 | 2.20E-04 | 1.22E-02 |
| STXB5_HUMAN\|Q5T5C0-2_S712 | 1.487844003 | 8.21E-05 | 6.64E-03 |
| ZN185_HUMAN\|O15231_S521 | 1.487819857 | 2.67E-04 | 4.49E-03 |
| IRS2_HUMAN\|Q9Y4H2_S1186_S1203 | 1.487814748 | 5.42E-04 | 1.03E-02 |
| K1C14_HUMAN\|P02533_S13 | 1.487717409 | 2.00E-04 | 4.67E-03 |
| DESP_HUMAN\|P15924_S2815 | 1.487685543 | 2.22E-04 | 1.69E-03 |
| RPP25_HUMAN\|Q9BUL9_S162 | 1.487625758 | 8.86E-04 | 1.33E-02 |
| IRS2_HUMAN\|Q9Y4H2_T350 | 1.487395253 | 1.60E-04 | 1.93E-03 |
| MK14_HUMAN\|Q16539_Y182 | 1.487362677 | 1.35E-03 | 1.90E-02 |
| PML_HUMAN\|P29590_S504 | 1.487337996 | 3.66E-04 | 4.41E-03 |
| SPTB2_HUMAN\|Q01082-3_S8 | 1.487152992 | 1.34E-03 | 1.59E-02 |
| UBXN7_HUMAN\|O94888_S280 | 1.487120971 | 9.26E-04 | 9.35E-03 |
| MTCL1_HUMAN\|Q9Y4B5_S549 | 1.486446526 | 2.42E-03 | 2.88E-02 |
| CAMP2_HUMAN\|Q08AD1_S980 | 1.486403066 | 2.61E-03 | 2.57E-02 |
| NIBL1_HUMAN\|Q96TA1_S692 | 1.486359882 | 3.52E-04 | 1.52E-02 |
| 2A5E_HUMAN\|Q16537_S33_S34 | 1.48622377 | 5.20E-05 | 2.29E-03 |
| NEB2_HUMAN\|Q96SB3_S99 | 1.486044725 | 7.79E-04 | 1.30E-02 |
| MAP1B_HUMAN\|P46821_S1797_Y1796 | 1.485937641 | 3.17E-03 | 1.39E-02 |
| DKC1_HUMAN\|O60832_S485_T496 | 1.485921883 | 3.85E-05 | 1.58E-03 |
| GYS1_HUMAN\|P13807_S727 | 1.485845086 | 2.28E-04 | 5.39E-03 |
| RABL6_HUMAN\|Q3YEC7_S492 | 1.485746404 | 1.75E-04 | 3.56E-03 |
| ROCK1_HUMAN\|Q13464_S1341 | 1.485720829 | 2.01E-03 | 2.10E-02 |
| DC1I2_HUMAN\|Q13409_S104 | 1.485641498 | 6.46E-03 | 3.74E-02 |

| | | | |
|---|---|---|---|
| PAXX_HUMAN\|Q9BUH6_S148 | 1.485517214 | 2.22E-04 | 5.49E-03 |
| NELFE_HUMAN\|P18615_S251 | 1.485503259 | 4.28E-03 | 4.24E-02 |
| SCRIB_HUMAN\|Q14160_S1486 | 1.485471583 | 6.84E-04 | 1.20E-02 |
| K1C14_HUMAN\|P02533_S435_S438 | 1.485375666 | 4.36E-03 | 3.98E-02 |
| GSK3A_HUMAN\|P49840_S21 | 1.485117396 | 3.27E-04 | 3.39E-03 |
| PKHG3_HUMAN\|A1L390_S1081 | 1.485006795 | 5.37E-05 | 1.03E-03 |
| ANR17_HUMAN\|O75179_S2041_S2045 | 1.485001479 | 1.44E-03 | 1.65E-02 |
| IASPP_HUMAN\|Q8WUF5_S74 | 1.484958866 | 4.82E-05 | 1.24E-03 |
| BZW2_HUMAN\|Q9Y6E2_S412 | 1.484851435 | 6.99E-03 | 3.01E-02 |
| RIN1_HUMAN\|Q13671_S333_S337 | 1.484643686 | 5.06E-04 | 5.66E-03 |
| LIPB1_HUMAN\|Q86W92_S1001 | 1.4845211 | 8.58E-03 | 4.04E-02 |
| T184B_HUMAN\|Q9Y519_S403 | 1.484500931 | 1.60E-03 | 1.15E-02 |
| CC186_HUMAN\|Q7Z3E2_S726 | 1.48438407 | 6.05E-04 | 1.07E-02 |
| RFIP5_HUMAN\|Q9BXF6_S395 | 1.484347262 | 4.11E-04 | 4.86E-03 |
| CTNA1_HUMAN\|P35221_S641_S652 | 1.484269321 | 1.06E-03 | 1.32E-02 |
| FA83H_HUMAN\|Q6ZRV2_S828 | 1.484226319 | 8.21E-05 | 1.12E-03 |
| TAB2_HUMAN\|Q9NYJ8_S450 | 1.484132052 | 8.25E-05 | 1.40E-03 |
| P33MX_HUMAN\|Q96A73_S183 | 1.484124625 | 2.69E-03 | 1.70E-02 |
| K2C5_HUMAN\|P13647_S50 | 1.483983488 | 2.91E-04 | 1.31E-02 |
| UHRF2_HUMAN\|Q96PU4_S667 | 1.483785321 | 4.87E-05 | 1.90E-03 |
| NUP50_HUMAN\|Q9UKX7_S221 | 1.483590961 | 1.85E-04 | 3.04E-02 |
| BABA1_HUMAN\|Q9NWV8_S57 | 1.483478052 | 2.74E-03 | 2.89E-02 |
| AFTIN_HUMAN\|Q6ULP2_S518 | 1.483280759 | 4.62E-04 | 1.73E-02 |
| E41L2_HUMAN\|O43491_S87 | 1.483068588 | 6.54E-04 | 2.20E-02 |
| ARAF_HUMAN\|P10398_S257_T253 | 1.483048525 | 1.29E-03 | 1.73E-02 |
| PPR18_HUMAN\|Q6NYC8_S224 | 1.482929836 | 1.86E-03 | 2.20E-02 |
| NDRG1_HUMAN\|Q92597_S333_S336 | 1.482912492 | 1.54E-03 | 1.12E-02 |
| ATX2L_HUMAN\|Q8WWM7_S339 | 1.482774497 | 9.85E-06 | 6.18E-04 |
| SF3B2_HUMAN\|Q13435_S309 | 1.48277335 | 1.26E-02 | 4.13E-02 |
| SMRC1_HUMAN\|Q92922_S328_S330 | 1.482706706 | 7.82E-04 | 1.00E-02 |
| CND1_HUMAN\|Q15021_S1333_T1339 | 1.482380488 | 3.78E-03 | 1.81E-02 |

| | | | |
|---|---|---|---|
| KC1D_HUMAN\|P48730_S331 | 1.48219949 | 3.76E-03 | 3.32E-02 |
| RBM27_HUMAN\|Q9P2N5_T447 | 1.482198063 | 8.06E-04 | 6.78E-03 |
| FA83H_HUMAN\|Q6ZRV2_S903 | 1.482184183 | 1.34E-03 | 7.72E-03 |
| AFTIN_HUMAN\|Q6ULP2_S518 | 1.481807109 | 4.75E-04 | 9.21E-03 |
| UT14A_HUMAN\|Q9BVJ6_S437 | 1.481730025 | 2.56E-03 | 3.01E-02 |
| PKHA1_HUMAN\|Q9HB21_S332 | 1.481692791 | 4.41E-05 | 1.32E-03 |
| WDR59_HUMAN\|Q6PJI9_S564 | 1.481587609 | 7.03E-03 | 2.58E-02 |
| VIP2_HUMAN\|O43314_S1006 | 1.481586298 | 1.31E-04 | 8.67E-03 |
| DEN4C_HUMAN\|Q5VZ89_S1046 | 1.481518101 | 1.30E-04 | 2.04E-03 |
| ZFP91_HUMAN\|Q96JP5_S82 | 1.481467873 | 8.73E-04 | 1.32E-02 |
| KLC3_HUMAN\|Q6P597_S497 | 1.481464673 | 8.82E-05 | 1.90E-03 |
| RHG12_HUMAN\|Q8IWW6_S215 | 1.481421874 | 5.08E-04 | 7.21E-03 |
| NUP98_HUMAN\|P52948_S612 | 1.481391754 | 5.01E-04 | 9.43E-03 |
| PDS5B_HUMAN\|Q9NTI5_S1165 | 1.481220859 | 3.78E-03 | 1.81E-02 |
| TBC9B_HUMAN\|Q66K14_S411 | 1.480989458 | 4.12E-03 | 3.46E-02 |
| 1B07_HUMAN\|P01889_S352 | 1.480728107 | 2.54E-03 | 1.64E-02 |
| AKAP2_HUMAN\|Q9Y2D5_S152_T157 | 1.480727303 | 5.42E-04 | 1.03E-02 |
| L7UQD0_HUMAN\|L7UQD0_S268 | 1.480689739 | 1.91E-03 | 4.22E-02 |
| ABCF1_HUMAN\|Q8NE71_T108 | 1.480665752 | 1.06E-03 | 6.40E-03 |
| DDX20_HUMAN\|Q9UHI6_S678_Y673 | 1.48061663 | 2.83E-03 | 1.43E-02 |
| SRSF4_HUMAN\|Q08170_S460 | 1.480393434 | 3.58E-03 | 1.54E-02 |
| DOCK9_HUMAN\|Q9BZ29-5_S37 | 1.480324103 | 3.88E-03 | 3.87E-02 |
| LAP2A_HUMAN\|P42166_S167_T160 | 1.480261865 | 1.05E-03 | 3.06E-02 |
| VIME_HUMAN\|P08670_S430 | 1.48012421 | 6.80E-04 | 5.97E-03 |
| M4K4_HUMAN\|O95819_S842 | 1.480069984 | 2.96E-04 | 3.14E-03 |
| BAD_HUMAN\|Q92934_Y76 | 1.480015981 | 1.70E-04 | 2.46E-03 |
| K1C14_HUMAN\|P02533_S432 | 1.479957816 | 5.37E-03 | 4.25E-02 |
| LAD1_HUMAN\|O00515_S38 | 1.479938477 | 8.22E-04 | 2.58E-02 |
| AKAP2_HUMAN\|Q9Y2D5_T157 | 1.479775414 | 1.11E-03 | 5.99E-03 |
| I2BP2_HUMAN\|Q7Z5L9_S460 | 1.479717745 | 1.76E-06 | 3.67E-04 |
| RMP_HUMAN\|O94763_S372 | 1.47967842 | 2.02E-03 | 2.11E-02 |

| | | | |
|---|---|---|---|
| JUN_HUMAN\|P05412_S63 | 1.479589224 | 3.48E-03 | 1.69E-02 |
| CCDC6_HUMAN\|Q16204_S244 | 1.47948882 | 2.63E-03 | 3.06E-02 |
| HDAC2_HUMAN\|Q92769_S422 | 1.479453324 | 5.70E-03 | 2.54E-02 |
| MON1A_HUMAN\|Q86VX9_S95 | 1.479357391 | 2.96E-04 | 3.80E-03 |
| ATRX_HUMAN\|P46100_S784 | 1.479149733 | 6.10E-03 | 3.75E-02 |
| NOP2_HUMAN\|P46087_T605 | 1.479094622 | 7.11E-04 | 7.28E-03 |
| LAD1_HUMAN\|O00515_S38 | 1.479054448 | 8.35E-04 | 1.36E-02 |
| DLGP4_HUMAN\|Q9Y2H0_S973 | 1.479050505 | 4.88E-03 | 4.81E-02 |
| NFIB_HUMAN\|O00712_S276_S295 | 1.479037194 | 3.47E-03 | 3.80E-02 |
| RTKN_HUMAN\|Q9BST9_S520_S529 | 1.478978672 | 1.47E-03 | 1.63E-02 |
| BRAF_HUMAN\|P15056_S446 | 1.478913217 | 3.36E-03 | 3.74E-02 |
| KHDR1_HUMAN\|Q07666_S18 | 1.478790522 | 1.48E-03 | 1.31E-02 |
| REQU_HUMAN\|Q92785_T149 | 1.478630552 | 1.42E-03 | 1.06E-02 |
| IWS1_HUMAN\|Q96ST2_S720 | 1.478549971 | 8.55E-05 | 1.43E-03 |
| PRSR2_HUMAN\|Q86WR7_S212 | 1.478407721 | 2.72E-04 | 3.58E-03 |
| SPD2A_HUMAN\|Q5TCZ1_S1002 | 1.478011714 | 1.98E-04 | 3.64E-03 |
| MSH6_HUMAN\|P52701_T139 | 1.477994022 | 4.76E-04 | 1.77E-02 |
| K2C5_HUMAN\|P13647_S62 | 1.47786898 | 1.14E-03 | 3.17E-02 |
| ZO2_HUMAN\|Q9UDY2_T1027 | 1.477722757 | 4.39E-04 | 7.07E-03 |
| XCT_HUMAN\|Q9UPY5_S26 | 1.477669967 | 6.61E-03 | 3.97E-02 |
| ANS1A_HUMAN\|Q92625_S649 | 1.477575406 | 4.61E-05 | 3.79E-03 |
| HTF4_HUMAN\|Q99081_S67 | 1.477536322 | 2.62E-03 | 1.98E-02 |
| RL15_HUMAN\|P61313_S34 | 1.477468948 | 9.16E-04 | 1.20E-02 |
| SCRIB_HUMAN\|Q14160_S1486 | 1.477437466 | 7.88E-04 | 4.56E-03 |
| GGYF1_HUMAN\|O75420_S638 | 1.477344463 | 8.47E-04 | 1.36E-02 |
| MK14_HUMAN\|Q16539_Y182 | 1.477306741 | 1.58E-03 | 1.96E-02 |
| ITA3_HUMAN\|P26006_T1044 | 1.477155868 | 8.35E-03 | 4.67E-02 |
| 1433S_HUMAN\|P31947_S74 | 1.477130399 | 2.39E-03 | 1.78E-02 |
| DOCK5_HUMAN\|Q9H7D0_S1805 | 1.477083013 | 3.27E-04 | 2.28E-03 |
| OTU7B_HUMAN\|Q6GQQ9_S467 | 1.477029701 | 9.46E-05 | 1.65E-03 |
| K1C14_HUMAN\|P02533_S28 | 1.476797843 | 8.81E-04 | 2.70E-02 |

| | | | |
|---|---|---|---|
| NKTR_HUMAN\|P30414_S463 | 1.47677348 | 1.55E-03 | 7.88E-03 |
| MB12A_HUMAN\|Q96EY5_S188 | 1.476576999 | 6.57E-03 | 3.28E-02 |
| CCDC6_HUMAN\|Q16204_S244 | 1.476440367 | 2.74E-03 | 1.97E-02 |
| K1C14_HUMAN\|P02533_S429_S432_S433 | 1.476062347 | 3.57E-03 | 2.10E-02 |
| I2BP2_HUMAN\|Q7Z5L9_S460 | 1.476005451 | 1.96E-06 | 1.71E-04 |
| BCL9L_HUMAN\|Q86UU0_S21_S25 | 1.475992005 | 1.70E-04 | 2.46E-03 |
| CCDC6_HUMAN\|Q16204_S244 | 1.475358254 | 2.78E-03 | 3.22E-02 |
| AMOL2_HUMAN\|Q9Y2J4_S759 | 1.475290374 | 2.22E-04 | 3.97E-03 |
| USO1_HUMAN\|O60763_S952 | 1.475289501 | 7.24E-03 | 3.10E-02 |
| HEXI1_HUMAN\|O94992_S237_S252 | 1.475214425 | 1.46E-02 | 4.67E-02 |
| PPM1G_HUMAN\|O15355_S183 | 1.475164108 | 7.26E-04 | 7.38E-03 |
| NASP_HUMAN\|P49321_S421 | 1.475159868 | 1.29E-04 | 2.02E-03 |
| SRPK2_HUMAN\|P78362_S494_S497 | 1.475082111 | 5.65E-04 | 1.06E-02 |
| MAP1B_HUMAN\|P46821_S1792 | 1.474836937 | 2.57E-03 | 2.76E-02 |
| P63_HUMAN\|Q9H3D4_S452_S463 | 1.47477686 | 6.04E-03 | 4.94E-02 |
| K1522_HUMAN\|Q9P206_T162 | 1.474492093 | 5.26E-03 | 4.20E-02 |
| DNLI1_HUMAN\|P18858_S66_S76 | 1.474407451 | 1.06E-02 | 3.64E-02 |
| NDK8_HUMAN\|O60361_S105 | 1.474325028 | 6.97E-04 | 6.06E-03 |
| ZCH18_HUMAN\|Q86VM9_S67 | 1.474186239 | 4.80E-04 | 5.45E-03 |
| OPTN_HUMAN\|Q96CV9_S177 | 1.474129978 | 5.04E-03 | 3.99E-02 |
| FRM4B_HUMAN\|Q9Y2L6_S628 | 1.473827448 | 1.69E-03 | 1.37E-02 |
| K2C6B_HUMAN\|P04259_S546 | 1.473450129 | 6.59E-04 | 5.85E-03 |
| PDLI2_HUMAN\|Q96JY6_S124_S127_S129 | 1.473433624 | 5.19E-03 | 4.48E-02 |
| MAP7_HUMAN\|Q14244_S315 | 1.473330186 | 5.54E-04 | 1.42E-02 |
| TPC12_HUMAN\|Q8WVT3_S184 | 1.473277846 | 4.81E-03 | 4.77E-02 |
| LAP2A_HUMAN\|P42166_S66_S67 | 1.47324997 | 1.42E-04 | 2.18E-03 |
| KC1D_HUMAN\|P48730_S331 | 1.473243327 | 4.24E-03 | 3.51E-02 |
| PKP3_HUMAN\|Q9Y446_S183 | 1.472727007 | 2.27E-03 | 4.99E-02 |
| LMNB2_HUMAN\|Q03252_S424 | 1.47258414 | 3.01E-03 | 2.77E-02 |
| ABCF1_HUMAN\|Q8NE71_T108 | 1.472425557 | 1.21E-03 | 1.09E-02 |
| CCDC6_HUMAN\|Q16204_S244 | 1.472318311 | 2.90E-03 | 3.15E-02 |

| | | | |
|---|---|---|---|
| AAPK1_HUMAN\|Q13131_S508 | 1.472281293 | 1.64E-05 | 2.16E-04 |
| AHNK_HUMAN\|Q09666_T4100 | 1.472180452 | 5.50E-03 | 4.32E-02 |
| RL19_HUMAN\|P84098_S13 | 1.472150974 | 1.29E-03 | 1.49E-02 |
| K1C14_HUMAN\|P02533_S433_S438 | 1.472004172 | 4.40E-03 | 3.70E-02 |
| YTDC2_HUMAN\|Q9H6S0_S1202 | 1.471932783 | 4.46E-04 | 1.69E-02 |
| STAT1_HUMAN\|P42224_S727 | 1.471909441 | 5.69E-04 | 1.06E-02 |
| TAOK3_HUMAN\|Q9H2K8_S442 | 1.471722451 | 2.61E-03 | 2.57E-02 |
| TRI29_HUMAN\|Q14134_S152 | 1.471628819 | 1.09E-03 | 1.63E-02 |
| IP3KB_HUMAN\|P27987_S43 | 1.471607618 | 4.08E-04 | 2.70E-03 |
| C2CD5_HUMAN\|Q86YS7_S307 | 1.471428797 | 9.22E-04 | 1.45E-02 |
| K2C80_HUMAN\|Q6KB66_S451 | 1.471375465 | 9.25E-04 | 5.69E-03 |
| K1671_HUMAN\|Q9BY89_S128 | 1.471351263 | 9.83E-05 | 2.39E-03 |
| ZCH18_HUMAN\|Q86VM9_S532_S534 | 1.4713337 | 2.23E-05 | 7.03E-04 |
| SGTA_HUMAN\|O43765_S305 | 1.471205898 | 8.00E-04 | 8.26E-03 |
| IF4G1_HUMAN\|Q04637_S1231 | 1.471168823 | 8.26E-04 | 1.12E-02 |
| COHA1_HUMAN\|Q9UMD9_S61 | 1.47102314 | 6.93E-03 | 5.00E-02 |
| KLC2_HUMAN\|Q9H0B6_S610 | 1.470933991 | 5.55E-05 | 1.39E-03 |
| PRC2C_HUMAN\|Q9Y520_S187_S189 | 1.470794039 | 1.94E-03 | 2.01E-02 |
| MY18A_HUMAN\|Q92614_S1998 | 1.470781732 | 3.64E-04 | 2.49E-03 |
| ARMX3_HUMAN\|Q9UH62_S110 | 1.470772333 | 5.96E-04 | 6.50E-03 |
| CING_HUMAN\|Q9P2M7_S129_S134 | 1.47076141 | 2.42E-03 | 1.79E-02 |
| CXA1_HUMAN\|P17302_S306 | 1.470611691 | 1.40E-03 | 7.98E-03 |
| AHNK_HUMAN\|Q09666_S5749_S5752 | 1.470585475 | 3.61E-04 | 6.07E-03 |
| SENP3_HUMAN\|Q9H4L4_S169 | 1.470555908 | 6.57E-04 | 1.11E-02 |
| PLEC_HUMAN\|Q15149_S4384 | 1.47055514 | 4.90E-03 | 2.24E-02 |
| TB22B_HUMAN\|Q9NU19_S132 | 1.470435871 | 6.19E-04 | 6.56E-03 |
| SRRM2_HUMAN\|Q9UQ35_S1482_S1483 | 1.470258974 | 2.12E-03 | 2.42E-02 |
| LARP6_HUMAN\|Q9BRS8_S447_T450 | 1.470255216 | 3.61E-03 | 2.11E-02 |
| OSTF1_HUMAN\|Q92882_S213 | 1.470082576 | 1.48E-03 | 1.09E-02 |
| EHBP1_HUMAN\|Q8NDI1_S428_S436 | 1.469804625 | 6.93E-05 | 1.28E-03 |
| RIPK2_HUMAN\|O43353_S531 | 1.469689496 | 2.76E-04 | 4.58E-03 |

| | | | |
|---|---|---|---|
| F120A_HUMAN\|Q9NZB2_S1023 | 1.469582883 | 4.40E-04 | 1.69E-02 |
| MPZL2_HUMAN\|O60487_S208_T214 | 1.469535003 | 1.97E-03 | 1.36E-02 |
| YBOX3_HUMAN\|P16989_S369_T366 | 1.469510132 | 4.30E-04 | 6.32E-03 |
| SRRM1_HUMAN\|Q8IYB3_S754_S756 | 1.469305067 | 6.78E-04 | 5.96E-03 |
| SRRM2_HUMAN\|Q9UQ35_S1582 | 1.469183152 | 9.65E-05 | 7.17E-03 |
| LMNA_HUMAN\|P02545_S414 | 1.46917727 | 1.35E-03 | 1.54E-02 |
| AHNK_HUMAN\|Q09666_T4766 | 1.469125397 | 1.44E-03 | 1.65E-02 |
| PER1_HUMAN\|O15534_S815 | 1.468914656 | 3.57E-03 | 2.10E-02 |
| TM1L1_HUMAN\|O75674_S321_S323 | 1.468894468 | 2.89E-03 | 4.34E-02 |
| HIRP3_HUMAN\|Q9BW71_S196_S199 | 1.468695747 | 4.34E-04 | 4.21E-03 |
| K1C16_HUMAN\|P08779_S447_S450 | 1.468662726 | 1.80E-03 | 2.15E-02 |
| H2AX_HUMAN\|P16104_S140 | 1.468661952 | 1.74E-04 | 2.49E-03 |
| ARFG1_HUMAN\|Q8N6T3_S378 | 1.468660231 | 1.52E-03 | 1.28E-02 |
| BZW2_HUMAN\|Q9Y6E2_S412 | 1.468639507 | 8.44E-03 | 3.02E-02 |
| TRI47_HUMAN\|Q96LD4_S588 | 1.46848922 | 1.81E-03 | 8.87E-03 |
| PKP2_HUMAN\|Q99959_S197 | 1.468413305 | 6.11E-04 | 8.32E-03 |
| K1C14_HUMAN\|P02533_S432_S435 | 1.4683843 | 8.36E-03 | 3.96E-02 |
| TBC9B_HUMAN\|Q66K14_S1241 | 1.468376355 | 6.60E-05 | 2.52E-03 |
| ARH40_HUMAN\|Q8TER5_S1513 | 1.468351948 | 5.10E-05 | 2.09E-03 |
| HMGA2_HUMAN\|P52926_S102 | 1.467968783 | 1.06E-02 | 4.23E-02 |
| ACINU_HUMAN\|Q9UKV3_S240_S243 | 1.467894433 | 2.36E-04 | 5.68E-03 |
| SLK_HUMAN\|Q9H2G2_T820 | 1.467378139 | 9.42E-05 | 1.24E-03 |
| NIBL1_HUMAN\|Q96TA1_S691 | 1.46716183 | 4.35E-05 | 3.70E-03 |
| RABE1_HUMAN\|Q15276_S407 | 1.467009199 | 4.23E-04 | 4.94E-03 |
| UBE3A_HUMAN\|Q05086_S218 | 1.46696892 | 9.44E-05 | 2.82E-03 |
| TB22B_HUMAN\|Q9NU19_S132 | 1.466906224 | 6.59E-04 | 7.01E-03 |
| PUM2_HUMAN\|Q8TB72_S182 | 1.46686364 | 5.63E-03 | 2.17E-02 |
| TM1L1_HUMAN\|O75674_S321 | 1.466601464 | 7.94E-03 | 4.51E-02 |
| MY18A_HUMAN\|Q92614_S1998 | 1.466458193 | 3.95E-04 | 1.12E-02 |
| AF1L2_HUMAN\|Q8N4X5_S344 | 1.466347531 | 9.15E-04 | 1.13E-02 |
| LAP2A_HUMAN\|P42166_S66_S67_T74 | 1.466247307 | 2.02E-03 | 1.58E-02 |

| | | | |
|---|---|---|---|
| CALD1_HUMAN\|Q05682_S789 | 1.466147184 | 2.73E-03 | 1.97E-02 |
| PLEC_HUMAN\|Q15149_S4673 | 1.466030746 | 9.88E-04 | 1.45E-02 |
| HDAC2_HUMAN\|Q92769_S422 | 1.465964765 | 6.71E-03 | 3.84E-02 |
| CBLB_HUMAN\|Q13191_S480_S484 | 1.465892546 | 2.13E-04 | 1.75E-03 |
| SRRM2_HUMAN\|Q9UQ35_S1482_S1483 | 1.465834505 | 2.27E-03 | 3.73E-02 |
| ZC3HE_HUMAN\|Q6PJT7_S390 | 1.465706797 | 1.99E-03 | 1.07E-02 |
| AHNK_HUMAN\|Q09666_S210_S216 | 1.465685337 | 7.15E-04 | 1.21E-02 |
| Q659E9_HUMAN\|Q659E9_S168 | 1.465454292 | 1.17E-04 | 3.52E-03 |
| KLC2_HUMAN\|Q9H0B6_S608_S611 | 1.465223279 | 1.01E-04 | 2.09E-03 |
| ABCF1_HUMAN\|Q8NE71_S109_T108 | 1.465212472 | 9.06E-05 | 2.25E-03 |
| AFTIN_HUMAN\|Q6ULP2_S518 | 1.46496767 | 6.38E-04 | 2.20E-02 |
| K1C14_HUMAN\|P02533_S435_S438 | 1.46486204 | 5.72E-03 | 4.45E-02 |
| TOM1_HUMAN\|O60784_S462 | 1.464640834 | 2.80E-03 | 3.17E-02 |
| PP14A_HUMAN\|Q96A00_S26 | 1.464631902 | 2.23E-04 | 2.52E-03 |
| ABI2_HUMAN\|Q9NYB9_S368 | 1.464490007 | 6.19E-05 | 9.03E-04 |
| K1522_HUMAN\|Q9P206_S161 | 1.464448685 | 3.77E-04 | 3.80E-03 |
| SI1L1_HUMAN\|O43166_S1647_S1649 | 1.464436177 | 2.24E-03 | 1.50E-02 |
| LASP1_HUMAN\|Q14847_T104 | 1.464399867 | 4.51E-04 | 7.15E-03 |
| EXOC1_HUMAN\|Q9NV70_T471 | 1.464374784 | 7.43E-05 | 1.71E-03 |
| N4BP1_HUMAN\|O75113_T242 | 1.464372795 | 4.68E-04 | 7.30E-03 |
| ASPP2_HUMAN\|Q13625_S558 | 1.464351646 | 1.93E-04 | 4.99E-03 |
| CTNA1_HUMAN\|P35221_T654_T658 | 1.464308297 | 3.13E-03 | 2.84E-02 |
| SFPQ_HUMAN\|P23246_T687 | 1.464256723 | 4.63E-03 | 3.75E-02 |
| CTR1_HUMAN\|P30825_T620 | 1.464201763 | 5.27E-03 | 4.50E-02 |
| SRC8_HUMAN\|Q14247_S418 | 1.464036864 | 3.54E-03 | 1.53E-02 |
| KSR1_HUMAN\|Q8IVT5_S406 | 1.463941498 | 7.73E-03 | 3.71E-02 |
| RRP1B_HUMAN\|Q14684_S245 | 1.463871862 | 1.56E-04 | 4.31E-03 |
| RHG05_HUMAN\|Q13017_S1173 | 1.463751732 | 6.33E-04 | 6.77E-03 |
| TB182_HUMAN\|Q9C0C2_S1103 | 1.46373329 | 1.88E-03 | 1.56E-02 |
| SARG_HUMAN\|Q9BW04_S429 | 1.463681778 | 5.63E-03 | 2.91E-02 |
| AHNK_HUMAN\|Q09666_S2397 | 1.463636392 | 4.05E-03 | 4.26E-02 |

| | | | |
|---|---|---|---|
| MAP1B_HUMAN\|P46821_S1427 | 1.46351992 | 1.55E-04 | 5.62E-03 |
| SSH2_HUMAN\|Q76I76_S38 | 1.463469717 | 7.46E-04 | 1.26E-02 |
| COBL1_HUMAN\|Q53SF7_T322 | 1.463456405 | 2.87E-04 | 1.31E-02 |
| ARP19_HUMAN\|P56211_S62 | 1.463048652 | 8.35E-04 | 6.92E-03 |
| SIK3_HUMAN\|Q9Y2K2_S493 | 1.462922699 | 1.36E-04 | 2.96E-03 |
| K1C14_HUMAN\|P02533_S429_S435 | 1.462909077 | 4.88E-03 | 2.64E-02 |
| P33MX_HUMAN\|Q96A73_S183_T179 | 1.462618458 | 5.46E-03 | 2.46E-02 |
| AHNK_HUMAN\|Q09666_S3412 | 1.462613548 | 1.43E-03 | 1.65E-02 |
| M3K3_HUMAN\|Q99759_S337 | 1.46251259 | 3.91E-03 | 2.59E-02 |
| M3K3_HUMAN\|Q99759_S340 | 1.462504407 | 7.83E-04 | 6.65E-03 |
| PLEC_HUMAN\|Q15149_S4389_S4396 | 1.462215572 | 6.62E-03 | 3.81E-02 |
| Z280C_HUMAN\|Q8ND82_S80 | 1.462147174 | 5.90E-04 | 6.44E-03 |
| PKHG5_HUMAN\|O94827_S959 | 1.461958344 | 5.82E-03 | 2.98E-02 |
| DDI2_HUMAN\|Q5TDH0_S194 | 1.461904446 | 2.06E-03 | 1.67E-02 |
| RHG05_HUMAN\|Q13017_S1173 | 1.461797717 | 6.55E-04 | 6.80E-03 |
| P4K2A_HUMAN\|Q9BTU6_S462_S468 | 1.461787976 | 3.76E-04 | 2.55E-03 |
| LAD1_HUMAN\|O00515_S394 | 1.461752513 | 9.99E-04 | 9.80E-03 |
| AAK1_HUMAN\|Q2M2I8_S678 | 1.461629966 | 9.98E-04 | 6.06E-03 |
| NEB2_HUMAN\|Q96SB3_S99 | 1.461621338 | 1.17E-03 | 1.37E-02 |
| PDLI2_HUMAN\|Q96JY6_S127_S129 | 1.461202528 | 6.65E-03 | 3.98E-02 |
| SGTA_HUMAN\|O43765_S305 | 1.461186859 | 9.53E-04 | 1.52E-02 |
| IASPP_HUMAN\|Q8WUF5_S395 | 1.461016937 | 5.99E-03 | 4.59E-02 |
| ICE1_HUMAN\|Q9Y2F5_S1712 | 1.461003797 | 1.42E-03 | 3.49E-02 |
| PPR18_HUMAN\|Q6NYC8_S490 | 1.460838299 | 1.43E-04 | 4.15E-03 |
| UT14A_HUMAN\|Q9BVJ6_S437 | 1.460764598 | 3.42E-03 | 3.77E-02 |
| COBL1_HUMAN\|Q53SF7_T298 | 1.460637192 | 2.03E-03 | 4.54E-02 |
| ARI1A_HUMAN\|O14497_S696_S698_S702 | 1.460634546 | 1.14E-04 | 6.96E-03 |
| AKAP2_HUMAN\|Q9Y2D5_S152_S155 | 1.460617537 | 7.18E-04 | 1.23E-02 |
| IPP2M_HUMAN\|Q6NXS1_S121 | 1.460516972 | 4.47E-03 | 3.00E-02 |
| IPP2M_HUMAN\|Q6NXS1_S122 | 1.460422417 | 1.82E-03 | 1.53E-02 |
| SPB1_HUMAN\|Q8IY81_T573 | 1.460125217 | 1.01E-03 | 9.86E-03 |

| | | | |
|---|---|---|---|
| ERF_HUMAN\|P50548_S24 | 1.459980186 | 6.65E-04 | 1.18E-02 |
| LAP2A_HUMAN\|P42166_S66_S67 | 1.459837742 | 1.86E-04 | 4.81E-03 |
| K1C14_HUMAN\|P02533_S28 | 1.45967734 | 1.19E-03 | 1.46E-02 |
| MBLC1_HUMAN\|A4D2B0_S252 | 1.459613994 | 3.46E-03 | 2.39E-02 |
| DDX10_HUMAN\|Q13206_S831 | 1.459456478 | 1.01E-03 | 6.09E-03 |
| BAIP2_HUMAN\|Q9UQB8-4_T520 | 1.459268248 | 3.35E-03 | 2.02E-02 |
| HMGA2_HUMAN\|P52926_S101_S105_T100 | 1.459135604 | 3.09E-03 | 2.81E-02 |
| PLEC_HUMAN\|Q15149_S4384_S4385 | 1.45913177 | 3.37E-03 | 1.47E-02 |
| ARVC_HUMAN\|O00192_S203 | 1.459078732 | 1.35E-03 | 1.60E-02 |
| LYAR_HUMAN\|Q9NX58_S276 | 1.458987724 | 1.70E-04 | 1.47E-03 |
| FRMD8_HUMAN\|Q9BZ67_S446 | 1.458811493 | 4.42E-03 | 2.06E-02 |
| LMNA_HUMAN\|P02545_S403_S407 | 1.458663975 | 2.33E-04 | 2.60E-03 |
| NSD3_HUMAN\|Q9BZ95_S457 | 1.458663122 | 6.89E-04 | 1.21E-02 |
| SCRIB_HUMAN\|Q14160_S1285 | 1.458481654 | 7.22E-03 | 4.22E-02 |
| LIPA3_HUMAN\|O75145_S1164 | 1.458410622 | 7.34E-04 | 4.29E-03 |
| EPN2_HUMAN\|O95208_S172 | 1.458377306 | 8.11E-03 | 3.87E-02 |
| CC186_HUMAN\|Q7Z3E2_S726 | 1.458357897 | 9.61E-04 | 9.56E-03 |
| PRC2A_HUMAN\|P48634_S1085 | 1.458135962 | 2.16E-03 | 1.73E-02 |
| E7ET48_HUMAN\|E7ET48_S441 | 1.458066636 | 2.32E-03 | 2.80E-02 |
| TACC2_HUMAN\|O95359_S2072 | 1.458059195 | 4.02E-04 | 4.76E-03 |
| RAI14_HUMAN\|Q9P0K7_S296 | 1.458041331 | 3.82E-03 | 3.63E-02 |
| NU107_HUMAN\|P57740_S37 | 1.458013345 | 9.61E-04 | 9.56E-03 |
| IRS2_HUMAN\|Q9Y4H2_S1185_S1203 | 1.457816113 | 2.27E-03 | 1.06E-02 |
| TFE3_HUMAN\|P19532_S568_S570 | 1.457679625 | 3.44E-04 | 2.52E-03 |
| RIPK2_HUMAN\|O43353_S531 | 1.457647362 | 3.47E-04 | 4.24E-03 |
| AKTS1_HUMAN\|Q96B36_S202_S203 | 1.457558211 | 1.95E-04 | 1.53E-03 |
| K1C17_HUMAN\|Q04695_S10 | 1.457479343 | 3.13E-03 | 2.95E-02 |
| K1522_HUMAN\|Q9P206_S404_S409 | 1.457198846 | 4.86E-04 | 9.08E-03 |
| TACC1_HUMAN\|O75410_S44 | 1.457058299 | 1.36E-03 | 1.01E-02 |
| T2FA_HUMAN\|P35269_T389 | 1.457049902 | 6.54E-03 | 2.85E-02 |
| SRRM2_HUMAN\|Q9UQ35_S871_S876 | 1.456942725 | 1.39E-03 | 1.81E-02 |

| | | | |
|---|---|---|---|
| AHNK_HUMAN\|Q09666_S212 | 1.456741962 | 3.88E-03 | 3.41E-02 |
| SRRM2_HUMAN\|Q9UQ35_S1014 | 1.456525295 | 8.80E-05 | 1.18E-03 |
| PKP2_HUMAN\|Q99959_S197 | 1.456437119 | 7.52E-04 | 1.05E-02 |
| SPS2L_HUMAN\|Q9NUQ6_S455 | 1.456224354 | 6.33E-03 | 3.69E-02 |
| RPB1_HUMAN\|P24928_S1847_T1856 | 1.456100303 | 7.78E-03 | 3.30E-02 |
| UBE3A_HUMAN\|Q05086_S218 | 1.456078016 | 1.20E-04 | 2.42E-03 |
| GYS1_HUMAN\|P13807_T703 | 1.456065943 | 1.74E-03 | 8.59E-03 |
| NIBL1_HUMAN\|Q96TA1_S665 | 1.456048531 | 2.69E-04 | 1.95E-03 |
| PER1_HUMAN\|O15534_S1007 | 1.455977076 | 6.06E-04 | 8.29E-03 |
| L7UQD0_HUMAN\|L7UQD0_S268 | 1.455956704 | 2.78E-03 | 1.74E-02 |
| CAMP2_HUMAN\|Q08AD1_S980 | 1.455692141 | 3.98E-03 | 3.73E-02 |
| CAVN3_HUMAN\|Q969G5_T205 | 1.455604383 | 4.29E-03 | 3.63E-02 |
| FKB15_HUMAN\|Q5T1M5_S344 | 1.455498494 | 9.64E-04 | 1.17E-02 |
| LIPA1_HUMAN\|Q13136_S666 | 1.45546163 | 8.43E-04 | 1.13E-02 |
| TWF1_HUMAN\|Q12792_S143 | 1.455448478 | 1.06E-03 | 6.38E-03 |
| CE170_HUMAN\|Q5SW79_S636_T628 | 1.455423697 | 3.44E-03 | 2.05E-02 |
| SPB1_HUMAN\|Q8IY81_T573 | 1.455385788 | 1.09E-03 | 1.30E-02 |
| K1C14_HUMAN\|P02533_S54 | 1.45518833 | 4.07E-04 | 4.01E-03 |
| UH1BL_HUMAN\|A0JNW5_S953 | 1.454982568 | 1.68E-04 | 4.13E-03 |
| SYNEM_HUMAN\|O15061_S1044_S1049 | 1.454930192 | 1.26E-03 | 9.57E-03 |
| RAD9A_HUMAN\|Q99638_S387 | 1.4548139 | 2.27E-03 | 1.71E-02 |
| K1522_HUMAN\|Q9P206_S404_S409 | 1.454645343 | 5.11E-04 | 1.90E-02 |
| SSH1_HUMAN\|Q8WYL5_S937 | 1.454620728 | 3.32E-04 | 1.00E-02 |
| PACN3_HUMAN\|Q9UKS6_S344_S354 | 1.454593567 | 6.60E-04 | 1.18E-02 |
| IF4B_HUMAN\|P23588_S495_S497 | 1.454558569 | 3.24E-04 | 4.03E-03 |
| SMRC1_HUMAN\|Q92922_S328_S330 | 1.454553321 | 1.23E-03 | 1.73E-02 |
| FA83H_HUMAN\|Q6ZRV2_S1024 | 1.454514035 | 5.70E-03 | 2.93E-02 |
| KDM3B_HUMAN\|Q7LBC6_S293 | 1.454427546 | 7.34E-05 | 2.49E-03 |
| ZN185_HUMAN\|O15231_S452 | 1.454419806 | 1.60E-03 | 1.73E-02 |
| PDS5A_HUMAN\|Q29RF7_S1276 | 1.454383535 | 4.45E-03 | 4.36E-02 |
| ICE1_HUMAN\|Q9Y2F5_S1712 | 1.454376488 | 1.59E-03 | 3.92E-02 |

| | | | |
|---|---|---|---|
| PDLI2_HUMAN\|Q96JY6_S123_S124 | 1.45420775 | 4.19E-04 | 4.91E-03 |
| SACS_HUMAN\|Q9NZJ4_S4264 | 1.454207033 | 4.81E-03 | 1.94E-02 |
| KI26B_HUMAN\|Q2KJY2_S1006_S1018_S1021 | 1.454146337 | 1.29E-04 | 1.11E-03 |
| WAC2A_HUMAN\|Q641Q2_S288 | 1.454089718 | 6.35E-03 | 4.70E-02 |
| ALKB5_HUMAN\|Q6P6C2_S384 | 1.453974517 | 1.43E-04 | 1.76E-03 |
| ZN185_HUMAN\|O15231_S153 | 1.453806622 | 5.05E-04 | 3.48E-03 |
| PRC2A_HUMAN\|P48634_S1147 | 1.45368904 | 2.27E-04 | 4.02E-03 |
| ASHWN_HUMAN\|Q9BVC5_S193 | 1.453563429 | 9.58E-05 | 2.82E-03 |
| PHLB1_HUMAN\|Q86UU1_S430 | 1.453552677 | 2.25E-03 | 2.25E-02 |
| MTAP2_HUMAN\|P11137_S1787_S1790_T1780 | 1.45335383 | 3.16E-04 | 5.51E-03 |
| BRD4_HUMAN\|O60885_S470 | 1.453330634 | 3.51E-03 | 3.18E-02 |
| UBP32_HUMAN\|Q8NFA0_S1393 | 1.453314208 | 4.87E-04 | 1.31E-02 |
| NGAP_HUMAN\|Q9UJF2-2_S134 | 1.453299336 | 1.17E-03 | 1.69E-02 |
| RHG32_HUMAN\|A7KAX9_S682 | 1.452885986 | 2.07E-04 | 2.91E-03 |
| EPN1_HUMAN\|Q9Y6I3_S454 | 1.452497815 | 2.82E-04 | 2.14E-03 |
| SRRM1_HUMAN\|Q8IYB3_S754_S756 | 1.452367888 | 9.23E-04 | 2.75E-02 |
| DDX3Y_HUMAN\|O15523_S607 | 1.452317856 | 6.96E-04 | 1.15E-02 |
| AFAD_HUMAN\|P55196_T1330 | 1.452291462 | 1.30E-03 | 1.56E-02 |
| IASPP_HUMAN\|Q8WUF5_Y109 | 1.452276819 | 8.01E-04 | 1.25E-02 |
| GIT1_HUMAN\|Q9Y2X7_S362 | 1.45213254 | 3.73E-04 | 7.83E-03 |
| KI26B_HUMAN\|Q2KJY2_S1006_S1018_S1021 | 1.452099885 | 1.35E-04 | 4.02E-03 |
| EP15R_HUMAN\|Q9UBC2_S229 | 1.452030601 | 3.37E-05 | 1.38E-03 |
| NOP2_HUMAN\|P46087_T605 | 1.451947514 | 1.14E-03 | 1.72E-02 |
| NUP98_HUMAN\|P52948_T1000 | 1.451852267 | 6.61E-03 | 3.81E-02 |
| PININ_HUMAN\|Q9H307_S690 | 1.451336592 | 7.00E-03 | 3.97E-02 |
| BICD2_HUMAN\|Q8TD16_S615 | 1.451237438 | 6.41E-04 | 1.11E-02 |
| TMF1_HUMAN\|P82094_S72 | 1.451143586 | 4.21E-04 | 8.71E-03 |
| BRE1A_HUMAN\|Q5VTR2_S136 | 1.450884757 | 1.14E-04 | 3.04E-03 |
| ASPP2_HUMAN\|Q13625_S558 | 1.450855811 | 2.56E-04 | 1.88E-03 |
| NDRG1_HUMAN\|Q92597_S364 | 1.450846936 | 7.90E-04 | 6.68E-03 |
| SKT_HUMAN\|Q5T5P2_S169 | 1.450835443 | 1.80E-03 | 1.93E-02 |

| | | | |
|---|---|---|---|
| K1C17_HUMAN\|Q04695_S11_S12 | 1.45077752 | 5.40E-03 | 4.58E-02 |
| PYGO2_HUMAN\|Q9BRQ0_T302 | 1.450691897 | 7.02E-04 | 2.37E-02 |
| CAMP2_HUMAN\|Q08AD1_S691 | 1.450593706 | 6.64E-03 | 3.29E-02 |
| TBD2A_HUMAN\|Q9BYX2_S920 | 1.450490815 | 6.83E-04 | 2.24E-02 |
| PAK2_HUMAN\|Q13177_Y139 | 1.450202125 | 4.81E-03 | 3.95E-02 |
| AHNK_HUMAN\|Q09666_S572 | 1.450101228 | 4.47E-03 | 4.36E-02 |
| FRMD8_HUMAN\|Q9BZ67_S446 | 1.450094824 | 4.96E-03 | 3.06E-02 |
| NFIC_HUMAN\|P08651_S339 | 1.45007974 | 5.76E-03 | 2.96E-02 |
| KLC2_HUMAN\|Q9H0B6_S608_S611 | 1.449907863 | 1.40E-04 | 3.59E-03 |
| CAVN1_HUMAN\|Q6NZI2_T302 | 1.449867568 | 2.46E-04 | 4.29E-03 |
| ITB4_HUMAN\|P16144_T1455 | 1.449838665 | 4.98E-03 | 2.68E-02 |
| PP1R7_HUMAN\|Q15435_S24 | 1.449837586 | 9.67E-03 | 3.92E-02 |
| SAMH1_HUMAN\|Q9Y3Z3_S33 | 1.449696783 | 1.67E-03 | 3.98E-02 |
| MSH6_HUMAN\|P52701_T139 | 1.449413068 | 8.14E-04 | 6.81E-03 |
| PCM1_HUMAN\|Q15154_S872 | 1.449036073 | 3.76E-04 | 7.46E-03 |
| ACINU_HUMAN\|Q9UKV3_S243 | 1.448974176 | 3.61E-04 | 5.59E-03 |
| HMGA2_HUMAN\|P52926_T100 | 1.44886051 | 1.57E-02 | 4.94E-02 |
| FXR2_HUMAN\|P51116_S453 | 1.448852059 | 1.62E-03 | 3.92E-02 |
| WDR44_HUMAN\|Q5JSH3_S561 | 1.448685282 | 6.55E-04 | 4.33E-03 |
| IF2P_HUMAN\|O60841_S182_S183 | 1.448607565 | 3.82E-04 | 2.75E-03 |
| CHSP1_HUMAN\|Q9Y2V2_S41 | 1.448593705 | 2.48E-03 | 1.61E-02 |
| PDLI2_HUMAN\|Q96JY6_S134_S137_S143 | 1.448557902 | 3.19E-03 | 3.48E-02 |
| ZN185_HUMAN\|O15231_S152 | 1.448496959 | 3.46E-04 | 1.47E-02 |
| TAB2_HUMAN\|Q9NYJ8_S450 | 1.448472538 | 1.78E-04 | 1.52E-03 |
| TBD2A_HUMAN\|Q9BYX2_S920 | 1.448355772 | 7.10E-04 | 1.67E-02 |
| RPP25_HUMAN\|Q9BUL9_S162 | 1.448208752 | 1.73E-03 | 2.28E-02 |
| DKC1_HUMAN\|O60832_S485_S494 | 1.448189397 | 2.35E-04 | 1.25E-02 |
| TB182_HUMAN\|Q9C0C2_S836 | 1.448065179 | 5.42E-04 | 3.38E-03 |
| CBLB_HUMAN\|Q13191_S480_S484 | 1.448028602 | 3.07E-04 | 6.33E-03 |
| SRC8_HUMAN\|Q14247_S405_T399 | 1.447940267 | 1.39E-03 | 2.01E-02 |
| KPCD_HUMAN\|Q05655_S307 | 1.447850351 | 5.63E-05 | 1.05E-03 |

| | | | |
|---|---|---|---|
| COBL1_HUMAN\|Q53SF7_S364 | 1.447713978 | 7.39E-04 | 9.70E-03 |
| DDI2_HUMAN\|Q5TDH0_S194 | 1.447624691 | 2.57E-03 | 2.55E-02 |
| K1C14_HUMAN\|P02533_S13 | 1.447587879 | 4.47E-04 | 1.78E-02 |
| PARP9_HUMAN\|Q8IXQ6_S851 | 1.447482545 | 3.01E-04 | 6.72E-03 |
| HNRPC_HUMAN\|P07910_S239 | 1.447435799 | 6.10E-04 | 8.32E-03 |
| GSK3A_HUMAN\|P49840_T19 | 1.447366132 | 3.18E-04 | 3.31E-03 |
| PDLI2_HUMAN\|Q96JY6_S74 | 1.447331866 | 5.28E-05 | 1.51E-03 |
| AHNK_HUMAN\|Q09666_S5749_S5752 | 1.447140907 | 5.67E-04 | 1.00E-02 |
| AHNK_HUMAN\|Q09666_S572 | 1.446983202 | 4.67E-03 | 2.96E-02 |
| SIPA1_HUMAN\|Q96FS4_S1036 | 1.446953995 | 1.50E-03 | 7.66E-03 |
| NELFE_HUMAN\|P18615_S51 | 1.446922522 | 8.76E-04 | 1.43E-02 |
| PML_HUMAN\|P29590_T867 | 1.446544339 | 1.29E-03 | 3.31E-02 |
| TPBG_HUMAN\|Q13641_S415_S418 | 1.446398568 | 5.56E-04 | 1.05E-02 |
| HSPB1_HUMAN\|P04792_S82 | 1.44637444 | 1.50E-04 | 1.23E-03 |
| SIK3_HUMAN\|Q9Y2K2_S493 | 1.446094951 | 1.95E-04 | 3.60E-03 |
| CING_HUMAN\|Q9P2M7_S129_S134 | 1.446066606 | 3.49E-03 | 2.07E-02 |
| TPD54_HUMAN\|O43399_S96 | 1.445406763 | 1.52E-02 | 4.83E-02 |
| KPCD_HUMAN\|Q05655_S307 | 1.44538218 | 5.96E-05 | 1.95E-03 |
| CNKR1_HUMAN\|Q969H4_S314 | 1.445345287 | 5.47E-04 | 9.95E-03 |
| TM1L1_HUMAN\|O75674_S321_S323 | 1.445279356 | 4.07E-03 | 4.27E-02 |
| RFIP5_HUMAN\|Q9BXF6_S397 | 1.445160506 | 1.58E-03 | 1.38E-02 |
| CND1_HUMAN\|Q15021_S1333 | 1.445100714 | 1.07E-03 | 1.65E-02 |
| LARP1_HUMAN\|Q6PKG0_S521 | 1.444738097 | 4.35E-03 | 2.04E-02 |
| PGAM1_HUMAN\|P18669_S14 | 1.44466384 | 2.54E-03 | 1.64E-02 |
| M3K3_HUMAN\|Q99759_S337_S340 | 1.444568423 | 4.82E-04 | 9.51E-03 |
| PLEC_HUMAN\|Q15149_S4626 | 1.444420385 | 4.16E-04 | 2.74E-03 |
| AT11C_HUMAN\|Q8NB49_T1124 | 1.444112273 | 1.30E-03 | 7.53E-03 |
| TCAL3_HUMAN\|Q969E4_S125 | 1.444013318 | 6.92E-03 | 3.40E-02 |
| APC_HUMAN\|P25054_S2242_S2244 | 1.444002073 | 1.24E-04 | 1.97E-03 |
| AKAP2_HUMAN\|Q9Y2D5_S152_S155 | 1.443950919 | 9.72E-04 | 1.25E-02 |
| PRR12_HUMAN\|Q9ULL5_S44 | 1.4438162 | 1.85E-03 | 1.97E-02 |

| | | | |
|---|---|---|---|
| K1C16_HUMAN\|P08779_S468 | 1.443792665 | 1.52E-03 | 1.92E-02 |
| AHNK_HUMAN\|Q09666_S5749 | 1.443743019 | 7.61E-05 | 6.27E-03 |
| PCM1_HUMAN\|Q15154_S93 | 1.443230885 | 1.55E-04 | 2.28E-03 |
| RNF31_HUMAN\|Q96EP0_S466 | 1.44320824 | 2.83E-04 | 3.69E-03 |
| TBCD4_HUMAN\|O60343_S318 | 1.443147413 | 9.26E-04 | 7.49E-03 |
| FLVC1_HUMAN\|Q9Y5Y0_S536 | 1.442851348 | 5.18E-03 | 4.80E-02 |
| ARP19_HUMAN\|P56211_Y59 | 1.442750682 | 9.56E-04 | 7.68E-03 |
| PHLB1_HUMAN\|Q86UU1_S430 | 1.44266519 | 2.66E-03 | 3.11E-02 |
| K2C8_HUMAN\|P05787_S21_S24_S27 | 1.442655337 | 3.57E-03 | 3.48E-02 |
| NELFE_HUMAN\|P18615_S51 | 1.442653647 | 9.45E-04 | 2.80E-02 |
| DTD1_HUMAN\|Q8TEA8_S196_S197 | 1.442333344 | 3.05E-04 | 2.17E-03 |
| REPS1_HUMAN\|Q96D71_S162_S166_T173 | 1.442286342 | 3.30E-04 | 6.87E-03 |
| SIK3_HUMAN\|Q9Y2K2_S808 | 1.442225803 | 3.39E-03 | 3.55E-02 |
| ARHG5_HUMAN\|Q12774_S445_S450 | 1.442212911 | 1.18E-04 | 3.54E-03 |
| SPTB2_HUMAN\|Q01082-3_S14_S8 | 1.442162043 | 8.88E-04 | 8.75E-03 |
| K2C6B_HUMAN\|P04259_S546 | 1.442116932 | 1.17E-03 | 1.61E-02 |
| AHNK2_HUMAN\|Q8IVF2_S5175 | 1.442012457 | 3.09E-03 | 3.16E-02 |
| PLEC_HUMAN\|Q15149_S4626 | 1.441927965 | 4.38E-04 | 8.99E-03 |
| RL37_HUMAN\|P61927_S95 | 1.441846859 | 5.82E-03 | 4.85E-02 |
| BCL9L_HUMAN\|Q86UU0_S21_S25 | 1.441835472 | 3.45E-04 | 6.88E-03 |
| TB10B_HUMAN\|Q4KMP7_S656 | 1.441762594 | 5.40E-03 | 4.17E-02 |
| PKP4_HUMAN\|Q99569_S236 | 1.44165209 | 1.28E-03 | 3.41E-02 |
| PKHG3_HUMAN\|A1L390_S1079 | 1.441423772 | 2.21E-03 | 2.74E-02 |
| LMNA_HUMAN\|P02545_S616 | 1.441275366 | 1.50E-03 | 1.65E-02 |
| LARP4_HUMAN\|Q71RC2_S597 | 1.441091408 | 1.32E-03 | 9.92E-03 |
| WAC2A_HUMAN\|Q641Q2_S390 | 1.441048521 | 6.38E-03 | 3.20E-02 |
| LMNA_HUMAN\|P02545_S616 | 1.440950416 | 1.51E-03 | 1.69E-02 |
| TLN1_HUMAN\|Q9Y490_T2270 | 1.440919445 | 2.75E-03 | 2.59E-02 |
| NIBL1_HUMAN\|Q96TA1_S692_S696 | 1.440849427 | 1.98E-03 | 1.62E-02 |
| RGS3_HUMAN\|P49796_S1044 | 1.440765645 | 8.17E-04 | 1.11E-02 |
| NFIC_HUMAN\|P08651_S305 | 1.440615988 | 3.11E-03 | 1.90E-02 |

| | | | |
|---|---|---|---|
| BCLF1_HUMAN|Q9NYF8_S397_T402 | 1.440458986 | 7.34E-03 | 3.55E-02 |
| K2C6B_HUMAN|P04259_T552 | 1.440425536 | 4.09E-03 | 3.44E-02 |
| YAP1_HUMAN|P46937_T110_T119 | 1.44025065 | 6.64E-03 | 4.85E-02 |
| AFAD_HUMAN|P55196_S1172_S1173_S1182 | 1.440060224 | 2.72E-04 | 5.04E-03 |
| CR025_HUMAN|Q96B23_S66 | 1.439934095 | 3.81E-04 | 4.58E-03 |
| LARP7_HUMAN|Q4G0J3_S298_S299 | 1.439905218 | 1.67E-04 | 4.13E-03 |
| ZBT21_HUMAN|Q9ULJ3_S435_T431 | 1.439873319 | 5.61E-04 | 1.00E-02 |
| NDRG1_HUMAN|Q92597_T328 | 1.439745758 | 1.70E-03 | 1.82E-02 |
| PER1_HUMAN|O15534_S809_S815 | 1.43962448 | 4.73E-03 | 4.20E-02 |
| ABLM3_HUMAN|O94929_S503_S504 | 1.43954381 | 2.94E-03 | 1.31E-02 |
| MK01_HUMAN|P28482_Y187 | 1.439483236 | 4.20E-03 | 3.89E-02 |
| CC186_HUMAN|Q7Z3E2_S726 | 1.439420784 | 1.35E-03 | 1.17E-02 |
| PRC2A_HUMAN|P48634_S1085_S1089_S1092 | 1.439360811 | 6.40E-04 | 8.67E-03 |
| S38A2_HUMAN|Q96QD8_S17_S21 | 1.439354727 | 1.10E-02 | 4.86E-02 |
| MICA1_HUMAN|Q8TDZ2_S960 | 1.439337085 | 3.42E-04 | 7.35E-03 |
| CPNE1_HUMAN|Q99829_S54 | 1.438908018 | 4.87E-03 | 2.64E-02 |
| VIME_HUMAN|P08670_S51 | 1.438835168 | 2.88E-03 | 2.99E-02 |
| SVIL_HUMAN|O95425_S63 | 1.438804575 | 1.23E-04 | 1.97E-03 |
| NOP56_HUMAN|O00567_S570 | 1.43869349 | 3.01E-05 | 1.34E-03 |
| ROA3_HUMAN|P51991_S366 | 1.438378987 | 5.23E-03 | 4.09E-02 |
| SARG_HUMAN|Q9BW04_S502 | 1.438336048 | 3.07E-04 | 5.48E-03 |
| SRRM1_HUMAN|Q8IYB3_S738_S740_S743 | 1.438120954 | 8.32E-04 | 1.12E-02 |
| NDRG1_HUMAN|Q92597_T366 | 1.437915621 | 3.68E-03 | 2.14E-02 |
| RHG12_HUMAN|Q8IWW6_S215 | 1.4378608 | 1.08E-03 | 3.09E-02 |
| TGS1_HUMAN|Q96RS0_S154 | 1.437655719 | 2.28E-03 | 1.52E-02 |
| ABCF1_HUMAN|Q8NE71_S109 | 1.437633238 | 1.56E-03 | 8.70E-03 |
| HMCS1_HUMAN|Q01581_S516 | 1.437298666 | 3.65E-03 | 3.52E-02 |
| TKT_HUMAN|P29401_S295 | 1.437257824 | 2.19E-04 | 4.28E-03 |
| HABP4_HUMAN|Q5JVS0_S108 | 1.437248509 | 3.24E-04 | 3.36E-03 |
| P66A_HUMAN|Q86YP4_S100_S107 | 1.436961665 | 3.24E-04 | 2.27E-03 |
| K2C75_HUMAN|O95678_S535 | 1.436952414 | 2.24E-04 | 1.11E-02 |

| | | | |
|---|---|---|---|
| KLF16_HUMAN\|Q9BXK1_S226 | 1.43693751 | 1.43E-02 | 4.60E-02 |
| TGS1_HUMAN\|Q96RS0_S55_T60 | 1.43690623 | 1.12E-04 | 2.57E-03 |
| SRRM2_HUMAN\|Q9UQ35_S1483 | 1.436783863 | 1.22E-04 | 1.55E-03 |
| NUP50_HUMAN\|Q9UKX7_S221 | 1.436693484 | 4.69E-04 | 8.73E-03 |
| VIME_HUMAN\|P08670_S47 | 1.436615 | 6.50E-04 | 9.41E-03 |
| PDE1C_HUMAN\|Q14123_S469 | 1.436577888 | 1.03E-03 | 1.23E-02 |
| TB182_HUMAN\|Q9C0C2_S1666 | 1.436462124 | 1.09E-04 | 3.46E-03 |
| TACC2_HUMAN\|O95359_S2072 | 1.436408356 | 6.14E-04 | 6.53E-03 |
| PDS5B_HUMAN\|Q9NTI5_S1383 | 1.436333492 | 3.56E-03 | 2.09E-02 |
| SPTB2_HUMAN\|Q01082-3_S14 | 1.436010581 | 3.85E-03 | 4.09E-02 |
| NDRG2_HUMAN\|Q9UN36_S332_T330 | 1.435981624 | 1.17E-03 | 1.37E-02 |
| EPIPL_HUMAN\|P58107_S2716 | 1.435979368 | 8.25E-05 | 2.54E-03 |
| SRRM2_HUMAN\|Q9UQ35_S778_S780 | 1.435961061 | 1.21E-03 | 1.77E-02 |
| RPGF6_HUMAN\|Q8TEU7_S1590 | 1.435838619 | 2.51E-03 | 2.51E-02 |
| VIME_HUMAN\|P08670_S51 | 1.435670818 | 3.03E-03 | 1.86E-02 |
| PRP4B_HUMAN\|Q13523_Y849 | 1.435641848 | 9.52E-03 | 4.37E-02 |
| K1C19_HUMAN\|P08727_S35 | 1.435423143 | 5.32E-03 | 2.82E-02 |
| FA83H_HUMAN\|Q6ZRV2_S1025 | 1.435399624 | 1.61E-03 | 3.79E-02 |
| KANK1_HUMAN\|Q14678_S144 | 1.435352 | 3.57E-05 | 1.14E-03 |
| ZCH18_HUMAN\|Q86VM9_S110_S95 | 1.435268039 | 8.81E-03 | 3.14E-02 |
| ZC3H4_HUMAN\|Q9UPT8_S1114_T1106 | 1.435153982 | 2.14E-03 | 2.18E-02 |
| NHSL1_HUMAN\|Q5SYE7_S722_S723 | 1.4349582 | 7.27E-03 | 4.24E-02 |
| RABE1_HUMAN\|Q15276_S407_S410 | 1.43494831 | 2.02E-05 | 9.65E-04 |
| SH24A_HUMAN\|Q9H788_S315 | 1.434780712 | 1.12E-03 | 1.07E-02 |
| 1B07_HUMAN\|P01889_S359 | 1.4344969 | 1.05E-03 | 1.25E-02 |
| KPCD2_HUMAN\|Q9BZL6_S212 | 1.434268678 | 3.64E-03 | 3.27E-02 |
| KI13A_HUMAN\|Q9H1H9_S1698 | 1.434139547 | 1.79E-03 | 2.15E-02 |
| GRSF1_HUMAN\|Q12849_S118 | 1.434028238 | 1.41E-03 | 2.02E-02 |
| AAK1_HUMAN\|Q2M2I8_S678 | 1.433928706 | 1.58E-03 | 2.11E-02 |
| CCDC9_HUMAN\|Q9Y3X0_S386_T381 | 1.433802843 | 1.76E-03 | 4.16E-02 |
| PEAK1_HUMAN\|Q9H792_T574 | 1.433657632 | 5.21E-04 | 9.98E-03 |

| | | | |
|---|---|---|---|
| PHF14_HUMAN\|O94880_S290 | 1.433610592 | 1.90E-03 | 1.57E-02 |
| LMNA_HUMAN\|P02545_S407_S414 | 1.433551322 | 2.59E-03 | 2.49E-02 |
| IP3KB_HUMAN\|P27987_S43 | 1.433149045 | 8.31E-04 | 1.36E-02 |
| PRC2B_HUMAN\|Q5JSZ5_S1136_T1134 | 1.433037598 | 9.26E-04 | 9.35E-03 |
| ARI1A_HUMAN\|O14497_S696_S698 | 1.432987874 | 4.52E-04 | 6.60E-03 |
| CO052_HUMAN\|Q6ZUT6_S193 | 1.432836811 | 2.79E-03 | 1.74E-02 |
| COHA1_HUMAN\|Q9UMD9_S174 | 1.432756051 | 6.10E-03 | 4.65E-02 |
| CL045_HUMAN\|Q8N5I9_S15 | 1.43271444 | 1.65E-03 | 1.18E-02 |
| K2C5_HUMAN\|P13647_S62 | 1.432710824 | 2.44E-03 | 2.38E-02 |
| EPN3_HUMAN\|Q9H201_S188_S191 | 1.432490326 | 3.46E-03 | 3.07E-02 |
| RHG05_HUMAN\|Q13017_S1173 | 1.432438961 | 1.10E-03 | 1.63E-02 |
| OPTN_HUMAN\|Q96CV9_S526 | 1.432390391 | 1.00E-03 | 1.57E-02 |
| CLIP2_HUMAN\|Q9UDT6_S207_T198 | 1.432265767 | 8.33E-03 | 2.98E-02 |
| RHG32_HUMAN\|A7KAX9_S682 | 1.4321515 | 3.16E-04 | 3.95E-03 |
| AN32A_HUMAN\|P39687_S17 | 1.431944213 | 7.63E-04 | 1.27E-02 |
| NUP98_HUMAN\|P52948_S623 | 1.431817835 | 5.16E-03 | 4.79E-02 |
| K1C14_HUMAN\|P02533_S11 | 1.431415283 | 8.23E-04 | 1.27E-02 |
| LUZP1_HUMAN\|Q86V48_S574 | 1.431399886 | 9.52E-04 | 9.53E-03 |
| HSPB1_HUMAN\|P04792_S82 | 1.431322818 | 2.09E-04 | 4.86E-03 |
| PSIP1_HUMAN\|O75475_S102 | 1.431246091 | 5.28E-03 | 4.50E-02 |
| ZFN2B_HUMAN\|Q8WV99_S183 | 1.431147365 | 3.03E-03 | 2.14E-02 |
| ICE1_HUMAN\|Q9Y2F5_S1712 | 1.431067418 | 2.36E-03 | 2.62E-02 |
| CCDC9_HUMAN\|Q9Y3X0_S386 | 1.430977293 | 1.10E-04 | 3.00E-03 |
| ARI1A_HUMAN\|O14497_S696_S698_S702 | 1.430968533 | 2.24E-04 | 5.50E-03 |
| CND1_HUMAN\|Q15021_S1333 | 1.430841254 | 1.39E-03 | 1.26E-02 |
| K1C16_HUMAN\|P08779_S449 | 1.430676122 | 6.13E-03 | 4.99E-02 |
| SPD2A_HUMAN\|Q5TCZ1_S1043 | 1.430628886 | 6.31E-03 | 4.69E-02 |
| FA83H_HUMAN\|Q6ZRV2_S1003 | 1.430614123 | 9.02E-05 | 7.09E-03 |
| ZCRB1_HUMAN\|Q8TBF4_S216 | 1.430546148 | 5.52E-04 | 1.02E-02 |
| G3P_HUMAN\|P04406_S210 | 1.430519464 | 3.07E-04 | 3.24E-03 |
| GIT1_HUMAN\|Q9Y2X7_S375_S385_S388 | 1.430417505 | 3.90E-03 | 4.12E-02 |

| | | | |
|---|---|---|---|
| CSN3_HUMAN\|Q9UNS2_S421 | 1.430363951 | 8.06E-05 | 2.50E-03 |
| FA83B_HUMAN\|Q5T0W9_S869 | 1.430354119 | 1.61E-05 | 3.26E-04 |
| ACTN4_HUMAN\|O43707_S909 | 1.430339346 | 6.53E-03 | 4.88E-02 |
| IRS2_HUMAN\|Q9Y4H2_S1186_T1202 | 1.430224795 | 2.64E-04 | 3.51E-03 |
| SRRM2_HUMAN\|Q9UQ35_S377_S387_S398 | 1.430031154 | 1.38E-03 | 1.19E-02 |
| P63_HUMAN\|Q9H3D4_S452_S463 | 1.429873865 | 1.07E-02 | 4.25E-02 |
| IRS2_HUMAN\|Q9Y4H2_T520 | 1.429801736 | 6.20E-03 | 3.13E-02 |
| SRG2B_HUMAN\|P0DMP2_T203 | 1.429743395 | 1.56E-04 | 1.27E-03 |
| SMG9_HUMAN\|Q9H0W8_S53 | 1.429418965 | 3.51E-04 | 3.57E-03 |
| NCOA7_HUMAN\|Q8NI08_S208_T210 | 1.429219992 | 1.10E-03 | 6.57E-03 |
| ARAF_HUMAN\|P10398_S157 | 1.429185385 | 5.89E-04 | 8.80E-03 |
| CDK9_HUMAN\|P50750_T186 | 1.429159986 | 2.34E-03 | 2.31E-02 |
| AHNK_HUMAN\|Q09666_S210_S216_S220 | 1.429082996 | 7.95E-04 | 1.25E-02 |
| SMC1A_HUMAN\|Q14683_S957 | 1.428911669 | 5.24E-04 | 5.83E-03 |
| RNF25_HUMAN\|Q96BH1_S450 | 1.428763005 | 4.51E-03 | 3.02E-02 |
| IRS1_HUMAN\|P35568_S1100 | 1.428748891 | 1.24E-03 | 3.33E-02 |
| NFIB_HUMAN\|O00712_S276_S295 | 1.428682059 | 7.03E-03 | 3.98E-02 |
| K2C5_HUMAN\|P13647_S574 | 1.428629669 | 1.51E-03 | 3.85E-02 |
| AHNK_HUMAN\|Q09666_S5749_S5752 | 1.428624523 | 8.11E-04 | 1.03E-02 |
| K1C16_HUMAN\|P08779_S448_S451 | 1.428606298 | 3.95E-03 | 3.44E-02 |
| FA83B_HUMAN\|Q5T0W9_S388 | 1.428335247 | 5.39E-03 | 4.26E-02 |
| P66B_HUMAN\|Q8WXI9_S122_T120 | 1.42825431 | 1.75E-03 | 1.88E-02 |
| I2BP2_HUMAN\|Q7Z5L9_S455_S460 | 1.428164872 | 2.56E-03 | 3.01E-02 |
| MISP_HUMAN\|Q8IVT2_S394 | 1.427861884 | 3.53E-03 | 3.45E-02 |
| SPTB2_HUMAN\|Q01082-3_S10 | 1.42769888 | 2.34E-03 | 2.38E-02 |
| K2C5_HUMAN\|P13647_S574 | 1.427588703 | 1.54E-03 | 1.68E-02 |
| FA76B_HUMAN\|Q5HYJ3_S338 | 1.42758555 | 4.45E-03 | 2.47E-02 |
| K1671_HUMAN\|Q9BY89_S1441 | 1.427525716 | 8.17E-03 | 3.89E-02 |
| PCY1A_HUMAN\|P49585_S343_S347 | 1.427476087 | 1.26E-03 | 9.57E-03 |
| MON1A_HUMAN\|Q86VX9_S95 | 1.427397129 | 8.03E-04 | 1.32E-02 |
| TFE3_HUMAN\|P19532_S568_S570 | 1.427327455 | 6.29E-04 | 6.59E-03 |

| | | | |
|---|---|---|---|
| IRS2_HUMAN\|Q9Y4H2_S679 | 1.427274366 | 1.49E-03 | 1.68E-02 |
| BCL10_HUMAN\|O95999_S138 | 1.427235256 | 2.92E-03 | 3.17E-02 |
| K2C5_HUMAN\|P13647_S82_T78 | 1.427111033 | 1.82E-03 | 4.25E-02 |
| EXOC7_HUMAN\|Q9UPT5_S250 | 1.427090732 | 1.29E-03 | 9.73E-03 |
| MISP_HUMAN\|Q8IVT2_S543 | 1.427084117 | 2.09E-03 | 3.48E-02 |
| DCAF6_HUMAN\|Q58WW2_S657 | 1.427027453 | 4.82E-03 | 3.19E-02 |
| BORG2_HUMAN\|Q9UKI2_S89 | 1.427007508 | 2.90E-04 | 4.72E-03 |
| CTR1_HUMAN\|P30825_T620 | 1.427003133 | 8.67E-03 | 4.67E-02 |
| SYNEM_HUMAN\|O15061_S1044_S1049 | 1.426793363 | 2.05E-03 | 3.47E-02 |
| CR025_HUMAN\|Q96B23_S143_S145_S147 | 1.426668555 | 5.89E-03 | 3.67E-02 |
| PARP4_HUMAN\|Q9UKK3_S1722 | 1.426553669 | 1.72E-03 | 8.51E-03 |
| ZC3H1_HUMAN\|O60293_S805_S809 | 1.42648629 | 6.01E-03 | 2.65E-02 |
| GIT1_HUMAN\|Q9Y2X7_S388_T392 | 1.42645436 | 9.02E-03 | 4.93E-02 |
| TCOF_HUMAN\|Q13428_T249 | 1.426450924 | 3.52E-03 | 2.08E-02 |
| ACINU_HUMAN\|Q9UKV3_S240 | 1.426057263 | 2.96E-03 | 3.32E-02 |
| SAMH1_HUMAN\|Q9Y3Z3_S33 | 1.426047324 | 2.46E-03 | 2.70E-02 |
| HAP28_HUMAN\|Q13442_S60 | 1.425992 | 1.25E-03 | 7.26E-03 |
| MILK1_HUMAN\|Q8N3F8_S578 | 1.425898569 | 3.92E-04 | 3.90E-03 |
| HNRPC_HUMAN\|P07910_S260 | 1.425889527 | 4.04E-04 | 3.99E-03 |
| PACS1_HUMAN\|Q6VY07_T504 | 1.425800065 | 5.35E-03 | 2.83E-02 |
| LAP2A_HUMAN\|P42166_S159 | 1.42566926 | 2.00E-03 | 2.06E-02 |
| RRP1B_HUMAN\|Q14684_S245 | 1.425647734 | 3.62E-04 | 7.29E-03 |
| APC_HUMAN\|P25054_S2242_S2244 | 1.42557873 | 1.91E-04 | 9.84E-03 |
| F120A_HUMAN\|Q9NZB2_S1023 | 1.425578449 | 9.70E-04 | 1.43E-02 |
| ATX2L_HUMAN\|Q8WWM7_S339 | 1.425455723 | 4.44E-05 | 3.70E-03 |
| NCK5L_HUMAN\|Q9HCH0_S489_S494 | 1.425449547 | 1.47E-03 | 1.67E-02 |
| SPTB2_HUMAN\|Q01082_S2341 | 1.424967356 | 3.99E-05 | 6.47E-04 |
| LMNB2_HUMAN\|Q03252_S421_S424 | 1.424890894 | 3.98E-03 | 3.46E-02 |
| AHNK_HUMAN\|Q09666_S212_S216 | 1.424804622 | 6.64E-04 | 2.20E-02 |
| IASPP_HUMAN\|Q8WUF5_S113 | 1.424784851 | 9.95E-04 | 1.27E-02 |
| ARI1A_HUMAN\|O14497_S696_S698_S702 | 1.424642256 | 2.58E-04 | 5.64E-03 |

| | | | |
|---|---|---|---|
| PKHG3_HUMAN\|A1L390_S640_S643 | 1.424582787 | 4.46E-03 | 2.07E-02 |
| RIOK2_HUMAN\|Q9BVS4_S382_S385 | 1.424582129 | 3.13E-04 | 3.28E-03 |
| PP1R7_HUMAN\|Q15435_S12 | 1.424523883 | 5.56E-03 | 4.69E-02 |
| VIME_HUMAN\|P08670_S66 | 1.424417077 | 9.73E-04 | 1.43E-02 |
| ACINU_HUMAN\|Q9UKV3_S240 | 1.42427943 | 3.05E-03 | 2.22E-02 |
| LMNB2_HUMAN\|Q03252_S420 | 1.424256563 | 1.34E-03 | 1.53E-02 |
| NDRG1_HUMAN\|Q92597_S336 | 1.424093064 | 1.36E-03 | 1.18E-02 |
| ARAF_HUMAN\|P10398_S157 | 1.424022062 | 6.51E-04 | 1.11E-02 |
| PLEC_HUMAN\|Q15149_S4385_S4386 | 1.423990268 | 2.86E-05 | 3.40E-04 |
| RL19_HUMAN\|P84098_S13 | 1.423971445 | 2.82E-03 | 2.73E-02 |
| FIL1L_HUMAN\|Q4L180_T986 | 1.423918139 | 3.43E-03 | 2.05E-02 |
| RAE1_HUMAN\|P24386_S651 | 1.423802258 | 5.33E-04 | 4.90E-02 |
| PAK2_HUMAN\|Q13177_S141 | 1.423631005 | 5.73E-03 | 4.38E-02 |
| ZN185_HUMAN\|O15231_S521 | 1.423392159 | 9.40E-04 | 1.22E-02 |
| NU153_HUMAN\|P49790_S334 | 1.42338827 | 2.20E-03 | 2.71E-02 |
| MPZL2_HUMAN\|O60487_S208 | 1.42334004 | 5.15E-03 | 2.75E-02 |
| CBX3_HUMAN\|Q13185_S176 | 1.423163787 | 5.61E-03 | 2.91E-02 |
| PP4R1_HUMAN\|Q8TF05_S948_T945 | 1.423120508 | 2.36E-03 | 2.62E-02 |
| HS90A_HUMAN\|P07900_S263 | 1.423081143 | 2.49E-03 | 1.61E-02 |
| CNN3_HUMAN\|Q15417_S323 | 1.423008234 | 3.46E-03 | 3.80E-02 |
| KMT2C_HUMAN\|Q8NEZ4_S111 | 1.422951205 | 1.39E-03 | 7.93E-03 |
| MICA1_HUMAN\|Q8TDZ2_S960 | 1.422867735 | 4.82E-04 | 9.51E-03 |
| ADRM1_HUMAN\|Q16186_S211 | 1.422765553 | 3.47E-03 | 3.16E-02 |
| MISP_HUMAN\|Q8IVT2_T377 | 1.422676749 | 2.55E-03 | 1.64E-02 |
| NIBL1_HUMAN\|Q96TA1_S696 | 1.422589351 | 1.80E-03 | 4.13E-02 |
| SGTA_HUMAN\|O43765_S77_T81 | 1.422262059 | 1.92E-03 | 1.33E-02 |
| KI13A_HUMAN\|Q9H1H9_S1698 | 1.42202603 | 2.18E-03 | 1.47E-02 |
| CTNB1_HUMAN\|P35222_S552 | 1.422000423 | 3.02E-03 | 2.77E-02 |
| K1671_HUMAN\|Q9BY89_S508 | 1.421943111 | 3.81E-03 | 3.86E-02 |
| IRS2_HUMAN\|Q9Y4H2_S620 | 1.421742642 | 4.74E-03 | 2.58E-02 |
| FA53C_HUMAN\|Q9NYF3_S122 | 1.421704627 | 8.06E-04 | 1.03E-02 |

| | | | |
|---|---|---|---|
| PKP4_HUMAN\|Q99569_S233 | 1.421522056 | 1.01E-03 | 1.22E-02 |
| K1C14_HUMAN\|P02533_S462 | 1.421430629 | 9.80E-03 | 4.48E-02 |
| K2C5_HUMAN\|P13647_Y66 | 1.421375625 | 4.81E-04 | 7.45E-03 |
| MICA3_HUMAN\|Q7RTP6_S977 | 1.421345748 | 3.55E-05 | 1.14E-03 |
| NDRG1_HUMAN\|Q92597_T366 | 1.421325805 | 4.73E-03 | 3.80E-02 |
| FA83H_HUMAN\|Q6ZRV2_S1098 | 1.421212583 | 1.27E-03 | 9.63E-03 |
| ACINU_HUMAN\|Q9UKV3_S243 | 1.42091358 | 6.32E-04 | 2.19E-02 |
| ZCH18_HUMAN\|Q86VM9_S532_S534 | 1.420904764 | 8.00E-05 | 5.42E-03 |
| SRRM2_HUMAN\|Q9UQ35_S1404 | 1.420707074 | 8.15E-04 | 5.18E-03 |
| SPTB2_HUMAN\|Q01082-3_S14 | 1.420600196 | 4.84E-03 | 4.54E-02 |
| PRSR2_HUMAN\|Q86WR7_S31 | 1.42059358 | 7.77E-04 | 8.10E-03 |
| ACSA_HUMAN\|Q9NR19_S267 | 1.420547031 | 2.52E-03 | 2.51E-02 |
| IF4B_HUMAN\|P23588_S489_S497 | 1.420524286 | 6.44E-03 | 3.73E-02 |
| AHNK2_HUMAN\|Q8IVF2_S5175 | 1.42051597 | 4.31E-03 | 2.94E-02 |
| PYGO2_HUMAN\|Q9BRQ0_T302 | 1.420509843 | 1.23E-03 | 9.43E-03 |
| SH24A_HUMAN\|Q9H788_S315 | 1.420284413 | 1.46E-03 | 1.93E-02 |
| UN45A_HUMAN\|Q9H3U1_S147_Y145 | 1.42025635 | 2.17E-03 | 4.52E-02 |
| RHG05_HUMAN\|Q13017_S1218 | 1.420051213 | 1.08E-03 | 3.09E-02 |
| CIRBP_HUMAN\|Q14011_S148 | 1.419989044 | 1.04E-03 | 1.31E-02 |
| NIBL1_HUMAN\|Q96TA1_S696 | 1.419916266 | 1.89E-03 | 2.22E-02 |
| CCNH_HUMAN\|P51946_T315 | 1.419368728 | 6.02E-03 | 4.54E-02 |
| TBCD4_HUMAN\|O60343_T752 | 1.419337572 | 6.36E-03 | 2.39E-02 |
| K1C14_HUMAN\|P02533_S12 | 1.41929328 | 1.44E-03 | 1.85E-02 |
| SPT5H_HUMAN\|O00267_S666 | 1.41928883 | 3.07E-04 | 5.48E-03 |
| MINT_HUMAN\|Q96T58_S727 | 1.419198684 | 2.84E-03 | 3.26E-02 |
| LAP2A_HUMAN\|P42166_S66_S67_T74 | 1.419056602 | 4.10E-03 | 2.32E-02 |
| PHIP_HUMAN\|Q8WWQ0_S1783 | 1.419012413 | 6.01E-04 | 3.65E-03 |
| RAI14_HUMAN\|Q9P0K7_S281 | 1.418800756 | 7.64E-04 | 1.24E-02 |
| GGA2_HUMAN\|Q9UJY4_S177 | 1.41873919 | 6.55E-03 | 4.89E-02 |
| PKP3_HUMAN\|Q9Y446_S314 | 1.418647053 | 8.30E-04 | 1.36E-02 |
| RGL2_HUMAN\|O15211_S761 | 1.418602041 | 2.29E-03 | 2.34E-02 |

| | | | |
|---|---|---|---|
| SMC1A_HUMAN\|Q14683_S957 | 1.418547725 | 6.46E-04 | 1.11E-02 |
| ZCRB1_HUMAN\|Q8TBF4_S216 | 1.418329518 | 7.05E-04 | 1.22E-02 |
| ABCF1_HUMAN\|Q8NE71_S109_T108 | 1.418328758 | 2.55E-04 | 4.39E-03 |
| RAD9A_HUMAN\|Q99638_S387 | 1.418318901 | 4.05E-03 | 3.79E-02 |
| RNF25_HUMAN\|Q96BH1_S450 | 1.418306867 | 5.24E-03 | 4.09E-02 |
| NFIC_HUMAN\|P08651_S333_S339 | 1.418012706 | 2.98E-03 | 1.84E-02 |
| DDX3Y_HUMAN\|O15523_S607 | 1.417923195 | 1.34E-03 | 1.53E-02 |
| SCAF8_HUMAN\|Q9UPN6_S617 | 1.417906759 | 2.31E-03 | 2.58E-02 |
| SKT_HUMAN\|Q5T5P2_S1794 | 1.417673368 | 2.36E-03 | 2.62E-02 |
| KLF16_HUMAN\|Q9BXK1_S238 | 1.417592694 | 5.82E-03 | 4.50E-02 |
| APC_HUMAN\|P25054_S2242_S2244 | 1.417240383 | 2.32E-04 | 5.61E-03 |
| PLEC_HUMAN\|Q15149_S4622_Y4615 | 1.416750553 | 9.14E-04 | 5.65E-03 |
| ABLM1_HUMAN\|O14639_S431 | 1.416649002 | 5.30E-03 | 2.08E-02 |
| KLC3_HUMAN\|Q6P597_S502 | 1.416581993 | 2.48E-05 | 1.14E-03 |
| IASPP_HUMAN\|Q8WUF5_S74 | 1.416493203 | 2.24E-04 | 4.35E-03 |
| STRN3_HUMAN\|Q13033_S202 | 1.416324238 | 4.45E-04 | 7.09E-03 |
| PRAG1_HUMAN\|Q86YV5_S826 | 1.416319432 | 1.69E-03 | 2.15E-02 |
| TB182_HUMAN\|Q9C0C2_S1158 | 1.416301115 | 4.55E-03 | 2.51E-02 |
| UN45A_HUMAN\|Q9H3U1_S147_Y145 | 1.416067145 | 2.32E-03 | 1.74E-02 |
| BAG3_HUMAN\|O95817_S289 | 1.416021276 | 6.11E-03 | 4.60E-02 |
| CE170_HUMAN\|Q5SW79_S636 | 1.416000193 | 3.60E-05 | 6.05E-04 |
| CE170_HUMAN\|Q5SW79_S636 | 1.415807178 | 3.62E-05 | 1.15E-03 |
| SRRM2_HUMAN\|Q9UQ35_S1404 | 1.415693315 | 8.99E-04 | 5.06E-03 |
| TPIS_HUMAN\|P60174_Y246 | 1.415666828 | 1.76E-03 | 2.13E-02 |
| SRSF1_HUMAN\|Q07955_S234_S238 | 1.415592697 | 8.76E-04 | 1.91E-02 |
| DC1L1_HUMAN\|Q9Y6G9_S510_S516 | 1.415471375 | 1.36E-03 | 1.91E-02 |
| TRI22_HUMAN\|Q8IYM9_S384 | 1.415446663 | 5.83E-03 | 4.50E-02 |
| PSF2_HUMAN\|Q9Y248_S179_S182 | 1.415446049 | 3.21E-03 | 3.22E-02 |
| CBLB_HUMAN\|Q13191_S480_S484 | 1.415125181 | 6.08E-04 | 6.56E-03 |
| PTN12_HUMAN\|Q05209_S571_T587 | 1.415027401 | 7.16E-04 | 1.21E-02 |
| SCAF8_HUMAN\|Q9UPN6_S617 | 1.415016901 | 2.42E-03 | 1.79E-02 |

| | | | |
|---|---|---|---|
| VIP2_HUMAN\|O43314_S1006 | 1.414703642 | 5.27E-04 | 5.83E-03 |
| K1C14_HUMAN\|P02533_S11 | 1.414690512 | 1.14E-03 | 3.17E-02 |
| NOS1_HUMAN\|P29475_S852 | 1.414573301 | 9.22E-03 | 4.85E-02 |
| SIPA1_HUMAN\|Q96FS4_S1036 | 1.41454811 | 2.57E-03 | 3.01E-02 |
| IF6_HUMAN\|P56537_T245 | 1.414344218 | 1.17E-03 | 6.23E-03 |
| IF2P_HUMAN\|O60841_S182_S183 | 1.414286843 | 7.49E-04 | 2.45E-02 |
| MYPT1_HUMAN\|O14974_S507 | 1.413884453 | 2.75E-03 | 2.67E-02 |
| TB182_HUMAN\|Q9C0C2_S494 | 1.413871513 | 3.54E-03 | 4.87E-02 |
| RBM27_HUMAN\|Q9P2N5_T447 | 1.413699009 | 2.47E-03 | 4.83E-02 |
| PHF2_HUMAN\|O75151_S1058 | 1.413690327 | 6.45E-03 | 2.82E-02 |
| NUMA1_HUMAN\|Q14980_S1757 | 1.413416333 | 1.85E-03 | 9.01E-03 |
| ASPP2_HUMAN\|Q13625_S558 | 1.413399076 | 5.64E-04 | 6.19E-03 |
| ARI1A_HUMAN\|O14497_S696_S698_S702 | 1.413198808 | 3.35E-04 | 5.79E-03 |
| SRPK2_HUMAN\|P78362_S497_T492 | 1.413146095 | 1.23E-03 | 1.77E-02 |
| NU188_HUMAN\|Q5SRE5_S1709 | 1.412842788 | 1.33E-03 | 1.83E-02 |
| TANC2_HUMAN\|Q9HCD6_T168 | 1.412774344 | 6.89E-04 | 4.06E-03 |
| ITBP1_HUMAN\|O14713_S10_S14 | 1.412740556 | 1.74E-03 | 1.88E-02 |
| K1522_HUMAN\|Q9P206_S929 | 1.412671823 | 6.23E-04 | 1.12E-02 |
| ARHGH_HUMAN\|Q96PE2_S619 | 1.412453713 | 2.50E-03 | 2.83E-02 |
| PHLB2_HUMAN\|Q86SQ0_S565 | 1.412425595 | 8.46E-04 | 8.65E-03 |
| MAP1B_HUMAN\|P46821_S1785 | 1.412321985 | 1.61E-04 | 4.02E-03 |
| PPR18_HUMAN\|Q6NYC8_S209_S213 | 1.412287431 | 9.43E-04 | 2.82E-02 |
| BABA1_HUMAN\|Q9NWV8_S57 | 1.412276822 | 7.32E-03 | 4.09E-02 |
| AHNK_HUMAN\|Q09666_S210 | 1.412270174 | 3.29E-04 | 4.04E-03 |
| PLEC_HUMAN\|Q15149_S4626_S4630 | 1.412238716 | 5.63E-04 | 3.46E-03 |
| UBP24_HUMAN\|Q9UPU5_S1143 | 1.412236235 | 1.96E-03 | 1.35E-02 |
| TBCD4_HUMAN\|O60343_T752 | 1.412169449 | 7.03E-03 | 3.98E-02 |
| SRC8_HUMAN\|Q14247_T399_T411 | 1.412054974 | 1.84E-03 | 2.43E-02 |
| FRM4B_HUMAN\|Q9Y2L6_S628 | 1.41199811 | 4.54E-03 | 4.33E-02 |
| MISP_HUMAN\|Q8IVT2_S376 | 1.411766785 | 2.94E-03 | 1.82E-02 |
| SGTA_HUMAN\|O43765_S305 | 1.41148364 | 2.28E-03 | 1.71E-02 |

| | | | |
|---|---|---|---|
| OPTN_HUMAN\|Q96CV9_T204 | 1.411022938 | 4.59E-04 | 5.23E-03 |
| SLK_HUMAN\|Q9H2G2_T820 | 1.410913179 | 3.22E-04 | 5.09E-03 |
| VIGLN_HUMAN\|Q00341_S31 | 1.410898669 | 1.04E-03 | 1.48E-02 |
| SRRM2_HUMAN\|Q9UQ35_S2449 | 1.410876503 | 3.46E-03 | 3.07E-02 |
| K1C14_HUMAN\|P02533_S53 | 1.410679777 | 1.31E-03 | 1.74E-02 |
| PLEC_HUMAN\|Q15149_S4386_S4389 | 1.410562847 | 1.66E-03 | 2.05E-02 |
| RABE1_HUMAN\|Q15276_T408 | 1.410556703 | 4.44E-04 | 5.11E-03 |
| DGC14_HUMAN\|Q96DF8_T386 | 1.410510124 | 9.99E-04 | 7.95E-03 |
| CUL4B_HUMAN\|Q13620_T49 | 1.410439002 | 5.32E-03 | 2.09E-02 |
| RRP1B_HUMAN\|Q14684_S245 | 1.410284757 | 5.08E-04 | 9.34E-03 |
| NDRG1_HUMAN\|Q92597_S330_T328 | 1.410034238 | 3.34E-03 | 2.01E-02 |
| RAI14_HUMAN\|Q9P0K7-4_S261 | 1.41000773 | 3.83E-03 | 1.63E-02 |
| ASAP1_HUMAN\|Q9ULH1_T58_T69 | 1.409986196 | 8.93E-03 | 3.66E-02 |
| K2C80_HUMAN\|Q6KB66_S451 | 1.409981876 | 2.72E-03 | 4.16E-02 |
| CCM2_HUMAN\|Q9BSQ5_S384 | 1.409797452 | 4.12E-03 | 3.55E-02 |
| IF2P_HUMAN\|O60841_S182_S183_S190 | 1.409787521 | 2.56E-03 | 1.94E-02 |
| BRAF_HUMAN\|P15056_S446 | 1.409678855 | 8.71E-03 | 3.60E-02 |
| GYS1_HUMAN\|P13807_T703 | 1.409673152 | 3.66E-03 | 1.76E-02 |
| AKTS1_HUMAN\|Q96B36_S202_S203 | 1.409638593 | 5.51E-04 | 9.96E-03 |
| PSIP1_HUMAN\|O75475_S273_T272 | 1.409473499 | 2.00E-03 | 1.37E-02 |
| COBL1_HUMAN\|Q53SF7_S1145 | 1.409376306 | 5.54E-03 | 2.49E-02 |
| ATX2L_HUMAN\|Q8WWM7_S680_S684 | 1.409349928 | 2.70E-04 | 4.53E-03 |
| SLK_HUMAN\|Q9H2G2_S655 | 1.409277966 | 1.00E-02 | 4.54E-02 |
| KLC3_HUMAN\|Q6P597_S497_S502 | 1.409196902 | 1.77E-04 | 4.64E-03 |
| T2FA_HUMAN\|P35269_S433 | 1.409190693 | 6.14E-04 | 5.57E-03 |
| KLC2_HUMAN\|Q9H0B6_S608_S611 | 1.409160876 | 3.36E-04 | 4.15E-03 |
| PDE1C_HUMAN\|Q14123_S469 | 1.408837403 | 1.73E-03 | 1.87E-02 |
| DBNL_HUMAN\|Q9UJU6_S269 | 1.408812332 | 9.40E-03 | 4.89E-02 |
| NIPA_HUMAN\|Q86WB0_S408_S409 | 1.408771314 | 1.66E-04 | 9.01E-03 |
| NRK_HUMAN\|Q7Z2Y5_S852_Y858 | 1.408733474 | 7.08E-03 | 2.60E-02 |
| CAMP2_HUMAN\|Q08AD1_S1313 | 1.408634715 | 2.97E-03 | 3.08E-02 |

| | | | |
|---|---|---|---|
| BAD_HUMAN\|Q92934_S74 | 1.408573695 | 4.00E-03 | 4.22E-02 |
| OPTN_HUMAN\|Q96CV9_T525 | 1.408374614 | 7.25E-04 | 1.26E-02 |
| K2C5_HUMAN\|P13647_S11_S8 | 1.40829432 | 1.47E-03 | 3.78E-02 |
| IASPP_HUMAN\|Q8WUF5_S567 | 1.408212126 | 2.11E-04 | 4.86E-03 |
| SH23A_HUMAN\|Q9BRG2_S147 | 1.408035639 | 3.28E-03 | 1.99E-02 |
| IP3KB_HUMAN\|P27987_S43 | 1.407873684 | 1.34E-03 | 1.23E-02 |
| DC1L1_HUMAN\|Q9Y6G9_S516 | 1.407829925 | 9.89E-04 | 6.01E-03 |
| LMNA_HUMAN\|P02545_S428 | 1.407475846 | 6.18E-04 | 2.19E-02 |
| CRBG1_HUMAN\|Q9Y4K1_S348 | 1.407431614 | 2.27E-03 | 2.55E-02 |
| MYO9A_HUMAN\|B2RTY4_S1733 | 1.407415113 | 8.84E-03 | 3.14E-02 |
| LMNB2_HUMAN\|Q03252_S421_S424 | 1.407193277 | 5.27E-03 | 2.80E-02 |
| PAXI_HUMAN\|P49023_S106 | 1.407112784 | 4.76E-03 | 4.55E-02 |
| IPP2M_HUMAN\|Q6NXS1_S121 | 1.407105381 | 9.28E-03 | 3.27E-02 |
| RLGPB_HUMAN\|Q86X10_S359 | 1.407089173 | 7.98E-03 | 3.36E-02 |
| K0930_HUMAN\|Q6ICG6_S267 | 1.407063173 | 2.27E-03 | 2.56E-02 |
| TB182_HUMAN\|Q9C0C2_S936 | 1.406878566 | 1.38E-03 | 1.81E-02 |
| SVIL_HUMAN\|O95425_T679 | 1.406792361 | 2.40E-04 | 1.17E-02 |
| CHM2A_HUMAN\|O43633_S203 | 1.406708887 | 1.20E-03 | 1.40E-02 |
| YTDC2_HUMAN\|Q9H6S0_S1202 | 1.406677114 | 1.51E-03 | 1.99E-02 |
| NIBL1_HUMAN\|Q96TA1_S692_S696 | 1.406632074 | 3.42E-03 | 3.57E-02 |
| PALLD_HUMAN\|Q8WX93_S893 | 1.406591797 | 5.08E-03 | 4.74E-02 |
| CCDC9_HUMAN\|Q9Y3X0_T389 | 1.406441288 | 2.10E-04 | 5.17E-03 |
| EGFR_HUMAN\|P00533_S991 | 1.406416689 | 2.97E-03 | 1.49E-02 |
| LARP1_HUMAN\|Q6PKG0_S1040 | 1.406416293 | 1.26E-03 | 9.59E-03 |
| NIBAN_HUMAN\|Q9BZQ8_S926 | 1.406390151 | 1.28E-03 | 1.89E-02 |
| TAOK3_HUMAN\|Q9H2K8_S442 | 1.406358388 | 6.63E-03 | 3.29E-02 |
| LUZP1_HUMAN\|Q86V48_S575 | 1.40613582 | 3.13E-04 | 5.49E-03 |
| STR3N_HUMAN\|O95772_S39 | 1.406126336 | 1.23E-02 | 4.05E-02 |
| TFE3_HUMAN\|P19532_S548_S556 | 1.406091562 | 4.95E-03 | 3.24E-02 |
| IWS1_HUMAN\|Q96ST2_S511 | 1.40585897 | 1.27E-03 | 7.35E-03 |
| AHNK_HUMAN\|Q09666_T4564 | 1.405808004 | 5.65E-03 | 4.41E-02 |

| | | | |
|---|---|---|---|
| K1C14_HUMAN|P02533_S54 | 1.405796235 | 1.10E-03 | 1.55E-02 |
| LARP1_HUMAN|Q6PKG0_S517_S521 | 1.405731927 | 3.23E-04 | 5.59E-03 |
| PAWR_HUMAN|Q96IZ0_S231 | 1.4056027 | 2.18E-03 | 4.81E-02 |
| KLC3_HUMAN|Q6P597_S167_S173 | 1.405325246 | 6.18E-03 | 3.13E-02 |
| IRS2_HUMAN|Q9Y4H2_T350 | 1.405300384 | 8.10E-04 | 1.10E-02 |
| PTN12_HUMAN|Q05209_S588 | 1.405132847 | 4.20E-03 | 4.36E-02 |
| NDRG1_HUMAN|Q92597_S330_S333 | 1.405121956 | 2.13E-03 | 2.20E-02 |
| AKAP2_HUMAN|Q9Y2D5_T157 | 1.404922896 | 3.79E-03 | 3.92E-02 |
| CLAP2_HUMAN|O75122_S1029 | 1.404723427 | 5.98E-03 | 4.59E-02 |
| ACINU_HUMAN|Q9UKV3_S243 | 1.404674821 | 8.76E-04 | 8.66E-03 |
| UHRF2_HUMAN|Q96PU4_S667 | 1.404648288 | 2.99E-04 | 3.83E-03 |
| CAAP1_HUMAN|Q9H8G2_S92 | 1.404542497 | 8.94E-04 | 5.56E-03 |
| NFIC_HUMAN|P08651_S294_S305 | 1.40450086 | 4.18E-03 | 3.88E-02 |
| PCM1_HUMAN|Q15154_S65_S68 | 1.404426001 | 1.67E-03 | 1.19E-02 |
| DKC1_HUMAN|O60832_S485_T496 | 1.404362578 | 2.57E-04 | 1.29E-02 |
| KC1D_HUMAN|P48730_S331 | 1.404255238 | 1.07E-02 | 4.78E-02 |
| NDRG1_HUMAN|Q92597_S330_S333 | 1.40421697 | 2.17E-03 | 2.20E-02 |
| CYTSA_HUMAN|Q69YQ0_S832 | 1.404194657 | 1.20E-04 | 1.87E-03 |
| YAP1_HUMAN|P46937_S109_T119 | 1.403979608 | 2.71E-04 | 1.32E-02 |
| AHNK_HUMAN|Q09666_S210_S216 | 1.403942786 | 2.16E-03 | 2.68E-02 |
| BUD13_HUMAN|Q9BRD0_T159 | 1.403835811 | 4.28E-03 | 2.01E-02 |
| MINT_HUMAN|Q96T58_S623 | 1.403776185 | 1.04E-04 | 9.39E-04 |
| PRAG1_HUMAN|Q86YV5_S826 | 1.403767587 | 2.11E-03 | 1.63E-02 |
| KLC3_HUMAN|Q6P597_S497_S502 | 1.403656956 | 2.03E-04 | 4.02E-03 |
| M4K4_HUMAN|O95819_S842 | 1.403587067 | 1.35E-03 | 1.60E-02 |
| TAF6_HUMAN|P49848_S532 | 1.403448317 | 6.41E-04 | 3.85E-03 |
| ACINU_HUMAN|Q9UKV3_S838 | 1.403424899 | 7.91E-04 | 1.25E-02 |
| KLC2_HUMAN|Q9H0B6_S610 | 1.403408886 | 2.52E-04 | 4.77E-03 |
| IF2P_HUMAN|O60841_S182_S183_S190 | 1.403398676 | 2.85E-03 | 1.28E-02 |
| ARHGG_HUMAN|Q5VV41_S227_T226 | 1.403287 | 1.15E-03 | 1.68E-02 |
| COMT_HUMAN|P21964_S267 | 1.403049142 | 1.90E-03 | 1.49E-02 |

| | | | |
|---|---|---|---|
| K1522_HUMAN\|Q9P206_S406 | 1.402962558 | 5.63E-03 | 2.91E-02 |
| AHNK_HUMAN\|Q09666_T4564 | 1.402944793 | 5.90E-03 | 4.88E-02 |
| NFIC_HUMAN\|P08651_S305 | 1.402887246 | 5.62E-03 | 4.73E-02 |
| EGFR_HUMAN\|P00533_S991 | 1.402844617 | 3.17E-03 | 3.47E-02 |
| DDX3Y_HUMAN\|O15523_S607 | 1.402829671 | 1.78E-03 | 1.41E-02 |
| K2C5_HUMAN\|P13647_S50 | 1.402720123 | 1.45E-03 | 1.86E-02 |
| IRS2_HUMAN\|Q9Y4H2_S608 | 1.402296434 | 1.07E-03 | 3.09E-02 |
| HSPB1_HUMAN\|P04792_S199 | 1.402291292 | 1.72E-03 | 1.39E-02 |
| PKP4_HUMAN\|Q99569_S336 | 1.402264725 | 1.53E-03 | 1.11E-02 |
| K1C16_HUMAN\|P08779_S468 | 1.402225466 | 3.12E-03 | 2.95E-02 |
| BRD4_HUMAN\|O60885_S470 | 1.402209013 | 7.28E-03 | 3.54E-02 |
| PKP3_HUMAN\|Q9Y446_Y176 | 1.402107129 | 4.53E-03 | 4.06E-02 |
| DDX3X_HUMAN\|O00571_S90 | 1.402052139 | 4.91E-03 | 4.01E-02 |
| 3BP1_HUMAN\|Q9Y3L3_S695 | 1.402050348 | 5.00E-03 | 4.06E-02 |
| GRAP1_HUMAN\|Q4V328_S666 | 1.401850546 | 6.63E-03 | 3.29E-02 |
| NIBL1_HUMAN\|Q96TA1_S691 | 1.401772477 | 1.98E-04 | 3.96E-03 |
| OSBP1_HUMAN\|P22059_S379_S382 | 1.401679103 | 7.66E-04 | 1.77E-02 |
| DDX20_HUMAN\|Q9UHI6_S677_Y673 | 1.401543579 | 2.42E-03 | 1.26E-02 |
| ARH40_HUMAN\|Q8TER5_S262 | 1.401478133 | 9.01E-04 | 2.75E-02 |
| NUP53_HUMAN\|Q8NFH5_T280 | 1.401458606 | 2.54E-04 | 4.39E-03 |
| DGC14_HUMAN\|Q96DF8_T386 | 1.401440838 | 1.20E-03 | 1.47E-02 |
| FA83H_HUMAN\|Q6ZRV2_Y875 | 1.401392717 | 2.96E-04 | 3.14E-03 |
| KLC2_HUMAN\|Q9H0B6_S611 | 1.400909698 | 4.41E-03 | 3.61E-02 |
| SMC1A_HUMAN\|Q14683_S957 | 1.400773128 | 9.27E-04 | 1.38E-02 |
| NIPA_HUMAN\|Q86WB0_S409 | 1.400672396 | 7.18E-05 | 6.89E-04 |
| NBN_HUMAN\|O60934_S343 | 1.40066411 | 3.18E-03 | 3.21E-02 |
| TB182_HUMAN\|Q9C0C2_S1666 | 1.400600569 | 2.61E-04 | 1.90E-03 |
| CPSF4_HUMAN\|O95639_S200 | 1.400483759 | 1.90E-03 | 4.22E-02 |
| PLEC_HUMAN\|Q15149_S4673 | 1.40043187 | 3.05E-03 | 2.79E-02 |
| ITBP1_HUMAN\|O14713_S10_S14 | 1.400367836 | 2.20E-03 | 2.22E-02 |
| FA83B_HUMAN\|Q5T0W9_S869 | 1.400352205 | 3.70E-05 | 3.51E-03 |

| | | | |
|---|---|---|---|
| BCL10_HUMAN\|O95999_S138 | 1.400262997 | 4.50E-03 | 4.06E-02 |
| TRAF7_HUMAN\|Q6Q0C0_S108 | 1.400211568 | 1.02E-02 | 4.10E-02 |
| NUFP2_HUMAN\|Q7Z417_S212_S214 | 1.40018967 | 4.11E-03 | 3.55E-02 |
| MYO1E_HUMAN\|Q12965_S980 | 1.400088061 | 6.13E-03 | 4.66E-02 |
| M3K3_HUMAN\|Q99759_S337_S340 | 1.400066354 | 1.19E-03 | 1.12E-02 |
| AAK1_HUMAN\|Q2M2I8_S678_T674 | 1.399989537 | 1.21E-03 | 1.65E-02 |
| K1C16_HUMAN\|P08779_S446_S449 | 1.399986125 | 6.72E-03 | 4.99E-02 |
| P66B_HUMAN\|Q8WXI9_S334 | 1.399943354 | 2.31E-04 | 1.14E-02 |
| LAD1_HUMAN\|O00515_S38 | 1.399717301 | 3.29E-03 | 3.05E-02 |
| K1C16_HUMAN\|P08779_S450 | 1.399388302 | 3.91E-03 | 3.42E-02 |
| AAPK1_HUMAN\|Q13131_S508 | 1.39922276 | 1.03E-04 | 2.89E-03 |
| LARP7_HUMAN\|Q4G0J3_S298_S299 | 1.399119388 | 4.09E-04 | 4.85E-03 |
| KLC2_HUMAN\|Q9H0B6_S608_S611 | 1.399024582 | 4.20E-04 | 4.93E-03 |
| TB182_HUMAN\|Q9C0C2_S920 | 1.398945004 | 5.51E-03 | 3.48E-02 |
| HIRP3_HUMAN\|Q9BW71_S196_S199 | 1.398873209 | 1.62E-03 | 1.75E-02 |
| TANC2_HUMAN\|Q9HCD6_S1827 | 1.398827463 | 7.35E-04 | 1.26E-02 |
| RL37_HUMAN\|P61927_S96 | 1.3987772 | 4.67E-03 | 2.96E-02 |
| SMRCD_HUMAN\|Q9H4L7_S96 | 1.39873808 | 3.17E-03 | 2.97E-02 |
| ARG28_HUMAN\|Q8N1W1_S513 | 1.398700781 | 3.31E-03 | 2.97E-02 |
| RAE1_HUMAN\|P24386_S651 | 1.39861543 | 8.87E-04 | 1.92E-02 |
| MAP7_HUMAN\|Q14244_S202 | 1.398518207 | 1.00E-03 | 1.45E-02 |
| FOXO3_HUMAN\|O43524_S425 | 1.398504683 | 1.25E-03 | 1.17E-02 |
| CND1_HUMAN\|Q15021_S1333_T1339 | 1.398303465 | 1.16E-02 | 3.88E-02 |
| MON1A_HUMAN\|Q86VX9_S56 | 1.398250752 | 1.05E-03 | 8.28E-03 |
| SRSF6_HUMAN\|Q13247_S303 | 1.398209493 | 7.20E-03 | 4.22E-02 |
| OSB10_HUMAN\|Q9BXB5_S26_S32 | 1.398178601 | 4.75E-04 | 4.51E-03 |
| PALLD_HUMAN\|Q8WX93_S1352 | 1.398026056 | 1.01E-03 | 1.45E-02 |
| PEAK1_HUMAN\|Q9H792_T574 | 1.397968862 | 1.09E-03 | 6.54E-03 |
| FRM4B_HUMAN\|Q9Y2L6_S628 | 1.397412679 | 5.74E-03 | 2.95E-02 |
| MMTA2_HUMAN\|Q9BU76_S216_S220 | 1.397364684 | 3.57E-03 | 4.87E-02 |
| MYO9B_HUMAN\|Q13459_T1346 | 1.397302681 | 4.29E-03 | 4.40E-02 |

| | | | |
|---|---|---|---|
| CTNA1_HUMAN\|P35221_S641_S652 | 1.397300113 | 4.27E-03 | 2.40E-02 |
| SRRM2_HUMAN\|Q9UQ35_S778_S780 | 1.397281439 | 2.41E-03 | 1.79E-02 |
| PLEC_HUMAN\|Q15149_S4389_S4390 | 1.397169576 | 1.30E-02 | 4.99E-02 |
| TRI16_HUMAN\|O95361_S62 | 1.39715356 | 4.75E-03 | 2.99E-02 |
| DEN4C_HUMAN\|Q5VZ89_S1046 | 1.396955038 | 7.97E-04 | 1.25E-02 |
| BICD2_HUMAN\|Q8TD16_S615 | 1.396771642 | 1.78E-03 | 2.34E-02 |
| LYAR_HUMAN\|Q9NX58_S276 | 1.39655653 | 6.56E-04 | 3.92E-03 |
| LYAR_HUMAN\|Q9NX58_S276 | 1.396324 | 6.60E-04 | 1.13E-02 |
| LMNA_HUMAN\|P02545_S18 | 1.396274314 | 1.44E-03 | 1.60E-02 |
| RU17_HUMAN\|P08621_S226 | 1.396243508 | 4.06E-03 | 2.30E-02 |
| EGFR_HUMAN\|P00533_S991 | 1.396102462 | 3.56E-03 | 3.67E-02 |
| TMF1_HUMAN\|P82094_S338_S344 | 1.39602796 | 9.79E-04 | 9.31E-03 |
| PLXA2_HUMAN\|O75051_S1633 | 1.395969205 | 7.23E-03 | 3.51E-02 |
| NIBL1_HUMAN\|Q96TA1_S665 | 1.395732858 | 8.91E-04 | 1.44E-02 |
| ARI1A_HUMAN\|O14497_S696_S698_S702 | 1.395707259 | 5.01E-04 | 9.73E-03 |
| NFKB1_HUMAN\|P19838_S907 | 1.395655189 | 1.60E-04 | 4.02E-03 |
| ZFP91_HUMAN\|Q96JP5_S82 | 1.395613259 | 3.68E-03 | 3.29E-02 |
| ARHG1_HUMAN\|Q92888_S786 | 1.395605481 | 1.83E-03 | 4.16E-02 |
| PKHA2_HUMAN\|Q9HB19_S184 | 1.395555296 | 5.08E-04 | 7.78E-03 |
| STAT3_HUMAN\|P40763_S727 | 1.395431662 | 7.42E-04 | 1.27E-02 |
| DKC1_HUMAN\|O60832_S485_S494 | 1.395156902 | 7.27E-04 | 6.23E-03 |
| M4K4_HUMAN\|O95819_S842 | 1.395156219 | 1.60E-03 | 1.73E-02 |
| NFIC_HUMAN\|P08651_S284_S304_T293 | 1.395136034 | 7.18E-03 | 2.63E-02 |
| NOG1_HUMAN\|Q9BZE4_S470_Y466 | 1.39490783 | 2.09E-03 | 2.44E-02 |
| LIPB1_HUMAN\|Q86W92_S582 | 1.394831981 | 5.86E-04 | 5.38E-03 |
| PRC2A_HUMAN\|P48634_S1219 | 1.394794911 | 1.45E-03 | 2.05E-02 |
| TB182_HUMAN\|Q9C0C2_S872 | 1.394665714 | 1.72E-03 | 1.87E-02 |
| REPS1_HUMAN\|Q96D71_S162_S170_S174 | 1.394638481 | 2.37E-03 | 2.63E-02 |
| ZFP91_HUMAN\|Q96JP5_S83 | 1.39462813 | 6.21E-04 | 2.17E-02 |
| KIF3C_HUMAN\|O14782_T173 | 1.394512795 | 1.16E-02 | 3.88E-02 |
| NIPA_HUMAN\|Q86WB0_S321 | 1.39448762 | 3.26E-04 | 2.28E-03 |

| | | | |
|---|---|---|---|
| KI21A_HUMAN\|Q7Z4S6_S855 | 1.394352007 | 3.00E-03 | 2.76E-02 |
| MA7D1_HUMAN\|Q3KQU3_S452 | 1.394298329 | 7.77E-03 | 4.28E-02 |
| PLEC_HUMAN\|Q15149_S4673_S4675 | 1.3941316 | 9.07E-03 | 4.20E-02 |
| SRRM2_HUMAN\|Q9UQ35_S2132 | 1.393708107 | 6.04E-04 | 5.51E-03 |
| RL22L_HUMAN\|Q6P5R6_S112_S118 | 1.39366959 | 8.76E-03 | 4.09E-02 |
| PLEC_HUMAN\|Q15149_S4626_S4630 | 1.393611124 | 8.41E-04 | 5.29E-03 |
| XPO4_HUMAN\|Q9C0E2_S521 | 1.393531164 | 3.16E-03 | 3.47E-02 |
| PDLI2_HUMAN\|Q96JY6_S123_S124 | 1.393507736 | 1.37E-03 | 1.86E-02 |
| MISP_HUMAN\|Q8IVT2_S543 | 1.393454508 | 3.64E-03 | 3.81E-02 |
| M3K3_HUMAN\|Q99759_S337_S340 | 1.393336207 | 1.37E-03 | 2.00E-02 |
| FA83G_HUMAN\|A6ND36_S726 | 1.393245995 | 8.08E-03 | 3.86E-02 |
| KLC1_HUMAN\|Q07866-4_S561 | 1.393237335 | 1.39E-03 | 1.63E-02 |
| MAP1B_HUMAN\|P46821_S1797_Y1796 | 1.393227622 | 1.11E-02 | 4.39E-02 |
| TCF20_HUMAN\|Q9UGU0_S538 | 1.39318872 | 3.86E-03 | 2.69E-02 |
| RABE1_HUMAN\|Q15276_S407 | 1.392952794 | 1.70E-03 | 2.15E-02 |
| AAK1_HUMAN\|Q2M2I8_S678 | 1.392819094 | 3.15E-03 | 1.38E-02 |
| PRAG1_HUMAN\|Q86YV5_Y413 | 1.392689682 | 1.59E-03 | 1.15E-02 |
| AAK1_HUMAN\|Q2M2I8_S21 | 1.392671756 | 6.63E-03 | 4.85E-02 |
| NFIC_HUMAN\|P08651_S333_S339 | 1.392350279 | 4.56E-03 | 3.81E-02 |
| ZO1_HUMAN\|Q07157_S1617 | 1.392256958 | 1.89E-03 | 9.17E-03 |
| HNRPC_HUMAN\|P07910_S240 | 1.39216392 | 4.63E-03 | 4.13E-02 |
| DESP_HUMAN\|P15924_S2815 | 1.39216024 | 1.49E-03 | 1.32E-02 |
| WDR13_HUMAN\|Q9H1Z4_S70_S74 | 1.392119474 | 3.47E-03 | 3.07E-02 |
| F118A_HUMAN\|Q9NWS6_S349 | 1.391976005 | 5.82E-04 | 3.55E-03 |
| BCLF1_HUMAN\|Q9NYF8_S222 | 1.39196511 | 6.28E-04 | 5.67E-03 |
| UBP24_HUMAN\|Q9UPU5_S1141 | 1.391948233 | 3.52E-03 | 2.08E-02 |
| MAP1B_HUMAN\|P46821_S1785 | 1.391718589 | 2.68E-04 | 2.91E-03 |
| T2FA_HUMAN\|P35269_S433 | 1.391618595 | 8.83E-04 | 1.33E-02 |
| AHNK_HUMAN\|Q09666_S210_S216_S220 | 1.391470385 | 1.66E-03 | 2.13E-02 |
| KC1D_HUMAN\|P48730_S383 | 1.391446289 | 5.19E-03 | 4.17E-02 |
| K1C19_HUMAN\|P08727_S14 | 1.391365571 | 2.73E-03 | 2.58E-02 |

| | | | |
|---|---|---|---|
| PKP3_HUMAN\|Q9Y446_Y176 | 1.391345011 | 5.36E-03 | 4.16E-02 |
| EFHD2_HUMAN\|Q96C19_S74 | 1.391333169 | 3.90E-03 | 2.24E-02 |
| SRRM1_HUMAN\|Q8IYB3_S738_S740_S743 | 1.391208528 | 2.06E-03 | 1.41E-02 |
| TB182_HUMAN\|Q9C0C2_S1029 | 1.390876552 | 1.60E-03 | 1.33E-02 |
| RMP_HUMAN\|O94763_S372 | 1.390856584 | 7.38E-03 | 4.12E-02 |
| PACN2_HUMAN\|Q9UNF0_S399_S425 | 1.390669428 | 2.46E-03 | 1.60E-02 |
| ARMX3_HUMAN\|Q9UH62_S110 | 1.390653473 | 2.62E-03 | 3.07E-02 |
| COBL1_HUMAN\|Q53SF7_S364 | 1.390618399 | 2.09E-03 | 2.44E-02 |
| SMC1A_HUMAN\|Q14683_S957 | 1.390613275 | 1.14E-03 | 3.17E-02 |
| ARAF_HUMAN\|P10398_S257_T253 | 1.390590537 | 5.32E-03 | 4.23E-02 |
| NCOA7_HUMAN\|Q8NI08_S208_T210 | 1.390534609 | 2.26E-03 | 2.78E-02 |
| H2AX_HUMAN\|P16104_S140 | 1.390373707 | 9.45E-04 | 7.60E-03 |
| AMOL2_HUMAN\|Q9Y2J4_S540 | 1.390012697 | 2.84E-04 | 3.05E-03 |
| P66A_HUMAN\|Q86YP4_S100_S107 | 1.389993797 | 8.82E-04 | 1.43E-02 |
| NF1_HUMAN\|P21359_S2515 | 1.389789215 | 1.47E-03 | 1.09E-02 |
| UBP32_HUMAN\|Q8NFA0_S1393 | 1.389740369 | 1.73E-03 | 2.33E-02 |
| STK10_HUMAN\|O94804_S448_S455 | 1.389733663 | 1.68E-03 | 1.84E-02 |
| PP1R7_HUMAN\|Q15435_S12 | 1.389632996 | 9.16E-03 | 4.82E-02 |
| PKHG3_HUMAN\|A1L390_S1081 | 1.389573814 | 4.55E-04 | 8.55E-03 |
| TB22B_HUMAN\|Q9NU19_S132 | 1.389452533 | 2.66E-03 | 1.21E-02 |
| PALLD_HUMAN\|Q8WX93_S1352 | 1.389091491 | 1.21E-03 | 1.77E-02 |
| MON1A_HUMAN\|Q86VX9_S56 | 1.388973302 | 1.27E-03 | 1.71E-02 |
| PEX1_HUMAN\|O43933_S1181 | 1.38891379 | 8.13E-03 | 3.41E-02 |
| KI21A_HUMAN\|Q7Z4S6_S1229 | 1.388825469 | 9.59E-04 | 5.87E-03 |
| EHBP1_HUMAN\|Q8NDI1_S428_S436 | 1.388808236 | 4.22E-04 | 8.14E-03 |
| COHA1_HUMAN\|Q9UMD9_S85_S93_S96 | 1.388645629 | 5.26E-03 | 4.20E-02 |
| FRYL_HUMAN\|O94915_T1959 | 1.388615506 | 2.78E-03 | 1.74E-02 |
| KI21A_HUMAN\|Q7Z4S6_S1229 | 1.388546483 | 9.65E-04 | 9.19E-03 |
| MTCL1_HUMAN\|Q9Y4B5_S776 | 1.388495132 | 8.24E-04 | 6.87E-03 |
| BCL9L_HUMAN\|Q86UU0_S21_S25 | 1.388433322 | 1.06E-03 | 2.92E-02 |
| RRP1B_HUMAN\|Q14684_S245 | 1.388041712 | 8.32E-04 | 5.26E-03 |

| | | | |
|---|---|---|---|
| SPB1_HUMAN\|Q8IY81_S333_S335 | 1.387987828 | 1.33E-03 | 6.91E-03 |
| RNF31_HUMAN\|Q96EP0_S466 | 1.387795485 | 8.64E-04 | 1.37E-02 |
| NELFE_HUMAN\|P18615_S181 | 1.387522882 | 5.60E-04 | 1.00E-02 |
| ACSA_HUMAN\|Q9NR19_S267 | 1.38750667 | 4.39E-03 | 2.05E-02 |
| IF4B_HUMAN\|P23588_T500 | 1.387488634 | 6.02E-03 | 3.54E-02 |
| VANG1_HUMAN\|Q8TAA9_S520_S523 | 1.387396963 | 4.45E-03 | 4.36E-02 |
| ARHGG_HUMAN\|Q5VV41_S227_T226 | 1.38733579 | 1.57E-03 | 1.31E-02 |
| EPN1_HUMAN\|Q9Y6I3_S454 | 1.386971722 | 1.04E-03 | 1.60E-02 |
| LARP1_HUMAN\|Q6PKG0_S517_S521 | 1.386753392 | 4.95E-04 | 4.68E-03 |
| ELF1_HUMAN\|P32519_S168 | 1.386620445 | 6.25E-04 | 1.09E-02 |
| SQSTM_HUMAN\|Q13501_S266 | 1.386468384 | 1.29E-03 | 9.70E-03 |
| PLCH2_HUMAN\|O75038_S595 | 1.386439691 | 2.10E-03 | 4.44E-02 |
| PUM1_HUMAN\|Q14671_S709 | 1.386400663 | 6.57E-04 | 9.45E-03 |
| DTD1_HUMAN\|Q8TEA8_S196_S197 | 1.386381249 | 1.03E-03 | 1.58E-02 |
| K1C17_HUMAN\|Q04695_S13 | 1.386354023 | 4.98E-03 | 4.36E-02 |
| HSPB1_HUMAN\|P04792_S82 | 1.38635192 | 5.78E-04 | 3.91E-03 |
| PRC2A_HUMAN\|P48634_S1004 | 1.386301655 | 4.19E-03 | 3.58E-02 |
| TLE3_HUMAN\|Q04726_S286 | 1.386255517 | 2.80E-03 | 1.75E-02 |
| CCD86_HUMAN\|Q9H6F5_S217 | 1.386200182 | 1.25E-02 | 4.82E-02 |
| LAD1_HUMAN\|O00515_S38 | 1.386146309 | 4.16E-03 | 2.72E-02 |
| SARG_HUMAN\|Q9BW04_S502 | 1.38611606 | 9.31E-04 | 7.50E-03 |
| SRRM1_HUMAN\|Q8IYB3_S754_S756 | 1.385885709 | 3.12E-03 | 3.17E-02 |
| AAPK1_HUMAN\|Q13131_S508 | 1.385789154 | 1.46E-04 | 1.21E-03 |
| SRSF1_HUMAN\|Q07955_S234_S238 | 1.385770955 | 1.60E-03 | 8.04E-03 |
| ARI1A_HUMAN\|O14497_S696_S698 | 1.385768183 | 1.21E-03 | 1.73E-02 |
| RAI1_HUMAN\|Q7Z5J4_S1108 | 1.385611544 | 1.81E-03 | 1.26E-02 |
| AF1L2_HUMAN\|Q8N4X5_S651 | 1.38549993 | 4.29E-04 | 8.85E-03 |
| PRC2A_HUMAN\|P48634_S1085_S1089 | 1.38519478 | 3.38E-03 | 1.65E-02 |
| EGFR_HUMAN\|P00533_S991 | 1.385142665 | 4.32E-03 | 2.94E-02 |
| TBA1C_HUMAN\|Q9BQE3_Y449 | 1.385130912 | 6.52E-03 | 3.76E-02 |
| PUM1_HUMAN\|Q14671_S709 | 1.385128838 | 6.76E-04 | 9.05E-03 |

| | | | |
|---|---|---|---|
| ZO1_HUMAN|Q07157_S1617 | 1.385092094 | 2.17E-03 | 2.52E-02 |
| DDX20_HUMAN|Q9UHI6_S677_Y673 | 1.385053787 | 3.24E-03 | 3.53E-02 |
| FA83H_HUMAN|Q6ZRV2_S1003 | 1.384931729 | 2.85E-04 | 3.71E-03 |
| LIPA1_HUMAN|Q13136_S1172 | 1.384912645 | 1.18E-03 | 1.45E-02 |
| LIPA3_HUMAN|O75145_S1164 | 1.38466092 | 2.92E-03 | 3.30E-02 |
| SRPK2_HUMAN|P78362_S496_T498 | 1.384658658 | 4.24E-03 | 4.22E-02 |
| KC1D_HUMAN|P48730_S383 | 1.384097471 | 5.85E-03 | 4.45E-02 |
| K2C5_HUMAN|P13647_S11_S14 | 1.384042608 | 1.45E-03 | 1.86E-02 |
| K1C16_HUMAN|P08779_S448 | 1.383769744 | 8.31E-03 | 3.94E-02 |
| ZC3HE_HUMAN|Q6PJT7_S390 | 1.383683905 | 6.90E-03 | 2.54E-02 |
| VIME_HUMAN|P08670_S430 | 1.383555419 | 3.74E-03 | 3.31E-02 |
| AHNK_HUMAN|Q09666_S3426 | 1.383506284 | 2.48E-03 | 1.83E-02 |
| RTKN_HUMAN|Q9BST9_S561 | 1.38337848 | 4.33E-03 | 1.79E-02 |
| KAP2_HUMAN|P13861_S78_S80 | 1.38315189 | 2.91E-03 | 1.30E-02 |
| FMNL2_HUMAN|Q96PY5_S1016 | 1.383071979 | 8.49E-03 | 4.73E-02 |
| PHIP_HUMAN|Q8WWQ0_S1783 | 1.38304892 | 1.27E-03 | 7.35E-03 |
| DEN1A_HUMAN|Q8TEH3_S536 | 1.382969618 | 3.91E-03 | 3.88E-02 |
| FA83H_HUMAN|Q6ZRV2_S1098 | 1.382881017 | 2.60E-03 | 2.80E-02 |
| H2AX_HUMAN|P16104_S140 | 1.382829043 | 1.11E-03 | 1.31E-02 |
| TBC9B_HUMAN|Q66K14_S1241 | 1.382722568 | 4.64E-04 | 9.15E-03 |
| PDLI2_HUMAN|Q96JY6_S137_S143 | 1.382708014 | 3.77E-03 | 3.32E-02 |
| SSH1_HUMAN|Q8WYL5_S937 | 1.382667236 | 1.44E-03 | 7.40E-03 |
| PEAK1_HUMAN|Q9H792_T574 | 1.382369535 | 1.51E-03 | 1.34E-02 |
| TISB_HUMAN|Q07352_S54 | 1.382361996 | 6.48E-04 | 2.27E-02 |
| MYPT1_HUMAN|O14974_S507 | 1.382290297 | 4.62E-03 | 3.74E-02 |
| MAP1B_HUMAN|P46821_S1501 | 1.382285267 | 9.11E-04 | 2.73E-02 |
| CLIP4_HUMAN|Q8N3C7_T582 | 1.382257143 | 3.83E-03 | 3.93E-02 |
| CRBG1_HUMAN|Q9Y4K1_S348 | 1.382183871 | 3.49E-03 | 2.07E-02 |
| SLU7_HUMAN|O95391_S215 | 1.382182967 | 3.93E-03 | 2.24E-02 |
| FA83H_HUMAN|Q6ZRV2_S1025 | 1.382023225 | 4.07E-03 | 3.80E-02 |
| CHRC1_HUMAN|Q9NRG0_S124 | 1.381988778 | 2.54E-03 | 1.86E-02 |

| | | | |
|---|---|---|---|
| RRP1B_HUMAN\|Q14684_S706 | 1.381965346 | 7.88E-03 | 4.49E-02 |
| EXPH5_HUMAN\|Q8NEV8_S341 | 1.381901482 | 3.57E-03 | 2.10E-02 |
| PRC2A_HUMAN\|P48634_S1089_S1092 | 1.381895566 | 2.72E-03 | 1.39E-02 |
| PPR18_HUMAN\|Q6NYC8_S490 | 1.381848398 | 7.92E-04 | 8.19E-03 |
| K2C5_HUMAN\|P13647_Y66 | 1.381655585 | 1.14E-03 | 3.17E-02 |
| NFIC_HUMAN\|P08651_S333_S339 | 1.381545745 | 5.46E-03 | 4.20E-02 |
| PCM1_HUMAN\|Q15154_S93 | 1.381526908 | 6.07E-04 | 4.07E-03 |
| CT2NL_HUMAN\|Q9P2B4_S527 | 1.381526039 | 1.54E-03 | 2.02E-02 |
| NFIC_HUMAN\|P08651_S284_S304_T293 | 1.381519259 | 8.84E-03 | 4.74E-02 |
| AFAP1_HUMAN\|Q8N556_S548 | 1.381435248 | 4.48E-04 | 7.12E-03 |
| BORG2_HUMAN\|Q9UKI2_T90 | 1.380948639 | 4.84E-03 | 3.86E-02 |
| IRS2_HUMAN\|Q9Y4H2_S679 | 1.380943069 | 3.35E-03 | 3.00E-02 |
| RNF25_HUMAN\|Q96BH1_S450 | 1.380900421 | 8.98E-03 | 4.76E-02 |
| MA7D1_HUMAN\|Q3KQU3_S113_S125 | 1.380881751 | 2.26E-03 | 1.79E-02 |
| NIPA_HUMAN\|Q86WB0_S407_S410_S411 | 1.380879842 | 6.70E-04 | 1.18E-02 |
| SRRM2_HUMAN\|Q9UQ35_S871_S876 | 1.38084976 | 4.91E-03 | 2.65E-02 |
| IF4B_HUMAN\|P23588_S497_S498 | 1.380289745 | 8.52E-03 | 4.75E-02 |
| NIPA_HUMAN\|Q86WB0_S407_S410_S411 | 1.380015412 | 6.83E-04 | 1.20E-02 |
| K1C14_HUMAN\|P02533_S53 | 1.379889769 | 2.40E-03 | 2.42E-02 |
| IF2A_HUMAN\|P05198_T279 | 1.3797122 | 9.86E-03 | 4.49E-02 |
| PKHG3_HUMAN\|A1L390_S638_S643 | 1.379712039 | 2.76E-04 | 3.61E-03 |
| PP4R1_HUMAN\|Q8TF05_S948_T945 | 1.37957529 | 4.82E-03 | 3.85E-02 |
| NDRG1_HUMAN\|Q92597_S336 | 1.379281726 | 3.14E-03 | 1.55E-02 |
| PRC2A_HUMAN\|P48634_S1004 | 1.379165365 | 4.72E-03 | 2.58E-02 |
| HMCS1_HUMAN\|Q01581_S516 | 1.379080483 | 8.96E-03 | 4.76E-02 |
| F118A_HUMAN\|Q9NWS6_S349 | 1.379027871 | 7.80E-04 | 1.30E-02 |
| TCF20_HUMAN\|Q9UGU0_S1322 | 1.378979463 | 1.88E-04 | 4.89E-03 |
| CDK17_HUMAN\|Q00537_S137 | 1.378858648 | 2.77E-04 | 1.32E-02 |
| GGYF2_HUMAN\|Q6Y7W6_S201 | 1.378855917 | 4.52E-03 | 4.06E-02 |
| YBOX3_HUMAN\|P16989_S369_S370 | 1.378833707 | 5.54E-03 | 2.89E-02 |
| CSN3_HUMAN\|Q9UNS2_S421 | 1.378746421 | 2.84E-04 | 5.20E-03 |

| | | | |
|---|---|---|---|
| ARI1A_HUMAN\|O14497_S696_S698_S702 | 1.378375407 | 7.46E-04 | 9.72E-03 |
| PRC2A_HUMAN\|P48634_S1004 | 1.378203963 | 4.80E-03 | 3.84E-02 |
| ZC3HD_HUMAN\|Q5T200_S265 | 1.378067078 | 5.97E-04 | 6.42E-03 |
| PLCH2_HUMAN\|O75038_S562 | 1.378024103 | 3.55E-04 | 6.00E-03 |
| OXR1_HUMAN\|Q8N573_S204_T203 | 1.377970683 | 1.95E-03 | 2.53E-02 |
| KLC3_HUMAN\|Q6P597_S497_S502 | 1.377824018 | 3.78E-04 | 4.18E-02 |
| IF2P_HUMAN\|O60841_S182_S183 | 1.377776208 | 1.55E-03 | 1.30E-02 |
| CTNA1_HUMAN\|P35221_S641_S652 | 1.377651689 | 5.88E-03 | 4.87E-02 |
| PDLI2_HUMAN\|Q96JY6_S117 | 1.377468724 | 9.79E-03 | 3.96E-02 |
| IF2P_HUMAN\|O60841_S182_S186_S190 | 1.377319778 | 4.52E-03 | 2.10E-02 |
| NASP_HUMAN\|P49321_S421 | 1.377269191 | 1.06E-03 | 1.26E-02 |
| LIMD1_HUMAN\|Q9UGP4_S277 | 1.377041311 | 2.11E-03 | 2.65E-02 |
| VIME_HUMAN\|P08670_S459 | 1.376874109 | 2.05E-03 | 2.10E-02 |
| T2FA_HUMAN\|P35269_S433 | 1.376833365 | 1.20E-03 | 2.37E-02 |
| DC1L2_HUMAN\|O43237_S383 | 1.376806915 | 4.29E-03 | 2.40E-02 |
| MACF1_HUMAN\|Q9UPN3_S4 | 1.37676524 | 3.08E-04 | 2.29E-03 |
| SRRM2_HUMAN\|Q9UQ35_S2694 | 1.376746966 | 2.12E-04 | 2.89E-03 |
| NFIB_HUMAN\|O00712_S295 | 1.376725057 | 1.25E-03 | 1.11E-02 |
| LMNA_HUMAN\|P02545_S428 | 1.37665927 | 1.21E-03 | 1.41E-02 |
| COMT_HUMAN\|P21964_S267 | 1.376630195 | 3.12E-03 | 3.17E-02 |
| RPGF6_HUMAN\|Q8TEU7_S1590 | 1.3765174 | 6.72E-03 | 4.90E-02 |
| MA7D1_HUMAN\|Q3KQU3_S70 | 1.37644528 | 3.33E-03 | 3.08E-02 |
| NCOA7_HUMAN\|Q8NI08_S208 | 1.376385167 | 2.86E-03 | 4.30E-02 |
| MAP4_HUMAN\|P27816_S1073 | 1.376331968 | 4.15E-03 | 3.56E-02 |
| SHRM2_HUMAN\|Q13796_S764 | 1.376317996 | 8.87E-03 | 4.12E-02 |
| APBB2_HUMAN\|Q92870_S409_S412 | 1.375986187 | 2.60E-03 | 1.66E-02 |
| NDRG2_HUMAN\|Q9UN36_S332 | 1.375951243 | 2.21E-03 | 1.48E-02 |
| RHG01_HUMAN\|Q07960_S51 | 1.375893621 | 4.05E-03 | 2.66E-02 |
| PRAG1_HUMAN\|Q86YV5_S148 | 1.375877793 | 3.39E-03 | 3.55E-02 |
| DDX50_HUMAN\|Q9BQ39_S683 | 1.375811185 | 1.77E-03 | 1.41E-02 |
| GYS1_HUMAN\|P13807_S727 | 1.375653289 | 1.94E-03 | 1.60E-02 |

| | | | |
|---|---|---|---|
| STMN1_HUMAN\|P16949_S16 | 1.375595159 | 6.64E-03 | 4.95E-02 |
| HNRPC_HUMAN\|P07910_S240 | 1.375511934 | 6.06E-03 | 3.08E-02 |
| PP1RA_HUMAN\|Q96QC0_S313 | 1.375446919 | 3.60E-03 | 1.55E-02 |
| AHNK_HUMAN\|Q09666_S5763 | 1.375229774 | 4.39E-04 | 8.74E-03 |
| HNRH3_HUMAN\|P31942_S298 | 1.375156544 | 4.10E-03 | 3.55E-02 |
| TB182_HUMAN\|Q9C0C2_S1029 | 1.375126526 | 2.19E-03 | 2.70E-02 |
| IASPP_HUMAN\|Q8WUF5_S113 | 1.374784686 | 2.54E-03 | 2.46E-02 |
| TENS4_HUMAN\|Q8IZW8_S253 | 1.37476806 | 1.38E-03 | 1.55E-02 |
| PAIRB_HUMAN\|Q8NC51_S392 | 1.374659508 | 8.33E-03 | 4.66E-02 |
| NFIX_HUMAN\|Q14938_S280 | 1.374600245 | 1.63E-03 | 8.16E-03 |
| BCL9_HUMAN\|O00512_S687 | 1.374510717 | 1.41E-03 | 2.02E-02 |
| MYPT1_HUMAN\|O14974_S507 | 1.374392997 | 5.26E-03 | 4.87E-02 |
| ACD_HUMAN\|Q96AP0_S425 | 1.374224038 | 2.43E-03 | 4.79E-02 |
| ZC3HD_HUMAN\|Q5T200_S64 | 1.374115207 | 6.89E-03 | 3.39E-02 |
| FA83H_HUMAN\|Q6ZRV2_S647 | 1.374068773 | 5.14E-04 | 4.84E-03 |
| MK14_HUMAN\|Q16539_Y182 | 1.37389398 | 7.86E-03 | 2.84E-02 |
| UBXN7_HUMAN\|O94888_S280 | 1.373858039 | 6.25E-03 | 2.35E-02 |
| MILK1_HUMAN\|Q8N3F8_S640_S644 | 1.373809591 | 4.31E-03 | 3.55E-02 |
| PAXX_HUMAN\|Q9BUH6_S148 | 1.373798714 | 1.93E-03 | 1.59E-02 |
| MSH6_HUMAN\|P52701_T139 | 1.373720441 | 3.41E-03 | 1.66E-02 |
| FRMD6_HUMAN\|Q96NE9_S375 | 1.373701048 | 6.57E-04 | 5.84E-03 |
| RRP8_HUMAN\|O43159_S104_S106 | 1.373694167 | 1.53E-03 | 3.85E-02 |
| GGYF2_HUMAN\|Q6Y7W6_S23 | 1.373563197 | 1.10E-03 | 5.93E-03 |
| CR025_HUMAN\|Q96B23_S145 | 1.37351976 | 1.04E-03 | 1.60E-02 |
| BAIP2_HUMAN\|Q9UQB8_S453 | 1.373503715 | 5.11E-03 | 2.74E-02 |
| MPLKI_HUMAN\|Q8TAP9_T110 | 1.373434296 | 1.04E-02 | 4.65E-02 |
| PML_HUMAN\|P29590_S504 | 1.373334693 | 3.09E-03 | 2.24E-02 |
| AHNK_HUMAN\|Q09666_S210 | 1.373266512 | 8.06E-04 | 1.03E-02 |
| K2C6B_HUMAN\|P04259_T552 | 1.373174008 | 1.13E-02 | 4.97E-02 |
| NEB2_HUMAN\|Q96SB3_S99 | 1.373167922 | 5.16E-03 | 4.47E-02 |
| PDLI4_HUMAN\|P50479_S116 | 1.37278175 | 8.47E-03 | 4.00E-02 |

| | | | |
|---|---|---|---|
| PKP3_HUMAN\|Q9Y446_S314 | 1.372752813 | 2.03E-03 | 4.38E-02 |
| DC1L2_HUMAN\|O43237_S194 | 1.372737202 | 1.74E-04 | 2.05E-03 |
| PHLB2_HUMAN\|Q86SQ0_Y567 | 1.372175126 | 1.88E-03 | 1.56E-02 |
| GIT1_HUMAN\|Q9Y2X7_S362 | 1.372135933 | 1.74E-03 | 2.18E-02 |
| TCF20_HUMAN\|Q9UGU0_S538 | 1.37182481 | 5.52E-03 | 3.31E-02 |
| AHNK_HUMAN\|Q09666_S2397 | 1.371804251 | 1.43E-02 | 4.61E-02 |
| ACAP2_HUMAN\|Q15057_S384_T382 | 1.371578891 | 5.24E-05 | 2.12E-03 |
| ACAP2_HUMAN\|Q15057_S384 | 1.371410871 | 5.60E-04 | 3.45E-03 |
| MDHC_HUMAN\|P40925_S241 | 1.371404041 | 4.33E-03 | 2.79E-02 |
| FCHO2_HUMAN\|Q0JRZ9_S403 | 1.371311147 | 6.14E-04 | 3.71E-03 |
| NDRG1_HUMAN\|Q92597_S364 | 1.371209696 | 3.51E-03 | 3.09E-02 |
| MILK1_HUMAN\|Q8N3F8_S578 | 1.37094362 | 1.23E-03 | 1.66E-02 |
| RHG12_HUMAN\|Q8IWW6_S213_S215 | 1.370923524 | 2.70E-03 | 1.70E-02 |
| DLG5_HUMAN\|Q8TDM6_S1264 | 1.370837859 | 4.69E-03 | 2.57E-02 |
| PRC2C_HUMAN\|Q9Y520_T2691 | 1.370785596 | 3.74E-03 | 3.58E-02 |
| NELFE_HUMAN\|P18615_S353 | 1.370734353 | 2.89E-04 | 5.24E-03 |
| RGL2_HUMAN\|O15211_S619 | 1.370732879 | 5.44E-03 | 3.27E-02 |
| MINT_HUMAN\|Q96T58_S623 | 1.370614715 | 2.50E-04 | 5.95E-03 |
| CASC3_HUMAN\|O15234_S148 | 1.37057741 | 6.64E-04 | 2.20E-02 |
| PCM1_HUMAN\|Q15154_S93 | 1.370568832 | 7.77E-04 | 1.29E-02 |
| ZYX_HUMAN\|Q15942_S259 | 1.37027703 | 2.65E-03 | 2.60E-02 |
| IF4G1_HUMAN\|Q04637_S1187 | 1.370158074 | 3.46E-03 | 3.07E-02 |
| AHNK2_HUMAN\|Q8IVF2_S4185 | 1.370042367 | 1.75E-03 | 2.34E-02 |
| SRRM1_HUMAN\|Q8IYB3_S874 | 1.37000746 | 1.04E-02 | 4.65E-02 |
| RBM7_HUMAN\|Q9Y580_S204 | 1.369992946 | 8.23E-03 | 3.44E-02 |
| SRRM1_HUMAN\|Q8IYB3_S754_S756 | 1.369910132 | 4.18E-03 | 2.73E-02 |
| K1468_HUMAN\|Q9P260_S180 | 1.36990674 | 2.31E-03 | 1.08E-02 |
| PRAG1_HUMAN\|Q86YV5_S712 | 1.369811483 | 1.25E-03 | 6.58E-03 |
| LIPB1_HUMAN\|Q86W92_S601 | 1.369797397 | 1.61E-03 | 3.79E-02 |
| BRAF_HUMAN\|P15056_S447 | 1.36979663 | 6.48E-03 | 2.42E-02 |
| K2C5_HUMAN\|P13647_S6 | 1.369673964 | 7.65E-03 | 4.22E-02 |

| | | | |
|---|---|---|---|
| ZN185_HUMAN\|O15231_T154_T158 | 1.369512185 | 1.72E-03 | 1.87E-02 |
| DEN1A_HUMAN\|Q8TEH3_S536 | 1.369491176 | 4.97E-03 | 3.24E-02 |
| NRK_HUMAN\|Q7Z2Y5_S854 | 1.36946438 | 5.25E-04 | 8.01E-03 |
| ATX2L_HUMAN\|Q8WWM7_S339 | 1.369390591 | 2.03E-04 | 3.71E-03 |
| 4EBP1_HUMAN\|Q13541_S44 | 1.368811559 | 5.58E-03 | 4.37E-02 |
| TRI47_HUMAN\|Q96LD4_S588 | 1.368806631 | 8.15E-03 | 4.60E-02 |
| NUCKS_HUMAN\|Q9H1E3_S54_S58_S61 | 1.368761455 | 2.12E-03 | 1.44E-02 |
| DDX50_HUMAN\|Q9BQ39_S683 | 1.368647398 | 2.04E-03 | 2.34E-02 |
| ZO2_HUMAN\|Q9UDY2_S170 | 1.368557515 | 1.84E-03 | 8.97E-03 |
| ELF1_HUMAN\|P32519_S163_S168 | 1.368354114 | 2.58E-03 | 1.65E-02 |
| ASPP2_HUMAN\|Q13625_S480 | 1.368323505 | 1.73E-03 | 2.11E-02 |
| HNRH3_HUMAN\|P31942_S298 | 1.368282293 | 4.62E-03 | 2.54E-02 |
| PLEC_HUMAN\|Q15149_S4622_Y4615 | 1.368103482 | 2.41E-03 | 1.79E-02 |
| PLEC_HUMAN\|Q15149_S4670 | 1.367974447 | 3.77E-03 | 1.80E-02 |
| PDS5A_HUMAN\|Q29RF7_S1276 | 1.367713649 | 1.54E-02 | 4.88E-02 |
| NCK5L_HUMAN\|Q9HCH0_S489_S494 | 1.3676831 | 4.18E-03 | 3.49E-02 |
| IF4B_HUMAN\|P23588_S489_S497_S498 | 1.367675351 | 3.29E-03 | 3.05E-02 |
| PLEC_HUMAN\|Q15149_S4626 | 1.367316369 | 2.02E-03 | 1.08E-02 |
| PARP4_HUMAN\|Q9UKK3_S1722 | 1.367277084 | 4.80E-03 | 3.18E-02 |
| P63_HUMAN\|Q9H3D4_S463 | 1.367146811 | 1.36E-03 | 1.79E-02 |
| NKAP_HUMAN\|Q8N5F7_S149 | 1.367099363 | 1.98E-03 | 2.08E-02 |
| NFIB_HUMAN\|O00712_S295 | 1.367051426 | 1.55E-03 | 3.71E-02 |
| AKTS1_HUMAN\|Q96B36_S202_S203 | 1.367012761 | 1.40E-03 | 1.21E-02 |
| CD2AP_HUMAN\|Q9Y5K6_S458 | 1.366976419 | 6.96E-03 | 3.42E-02 |
| LMNA_HUMAN\|P02545_S428 | 1.366951014 | 1.50E-03 | 1.69E-02 |
| SZRD1_HUMAN\|Q7Z422_S107 | 1.366794155 | 2.49E-03 | 2.50E-02 |
| Q659E9_HUMAN\|Q659E9_S168 | 1.366652639 | 1.10E-03 | 1.63E-02 |
| TPBG_HUMAN\|Q13641_S415_S418 | 1.366429616 | 2.72E-03 | 1.23E-02 |
| SCAF8_HUMAN\|Q9UPN6_S617 | 1.366388973 | 5.46E-03 | 4.20E-02 |
| DCA10_HUMAN\|Q5QP82_S350 | 1.366230107 | 7.81E-03 | 2.83E-02 |
| SOS1_HUMAN\|Q07889_S1078_S1082 | 1.366205649 | 1.05E-02 | 4.20E-02 |

| | | | |
|---|---|---|---|
| ZBT21_HUMAN\|Q9ULJ3_S435_T431 | 1.36617639 | 2.41E-03 | 2.42E-02 |
| SMCA4_HUMAN\|P51532_S1570_S1575_S1586 | 1.366140742 | 8.26E-03 | 3.93E-02 |
| LUZP1_HUMAN\|Q86V48_S574 | 1.366050774 | 3.26E-03 | 3.45E-02 |
| NFIC_HUMAN\|P08651_S284_S304_T293 | 1.366020242 | 1.12E-02 | 4.43E-02 |
| STMN1_HUMAN\|P16949_S25 | 1.365985616 | 2.39E-03 | 1.24E-02 |
| SLK_HUMAN\|Q9H2G2_S372 | 1.365888182 | 2.61E-04 | 2.85E-03 |
| IF4B_HUMAN\|P23588_S495_S497 | 1.36586907 | 2.04E-03 | 2.58E-02 |
| TB182_HUMAN\|Q9C0C2_S836 | 1.365844107 | 2.72E-03 | 3.10E-02 |
| ARI1A_HUMAN\|O14497_S381 | 1.365782646 | 1.66E-03 | 1.42E-02 |
| CSN1_HUMAN\|Q13098_S483_T479 | 1.365766281 | 3.09E-03 | 2.18E-02 |
| NAV1_HUMAN\|Q8NEY1_S1000 | 1.365746448 | 2.29E-03 | 1.21E-02 |
| PCSK9_HUMAN\|Q8NBP7_S688 | 1.365708323 | 1.18E-02 | 3.91E-02 |
| CHRC1_HUMAN\|Q9NRG0_S124 | 1.365303605 | 3.50E-03 | 3.44E-02 |
| DCTN1_HUMAN\|Q14203_S105 | 1.365149238 | 2.97E-03 | 2.11E-02 |
| IASPP_HUMAN\|Q8WUF5_S567 | 1.365148708 | 5.89E-04 | 1.09E-02 |
| CDK16_HUMAN\|Q00536_S95 | 1.365110041 | 4.95E-03 | 2.67E-02 |
| PLEC_HUMAN\|Q15149_S4626 | 1.364956996 | 2.12E-03 | 1.63E-02 |
| NU107_HUMAN\|P57740_S37 | 1.364956156 | 5.19E-03 | 2.35E-02 |
| C170B_HUMAN\|Q9Y4F5_S1548 | 1.364894994 | 6.89E-04 | 9.78E-03 |
| AFAD_HUMAN\|P55196_S1696 | 1.364760745 | 5.68E-03 | 4.43E-02 |
| SMRCD_HUMAN\|Q9H4L7_S96 | 1.364592918 | 5.81E-03 | 4.43E-02 |
| P63_HUMAN\|Q9H3D4_S463 | 1.364581668 | 1.44E-03 | 3.73E-02 |
| K1C16_HUMAN\|P08779_S447_S450 | 1.364579791 | 9.56E-03 | 4.38E-02 |
| LUZP1_HUMAN\|Q86V48_S575 | 1.364540356 | 8.29E-04 | 1.05E-02 |
| ML12A_HUMAN\|P19105_S19 | 1.364495906 | 3.57E-03 | 3.13E-02 |
| PAIRB_HUMAN\|Q8NC51_S392 | 1.364284163 | 9.77E-03 | 3.39E-02 |
| PCM1_HUMAN\|Q15154_S872 | 1.364056544 | 2.02E-03 | 3.46E-02 |
| VIME_HUMAN\|P08670_S66 | 1.363971564 | 3.16E-03 | 2.96E-02 |
| LMNA_HUMAN\|P02545_S403_S407 | 1.363883332 | 1.78E-03 | 2.15E-02 |
| MPDZ_HUMAN\|O75970_S1595 | 1.363740875 | 2.45E-03 | 1.59E-02 |
| SRRM2_HUMAN\|Q9UQ35_S2694 | 1.363682077 | 3.01E-04 | 6.34E-03 |

| | | | |
|---|---|---|---|
| URFB1_HUMAN\|Q6BDS2_S1106 | 1.363332612 | 3.43E-03 | 1.49E-02 |
| ACINU_HUMAN\|Q9UKV3_S838 | 1.363260639 | 1.81E-03 | 1.92E-02 |
| SLU7_HUMAN\|O95391_S215 | 1.363221797 | 5.44E-03 | 3.27E-02 |
| EFHD2_HUMAN\|Q96C19_S76 | 1.363112724 | 9.44E-03 | 4.34E-02 |
| LAR4B_HUMAN\|Q92615_S601 | 1.363069405 | 3.81E-03 | 2.19E-02 |
| SRGP3_HUMAN\|O43295_S858 | 1.362920543 | 7.14E-03 | 3.48E-02 |
| FA83H_HUMAN\|Q6ZRV2_S914 | 1.362911024 | 5.47E-03 | 2.86E-02 |
| MAP1B_HUMAN\|P46821_S1501 | 1.362820804 | 1.40E-03 | 1.81E-02 |
| SI1L3_HUMAN\|O60292_S172 | 1.362400579 | 1.64E-04 | 4.09E-03 |
| CCDC9_HUMAN\|Q9Y3X0_S376_S386_T381 | 1.362269629 | 5.18E-03 | 4.48E-02 |
| COBL1_HUMAN\|Q53SF7_S294 | 1.362148786 | 1.38E-03 | 1.62E-02 |
| NCOA7_HUMAN\|Q8NI08_S208 | 1.362083155 | 3.75E-03 | 2.17E-02 |
| NUP53_HUMAN\|Q8NFH5_T280 | 1.361953242 | 6.52E-04 | 9.41E-03 |
| MATR3_HUMAN\|P43243_S598 | 1.361923514 | 3.15E-04 | 3.29E-03 |
| K1468_HUMAN\|Q9P260_S180 | 1.36192039 | 2.71E-03 | 3.10E-02 |
| AHNK_HUMAN\|Q09666_S5749 | 1.361707632 | 5.80E-04 | 5.34E-03 |
| KMT2D_HUMAN\|O14686_S2274 | 1.361493526 | 5.99E-03 | 3.69E-02 |
| JUNB_HUMAN\|P17275_T255 | 1.361490548 | 6.97E-04 | 9.87E-03 |
| OTU7B_HUMAN\|Q6GQQ9_S467 | 1.361225401 | 1.19E-03 | 1.62E-02 |
| SPIR1_HUMAN\|Q08AE8_T509 | 1.361061846 | 5.31E-03 | 4.23E-02 |
| UN45A_HUMAN\|Q9H3U1_S147_Y145 | 1.361013399 | 5.88E-03 | 3.67E-02 |
| BCL9L_HUMAN\|Q86UU0_S118_S121 | 1.360980239 | 6.62E-04 | 2.20E-02 |
| PKP3_HUMAN\|Q9Y446_S313_S314 | 1.360935009 | 2.56E-03 | 1.94E-02 |
| SSH2_HUMAN\|Q76I76_S38 | 1.360864663 | 4.81E-03 | 4.77E-02 |
| KLC3_HUMAN\|Q6P597_S502 | 1.360371747 | 1.25E-04 | 3.70E-03 |
| MACF1_HUMAN\|Q9UPN3_S3 | 1.360357578 | 1.50E-03 | 7.66E-03 |
| MSH6_HUMAN\|P52701_T139 | 1.360346674 | 4.38E-03 | 4.24E-02 |
| TB22B_HUMAN\|Q9NU19_S132 | 1.360008344 | 4.54E-03 | 4.42E-02 |
| NDK8_HUMAN\|O60361_S105 | 1.35995721 | 5.15E-03 | 4.47E-02 |
| PDLI2_HUMAN\|Q96JY6_S123_S124 | 1.35976129 | 2.66E-03 | 1.36E-02 |
| CAAP1_HUMAN\|Q9H8G2_S92 | 1.359402957 | 2.25E-03 | 2.74E-02 |

| | | | |
|---|---|---|---|
| ARI1A_HUMAN\|O14497_S233 | 1.359316342 | 1.46E-03 | 1.24E-02 |
| IASPP_HUMAN\|Q8WUF5_Y109 | 1.359257913 | 4.27E-03 | 3.63E-02 |
| FA83B_HUMAN\|Q5T0W9_S731 | 1.359148603 | 3.79E-03 | 2.19E-02 |
| KPCD_HUMAN\|Q05655_S304 | 1.359098592 | 1.00E-03 | 9.81E-03 |
| HMGA2_HUMAN\|P52926_S14 | 1.358734319 | 5.06E-03 | 4.08E-02 |
| MILK1_HUMAN\|Q8N3F8_S486 | 1.358698849 | 2.09E-03 | 2.39E-02 |
| PACN2_HUMAN\|Q9UNF0_S399_S425 | 1.358489657 | 4.40E-03 | 3.61E-02 |
| TMF1_HUMAN\|P82094_S72 | 1.358388966 | 2.77E-03 | 1.41E-02 |
| RABE1_HUMAN\|Q15276_S407 | 1.358161498 | 3.31E-03 | 1.44E-02 |
| TX1B3_HUMAN\|O14907_S124 | 1.358087841 | 1.99E-03 | 9.59E-03 |
| AFAD_HUMAN\|P55196_S1173_S1182 | 1.358071055 | 6.24E-03 | 4.66E-02 |
| ACINU_HUMAN\|Q9UKV3_S243 | 1.357905082 | 2.27E-03 | 4.60E-02 |
| ROA3_HUMAN\|P51991_S355 | 1.357528263 | 1.27E-03 | 1.71E-02 |
| RHG01_HUMAN\|Q07960_S51 | 1.357425489 | 5.61E-03 | 2.50E-02 |
| RB6I2_HUMAN\|Q8IUD2_S37_S42_S45 | 1.357351138 | 7.28E-04 | 6.23E-03 |
| KAZRN_HUMAN\|Q674X7-3_S8 | 1.357187442 | 4.45E-03 | 2.47E-02 |
| RIOK2_HUMAN\|Q9BVS4_S380_S382_S385 | 1.357180664 | 4.48E-03 | 2.48E-02 |
| ZN185_HUMAN\|O15231_S155_T158 | 1.357120226 | 3.75E-03 | 3.31E-02 |
| MED1_HUMAN\|Q15648_Y1431 | 1.357091075 | 1.52E-03 | 1.11E-02 |
| MAP1B_HUMAN\|P46821_S1793_S1797 | 1.357026047 | 2.94E-03 | 2.81E-02 |
| SH24A_HUMAN\|Q9H788_S315 | 1.356850934 | 4.60E-03 | 3.82E-02 |
| KMT2C_HUMAN\|Q8NEZ4_S111 | 1.356677166 | 4.84E-03 | 3.19E-02 |
| CSKI2_HUMAN\|Q8WXE0_S403_S406 | 1.356624073 | 2.18E-04 | 2.48E-03 |
| F169A_HUMAN\|Q9Y6X4_T448 | 1.356485265 | 4.91E-03 | 4.01E-02 |
| FA83H_HUMAN\|Q6ZRV2_S998 | 1.356296384 | 6.79E-04 | 9.67E-03 |
| E41L2_HUMAN\|O43491_S87 | 1.356266862 | 5.84E-03 | 4.85E-02 |
| AHNK2_HUMAN\|Q8IVF2_S294_S297 | 1.356182156 | 1.04E-03 | 1.31E-02 |
| ABCF1_HUMAN\|Q8NE71_S109_T108 | 1.356078279 | 1.04E-03 | 8.21E-03 |
| FA83B_HUMAN\|Q5T0W9_S731 | 1.356061386 | 4.01E-03 | 3.41E-02 |
| C2CD5_HUMAN\|Q86YS7_S307 | 1.35578476 | 6.92E-03 | 4.09E-02 |
| SRRM2_HUMAN\|Q9UQ35_S377_S387_S398 | 1.355690804 | 5.21E-03 | 4.08E-02 |

| | | | |
|---|---|---|---|
| GGYF2_HUMAN\|Q6Y7W6_S26 | 1.355613243 | 1.12E-03 | 6.00E-03 |
| FMNL2_HUMAN\|Q96PY5_S1016 | 1.355374263 | 1.28E-02 | 4.20E-02 |
| RPRD2_HUMAN\|Q5VT52_S485 | 1.355367907 | 1.48E-03 | 2.08E-02 |
| IRS2_HUMAN\|Q9Y4H2_S1203 | 1.355313536 | 6.87E-04 | 1.20E-02 |
| RABL6_HUMAN\|Q3YEC7_S492 | 1.355156185 | 2.58E-03 | 1.65E-02 |
| KANK1_HUMAN\|Q14678_S144 | 1.354901941 | 2.95E-04 | 4.73E-03 |
| SRG2B_HUMAN\|P0DMP2_S206 | 1.354835836 | 8.90E-03 | 3.16E-02 |
| CYTSA_HUMAN\|Q69YQ0_S832 | 1.354677313 | 4.32E-04 | 8.25E-03 |
| PTN13_HUMAN\|Q12923_S1630 | 1.354619144 | 1.27E-03 | 1.71E-02 |
| FBXW9_HUMAN\|Q5XUX1_S18_T24 | 1.35457979 | 8.91E-03 | 4.74E-02 |
| SRRM2_HUMAN\|Q9UQ35_S2694 | 1.354467036 | 3.85E-04 | 1.57E-02 |
| TGS1_HUMAN\|Q96RS0_S55_T60 | 1.354338481 | 7.82E-04 | 1.25E-02 |
| RPB1_HUMAN\|P24928_S1845_S1857 | 1.354324323 | 8.82E-03 | 3.14E-02 |
| APBB2_HUMAN\|Q92870_S409_S412 | 1.3542194 | 3.94E-03 | 3.37E-02 |
| CCDC9_HUMAN\|Q9Y3X0_S386 | 1.354154331 | 7.29E-04 | 4.73E-03 |
| TBC9B_HUMAN\|Q66K14_S1241 | 1.354079362 | 9.06E-04 | 5.61E-03 |
| KI21A_HUMAN\|Q7Z4S6_S1229 | 1.353935307 | 2.04E-03 | 2.41E-02 |
| KMT2C_HUMAN\|Q8NEZ4_S111 | 1.353913217 | 5.10E-03 | 2.02E-02 |
| NIBL1_HUMAN\|Q96TA1_S691_S696 | 1.353899488 | 5.36E-03 | 3.41E-02 |
| CR025_HUMAN\|Q96B23_S145 | 1.353860231 | 1.62E-03 | 1.40E-02 |
| ZFP91_HUMAN\|Q96JP5_S83 | 1.35384047 | 1.54E-03 | 1.94E-02 |
| DEN4C_HUMAN\|Q5VZ89_S1046 | 1.35360106 | 2.04E-03 | 2.10E-02 |
| PTN12_HUMAN\|Q05209_S571_T587 | 1.353454332 | 2.46E-03 | 2.92E-02 |
| UBE3A_HUMAN\|Q05086_S218 | 1.353196673 | 1.26E-03 | 1.11E-02 |
| IASPP_HUMAN\|Q8WUF5_S567 | 1.353086362 | 7.89E-04 | 1.09E-02 |
| ARAF_HUMAN\|P10398_S157 | 1.352947378 | 2.62E-03 | 1.67E-02 |
| RHG23_HUMAN\|Q9P227_S351 | 1.352639874 | 6.77E-03 | 3.34E-02 |
| LAP2A_HUMAN\|P42166_S66_S67 | 1.352577395 | 1.76E-03 | 1.87E-02 |
| ZCH18_HUMAN\|Q86VM9_S67 | 1.352549768 | 4.52E-03 | 4.33E-02 |
| RBM10_HUMAN\|P98175_S61_S65_Y70 | 1.352517257 | 1.18E-02 | 3.92E-02 |
| AHNK2_HUMAN\|Q8IVF2_S278_S294 | 1.352091562 | 3.64E-03 | 2.12E-02 |

| | | | |
|---|---|---|---|
| HNRPC_HUMAN\|P07910_S260 | 1.352085753 | 1.95E-03 | 4.29E-02 |
| CC117_HUMAN\|Q8IWD4_S53 | 1.352027658 | 6.85E-03 | 4.97E-02 |
| CRBG1_HUMAN\|Q9Y4K1_S348 | 1.35185637 | 5.90E-03 | 3.48E-02 |
| OPTN_HUMAN\|Q96CV9_T525 | 1.35178561 | 2.40E-03 | 1.86E-02 |
| SRRM1_HUMAN\|Q8IYB3_S638 | 1.351763573 | 2.85E-03 | 1.77E-02 |
| CSN1_HUMAN\|Q13098_S474_T479 | 1.351628859 | 6.91E-03 | 3.93E-02 |
| PML_HUMAN\|P29590_S504 | 1.35157469 | 4.66E-03 | 1.89E-02 |
| PRSR2_HUMAN\|Q86WR7_S31 | 1.351466149 | 3.13E-03 | 1.55E-02 |
| CCDC9_HUMAN\|Q9Y3X0_S386 | 1.35136819 | 7.81E-04 | 1.30E-02 |
| F126B_HUMAN\|Q8IXS8_S430 | 1.35129836 | 1.62E-03 | 1.40E-02 |
| IF4B_HUMAN\|P23588_S489_S497_S498 | 1.35109329 | 4.53E-03 | 4.33E-02 |
| RB6I2_HUMAN\|Q8IUD2_S37_S42_S45 | 1.350965661 | 8.53E-04 | 1.07E-02 |
| NIBAN_HUMAN\|Q9BZQ8_S926 | 1.35095647 | 3.75E-03 | 3.31E-02 |
| ZN185_HUMAN\|O15231-2_T64 | 1.350875696 | 1.52E-03 | 1.11E-02 |
| BCL9L_HUMAN\|Q86UU0_S21_S25 | 1.350870517 | 2.36E-03 | 1.84E-02 |
| ZCCHV_HUMAN\|Q7Z2W4_S284 | 1.350788798 | 2.85E-03 | 3.26E-02 |
| SQSTM_HUMAN\|Q13501_T269 | 1.350683381 | 6.31E-03 | 3.17E-02 |
| KI21A_HUMAN\|Q7Z4S6_S855 | 1.350657497 | 6.52E-03 | 4.88E-02 |
| ZN185_HUMAN\|O15231_S153 | 1.35062491 | 3.87E-03 | 3.87E-02 |
| MATR3_HUMAN\|P43243_S598 | 1.350541545 | 4.23E-04 | 1.65E-02 |
| LMNA_HUMAN\|P02545_S18 | 1.35047186 | 3.61E-03 | 2.45E-02 |
| FA83B_HUMAN\|Q5T0W9_S869 | 1.350447409 | 1.54E-04 | 5.62E-03 |
| RPRD2_HUMAN\|Q5VT52_S485 | 1.350111226 | 1.66E-03 | 8.30E-03 |
| CBX3_HUMAN\|Q13185_S99 | 1.350063139 | 1.03E-03 | 8.15E-03 |
| PIEZ1_HUMAN\|Q92508_S1395_S1396 | 1.349900464 | 7.10E-03 | 2.60E-02 |
| NHRF1_HUMAN\|O14745_S280 | 1.34962509 | 5.07E-03 | 4.92E-02 |
| BCLF1_HUMAN\|Q9NYF8_S222 | 1.349501798 | 1.60E-03 | 3.92E-02 |
| ANR17_HUMAN\|O75179_S2045_S2047 | 1.349418551 | 1.54E-03 | 8.60E-03 |
| CE170_HUMAN\|Q5SW79_S971 | 1.349399218 | 5.97E-03 | 4.52E-02 |
| IRS1_HUMAN\|P35568_S1101 | 1.349273441 | 3.07E-03 | 2.79E-02 |
| R3HD1_HUMAN\|Q15032_S973 | 1.349119616 | 1.98E-03 | 2.35E-02 |

| | | | |
|---|---|---|---|
| S11IP_HUMAN\|Q8N1F8_S398 | 1.349017657 | 3.83E-03 | 3.87E-02 |
| GIT1_HUMAN\|Q9Y2X7_S362 | 1.348981066 | 2.75E-03 | 2.67E-02 |
| NDRG2_HUMAN\|Q9UN36_S332_T330 | 1.348799694 | 5.86E-03 | 4.52E-02 |
| CAVN1_HUMAN\|Q6NZI2_T302 | 1.34869845 | 2.09E-03 | 1.11E-02 |
| HELZ_HUMAN\|P42694_T1163 | 1.348623143 | 3.50E-04 | 5.97E-03 |
| PLEC_HUMAN\|Q15149_S4385_S4386 | 1.348392032 | 2.48E-04 | 1.84E-03 |
| RBMX_HUMAN\|P38159_S175 | 1.347913682 | 3.82E-03 | 2.20E-02 |
| TBC15_HUMAN\|Q8TC07_S675 | 1.34766325 | 3.11E-03 | 1.90E-02 |
| HDAC2_HUMAN\|Q92769_S394 | 1.347524782 | 6.28E-03 | 4.76E-02 |
| MON1A_HUMAN\|Q86VX9_Y57 | 1.347462848 | 3.04E-03 | 3.12E-02 |
| RABE1_HUMAN\|Q15276_S410_T408 | 1.347325399 | 3.75E-03 | 3.31E-02 |
| LMNA_HUMAN\|P02545_S407_S414 | 1.347288593 | 1.06E-02 | 4.74E-02 |
| LMNA_HUMAN\|P02545_S22 | 1.347233679 | 1.52E-03 | 2.05E-02 |
| EI2BE_HUMAN\|Q13144_S544 | 1.347174323 | 8.61E-03 | 4.65E-02 |
| NRK_HUMAN\|Q7Z2Y5_S854_Y858 | 1.347143635 | 4.33E-03 | 3.56E-02 |
| NMD3_HUMAN\|Q96D46_S468_T470 | 1.347116853 | 6.99E-04 | 6.06E-03 |
| SVIL_HUMAN\|O95425_T679 | 1.346924239 | 9.54E-04 | 1.23E-02 |
| CPSF4_HUMAN\|O95639_S200_S212 | 1.346558264 | 5.93E-03 | 3.49E-02 |
| ABI2_HUMAN\|Q9NYB9_S368 | 1.346512054 | 9.94E-04 | 1.20E-02 |
| PAXX_HUMAN\|Q9BUH6_S148 | 1.34647933 | 3.31E-03 | 1.44E-02 |
| CLASR_HUMAN\|Q8N2M8_S547 | 1.346149964 | 1.14E-03 | 8.85E-03 |
| SVIL_HUMAN\|O95425_S63 | 1.345978794 | 1.02E-03 | 2.86E-02 |
| MY18A_HUMAN\|Q92614_S1998 | 1.345937834 | 3.87E-03 | 3.87E-02 |
| PLEC_HUMAN\|Q15149_S4670 | 1.345787321 | 5.68E-03 | 3.37E-02 |
| KLC2_HUMAN\|Q9H0B6_S611 | 1.345752174 | 1.05E-02 | 4.69E-02 |
| PFD2_HUMAN\|Q9UHV9_S147 | 1.345501867 | 2.64E-03 | 2.51E-02 |
| KIF1C_HUMAN\|O43896_S676 | 1.345475523 | 2.62E-03 | 1.35E-02 |
| AKAP8_HUMAN\|O43823_S328 | 1.345340904 | 1.08E-02 | 4.80E-02 |
| PRP31_HUMAN\|Q8WWY3_S498 | 1.345257429 | 6.13E-04 | 5.57E-03 |
| SWAHC_HUMAN\|Q53LP3_S212 | 1.345087148 | 1.58E-03 | 1.71E-02 |
| ADNP_HUMAN\|Q9H2P0_S921 | 1.345010148 | 3.72E-03 | 3.23E-02 |

| | | | |
|---|---|---|---|
| PP4R1_HUMAN\|Q8TF05_S947_S948 | 1.344870807 | 6.07E-03 | 4.95E-02 |
| PLEC_HUMAN\|Q15149_S4386_S4389 | 1.344869491 | 5.62E-03 | 3.35E-02 |
| PTN12_HUMAN\|Q05209_S571_T587 | 1.344687602 | 2.94E-03 | 2.09E-02 |
| LASP1_HUMAN\|Q14847_T104 | 1.344651208 | 4.47E-03 | 4.06E-02 |
| P66B_HUMAN\|Q8WXI9_S129_T120 | 1.344646869 | 3.01E-03 | 2.86E-02 |
| VIME_HUMAN\|P08670_T458 | 1.344624784 | 4.00E-03 | 3.40E-02 |
| ZN185_HUMAN\|O15231_S153 | 1.34451195 | 4.37E-03 | 2.96E-02 |
| INT12_HUMAN\|Q96CB8_S378 | 1.34442265 | 2.08E-03 | 2.44E-02 |
| IF4G1_HUMAN\|Q04637_S1187 | 1.344336899 | 5.66E-03 | 4.41E-02 |
| R3HD1_HUMAN\|Q15032_S973 | 1.344268639 | 2.20E-03 | 2.26E-02 |
| TPIS_HUMAN\|P60174_Y246 | 1.344262207 | 6.36E-03 | 3.19E-02 |
| SRPK2_HUMAN\|P78362_S496_T498 | 1.344199077 | 8.31E-03 | 4.65E-02 |
| AFAD_HUMAN\|P55196_S1182 | 1.344138803 | 7.01E-04 | 4.58E-03 |
| SCML2_HUMAN\|Q9UQR0_S590 | 1.344010386 | 1.17E-03 | 1.75E-02 |
| TGS1_HUMAN\|Q96RS0_S55 | 1.343830819 | 8.84E-03 | 4.85E-02 |
| MYPT1_HUMAN\|O14974_S507 | 1.343681304 | 8.75E-03 | 4.70E-02 |
| MFAP1_HUMAN\|P55081_S80 | 1.343274171 | 2.18E-03 | 2.20E-02 |
| MMTA2_HUMAN\|Q9BU76_S216_S220 | 1.343271639 | 8.87E-03 | 4.74E-02 |
| P66B_HUMAN\|Q8WXI9_S122_S129_T120 | 1.342975163 | 1.30E-03 | 1.56E-02 |
| LAP2A_HUMAN\|P42166_T160 | 1.342971364 | 3.62E-03 | 3.51E-02 |
| AAK1_HUMAN\|Q2M2I8_S678_T674 | 1.342891593 | 3.65E-03 | 2.12E-02 |
| DC1L2_HUMAN\|O43237_S194 | 1.342792097 | 3.89E-04 | 7.68E-03 |
| CDK18_HUMAN\|Q07002_S87 | 1.342571087 | 2.80E-04 | 5.97E-03 |
| SPIR1_HUMAN\|Q08AE8_T509 | 1.342514704 | 7.33E-03 | 3.55E-02 |
| DNLI1_HUMAN\|P18858_S66 | 1.342185943 | 1.02E-02 | 3.50E-02 |
| AF1L2_HUMAN\|Q8N4X5_S651 | 1.342104153 | 1.20E-03 | 1.47E-02 |
| ARH40_HUMAN\|Q8TER5_S1513 | 1.342082699 | 9.72E-04 | 1.55E-02 |
| NMD3_HUMAN\|Q96D46_S468_T470 | 1.341862985 | 7.94E-04 | 1.09E-02 |
| STMN1_HUMAN\|P16949_S25 | 1.341693948 | 3.88E-03 | 2.58E-02 |
| PER1_HUMAN\|O15534_S1007 | 1.341683601 | 5.25E-03 | 4.50E-02 |
| SRRM2_HUMAN\|Q9UQ35_S2694 | 1.341613577 | 5.43E-04 | 5.08E-03 |

| | | | |
|---|---|---|---|
| P66B_HUMAN\|Q8WXI9_S334 | 1.341441908 | 9.70E-04 | 1.25E-02 |
| IF2P_HUMAN\|O60841_S182_S183_S186 | 1.341287383 | 7.12E-03 | 4.01E-02 |
| IRS1_HUMAN\|P35568_S1100 | 1.341106391 | 6.15E-03 | 5.00E-02 |
| PKHG3_HUMAN\|A1L390_S638_S643 | 1.340950871 | 7.36E-04 | 1.24E-02 |
| MA7D1_HUMAN\|Q3KQU3_S113 | 1.34086537 | 6.47E-03 | 4.86E-02 |
| CPSF4_HUMAN\|O95639_S200 | 1.340754154 | 5.58E-03 | 4.28E-02 |
| SAMH1_HUMAN\|Q9Y3Z3_S33 | 1.340700152 | 1.00E-02 | 4.54E-02 |
| TPBG_HUMAN\|Q13641_S415_S418 | 1.340682594 | 4.53E-03 | 3.02E-02 |
| CCDC9_HUMAN\|Q9Y3X0_S386 | 1.340599906 | 1.02E-03 | 9.95E-03 |
| ASHWN_HUMAN\|Q9BVC5_S193 | 1.340553703 | 1.41E-03 | 2.67E-02 |
| KLC3_HUMAN\|Q6P597_S497_S502 | 1.340457159 | 9.41E-04 | 9.06E-03 |
| VIME_HUMAN\|P08670_S65 | 1.340381942 | 5.93E-03 | 4.88E-02 |
| ZN185_HUMAN\|O15231-2_T64 | 1.340359067 | 1.93E-03 | 4.44E-02 |
| TPBG_HUMAN\|Q13641_S415_S418 | 1.340325969 | 4.56E-03 | 4.61E-02 |
| ARHGG_HUMAN\|Q5VV41_S227_T226 | 1.340242208 | 3.98E-03 | 3.73E-02 |
| NRK_HUMAN\|Q7Z2Y5_S854_Y858 | 1.340094724 | 4.96E-03 | 4.62E-02 |
| OXR1_HUMAN\|Q8N573_S201_S202 | 1.340001454 | 3.92E-03 | 3.43E-02 |
| JUNB_HUMAN\|P17275_T255 | 1.339704595 | 1.18E-03 | 3.18E-02 |
| MTAP2_HUMAN\|P11137_S1591_T1602_T1605 | 1.339672285 | 2.87E-03 | 1.78E-02 |
| INADL_HUMAN\|Q8NI35_S645 | 1.339555704 | 3.34E-03 | 3.31E-02 |
| UFL1_HUMAN\|O94874_S458 | 1.339546384 | 5.25E-04 | 5.83E-03 |
| DDB2_HUMAN\|Q92466_S24_S26 | 1.339483809 | 4.40E-04 | 9.01E-03 |
| UT14A_HUMAN\|Q9BVJ6_S434 | 1.338960758 | 9.71E-03 | 3.37E-02 |
| RHG23_HUMAN\|Q9P227_S611 | 1.33891569 | 4.77E-03 | 3.93E-02 |
| AFTIN_HUMAN\|Q6ULP2_S518 | 1.338410899 | 6.23E-03 | 2.74E-02 |
| MTAP2_HUMAN\|P11137_S1782_S1786_S1790 | 1.338014926 | 7.02E-03 | 3.43E-02 |
| RBCC1_HUMAN\|Q8TDY2_S653 | 1.337988193 | 1.26E-02 | 4.15E-02 |
| AHNK_HUMAN\|Q09666_S216_S220 | 1.337899377 | 3.51E-03 | 3.18E-02 |
| PALLD_HUMAN\|Q8WX93_S1104 | 1.337874238 | 5.12E-03 | 4.46E-02 |
| YBOX3_HUMAN\|P16989_S203_S204 | 1.337835896 | 6.83E-03 | 4.06E-02 |
| SRGP1_HUMAN\|Q7Z6B7_S906 | 1.337707985 | 1.06E-02 | 4.72E-02 |

| | | | |
|---|---|---|---|
| TLE3_HUMAN\|Q04726_S203 | 1.337461508 | 9.73E-03 | 4.45E-02 |
| ZYX_HUMAN\|Q15942_S259 | 1.337444426 | 5.04E-03 | 2.71E-02 |
| SCRIB_HUMAN\|Q14160_S1448 | 1.337379093 | 1.09E-02 | 3.71E-02 |
| TACC2_HUMAN\|O95359_S2072 | 1.337286467 | 4.39E-03 | 4.24E-02 |
| DCTN1_HUMAN\|Q14203_S105 | 1.337206307 | 5.13E-03 | 2.33E-02 |
| OTUL_HUMAN\|Q96BN8_S351 | 1.337201046 | 6.47E-04 | 1.11E-02 |
| ARVC_HUMAN\|O00192_T642 | 1.337121085 | 1.04E-02 | 4.65E-02 |
| RPRD2_HUMAN\|Q5VT52_T723 | 1.337033653 | 1.48E-03 | 1.63E-02 |
| RRP8_HUMAN\|O43159_S104_S106 | 1.337019457 | 3.22E-03 | 3.22E-02 |
| UBP20_HUMAN\|Q9Y2K6_S132 | 1.336849606 | 4.35E-03 | 3.67E-02 |
| PKP3_HUMAN\|Q9Y446_S313 | 1.336817852 | 7.96E-04 | 1.09E-02 |
| RIOK1_HUMAN\|Q9BRS2_S22 | 1.336771796 | 2.96E-04 | 3.14E-03 |
| PACN3_HUMAN\|Q9UKS6_S344_S354 | 1.336762275 | 5.73E-03 | 2.55E-02 |
| ASHWN_HUMAN\|Q9BVC5_S193 | 1.336697896 | 1.54E-03 | 1.12E-02 |
| RFIP5_HUMAN\|Q9BXF6_S396 | 1.336632565 | 7.73E-03 | 4.44E-02 |
| LYAR_HUMAN\|Q9NX58_S276 | 1.336574234 | 2.46E-03 | 2.92E-02 |
| ASPP2_HUMAN\|Q13625_S480 | 1.336097617 | 3.40E-03 | 2.03E-02 |
| AHNK_HUMAN\|Q09666_S210_S216_S220 | 1.3360299 | 4.91E-03 | 4.82E-02 |
| RHG21_HUMAN\|Q5T5U3_S615 | 1.335994067 | 3.67E-03 | 3.29E-02 |
| CCSE2_HUMAN\|Q9H7U1_S197 | 1.335965728 | 9.17E-04 | 7.44E-03 |
| TLN1_HUMAN\|Q9Y490_S1641 | 1.335921713 | 1.79E-03 | 1.26E-02 |
| F16B2_HUMAN\|Q86V87_S736 | 1.335842074 | 9.44E-03 | 3.30E-02 |
| IP3KB_HUMAN\|P27987_S43 | 1.335788809 | 5.26E-03 | 3.18E-02 |
| PLEC_HUMAN\|Q15149_S4385_S4386 | 1.335724279 | 3.57E-04 | 1.07E-02 |
| DC1L1_HUMAN\|Q9Y6G9_S510 | 1.335652333 | 6.02E-04 | 2.12E-02 |
| CR025_HUMAN\|Q96B23_S66 | 1.335626922 | 3.49E-03 | 1.51E-02 |
| OXR1_HUMAN\|Q8N573_S201_S202 | 1.335603291 | 4.28E-03 | 3.53E-02 |
| CI078_HUMAN\|Q9NZ63_S261 | 1.335501988 | 5.46E-03 | 2.86E-02 |
| OTU7B_HUMAN\|Q6GQQ9_S467 | 1.335423237 | 2.12E-03 | 2.46E-02 |
| RIOK1_HUMAN\|Q9BRS2_S21 | 1.33540541 | 1.09E-03 | 8.50E-03 |
| OSBP1_HUMAN\|P22059_S379_S382 | 1.33532715 | 2.99E-03 | 1.33E-02 |

| | | | |
|---|---|---|---|
| SH23A_HUMAN\|Q9BRG2_S180 | 1.335318537 | 1.48E-03 | 8.33E-03 |
| NIBL1_HUMAN\|Q96TA1_S691 | 1.335235185 | 9.85E-04 | 1.53E-02 |
| 1B07_HUMAN\|P01889_S359 | 1.335140817 | 6.40E-03 | 4.82E-02 |
| DC1L1_HUMAN\|Q9Y6G9_S510 | 1.335115033 | 6.10E-04 | 5.55E-03 |
| NRK_HUMAN\|Q7Z2Y5_S854 | 1.334908027 | 1.23E-03 | 1.43E-02 |
| RPC5_HUMAN\|Q9NVU0_S161 | 1.334905952 | 1.51E-02 | 4.79E-02 |
| NRK_HUMAN\|Q7Z2Y5_S854 | 1.334768648 | 1.23E-03 | 3.20E-02 |
| RANB3_HUMAN\|Q9H6Z4_S100_S101_S108 | 1.33451632 | 5.39E-03 | 2.84E-02 |
| DYST_HUMAN\|Q03001_S1382 | 1.334472389 | 6.60E-04 | 4.35E-03 |
| BRE1A_HUMAN\|Q5VTR2_S136 | 1.334468539 | 1.60E-03 | 2.00E-02 |
| APC_HUMAN\|P25054_S2242_S2244 | 1.333918793 | 1.68E-03 | 2.07E-02 |
| HNRL2_HUMAN\|Q1KMD3_S161 | 1.333917965 | 5.14E-03 | 3.14E-02 |
| SON_HUMAN\|P18583_S94 | 1.333758423 | 1.94E-03 | 9.39E-03 |
| CAMP3_HUMAN\|Q9P1Y5_S553 | 1.333691776 | 6.38E-04 | 1.14E-02 |
| PHIP_HUMAN\|Q8WWQ0_S1783 | 1.333655439 | 3.55E-03 | 4.87E-02 |
| LARP6_HUMAN\|Q9BRS8_S56_S58 | 1.333548731 | 1.13E-02 | 4.98E-02 |
| PDS5B_HUMAN\|Q9NTI5_S1257_S1259 | 1.333361935 | 5.94E-03 | 2.62E-02 |
| OSTF1_HUMAN\|Q92882_S202_S213 | 1.333299617 | 2.12E-03 | 2.19E-02 |
| CBX3_HUMAN\|Q13185_S95 | 1.333104525 | 7.32E-04 | 7.41E-03 |
| K1671_HUMAN\|Q9BY89_S127 | 1.333060479 | 5.21E-03 | 4.08E-02 |
| TP53B_HUMAN\|Q12888_S525 | 1.332907422 | 3.27E-03 | 1.61E-02 |
| PDS5B_HUMAN\|Q9NTI5_S1362 | 1.332740309 | 4.41E-03 | 1.82E-02 |
| BORG5_HUMAN\|Q00587_S303 | 1.3326978 | 5.40E-03 | 2.84E-02 |
| AHNK_HUMAN\|Q09666_S5763 | 1.332614258 | 1.26E-03 | 6.65E-03 |
| PRSR2_HUMAN\|Q86WR7_S18 | 1.33254339 | 8.11E-04 | 6.80E-03 |
| ARH40_HUMAN\|Q8TER5_S1474 | 1.33254083 | 6.84E-03 | 4.06E-02 |
| KHDR1_HUMAN\|Q07666_S20 | 1.332449737 | 6.20E-03 | 4.64E-02 |
| CBX3_HUMAN\|Q13185_S99 | 1.332308179 | 1.57E-03 | 3.76E-02 |
| HABP4_HUMAN\|Q5JVS0_S108 | 1.332295169 | 3.01E-03 | 2.86E-02 |
| GGYF2_HUMAN\|Q6Y7W6_S23 | 1.332234326 | 2.65E-03 | 1.99E-02 |
| RIPR1_HUMAN\|Q6ZS17_S351 | 1.332040856 | 2.07E-03 | 2.44E-02 |

| | | | |
|---|---|---|---|
| MK06_HUMAN\|Q16659_S386_T389 | 1.331990056 | 6.45E-04 | 5.76E-03 |
| CTNA1_HUMAN\|P35221_T654 | 1.331882485 | 6.89E-03 | 4.08E-02 |
| TENS4_HUMAN\|Q8IZW8_S253 | 1.331879628 | 3.38E-03 | 3.11E-02 |
| ZO2_HUMAN\|Q9UDY2_S170 | 1.33186612 | 3.88E-03 | 4.11E-02 |
| MI4GD_HUMAN\|A9UHW6_S221 | 1.331846816 | 1.36E-03 | 1.61E-02 |
| MTAP2_HUMAN\|P11137_S1787_S1790_T1780 | 1.33180674 | 4.00E-03 | 2.28E-02 |
| MYO9B_HUMAN\|Q13459_S1992 | 1.331533317 | 9.43E-03 | 3.30E-02 |
| F118A_HUMAN\|Q9NWS6_S349 | 1.331441868 | 2.30E-03 | 2.79E-02 |
| ZFP91_HUMAN\|Q96JP5_S83 | 1.331329581 | 2.56E-03 | 4.90E-02 |
| NIPA_HUMAN\|Q86WB0_S407_S410_S411 | 1.331315635 | 1.99E-03 | 9.59E-03 |
| CD2B2_HUMAN\|O95400_S195 | 1.331101569 | 3.47E-04 | 3.54E-03 |
| TISB_HUMAN\|Q07352_S54 | 1.330982583 | 2.11E-03 | 1.44E-02 |
| NAA15_HUMAN\|Q9BXJ9_S855 | 1.330975152 | 1.06E-02 | 3.62E-02 |
| 2A5E_HUMAN\|Q16537_S33_S34 | 1.330819059 | 1.98E-03 | 4.49E-02 |
| PKHA2_HUMAN\|Q9HB19_S184 | 1.330751382 | 2.17E-03 | 1.66E-02 |
| SIPA1_HUMAN\|Q96FS4_S55 | 1.330641677 | 3.75E-03 | 3.89E-02 |
| DKC1_HUMAN\|O60832_T496 | 1.330617941 | 1.23E-03 | 3.20E-02 |
| MAP1B_HUMAN\|P46821_Y1796 | 1.330440535 | 5.33E-03 | 4.53E-02 |
| SLK_HUMAN\|Q9H2G2_T820 | 1.330209765 | 1.98E-03 | 2.31E-02 |
| NELFE_HUMAN\|P18615_S353 | 1.330015758 | 8.23E-04 | 6.87E-03 |
| KS6A1_HUMAN\|Q15418_T359 | 1.329961358 | 1.46E-04 | 4.09E-03 |
| LIPB1_HUMAN\|Q86W92_T603 | 1.329879685 | 1.04E-02 | 4.65E-02 |
| CHM2A_HUMAN\|O43633_S203 | 1.329576345 | 5.32E-03 | 2.82E-02 |
| BCL9L_HUMAN\|Q86UU0_S21_S25 | 1.329556743 | 3.71E-03 | 1.78E-02 |
| BICD2_HUMAN\|Q8TD16_S615 | 1.329474253 | 6.35E-03 | 3.69E-02 |
| K1468_HUMAN\|Q9P260_S180 | 1.329369423 | 5.19E-03 | 3.35E-02 |
| 1433G_HUMAN\|P61981_T234 | 1.32935829 | 6.71E-03 | 4.99E-02 |
| KLC1_HUMAN\|Q07866-4_S631 | 1.329350033 | 2.63E-03 | 1.67E-02 |
| AFAD_HUMAN\|P55196_S1182 | 1.329223658 | 1.01E-03 | 9.54E-03 |
| COBL1_HUMAN\|Q53SF7_S294 | 1.329149685 | 2.82E-03 | 1.76E-02 |
| DESP_HUMAN\|P15924_S2815_S2820 | 1.329008849 | 5.22E-03 | 2.06E-02 |

| | | | |
|---|---|---|---|
| ANKH1_HUMAN\|Q8IWZ3_S1591_S1593 | 1.328961689 | 1.17E-03 | 1.62E-02 |
| EPIPL_HUMAN\|P58107_S2716 | 1.328936305 | 1.16E-03 | 1.74E-02 |
| NDRG2_HUMAN\|Q9UN36_S332 | 1.328915168 | 5.66E-03 | 4.33E-02 |
| TLN1_HUMAN\|Q9Y490_S729 | 1.328631455 | 9.11E-03 | 4.21E-02 |
| CDK18_HUMAN\|Q07002_S87 | 1.328485749 | 4.17E-04 | 1.64E-02 |
| PKHG3_HUMAN\|A1L390_S1081 | 1.328106692 | 1.89E-03 | 2.46E-02 |
| 3BP5L_HUMAN\|Q7L8J4_S343 | 1.328082791 | 3.94E-03 | 3.44E-02 |
| BCL9L_HUMAN\|Q86UU0_S118_S121 | 1.32805835 | 1.49E-03 | 2.09E-02 |
| TBC9B_HUMAN\|Q66K14_S1241 | 1.327614785 | 1.69E-03 | 1.44E-02 |
| LAR4B_HUMAN\|Q92615_T732 | 1.327405564 | 1.21E-03 | 3.17E-02 |
| MAP1B_HUMAN\|P46821_S1785 | 1.32723907 | 1.34E-03 | 2.57E-02 |
| RAE1_HUMAN\|P24386_S651 | 1.3271929 | 3.84E-03 | 3.93E-02 |
| MVD1_HUMAN\|P53602_S96 | 1.32717325 | 2.03E-03 | 4.54E-02 |
| NASP_HUMAN\|P49321_S421 | 1.327104155 | 3.16E-03 | 1.93E-02 |
| CACO1_HUMAN\|Q9P1Z2_S563 | 1.32707953 | 1.52E-03 | 1.11E-02 |
| RTKN_HUMAN\|Q9BST9_S106 | 1.327058105 | 1.40E-02 | 4.53E-02 |
| RPRD2_HUMAN\|Q5VT52_T723 | 1.327015179 | 1.87E-03 | 4.19E-02 |
| NGAP_HUMAN\|Q9UJF2_S802 | 1.326979274 | 1.59E-03 | 3.92E-02 |
| EP15R_HUMAN\|Q9UBC2_S229 | 1.326754161 | 9.06E-04 | 1.20E-02 |
| PKHG3_HUMAN\|A1L390_S617 | 1.326547334 | 5.05E-03 | 4.08E-02 |
| RABE1_HUMAN\|Q15276_S407_S410 | 1.326441715 | 4.37E-04 | 7.07E-03 |
| PKHG3_HUMAN\|A1L390_S640 | 1.326409823 | 1.10E-02 | 4.85E-02 |
| TP53B_HUMAN\|Q12888_S366 | 1.326156128 | 2.24E-03 | 1.78E-02 |
| APC_HUMAN\|P25054_S2242_S2244 | 1.326116572 | 2.02E-03 | 2.57E-02 |
| SRRM2_HUMAN\|Q9UQ35_S1014_T1003 | 1.326053509 | 2.35E-03 | 1.55E-02 |
| DAPK3_HUMAN\|O43293_S311_S312 | 1.325968684 | 6.62E-03 | 4.93E-02 |
| IRS2_HUMAN\|Q9Y4H2_S1203 | 1.32595369 | 1.41E-03 | 2.02E-02 |
| RRBP1_HUMAN\|Q9P2E9_S615 | 1.325447059 | 1.42E-02 | 4.59E-02 |
| HNRPC_HUMAN\|P07910_S233_S238 | 1.325366392 | 9.15E-03 | 4.82E-02 |
| RHG32_HUMAN\|A7KAX9_S952 | 1.325101906 | 7.30E-04 | 1.23E-02 |
| ARHGB_HUMAN\|O15085_T668_T672 | 1.324878238 | 4.86E-03 | 2.63E-02 |

| | | | |
|---|---|---|---|
| HELZ_HUMAN\|P42694_T1163 | 1.324852856 | 6.73E-04 | 5.94E-03 |
| GGYF2_HUMAN\|Q6Y7W6_S26 | 1.32464132 | 2.25E-03 | 2.77E-02 |
| YTDC2_HUMAN\|Q9H6S0_S1090 | 1.324635462 | 1.11E-03 | 6.60E-03 |
| ABI2_HUMAN\|Q9NYB9_Y371 | 1.324548728 | 4.66E-03 | 3.76E-02 |
| IF2P_HUMAN\|O60841_S182_S183 | 1.324468331 | 4.54E-03 | 4.33E-02 |
| LIPA1_HUMAN\|Q13136_S1172 | 1.324378056 | 4.22E-03 | 2.38E-02 |
| ARMX3_HUMAN\|Q9UH62_S110 | 1.324172899 | 8.95E-03 | 3.17E-02 |
| I2BPL_HUMAN\|Q9H1B7_S657_S662 | 1.324049832 | 5.34E-03 | 3.41E-02 |
| CCD86_HUMAN\|Q9H6F5_S91 | 1.324000571 | 3.00E-03 | 4.44E-02 |
| AT11C_HUMAN\|Q8NB49_T1124 | 1.323991425 | 1.09E-02 | 3.71E-02 |
| VIGLN_HUMAN\|Q00341_S31 | 1.323878446 | 5.71E-03 | 3.38E-02 |
| DEN4C_HUMAN\|Q5VZ89_S1184 | 1.323838629 | 6.23E-03 | 4.73E-02 |
| GPTC2_HUMAN\|Q9NW75_S115_S117 | 1.323787633 | 2.78E-03 | 2.62E-02 |
| KS6A1_HUMAN\|Q15418_S363 | 1.323531045 | 1.98E-03 | 2.54E-02 |
| LMNA_HUMAN\|P02545_S428 | 1.323445191 | 3.87E-03 | 3.67E-02 |
| PKP3_HUMAN\|Q9Y446_S313_S314 | 1.323347381 | 5.36E-03 | 4.93E-02 |
| NOC2L_HUMAN\|Q9Y3T9_S56 | 1.323253764 | 1.11E-03 | 1.02E-02 |
| SH23A_HUMAN\|Q9BRG2_S180 | 1.323213985 | 1.98E-03 | 2.35E-02 |
| LIPB1_HUMAN\|Q86W92_S579 | 1.323087313 | 4.29E-03 | 2.40E-02 |
| PLEC_HUMAN\|Q15149_S4626 | 1.322575648 | 5.07E-03 | 4.92E-02 |
| ACAP2_HUMAN\|Q15057_T382 | 1.322505811 | 1.05E-03 | 2.17E-02 |
| MON1A_HUMAN\|Q86VX9_Y57 | 1.322483292 | 5.07E-03 | 3.11E-02 |
| ROA3_HUMAN\|P51991_S356 | 1.322324653 | 5.59E-03 | 2.90E-02 |
| CE170_HUMAN\|Q5SW79_S630_S633 | 1.322323031 | 5.63E-03 | 4.73E-02 |
| SH23A_HUMAN\|Q9BRG2_S125 | 1.32201334 | 2.21E-03 | 2.27E-02 |
| TWF1_HUMAN\|Q12792_T349 | 1.321825165 | 3.36E-03 | 2.02E-02 |
| RABE1_HUMAN\|Q15276_S410_T408 | 1.32181523 | 6.15E-03 | 3.11E-02 |
| DKC1_HUMAN\|O60832_S485_T496 | 1.321596905 | 1.87E-03 | 1.30E-02 |
| AHNK_HUMAN\|Q09666_S216 | 1.321535352 | 3.52E-03 | 2.08E-02 |
| SPE39_HUMAN\|Q9H9C1_S121 | 1.321488699 | 1.59E-03 | 3.92E-02 |
| CNOT2_HUMAN\|Q9NZN8_S165 | 1.321352903 | 5.08E-03 | 2.72E-02 |

| | | | |
|---|---|---|---|
| ASPP2_HUMAN\|Q13625_S558 | 1.320610189 | 4.10E-03 | 4.05E-02 |
| EF2K_HUMAN\|O00418_S27_S31 | 1.320354725 | 1.82E-03 | 2.17E-02 |
| AGFG1_HUMAN\|P52594_S181 | 1.320165063 | 2.18E-03 | 1.47E-02 |
| IF4G2_HUMAN\|P78344_S17 | 1.320041375 | 6.01E-04 | 8.92E-03 |
| CEP89_HUMAN\|Q96ST8_S40_S44 | 1.320040825 | 8.16E-03 | 4.46E-02 |
| AHNK2_HUMAN\|Q8IVF2_S4185 | 1.319962228 | 5.02E-03 | 2.00E-02 |
| DC1L2_HUMAN\|O43237_S194 | 1.319933061 | 7.24E-04 | 1.69E-02 |
| NOG1_HUMAN\|Q9BZE4_S470_Y466 | 1.319924076 | 8.73E-03 | 3.60E-02 |
| NFIB_HUMAN\|O00712_S295 | 1.31969214 | 4.28E-03 | 3.94E-02 |
| RB6I2_HUMAN\|Q8IUD2_S37_S42_S45 | 1.319626321 | 1.86E-03 | 2.20E-02 |
| RABE1_HUMAN\|Q15276_S407_S410 | 1.319605718 | 5.33E-04 | 5.00E-03 |
| RHG23_HUMAN\|Q9P227_S361_S372 | 1.319604044 | 2.03E-03 | 2.12E-02 |
| SRG2B_HUMAN\|P0DMP2_T203 | 1.319503586 | 1.98E-03 | 2.55E-02 |
| KMT2D_HUMAN\|O14686_S2274 | 1.319432466 | 1.25E-02 | 4.81E-02 |
| BAD_HUMAN\|Q92934_Y76 | 1.319386774 | 4.66E-03 | 4.40E-02 |
| PTN14_HUMAN\|Q15678_S593_S594 | 1.319182617 | 2.62E-03 | 1.67E-02 |
| PDS5B_HUMAN\|Q9NTI5_S1259 | 1.319159311 | 6.92E-03 | 2.55E-02 |
| EPN1_HUMAN\|Q9Y6I3_S454 | 1.319037965 | 4.17E-03 | 4.20E-02 |
| NIBL1_HUMAN\|Q96TA1_S696 | 1.318988755 | 1.07E-02 | 4.76E-02 |
| LARP1_HUMAN\|Q6PKG0_S517_S521 | 1.31889472 | 2.34E-03 | 2.31E-02 |
| PRSR2_HUMAN\|Q86WR7_S43 | 1.318836867 | 2.86E-03 | 2.11E-02 |
| ATX2L_HUMAN\|Q8WWM7_S111 | 1.318786228 | 3.46E-03 | 3.59E-02 |
| NIPA_HUMAN\|Q86WB0_S409 | 1.318383431 | 6.47E-04 | 4.29E-03 |
| ARI1A_HUMAN\|O14497_S696_S698 | 1.318347297 | 5.00E-03 | 3.08E-02 |
| SVIL_HUMAN\|O95425-2_S270 | 1.318223414 | 4.51E-03 | 2.50E-02 |
| OPTN_HUMAN\|Q96CV9_T525 | 1.318072499 | 4.90E-03 | 1.97E-02 |
| FOXJ3_HUMAN\|Q9UPW0_S223 | 1.317919375 | 9.22E-03 | 4.26E-02 |
| BAIP2_HUMAN\|Q9UQB8_S454 | 1.31779816 | 7.22E-03 | 3.51E-02 |
| EHBP1_HUMAN\|Q8NDI1_S428_S436 | 1.317661124 | 2.19E-03 | 1.75E-02 |
| K2C80_HUMAN\|Q6KB66_S451 | 1.317603213 | 1.36E-02 | 4.42E-02 |
| NFIB_HUMAN\|O00712_S295 | 1.317366326 | 4.50E-03 | 4.32E-02 |

| | | | |
|---|---|---|---|
| SMCA1_HUMAN\|P28370_S119 | 1.317332729 | 1.26E-03 | 6.63E-03 |
| MAST2_HUMAN\|Q6P0Q8_S148 | 1.317227073 | 8.37E-03 | 3.96E-02 |
| AAPK1_HUMAN\|Q13131_S508 | 1.317022592 | 8.95E-04 | 1.44E-02 |
| MTCL1_HUMAN\|Q9Y4B5_S776 | 1.316992833 | 3.76E-03 | 2.52E-02 |
| GRDN_HUMAN\|Q3V6T2_T1559 | 1.316941056 | 7.08E-03 | 2.60E-02 |
| TMF1_HUMAN\|P82094_S338_S344 | 1.316940647 | 5.21E-03 | 4.19E-02 |
| KI21A_HUMAN\|Q7Z4S6_S1229 | 1.316815658 | 4.53E-03 | 3.02E-02 |
| SSH2_HUMAN\|Q76I76_S38 | 1.316590177 | 1.07E-02 | 3.66E-02 |
| ATX2L_HUMAN\|Q8WWM7_S339 | 1.316454834 | 8.73E-04 | 1.16E-02 |
| ACAP2_HUMAN\|Q15057_T382 | 1.316432645 | 1.23E-03 | 7.16E-03 |
| SRSF1_HUMAN\|Q07955_S234_S238 | 1.316242649 | 6.49E-03 | 2.83E-02 |
| MACF1_HUMAN\|Q9UPN3_S3 | 1.316120361 | 3.94E-03 | 1.87E-02 |
| ROA3_HUMAN\|P51991_S356 | 1.316064101 | 6.34E-03 | 4.70E-02 |
| XRCC4_HUMAN\|Q13426_S328 | 1.315732388 | 4.39E-03 | 1.81E-02 |
| PI4KB_HUMAN\|Q9UBF8-2_T292 | 1.315517895 | 2.36E-03 | 1.55E-02 |
| CIC_HUMAN\|Q96RK0_S904 | 1.315287133 | 8.88E-03 | 4.12E-02 |
| DC1L1_HUMAN\|Q9Y6G9_S510 | 1.314917511 | 1.04E-03 | 1.31E-02 |
| YTDC2_HUMAN\|Q9H6S0_S1090 | 1.3148889 | 1.43E-03 | 1.22E-02 |
| BORG2_HUMAN\|Q9UKI2_S89 | 1.314666568 | 3.71E-03 | 2.49E-02 |
| NELFE_HUMAN\|P18615_S181 | 1.314612833 | 2.89E-03 | 1.79E-02 |
| HDAC2_HUMAN\|Q92769_S422_S424 | 1.314532765 | 8.00E-04 | 6.74E-03 |
| UBE3A_HUMAN\|Q05086_S218 | 1.314474484 | 3.07E-03 | 1.89E-02 |
| HDAC4_HUMAN\|P56524_S266 | 1.314350937 | 8.92E-03 | 4.14E-02 |
| PKHG3_HUMAN\|A1L390_S638_S643 | 1.314289399 | 1.45E-03 | 8.20E-03 |
| NFKB1_HUMAN\|P19838_S907 | 1.314191614 | 1.27E-03 | 1.53E-02 |
| DEK_HUMAN\|P35659_S32 | 1.3139986 | 5.75E-03 | 2.56E-02 |
| ACINU_HUMAN\|Q9UKV3_T563 | 1.313994812 | 2.28E-03 | 1.07E-02 |
| RNF31_HUMAN\|Q96EP0_S466 | 1.313760608 | 4.04E-03 | 2.66E-02 |
| LMNB1_HUMAN\|P20700_T19 | 1.313749173 | 6.02E-03 | 2.29E-02 |
| PSA5_HUMAN\|P28066_T55 | 1.313744081 | 5.36E-03 | 4.16E-02 |
| TANC2_HUMAN\|Q9HCD6_S1579 | 1.313734387 | 1.45E-03 | 1.23E-02 |

| | | | |
|---|---|---|---|
| BD1L1_HUMAN\|Q8NFC6_S484_Y480 | 1.313700925 | 1.12E-02 | 4.92E-02 |
| TWF1_HUMAN\|Q12792_T348 | 1.313596534 | 1.67E-03 | 1.19E-02 |
| TAB2_HUMAN\|Q9NYJ8_S450 | 1.313529005 | 3.55E-03 | 3.66E-02 |
| C170B_HUMAN\|Q9Y4F5_S1548 | 1.313321845 | 2.39E-03 | 2.73E-02 |
| MILK1_HUMAN\|Q8N3F8_S486 | 1.31293505 | 5.29E-03 | 4.12E-02 |
| IASPP_HUMAN\|Q8WUF5_S74 | 1.312868427 | 2.52E-03 | 2.73E-02 |
| IWS1_HUMAN\|Q96ST2_S511 | 1.312855963 | 7.53E-03 | 4.17E-02 |
| DC1L1_HUMAN\|Q9Y6G9_S510_S516 | 1.312847324 | 8.89E-03 | 4.87E-02 |
| SH23A_HUMAN\|Q9BRG2_S180 | 1.312708367 | 2.54E-03 | 1.93E-02 |
| CTU2_HUMAN\|Q2VPK5_S419 | 1.312691399 | 2.90E-03 | 2.70E-02 |
| PTN12_HUMAN\|Q05209_S571_T587 | 1.312556091 | 5.64E-03 | 2.51E-02 |
| MACF1_HUMAN\|Q9UPN3_S4 | 1.312493637 | 1.60E-03 | 8.06E-03 |
| NFKB1_HUMAN\|P19838_S907 | 1.31192081 | 1.34E-03 | 1.23E-02 |
| SRRM2_HUMAN\|Q9UQ35_S1404 | 1.311743234 | 7.07E-03 | 3.99E-02 |
| LUZP1_HUMAN\|Q86V48_S575 | 1.311667198 | 2.88E-03 | 2.12E-02 |
| FA83H_HUMAN\|Q6ZRV2_S925 | 1.311644254 | 1.74E-03 | 1.23E-02 |
| MDHC_HUMAN\|P40925_S241 | 1.311510094 | 1.23E-02 | 4.77E-02 |
| BAD_HUMAN\|Q92934_Y76 | 1.311419734 | 5.50E-03 | 2.13E-02 |
| UFL1_HUMAN\|O94874_S458 | 1.311380802 | 1.10E-03 | 2.97E-02 |
| I2BP2_HUMAN\|Q7Z5L9_S460 | 1.311319218 | 2.94E-04 | 1.35E-02 |
| RRP8_HUMAN\|O43159_S104_S106 | 1.310853666 | 5.49E-03 | 2.87E-02 |
| TFE3_HUMAN\|P19532_S568_S570 | 1.310830267 | 6.58E-03 | 2.45E-02 |
| AHNK2_HUMAN\|Q8IVF2_S294_S297 | 1.310713767 | 3.06E-03 | 2.79E-02 |
| LIPB1_HUMAN\|Q86W92_S601 | 1.310572822 | 5.42E-03 | 2.85E-02 |
| CCDC9_HUMAN\|Q9Y3X0_T389 | 1.310519179 | 2.23E-03 | 2.72E-02 |
| NFIB_HUMAN\|O00712_S295 | 1.310419254 | 5.23E-03 | 2.78E-02 |
| SRSF9_HUMAN\|Q13242_S189 | 1.310261488 | 2.47E-03 | 1.60E-02 |
| CLAP2_HUMAN\|O75122_S455 | 1.310224324 | 8.54E-03 | 3.54E-02 |
| ZN185_HUMAN\|O15231_S521 | 1.31013872 | 8.72E-03 | 4.69E-02 |
| MILK1_HUMAN\|Q8N3F8_S578 | 1.3100068 | 4.46E-03 | 3.65E-02 |
| SON_HUMAN\|P18583_S94 | 1.309615007 | 3.36E-03 | 3.74E-02 |

| | | | |
|---|---|---|---|
| NOP56_HUMAN\|O00567_S570 | 1.309417849 | 1.03E-03 | 1.58E-02 |
| CYTSA_HUMAN\|Q69YQ0_S832 | 1.309312901 | 1.41E-03 | 1.64E-02 |
| NIPA_HUMAN\|Q86WB0_S344 | 1.309140449 | 3.41E-03 | 3.77E-02 |
| UH1BL_HUMAN\|A0JNW5_S953 | 1.308867511 | 4.07E-03 | 2.30E-02 |
| NUP50_HUMAN\|Q9UKX7_S221 | 1.308690758 | 6.33E-03 | 2.78E-02 |
| SRRM1_HUMAN\|Q8IYB3_S653 | 1.308545476 | 6.49E-03 | 2.83E-02 |
| IF4G3_HUMAN\|O43432_S495 | 1.308501722 | 1.00E-02 | 4.53E-02 |
| ASPP2_HUMAN\|Q13625_S558 | 1.308439112 | 5.31E-03 | 2.40E-02 |
| TKT_HUMAN\|P29401_S295 | 1.308200204 | 3.81E-03 | 3.28E-02 |
| PCM1_HUMAN\|Q15154_S65 | 1.308167029 | 5.67E-04 | 1.02E-02 |
| STXB5_HUMAN\|Q5T5C0-2_S712 | 1.308069473 | 3.89E-03 | 3.96E-02 |
| CIRBP_HUMAN\|Q14011_S148 | 1.307897954 | 9.24E-03 | 4.26E-02 |
| NAV1_HUMAN\|Q8NEY1_S1000 | 1.307830845 | 7.42E-03 | 2.70E-02 |
| SPD2A_HUMAN\|Q5TCZ1_S1038 | 1.307658914 | 9.16E-04 | 1.20E-02 |
| ZN185_HUMAN\|O15231_S153_S155 | 1.307567415 | 3.97E-03 | 3.39E-02 |
| ZCH18_HUMAN\|Q86VM9_S67 | 1.307277685 | 1.05E-02 | 4.20E-02 |
| SMG9_HUMAN\|Q9H0W8_S53 | 1.307009975 | 4.91E-03 | 3.91E-02 |
| NIBL1_HUMAN\|Q96TA1_S665 | 1.306849963 | 5.51E-03 | 2.48E-02 |
| SRRM2_HUMAN\|Q9UQ35_S1621 | 1.306846633 | 6.32E-03 | 3.18E-02 |
| 2A5E_HUMAN\|Q16537_S33_S34 | 1.306784146 | 3.48E-03 | 2.39E-02 |
| GGYF2_HUMAN\|Q6Y7W6_S26 | 1.306731482 | 3.37E-03 | 2.40E-02 |
| GSK3A_HUMAN\|P49840_S282_Y279 | 1.306565671 | 1.01E-02 | 4.56E-02 |
| SIPA1_HUMAN\|Q96FS4_S55 | 1.306523807 | 6.16E-03 | 2.32E-02 |
| ELF1_HUMAN\|P32519_S432 | 1.306343661 | 9.13E-04 | 5.12E-03 |
| SMG9_HUMAN\|Q9H0W8_T56 | 1.305919009 | 1.07E-02 | 4.78E-02 |
| SMG9_HUMAN\|Q9H0W8_S53 | 1.305779696 | 5.04E-03 | 4.08E-02 |
| FA83B_HUMAN\|Q5T0W9_S802 | 1.305754276 | 1.33E-02 | 4.34E-02 |
| TBCD5_HUMAN\|Q92609_S541 | 1.305655363 | 6.55E-04 | 1.12E-02 |
| NHS_HUMAN\|Q6T4R5_S739 | 1.305140565 | 1.56E-02 | 4.92E-02 |
| CCD86_HUMAN\|Q9H6F5_S91 | 1.305133888 | 4.53E-03 | 2.50E-02 |
| NAA10_HUMAN\|P41227_S228_S235 | 1.304979645 | 1.61E-03 | 1.77E-02 |

| | | | |
|---|---|---|---|
| AHNK_HUMAN\|Q09666_S5752 | 1.304928942 | 3.75E-03 | 2.17E-02 |
| RBM10_HUMAN\|P98175_S60_S61_S65 | 1.304856397 | 1.74E-03 | 2.11E-02 |
| PSB5_HUMAN\|P28074_S261 | 1.304770926 | 3.27E-03 | 2.28E-02 |
| CAAP1_HUMAN\|Q9H8G2_S93 | 1.304733696 | 2.40E-03 | 1.25E-02 |
| SRRM2_HUMAN\|Q9UQ35_S2272 | 1.304659213 | 2.94E-03 | 1.48E-02 |
| PHLP_HUMAN\|Q13371_S293_S296 | 1.304576581 | 1.15E-03 | 1.60E-02 |
| SPTB2_HUMAN\|Q01082_S2165_S2169 | 1.304282448 | 2.91E-03 | 3.30E-02 |
| IF2B3_HUMAN\|O00425_S184 | 1.304188307 | 1.21E-03 | 1.09E-02 |
| MTCL1_HUMAN\|Q9Y4B5_S776 | 1.304183518 | 4.95E-03 | 3.94E-02 |
| NMD3_HUMAN\|Q96D46_S468_T470 | 1.304129407 | 1.99E-03 | 4.50E-02 |
| TENS4_HUMAN\|Q8IZW8_S416 | 1.304011841 | 3.20E-03 | 2.89E-02 |
| EHBP1_HUMAN\|Q8NDI1_S428_S436 | 1.304002905 | 3.01E-03 | 1.34E-02 |
| ZCCHV_HUMAN\|Q7Z2W4_S378 | 1.303417149 | 1.69E-03 | 1.81E-02 |
| M3K7_HUMAN\|O43318_S439 | 1.303361368 | 1.08E-02 | 4.30E-02 |
| IF4G2_HUMAN\|P78344_S17 | 1.3033304 | 9.70E-04 | 7.76E-03 |
| REPS1_HUMAN\|Q96D71_S162_S166_T173 | 1.303298098 | 5.97E-03 | 4.59E-02 |
| DKC1_HUMAN\|O60832_S485_S494 | 1.303049012 | 5.32E-03 | 4.23E-02 |
| MFAP1_HUMAN\|P55081_S80 | 1.302924646 | 5.32E-03 | 4.23E-02 |
| K1671_HUMAN\|Q9BY89_S1019 | 1.302729558 | 1.32E-03 | 1.58E-02 |
| IRS2_HUMAN\|Q9Y4H2_T1202 | 1.302624396 | 6.24E-04 | 1.12E-02 |
| ZN185_HUMAN\|O15231_S152 | 1.302533717 | 7.39E-03 | 3.16E-02 |
| SRRM2_HUMAN\|Q9UQ35_S1014 | 1.301913106 | 3.36E-03 | 3.31E-02 |
| MVD1_HUMAN\|P53602_S96 | 1.30181897 | 3.66E-03 | 2.13E-02 |
| NFKB1_HUMAN\|P19838_S907 | 1.301579628 | 1.75E-03 | 1.41E-02 |
| BRE1A_HUMAN\|Q5VTR2_S136 | 1.301410671 | 3.46E-03 | 1.68E-02 |
| VIP2_HUMAN\|O43314_S1006_S1016 | 1.301375568 | 1.45E-03 | 2.05E-02 |
| SLK_HUMAN\|Q9H2G2_S372 | 1.301206317 | 1.45E-03 | 1.61E-02 |
| XRCC6_HUMAN\|P12956_T455 | 1.300890598 | 4.65E-03 | 3.84E-02 |
| MAST4_HUMAN\|O15021_S1387_S1394 | 1.300823376 | 8.18E-03 | 2.94E-02 |
| DDX20_HUMAN\|Q9UHI6_S677_Y673 | 1.300492351 | 1.45E-02 | 4.65E-02 |
| KLC1_HUMAN\|Q07866-4_S600 | 1.30044967 | 3.67E-03 | 3.20E-02 |

| | | | |
|---|---|---|---|
| HAP28_HUMAN\|Q13442_S60 | 1.300442412 | 1.30E-02 | 4.26E-02 |
| K1671_HUMAN\|Q9BY89_S1019 | 1.300235817 | 1.41E-03 | 1.89E-02 |
| PKHG3_HUMAN\|A1L390_S1081 | 1.300166697 | 3.61E-03 | 1.55E-02 |
| DKC1_HUMAN\|O60832_T496 | 1.30014692 | 2.62E-03 | 1.67E-02 |
| IP3KB_HUMAN\|P27987_S43 | 1.299893913 | 1.04E-02 | 4.17E-02 |
| ANM3_HUMAN\|O60678_S25_S27 | 1.299884092 | 5.44E-03 | 2.85E-02 |
| NAV1_HUMAN\|Q8NEY1_S1253_T1245 | 1.299508568 | 1.13E-02 | 4.45E-02 |
| MY18A_HUMAN\|Q92614_S2041 | 1.299421452 | 1.29E-02 | 4.94E-02 |
| MILK1_HUMAN\|Q8N3F8_S640 | 1.299151433 | 3.22E-03 | 1.96E-02 |
| SVIL_HUMAN\|O95425_S63 | 1.299138197 | 3.04E-03 | 3.12E-02 |
| VIP2_HUMAN\|O43314_S1006_S1016 | 1.299135386 | 1.54E-03 | 2.07E-02 |
| SIR2_HUMAN\|Q8IXJ6_S23_S25 | 1.298508055 | 1.56E-02 | 4.92E-02 |
| NAV1_HUMAN\|Q8NEY1_S1000 | 1.298288199 | 9.00E-03 | 4.92E-02 |
| NIPA_HUMAN\|Q86WB0_S321 | 1.29822683 | 3.05E-03 | 3.42E-02 |
| PTN13_HUMAN\|Q12923_S1630 | 1.297610825 | 4.69E-03 | 2.57E-02 |
| AHNK_HUMAN\|Q09666_S210 | 1.297482865 | 4.62E-03 | 4.64E-02 |
| ACINU_HUMAN\|Q9UKV3_T563 | 1.297421677 | 3.39E-03 | 3.76E-02 |
| LIMD1_HUMAN\|Q9UGP4_S277 | 1.297021034 | 9.87E-03 | 3.42E-02 |
| MAP1B_HUMAN\|P46821_S1785 | 1.296952617 | 2.85E-03 | 2.04E-02 |
| RIPR1_HUMAN\|Q6ZS17_S351 | 1.296936071 | 4.69E-03 | 2.97E-02 |
| BICD2_HUMAN\|Q8TD16_S615 | 1.296872147 | 1.17E-02 | 3.91E-02 |
| ES8L1_HUMAN\|Q8TE68_S182_T187 | 1.296321625 | 4.76E-03 | 2.59E-02 |
| PELP1_HUMAN\|Q8IZL8_S481 | 1.29587458 | 1.03E-02 | 4.63E-02 |
| TBC15_HUMAN\|Q8TC07_T677 | 1.295055708 | 1.87E-03 | 1.30E-02 |
| PPR18_HUMAN\|Q6NYC8_S490 | 1.294858095 | 5.43E-03 | 2.12E-02 |
| JUNB_HUMAN\|P17275_T255 | 1.294502522 | 3.52E-03 | 3.09E-02 |
| SWAHC_HUMAN\|Q53LP3_S212 | 1.294402753 | 4.85E-03 | 3.98E-02 |
| AFAD_HUMAN\|P55196_S1182 | 1.294115488 | 2.42E-03 | 2.74E-02 |
| NIBL1_HUMAN\|Q96TA1_S665 | 1.294055847 | 7.17E-03 | 4.21E-02 |
| LIPB1_HUMAN\|Q86W92_S582 | 1.294016489 | 5.31E-03 | 4.53E-02 |
| ZN185_HUMAN\|O15231_S152 | 1.293970138 | 8.82E-03 | 4.84E-02 |

| | | | |
|---|---|---|---|
| CAAP1_HUMAN\|Q9H8G2_S93 | 1.292926011 | 3.21E-03 | 1.41E-02 |
| RB6I2_HUMAN\|Q8IUD2_S37_S42_S45 | 1.292642157 | 3.62E-03 | 3.26E-02 |
| SMC1A_HUMAN\|Q14683_S957 | 1.292429174 | 8.57E-03 | 3.55E-02 |
| TBC9B_HUMAN\|Q66K14_S1241 | 1.29232132 | 3.89E-03 | 2.58E-02 |
| ZFP91_HUMAN\|Q96JP5_S83 | 1.292246182 | 6.11E-03 | 4.65E-02 |
| ZN185_HUMAN\|O15231-2_S63 | 1.292064591 | 4.62E-03 | 3.83E-02 |
| LUZP1_HUMAN\|Q86V48_S575 | 1.292034626 | 4.56E-03 | 4.34E-02 |
| PKHA2_HUMAN\|Q9HB19_S184 | 1.291548912 | 5.26E-03 | 4.50E-02 |
| COHA1_HUMAN\|Q9UMD9_S172 | 1.29148598 | 5.01E-03 | 4.06E-02 |
| SFR19_HUMAN\|Q9H7N4_S965 | 1.291292104 | 8.66E-03 | 4.06E-02 |
| MATR3_HUMAN\|P43243_S598 | 1.291175414 | 2.01E-03 | 2.33E-02 |
| DDB2_HUMAN\|Q92466_S24_S26 | 1.291134325 | 1.67E-03 | 8.32E-03 |
| ES8L1_HUMAN\|Q8TE68_S182 | 1.291114696 | 9.93E-03 | 4.01E-02 |
| ACAP2_HUMAN\|Q15057_T382 | 1.291102271 | 2.32E-03 | 2.31E-02 |
| CRK_HUMAN\|P46108_S304 | 1.291096568 | 4.29E-03 | 4.40E-02 |
| HDAC2_HUMAN\|Q92769_S424 | 1.290902827 | 8.67E-03 | 4.06E-02 |
| IF2B3_HUMAN\|O00425_S180_S184 | 1.290840072 | 1.75E-03 | 1.24E-02 |
| MEPCE_HUMAN\|Q7L2J0_S60 | 1.290698894 | 8.55E-03 | 4.03E-02 |
| XRCC6_HUMAN\|P12956_T455 | 1.290521451 | 5.87E-03 | 3.00E-02 |
| BCLF1_HUMAN\|Q9NYF8_S222 | 1.290505815 | 5.93E-03 | 4.88E-02 |
| IRS2_HUMAN\|Q9Y4H2_S560 | 1.290439618 | 5.99E-03 | 3.69E-02 |
| M3K3_HUMAN\|Q99759_S337_S340 | 1.2904331 | 1.11E-02 | 4.38E-02 |
| PHF8_HUMAN\|Q9UPP1_S804 | 1.290317682 | 1.16E-02 | 3.88E-02 |
| OTUL_HUMAN\|Q96BN8_S351 | 1.290262577 | 2.19E-03 | 2.53E-02 |
| UBP47_HUMAN\|Q96K76_S933 | 1.290114823 | 5.92E-03 | 4.88E-02 |
| PLEC_HUMAN\|Q15149_S4670 | 1.289445629 | 1.59E-02 | 4.99E-02 |
| ARI1A_HUMAN\|O14497_S233 | 1.289213631 | 6.92E-03 | 4.09E-02 |
| KPCD_HUMAN\|Q05655_S304 | 1.289033061 | 4.90E-03 | 4.58E-02 |
| ZN185_HUMAN\|O15231_S153_T158 | 1.288991418 | 5.67E-03 | 4.34E-02 |
| ATX2L_HUMAN\|Q8WWM7_S680_S684 | 1.288809466 | 4.64E-03 | 3.84E-02 |
| CP131_HUMAN\|Q9UPN4_S78 | 1.288629972 | 9.31E-03 | 4.87E-02 |

| | | | |
|---|---|---|---|
| PYR1_HUMAN\|P27708_T1037 | 1.288427371 | 1.16E-02 | 3.88E-02 |
| OAS1_HUMAN\|P00973_S50 | 1.288142845 | 6.74E-03 | 2.50E-02 |
| P66B_HUMAN\|Q8WXI9_S122_S129_T120 | 1.288099593 | 4.73E-03 | 4.45E-02 |
| OTUL_HUMAN\|Q96BN8_S351 | 1.288094235 | 2.32E-03 | 2.31E-02 |
| APC_HUMAN\|P25054_S2242_S2244 | 1.287939373 | 5.00E-03 | 2.28E-02 |
| PSA5_HUMAN\|P28066_T55 | 1.287808593 | 9.25E-03 | 4.86E-02 |
| CARL1_HUMAN\|Q5VZK9_S1288 | 1.287642587 | 3.76E-03 | 3.59E-02 |
| RBM39_HUMAN\|Q14498_S97 | 1.287635683 | 1.79E-03 | 1.26E-02 |
| ANS1A_HUMAN\|Q92625_S649 | 1.287423441 | 4.14E-03 | 3.47E-02 |
| MON1A_HUMAN\|Q86VX9_Y57 | 1.287196923 | 1.04E-02 | 4.65E-02 |
| MON2_HUMAN\|Q7Z3U7_S1214 | 1.287026446 | 1.21E-02 | 4.70E-02 |
| MILK1_HUMAN\|Q8N3F8_S486 | 1.287007789 | 8.97E-03 | 4.16E-02 |
| TANC2_HUMAN\|Q9HCD6_S1579 | 1.286949082 | 2.83E-03 | 3.09E-02 |
| BCL9L_HUMAN\|Q86UU0_S118_S121 | 1.286868403 | 4.12E-03 | 3.84E-02 |
| CNKR1_HUMAN\|Q969H4_S314 | 1.286654199 | 1.24E-02 | 4.79E-02 |
| ELF1_HUMAN\|P32519_S168 | 1.286537655 | 6.10E-03 | 3.58E-02 |
| DCP1A_HUMAN\|Q9NPI6_S525 | 1.285927846 | 2.51E-03 | 2.73E-02 |
| ELF1_HUMAN\|P32519_S163 | 1.285892207 | 4.59E-03 | 2.53E-02 |
| PKP3_HUMAN\|Q9Y446_S313 | 1.285769114 | 2.93E-03 | 2.72E-02 |
| BD1L1_HUMAN\|Q8NFC6_S2964 | 1.285662112 | 5.46E-03 | 2.86E-02 |
| UHRF2_HUMAN\|Q96PU4_S667 | 1.28564735 | 4.96E-03 | 2.67E-02 |
| TB182_HUMAN\|Q9C0C2_S1666 | 1.285611913 | 4.40E-03 | 4.32E-02 |
| BRD2_HUMAN\|P25440_S298 | 1.28522016 | 3.76E-03 | 3.25E-02 |
| KS6A1_HUMAN\|Q15418_S363 | 1.285168715 | 4.77E-03 | 4.21E-02 |
| CAAP1_HUMAN\|Q9H8G2_S89 | 1.284621547 | 3.58E-03 | 1.73E-02 |
| C170B_HUMAN\|Q9Y4F5_S1548 | 1.283831553 | 4.86E-03 | 4.80E-02 |
| FOXO3_HUMAN\|O43524_S253 | 1.283150967 | 5.61E-03 | 2.91E-02 |
| PRP4B_HUMAN\|Q13523_S257 | 1.282852432 | 6.09E-03 | 3.09E-02 |
| SRPK2_HUMAN\|P78362_S497 | 1.282758108 | 1.56E-02 | 4.91E-02 |
| SRRM2_HUMAN\|Q9UQ35_S1219 | 1.282467677 | 6.97E-03 | 3.42E-02 |
| TCF20_HUMAN\|Q9UGU0_T1671 | 1.282407692 | 2.99E-03 | 2.11E-02 |

| | | | |
|---|---|---|---|
| HNRL2_HUMAN\|Q1KMD3_S161_S168 | 1.282259439 | 4.28E-03 | 2.40E-02 |
| CBX3_HUMAN\|Q13185_S95 | 1.282196912 | 2.83E-03 | 2.96E-02 |
| SRRM2_HUMAN\|Q9UQ35_S2692_S2694 | 1.282045745 | 4.74E-03 | 2.58E-02 |
| TBCD5_HUMAN\|Q92609_S541 | 1.281989856 | 1.30E-03 | 3.32E-02 |
| UFL1_HUMAN\|O94874_S458 | 1.281793883 | 2.42E-03 | 2.37E-02 |
| PCM1_HUMAN\|Q15154_S65 | 1.281759018 | 1.24E-03 | 7.22E-03 |
| TP53B_HUMAN\|Q12888_S831_S834 | 1.28173276 | 9.81E-03 | 4.48E-02 |
| IF4G2_HUMAN\|P78344_T508 | 1.281724585 | 4.49E-03 | 3.76E-02 |
| LIPB1_HUMAN\|Q86W92_T602 | 1.281083084 | 2.27E-03 | 4.60E-02 |
| ACINU_HUMAN\|Q9UKV3_T563 | 1.281083043 | 5.01E-03 | 4.69E-02 |
| NAA10_HUMAN\|P41227_S228_S235 | 1.281038575 | 2.99E-03 | 2.75E-02 |
| UBP8_HUMAN\|P40818_T577 | 1.281002856 | 6.49E-03 | 3.24E-02 |
| CDK13_HUMAN\|Q14004_T1246 | 1.280879897 | 1.29E-03 | 1.55E-02 |
| DNPH1_HUMAN\|O43598_S169 | 1.280582714 | 1.33E-02 | 4.33E-02 |
| ZC3H8_HUMAN\|Q8N5P1_S59 | 1.280440151 | 7.27E-03 | 4.24E-02 |
| MATR3_HUMAN\|P43243_S598 | 1.280384706 | 2.68E-03 | 1.95E-02 |
| MMTA2_HUMAN\|Q9BU76_S216_S217_S220 | 1.28037389 | 3.12E-03 | 2.19E-02 |
| MON1A_HUMAN\|Q86VX9_S56 | 1.279669659 | 1.19E-02 | 4.63E-02 |
| TBA1C_HUMAN\|Q9BQE3_S439 | 1.279646852 | 4.26E-03 | 2.76E-02 |
| K1522_HUMAN\|Q9P206_S327 | 1.279544482 | 8.28E-03 | 3.94E-02 |
| UBP47_HUMAN\|Q96K76_S1013 | 1.279508564 | 6.60E-03 | 2.87E-02 |
| TENS4_HUMAN\|Q8IZW8_S350 | 1.279463044 | 1.57E-02 | 4.94E-02 |
| SHOT1_HUMAN\|A0MZ66_S494 | 1.278691033 | 8.14E-03 | 4.60E-02 |
| VIP2_HUMAN\|O43314_S1006_S1016 | 1.278363311 | 2.71E-03 | 2.98E-02 |
| SI1L3_HUMAN\|O60292_S172 | 1.278359527 | 1.87E-03 | 1.96E-02 |
| KLC3_HUMAN\|Q6P597_S502 | 1.278087892 | 1.42E-03 | 1.89E-02 |
| SPT5H_HUMAN\|O00267_S666 | 1.278064707 | 7.19E-03 | 3.50E-02 |
| ACAP2_HUMAN\|Q15057_S384 | 1.277584734 | 4.95E-03 | 4.36E-02 |
| SPE39_HUMAN\|Q9H9C1_S121 | 1.277083239 | 4.71E-03 | 4.19E-02 |
| UFL1_HUMAN\|O94874_S462 | 1.276978072 | 9.89E-03 | 4.49E-02 |
| HEAT6_HUMAN\|Q6AI08_S643 | 1.276801069 | 5.55E-03 | 2.89E-02 |

| | | | |
|---|---|---|---|
| DKC1_HUMAN\|O60832_T496 | 1.276754949 | 4.67E-03 | 2.96E-02 |
| SENP3_HUMAN\|Q9H4L4_S169_S181 | 1.276278858 | 1.10E-02 | 4.86E-02 |
| VIP2_HUMAN\|O43314_S1006_S1016 | 1.276162742 | 2.87E-03 | 2.05E-02 |
| LMNA_HUMAN\|P02545_S22 | 1.275588642 | 7.92E-03 | 4.34E-02 |
| TKT_HUMAN\|P29401_S295 | 1.275403092 | 7.91E-03 | 3.79E-02 |
| PKP4_HUMAN\|Q99569_S314 | 1.275051938 | 5.64E-03 | 2.17E-02 |
| SVIL_HUMAN\|O95425_S63 | 1.274560975 | 5.42E-03 | 2.45E-02 |
| EXOC1_HUMAN\|Q9NV70_T468 | 1.274371018 | 3.03E-03 | 2.87E-02 |
| ZC3HD_HUMAN\|Q5T200_S242 | 1.274231057 | 8.16E-03 | 3.89E-02 |
| NMD3_HUMAN\|Q96D46_S468_T470 | 1.273879865 | 4.19E-03 | 3.49E-02 |
| JUNB_HUMAN\|P17275_T255 | 1.27378848 | 5.78E-03 | 4.84E-02 |
| NOP56_HUMAN\|O00567_S570 | 1.273668377 | 2.77E-03 | 3.04E-02 |
| EXOC1_HUMAN\|Q9NV70_T471 | 1.27345799 | 6.05E-03 | 4.64E-02 |
| TCF20_HUMAN\|Q9UGU0_S574_S583 | 1.27284745 | 9.93E-03 | 4.51E-02 |
| SETLP_HUMAN\|P0DME0_S171 | 1.272230067 | 5.40E-03 | 4.17E-02 |
| RIPR1_HUMAN\|Q6ZS17_S351 | 1.272103501 | 8.29E-03 | 3.46E-02 |
| KPCD_HUMAN\|Q05655_S307 | 1.27210001 | 4.23E-03 | 3.60E-02 |
| ARH40_HUMAN\|Q8TER5_T371 | 1.271737549 | 5.31E-03 | 2.82E-02 |
| IF2B3_HUMAN\|O00425_S179_S180 | 1.271532513 | 1.93E-03 | 1.33E-02 |
| STRP1_HUMAN\|Q5VSL9_S339 | 1.271262488 | 9.86E-03 | 4.49E-02 |
| MK06_HUMAN\|Q16659_S386_T389 | 1.271117365 | 3.30E-03 | 3.28E-02 |
| RPB1_HUMAN\|P24928_S1892 | 1.270956772 | 8.03E-03 | 3.37E-02 |
| I2BPL_HUMAN\|Q9H1B7_S657_S662 | 1.270251987 | 1.57E-02 | 4.94E-02 |
| SPD2A_HUMAN\|Q5TCZ1_S1038 | 1.269973759 | 2.61E-03 | 2.49E-02 |
| SMN_HUMAN\|Q16637_S28 | 1.269930899 | 6.74E-03 | 2.50E-02 |
| AFAD_HUMAN\|P55196_S1182 | 1.269565233 | 4.45E-03 | 3.00E-02 |
| CAVN3_HUMAN\|Q969G5_S165_S166 | 1.269469884 | 5.88E-03 | 3.00E-02 |
| M3K20_HUMAN\|Q9NYL2_S727 | 1.269298716 | 5.46E-03 | 2.86E-02 |
| EDC3_HUMAN\|Q96F86_S131 | 1.269158646 | 6.62E-03 | 3.29E-02 |
| LIPB1_HUMAN\|Q86W92_S794 | 1.269077603 | 2.30E-03 | 2.65E-02 |
| FIP1_HUMAN\|Q6UN15_S304 | 1.268770296 | 1.22E-03 | 9.34E-03 |

| | | | |
|---|---|---|---|
| CSN3_HUMAN\|Q9UNS2_S407 | 1.267776103 | 2.11E-03 | 1.63E-02 |
| LIMA1_HUMAN\|Q9UHB6_S374 | 1.267313551 | 3.33E-03 | 2.01E-02 |
| PHLP_HUMAN\|Q13371_S293_S296 | 1.266078911 | 3.34E-03 | 3.59E-02 |
| CARL1_HUMAN\|Q5VZK9_S1288 | 1.265974669 | 6.35E-03 | 4.70E-02 |
| NOP56_HUMAN\|O00567_S570 | 1.265549234 | 3.46E-03 | 3.07E-02 |
| PININ_HUMAN\|Q9H307_T657 | 1.265353221 | 2.65E-03 | 1.68E-02 |
| K1671_HUMAN\|Q9BY89_S1019 | 1.265153584 | 3.65E-03 | 3.92E-02 |
| YBOX3_HUMAN\|P16989_S370 | 1.264999619 | 1.18E-02 | 4.61E-02 |
| HNRL2_HUMAN\|Q1KMD3_S161_S168 | 1.264631272 | 6.50E-03 | 4.77E-02 |
| CARL1_HUMAN\|Q5VZK9_S1288 | 1.264229434 | 6.62E-03 | 3.81E-02 |
| ELF1_HUMAN\|P32519_S432 | 1.263817676 | 3.07E-03 | 2.23E-02 |
| VIP2_HUMAN\|O43314_S1006_S1016 | 1.263124642 | 4.08E-03 | 2.82E-02 |
| RTKN_HUMAN\|Q9BST9_S529_T518 | 1.263078985 | 6.31E-03 | 4.69E-02 |
| RHG32_HUMAN\|A7KAX9_S682 | 1.263017922 | 1.15E-02 | 4.52E-02 |
| NFKB1_HUMAN\|P19838_S907 | 1.261963312 | 4.85E-03 | 1.95E-02 |
| RPB1_HUMAN\|P24928_S1892 | 1.261244782 | 1.00E-02 | 3.46E-02 |
| BRD2_HUMAN\|P25440_S298 | 1.260591476 | 6.88E-03 | 3.39E-02 |
| BRD8_HUMAN\|Q9H0E9-2_S284 | 1.260547369 | 1.08E-02 | 3.68E-02 |
| SRS11_HUMAN\|Q05519_S414 | 1.260081195 | 5.21E-03 | 4.84E-02 |
| MICA3_HUMAN\|Q7RTP6_S1143 | 1.259434425 | 9.54E-03 | 3.33E-02 |
| EP15R_HUMAN\|Q9UBC2_S229 | 1.25938207 | 5.39E-03 | 3.43E-02 |
| STXB5_HUMAN\|Q5T5C0-2_S712 | 1.25892317 | 1.14E-02 | 4.50E-02 |
| PDLI2_HUMAN\|Q96JY6_S74 | 1.258391999 | 6.26E-03 | 3.16E-02 |
| KLC3_HUMAN\|Q6P597_S466 | 1.257892047 | 5.37E-03 | 4.16E-02 |
| AFAD_HUMAN\|P55196_S1182 | 1.257814884 | 5.96E-03 | 2.27E-02 |
| MACF1_HUMAN\|Q9UPN3_S4 | 1.257106289 | 6.60E-03 | 3.97E-02 |
| KLC3_HUMAN\|Q6P597_S466 | 1.256689885 | 5.53E-03 | 4.67E-02 |
| ARI1A_HUMAN\|O14497_S382 | 1.256348607 | 5.93E-03 | 3.67E-02 |
| I2BP2_HUMAN\|Q7Z5L9_S460 | 1.256101585 | 1.71E-03 | 3.06E-02 |
| PP12C_HUMAN\|Q9BZL4_S453 | 1.255751844 | 7.44E-03 | 3.59E-02 |
| NGDN_HUMAN\|Q8NEJ9_S143 | 1.255392249 | 3.75E-03 | 3.59E-02 |

| | | | |
|---|---|---|---|
| JHD2C_HUMAN\|Q15652_S943 | 1.255092134 | 1.12E-02 | 4.93E-02 |
| SYEP_HUMAN\|P07814_S882_S886 | 1.255013139 | 1.46E-03 | 1.08E-02 |
| SMCA1_HUMAN\|P28370_S119 | 1.253963219 | 6.61E-03 | 2.87E-02 |
| CAMP3_HUMAN\|Q9P1Y5_S553 | 1.252621983 | 5.44E-03 | 4.29E-02 |
| ACAP2_HUMAN\|Q15057_S384 | 1.252575288 | 8.90E-03 | 4.74E-02 |
| IRS2_HUMAN\|Q9Y4H2_S1203 | 1.251653039 | 8.60E-03 | 3.07E-02 |
| CDK18_HUMAN\|Q07002_S87 | 1.251647845 | 3.72E-03 | 2.49E-02 |
| TDIF2_HUMAN\|Q5QJE6_S117 | 1.250915857 | 6.89E-03 | 3.39E-02 |
| KIF1C_HUMAN\|O43896_S1092 | 1.249974659 | 8.48E-03 | 3.52E-02 |
| LIPB1_HUMAN\|Q86W92_T602 | 1.249969246 | 5.24E-03 | 2.79E-02 |
| PFKAP_HUMAN\|Q01813_S386 | 1.249150291 | 4.11E-03 | 2.32E-02 |
| CE170_HUMAN\|Q5SW79_S636 | 1.247985222 | 4.49E-03 | 4.06E-02 |
| BCLF1_HUMAN\|Q9NYF8_S531 | 1.247778431 | 4.66E-03 | 2.56E-02 |
| RPB1_HUMAN\|P24928_S1899 | 1.247677943 | 8.16E-03 | 3.41E-02 |
| KLC3_HUMAN\|Q6P597_S502 | 1.246713185 | 3.58E-03 | 1.54E-02 |
| DCAF6_HUMAN\|Q58WW2_S847_S850 | 1.246105184 | 5.91E-03 | 3.02E-02 |
| TJAP1_HUMAN\|Q5JTD0_S545 | 1.245754171 | 2.52E-03 | 2.84E-02 |
| PLCH2_HUMAN\|O75038_S562 | 1.244398245 | 9.84E-03 | 4.49E-02 |
| IRS2_HUMAN\|Q9Y4H2_S1186_T1202 | 1.243934103 | 1.59E-02 | 4.97E-02 |
| PHF14_HUMAN\|O94880_T292 | 1.24322253 | 1.56E-02 | 4.91E-02 |
| CDK18_HUMAN\|Q07002_S87 | 1.241145225 | 5.00E-03 | 3.26E-02 |
| RB15B_HUMAN\|Q8NDT2_S109 | 1.240614937 | 5.37E-03 | 2.10E-02 |
| TANC2_HUMAN\|Q9HCD6_S1579 | 1.240281252 | 9.08E-03 | 3.72E-02 |
| KS6A1_HUMAN\|Q15418_T359 | 1.240132226 | 2.47E-03 | 2.71E-02 |
| TJAP1_HUMAN\|Q5JTD0_S545 | 1.239309575 | 3.08E-03 | 2.91E-02 |
| RBM10_HUMAN\|P98175_S60_S61_S65 | 1.238803289 | 9.45E-03 | 4.35E-02 |
| SYEP_HUMAN\|P07814_S885_S886 | 1.236757067 | 5.75E-03 | 2.21E-02 |
| IF4G2_HUMAN\|P78344_S17 | 1.236650532 | 6.45E-03 | 4.75E-02 |
| TFE3_HUMAN\|P19532_S564_S568 | 1.23653633 | 7.07E-03 | 4.17E-02 |
| LIPB1_HUMAN\|Q86W92_T602 | 1.236417196 | 7.52E-03 | 4.17E-02 |
| IRS2_HUMAN\|Q9Y4H2_T1202 | 1.233536644 | 4.92E-03 | 4.63E-02 |

| | | | |
|---|---|---|---|
| ZO2_HUMAN\|Q9UDY2_S966 | 1.23331706 | 9.21E-03 | 4.25E-02 |
| IRS2_HUMAN\|Q9Y4H2_T1202 | 1.232292358 | 5.11E-03 | 2.02E-02 |
| EIF3C_HUMAN\|Q99613_S39 | 1.232182361 | 6.58E-03 | 3.28E-02 |
| MARE3_HUMAN\|Q9UPY8-2_S147 | 1.231655658 | 6.61E-03 | 3.28E-02 |
| RHG32_HUMAN\|A7KAX9_S952 | 1.231062363 | 8.92E-03 | 4.74E-02 |
| KLC3_HUMAN\|Q6P597_S502 | 1.227373119 | 6.29E-03 | 3.67E-02 |
| M3K20_HUMAN\|Q9NYL2_S667 | 1.222336258 | 5.23E-03 | 4.20E-02 |
| UBP31_HUMAN\|Q70CQ4_S1088 | 1.222025647 | 9.88E-03 | 4.49E-02 |
| AK17A_HUMAN\|Q02040_S537 | 1.221564508 | 3.73E-03 | 2.16E-02 |
| TCF20_HUMAN\|Q9UGU0_S1322 | 1.220871802 | 1.29E-02 | 4.24E-02 |
| RS3_HUMAN\|P23396_T221 | 1.218665336 | 8.79E-03 | 4.10E-02 |
| KS6A1_HUMAN\|Q15418_T359 | 1.218270548 | 4.92E-03 | 3.04E-02 |
| VIP2_HUMAN\|O43314_S1006_S1016 | 1.212710707 | 1.55E-02 | 4.89E-02 |
| SYEP_HUMAN\|P07814_S885_S886 | 1.209467159 | 1.23E-02 | 4.77E-02 |
| NIPA_HUMAN\|Q86WB0_S395 | 1.205762314 | 1.53E-02 | 4.85E-02 |
| PYRG2_HUMAN\|Q9NRF8_S568_S571_S574 | 0.85133805 | 1.58E-02 | 4.96E-02 |
| TJAP1_HUMAN\|Q5JTD0_S545 | 0.833952298 | 1.05E-02 | 3.61E-02 |
| EIF3C_HUMAN\|Q99613_S39 | 0.832019401 | 1.54E-02 | 4.86E-02 |
| SYEP_HUMAN\|P07814_S882_S886 | 0.83120637 | 7.85E-03 | 4.49E-02 |
| INADL_HUMAN\|Q8NI35_S1212_T1209 | 0.828475326 | 1.32E-02 | 4.32E-02 |
| TBC15_HUMAN\|Q8TC07_T677 | 0.825282001 | 1.59E-02 | 4.98E-02 |
| IF2P_HUMAN\|O60841_S214 | 0.824873624 | 1.55E-02 | 4.89E-02 |
| SYEP_HUMAN\|P07814_S882_S886 | 0.823971868 | 5.61E-03 | 3.35E-02 |
| KLC3_HUMAN\|Q6P597_S466 | 0.821164479 | 1.49E-02 | 4.76E-02 |
| PCBP1_HUMAN\|Q15365_S190 | 0.818967264 | 8.72E-03 | 4.82E-02 |
| DCAF6_HUMAN\|Q58WW2_S847_S850 | 0.81841074 | 1.14E-02 | 3.81E-02 |
| RENT1_HUMAN\|Q92900_S1107 | 0.818197855 | 1.13E-02 | 3.81E-02 |
| AK17A_HUMAN\|Q02040_S537 | 0.817950117 | 3.61E-03 | 1.55E-02 |
| PCBP1_HUMAN\|Q15365_S190 | 0.816728858 | 7.94E-03 | 3.35E-02 |
| LARP1_HUMAN\|Q6PKG0_S143 | 0.816326531 | 6.32E-03 | 2.77E-02 |
| M3K20_HUMAN\|Q9NYL2_S667 | 0.816211872 | 4.80E-03 | 1.94E-02 |

| | | | |
|---|---|---|---|
| IF4G2_HUMAN\|P78344_S17 | 0.813974817 | 8.08E-03 | 4.42E-02 |
| RENT1_HUMAN\|Q92900_S1107 | 0.813661261 | 9.46E-03 | 3.86E-02 |
| NIPA_HUMAN\|Q86WB0_S394 | 0.813112659 | 9.21E-03 | 4.25E-02 |
| LARP1_HUMAN\|Q6PKG0_S143 | 0.813071494 | 5.47E-03 | 2.13E-02 |
| SNW1_HUMAN\|Q13573_S224_S232 | 0.813063592 | 1.08E-02 | 3.66E-02 |
| PININ_HUMAN\|Q9H307_T657 | 0.812925226 | 7.22E-03 | 4.04E-02 |
| ATX2_HUMAN\|Q99700_T741 | 0.812624344 | 4.88E-03 | 3.20E-02 |
| ZCCHV_HUMAN\|Q7Z2W4_S459 | 0.812480583 | 9.40E-03 | 3.84E-02 |
| IF2B3_HUMAN\|O00425_S180_S184 | 0.81241655 | 8.63E-03 | 3.57E-02 |
| I2BP2_HUMAN\|Q7Z5L9_S460 | 0.812075233 | 3.73E-03 | 3.98E-02 |
| MATR3_HUMAN\|P43243_S598 | 0.810765919 | 9.16E-03 | 3.23E-02 |
| EF1D_HUMAN\|P29692_S133 | 0.810248304 | 4.23E-03 | 1.76E-02 |
| I2BP2_HUMAN\|Q7Z5L9_S460 | 0.81003791 | 3.38E-03 | 3.62E-02 |
| MK06_HUMAN\|Q16659_S386_T389 | 0.808888317 | 8.32E-03 | 2.98E-02 |
| CYTSA_HUMAN\|Q69YQ0_S832 | 0.808473908 | 9.30E-03 | 3.27E-02 |
| MILK1_HUMAN\|Q8N3F8_S391 | 0.806810523 | 6.65E-03 | 2.47E-02 |
| SH23A_HUMAN\|Q9BRG2_S125 | 0.806514942 | 1.50E-02 | 4.77E-02 |
| CACO1_HUMAN\|Q9P1Z2_S563 | 0.806483432 | 1.24E-02 | 4.08E-02 |
| CDK13_HUMAN\|Q14004_T1246 | 0.806451711 | 4.31E-03 | 4.25E-02 |
| F122B_HUMAN\|Q7Z309_S115 | 0.806430987 | 1.07E-02 | 3.65E-02 |
| PCM1_HUMAN\|Q15154_S65_S69 | 0.806296804 | 5.94E-03 | 3.67E-02 |
| DEFI6_HUMAN\|Q9H4E7_S597 | 0.806259262 | 1.02E-02 | 3.51E-02 |
| PHLP_HUMAN\|Q13371_S296 | 0.806180737 | 1.38E-02 | 4.48E-02 |
| LIPA1_HUMAN\|Q13136_S763 | 0.805543171 | 1.37E-02 | 4.44E-02 |
| TANC1_HUMAN\|Q9C0D5_S444 | 0.805383045 | 1.11E-02 | 3.74E-02 |
| EF1D_HUMAN\|P29692_S133 | 0.804707545 | 3.26E-03 | 3.53E-02 |
| PININ_HUMAN\|Q9H307_T657 | 0.804341752 | 4.98E-03 | 4.64E-02 |
| PDLI2_HUMAN\|Q96JY6_S124_S129 | 0.80379268 | 9.13E-03 | 4.82E-02 |
| PKP4_HUMAN\|Q99569_S337 | 0.803633347 | 1.35E-02 | 4.39E-02 |
| PCY1A_HUMAN\|P49585_S362 | 0.80355008 | 8.16E-03 | 3.41E-02 |
| MILK1_HUMAN\|Q8N3F8_S640 | 0.803515497 | 1.16E-02 | 4.53E-02 |

| | | | |
|---|---|---|---|
| SHRM2_HUMAN\|Q13796_S1039 | 0.80336413 | 3.50E-03 | 3.69E-02 |
| VIP2_HUMAN\|O43314_S1006_S1013_S1016 | 0.803192915 | 1.49E-02 | 4.76E-02 |
| SMN_HUMAN\|Q16637_S31_T25 | 0.803189064 | 8.59E-03 | 4.77E-02 |
| AKTS1_HUMAN\|Q96B36_S203_S211_T198 | 0.803072475 | 1.42E-02 | 4.57E-02 |
| FA83H_HUMAN\|Q6ZRV2_T894 | 0.802862159 | 5.96E-03 | 2.63E-02 |
| SRRM2_HUMAN\|Q9UQ35_S2692_S2694 | 0.802747924 | 1.13E-02 | 3.81E-02 |
| DPYL2_HUMAN\|Q16555_T514 | 0.802556208 | 7.95E-03 | 4.51E-02 |
| LARP1_HUMAN\|Q6PKG0_S143 | 0.802539988 | 3.40E-03 | 4.81E-02 |
| SHRM2_HUMAN\|Q13796_S1039 | 0.802470165 | 3.36E-03 | 3.54E-02 |
| SETLP_HUMAN\|P0DME0_S171 | 0.802233127 | 1.01E-02 | 3.50E-02 |
| ZN185_HUMAN\|O15231-2_T64 | 0.801933399 | 1.57E-02 | 4.94E-02 |
| NFRKB_HUMAN\|Q6P4R8_S1291 | 0.801837907 | 3.94E-03 | 1.87E-02 |
| IRS2_HUMAN\|Q9Y4H2_S560 | 0.801724065 | 1.53E-02 | 4.86E-02 |
| MAGI3_HUMAN\|Q5TCQ9_S724 | 0.8013317 | 8.88E-03 | 4.74E-02 |
| HNRPK_HUMAN\|P61978_S284 | 0.801098393 | 4.23E-03 | 4.14E-02 |
| MINT_HUMAN\|Q96T58_S623 | 0.800967371 | 6.54E-03 | 3.26E-02 |
| ANXA2_HUMAN\|P07355_T19 | 0.800434422 | 8.02E-03 | 2.90E-02 |
| K1671_HUMAN\|Q9BY89_S1366 | 0.800247173 | 5.06E-03 | 2.01E-02 |
| TBC15_HUMAN\|Q8TC07_T677 | 0.800214353 | 6.04E-03 | 3.72E-02 |
| AFF4_HUMAN\|Q9UHB7_S1055_S1058 | 0.800096062 | 7.11E-03 | 3.06E-02 |
| IWS1_HUMAN\|Q96ST2_S438_S440 | 0.800062121 | 9.43E-03 | 3.30E-02 |
| IWS1_HUMAN\|Q96ST2_S511_S513 | 0.799670721 | 7.88E-03 | 2.85E-02 |
| RHG35_HUMAN\|Q9NRY4_S1134 | 0.799507923 | 1.15E-02 | 3.84E-02 |
| SI1L3_HUMAN\|O60292_S172 | 0.799452977 | 4.11E-03 | 4.31E-02 |
| UBP31_HUMAN\|Q70CQ4_S1088 | 0.799266544 | 4.46E-03 | 4.30E-02 |
| RALY_HUMAN\|Q9UKM9_S288_T286 | 0.799266129 | 1.19E-02 | 4.63E-02 |
| PININ_HUMAN\|Q9H307_T657 | 0.799228118 | 3.97E-03 | 1.88E-02 |
| MILK1_HUMAN\|Q8N3F8_S391 | 0.799188896 | 4.84E-03 | 2.22E-02 |
| NIPA_HUMAN\|Q86WB0_S409 | 0.799012706 | 4.06E-03 | 3.52E-02 |
| MINT_HUMAN\|Q96T58_S623 | 0.798711193 | 5.96E-03 | 4.52E-02 |
| NUFP2_HUMAN\|Q7Z417_S113 | 0.798621208 | 8.48E-03 | 3.52E-02 |

| | | | |
|---|---|---|---|
| NIPA_HUMAN\|Q86WB0_S409 | 0.798606602 | 3.99E-03 | 3.40E-02 |
| RFIP5_HUMAN\|Q9BXF6_S296 | 0.798495215 | 9.38E-03 | 3.29E-02 |
| ZN687_HUMAN\|Q8N1G0_T432 | 0.798491404 | 5.02E-03 | 3.99E-02 |
| DCAF6_HUMAN\|Q58WW2_S847_S850 | 0.798032059 | 4.89E-03 | 4.81E-02 |
| FBXW9_HUMAN\|Q5XUX1_S18 | 0.79802865 | 1.45E-02 | 4.66E-02 |
| PSB5_HUMAN\|P28074_S261 | 0.797851363 | 1.08E-02 | 3.67E-02 |
| TDIF2_HUMAN\|Q5QJE6_S117 | 0.797747135 | 6.44E-03 | 3.90E-02 |
| NEDD1_HUMAN\|Q8NHV4_S516 | 0.797164609 | 4.02E-03 | 3.41E-02 |
| DPYL2_HUMAN\|Q16555_T509 | 0.797045972 | 5.56E-03 | 2.49E-02 |
| IRS2_HUMAN\|Q9Y4H2_T1202 | 0.797012575 | 2.63E-03 | 4.98E-02 |
| FA83H_HUMAN\|Q6ZRV2_S647 | 0.796989142 | 9.40E-03 | 4.89E-02 |
| RIOK1_HUMAN\|Q9BRS2_S22 | 0.796926471 | 3.27E-03 | 3.53E-02 |
| UFL1_HUMAN\|O94874_S458 | 0.796912686 | 4.94E-03 | 1.98E-02 |
| PCBP1_HUMAN\|Q15365_S190 | 0.796744852 | 3.32E-03 | 2.30E-02 |
| ZO2_HUMAN\|Q9UDY2_S966 | 0.796083432 | 5.05E-03 | 4.91E-02 |
| MILK1_HUMAN\|Q8N3F8_S640 | 0.795931546 | 8.77E-03 | 4.83E-02 |
| K0930_HUMAN\|Q6ICG6_S267_S279 | 0.79559356 | 1.08E-02 | 4.79E-02 |
| ACAP2_HUMAN\|Q15057_T382 | 0.795433579 | 5.46E-03 | 4.30E-02 |
| KANK1_HUMAN\|Q14678_S144 | 0.795273232 | 4.15E-03 | 2.72E-02 |
| HNRPK_HUMAN\|P61978_S284 | 0.795158039 | 3.26E-03 | 3.53E-02 |
| ATX2L_HUMAN\|Q8WWM7_S680_S684 | 0.795142552 | 9.63E-03 | 3.35E-02 |
| NELFE_HUMAN\|P18615_S353 | 0.794998556 | 5.34E-03 | 3.41E-02 |
| KLC3_HUMAN\|Q6P597_S497_S502 | 0.794800851 | 7.11E-03 | 4.18E-02 |
| PLCH2_HUMAN\|O75038_S982 | 0.794723631 | 7.82E-03 | 2.83E-02 |
| MILK1_HUMAN\|Q8N3F8_S471 | 0.794451565 | 8.89E-03 | 4.74E-02 |
| ATX2_HUMAN\|Q99700_T741 | 0.794028195 | 2.04E-03 | 2.58E-02 |
| VIP2_HUMAN\|O43314_S1006_S1013_S1016 | 0.793890428 | 1.08E-02 | 4.79E-02 |
| SRRM2_HUMAN\|Q9UQ35_S1014 | 0.793676775 | 9.02E-03 | 3.19E-02 |
| YAP1_HUMAN\|P46937_S109_T110 | 0.79334808 | 8.12E-03 | 4.59E-02 |
| PDLI2_HUMAN\|Q96JY6_S74 | 0.793319939 | 5.93E-03 | 3.67E-02 |
| FIP1_HUMAN\|Q6UN15_S304 | 0.793128957 | 1.56E-03 | 7.92E-03 |

| | | | |
|---|---|---|---|
| TWF1_HUMAN|Q12792_T348 | 0.793117287 | 6.21E-03 | 3.63E-02 |
| CLASR_HUMAN|Q8N2M8_S547 | 0.793101819 | 8.61E-03 | 4.77E-02 |
| ACLY_HUMAN|P53396_S455 | 0.793097555 | 5.80E-03 | 3.63E-02 |
| SPD2A_HUMAN|Q5TCZ1_S567 | 0.792936644 | 6.95E-03 | 2.56E-02 |
| LIPB1_HUMAN|Q86W92_T602 | 0.792921392 | 3.88E-03 | 1.65E-02 |
| RIOK1_HUMAN|Q9BRS2_S21 | 0.792816424 | 6.83E-03 | 2.52E-02 |
| BRE1A_HUMAN|Q5VTR2_S138 | 0.791987309 | 5.41E-03 | 2.11E-02 |
| SMN_HUMAN|Q16637_S31_T25 | 0.791925089 | 5.49E-03 | 3.30E-02 |
| DLG5_HUMAN|Q8TDM6_S1209 | 0.791817215 | 2.39E-03 | 1.78E-02 |
| RUFY3_HUMAN|Q7L099-3_S618 | 0.791752687 | 5.78E-03 | 3.62E-02 |
| TFE3_HUMAN|P19532_S564_S568 | 0.791729814 | 3.40E-03 | 4.81E-02 |
| EXOC1_HUMAN|Q9NV70_T471 | 0.791279526 | 7.59E-03 | 3.23E-02 |
| RABE1_HUMAN|Q15276_S407_S410 | 0.79115547 | 2.66E-03 | 2.00E-02 |
| NFRKB_HUMAN|Q6P4R8_S1291 | 0.790966086 | 2.39E-03 | 1.11E-02 |
| COHA1_HUMAN|Q9UMD9_S172 | 0.790867024 | 9.37E-03 | 4.89E-02 |
| AF1L2_HUMAN|Q8N4X5_S640 | 0.790690403 | 7.11E-03 | 3.05E-02 |
| NEDD1_HUMAN|Q8NHV4_S516 | 0.790239326 | 2.96E-03 | 2.10E-02 |
| SYEP_HUMAN|P07814_S882_S886 | 0.789869326 | 1.02E-03 | 1.57E-02 |
| SMN_HUMAN|Q16637_S31_T25 | 0.789791592 | 5.03E-03 | 2.29E-02 |
| CARL1_HUMAN|Q5VZK9_S1315 | 0.789158163 | 9.67E-03 | 3.92E-02 |
| MILK1_HUMAN|Q8N3F8_S391 | 0.788748417 | 3.10E-03 | 3.42E-02 |
| FA83H_HUMAN|Q6ZRV2_S892 | 0.788598885 | 5.81E-03 | 2.23E-02 |
| RBM15_HUMAN|Q96T37_S109 | 0.788588603 | 2.30E-03 | 2.58E-02 |
| CTU2_HUMAN|Q2VPK5_S419 | 0.788566241 | 8.15E-03 | 2.94E-02 |
| NIPA_HUMAN|Q86WB0_S321 | 0.788538529 | 6.18E-03 | 4.63E-02 |
| F169A_HUMAN|Q9Y6X4_S615_S618_S619 | 0.788500433 | 1.47E-02 | 4.71E-02 |
| SNAPN_HUMAN|O95295_S133 | 0.788400857 | 6.83E-03 | 2.52E-02 |
| SLK_HUMAN|Q9H2G2_S372 | 0.7883673 | 3.48E-03 | 4.87E-02 |
| MINK1_HUMAN|Q8N4C8_S778 | 0.788145279 | 8.70E-03 | 4.81E-02 |
| CAVN3_HUMAN|Q969G5_S165_S166 | 0.787988534 | 5.94E-03 | 2.27E-02 |
| ACAP2_HUMAN|Q15057_S384_T382 | 0.787870341 | 1.15E-03 | 6.82E-03 |

| | | | |
|---|---|---|---|
| PLCH2_HUMAN\|O75038_S984 | 0.787728633 | 9.49E-03 | 3.86E-02 |
| ANKZ1_HUMAN\|Q9H8Y5_S675 | 0.787661838 | 9.69E-03 | 3.37E-02 |
| YLPM1_HUMAN\|P49750_S634 | 0.787613444 | 6.84E-03 | 3.90E-02 |
| R3HD1_HUMAN\|Q15032_S973 | 0.787593262 | 1.12E-02 | 3.77E-02 |
| NUMA1_HUMAN\|Q14980_T2000 | 0.787429729 | 1.58E-02 | 4.95E-02 |
| YBOX1_HUMAN\|P67809_S174 | 0.787348854 | 6.56E-03 | 3.78E-02 |
| SYEP_HUMAN\|P07814_S885_S886 | 0.786999997 | 2.19E-03 | 2.21E-02 |
| RHG12_HUMAN\|Q8IWW6_S240 | 0.786966077 | 9.15E-03 | 3.23E-02 |
| LMNA_HUMAN\|P02545_S403_S406_S407 | 0.786877805 | 1.25E-02 | 4.82E-02 |
| CENPB_HUMAN\|P07199_S156 | 0.786784064 | 1.04E-02 | 3.59E-02 |
| KANK1_HUMAN\|Q14678_S144 | 0.786603421 | 2.85E-03 | 2.11E-02 |
| IWS1_HUMAN\|Q96ST2_S400 | 0.786569846 | 1.21E-02 | 4.00E-02 |
| PDLI2_HUMAN\|Q96JY6_S124_S129 | 0.786468821 | 4.59E-03 | 4.44E-02 |
| DEFI6_HUMAN\|Q9H4E7_S597_T595 | 0.786460491 | 7.60E-03 | 4.39E-02 |
| PCM1_HUMAN\|Q15154_S65_S69 | 0.786384968 | 2.50E-03 | 3.00E-02 |
| RBM8A_HUMAN\|Q9Y5S9_S56 | 0.786314433 | 4.30E-03 | 1.78E-02 |
| CD2B2_HUMAN\|O95400_S195 | 0.786303599 | 1.96E-03 | 9.47E-03 |
| VINEX_HUMAN\|O60504_S530 | 0.786149967 | 9.84E-03 | 3.40E-02 |
| CCDC9_HUMAN\|Q9Y3X0_T389 | 0.785862708 | 5.66E-03 | 2.91E-02 |
| TWF1_HUMAN\|Q12792_T348 | 0.785634556 | 4.61E-03 | 4.45E-02 |
| FRMD6_HUMAN\|Q96NE9_S375 | 0.785593391 | 6.52E-03 | 2.43E-02 |
| TENS4_HUMAN\|Q8IZW8_T194 | 0.785506636 | 8.02E-03 | 3.37E-02 |
| FOXK2_HUMAN\|Q01167_S428 | 0.785502847 | 5.03E-03 | 4.90E-02 |
| G3P_HUMAN\|P04406_S210 | 0.785227101 | 9.53E-03 | 3.87E-02 |
| MGMT_HUMAN\|P16455_S201 | 0.785207751 | 3.04E-03 | 1.35E-02 |
| YJ005_HUMAN\|Q6ZSR9_S93 | 0.784984918 | 2.47E-03 | 2.93E-02 |
| ANR17_HUMAN\|O75179_S2045_S2047 | 0.784866746 | 8.48E-03 | 4.00E-02 |
| PUM1_HUMAN\|Q14671_S709 | 0.78459495 | 8.21E-03 | 3.43E-02 |
| NOC3L_HUMAN\|Q8WTT2_S787 | 0.784563654 | 5.81E-03 | 3.63E-02 |
| MI4GD_HUMAN\|A9UHW6_S221 | 0.784444125 | 5.47E-03 | 2.46E-02 |
| FOXO3_HUMAN\|O43524_S253 | 0.784318943 | 6.80E-03 | 4.05E-02 |

| | | | |
|---|---|---|---|
| SI1L3_HUMAN\|O60292_S1364 | 0.784286521 | 6.53E-03 | 2.85E-02 |
| IRS2_HUMAN\|Q9Y4H2_S736 | 0.784274167 | 3.53E-03 | 1.53E-02 |
| CCNL2_HUMAN\|Q96S94_S330 | 0.784102831 | 7.86E-03 | 4.49E-02 |
| CACO1_HUMAN\|Q9P1Z2_S563 | 0.784056096 | 5.31E-03 | 3.21E-02 |
| SAFB1_HUMAN\|Q15424_S604 | 0.783993252 | 4.65E-03 | 4.66E-02 |
| DIDO1_HUMAN\|Q9BTC0_S523 | 0.783778317 | 6.10E-03 | 2.30E-02 |
| PHIP_HUMAN\|Q8WWQ0_S1315 | 0.783555788 | 3.70E-03 | 2.15E-02 |
| CTU2_HUMAN\|Q2VPK5_S419 | 0.783487309 | 6.74E-03 | 3.85E-02 |
| SNIP1_HUMAN\|Q8TAD8_S52_S54_S58 | 0.783345988 | 7.06E-03 | 4.17E-02 |
| TR150_HUMAN\|Q9Y2W1_S682 | 0.783177174 | 5.20E-03 | 3.16E-02 |
| AHNK_HUMAN\|Q09666_S5752 | 0.783067481 | 7.14E-03 | 4.02E-02 |
| EXOC1_HUMAN\|Q9NV70_T471 | 0.782826718 | 5.52E-03 | 2.13E-02 |
| MILK1_HUMAN\|Q8N3F8_S578 | 0.782758403 | 8.89E-03 | 4.74E-02 |
| SYEP_HUMAN\|P07814_S885_S886 | 0.782232399 | 1.75E-03 | 1.88E-02 |
| MTAP2_HUMAN\|P11137_S1591_T1602_T1605 | 0.782084986 | 1.05E-02 | 3.61E-02 |
| SRRM2_HUMAN\|Q9UQ35_S2132 | 0.781958528 | 7.85E-03 | 2.84E-02 |
| LIPB1_HUMAN\|Q86W92_S582 | 0.781815573 | 7.37E-03 | 2.68E-02 |
| AHNK_HUMAN\|Q09666_S5841 | 0.781777377 | 4.19E-03 | 1.97E-02 |
| TDIF2_HUMAN\|Q5QJE6_S236_S239_T232 | 0.781727076 | 6.33E-03 | 2.38E-02 |
| MGMT_HUMAN\|P16455_S201 | 0.781667756 | 2.60E-03 | 2.49E-02 |
| PURB_HUMAN\|Q96QR8_S304 | 0.781659041 | 1.07E-02 | 4.27E-02 |
| UBP31_HUMAN\|Q70CQ4_S1088 | 0.781436509 | 2.03E-03 | 1.58E-02 |
| AFF4_HUMAN\|Q9UHB7_S1043 | 0.781304078 | 9.43E-03 | 4.90E-02 |
| MILK1_HUMAN\|Q8N3F8_S391 | 0.781297417 | 2.23E-03 | 1.69E-02 |
| PDLI2_HUMAN\|Q96JY6_S124_S129 | 0.781289212 | 3.71E-03 | 3.22E-02 |
| MPDZ_HUMAN\|O75970_S1595 | 0.78111264 | 1.28E-02 | 4.20E-02 |
| AF1L2_HUMAN\|Q8N4X5_S651 | 0.780981302 | 5.21E-03 | 2.78E-02 |
| ZCH18_HUMAN\|Q86VM9_S532_S534 | 0.78078497 | 3.14E-03 | 2.26E-02 |
| TENS4_HUMAN\|Q8IZW8_S416 | 0.780748221 | 5.51E-03 | 2.48E-02 |
| KS6A3_HUMAN\|P51812_S715 | 0.78061623 | 4.42E-03 | 2.99E-02 |
| LUZP1_HUMAN\|Q86V48_S659 | 0.780369483 | 1.05E-02 | 4.21E-02 |

| | | | |
|---|---|---|---|
| MK06_HUMAN\|Q16659_S386_T389 | 0.78024732 | 2.49E-03 | 1.91E-02 |
| ELF1_HUMAN\|P32519_S163 | 0.779797639 | 4.99E-03 | 2.00E-02 |
| AFF4_HUMAN\|Q9UHB7_S1055_S1058 | 0.779782431 | 3.05E-03 | 2.15E-02 |
| RPRD2_HUMAN\|Q5VT52_T723 | 0.779603804 | 5.28E-03 | 3.39E-02 |
| AFAP1_HUMAN\|Q8N556_S548 | 0.779412229 | 4.96E-03 | 3.06E-02 |
| PPHLN_HUMAN\|Q8NEY8_S205 | 0.779307306 | 1.50E-02 | 4.77E-02 |
| SSH2_HUMAN\|Q76I76_S487 | 0.779167145 | 3.63E-03 | 3.91E-02 |
| IRS2_HUMAN\|Q9Y4H2_T1202 | 0.779150256 | 1.11E-03 | 8.62E-03 |
| JIP4_HUMAN\|O60271_S815 | 0.779114131 | 1.10E-02 | 3.74E-02 |
| MACF1_HUMAN\|Q9UPN3_S4 | 0.779067123 | 3.37E-03 | 3.10E-02 |
| OSTF1_HUMAN\|Q92882_S202_S213 | 0.778946561 | 6.34E-03 | 3.18E-02 |
| PPHLN_HUMAN\|Q8NEY8_S140 | 0.778932501 | 9.18E-03 | 4.83E-02 |
| AMOL2_HUMAN\|Q9Y2J4_S540 | 0.778904128 | 4.00E-03 | 4.05E-02 |
| SSH2_HUMAN\|Q76I76_S487 | 0.778664701 | 3.55E-03 | 1.72E-02 |
| RBM26_HUMAN\|Q5T8P6_S616 | 0.778581387 | 1.13E-02 | 3.80E-02 |
| CAR11_HUMAN\|Q9BXL7_S593 | 0.778548759 | 8.76E-03 | 3.60E-02 |
| DLG5_HUMAN\|Q8TDM6_S1209 | 0.778545035 | 1.27E-03 | 7.37E-03 |
| FA83H_HUMAN\|Q6ZRV2_S998 | 0.778430751 | 4.06E-03 | 1.91E-02 |
| AFF4_HUMAN\|Q9UHB7_S1055_S1058 | 0.778316733 | 2.86E-03 | 1.29E-02 |
| YLPM1_HUMAN\|P49750_S634 | 0.777962865 | 4.69E-03 | 4.50E-02 |
| BAD_HUMAN\|Q92934_S99 | 0.777922659 | 5.83E-03 | 2.59E-02 |
| IF2B3_HUMAN\|O00425_S179_S180 | 0.777771217 | 1.29E-03 | 1.89E-02 |
| VIP2_HUMAN\|O43314_S1006_S1016 | 0.777688078 | 2.21E-03 | 1.48E-02 |
| HEAT6_HUMAN\|Q6AI08_S643 | 0.777599207 | 4.44E-03 | 3.00E-02 |
| EF2K_HUMAN\|O00418_S27_S31 | 0.777589595 | 4.18E-03 | 2.87E-02 |
| SSH2_HUMAN\|Q76I76_S487 | 0.777530218 | 3.39E-03 | 3.62E-02 |
| G3P_HUMAN\|P04406_S210 | 0.777504901 | 7.24E-03 | 4.23E-02 |
| PLEC_HUMAN\|Q15149_S4613_S4618_T4623 | 0.777349255 | 6.43E-03 | 2.41E-02 |
| PGK1_HUMAN\|P00558_S203 | 0.777295397 | 1.35E-02 | 4.39E-02 |
| LIPB1_HUMAN\|Q86W92_S582 | 0.777256175 | 6.25E-03 | 3.81E-02 |
| FA83H_HUMAN\|Q6ZRV2_S523 | 0.777222192 | 2.64E-03 | 2.94E-02 |

| | | | |
|---|---|---|---|
| GIT1_HUMAN\|Q9Y2X7_T480 | 0.777197946 | 1.71E-03 | 1.38E-02 |
| MA7D1_HUMAN\|Q3KQU3_S366 | 0.77718581 | 4.81E-03 | 3.01E-02 |
| PKHG3_HUMAN\|A1L390_S1081 | 0.777106025 | 4.93E-03 | 2.66E-02 |
| MILK1_HUMAN\|Q8N3F8_S640 | 0.777067531 | 4.29E-03 | 1.78E-02 |
| KS6A3_HUMAN\|P51812_S715 | 0.777012913 | 3.82E-03 | 4.07E-02 |
| SRRM2_HUMAN\|Q9UQ35_S2272 | 0.776978309 | 4.48E-03 | 4.06E-02 |
| NU214_HUMAN\|P35658_S985_S986 | 0.776974083 | 1.07E-02 | 4.78E-02 |
| MEPCE_HUMAN\|Q7L2J0_S216 | 0.776957215 | 6.01E-03 | 2.29E-02 |
| P63_HUMAN\|Q9H3D4_S463 | 0.776908561 | 7.90E-03 | 4.50E-02 |
| SPTB2_HUMAN\|Q01082_S2138 | 0.776891157 | 4.49E-03 | 3.01E-02 |
| M4K4_HUMAN\|O95819_S580 | 0.776822146 | 8.64E-03 | 3.08E-02 |
| RABX5_HUMAN\|Q9UJ41_S607 | 0.776800304 | 4.82E-03 | 3.01E-02 |
| K1671_HUMAN\|Q9BY89_S404 | 0.776262551 | 6.01E-03 | 2.29E-02 |
| DEFI6_HUMAN\|Q9H4E7_S597_T595 | 0.776153029 | 5.17E-03 | 3.15E-02 |
| KIF1B_HUMAN\|O60333_S1057 | 0.776090858 | 1.44E-02 | 4.63E-02 |
| CAMP3_HUMAN\|Q9P1Y5_S334 | 0.77600842 | 2.76E-03 | 1.74E-02 |
| PURB_HUMAN\|Q96QR8_S304 | 0.775839785 | 8.82E-03 | 3.14E-02 |
| STRP1_HUMAN\|Q5VSL9_S339 | 0.775803629 | 6.68E-03 | 2.49E-02 |
| CDK13_HUMAN\|Q14004_T1246 | 0.775750104 | 1.01E-03 | 6.10E-03 |
| SPD2A_HUMAN\|Q5TCZ1_S1002 | 0.775453659 | 8.62E-03 | 4.77E-02 |
| NU214_HUMAN\|P35658_S678 | 0.775389165 | 4.20E-03 | 3.89E-02 |
| SRRM2_HUMAN\|Q9UQ35_S1519 | 0.775368803 | 1.36E-02 | 4.42E-02 |
| KLC1_HUMAN\|Q07866-4_S631 | 0.775353363 | 6.37E-03 | 3.86E-02 |
| MLF2_HUMAN\|Q15773_S238 | 0.775319125 | 1.26E-02 | 4.87E-02 |
| RBM15_HUMAN\|Q96T37_S294 | 0.775258119 | 7.37E-03 | 4.11E-02 |
| RTKN_HUMAN\|Q9BST9_S543 | 0.775214494 | 1.07E-02 | 3.66E-02 |
| MDC1_HUMAN\|Q14676_S453_T455 | 0.775203863 | 1.59E-02 | 4.98E-02 |
| T2FA_HUMAN\|P35269_S433 | 0.775203561 | 7.43E-03 | 4.32E-02 |
| ZC11A_HUMAN\|O75152_S132 | 0.775194011 | 4.66E-03 | 4.66E-02 |
| AHNK_HUMAN\|Q09666_S177 | 0.775044925 | 5.57E-03 | 2.90E-02 |
| KIZ_HUMAN\|Q2M2Z5_S668_T665 | 0.775029626 | 5.41E-03 | 4.58E-02 |

| | | | |
|---|---|---|---|
| AHNK_HUMAN\|Q09666_S559 | 0.774851171 | 6.36E-03 | 2.79E-02 |
| LMNA_HUMAN\|P02545_S22 | 0.774842957 | 5.62E-03 | 2.91E-02 |
| FA83H_HUMAN\|Q6ZRV2_S892 | 0.7747742 | 3.30E-03 | 3.55E-02 |
| SRRM2_HUMAN\|Q9UQ35_S1443 | 0.774672086 | 6.09E-03 | 2.30E-02 |
| R113A_HUMAN\|O15541_S329 | 0.774567326 | 4.63E-03 | 2.14E-02 |
| ZN687_HUMAN\|Q8N1G0_S433 | 0.774524307 | 1.31E-03 | 1.51E-02 |
| VIP2_HUMAN\|O43314_S1006_S1013_S1016 | 0.774410851 | 5.28E-03 | 4.87E-02 |
| ASPC1_HUMAN\|Q9BZE9_S275 | 0.774398234 | 1.47E-02 | 4.71E-02 |
| VIME_HUMAN\|P08670_S47 | 0.774363658 | 1.16E-02 | 4.55E-02 |
| YAP1_HUMAN\|P46937_S109_T110 | 0.774311066 | 3.84E-03 | 2.56E-02 |
| ZCH18_HUMAN\|Q86VM9_S532_S534 | 0.774243787 | 2.36E-03 | 2.86E-02 |
| CAP1_HUMAN\|Q01518_S308 | 0.774223397 | 1.18E-02 | 3.92E-02 |
| PTN14_HUMAN\|Q15678_S593_S594 | 0.774208167 | 4.85E-03 | 3.19E-02 |
| NAA10_HUMAN\|P41227_S228_S235 | 0.774043891 | 2.26E-03 | 1.06E-02 |
| ZN687_HUMAN\|Q8N1G0_T432 | 0.774003898 | 1.68E-03 | 1.37E-02 |
| SPTB2_HUMAN\|Q01082_S2341 | 0.77399234 | 1.32E-03 | 3.47E-02 |
| SPD2A_HUMAN\|Q5TCZ1_S724 | 0.773946109 | 8.72E-03 | 4.69E-02 |
| SOX15_HUMAN\|O60248_S37 | 0.773900081 | 4.75E-03 | 1.93E-02 |
| VIP2_HUMAN\|O43314_S1006 | 0.773820151 | 7.18E-03 | 2.63E-02 |
| ADNP_HUMAN\|Q9H2P0_S709 | 0.773708403 | 3.52E-03 | 1.71E-02 |
| RBM15_HUMAN\|Q96T37_S109 | 0.773678976 | 1.17E-03 | 1.06E-02 |
| SHRM2_HUMAN\|Q13796_S1039 | 0.773594416 | 8.33E-04 | 5.26E-03 |
| DPYL3_HUMAN\|Q14195_T509 | 0.773561849 | 1.90E-03 | 2.46E-02 |
| IF2B3_HUMAN\|O00425_S179_S180 | 0.773514673 | 1.05E-03 | 1.60E-02 |
| ANKZ1_HUMAN\|Q9H8Y5_S675 | 0.77343135 | 5.82E-03 | 2.58E-02 |
| NGAP_HUMAN\|Q9UJF2_S802 | 0.773126338 | 3.57E-03 | 2.52E-02 |
| LUZP1_HUMAN\|Q86V48_S659 | 0.772957764 | 8.19E-03 | 4.46E-02 |
| P63_HUMAN\|Q9H3D4_S463 | 0.772778942 | 6.81E-03 | 2.52E-02 |
| SARG_HUMAN\|Q9BW04_S502 | 0.772544373 | 6.69E-03 | 4.00E-02 |
| KLC1_HUMAN\|Q07866-4_S599 | 0.772385415 | 9.80E-03 | 3.39E-02 |
| LARP1_HUMAN\|Q6PKG0_S546 | 0.772357638 | 1.29E-02 | 4.94E-02 |

| | | | |
|---|---|---|---|
| COBL1_HUMAN\|Q53SF7_S294 | 0.772245189 | 5.90E-03 | 2.61E-02 |
| SARG_HUMAN\|Q9BW04_T456 | 0.77218633 | 3.18E-03 | 3.38E-02 |
| NEDD1_HUMAN\|Q8NHV4_S516 | 0.771612072 | 1.25E-03 | 1.80E-02 |
| MTAP2_HUMAN\|P11137_S1787_S1790_T1780 | 0.771567461 | 8.41E-03 | 4.57E-02 |
| ANS1A_HUMAN\|Q92625_S649 | 0.771522984 | 3.36E-03 | 3.54E-02 |
| WDR47_HUMAN\|O94967_S422 | 0.771515246 | 4.96E-03 | 4.66E-02 |
| OSB10_HUMAN\|Q9BXB5_S26_S32 | 0.771492514 | 4.87E-03 | 2.23E-02 |
| ZCHC8_HUMAN\|Q6NZY4_S500 | 0.771401184 | 4.09E-03 | 1.71E-02 |
| SP100_HUMAN\|P23497_S171 | 0.771366336 | 1.10E-02 | 4.35E-02 |
| PCY1A_HUMAN\|P49585_S362 | 0.771350859 | 2.15E-03 | 1.02E-02 |
| RIOK2_HUMAN\|Q9BVS4_S382_S385 | 0.77127418 | 5.28E-03 | 2.39E-02 |
| GRDN_HUMAN\|Q3V6T2_S1653 | 0.771182012 | 4.30E-03 | 4.19E-02 |
| FA83H_HUMAN\|Q6ZRV2_Y875 | 0.771170762 | 3.69E-03 | 3.84E-02 |
| DC1L2_HUMAN\|O43237_S194 | 0.77111781 | 1.36E-03 | 1.25E-02 |
| RALY_HUMAN\|Q9UKM9_S295_T286_T298 | 0.77080478 | 8.21E-03 | 2.95E-02 |
| TMF1_HUMAN\|P82094_S338_S344 | 0.770781212 | 7.85E-03 | 3.32E-02 |
| TDIF2_HUMAN\|Q5QJE6_S117 | 0.770561591 | 2.02E-03 | 9.70E-03 |
| SRG2B_HUMAN\|P0DMP2_T203 | 0.770498585 | 3.30E-03 | 2.97E-02 |
| DKC1_HUMAN\|O60832_T496 | 0.770490433 | 2.77E-03 | 1.25E-02 |
| AHNK_HUMAN\|Q09666_S216 | 0.770229209 | 5.86E-03 | 2.25E-02 |
| FA83H_HUMAN\|Q6ZRV2_S1003 | 0.77014633 | 2.57E-03 | 1.18E-02 |
| MI4GD_HUMAN\|A9UHW6_S221 | 0.770067364 | 3.07E-03 | 1.35E-02 |
| KLC2_HUMAN\|Q9H0B6_S581 | 0.770064883 | 2.31E-03 | 1.08E-02 |
| ABI2_HUMAN\|Q9NYB9_S368 | 0.769983161 | 3.14E-03 | 4.58E-02 |
| SEPT9_HUMAN\|Q9UHD8_S82 | 0.769978515 | 1.25E-02 | 4.12E-02 |
| CRBG1_HUMAN\|Q9Y4K1_S348 | 0.769937041 | 1.48E-02 | 4.73E-02 |
| NELFE_HUMAN\|P18615_S353 | 0.769924003 | 1.85E-03 | 2.26E-02 |
| SAV1_HUMAN\|Q9H4B6_S27 | 0.769892442 | 5.95E-03 | 4.89E-02 |
| HELZ_HUMAN\|P42694_T1163 | 0.769886004 | 1.38E-03 | 1.25E-02 |
| MPRIP_HUMAN\|Q6WCQ1_S365 | 0.769859992 | 2.44E-03 | 2.69E-02 |
| EDC3_HUMAN\|Q96F86_S131 | 0.769793168 | 3.26E-03 | 3.25E-02 |

| | | | |
|---|---|---|---|
| MTAP2_HUMAN\|P11137_S1787_S1790_T1780 | 0.76976434 | 7.90E-03 | 4.50E-02 |
| ATX2_HUMAN\|Q99700_T741 | 0.769547101 | 5.97E-04 | 1.10E-02 |
| TDIF2_HUMAN\|Q5QJE6_S117 | 0.769415651 | 1.92E-03 | 2.50E-02 |
| TB10B_HUMAN\|Q4KMP7_S678 | 0.76931137 | 2.33E-03 | 1.09E-02 |
| 3BP5L_HUMAN\|Q7L8J4_S343 | 0.769283561 | 7.13E-03 | 4.02E-02 |
| F126B_HUMAN\|Q8IXS8_S430 | 0.76903154 | 4.97E-03 | 4.36E-02 |
| FA83H_HUMAN\|Q6ZRV2_S998 | 0.769013126 | 2.75E-03 | 3.12E-02 |
| ZN185_HUMAN\|O15231-2_S63 | 0.768990224 | 3.79E-03 | 2.53E-02 |
| TDIF2_HUMAN\|Q5QJE6_S117 | 0.768954616 | 1.88E-03 | 1.02E-02 |
| SRRM1_HUMAN\|Q8IYB3_S752_S754_S756 | 0.7688843 | 6.33E-03 | 3.84E-02 |
| MAST2_HUMAN\|Q6P0Q8_S148 | 0.768806192 | 1.15E-02 | 3.84E-02 |
| PP12C_HUMAN\|Q9BZL4_S453 | 0.76878089 | 2.40E-03 | 2.73E-02 |
| MACF1_HUMAN\|Q9UPN3_S3 | 0.768712266 | 5.49E-03 | 4.22E-02 |
| ZC3H8_HUMAN\|Q8N5P1_S59 | 0.768680978 | 4.56E-03 | 2.51E-02 |
| TWF1_HUMAN\|Q12792_T348 | 0.768519416 | 2.27E-03 | 1.79E-02 |
| BRE1A_HUMAN\|Q5VTR2_S136_S138 | 0.768485508 | 1.47E-03 | 2.01E-02 |
| AHNK_HUMAN\|Q09666_S177 | 0.768404468 | 4.32E-03 | 4.20E-02 |
| BRE1A_HUMAN\|Q5VTR2_S136_S138 | 0.768269419 | 1.45E-03 | 1.23E-02 |
| IRS2_HUMAN\|Q9Y4H2_S1100 | 0.768198035 | 5.00E-03 | 2.00E-02 |
| SI1L1_HUMAN\|O43166_S1433 | 0.768172793 | 4.86E-03 | 4.60E-02 |
| K1671_HUMAN\|Q9BY89_S404 | 0.768125131 | 4.43E-03 | 4.51E-02 |
| IASPP_HUMAN\|Q8WUF5_S74 | 0.768103931 | 3.26E-03 | 3.53E-02 |
| RBMX_HUMAN\|P38159_S175 | 0.767920211 | 9.50E-03 | 3.32E-02 |
| SRRM2_HUMAN\|Q9UQ35_T1492 | 0.767909595 | 1.37E-02 | 4.43E-02 |
| TDIF2_HUMAN\|Q5QJE6_S117 | 0.767811066 | 1.79E-03 | 2.22E-02 |
| SEPT9_HUMAN\|Q9UHD8_S85 | 0.767699279 | 4.14E-03 | 1.73E-02 |
| ACLY_HUMAN\|P53396_S455 | 0.767666985 | 2.04E-03 | 2.58E-02 |
| PFKAP_HUMAN\|Q01813_S386 | 0.767531213 | 9.19E-04 | 5.13E-03 |
| SDCG3_HUMAN\|Q96C92_S243_S249 | 0.767085759 | 3.65E-03 | 3.52E-02 |
| 2A5E_HUMAN\|Q16537_S33_S34 | 0.76701916 | 3.74E-03 | 2.62E-02 |
| 3BP5L_HUMAN\|Q7L8J4_S343 | 0.766978364 | 6.57E-03 | 2.45E-02 |

| | | | |
|---|---|---|---|
| MAP1B_HUMAN\|P46821_Y1796 | 0.766874797 | 8.93E-03 | 3.16E-02 |
| TCOF_HUMAN\|Q13428_S1230 | 0.766855342 | 8.31E-03 | 4.65E-02 |
| PLEC_HUMAN\|Q15149_S4386_S4396 | 0.766709804 | 4.81E-03 | 3.01E-02 |
| OPTN_HUMAN\|Q96CV9_T204 | 0.766697822 | 4.97E-03 | 4.36E-02 |
| NUCKS_HUMAN\|Q9H1E3_S214 | 0.766610155 | 5.20E-03 | 2.05E-02 |
| TIAM1_HUMAN\|Q13009_S358 | 0.766550648 | 9.11E-03 | 3.22E-02 |
| UBE3A_HUMAN\|Q05086_S218 | 0.766450492 | 3.85E-03 | 1.64E-02 |
| PLCH2_HUMAN\|O75038_S982 | 0.766404732 | 2.53E-03 | 1.93E-02 |
| KLC1_HUMAN\|Q07866-4_S599 | 0.766376212 | 8.00E-03 | 3.36E-02 |
| ZCCHV_HUMAN\|Q7Z2W4_S378 | 0.76629106 | 1.63E-03 | 1.17E-02 |
| AKAP2_HUMAN\|Q9Y2D5_S152_T157 | 0.76619606 | 1.30E-02 | 4.25E-02 |
| TCPA_HUMAN\|P17987_S544 | 0.766186003 | 4.04E-03 | 1.70E-02 |
| SRGP3_HUMAN\|O43295_S1070 | 0.766169507 | 7.64E-03 | 4.40E-02 |
| PLEC_HUMAN\|Q15149_S4386_S4396 | 0.766135893 | 4.71E-03 | 3.13E-02 |
| FA83H_HUMAN\|Q6ZRV2_S647 | 0.766095061 | 2.77E-03 | 2.07E-02 |
| SRRM2_HUMAN\|Q9UQ35_S1621 | 0.765969651 | 6.50E-03 | 2.42E-02 |
| BD1L1_HUMAN\|Q8NFC6_S2964 | 0.765948654 | 3.43E-03 | 2.43E-02 |
| SRRM2_HUMAN\|Q9UQ35_S1483 | 0.765531061 | 2.78E-03 | 1.26E-02 |
| F120A_HUMAN\|Q9NZB2_S1023 | 0.765452147 | 8.88E-03 | 3.65E-02 |
| NUP53_HUMAN\|Q8NFH5_T280 | 0.765437335 | 2.49E-03 | 1.29E-02 |
| ADNP_HUMAN\|Q9H2P0_S921 | 0.765429757 | 7.94E-03 | 3.35E-02 |
| TE2IP_HUMAN\|Q9NYB0_S203 | 0.765424338 | 2.17E-03 | 2.47E-02 |
| BAZ2A_HUMAN\|Q9UIF9_S1397 | 0.765312827 | 8.72E-03 | 4.08E-02 |
| DDB2_HUMAN\|Q92466_S24_S26 | 0.765270031 | 1.09E-03 | 8.50E-03 |
| RABL6_HUMAN\|Q3YEC7_S425_S427 | 0.765246086 | 4.45E-03 | 3.00E-02 |
| FOXK2_HUMAN\|Q01167_S428 | 0.765222152 | 2.20E-03 | 2.74E-02 |
| PSD3_HUMAN\|Q9NYI0_S1012_S1020 | 0.7651064 | 4.25E-03 | 3.51E-02 |
| PTN14_HUMAN\|Q15678_S593_S594 | 0.76500405 | 3.43E-03 | 3.57E-02 |
| RB6I2_HUMAN\|Q8IUD2_S37_S42_S45 | 0.764952207 | 2.52E-03 | 1.93E-02 |
| STK10_HUMAN\|O94804_S448_S455 | 0.76483906 | 8.18E-03 | 2.94E-02 |
| KS6A3_HUMAN\|P51812_S715 | 0.764810091 | 2.30E-03 | 1.73E-02 |

| | | | |
|---|---|---|---|
| CAAP1_HUMAN\|Q9H8G2_S92 | 0.764788176 | 6.54E-03 | 4.79E-02 |
| GIT1_HUMAN\|Q9Y2X7_S385_S388_T392 | 0.764745961 | 1.14E-02 | 4.48E-02 |
| TRI16_HUMAN\|O95361_S60 | 0.764499019 | 4.36E-03 | 4.46E-02 |
| ES8L1_HUMAN\|Q8TE68_S182_T187 | 0.764193248 | 3.59E-03 | 3.49E-02 |
| ZCCHV_HUMAN\|Q7Z2W4_S378 | 0.764164315 | 1.48E-03 | 7.62E-03 |
| RPTOR_HUMAN\|Q8N122_S863 | 0.764093876 | 3.52E-03 | 1.70E-02 |
| ROA3_HUMAN\|P51991_S355 | 0.764012882 | 3.86E-03 | 2.69E-02 |
| KLC2_HUMAN\|Q9H0B6_S610 | 0.764006713 | 2.32E-03 | 1.22E-02 |
| DCP1A_HUMAN\|Q9NPI6_S525 | 0.763826397 | 1.39E-03 | 1.26E-02 |
| SAV1_HUMAN\|Q9H4B6_S27 | 0.763825789 | 4.78E-03 | 4.76E-02 |
| SAFB1_HUMAN\|Q15424_S604 | 0.763720431 | 1.98E-03 | 2.31E-02 |
| PRAG1_HUMAN\|Q86YV5_S712 | 0.763666702 | 4.57E-03 | 2.92E-02 |
| DEK_HUMAN\|P35659_S32 | 0.763462949 | 6.26E-03 | 4.74E-02 |
| PDS5B_HUMAN\|Q9NTI5_S1259 | 0.763449049 | 8.32E-03 | 3.94E-02 |
| YAP1_HUMAN\|P46937_S105_S109 | 0.763375338 | 2.11E-03 | 1.63E-02 |
| AF1L2_HUMAN\|Q8N4X5_S640 | 0.763344817 | 2.39E-03 | 2.73E-02 |
| ZCCHV_HUMAN\|Q7Z2W4_S378 | 0.763244956 | 1.43E-03 | 1.84E-02 |
| CE170_HUMAN\|Q5SW79_S630_S636 | 0.763143115 | 6.64E-03 | 3.81E-02 |
| ALKB5_HUMAN\|Q6P6C2_S384 | 0.763132495 | 3.42E-03 | 2.37E-02 |
| CAMP3_HUMAN\|Q9P1Y5_S553 | 0.763122363 | 1.18E-03 | 1.62E-02 |
| NDRG2_HUMAN\|Q9UN36_S332 | 0.763083959 | 8.16E-03 | 4.60E-02 |
| PMS2L_HUMAN\|Q68D20_S59 | 0.763058676 | 5.12E-03 | 3.33E-02 |
| PLEC_HUMAN\|Q15149_S4613_S4618 | 0.76275914 | 1.11E-02 | 3.75E-02 |
| LMO7_HUMAN\|Q8WWI1_S1586 | 0.762678444 | 8.75E-03 | 3.60E-02 |
| PLEC_HUMAN\|Q15149_Y4615 | 0.762579126 | 1.51E-02 | 4.80E-02 |
| PYRG1_HUMAN\|P17812_S573_S575 | 0.762550182 | 3.13E-03 | 2.26E-02 |
| NAA30_HUMAN\|Q147X3_S152 | 0.762454095 | 2.76E-03 | 1.25E-02 |
| PPR18_HUMAN\|Q6NYC8_S530 | 0.762376168 | 4.17E-03 | 4.34E-02 |
| SP100_HUMAN\|P23497_S171 | 0.762295173 | 8.10E-03 | 4.43E-02 |
| KRI1_HUMAN\|Q8N9T8_S171 | 0.7622666 | 5.70E-04 | 3.50E-03 |
| PSD3_HUMAN\|Q9NYI0_S1012_S1020 | 0.762221486 | 3.81E-03 | 3.93E-02 |

| | | | |
|---|---|---|---|
| BICD2_HUMAN\|Q8TD16_S615 | 0.762207131 | 8.83E-03 | 4.11E-02 |
| MILK1_HUMAN\|Q8N3F8_S471 | 0.762076319 | 2.58E-03 | 1.33E-02 |
| SI1L1_HUMAN\|O43166_S1433 | 0.762052085 | 3.87E-03 | 1.64E-02 |
| TP53B_HUMAN\|Q12888_S831_S834 | 0.761928226 | 5.23E-03 | 2.06E-02 |
| RPB1_HUMAN\|P24928_T1877_T1885 | 0.761909017 | 4.88E-03 | 4.31E-02 |
| CLASR_HUMAN\|Q8N2M8_S547 | 0.761900389 | 2.54E-03 | 1.86E-02 |
| MAP1B_HUMAN\|P46821_S1793_S1797 | 0.761761798 | 7.08E-03 | 3.46E-02 |
| PHF14_HUMAN\|O94880_T292 | 0.761755841 | 3.36E-03 | 3.31E-02 |
| JHD2C_HUMAN\|Q15652_S943 | 0.761731118 | 2.99E-03 | 3.33E-02 |
| CSKI2_HUMAN\|Q8WXE0_S406 | 0.761506432 | 1.93E-03 | 1.59E-02 |
| IRS2_HUMAN\|Q9Y4H2_S560 | 0.761368849 | 3.62E-03 | 4.87E-02 |
| PRC2A_HUMAN\|P48634_S1147 | 0.761272628 | 4.38E-03 | 4.47E-02 |
| FBXW9_HUMAN\|Q5XUX1_S18 | 0.76119556 | 3.98E-03 | 2.76E-02 |
| PTN13_HUMAN\|Q12923_S1630 | 0.761132274 | 3.24E-03 | 2.31E-02 |
| CR025_HUMAN\|Q96B23_S145 | 0.761120321 | 3.98E-03 | 3.73E-02 |
| MICA3_HUMAN\|Q7RTP6_S977 | 0.761097053 | 6.79E-04 | 1.19E-02 |
| ZN185_HUMAN\|O15231_S155_T158 | 0.761011939 | 8.63E-03 | 3.57E-02 |
| ELF1_HUMAN\|P32519_S432 | 0.760951676 | 7.30E-04 | 1.19E-02 |
| TWF1_HUMAN\|Q12792_T349 | 0.760908182 | 3.96E-03 | 4.01E-02 |
| DC1L1_HUMAN\|Q9Y6G9_S510 | 0.760810035 | 1.05E-03 | 5.75E-03 |
| MTAP2_HUMAN\|P11137_S1591_T1602_T1605 | 0.760716137 | 4.92E-03 | 3.04E-02 |
| PDLI2_HUMAN\|Q96JY6_S134 | 0.760671964 | 1.08E-03 | 1.00E-02 |
| ELF1_HUMAN\|P32519_S163 | 0.760547259 | 2.29E-03 | 2.80E-02 |
| PRSR2_HUMAN\|Q86WR7_S18 | 0.760502808 | 1.29E-03 | 1.21E-02 |
| RBM7_HUMAN\|Q9Y580_S137 | 0.760483314 | 4.85E-03 | 3.19E-02 |
| LMNA_HUMAN\|P02545_S428 | 0.760473376 | 4.65E-03 | 2.15E-02 |
| RBM7_HUMAN\|Q9Y580_S137 | 0.760388861 | 4.83E-03 | 4.79E-02 |
| PAN3_HUMAN\|Q58A45_S354 | 0.760363306 | 9.23E-03 | 3.78E-02 |
| DIDO1_HUMAN\|Q9BTC0_S523 | 0.76030809 | 2.39E-03 | 2.87E-02 |
| NUMA1_HUMAN\|Q14980_S1757 | 0.760178211 | 1.09E-02 | 4.83E-02 |
| MOES_HUMAN\|P26038_S576 | 0.760094307 | 5.87E-03 | 2.60E-02 |

| | | | |
|---|---|---|---|
| TAB2_HUMAN\|Q9NYJ8_S372 | 0.759899295 | 7.97E-03 | 4.52E-02 |
| DESP_HUMAN\|P15924_S2815_S2820 | 0.759841598 | 6.76E-03 | 4.91E-02 |
| K1671_HUMAN\|Q9BY89_S127 | 0.759787207 | 7.21E-03 | 4.04E-02 |
| MILK1_HUMAN\|Q8N3F8_S470 | 0.759759352 | 9.14E-03 | 4.99E-02 |
| DEFI6_HUMAN\|Q9H4E7_S597 | 0.759646678 | 1.52E-03 | 1.28E-02 |
| CDK17_HUMAN\|Q00537_S137 | 0.759614621 | 1.37E-03 | 7.09E-03 |
| SHRM2_HUMAN\|Q13796_S1036_S1039 | 0.7595831 | 2.61E-03 | 1.34E-02 |
| AHNK_HUMAN\|Q09666_S5841 | 0.759331488 | 1.61E-03 | 8.07E-03 |
| AHNK_HUMAN\|Q09666_S216 | 0.75919362 | 3.87E-03 | 3.96E-02 |
| DEN4C_HUMAN\|Q5VZ89_S1184 | 0.759116739 | 7.09E-03 | 3.05E-02 |
| RHG35_HUMAN\|Q9NRY4_S1134 | 0.759094156 | 2.53E-03 | 1.30E-02 |
| YBOX1_HUMAN\|P67809_S165_S176 | 0.758972894 | 1.11E-02 | 3.75E-02 |
| ZC3HD_HUMAN\|Q5T200_S1292 | 0.75895487 | 4.53E-03 | 2.90E-02 |
| F169A_HUMAN\|Q9Y6X4_S399 | 0.758790631 | 9.27E-03 | 3.27E-02 |
| PDIP3_HUMAN\|Q9BY77_S127 | 0.758715426 | 6.01E-03 | 2.29E-02 |
| RHG23_HUMAN\|Q9P227_S611 | 0.758699132 | 7.20E-03 | 4.04E-02 |
| ZN687_HUMAN\|Q8N1G0_S433 | 0.758660057 | 6.00E-04 | 6.44E-03 |
| STX7_HUMAN\|O15400_S131 | 0.758641344 | 5.45E-03 | 4.61E-02 |
| AGFG1_HUMAN\|P52594_S181 | 0.758591894 | 2.29E-03 | 1.07E-02 |
| AFF4_HUMAN\|Q9UHB7_S1055_S1058 | 0.758556057 | 1.19E-03 | 1.75E-02 |
| TFE3_HUMAN\|P19532_S564_S568 | 0.75846351 | 7.17E-04 | 1.23E-02 |
| TAOK1_HUMAN\|Q7L7X3_S421 | 0.758418728 | 5.63E-03 | 2.91E-02 |
| CCNL2_HUMAN\|Q96S94_S330 | 0.758357897 | 2.93E-03 | 3.30E-02 |
| SPTB2_HUMAN\|Q01082_S2341 | 0.75835413 | 6.40E-04 | 3.85E-03 |
| RHG23_HUMAN\|Q9P227_S361_S372 | 0.758338641 | 2.08E-03 | 3.48E-02 |
| ARHGB_HUMAN\|O15085_S1519 | 0.758244668 | 1.12E-02 | 4.94E-02 |
| MYPT1_HUMAN\|O14974_S445 | 0.758144816 | 7.70E-03 | 4.43E-02 |
| PCM1_HUMAN\|Q15154_S65_S68 | 0.757939131 | 7.96E-03 | 3.35E-02 |
| CDK13_HUMAN\|Q14004_T1246 | 0.757934298 | 4.10E-04 | 2.70E-03 |
| ZN185_HUMAN\|O15231_S153_T158 | 0.757893311 | 2.80E-03 | 1.42E-02 |
| PTN14_HUMAN\|Q15678_S809 | 0.75788475 | 2.09E-03 | 1.11E-02 |

| | | | |
|---|---|---|---|
| CARL1_HUMAN|Q5VZK9_S1290_T1278 | 0.75772336 | 7.94E-03 | 3.35E-02 |
| K1C14_HUMAN|P02533_S54 | 0.757718675 | 6.27E-03 | 2.36E-02 |
| PRC2C_HUMAN|Q9Y520_T2691 | 0.75770891 | 9.65E-03 | 3.36E-02 |
| DGC14_HUMAN|Q96DF8_T386 | 0.7576727 | 6.21E-03 | 3.79E-02 |
| UBP8_HUMAN|P40818_T577 | 0.757609391 | 2.68E-03 | 1.95E-02 |
| TBC15_HUMAN|Q8TC07_S675 | 0.757549721 | 5.34E-03 | 3.41E-02 |
| NCK5L_HUMAN|Q9HCH0_S494 | 0.757455731 | 7.47E-03 | 3.19E-02 |
| PRSR2_HUMAN|Q86WR7_S43 | 0.757375119 | 2.77E-03 | 1.74E-02 |
| TFAP4_HUMAN|Q01664_S124 | 0.757372916 | 6.84E-03 | 2.53E-02 |
| LIPB1_HUMAN|Q86W92_S582 | 0.7573021 | 2.97E-03 | 3.34E-02 |
| PLEC_HUMAN|Q15149_S4626 | 0.75726362 | 5.29E-03 | 2.80E-02 |
| MLF2_HUMAN|Q15773_S240 | 0.757229628 | 1.12E-02 | 3.78E-02 |
| OAS1_HUMAN|P00973_S50 | 0.757221257 | 3.30E-03 | 2.00E-02 |
| FA83H_HUMAN|Q6ZRV2_S647 | 0.757113992 | 1.90E-03 | 2.46E-02 |
| NKAP_HUMAN|Q8N5F7_S149 | 0.757039101 | 5.05E-03 | 2.30E-02 |
| BRE1A_HUMAN|Q5VTR2_S138 | 0.757025647 | 1.23E-03 | 7.19E-03 |
| ACLY_HUMAN|P53396_S455 | 0.757005852 | 1.29E-03 | 6.75E-03 |
| WWP2_HUMAN|O00308_S211 | 0.756944196 | 5.25E-03 | 3.38E-02 |
| CTU2_HUMAN|Q2VPK5_S419 | 0.756887173 | 2.38E-03 | 4.72E-02 |
| PPR18_HUMAN|Q6NYC8_S209_S213 | 0.756851465 | 5.67E-03 | 3.37E-02 |
| SENP3_HUMAN|Q9H4L4_S169_S181 | 0.756830158 | 4.21E-03 | 2.89E-02 |
| PRP31_HUMAN|Q8WWY3_S498 | 0.756829394 | 1.14E-03 | 1.72E-02 |
| P4K2A_HUMAN|Q9BTU6_S462_S468 | 0.756775831 | 6.03E-03 | 3.07E-02 |
| PCNP_HUMAN|Q8WW12_S119 | 0.756750135 | 5.81E-03 | 3.43E-02 |
| ARHGB_HUMAN|O15085_T668_T672 | 0.756668529 | 5.19E-03 | 2.05E-02 |
| TLE1_HUMAN|Q04724_S284 | 0.756610871 | 2.99E-03 | 3.33E-02 |
| SH24A_HUMAN|Q9H788_S315 | 0.756553684 | 8.70E-03 | 4.07E-02 |
| SMAG1_HUMAN|Q9UPU9_S420_T424 | 0.756364758 | 3.60E-03 | 3.25E-02 |
| LRRF2_HUMAN|Q9Y608_S323 | 0.756286646 | 3.64E-03 | 3.18E-02 |
| FOXO3_HUMAN|O43524_S253 | 0.755920752 | 2.19E-03 | 1.16E-02 |
| PAXI_HUMAN|P49023_S258 | 0.755858305 | 5.10E-03 | 4.45E-02 |

| | | | |
|---|---|---|---|
| SRRM2_HUMAN|Q9UQ35_S1582 | 0.755790948 | 2.60E-03 | 3.04E-02 |
| BD1L1_HUMAN|Q8NFC6_S659_T658 | 0.755685184 | 6.68E-03 | 3.82E-02 |
| NOL8_HUMAN|Q76FK4_S890_T888 | 0.755588377 | 4.19E-03 | 2.87E-02 |
| K1C19_HUMAN|P08727_S14 | 0.755494793 | 9.51E-03 | 3.32E-02 |
| DIDO1_HUMAN|Q9BTC0_S523 | 0.755439493 | 1.95E-03 | 1.53E-02 |
| SHRM2_HUMAN|Q13796_S1036_S1039 | 0.755413664 | 2.21E-03 | 1.76E-02 |
| MILK1_HUMAN|Q8N3F8_S578 | 0.755385046 | 3.33E-03 | 3.72E-02 |
| KAZRN_HUMAN|Q674X7-3_S8 | 0.755364106 | 8.24E-03 | 4.63E-02 |
| RAB8A_HUMAN|P61006_S185 | 0.755304845 | 1.17E-02 | 3.90E-02 |
| OGFR_HUMAN|Q9NZT2_S349 | 0.75526197 | 6.52E-03 | 3.76E-02 |
| PHF14_HUMAN|O94880_T292 | 0.755185065 | 2.60E-03 | 2.49E-02 |
| AKP13_HUMAN|Q12802_S983 | 0.755105829 | 4.03E-03 | 3.41E-02 |
| 4EBP1_HUMAN|Q13541_S44 | 0.755087765 | 1.19E-02 | 4.63E-02 |
| OSB10_HUMAN|Q9BXB5_S26_S32 | 0.755079762 | 2.56E-03 | 3.01E-02 |
| TDIF2_HUMAN|Q5QJE6_S117 | 0.754991465 | 9.95E-04 | 1.55E-02 |
| NAV1_HUMAN|Q8NEY1_S760 | 0.754960601 | 6.23E-03 | 3.14E-02 |
| ADNP_HUMAN|Q9H2P0_S709 | 0.754921981 | 1.61E-03 | 2.22E-02 |
| AFAD_HUMAN|P55196_S1172_S1173_S1182 | 0.754829985 | 3.00E-03 | 4.44E-02 |
| IF6_HUMAN|P56537_T245 | 0.754809915 | 6.59E-03 | 3.28E-02 |
| G3BP1_HUMAN|Q13283_S231 | 0.754737097 | 3.96E-03 | 1.87E-02 |
| YAP1_HUMAN|P46937_S109_T119 | 0.754555133 | 1.82E-03 | 2.24E-02 |
| SRG2B_HUMAN|P0DMP2_T203 | 0.754446774 | 1.72E-03 | 1.22E-02 |
| MISP_HUMAN|Q8IVT2_S675 | 0.754288984 | 5.21E-03 | 3.16E-02 |
| IRS2_HUMAN|Q9Y4H2_S1100 | 0.754286721 | 2.97E-03 | 4.43E-02 |
| AGFG1_HUMAN|P52594_S181 | 0.754213858 | 1.90E-03 | 2.49E-02 |
| SPD2A_HUMAN|Q5TCZ1_S1038 | 0.754197105 | 5.52E-04 | 5.14E-03 |
| SAV1_HUMAN|Q9H4B6_S94 | 0.753861631 | 2.83E-03 | 1.43E-02 |
| RBCC1_HUMAN|Q8TDY2_S257 | 0.753841532 | 8.89E-03 | 3.65E-02 |
| ALKB5_HUMAN|Q6P6C2_S384 | 0.753792766 | 2.37E-03 | 2.87E-02 |
| PDLI2_HUMAN|Q96JY6_S123_S129 | 0.753616225 | 2.85E-03 | 2.67E-02 |
| CNKR1_HUMAN|Q969H4_S314 | 0.753521139 | 5.61E-03 | 2.16E-02 |

| | | | |
|---|---|---|---|
| TDIF2_HUMAN\|Q5QJE6_S117 | 0.753416961 | 9.25E-04 | 8.93E-03 |
| INT12_HUMAN\|Q96CB8_S128 | 0.753394949 | 3.12E-03 | 3.44E-02 |
| MTAP2_HUMAN\|P11137_S1591_T1602_T1605 | 0.753354384 | 3.74E-03 | 1.79E-02 |
| CSN3_HUMAN\|Q9UNS2_S421 | 0.753352649 | 1.03E-03 | 1.46E-02 |
| CRBG1_HUMAN\|Q9Y4K1_S75 | 0.753292202 | 7.02E-03 | 2.58E-02 |
| HDAC2_HUMAN\|Q92769_S394 | 0.753270859 | 8.92E-03 | 3.16E-02 |
| FOXK2_HUMAN\|Q01167_S428 | 0.753270404 | 1.33E-03 | 6.91E-03 |
| SWAHC_HUMAN\|Q53LP3_S212 | 0.753070701 | 2.31E-03 | 2.79E-02 |
| NF2IP_HUMAN\|Q8NCF5_S204 | 0.752868543 | 1.09E-02 | 3.69E-02 |
| BL1S3_HUMAN\|Q6QNY0_S65_T63 | 0.752857968 | 7.85E-03 | 3.32E-02 |
| IRS2_HUMAN\|Q9Y4H2_S608 | 0.752791469 | 4.62E-03 | 4.64E-02 |
| HELZ_HUMAN\|P42694_T1163 | 0.752752086 | 6.09E-04 | 6.52E-03 |
| EXOC2_HUMAN\|Q96KP1_S432 | 0.75275179 | 2.97E-03 | 2.74E-02 |
| SPD2A_HUMAN\|Q5TCZ1_S567 | 0.752700028 | 1.36E-03 | 7.78E-03 |
| ANXA2_HUMAN\|P07355_T19 | 0.752646797 | 1.12E-03 | 1.03E-02 |
| MTAP2_HUMAN\|P11137_S1787_S1790_T1780 | 0.752618531 | 4.27E-03 | 4.24E-02 |
| SRRM2_HUMAN\|Q9UQ35_S2272 | 0.752590244 | 1.64E-03 | 1.18E-02 |
| MAP1B_HUMAN\|P46821_S1427 | 0.752406064 | 2.90E-03 | 3.15E-02 |
| KLC1_HUMAN\|Q07866-4_S561 | 0.752365397 | 5.02E-03 | 2.00E-02 |
| TP53B_HUMAN\|Q12888_S366 | 0.752289786 | 2.09E-03 | 4.64E-02 |
| AKT1_HUMAN\|P31749_S122 | 0.752287472 | 1.68E-03 | 8.36E-03 |
| OPTN_HUMAN\|Q96CV9_S528 | 0.752210913 | 2.20E-03 | 1.48E-02 |
| APBB2_HUMAN\|Q92870_S409_S412 | 0.752017654 | 6.27E-03 | 3.82E-02 |
| TAF6_HUMAN\|P49848_S532 | 0.751911355 | 3.18E-03 | 3.21E-02 |
| KS6A3_HUMAN\|P51812_S715 | 0.751890532 | 1.32E-03 | 7.61E-03 |
| CAR11_HUMAN\|Q9BXL7_S593 | 0.751662273 | 3.31E-03 | 2.30E-02 |
| MINT_HUMAN\|Q96T58_S2101 | 0.751610393 | 8.96E-04 | 8.78E-03 |
| PRAG1_HUMAN\|Q86YV5_S712 | 0.751608838 | 2.91E-03 | 2.71E-02 |
| MEPCE_HUMAN\|Q7L2J0_S217 | 0.751532416 | 3.08E-03 | 1.36E-02 |
| RABE1_HUMAN\|Q15276_S410_T408 | 0.751521315 | 5.18E-03 | 3.35E-02 |
| ARI1A_HUMAN\|O14497_S381 | 0.751363689 | 3.50E-03 | 2.07E-02 |

| | | | |
|---|---|---|---|
| CF132_HUMAN\|Q5T0Z8_S738 | 0.751252254 | 1.70E-03 | 2.15E-02 |
| ARHGG_HUMAN\|Q5VV41_S107 | 0.75121594 | 1.50E-03 | 8.44E-03 |
| FOXO3_HUMAN\|O43524_S253 | 0.751113916 | 1.79E-03 | 8.80E-03 |
| SLK_HUMAN\|Q9H2G2_S372 | 0.751034033 | 6.47E-04 | 1.16E-02 |
| IF4G2_HUMAN\|P78344_S17 | 0.751021085 | 4.32E-04 | 5.02E-03 |
| BRD2_HUMAN\|P25440_S298 | 0.750637671 | 1.18E-03 | 6.26E-03 |
| DDX17_HUMAN\|Q92841_S599 | 0.750602269 | 2.69E-03 | 2.87E-02 |
| PARP4_HUMAN\|Q9UKK3_T101 | 0.750597318 | 5.48E-03 | 2.13E-02 |
| ACINU_HUMAN\|Q9UKV3_T563 | 0.750563206 | 1.47E-03 | 2.73E-02 |
| MINT_HUMAN\|Q96T58_S623 | 0.750525125 | 6.96E-04 | 2.37E-02 |
| NU214_HUMAN\|P35658_S678 | 0.750509879 | 1.48E-03 | 1.09E-02 |
| 1B07_HUMAN\|P01889_S356 | 0.750426253 | 6.83E-03 | 2.52E-02 |
| YBOX3_HUMAN\|P16989_S203_S204 | 0.750414019 | 7.51E-03 | 3.62E-02 |
| RTKN_HUMAN\|Q9BST9_S520 | 0.750408048 | 2.60E-03 | 1.19E-02 |
| SNAPN_HUMAN\|O95295_S133 | 0.750394247 | 1.45E-03 | 1.23E-02 |
| PDLI2_HUMAN\|Q96JY6_S123_S129 | 0.750349411 | 2.51E-03 | 3.00E-02 |
| PKHG3_HUMAN\|A1L390_S617 | 0.750230369 | 4.45E-03 | 1.83E-02 |
| AR6P4_HUMAN\|Q66PJ3_S239 | 0.750058043 | 9.85E-03 | 3.98E-02 |
| ATX2L_HUMAN\|Q8WWM7_S680_S684 | 0.75003486 | 1.62E-03 | 9.00E-03 |
| PDLI2_HUMAN\|Q96JY6_S134_T146 | 0.750018843 | 1.10E-02 | 3.73E-02 |
| RIOK1_HUMAN\|Q9BRS2_S22 | 0.749986887 | 3.27E-04 | 4.04E-03 |
| LIPB1_HUMAN\|Q86W92_S579 | 0.749978161 | 3.48E-03 | 1.69E-02 |
| EXOC1_HUMAN\|Q9NV70_T468 | 0.749901215 | 6.32E-04 | 4.20E-03 |
| C170B_HUMAN\|Q9Y4F5_S1179 | 0.749842238 | 2.49E-03 | 1.29E-02 |
| MEPCE_HUMAN\|Q7L2J0_S216 | 0.749761729 | 2.16E-03 | 2.71E-02 |
| TKT_HUMAN\|P29401_T287 | 0.749565546 | 4.56E-03 | 2.11E-02 |
| ZN185_HUMAN\|O15231_S521 | 0.749533169 | 5.45E-03 | 2.12E-02 |
| IF4E_HUMAN\|P06730_T211 | 0.749532962 | 4.45E-03 | 2.07E-02 |
| AKTS1_HUMAN\|Q96B36_S202_S203 | 0.749444223 | 2.88E-03 | 2.76E-02 |
| CACO1_HUMAN\|Q9P1Z2_S563 | 0.749443583 | 1.28E-03 | 1.78E-02 |
| SLK_HUMAN\|Q9H2G2_T820 | 0.749302837 | 1.79E-03 | 9.76E-03 |

| | | | |
|---|---|---|---|
| NUMB_HUMAN|P49757_T636 | 0.749297679 | 8.67E-03 | 4.67E-02 |
| PLEC_HUMAN|Q15149_S4386_S4396 | 0.749264949 | 2.45E-03 | 1.13E-02 |
| SRC8_HUMAN|Q14247_T401 | 0.749235299 | 6.12E-03 | 3.75E-02 |
| HNRPK_HUMAN|P61978_S284 | 0.749176245 | 3.80E-04 | 4.56E-03 |
| CLAP2_HUMAN|O75122_S360 | 0.749048663 | 8.75E-03 | 4.83E-02 |
| PTN14_HUMAN|Q15678_S593_S594 | 0.749033197 | 1.84E-03 | 2.43E-02 |
| RBBP6_HUMAN|Q7Z6E9_S516 | 0.748944858 | 2.74E-03 | 1.97E-02 |
| SRRM1_HUMAN|Q8IYB3_S638 | 0.748944722 | 4.01E-03 | 1.69E-02 |
| ZC11A_HUMAN|O75152_S132 | 0.748899095 | 1.57E-03 | 2.05E-02 |
| COHA1_HUMAN|Q9UMD9_S148_S150 | 0.748678739 | 7.93E-03 | 4.51E-02 |
| SPD2A_HUMAN|Q5TCZ1_S724 | 0.748657903 | 3.62E-03 | 4.87E-02 |
| SOX15_HUMAN|O60248_S37 | 0.748586353 | 1.69E-03 | 9.28E-03 |
| ASPP2_HUMAN|Q13625_S480 | 0.748443826 | 3.40E-03 | 2.41E-02 |
| DESP_HUMAN|P15924_S2815_S2820 | 0.748419303 | 4.52E-03 | 4.06E-02 |
| STRN3_HUMAN|Q13033_S202 | 0.748411506 | 2.48E-03 | 3.97E-02 |
| SRSF9_HUMAN|Q13242_S189 | 0.748366634 | 1.33E-03 | 6.91E-03 |
| SMCA1_HUMAN|P28370_S119 | 0.748333459 | 7.63E-04 | 1.06E-02 |
| APC_HUMAN|P25054_S2283 | 0.74829807 | 3.55E-03 | 2.43E-02 |
| GPTC2_HUMAN|Q9NW75_S115_S117 | 0.748260914 | 2.08E-03 | 4.42E-02 |
| PUM1_HUMAN|Q14671_S709 | 0.748150558 | 2.03E-03 | 3.46E-02 |
| SQSTM_HUMAN|Q13501_S266 | 0.748058565 | 3.63E-03 | 3.80E-02 |
| K2C5_HUMAN|P13647_S571 | 0.748018135 | 4.87E-03 | 3.03E-02 |
| GRDN_HUMAN|Q3V6T2_S1653 | 0.747994689 | 1.72E-03 | 1.46E-02 |
| RTKN_HUMAN|Q9BST9_S543 | 0.747960887 | 4.31E-03 | 4.25E-02 |
| ROA3_HUMAN|P51991_S356 | 0.747955957 | 4.15E-03 | 1.95E-02 |
| IWS1_HUMAN|Q96ST2_S720 | 0.747954121 | 1.90E-03 | 4.38E-02 |
| AHNK_HUMAN|Q09666_S5749_S5752 | 0.747934912 | 4.86E-03 | 3.03E-02 |
| DESP_HUMAN|P15924_S2820 | 0.747928021 | 5.07E-03 | 4.00E-02 |
| LARP1_HUMAN|Q6PKG0_S517_S521 | 0.747913849 | 1.54E-03 | 1.35E-02 |
| TWF1_HUMAN|Q12792_T349 | 0.747889372 | 2.41E-03 | 2.88E-02 |
| K2C5_HUMAN|P13647_Y66 | 0.747871033 | 2.99E-03 | 2.18E-02 |

| | | | |
|---|---|---|---|
| IF2B3_HUMAN\|O00425_S180_S184 | 0.747767931 | 5.10E-04 | 9.87E-03 |
| PKP4_HUMAN\|Q99569_S314 | 0.74776669 | 1.26E-03 | 9.57E-03 |
| TENS4_HUMAN\|Q8IZW8_S416 | 0.747765844 | 1.46E-03 | 7.51E-03 |
| SI1L3_HUMAN\|O60292_S1364 | 0.747723682 | 1.50E-03 | 7.66E-03 |
| CACO1_HUMAN\|Q9P1Z2_S563 | 0.747595207 | 1.18E-03 | 6.94E-03 |
| ACD_HUMAN\|Q96AP0_S425 | 0.747483326 | 4.99E-03 | 4.88E-02 |
| SRRM1_HUMAN\|Q8IYB3_S752_S754_S756 | 0.747469471 | 2.82E-03 | 3.18E-02 |
| RAI14_HUMAN\|Q9P0K7_S281 | 0.747438964 | 3.94E-03 | 3.37E-02 |
| PRC2A_HUMAN\|P48634_S1014 | 0.747311409 | 7.31E-03 | 2.67E-02 |
| IASPP_HUMAN\|Q8WUF5_S74 | 0.747302332 | 1.39E-03 | 1.26E-02 |
| RB6I2_HUMAN\|Q8IUD2_S37_S42_S45 | 0.747207051 | 1.17E-03 | 6.92E-03 |
| AKP13_HUMAN\|Q12802_S2728 | 0.7471974 | 1.81E-03 | 2.24E-02 |
| AR6P4_HUMAN\|Q66PJ3_S239 | 0.746892382 | 8.92E-03 | 4.74E-02 |
| MFAP1_HUMAN\|P55081_S116_S118 | 0.746824911 | 4.82E-03 | 2.22E-02 |
| PTN14_HUMAN\|Q15678_S594 | 0.746647566 | 2.45E-04 | 1.93E-03 |
| NEDD1_HUMAN\|Q8NHV4_S516 | 0.746618228 | 3.69E-04 | 7.38E-03 |
| OSBP1_HUMAN\|P22059_S382 | 0.746539046 | 1.18E-03 | 6.94E-03 |
| GGYF1_HUMAN\|O75420_S24 | 0.746261009 | 5.15E-03 | 4.47E-02 |
| KANK1_HUMAN\|Q14678_S144 | 0.746195576 | 4.39E-04 | 1.20E-02 |
| PHF20_HUMAN\|Q9BVI0_S878_S880 | 0.746190516 | 5.10E-03 | 4.74E-02 |
| PKP2_HUMAN\|Q99959_S151_S155 | 0.746155642 | 8.42E-03 | 4.57E-02 |
| 3BP1_HUMAN\|Q9Y3L3_S695 | 0.746135291 | 1.29E-02 | 4.23E-02 |
| XPR1_HUMAN\|Q9UBH6_T690 | 0.746116252 | 9.85E-03 | 3.98E-02 |
| TB182_HUMAN\|Q9C0C2_S872 | 0.746108464 | 4.92E-03 | 4.63E-02 |
| RALY_HUMAN\|Q9UKM9_S135 | 0.746094462 | 9.33E-04 | 5.20E-03 |
| K1671_HUMAN\|Q9BY89_S404 | 0.746004082 | 1.88E-03 | 2.45E-02 |
| ACINU_HUMAN\|Q9UKV3_S240_S243 | 0.74597256 | 3.12E-03 | 3.17E-02 |
| AHNK_HUMAN\|Q09666_S5731 | 0.74589736 | 5.13E-04 | 7.26E-03 |
| PHLB1_HUMAN\|Q86UU1_S461 | 0.745879856 | 1.13E-03 | 6.05E-03 |
| EMAL4_HUMAN\|Q9HC35_S981 | 0.745868473 | 1.37E-02 | 4.44E-02 |
| MTAP2_HUMAN\|P11137_S1782_S1786_S1790 | 0.745605971 | 6.62E-03 | 3.97E-02 |

| | | | |
|---|---|---|---|
| AGM1_HUMAN|O95394_S64 | 0.745536767 | 3.78E-03 | 2.53E-02 |
| GUAA_HUMAN|P49915_S332 | 0.745490634 | 2.64E-03 | 4.98E-02 |
| AGFG1_HUMAN|P52594_S181 | 0.745484099 | 1.31E-03 | 1.15E-02 |
| NUMB_HUMAN|P49757_T636 | 0.745451422 | 7.69E-03 | 3.26E-02 |
| SHRM2_HUMAN|Q13796_S764 | 0.745350856 | 1.49E-02 | 4.74E-02 |
| PLCH2_HUMAN|O75038_S982 | 0.745283287 | 1.02E-03 | 9.64E-03 |
| SRRM2_HUMAN|Q9UQ35_S1483 | 0.745168763 | 1.18E-03 | 1.76E-02 |
| 1433Z_HUMAN|P63104_S230 | 0.745034138 | 4.64E-03 | 1.89E-02 |
| SRSF7_HUMAN|Q16629_S192_S194_S196 | 0.744962324 | 2.84E-03 | 2.66E-02 |
| PRC2A_HUMAN|P48634_S761_S764 | 0.744858769 | 1.61E-03 | 1.33E-02 |
| PP14A_HUMAN|Q96A00_S128_S134_S136 | 0.744607205 | 3.17E-03 | 3.21E-02 |
| TB182_HUMAN|Q9C0C2_S1711 | 0.744592307 | 5.44E-03 | 3.45E-02 |
| ROA3_HUMAN|P51991_S356 | 0.744585721 | 3.67E-03 | 3.93E-02 |
| CENPB_HUMAN|P07199_S156 | 0.744561103 | 2.18E-03 | 2.72E-02 |
| RBMX2_HUMAN|Q9Y388_S188 | 0.74453012 | 2.25E-03 | 2.25E-02 |
| MYPT1_HUMAN|O14974_S445 | 0.744523065 | 4.92E-03 | 2.25E-02 |
| I2BPL_HUMAN|Q9H1B7_S657_S662 | 0.744502514 | 3.62E-03 | 2.12E-02 |
| GGYF2_HUMAN|Q6Y7W6_S23 | 0.744328639 | 2.08E-03 | 3.48E-02 |
| DPYL2_HUMAN|Q16555_T509 | 0.74419433 | 5.65E-04 | 1.04E-02 |
| SMG9_HUMAN|Q9H0W8_S53 | 0.744141646 | 2.21E-03 | 1.76E-02 |
| DDI2_HUMAN|Q5TDH0_S194 | 0.744073868 | 1.31E-02 | 5.00E-02 |
| RGS3_HUMAN|P49796_S1044 | 0.744051595 | 5.25E-03 | 4.86E-02 |
| CNOT2_HUMAN|Q9NZN8_S165 | 0.744032825 | 3.23E-03 | 3.42E-02 |
| SI1L3_HUMAN|O60292_S172 | 0.743977152 | 2.78E-04 | 2.12E-03 |
| IASPP_HUMAN|Q8WUF5_Y109 | 0.74394951 | 5.62E-03 | 3.54E-02 |
| 3BP5L_HUMAN|Q7L8J4_S343 | 0.743896695 | 2.80E-03 | 2.09E-02 |
| BIG1_HUMAN|Q9Y6D6_S397 | 0.743849517 | 5.64E-03 | 3.55E-02 |
| TBCD4_HUMAN|O60343_S318 | 0.743836288 | 5.37E-03 | 3.24E-02 |
| LMNA_HUMAN|P02545_S428 | 0.74382287 | 2.45E-03 | 2.95E-02 |
| C170B_HUMAN|Q9Y4F5_S1179 | 0.743802881 | 1.96E-03 | 9.46E-03 |
| NCOR1_HUMAN|O75376_S1472 | 0.743783572 | 1.01E-02 | 3.49E-02 |

| | | | |
|---|---|---|---|
| BD1L1_HUMAN\|Q8NFC6_T658 | 0.743673569 | 6.50E-03 | 3.24E-02 |
| FCHO2_HUMAN\|Q0JRZ9_S403 | 0.74366197 | 1.17E-03 | 3.08E-02 |
| RHG21_HUMAN\|Q5T5U3_S615 | 0.743631534 | 3.07E-03 | 3.41E-02 |
| ANKZ1_HUMAN\|Q9H8Y5_S675 | 0.743616902 | 1.86E-03 | 3.27E-02 |
| SFR19_HUMAN\|Q9H7N4_S498_S500 | 0.743514389 | 2.66E-03 | 4.12E-02 |
| PCM1_HUMAN\|Q15154_S65_S69 | 0.743500076 | 3.27E-04 | 4.04E-03 |
| SLK_HUMAN\|Q9H2G2_T820 | 0.743465077 | 1.41E-03 | 2.02E-02 |
| AKP13_HUMAN\|Q12802_S2728 | 0.743369616 | 1.55E-03 | 2.15E-02 |
| DPYL3_HUMAN\|Q14195-2_S102 | 0.743360897 | 7.28E-03 | 3.11E-02 |
| VIME_HUMAN\|P08670_S56 | 0.743351469 | 3.85E-03 | 3.93E-02 |
| SARG_HUMAN\|Q9BW04_T456 | 0.743337913 | 9.20E-04 | 5.68E-03 |
| PDIP3_HUMAN\|Q9BY77_S127 | 0.743229656 | 3.47E-03 | 3.68E-02 |
| URFB1_HUMAN\|Q6BDS2_S1106 | 0.743180212 | 4.85E-03 | 3.03E-02 |
| SRGP3_HUMAN\|O43295_S1070 | 0.743136909 | 3.35E-03 | 1.64E-02 |
| CLAP2_HUMAN\|O75122_S360 | 0.743109485 | 7.25E-03 | 3.10E-02 |
| KMT2D_HUMAN\|O14686_S2274 | 0.743101776 | 7.89E-03 | 3.78E-02 |
| F169A_HUMAN\|Q9Y6X4_T448 | 0.742979871 | 5.95E-03 | 2.27E-02 |
| BRAF_HUMAN\|P15056_S447 | 0.742941006 | 9.62E-03 | 4.40E-02 |
| IF4E_HUMAN\|P06730_T211 | 0.742918358 | 3.50E-03 | 4.87E-02 |
| CBX3_HUMAN\|Q13185_S99 | 0.742907722 | 1.13E-03 | 6.06E-03 |
| SRGP3_HUMAN\|O43295_S1070 | 0.742864529 | 3.31E-03 | 3.50E-02 |
| NDRG1_HUMAN\|Q92597_S330_S333 | 0.742858793 | 6.28E-03 | 3.82E-02 |
| PRR12_HUMAN\|Q9ULL5_S44 | 0.742855391 | 9.64E-03 | 3.91E-02 |
| YBOX3_HUMAN\|P16989_S203_S204 | 0.742663229 | 5.84E-03 | 4.85E-02 |
| ZYX_HUMAN\|Q15942_S259 | 0.7426243 | 4.21E-03 | 2.75E-02 |
| IRS2_HUMAN\|Q9Y4H2_S1186_T1202 | 0.742561514 | 1.64E-03 | 2.98E-02 |
| RASA3_HUMAN\|Q14644_S833 | 0.742553454 | 3.39E-03 | 2.35E-02 |
| DESP_HUMAN\|P15924_S2821_S2825 | 0.742369013 | 6.09E-03 | 2.30E-02 |
| ARI1A_HUMAN\|O14497_S696_S698 | 0.74222656 | 2.73E-03 | 1.24E-02 |
| KIF1C_HUMAN\|O43896_S817 | 0.742170767 | 5.67E-03 | 2.18E-02 |
| PRP31_HUMAN\|Q8WWY3_S498 | 0.742147968 | 5.80E-04 | 3.91E-03 |

| | | | |
|---|---|---|---|
| PHLB1_HUMAN\|Q86UU1_S461 | 0.742146985 | 9.57E-04 | 9.55E-03 |
| DPYL2_HUMAN\|Q16555_T509 | 0.742049789 | 5.12E-04 | 9.87E-03 |
| KLC2_HUMAN\|Q9H0B6_S608_S611 | 0.742049209 | 1.32E-03 | 6.88E-03 |
| CARL1_HUMAN\|Q5VZK9_S1315 | 0.742006902 | 1.64E-03 | 1.40E-02 |
| CTNB1_HUMAN\|P35222_S552 | 0.741973903 | 1.04E-02 | 4.18E-02 |
| P66B_HUMAN\|Q8WXI9_S135_T120 | 0.741939994 | 5.01E-03 | 4.69E-02 |
| GMIP_HUMAN\|Q9P107_S914_S919 | 0.741935688 | 3.82E-03 | 4.07E-02 |
| PCM1_HUMAN\|Q15154_S65_S69 | 0.741879936 | 3.01E-04 | 6.72E-03 |
| NUCKS_HUMAN\|Q9H1E3_S19_S30 | 0.741833301 | 1.22E-02 | 4.03E-02 |
| PHIP_HUMAN\|Q8WWQ0_S1783 | 0.741807261 | 2.63E-03 | 2.58E-02 |
| PRC2A_HUMAN\|P48634_S1085_S1089_S1092 | 0.741794574 | 3.71E-03 | 3.85E-02 |
| CE170_HUMAN\|Q5SW79_S630_S636 | 0.741770748 | 3.09E-03 | 3.41E-02 |
| PRAG1_HUMAN\|Q86YV5_S712 | 0.741739101 | 1.99E-03 | 4.32E-02 |
| PP1RA_HUMAN\|Q96QC0_S313 | 0.741722004 | 5.91E-03 | 3.02E-02 |
| NU153_HUMAN\|P49790_S334 | 0.741694885 | 7.82E-03 | 4.30E-02 |
| HNRPC_HUMAN\|P07910_S239 | 0.741654978 | 4.03E-03 | 2.80E-02 |
| FA83H_HUMAN\|Q6ZRV2_S828 | 0.741641307 | 1.69E-03 | 2.30E-02 |
| NAA30_HUMAN\|Q147X3_S152 | 0.741614963 | 1.16E-03 | 1.73E-02 |
| SRRM2_HUMAN\|Q9UQ35_S1404 | 0.741511081 | 3.40E-03 | 3.12E-02 |
| CRBG1_HUMAN\|Q9Y4K1_S75 | 0.741507476 | 4.78E-03 | 2.20E-02 |
| 1B07_HUMAN\|P01889_S356 | 0.741503755 | 5.09E-03 | 4.74E-02 |
| CRBG1_HUMAN\|Q9Y4K1_S75 | 0.741337841 | 4.76E-03 | 4.21E-02 |
| ITB4_HUMAN\|P16144_S1515_S1518 | 0.741322433 | 4.08E-03 | 2.82E-02 |
| FKB15_HUMAN\|Q5T1M5_S344 | 0.741315079 | 5.97E-03 | 2.27E-02 |
| TOX4_HUMAN\|O94842_S178_S182_T176 | 0.741203464 | 2.54E-03 | 1.17E-02 |
| SARG_HUMAN\|Q9BW04_T456 | 0.741189038 | 8.36E-04 | 1.29E-02 |
| DESP_HUMAN\|P15924_S2815_S2820 | 0.7411294 | 3.47E-03 | 2.06E-02 |
| SLAI2_HUMAN\|Q9P270_S63 | 0.741117769 | 9.61E-03 | 3.35E-02 |
| C170B_HUMAN\|Q9Y4F5_S1179 | 0.741113374 | 1.76E-03 | 2.20E-02 |
| ACINU_HUMAN\|Q9UKV3_T563 | 0.741096513 | 9.79E-04 | 2.90E-02 |
| F122B_HUMAN\|Q7Z309_S115_S119 | 0.741072744 | 2.71E-03 | 2.98E-02 |

| | | | |
|---|---|---|---|
| STK10_HUMAN\|O94804_T459 | 0.741065794 | 1.20E-03 | 7.06E-03 |
| VIME_HUMAN\|P08670_S459 | 0.741022776 | 3.55E-03 | 3.85E-02 |
| AHNK_HUMAN\|Q09666_S177 | 0.740961477 | 1.43E-03 | 8.09E-03 |
| ES8L2_HUMAN\|Q9H6S3_S28 | 0.740961291 | 5.74E-04 | 8.65E-03 |
| TAB2_HUMAN\|Q9NYJ8_S372 | 0.740949946 | 4.21E-03 | 4.13E-02 |
| NF1_HUMAN\|P21359_S2515 | 0.740922779 | 3.27E-03 | 1.61E-02 |
| LAP2A_HUMAN\|P42166_T160 | 0.740905737 | 3.17E-03 | 2.28E-02 |
| VCIP1_HUMAN\|Q96JH7_S1198 | 0.740875764 | 3.39E-03 | 2.03E-02 |
| PDLI4_HUMAN\|P50479_T114 | 0.740824935 | 4.21E-03 | 4.21E-02 |
| PHLB2_HUMAN\|Q86SQ0_S402 | 0.74074305 | 6.14E-03 | 3.59E-02 |
| RING1_HUMAN\|Q06587_S38 | 0.740686322 | 8.76E-03 | 3.60E-02 |
| ZCCHV_HUMAN\|Q7Z2W4_T380 | 0.740413464 | 7.94E-03 | 3.80E-02 |
| PKP3_HUMAN\|Q9Y446_Y176 | 0.740380401 | 1.02E-02 | 4.09E-02 |
| TB10B_HUMAN\|Q4KMP7_S22 | 0.740334343 | 3.69E-03 | 3.55E-02 |
| CLASR_HUMAN\|Q8N2M8_S547 | 0.740260215 | 1.02E-03 | 6.17E-03 |
| MAGI3_HUMAN\|Q5TCQ9_S724 | 0.740236323 | 6.49E-04 | 3.88E-03 |
| IASPP_HUMAN\|Q8WUF5_Y109 | 0.740103111 | 4.95E-03 | 4.85E-02 |
| ARHGG_HUMAN\|Q5VV41_S107 | 0.740001354 | 9.28E-04 | 1.47E-02 |
| AAPK1_HUMAN\|Q13131_T488_T490 | 0.739975591 | 6.04E-03 | 4.94E-02 |
| NECP2_HUMAN\|Q9NVZ3_T182 | 0.739811593 | 3.06E-03 | 3.40E-02 |
| MPRIP_HUMAN\|Q6WCQ1_S365 | 0.739780587 | 6.58E-04 | 6.82E-03 |
| NU214_HUMAN\|P35658_T987 | 0.739741192 | 5.02E-03 | 3.09E-02 |
| TLN1_HUMAN\|Q9Y490_S729 | 0.73972198 | 6.09E-03 | 2.30E-02 |
| CHAP1_HUMAN\|Q96JM3_S542 | 0.739547639 | 6.82E-04 | 5.97E-03 |
| UT14A_HUMAN\|Q9BVJ6_S434 | 0.739502041 | 7.80E-03 | 3.74E-02 |
| SPTB2_HUMAN\|Q01082_S2160_S2161_S2169 | 0.739499918 | 2.28E-03 | 1.72E-02 |
| GSK3A_HUMAN\|P49840_S282_Y279 | 0.739490851 | 4.34E-03 | 2.95E-02 |
| ES8L2_HUMAN\|Q9H6S3_S28 | 0.739478078 | 5.38E-04 | 1.98E-02 |
| IASPP_HUMAN\|Q8WUF5_S113 | 0.739413417 | 3.88E-03 | 1.84E-02 |
| FBXW9_HUMAN\|Q5XUX1_S18 | 0.739407037 | 1.74E-03 | 9.50E-03 |
| PLEC_HUMAN\|Q15149_S4385_S4386 | 0.739386999 | 2.20E-04 | 3.96E-03 |

| | | | |
|---|---|---|---|
| NHRF1_HUMAN\|O14745_S280 | 0.739358447 | 4.80E-03 | 3.01E-02 |
| SRRM1_HUMAN\|Q8IYB3_S752_S754_S756 | 0.739328112 | 2.04E-03 | 2.58E-02 |
| YBOX1_HUMAN\|P67809_S174_S176 | 0.739131209 | 4.22E-03 | 2.75E-02 |
| PKP2_HUMAN\|Q99959_S312 | 0.739105387 | 1.34E-03 | 1.17E-02 |
| RABL6_HUMAN\|Q3YEC7_S492 | 0.739083247 | 2.70E-03 | 2.02E-02 |
| ABLM1_HUMAN\|O14639_S655 | 0.739031203 | 7.44E-03 | 4.14E-02 |
| K2C5_HUMAN\|P13647_S6 | 0.738914481 | 9.99E-03 | 3.45E-02 |
| PININ_HUMAN\|Q9H307_T657 | 0.738871123 | 2.09E-04 | 5.17E-03 |
| FA83H_HUMAN\|Q6ZRV2_S998 | 0.738835935 | 7.29E-04 | 4.26E-03 |
| GRDN_HUMAN\|Q3V6T2_S1653 | 0.738787543 | 1.18E-03 | 1.75E-02 |
| NU214_HUMAN\|P35658_S985_S986 | 0.738736905 | 2.72E-03 | 2.89E-02 |
| SPTN1_HUMAN\|Q13813_S1190 | 0.738723115 | 2.62E-03 | 2.92E-02 |
| FA83H_HUMAN\|Q6ZRV2_S514 | 0.738721121 | 1.98E-03 | 1.62E-02 |
| PPR18_HUMAN\|Q6NYC8_S490 | 0.738653301 | 1.47E-03 | 1.87E-02 |
| DDI2_HUMAN\|Q5TDH0_S106 | 0.73861532 | 2.61E-03 | 1.19E-02 |
| PHLB1_HUMAN\|Q86UU1_S430 | 0.738581474 | 1.07E-02 | 4.25E-02 |
| LMNA_HUMAN\|P02545_S428 | 0.738577401 | 1.99E-03 | 2.56E-02 |
| AHNK_HUMAN\|Q09666_S5763 | 0.738522165 | 7.42E-04 | 9.71E-03 |
| EP15R_HUMAN\|Q9UBC2_S229 | 0.738458823 | 4.37E-04 | 4.22E-03 |
| BORG4_HUMAN\|Q9H3Q1_S64 | 0.738455542 | 4.73E-03 | 4.45E-02 |
| MYPT1_HUMAN\|O14974_T443 | 0.738449697 | 5.84E-03 | 2.59E-02 |
| NONO_HUMAN\|Q15233_T450 | 0.738416628 | 4.49E-03 | 4.06E-02 |
| EPN2_HUMAN\|O95208_S173 | 0.738411505 | 1.29E-02 | 4.95E-02 |
| NU107_HUMAN\|P57740_S37 | 0.738362464 | 6.28E-03 | 3.16E-02 |
| SAV1_HUMAN\|Q9H4B6_S27 | 0.73835449 | 1.82E-03 | 4.25E-02 |
| ZC3HD_HUMAN\|Q5T200_S242 | 0.738255108 | 1.33E-03 | 1.16E-02 |
| DEFI6_HUMAN\|Q9H4E7_S597_T595 | 0.738215864 | 1.13E-03 | 1.66E-02 |
| M4K4_HUMAN\|O95819_S842 | 0.738198932 | 3.59E-03 | 2.53E-02 |
| AMOL2_HUMAN\|Q9Y2J4_S540 | 0.738117512 | 6.82E-04 | 4.04E-03 |
| TLN1_HUMAN\|Q9Y490_S1641 | 0.738085594 | 1.17E-03 | 6.22E-03 |
| HELZ_HUMAN\|P42694_T1163 | 0.738002609 | 2.94E-04 | 6.55E-03 |

| | | | |
|---|---|---|---|
| PKHF2_HUMAN\|Q9H8W4_S239 | 0.73799057 | 6.36E-03 | 3.70E-02 |
| MGMT_HUMAN\|P16455_S201 | 0.737974024 | 3.28E-04 | 2.42E-03 |
| GOGA4_HUMAN\|Q13439_T43 | 0.73785863 | 1.32E-03 | 6.90E-03 |
| RGS3_HUMAN\|P49796_S1044 | 0.737826291 | 4.22E-03 | 4.36E-02 |
| MAP1B_HUMAN\|P46821_S1427 | 0.737629432 | 1.61E-03 | 2.22E-02 |
| IASPP_HUMAN\|Q8WUF5_S567 | 0.737575192 | 7.39E-04 | 6.30E-03 |
| BI2L1_HUMAN\|Q9UHR4_S261_T257 | 0.737370012 | 2.56E-03 | 4.90E-02 |
| SSH3_HUMAN\|Q8TE77_S649_S653 | 0.737321814 | 2.10E-03 | 1.70E-02 |
| BD1L1_HUMAN\|Q8NFC6_S659_T658 | 0.737271272 | 3.51E-03 | 3.09E-02 |
| SNIP1_HUMAN\|Q8TAD8_S52_S54_S58 | 0.737238534 | 1.08E-03 | 1.00E-02 |
| M3K20_HUMAN\|Q9NYL2_S727 | 0.737208263 | 6.79E-04 | 7.19E-03 |
| LARP4_HUMAN\|Q71RC2_S597 | 0.737200227 | 6.05E-03 | 2.67E-02 |
| SAFB1_HUMAN\|Q15424_S604 | 0.737189672 | 6.00E-04 | 1.10E-02 |
| ABCF1_HUMAN\|Q8NE71_S109_T108 | 0.737184178 | 1.03E-03 | 1.00E-02 |
| SPB1_HUMAN\|Q8IY81_S333_S335 | 0.737133146 | 2.51E-03 | 2.44E-02 |
| NDRG1_HUMAN\|Q92597_S364 | 0.737095963 | 4.61E-03 | 4.64E-02 |
| TB22B_HUMAN\|Q9NU19_S132 | 0.737058848 | 4.82E-03 | 2.61E-02 |
| PLEC_HUMAN\|Q15149_S4613_S4618_T4623 | 0.736948169 | 1.29E-03 | 1.84E-02 |
| ABLM1_HUMAN\|O14639_S431_T429 | 0.736896979 | 3.27E-03 | 1.61E-02 |
| RB6I2_HUMAN\|Q8IUD2_S17_S21 | 0.736850752 | 4.89E-03 | 3.03E-02 |
| CYTSA_HUMAN\|Q69YQ0_S832 | 0.736826258 | 4.05E-04 | 1.14E-02 |
| LAP2A_HUMAN\|P42166_S66_S67_T74 | 0.736807953 | 1.05E-02 | 4.19E-02 |
| CL045_HUMAN\|Q8N5I9_S15 | 0.736791758 | 5.97E-03 | 2.27E-02 |
| BAIP2_HUMAN\|Q9UQB8-4_T520 | 0.736791504 | 1.52E-02 | 4.82E-02 |
| LARP1_HUMAN\|Q6PKG0_S774 | 0.736668593 | 9.26E-04 | 1.45E-02 |
| PCM1_HUMAN\|Q15154_S65_S68 | 0.736640038 | 3.94E-03 | 1.67E-02 |
| TANC2_HUMAN\|Q9HCD6_T168 | 0.736585295 | 2.07E-03 | 1.42E-02 |
| YAP1_HUMAN\|P46937_S164 | 0.736532508 | 1.17E-02 | 3.90E-02 |
| CCD97_HUMAN\|Q96F63_S337 | 0.736313826 | 1.67E-03 | 2.29E-02 |
| VIP2_HUMAN\|O43314_S1006_S1013_S1016 | 0.736310936 | 1.15E-03 | 1.72E-02 |
| SSH1_HUMAN\|Q8WYL5_S937 | 0.736216793 | 2.37E-03 | 2.40E-02 |

| | | | |
|---|---|---|---|
| SH3K1_HUMAN\|Q96B97_S587 | 0.736181181 | 4.15E-03 | 1.73E-02 |
| KI16B_HUMAN\|Q96L93_S662 | 0.736119281 | 3.14E-03 | 3.18E-02 |
| K1671_HUMAN\|Q9BY89_S127 | 0.736072766 | 3.19E-03 | 3.40E-02 |
| CE170_HUMAN\|Q5SW79_S630_S636 | 0.736037319 | 2.49E-03 | 3.97E-02 |
| DESP_HUMAN\|P15924_S2821_S2825 | 0.735973483 | 4.91E-03 | 3.22E-02 |
| AF1L2_HUMAN\|Q8N4X5_S640 | 0.735820043 | 7.35E-04 | 1.25E-02 |
| SPTN1_HUMAN\|Q13813_S1190 | 0.735783623 | 2.34E-03 | 2.81E-02 |
| ANR54_HUMAN\|Q6NXT1_S63 | 0.73577028 | 1.03E-02 | 4.15E-02 |
| K2C5_HUMAN\|P13647_S574 | 0.735618616 | 5.35E-03 | 4.93E-02 |
| CCD43_HUMAN\|Q96MW1_T139 | 0.735570268 | 1.18E-02 | 3.92E-02 |
| PRAG1_HUMAN\|Q86YV5_Y413 | 0.735525489 | 3.01E-03 | 1.34E-02 |
| NUP50_HUMAN\|Q9UKX7_S221 | 0.735307718 | 2.22E-03 | 2.72E-02 |
| COIL_HUMAN\|P38432_S566_S571 | 0.735273971 | 6.41E-03 | 4.74E-02 |
| C170B_HUMAN\|Q9Y4F5_S1179 | 0.735144322 | 1.38E-03 | 2.00E-02 |
| PLEC_HUMAN\|Q15149_S4622_Y4612 | 0.735097621 | 9.51E-03 | 3.32E-02 |
| TM1L1_HUMAN\|O75674_S323 | 0.735077282 | 2.05E-03 | 1.59E-02 |
| MAP1B_HUMAN\|P46821_S1792 | 0.735017094 | 1.34E-02 | 4.36E-02 |
| YBOX1_HUMAN\|P67809_S165_S174_S176 | 0.73499781 | 7.93E-04 | 7.97E-03 |
| IASPP_HUMAN\|Q8WUF5_S335 | 0.734993597 | 2.63E-03 | 4.09E-02 |
| VIME_HUMAN\|P08670_S56 | 0.734911381 | 2.82E-03 | 1.43E-02 |
| AHNK2_HUMAN\|Q8IVF2_S4185 | 0.73489287 | 2.13E-03 | 1.45E-02 |
| NFIX_HUMAN\|Q14938_S280 | 0.734829867 | 2.18E-03 | 4.52E-02 |
| SRGP3_HUMAN\|O43295_S1073 | 0.734803553 | 4.65E-03 | 4.40E-02 |
| AHNK2_HUMAN\|Q8IVF2_S278_S294 | 0.734798487 | 3.06E-03 | 1.35E-02 |
| SRRM2_HUMAN\|Q9UQ35_T1531 | 0.734763947 | 3.05E-03 | 1.35E-02 |
| NAV1_HUMAN\|Q8NEY1_S760 | 0.734745694 | 3.04E-03 | 3.12E-02 |
| HNRPC_HUMAN\|P07910_S260 | 0.734741153 | 1.61E-03 | 8.06E-03 |
| LARP1_HUMAN\|Q6PKG0_S774 | 0.73473202 | 8.53E-04 | 7.03E-03 |
| FOXO3_HUMAN\|O43524_S284 | 0.73472312 | 9.77E-03 | 3.95E-02 |
| PKP3_HUMAN\|Q9Y446_Y176 | 0.734697482 | 8.62E-03 | 4.77E-02 |
| LARP1_HUMAN\|Q6PKG0_S774 | 0.734643549 | 8.50E-04 | 1.40E-02 |

| | | | |
|---|---|---|---|
| UBP20_HUMAN\|Q9Y2K6_S132 | 0.734574533 | 2.66E-03 | 1.36E-02 |
| M3K3_HUMAN\|Q99759_S340 | 0.734316015 | 4.92E-03 | 4.63E-02 |
| MAP1B_HUMAN\|P46821_Y1796 | 0.734247898 | 2.87E-03 | 3.27E-02 |
| SPD2A_HUMAN\|Q5TCZ1_S724 | 0.73406131 | 2.12E-03 | 2.66E-02 |
| CDC23_HUMAN\|Q9UJX2_S588 | 0.734042706 | 8.39E-03 | 4.68E-02 |
| SRGP3_HUMAN\|O43295_S1073 | 0.734030629 | 4.53E-03 | 1.86E-02 |
| IWS1_HUMAN\|Q96ST2_S400 | 0.733939949 | 1.84E-03 | 2.43E-02 |
| MILK1_HUMAN\|Q8N3F8_S578 | 0.733934683 | 1.47E-03 | 7.53E-03 |
| TB182_HUMAN\|Q9C0C2_S1666 | 0.733899802 | 6.64E-04 | 2.31E-02 |
| PAXI_HUMAN\|P49023_S85 | 0.733882665 | 2.41E-03 | 2.67E-02 |
| LIMD1_HUMAN\|Q9UGP4_S424 | 0.733880938 | 4.28E-03 | 2.40E-02 |
| SRRM2_HUMAN\|Q9UQ35_S1541 | 0.733870478 | 3.96E-03 | 1.67E-02 |
| MA7D1_HUMAN\|Q3KQU3_S460 | 0.733719769 | 5.22E-03 | 2.06E-02 |
| K1522_HUMAN\|Q9P206_S327 | 0.733646764 | 1.27E-03 | 1.78E-02 |
| MILK1_HUMAN\|Q8N3F8_S644 | 0.733635416 | 5.31E-03 | 2.08E-02 |
| PLXA2_HUMAN\|O75051_S1633 | 0.733557447 | 1.18E-02 | 4.62E-02 |
| SUGP1_HUMAN\|Q8IWZ8_S411 | 0.733487305 | 5.31E-03 | 2.08E-02 |
| BI2L1_HUMAN\|Q9UHR4_S261_T257 | 0.733383839 | 2.20E-03 | 1.67E-02 |
| K1522_HUMAN\|Q9P206_S327 | 0.733380373 | 1.26E-03 | 7.32E-03 |
| SRRM2_HUMAN\|Q9UQ35_S1582 | 0.733355296 | 1.03E-03 | 1.60E-02 |
| PLEC_HUMAN\|Q15149_S4613_S4618_T4623 | 0.733211344 | 1.10E-03 | 1.06E-02 |
| COHA1_HUMAN\|Q9UMD9_S85_S93_S96 | 0.733160687 | 7.87E-03 | 4.32E-02 |
| UBP8_HUMAN\|P40818_T577 | 0.733134734 | 9.87E-04 | 9.73E-03 |
| GPTC2_HUMAN\|Q9NW75_S115_S117 | 0.73291084 | 1.09E-03 | 1.63E-02 |
| MISP_HUMAN\|Q8IVT2_S376 | 0.732894475 | 6.38E-03 | 2.39E-02 |
| ACINU_HUMAN\|Q9UKV3_S240_S243 | 0.732881232 | 1.91E-03 | 4.22E-02 |
| BD1L1_HUMAN\|Q8NFC6_S659_T658 | 0.732737235 | 2.99E-03 | 1.84E-02 |
| IASPP_HUMAN\|Q8WUF5_S74 | 0.732689654 | 7.43E-04 | 1.27E-02 |
| NECP2_HUMAN\|Q9NVZ3_T182 | 0.732659703 | 2.34E-03 | 1.23E-02 |
| KHDR1_HUMAN\|Q07666_S20 | 0.73263644 | 3.30E-03 | 3.28E-02 |
| DUSTY_HUMAN\|Q6XUX3_S928 | 0.732632513 | 2.02E-03 | 9.70E-03 |

| | | | |
|---|---|---|---|
| PCM1_HUMAN\|Q15154_S65_S68 | 0.732622325 | 3.44E-03 | 2.43E-02 |
| GGYF2_HUMAN\|Q6Y7W6_S26 | 0.732610767 | 9.05E-04 | 1.95E-02 |
| CSKI2_HUMAN\|Q8WXE0_S403_S406 | 0.73256923 | 1.72E-04 | 1.37E-03 |
| YJ005_HUMAN\|Q6ZSR9_S93 | 0.73250001 | 2.15E-04 | 4.92E-03 |
| ACAP2_HUMAN\|Q15057_S384_T382 | 0.732449217 | 6.30E-05 | 1.14E-03 |
| LIPA1_HUMAN\|Q13136_S1172 | 0.732440629 | 1.78E-03 | 1.41E-02 |
| LIMD1_HUMAN\|Q9UGP4_S277 | 0.732434633 | 2.64E-03 | 1.68E-02 |
| DC1L1_HUMAN\|Q9Y6G9_S516 | 0.732415327 | 2.26E-03 | 4.60E-02 |
| LARP1_HUMAN\|Q6PKG0_S548 | 0.73239342 | 1.18E-02 | 4.60E-02 |
| BI2L1_HUMAN\|Q9UHR4_S261_T257 | 0.732359436 | 2.12E-03 | 1.71E-02 |
| PPM1G_HUMAN\|O15355_S183 | 0.732340804 | 4.62E-03 | 1.88E-02 |
| CC117_HUMAN\|Q8IWD4_S53 | 0.732304138 | 5.43E-03 | 2.12E-02 |
| MICLK_HUMAN\|Q6ZW33_S59_T62 | 0.732180974 | 1.56E-03 | 1.31E-02 |
| TM1L1_HUMAN\|O75674_S323 | 0.732127399 | 1.83E-03 | 9.92E-03 |
| CSKI2_HUMAN\|Q8WXE0_S403_S406 | 0.732046236 | 1.68E-04 | 4.60E-03 |
| TRI16_HUMAN\|O95361_S60 | 0.732019056 | 1.17E-03 | 6.24E-03 |
| SRF_HUMAN\|P11831_S224 | 0.731971495 | 7.57E-03 | 3.22E-02 |
| G3BP1_HUMAN\|Q13283_S231 | 0.731968814 | 1.69E-03 | 2.08E-02 |
| AHNK_HUMAN\|Q09666_T566 | 0.731927403 | 4.38E-03 | 2.82E-02 |
| RBMX2_HUMAN\|Q9Y388_S188 | 0.731866663 | 1.34E-03 | 1.17E-02 |
| ACINU_HUMAN\|Q9UKV3_T563 | 0.73176377 | 6.49E-04 | 1.11E-02 |
| FCHO2_HUMAN\|Q0JRZ9_S509 | 0.731753514 | 1.69E-03 | 1.37E-02 |
| 3BP1_HUMAN\|Q9Y3L3_S695 | 0.73173957 | 8.61E-03 | 4.65E-02 |
| PSF2_HUMAN\|Q9Y248_S179_S182 | 0.731739385 | 7.27E-03 | 2.66E-02 |
| NAV1_HUMAN\|Q8NEY1_S760 | 0.731670813 | 2.71E-03 | 1.96E-02 |
| PKP2_HUMAN\|Q99959_S132_S135 | 0.731570883 | 1.24E-02 | 4.09E-02 |
| ATAT_HUMAN\|Q5SQI0_S315 | 0.731552018 | 8.74E-04 | 2.71E-02 |
| KI16B_HUMAN\|Q96L93_S662 | 0.731528714 | 2.67E-03 | 2.96E-02 |
| PLEC_HUMAN\|Q15149_S4382_S4396 | 0.731470944 | 5.40E-03 | 3.25E-02 |
| FA83H_HUMAN\|Q6ZRV2_S925 | 0.731436704 | 4.17E-04 | 2.74E-03 |
| JIP4_HUMAN\|O60271_S815 | 0.73134441 | 2.14E-03 | 3.56E-02 |

| | | | |
|---|---|---|---|
| RPRD2_HUMAN\|Q5VT52_T723 | 0.73128305 | 7.25E-04 | 1.26E-02 |
| SRGP3_HUMAN\|O43295_S1070 | 0.731152052 | 2.13E-03 | 1.63E-02 |
| PININ_HUMAN\|Q9H307_T657 | 0.731069568 | 1.38E-04 | 4.02E-03 |
| HMGA2_HUMAN\|P52926_S14 | 0.730989351 | 4.26E-03 | 1.77E-02 |
| DESP_HUMAN\|P15924_S2815 | 0.730864048 | 2.41E-03 | 1.57E-02 |
| PEPL_HUMAN\|O60437_S1657 | 0.730862729 | 4.00E-03 | 1.69E-02 |
| MD1L1_HUMAN\|Q9Y6D9_S428 | 0.730825912 | 6.64E-04 | 6.86E-03 |
| IF4G3_HUMAN\|O43432_S495 | 0.730825225 | 3.27E-03 | 2.33E-02 |
| RIPK2_HUMAN\|O43353_S531 | 0.730824727 | 2.00E-03 | 3.45E-02 |
| MA7D1_HUMAN\|Q3KQU3_S446 | 0.730809034 | 1.76E-03 | 2.34E-02 |
| IRS2_HUMAN\|Q9Y4H2_S730_S736 | 0.730771716 | 2.97E-03 | 3.32E-02 |
| RHG23_HUMAN\|Q9P227_S361_S372 | 0.730768565 | 6.14E-04 | 3.71E-03 |
| PDLI4_HUMAN\|P50479_S116 | 0.730757362 | 9.05E-03 | 4.79E-02 |
| DKC1_HUMAN\|O60832_T496 | 0.73075155 | 4.80E-04 | 9.21E-03 |
| BIG1_HUMAN\|Q9Y6D6_S394 | 0.730647712 | 4.65E-03 | 1.89E-02 |
| MA7D1_HUMAN\|Q3KQU3_S460 | 0.730450739 | 4.68E-03 | 4.50E-02 |
| IF4G2_HUMAN\|P78344_T508 | 0.73040832 | 5.24E-04 | 9.79E-03 |
| MAST4_HUMAN\|O15021_S1387_S1394 | 0.730335494 | 2.04E-03 | 2.10E-02 |
| YBOX3_HUMAN\|P16989_S369_T366 | 0.730280811 | 2.86E-03 | 4.30E-02 |
| DKC1_HUMAN\|O60832_S485_T496 | 0.730261128 | 5.92E-04 | 3.60E-03 |
| SPD2A_HUMAN\|Q5TCZ1_S1038 | 0.730230695 | 1.67E-04 | 1.45E-03 |
| PEAK1_HUMAN\|Q9H792_S1035 | 0.730210794 | 2.55E-03 | 2.86E-02 |
| K2C6B_HUMAN\|P04259_S546 | 0.73019819 | 4.40E-03 | 4.25E-02 |
| MAP1B_HUMAN\|P46821_T1503 | 0.730114489 | 4.61E-03 | 1.88E-02 |
| RAI14_HUMAN\|Q9P0K7_S281 | 0.730041055 | 2.06E-03 | 4.41E-02 |
| RRP8_HUMAN\|O43159_S104_S106 | 0.730001336 | 1.66E-03 | 1.42E-02 |
| YBOX3_HUMAN\|P16989_S203 | 0.729961923 | 3.73E-03 | 4.96E-02 |
| KI26B_HUMAN\|Q2KJY2_S1724 | 0.729903573 | 1.15E-02 | 3.84E-02 |
| SQSTM_HUMAN\|Q13501_S272_T269 | 0.72978699 | 3.49E-03 | 3.09E-02 |
| OSBP1_HUMAN\|P22059_S379_S382 | 0.729612554 | 1.45E-03 | 1.61E-02 |
| SH24A_HUMAN\|Q9H788_S315 | 0.729554354 | 3.58E-03 | 4.87E-02 |

| | | | |
|---|---|---|---|
| ELF1_HUMAN\|P32519_S432 | 0.729553084 | 1.46E-04 | 8.31E-03 |
| PLEC_HUMAN\|Q15149_S4382_S4396 | 0.729444382 | 5.05E-03 | 3.28E-02 |
| SERF2_HUMAN\|P84101_S21 | 0.729429608 | 1.13E-02 | 3.79E-02 |
| PRC2A_HUMAN\|P48634_S1147 | 0.729255216 | 1.28E-03 | 1.83E-02 |
| CE170_HUMAN\|Q5SW79_S630_S636 | 0.729201421 | 1.92E-03 | 2.33E-02 |
| TP53B_HUMAN\|Q12888_S831 | 0.729137645 | 6.55E-04 | 4.33E-03 |
| TB10B_HUMAN\|Q4KMP7_S22 | 0.729086268 | 2.48E-03 | 1.29E-02 |
| K1522_HUMAN\|Q9P206_S404_S409 | 0.729080457 | 2.59E-03 | 1.96E-02 |
| T184B_HUMAN\|Q9Y519_S403 | 0.729021879 | 9.38E-03 | 3.29E-02 |
| SCAF8_HUMAN\|Q9UPN6_S617 | 0.72897164 | 4.98E-03 | 4.88E-02 |
| CAAP1_HUMAN\|Q9H8G2_S92 | 0.72896383 | 1.75E-03 | 1.23E-02 |
| PRC2A_HUMAN\|P48634_S1085_S1089_S1092 | 0.728926674 | 2.34E-03 | 1.75E-02 |
| DLG5_HUMAN\|Q8TDM6_S1264 | 0.728907796 | 4.61E-03 | 4.37E-02 |
| DOCK5_HUMAN\|Q9H7D0_S1805 | 0.728876115 | 2.49E-03 | 4.83E-02 |
| PKHG3_HUMAN\|A1L390_S617 | 0.728800308 | 2.05E-03 | 3.47E-02 |
| LARP6_HUMAN\|Q9BRS8_S56_S58 | 0.728794485 | 5.86E-03 | 2.60E-02 |
| M3K20_HUMAN\|Q9NYL2_S667 | 0.728681817 | 5.57E-05 | 1.05E-03 |
| KLC3_HUMAN\|Q6P597_S497_S502 | 0.728646606 | 4.31E-04 | 2.81E-03 |
| AKAP2_HUMAN\|Q9Y2D5_S152_T157 | 0.728625165 | 3.84E-03 | 2.68E-02 |
| KIF1C_HUMAN\|O43896_S817 | 0.728570863 | 3.64E-03 | 3.72E-02 |
| PRSR2_HUMAN\|Q86WR7_S18 | 0.728542266 | 2.80E-04 | 2.13E-03 |
| ITBP1_HUMAN\|O14713_S10_S14 | 0.728539444 | 3.69E-03 | 1.78E-02 |
| FA83H_HUMAN\|Q6ZRV2_S998 | 0.728514134 | 4.51E-04 | 8.70E-03 |
| AHNK_HUMAN\|Q09666_S559 | 0.728363632 | 1.01E-03 | 1.45E-02 |
| ARHG1_HUMAN\|Q92888_S786 | 0.728337927 | 2.79E-03 | 1.26E-02 |
| ANR17_HUMAN\|O75179_S1693 | 0.728219686 | 2.49E-03 | 2.73E-02 |
| BRD4_HUMAN\|O60885_S470 | 0.728205964 | 1.10E-02 | 4.37E-02 |
| CING_HUMAN\|Q9P2M7_S129_S134 | 0.728205161 | 1.03E-02 | 4.16E-02 |
| NELFE_HUMAN\|P18615_S353 | 0.728156504 | 2.71E-04 | 6.27E-03 |
| MISP_HUMAN\|Q8IVT2_S376 | 0.728121893 | 5.50E-03 | 3.30E-02 |
| NDRG1_HUMAN\|Q92597_S330_S333 | 0.728103878 | 3.82E-03 | 3.87E-02 |

| | | | |
|---|---|---|---|
| KAP2_HUMAN\|P13861_S78_S80 | 0.7280177 | 3.44E-03 | 2.43E-02 |
| UBP20_HUMAN\|Q9Y2K6_S132 | 0.728010985 | 2.08E-03 | 1.68E-02 |
| ACTN1_HUMAN\|P12814_S140 | 0.728010224 | 1.35E-02 | 4.39E-02 |
| STAT3_HUMAN\|P40763_S727 | 0.728001723 | 1.17E-03 | 9.05E-03 |
| SH3K1_HUMAN\|Q96B97_S587 | 0.727872259 | 3.10E-03 | 1.54E-02 |
| SPT5H_HUMAN\|O00267_S666 | 0.727853571 | 8.38E-04 | 8.35E-03 |
| LAR4B_HUMAN\|Q92615_T732 | 0.727853195 | 3.81E-04 | 4.56E-03 |
| NCOA7_HUMAN\|Q8NI08_S384 | 0.727699593 | 1.38E-03 | 7.14E-03 |
| SRC8_HUMAN\|Q14247_T401 | 0.72767953 | 2.86E-03 | 3.21E-02 |
| NDRG1_HUMAN\|Q92597_S330_S333 | 0.727634933 | 3.75E-03 | 2.51E-02 |
| SNX29_HUMAN\|Q8TEQ0_S810 | 0.72756819 | 6.70E-03 | 2.90E-02 |
| ZN185_HUMAN\|O15231_S519_T513 | 0.727510298 | 1.78E-03 | 2.35E-02 |
| COBL1_HUMAN\|Q53SF7_T322 | 0.727441712 | 1.65E-03 | 2.27E-02 |
| SRRM1_HUMAN\|Q8IYB3_S740_S743 | 0.727387286 | 7.57E-03 | 4.18E-02 |
| WWC3_HUMAN\|Q9ULE0_T909 | 0.727347597 | 7.02E-03 | 2.58E-02 |
| TM1L1_HUMAN\|O75674_S323 | 0.72721534 | 1.50E-03 | 1.99E-02 |
| LMNB2_HUMAN\|Q03252_S420 | 0.727206325 | 3.33E-03 | 4.77E-02 |
| HNRL2_HUMAN\|Q1KMD3_S161_S168 | 0.727181269 | 4.81E-04 | 3.35E-03 |
| AHNK_HUMAN\|Q09666_S216 | 0.727180453 | 1.08E-03 | 1.63E-02 |
| EFHD2_HUMAN\|Q96C19_S74 | 0.72717361 | 5.11E-03 | 4.94E-02 |
| RANB3_HUMAN\|Q9H6Z4_S125 | 0.727010866 | 2.99E-03 | 1.49E-02 |
| K2C5_HUMAN\|P13647_Y66 | 0.726971936 | 1.30E-03 | 7.51E-03 |
| PDLI2_HUMAN\|Q96JY6_S134 | 0.726919605 | 2.12E-04 | 1.74E-03 |
| RAVR1_HUMAN\|Q8IY67_T463 | 0.726916671 | 5.15E-03 | 2.75E-02 |
| DAPK3_HUMAN\|O43293_S311_S312 | 0.726884096 | 2.62E-03 | 2.92E-02 |
| CCSE2_HUMAN\|Q9H7U1_S197 | 0.726880051 | 3.35E-04 | 7.28E-03 |
| ZBT21_HUMAN\|Q9ULJ3_S435_T431 | 0.72680511 | 1.98E-03 | 9.55E-03 |
| ADNP_HUMAN\|Q9H2P0_S709 | 0.726760892 | 4.65E-04 | 1.82E-02 |
| IRS2_HUMAN\|Q9Y4H2_S620 | 0.726691167 | 9.27E-03 | 4.86E-02 |
| COBL1_HUMAN\|Q53SF7_T322 | 0.726654695 | 1.60E-03 | 2.07E-02 |
| K2C5_HUMAN\|P13647_S18 | 0.726532693 | 6.33E-03 | 3.84E-02 |

| | | | |
|---|---|---|---|
| PININ_HUMAN\|Q9H307_T657 | 0.726421765 | 1.07E-04 | 9.64E-04 |
| HDAC4_HUMAN\|P56524_S266 | 0.726400085 | 2.68E-03 | 1.95E-02 |
| SHRM2_HUMAN\|Q13796_S1039 | 0.726358717 | 6.81E-05 | 9.64E-04 |
| PKP3_HUMAN\|Q9Y446_S238 | 0.726314758 | 5.54E-04 | 9.98E-03 |
| KLC3_HUMAN\|Q6P597_S167_S173 | 0.7262736 | 9.38E-03 | 3.83E-02 |
| MICLK_HUMAN\|Q6ZW33_S59_T62 | 0.726263958 | 1.23E-03 | 1.77E-02 |
| SH23A_HUMAN\|Q9BRG2_S147 | 0.726261559 | 5.49E-03 | 2.13E-02 |
| XRCC6_HUMAN\|P12956_T455 | 0.7262143 | 8.34E-04 | 4.75E-03 |
| LARP4_HUMAN\|Q71RC2_S597 | 0.726016914 | 4.16E-03 | 2.87E-02 |
| GRDN_HUMAN\|Q3V6T2_S1653 | 0.725970726 | 6.88E-04 | 4.06E-03 |
| PTN13_HUMAN\|Q12923_Y344 | 0.725920539 | 9.05E-03 | 4.79E-02 |
| ZN687_HUMAN\|Q8N1G0_T432 | 0.72578812 | 1.55E-04 | 3.96E-03 |
| FRYL_HUMAN\|O94915_T1959 | 0.72575742 | 3.38E-03 | 3.55E-02 |
| RUFY3_HUMAN\|Q7L099-3_S618 | 0.725756593 | 3.43E-04 | 6.86E-03 |
| RABE1_HUMAN\|Q15276_S407_S410 | 0.725673436 | 9.95E-05 | 9.61E-04 |
| NRK_HUMAN\|Q7Z2Y5_S854 | 0.725524279 | 4.22E-04 | 1.16E-02 |
| NIPA_HUMAN\|Q86WB0_S321 | 0.725470624 | 4.70E-04 | 8.93E-03 |
| CRBG1_HUMAN\|Q9Y4K1_S22 | 0.72546021 | 8.28E-04 | 4.75E-03 |
| SRRM2_HUMAN\|Q9UQ35_S1621 | 0.72537477 | 1.43E-03 | 1.98E-02 |
| RIOK1_HUMAN\|Q9BRS2_S21 | 0.725223023 | 3.69E-04 | 4.43E-03 |
| NIPA_HUMAN\|Q86WB0_S395 | 0.725184993 | 1.50E-04 | 1.83E-03 |
| SCRIB_HUMAN\|Q14160_S1448 | 0.725126682 | 5.39E-03 | 2.84E-02 |
| K1C16_HUMAN\|P08779_S447_S450 | 0.72498094 | 7.55E-03 | 3.22E-02 |
| PKHA1_HUMAN\|Q9HB21_S332 | 0.724888613 | 4.57E-04 | 9.28E-03 |
| LAD1_HUMAN\|O00515_T358 | 0.724882772 | 5.88E-03 | 4.87E-02 |
| HDAC4_HUMAN\|P56524_S266 | 0.724808008 | 2.53E-03 | 1.93E-02 |
| NCOA7_HUMAN\|Q8NI08_S384 | 0.724668691 | 1.22E-03 | 1.77E-02 |
| K2C5_HUMAN\|P13647_S571 | 0.724633479 | 2.05E-03 | 1.09E-02 |
| SRGP3_HUMAN\|O43295_S858 | 0.724549516 | 5.35E-03 | 2.41E-02 |
| UBP31_HUMAN\|Q70CQ4_S1088 | 0.724484649 | 1.28E-04 | 1.17E-03 |
| LIPA1_HUMAN\|Q13136_S666 | 0.724407871 | 3.27E-03 | 4.71E-02 |

| | | | |
|---|---|---|---|
| T106B_HUMAN\|Q9NUM4_S33 | 0.724390996 | 2.47E-03 | 4.83E-02 |
| NDRG2_HUMAN\|Q9UN36_S332 | 0.724372475 | 2.02E-03 | 1.08E-02 |
| PLEC_HUMAN\|Q15149_S4386_S4396 | 0.724286042 | 8.74E-04 | 1.42E-02 |
| MYO9B_HUMAN\|Q13459_S1992 | 0.724285693 | 4.04E-03 | 3.41E-02 |
| MFAP1_HUMAN\|P55081_T267 | 0.724245718 | 1.20E-02 | 4.65E-02 |
| YBOX1_HUMAN\|P67809_S165 | 0.72418156 | 1.78E-04 | 2.53E-03 |
| TCOF_HUMAN\|Q13428_S156 | 0.724098498 | 1.42E-03 | 1.22E-02 |
| FA83H_HUMAN\|Q6ZRV2_Y875 | 0.724036567 | 4.81E-04 | 9.21E-03 |
| PEAK1_HUMAN\|Q9H792_S1035 | 0.724033632 | 2.02E-03 | 2.57E-02 |
| TGS1_HUMAN\|Q96RS0_S154 | 0.724008026 | 5.61E-03 | 3.54E-02 |
| RIOK1_HUMAN\|Q9BRS2_S22 | 0.724003449 | 8.11E-05 | 7.61E-04 |
| TPMT_HUMAN\|P51580_S14 | 0.724000558 | 1.50E-03 | 8.41E-03 |
| ANS1A_HUMAN\|Q92625_S649 | 0.723969995 | 4.34E-04 | 2.83E-03 |
| SRSF9_HUMAN\|Q13242_S193 | 0.723929886 | 2.54E-03 | 2.86E-02 |
| SARG_HUMAN\|Q9BW04_T456 | 0.723825868 | 3.77E-04 | 7.87E-03 |
| DPYL2_HUMAN\|Q16555_T509 | 0.723729458 | 2.17E-04 | 1.66E-03 |
| RTKN_HUMAN\|Q9BST9_T518 | 0.723726924 | 4.42E-03 | 1.82E-02 |
| ZC3HD_HUMAN\|Q5T200_S242 | 0.723646909 | 7.16E-04 | 1.23E-02 |
| RBM39_HUMAN\|Q14498_S97 | 0.72360064 | 1.27E-04 | 1.10E-03 |
| 1B07_HUMAN\|P01889_S359 | 0.723593038 | 2.72E-03 | 3.10E-02 |
| CR025_HUMAN\|Q96B23_S145 | 0.723577689 | 8.57E-04 | 1.89E-02 |
| IWS1_HUMAN\|Q96ST2_S720 | 0.723553109 | 6.93E-04 | 1.66E-02 |
| SEPT9_HUMAN\|Q9UHD8_S30 | 0.72352732 | 8.68E-03 | 3.09E-02 |
| SMAG1_HUMAN\|Q9UPU9_S420_T424 | 0.723449828 | 9.45E-04 | 2.82E-02 |
| PHLB2_HUMAN\|Q86SQ0_T404 | 0.723300418 | 6.58E-03 | 3.97E-02 |
| OXR1_HUMAN\|Q8N573_S204_T203 | 0.723289611 | 1.78E-03 | 1.25E-02 |
| MISP_HUMAN\|Q8IVT2_S575 | 0.723284785 | 4.74E-03 | 4.21E-02 |
| CAAP1_HUMAN\|Q9H8G2_S89 | 0.723269013 | 2.99E-04 | 3.17E-03 |
| LMNA_HUMAN\|P02545_S404_S406 | 0.72324619 | 4.93E-03 | 1.98E-02 |
| K1522_HUMAN\|Q9P206_S929 | 0.723203706 | 1.18E-03 | 2.35E-02 |
| ES8L2_HUMAN\|Q9H6S3_S28 | 0.723197936 | 2.57E-04 | 6.04E-03 |

| | | | |
|---|---|---|---|
| CSN1_HUMAN\|Q13098_S483_T479 | 0.723098045 | 2.23E-03 | 1.05E-02 |
| MA7D1_HUMAN\|Q3KQU3_S460 | 0.723052812 | 3.65E-03 | 3.52E-02 |
| MAGI3_HUMAN\|Q5TCQ9_S724 | 0.723039941 | 2.86E-04 | 6.40E-03 |
| FXR2_HUMAN\|P51116_S453 | 0.723007729 | 4.96E-03 | 3.24E-02 |
| SRRM1_HUMAN\|Q8IYB3_S738_S740_S743 | 0.722917748 | 2.40E-03 | 1.86E-02 |
| PP14A_HUMAN\|Q96A00_S128_S134_S136 | 0.722775442 | 1.36E-03 | 1.90E-02 |
| RABL6_HUMAN\|Q3YEC7_S425_S427 | 0.722768302 | 7.85E-04 | 1.80E-02 |
| ITB1_HUMAN\|P05556_T789 | 0.7227662 | 1.29E-02 | 4.23E-02 |
| FA83H_HUMAN\|Q6ZRV2_S828 | 0.722710911 | 7.63E-04 | 4.92E-03 |
| RALY_HUMAN\|Q9UKM9_S288_T286 | 0.722610691 | 5.37E-04 | 3.36E-03 |
| CSKI2_HUMAN\|Q8WXE0_S403 | 0.722584866 | 8.77E-04 | 8.94E-03 |
| RTKN_HUMAN\|Q9BST9_T518 | 0.722476324 | 4.25E-03 | 4.15E-02 |
| BI2L1_HUMAN\|Q9UHR4_S261 | 0.722439196 | 4.92E-03 | 4.02E-02 |
| AHNK_HUMAN\|Q09666_S5731 | 0.722437362 | 1.57E-04 | 1.28E-03 |
| STK10_HUMAN\|O94804_S455 | 0.722406662 | 2.03E-03 | 1.58E-02 |
| PARD3_HUMAN\|Q8TEW0_S225 | 0.722243316 | 6.38E-03 | 3.86E-02 |
| CWC25_HUMAN\|Q9NXE8_S218 | 0.722207601 | 3.47E-03 | 2.39E-02 |
| NU153_HUMAN\|P49790_S334 | 0.722193318 | 4.23E-03 | 2.38E-02 |
| PP12C_HUMAN\|Q9BZL4_S453 | 0.722072538 | 2.83E-04 | 3.69E-03 |
| DGC14_HUMAN\|Q96DF8_T386 | 0.72203171 | 1.67E-03 | 9.22E-03 |
| PLEC_HUMAN\|Q15149_S4386 | 0.721969977 | 2.09E-03 | 1.61E-02 |
| GGYF2_HUMAN\|Q6Y7W6_S23 | 0.721932682 | 8.58E-04 | 2.67E-02 |
| APBB2_HUMAN\|Q92870_S409_S412 | 0.721932555 | 2.19E-03 | 1.67E-02 |
| PEAK1_HUMAN\|Q9H792_S1035 | 0.721854791 | 1.86E-03 | 9.03E-03 |
| 4EBP1_HUMAN\|Q13541_S44 | 0.721763733 | 4.20E-03 | 1.75E-02 |
| YAP1_HUMAN\|P46937_S367 | 0.721733335 | 1.04E-03 | 2.16E-02 |
| ELF1_HUMAN\|P32519_S163_S167_S168 | 0.721683631 | 1.65E-03 | 1.36E-02 |
| OPTN_HUMAN\|Q96CV9_T204 | 0.721677947 | 8.08E-04 | 6.79E-03 |
| ANXA2_HUMAN\|P07355_T19 | 0.721660257 | 2.78E-04 | 2.12E-03 |
| ARH40_HUMAN\|Q8TER5_S1513 | 0.721659279 | 3.43E-04 | 1.47E-02 |
| SRPK2_HUMAN\|P78362_S496_T498 | 0.721508333 | 4.15E-03 | 3.86E-02 |

| | | | |
|---|---|---|---|
| BL1S3_HUMAN\|Q6QNY0_S65_T63 | 0.721505183 | 2.73E-03 | 3.10E-02 |
| RALY_HUMAN\|Q9UKM9_S135 | 0.721455933 | 2.96E-04 | 6.68E-03 |
| NMD3_HUMAN\|Q96D46_S468_T470 | 0.721451674 | 2.76E-04 | 1.99E-03 |
| ZN687_HUMAN\|Q8N1G0_S433 | 0.721444906 | 8.46E-05 | 2.63E-03 |
| LIPA1_HUMAN\|Q13136_S666 | 0.721440543 | 2.95E-03 | 1.31E-02 |
| ARHGB_HUMAN\|O15085_T668_T672 | 0.721344758 | 1.39E-03 | 1.93E-02 |
| TM1L1_HUMAN\|O75674_S323 | 0.721260824 | 1.18E-03 | 1.12E-02 |
| NSD3_HUMAN\|Q9BZ95_S585 | 0.721177364 | 3.59E-03 | 3.49E-02 |
| MTCL1_HUMAN\|Q9Y4B5_S263 | 0.721079144 | 9.64E-03 | 3.35E-02 |
| SRF_HUMAN\|P11831_S224 | 0.721050779 | 5.45E-03 | 3.45E-02 |
| CND1_HUMAN\|Q15021_S1333 | 0.721002769 | 3.06E-03 | 3.14E-02 |
| SAFB1_HUMAN\|Q15424_S247 | 0.720996333 | 1.98E-03 | 2.35E-02 |
| PKP4_HUMAN\|Q99569_S336 | 0.720940793 | 2.03E-03 | 9.73E-03 |
| ITSN1_HUMAN\|Q15811_S315 | 0.720823299 | 1.91E-03 | 1.03E-02 |
| BAIP2_HUMAN\|Q9UQB8_S453 | 0.720724519 | 4.02E-03 | 1.89E-02 |
| HABP4_HUMAN\|Q5JVS0_S108 | 0.720690458 | 9.25E-04 | 5.69E-03 |
| TP53B_HUMAN\|Q12888_S831 | 0.720679386 | 4.54E-04 | 2.92E-03 |
| MICA1_HUMAN\|Q8TDZ2_S960 | 0.720676406 | 1.01E-03 | 2.86E-02 |
| IRS1_HUMAN\|P35568_S1100 | 0.720664002 | 2.63E-03 | 1.20E-02 |
| BRD1_HUMAN\|O95696_S1055_T1051 | 0.720629573 | 1.53E-02 | 4.86E-02 |
| RPB1_HUMAN\|P24928_T1877_T1885 | 0.720613213 | 9.72E-04 | 5.93E-03 |
| M3K3_HUMAN\|Q99759_S340 | 0.72060838 | 3.07E-03 | 2.79E-02 |
| KAP2_HUMAN\|P13861_S78_S80 | 0.720551826 | 2.68E-03 | 2.54E-02 |
| YAP1_HUMAN\|P46937_S105_S109 | 0.720550944 | 2.99E-04 | 6.72E-03 |
| K2C5_HUMAN\|P13647_S31 | 0.720507199 | 1.53E-02 | 4.86E-02 |
| NECP2_HUMAN\|Q9NVZ3_T182 | 0.720494692 | 1.47E-03 | 1.87E-02 |
| YBOX1_HUMAN\|P67809_S165_S176 | 0.720433528 | 3.17E-03 | 2.28E-02 |
| MILK1_HUMAN\|Q8N3F8_S644 | 0.720405356 | 3.47E-03 | 3.59E-02 |
| RU17_HUMAN\|P08621_S226 | 0.720381588 | 4.63E-03 | 1.89E-02 |
| EXPH5_HUMAN\|Q8NEV8_S341 | 0.720376587 | 3.20E-03 | 2.30E-02 |
| AHNK_HUMAN\|Q09666_S5749 | 0.720365848 | 2.99E-04 | 3.65E-02 |

| | | | |
|---|---|---|---|
| FA76A_HUMAN\|Q8TAV0_S306 | 0.720229458 | 1.87E-03 | 1.01E-02 |
| RFIP5_HUMAN\|Q9BXF6_S395 | 0.720077316 | 2.45E-03 | 2.70E-02 |
| CD2AP_HUMAN\|Q9Y5K6_S458 | 0.720071925 | 4.84E-03 | 3.19E-02 |
| MYO1E_HUMAN\|Q12965_S980 | 0.720017356 | 7.24E-03 | 2.65E-02 |
| DCAF5_HUMAN\|Q96JK2_T497_T500 | 0.720013163 | 2.54E-03 | 2.99E-02 |
| EMAL4_HUMAN\|Q9HC35_S146 | 0.719988978 | 4.62E-03 | 1.88E-02 |
| RIOK1_HUMAN\|Q9BRS2_S21 | 0.719903312 | 2.87E-04 | 2.17E-03 |
| MTAP2_HUMAN\|P11137_S1782_S1787_S1790 | 0.719868525 | 2.34E-03 | 1.75E-02 |
| LIPA1_HUMAN\|Q13136_S666 | 0.719851375 | 2.79E-03 | 2.93E-02 |
| PRC2A_HUMAN\|P48634_S1219 | 0.719851025 | 1.62E-03 | 1.33E-02 |
| KLC4_HUMAN\|Q9NSK0_S590 | 0.719805861 | 3.12E-03 | 1.37E-02 |
| UBP32_HUMAN\|Q8NFA0_S1393 | 0.719728882 | 1.74E-03 | 1.23E-02 |
| ZO2_HUMAN\|Q9UDY2_T1027 | 0.71966492 | 2.19E-03 | 1.67E-02 |
| I2BP2_HUMAN\|Q7Z5L9_S460 | 0.719657423 | 2.54E-05 | 5.97E-04 |
| RB6I2_HUMAN\|Q8IUD2_S37_S42_S45 | 0.719638596 | 3.27E-04 | 2.28E-03 |
| SEPT9_HUMAN\|Q9UHD8_S30 | 0.719576231 | 7.76E-03 | 3.29E-02 |
| SSH2_HUMAN\|Q76I76_T486 | 0.719523379 | 2.12E-03 | 1.12E-02 |
| SLAI2_HUMAN\|Q9P270_S349_S353_S357 | 0.719509506 | 1.21E-03 | 1.73E-02 |
| FA83H_HUMAN\|Q6ZRV2_S523 | 0.719442105 | 1.69E-04 | 2.45E-03 |
| MED1_HUMAN\|Q15648_Y1431 | 0.719415823 | 7.50E-04 | 4.36E-03 |
| STK10_HUMAN\|O94804_S455 | 0.719372628 | 1.82E-03 | 8.91E-03 |
| LARP1_HUMAN\|Q6PKG0_S546 | 0.719328568 | 2.25E-03 | 1.06E-02 |
| MTUS1_HUMAN\|Q9ULD2_S1264_S1268 | 0.719280591 | 7.69E-03 | 3.69E-02 |
| MILK1_HUMAN\|Q8N3F8_S471 | 0.719237697 | 4.29E-04 | 8.22E-03 |
| PRC2A_HUMAN\|P48634_S1014 | 0.719227683 | 2.91E-03 | 1.46E-02 |
| ITBP1_HUMAN\|O14713_S10_S14 | 0.719225931 | 2.65E-03 | 4.12E-02 |
| TB182_HUMAN\|Q9C0C2_S1029 | 0.719216982 | 1.62E-03 | 3.92E-02 |
| ABLM3_HUMAN\|O94929_S372_S379 | 0.719166495 | 9.05E-03 | 4.95E-02 |
| ITB4_HUMAN\|P16144_T1455 | 0.719140277 | 1.15E-02 | 3.84E-02 |
| PKHA1_HUMAN\|Q9HB21_S332 | 0.719091034 | 3.52E-04 | 4.27E-03 |
| AAK1_HUMAN\|Q2M2I8_S678_T674 | 0.719064396 | 1.45E-03 | 1.30E-02 |

| | | | |
|---|---|---|---|
| MTAP2_HUMAN\|P11137_S1591_T1602_T1605 | 0.719029648 | 9.54E-04 | 9.54E-03 |
| MYO9B_HUMAN\|Q13459_T1346 | 0.719004071 | 4.77E-03 | 2.59E-02 |
| PRSR2_HUMAN\|Q86WR7_S18 | 0.718906767 | 1.73E-04 | 1.37E-03 |
| SRPK2_HUMAN\|P78362_S497_T492 | 0.718831761 | 1.84E-03 | 2.19E-02 |
| HNRPC_HUMAN\|P07910_S260 | 0.718813231 | 8.43E-04 | 1.39E-02 |
| GBG12_HUMAN\|Q9UBI6_S26 | 0.718805845 | 6.77E-03 | 2.51E-02 |
| DEFI6_HUMAN\|Q9H4E7_S590_S597_T595 | 0.718763079 | 6.62E-03 | 2.46E-02 |
| CAVN1_HUMAN\|Q6NZI2_T302 | 0.718699902 | 8.40E-04 | 1.13E-02 |
| UBP24_HUMAN\|Q9UPU5_S1141 | 0.718699565 | 3.55E-03 | 2.43E-02 |
| AGM1_HUMAN\|O95394_S64 | 0.718696225 | 1.42E-03 | 1.83E-02 |
| LAR4B_HUMAN\|Q92615_S601 | 0.718695144 | 2.27E-03 | 1.79E-02 |
| MILK1_HUMAN\|Q8N3F8_T469 | 0.718682703 | 8.03E-04 | 8.28E-03 |
| RBBP6_HUMAN\|Q7Z6E9_S516 | 0.718655163 | 8.04E-04 | 5.15E-03 |
| MICLK_HUMAN\|Q6ZW33_S59_T62 | 0.718611221 | 8.91E-04 | 5.55E-03 |
| NSD3_HUMAN\|Q9BZ95_S585 | 0.718568855 | 3.27E-03 | 1.99E-02 |
| FA83H_HUMAN\|Q6ZRV2_S523 | 0.718557034 | 1.62E-04 | 4.50E-03 |
| TKT_HUMAN\|P29401_S295 | 0.718494215 | 5.92E-04 | 3.98E-03 |
| LARP6_HUMAN\|Q9BRS8_S56_S58 | 0.718463417 | 4.17E-03 | 4.20E-02 |
| PDLI2_HUMAN\|Q96JY6_S137 | 0.71841587 | 1.70E-03 | 3.96E-02 |
| RBM27_HUMAN\|Q9P2N5_T447 | 0.718354251 | 3.55E-03 | 3.85E-02 |
| ARVC_HUMAN\|O00192_T642 | 0.718238928 | 3.99E-03 | 1.68E-02 |
| IRS2_HUMAN\|Q9Y4H2_S608 | 0.718206867 | 1.30E-03 | 1.21E-02 |
| NCOA7_HUMAN\|Q8NI08_S208_T210 | 0.718174448 | 2.18E-03 | 4.81E-02 |
| BI2L1_HUMAN\|Q9UHR4_S414 | 0.718169014 | 8.10E-03 | 4.59E-02 |
| TFE3_HUMAN\|P19532_S568_S570 | 0.718163828 | 1.27E-03 | 9.59E-03 |
| DESP_HUMAN\|P15924_S2820_S2821 | 0.71815071 | 5.94E-03 | 3.67E-02 |
| PRC2A_HUMAN\|P48634_S1219 | 0.718094588 | 1.52E-03 | 1.11E-02 |
| TISB_HUMAN\|Q07352_S336 | 0.718070171 | 1.17E-02 | 3.89E-02 |
| RHG21_HUMAN\|Q5T5U3_S615 | 0.718043512 | 1.17E-03 | 3.08E-02 |
| FA83H_HUMAN\|Q6ZRV2_S1009 | 0.717901298 | 1.26E-02 | 4.14E-02 |
| RGL2_HUMAN\|O15211_S761 | 0.717850791 | 3.53E-03 | 2.42E-02 |

| | | | |
|---|---|---|---|
| TCOF_HUMAN\|Q13428_S156 | 0.71782068 | 1.10E-03 | 1.63E-02 |
| RU17_HUMAN\|P08621_S226 | 0.717791387 | 4.27E-03 | 4.39E-02 |
| ADNP_HUMAN\|Q9H2P0_S709 | 0.717719473 | 3.07E-04 | 6.33E-03 |
| FA83H_HUMAN\|Q6ZRV2_S1025 | 0.717616202 | 3.33E-03 | 3.72E-02 |
| ZN593_HUMAN\|O00488_T134 | 0.717522845 | 4.91E-03 | 1.97E-02 |
| TFE3_HUMAN\|P19532_S548_S556 | 0.717492103 | 5.99E-03 | 4.91E-02 |
| IASPP_HUMAN\|Q8WUF5_S332 | 0.717447846 | 5.09E-03 | 2.02E-02 |
| TENS4_HUMAN\|Q8IZW8_S253 | 0.717410276 | 9.27E-04 | 1.98E-02 |
| IF4E_HUMAN\|P06730_T203 | 0.717406212 | 3.68E-03 | 1.77E-02 |
| KDM3B_HUMAN\|Q7LBC6_S293 | 0.717371868 | 3.05E-04 | 1.37E-02 |
| ES8L1_HUMAN\|Q8TE68_S182_T187 | 0.717309425 | 4.97E-04 | 1.90E-02 |
| CR025_HUMAN\|Q96B23_S145_S147 | 0.717229898 | 1.11E-03 | 8.60E-03 |
| SCRIB_HUMAN\|Q14160_S1378 | 0.717199269 | 2.04E-03 | 2.59E-02 |
| IF4G2_HUMAN\|P78344_T508 | 0.717191411 | 2.82E-04 | 5.98E-03 |
| ARFG1_HUMAN\|Q8N6T3_T135 | 0.71719093 | 1.80E-03 | 2.15E-02 |
| KAP2_HUMAN\|P13861_S99 | 0.716998825 | 6.40E-04 | 6.83E-03 |
| LIPA1_HUMAN\|Q13136_S666 | 0.716902712 | 2.51E-03 | 3.00E-02 |
| PAXI_HUMAN\|P49023_S85 | 0.716882543 | 1.24E-03 | 9.46E-03 |
| RHG29_HUMAN\|Q52LW3_S1029 | 0.716873979 | 3.70E-03 | 2.49E-02 |
| LAD1_HUMAN\|O00515_S512 | 0.716790618 | 2.62E-03 | 1.20E-02 |
| KDM3B_HUMAN\|Q7LBC6_S289 | 0.716677922 | 2.55E-03 | 1.64E-02 |
| IRS2_HUMAN\|Q9Y4H2_S1100 | 0.716635823 | 6.61E-04 | 4.35E-03 |
| CCNL2_HUMAN\|Q96S94_S330 | 0.716538256 | 4.98E-04 | 3.16E-03 |
| CDK13_HUMAN\|Q14004_T1056 | 0.716462754 | 4.49E-03 | 2.88E-02 |
| AFAD_HUMAN\|P55196_S1173_S1182 | 0.716450539 | 3.34E-03 | 1.45E-02 |
| ARH40_HUMAN\|Q8TER5_S262 | 0.716392107 | 1.01E-03 | 1.56E-02 |
| SSH3_HUMAN\|Q8TE77_S649_S653 | 0.716334749 | 9.40E-04 | 5.77E-03 |
| SCRIB_HUMAN\|Q14160_S1486 | 0.716260138 | 3.31E-03 | 2.00E-02 |
| SQSTM_HUMAN\|Q13501_T269 | 0.71625509 | 2.80E-03 | 3.17E-02 |
| SPTB2_HUMAN\|Q01082_S2341 | 0.716241845 | 8.27E-05 | 2.10E-03 |
| ZYX_HUMAN\|Q15942_S259_S267 | 0.71621976 | 4.16E-03 | 4.34E-02 |

| | | | |
|---|---|---|---|
| DOCK6_HUMAN\|Q96HP0_S882 | 0.716082759 | 7.95E-03 | 3.35E-02 |
| CNOT2_HUMAN\|Q9NZN8_S165 | 0.71607392 | 1.14E-03 | 1.03E-02 |
| K1671_HUMAN\|Q9BY89_S128 | 0.715908744 | 4.98E-04 | 4.69E-02 |
| GGYF2_HUMAN\|Q6Y7W6_S26 | 0.715872685 | 4.45E-04 | 8.61E-03 |
| TX1B3_HUMAN\|O14907_S124 | 0.715854827 | 8.82E-04 | 7.21E-03 |
| CBX3_HUMAN\|Q13185_S99 | 0.71574111 | 3.36E-04 | 7.28E-03 |
| SRGP1_HUMAN\|Q7Z6B7_S906 | 0.715693154 | 4.00E-03 | 4.05E-02 |
| ML12A_HUMAN\|P19105_T18 | 0.715689807 | 5.22E-03 | 3.16E-02 |
| AHNK_HUMAN\|Q09666_S5763 | 0.715636896 | 2.54E-04 | 3.40E-02 |
| PKHG3_HUMAN\|A1L390_S1079 | 0.715626091 | 4.44E-03 | 3.73E-02 |
| CNKR1_HUMAN\|Q969H4_S314 | 0.715485087 | 1.39E-03 | 2.01E-02 |
| CDK13_HUMAN\|Q14004_T1056 | 0.715474759 | 4.35E-03 | 2.96E-02 |
| OSTF1_HUMAN\|Q92882_S202_S213 | 0.715440807 | 5.18E-04 | 1.37E-02 |
| CUL4B_HUMAN\|Q13620_T49 | 0.715408023 | 6.45E-03 | 4.85E-02 |
| ZCHC8_HUMAN\|Q6NZY4_S500_T492_T496 | 0.715398347 | 5.94E-03 | 2.27E-02 |
| AF1L2_HUMAN\|Q8N4X5_S640 | 0.715383123 | 2.91E-04 | 3.75E-03 |
| PER1_HUMAN\|O15534_S1007 | 0.715345023 | 1.81E-03 | 8.87E-03 |
| GMIP_HUMAN\|Q9P107_S914_S919 | 0.715286845 | 1.48E-03 | 1.25E-02 |
| MISP_HUMAN\|Q8IVT2_T377 | 0.715283903 | 3.80E-03 | 2.67E-02 |
| T106B_HUMAN\|Q9NUM4_S33 | 0.715232841 | 1.76E-03 | 1.41E-02 |
| VIP2_HUMAN\|O43314_S1006 | 0.715232003 | 7.49E-04 | 7.82E-03 |
| SIK3_HUMAN\|Q9Y2K2_S493 | 0.715051222 | 5.47E-04 | 1.42E-02 |
| CF132_HUMAN\|Q5T0Z8_S738 | 0.715036072 | 3.45E-04 | 4.19E-03 |
| K2C5_HUMAN\|P13647_S571 | 0.715022007 | 1.41E-03 | 1.89E-02 |
| RFIP5_HUMAN\|Q9BXF6_S396 | 0.715005384 | 2.52E-03 | 4.00E-02 |
| PRSR2_HUMAN\|Q86WR7_S18 | 0.714987561 | 1.41E-04 | 2.17E-03 |
| SRRM2_HUMAN\|Q9UQ35_T1492 | 0.714967471 | 2.44E-03 | 1.27E-02 |
| MA7D1_HUMAN\|Q3KQU3_S446 | 0.714925741 | 9.28E-04 | 1.47E-02 |
| ZCCHV_HUMAN\|Q7Z2W4_T380 | 0.71492479 | 3.63E-03 | 1.56E-02 |
| AFAD_HUMAN\|P55196_S1172_S1173_S1182 | 0.71479759 | 6.30E-04 | 1.10E-02 |
| ZC3HD_HUMAN\|Q5T200_S1292 | 0.714781828 | 8.11E-04 | 5.18E-03 |

| | | | |
|---|---|---|---|
| KLC1_HUMAN\|Q07866-4_S631 | 0.714729423 | 5.51E-04 | 1.42E-02 |
| TB182_HUMAN\|Q9C0C2_S1158 | 0.714681492 | 5.92E-03 | 3.67E-02 |
| PKHA4_HUMAN\|Q9H4M7_S194 | 0.714676562 | 5.20E-03 | 3.16E-02 |
| SSH2_HUMAN\|Q76I76_T486 | 0.714620353 | 1.77E-03 | 1.49E-02 |
| CAMP3_HUMAN\|Q9P1Y5_S553 | 0.714607496 | 1.15E-04 | 4.35E-03 |
| SGTA_HUMAN\|O43765_S77_T81 | 0.714585147 | 2.87E-03 | 1.29E-02 |
| ABLM1_HUMAN\|O14639_S655 | 0.714521494 | 3.30E-03 | 1.44E-02 |
| YBOX1_HUMAN\|P67809_S165 | 0.714503 | 1.07E-04 | 1.02E-03 |
| KLC1_HUMAN\|Q07866-4_S600 | 0.714497783 | 3.51E-04 | 2.42E-03 |
| ASPC1_HUMAN\|Q9BZE9_S275 | 0.714422995 | 2.02E-03 | 2.34E-02 |
| SRRM1_HUMAN\|Q8IYB3_S874 | 0.714406575 | 6.59E-03 | 3.97E-02 |
| IF2B3_HUMAN\|O00425_S180_S184 | 0.714345059 | 1.01E-04 | 9.20E-04 |
| 4ET_HUMAN\|Q9NRA8_S587 | 0.714239872 | 4.48E-03 | 1.84E-02 |
| YJ005_HUMAN\|Q6ZSR9_S93 | 0.714198976 | 8.67E-05 | 2.18E-03 |
| SHRM2_HUMAN\|Q13796_S1039 | 0.714087498 | 3.42E-05 | 3.37E-03 |
| ABLM1_HUMAN\|O14639_S431_T429 | 0.71407376 | 1.37E-03 | 3.40E-02 |
| LUZP1_HUMAN\|Q86V48_S575 | 0.714019216 | 3.57E-04 | 2.45E-03 |
| NUP53_HUMAN\|Q8NFH5_T280 | 0.714009931 | 2.59E-04 | 1.90E-03 |
| LAD1_HUMAN\|O00515_S38 | 0.714003219 | 3.24E-03 | 1.42E-02 |
| IF2B3_HUMAN\|O00425_S179_S180 | 0.713993739 | 5.18E-05 | 5.34E-04 |
| PDLI4_HUMAN\|P50479_T114 | 0.713979448 | 1.62E-03 | 1.40E-02 |
| RAE1_HUMAN\|P24386_S651 | 0.71397911 | 8.52E-04 | 1.37E-02 |
| TWF1_HUMAN\|Q12792_T349 | 0.713971747 | 6.18E-04 | 6.56E-03 |
| ACINU_HUMAN\|Q9UKV3_S243 | 0.713913736 | 9.49E-04 | 5.28E-03 |
| PLEC_HUMAN\|Q15149_Y4615 | 0.713865573 | 3.29E-03 | 2.35E-02 |
| SWAHC_HUMAN\|Q53LP3_S212 | 0.713864963 | 4.66E-04 | 1.26E-02 |
| DBNL_HUMAN\|Q9UJU6_T271 | 0.713797505 | 2.88E-03 | 2.12E-02 |
| WWP2_HUMAN\|O00308_S211 | 0.713758133 | 9.96E-04 | 9.42E-03 |
| FGD6_HUMAN\|Q6ZV73_T693 | 0.71374301 | 3.81E-03 | 2.54E-02 |
| LAP2A_HUMAN\|P42166_S159 | 0.71373201 | 3.01E-03 | 2.19E-02 |
| CLAP2_HUMAN\|O75122_S1029 | 0.713657644 | 6.31E-03 | 2.77E-02 |

| | | | |
|---|---|---|---|
| CLASR_HUMAN\|Q8N2M8_S547 | 0.713634447 | 3.12E-04 | 6.74E-03 |
| F122B_HUMAN\|Q7Z309_S115_S119 | 0.713566541 | 9.38E-04 | 1.15E-02 |
| IRS2_HUMAN\|Q9Y4H2_S1203 | 0.713549167 | 2.26E-04 | 2.54E-03 |
| RAI14_HUMAN\|Q9P0K7_T251 | 0.713513967 | 6.82E-03 | 2.94E-02 |
| NOP2_HUMAN\|P46087_T605 | 0.713420612 | 2.72E-03 | 1.72E-02 |
| AMOL2_HUMAN\|Q9Y2J4_S759 | 0.713299443 | 9.70E-04 | 5.92E-03 |
| SI1L3_HUMAN\|O60292_S172 | 0.713283987 | 5.31E-05 | 5.39E-04 |
| PHF14_HUMAN\|O94880_S290 | 0.713273089 | 3.19E-03 | 3.21E-02 |
| AHNK_HUMAN\|Q09666_T566 | 0.713265463 | 2.30E-03 | 2.80E-02 |
| PEPL_HUMAN\|O60437_S1657 | 0.713153251 | 2.13E-03 | 1.13E-02 |
| RABE1_HUMAN\|Q15276_S410_T408 | 0.713125023 | 1.29E-03 | 6.75E-03 |
| DBNL_HUMAN\|Q9UJU6_S275 | 0.713070783 | 1.42E-02 | 4.57E-02 |
| MA7D1_HUMAN\|Q3KQU3_S446 | 0.713037723 | 8.59E-04 | 4.86E-03 |
| AAPK1_HUMAN\|Q13131_S496 | 0.712978843 | 2.34E-03 | 1.09E-02 |
| PCM1_HUMAN\|Q15154_S872 | 0.712964599 | 9.36E-04 | 9.41E-03 |
| SWAHC_HUMAN\|Q53LP3_S212 | 0.71293924 | 4.48E-04 | 5.13E-03 |
| MICLK_HUMAN\|Q6ZW33_S59_T62 | 0.712803867 | 6.96E-04 | 4.10E-03 |
| RGS3_HUMAN\|P49796_S946_T941 | 0.71280326 | 1.38E-03 | 7.90E-03 |
| KLC2_HUMAN\|Q9H0B6_S611 | 0.712713803 | 4.26E-03 | 1.77E-02 |
| PHLB1_HUMAN\|Q86UU1_S461 | 0.712711104 | 2.46E-04 | 3.30E-03 |
| RRP1B_HUMAN\|Q14684_S706 | 0.71268871 | 5.68E-03 | 4.76E-02 |
| CNOT2_HUMAN\|Q9NZN8_S165 | 0.712681425 | 9.96E-04 | 5.50E-03 |
| PRC2A_HUMAN\|P48634_S1085 | 0.712679509 | 5.01E-03 | 4.06E-02 |
| SRC8_HUMAN\|Q14247_T399_T411 | 0.712637025 | 2.17E-03 | 4.52E-02 |
| DDI2_HUMAN\|Q5TDH0_S106 | 0.712631041 | 9.33E-04 | 2.82E-02 |
| COHA1_HUMAN\|Q9UMD9_T387 | 0.712589956 | 2.10E-03 | 1.00E-02 |
| SNP23_HUMAN\|O00161_S20 | 0.712566947 | 1.20E-02 | 4.65E-02 |
| ATAT_HUMAN\|Q5SQI0_S315 | 0.712492591 | 3.70E-04 | 7.81E-03 |
| SI1L1_HUMAN\|O43166_S1647_S1650 | 0.71243827 | 5.97E-03 | 2.63E-02 |
| AHNK_HUMAN\|Q09666_S177 | 0.712425974 | 4.13E-04 | 1.64E-02 |
| MFAP1_HUMAN\|P55081_S80 | 0.712370879 | 5.66E-04 | 1.44E-02 |

| | | | |
|---|---|---|---|
| SF3B1_HUMAN\|O75533_S194 | 0.712322709 | 4.60E-03 | 2.93E-02 |
| CYBR1_HUMAN\|Q53TN4_T285 | 0.712313868 | 1.14E-02 | 4.50E-02 |
| PHF14_HUMAN\|O94880_S290_T287 | 0.712212499 | 6.78E-03 | 2.51E-02 |
| DESP_HUMAN\|P15924_S2815 | 0.712093094 | 1.17E-03 | 3.18E-02 |
| TBCD5_HUMAN\|Q92609_S43 | 0.712002459 | 4.32E-03 | 2.79E-02 |
| HEAT6_HUMAN\|Q6AI08_S643 | 0.711947813 | 2.44E-04 | 5.72E-03 |
| K1522_HUMAN\|Q9P206_S327 | 0.711904506 | 4.98E-04 | 5.59E-03 |
| NU107_HUMAN\|P57740_S37 | 0.71181647 | 2.52E-03 | 2.51E-02 |
| PTN14_HUMAN\|Q15678_S578 | 0.711777758 | 3.94E-03 | 3.73E-02 |
| NIBL1_HUMAN\|Q96TA1_S696 | 0.711689225 | 2.43E-03 | 1.12E-02 |
| PHLB2_HUMAN\|Q86SQ0_T404 | 0.711677334 | 4.65E-03 | 1.89E-02 |
| SPTB2_HUMAN\|Q01082-3_S10 | 0.711610886 | 3.39E-03 | 4.81E-02 |
| VIME_HUMAN\|P08670_S459 | 0.711609043 | 1.16E-03 | 3.17E-02 |
| RBP56_HUMAN\|Q92804_T222 | 0.71152044 | 7.37E-03 | 2.68E-02 |
| TPR_HUMAN\|P12270_S2155 | 0.711417118 | 4.25E-03 | 4.38E-02 |
| SCRIB_HUMAN\|Q14160_S1378 | 0.711327565 | 1.63E-03 | 8.16E-03 |
| IWS1_HUMAN\|Q96ST2_S720 | 0.7112953 | 4.10E-04 | 4.03E-03 |
| PPHLN_HUMAN\|Q8NEY8_S140 | 0.711288256 | 6.45E-04 | 1.16E-02 |
| DESP_HUMAN\|P15924_S2820_S2821 | 0.711283283 | 4.80E-03 | 1.94E-02 |
| PKP3_HUMAN\|Q9Y446_S238 | 0.711037437 | 2.80E-04 | 3.68E-03 |
| PDLI5_HUMAN\|Q96HC4_S111 | 0.710912609 | 2.73E-03 | 3.17E-02 |
| CCDC9_HUMAN\|Q9Y3X0_S376 | 0.710905343 | 1.72E-03 | 2.16E-02 |
| SND1_HUMAN\|Q7KZF4_S426 | 0.710689531 | 4.37E-03 | 1.81E-02 |
| PKHA2_HUMAN\|Q9HB19_S184 | 0.710682531 | 3.93E-04 | 2.62E-03 |
| AFAD_HUMAN\|P55196_S1696 | 0.710663476 | 2.82E-03 | 4.28E-02 |
| HIRP3_HUMAN\|Q9BW71_S196_S199 | 0.710633072 | 1.39E-03 | 1.26E-02 |
| PARP9_HUMAN\|Q8IXQ6_S851 | 0.710606131 | 6.99E-04 | 6.06E-03 |
| AHNK_HUMAN\|Q09666_T566 | 0.710596986 | 2.09E-03 | 1.11E-02 |
| EPN2_HUMAN\|O95208-2_S192_Y196 | 0.710509189 | 1.21E-02 | 4.00E-02 |
| XRCC4_HUMAN\|Q13426_S328 | 0.710498213 | 6.54E-04 | 8.80E-03 |
| NCOA2_HUMAN\|Q15596_S699 | 0.710479563 | 7.28E-03 | 2.66E-02 |

| | | | |
|---|---|---|---|
| CO052_HUMAN\|Q6ZUT6_S193 | 0.710454779 | 4.22E-03 | 2.75E-02 |
| YLPM1_HUMAN\|P49750_S634 | 0.71038899 | 2.41E-04 | 5.43E-03 |
| ZC11A_HUMAN\|O75152_S132 | 0.710387645 | 2.76E-04 | 4.58E-03 |
| RUFY3_HUMAN\|Q7L099-3_S618 | 0.710354282 | 1.68E-04 | 1.45E-03 |
| F193A_HUMAN\|P78312_S1144 | 0.710330252 | 2.78E-03 | 4.23E-02 |
| K2C5_HUMAN\|P13647_S571 | 0.710299011 | 1.17E-03 | 1.74E-02 |
| TGS1_HUMAN\|Q96RS0_S89 | 0.710292663 | 4.98E-03 | 1.99E-02 |
| MAP7_HUMAN\|Q14244_S315 | 0.710206339 | 1.84E-03 | 2.25E-02 |
| PKHG3_HUMAN\|A1L390_S1079 | 0.710052856 | 3.73E-03 | 2.16E-02 |
| K1C14_HUMAN\|P02533_S53 | 0.710024228 | 1.37E-03 | 1.25E-02 |
| BRAF_HUMAN\|P15056_S446 | 0.709935834 | 8.85E-03 | 4.11E-02 |
| TWF1_HUMAN\|Q12792_T349 | 0.709863354 | 5.21E-04 | 5.77E-03 |
| FA83H_HUMAN\|Q6ZRV2_S828 | 0.709839038 | 4.34E-04 | 5.03E-03 |
| KI21A_HUMAN\|Q7Z4S6_S855 | 0.709837587 | 2.33E-03 | 3.80E-02 |
| IRS2_HUMAN\|Q9Y4H2_S679 | 0.709724346 | 2.05E-03 | 2.35E-02 |
| SMCA4_HUMAN\|P51532_S1570_S1575_S1586 | 0.709594178 | 4.06E-03 | 1.70E-02 |
| REPS1_HUMAN\|Q96D71_S162_S170_S174 | 0.709570387 | 1.82E-03 | 2.24E-02 |
| PHLB2_HUMAN\|Q86SQ0_S402 | 0.709451706 | 2.18E-03 | 3.61E-02 |
| CDK9_HUMAN\|P50750_T186 | 0.709374781 | 3.21E-03 | 1.41E-02 |
| 4EBP1_HUMAN\|Q13541_S44 | 0.709334636 | 2.78E-03 | 1.99E-02 |
| VIME_HUMAN\|P08670_T458 | 0.709324688 | 1.09E-03 | 3.10E-02 |
| ARH40_HUMAN\|Q8TER5_T371 | 0.709147461 | 1.93E-04 | 1.12E-02 |
| K1671_HUMAN\|Q9BY89_S127 | 0.70905082 | 1.19E-03 | 1.12E-02 |
| CSKI2_HUMAN\|Q8WXE0_S403_S406 | 0.709034734 | 4.87E-05 | 2.04E-03 |
| LAR4B_HUMAN\|Q92615_S731_S736 | 0.709020424 | 3.81E-03 | 2.67E-02 |
| ENOA_HUMAN\|P06733_S40 | 0.708959623 | 9.00E-03 | 3.69E-02 |
| APBB2_HUMAN\|Q92870_S409_S412 | 0.708891158 | 1.35E-03 | 2.58E-02 |
| OPTN_HUMAN\|Q96CV9_T204 | 0.708859276 | 4.62E-04 | 1.82E-02 |
| CE170_HUMAN\|Q5SW79_S630_S636 | 0.708779614 | 8.65E-04 | 1.42E-02 |
| CSKI2_HUMAN\|Q8WXE0_S396_S403 | 0.708777163 | 3.59E-03 | 4.87E-02 |
| ACAP2_HUMAN\|Q15057_S384_T382 | 0.708728405 | 1.69E-05 | 8.55E-04 |

| | | | |
|---|---|---|---|
| EFHD2_HUMAN\|Q96C19_S76 | 0.70868593 | 4.52E-03 | 4.59E-02 |
| PEAK1_HUMAN\|Q9H792_S1036 | 0.708678423 | 4.15E-04 | 8.49E-03 |
| SNAPN_HUMAN\|O95295_S133 | 0.708628221 | 2.17E-04 | 1.77E-03 |
| COBL1_HUMAN\|Q53SF7_S294_T298 | 0.708471332 | 3.67E-03 | 4.92E-02 |
| EDC3_HUMAN\|Q96F86_S131 | 0.708468474 | 2.28E-04 | 5.13E-03 |
| PRR12_HUMAN\|Q9ULL5_S44 | 0.708422134 | 3.20E-03 | 4.63E-02 |
| RABE1_HUMAN\|Q15276_S410_T408 | 0.708398658 | 1.08E-03 | 6.46E-03 |
| RHG23_HUMAN\|Q9P227_S351 | 0.708377098 | 2.38E-03 | 1.85E-02 |
| CSKI2_HUMAN\|Q8WXE0_S406 | 0.708231204 | 1.70E-04 | 1.36E-03 |
| OXR1_HUMAN\|Q8N573_S201_S202 | 0.708228878 | 9.35E-04 | 2.82E-02 |
| SNX29_HUMAN\|Q8TEQ0_S810 | 0.708157928 | 3.63E-03 | 2.45E-02 |
| UBP5_HUMAN\|P45974_S779 | 0.708157617 | 9.98E-03 | 4.02E-02 |
| R113A_HUMAN\|O15541_S329 | 0.708145451 | 2.63E-04 | 1.92E-03 |
| OSB10_HUMAN\|Q9BXB5_S32 | 0.70811531 | 2.55E-03 | 1.17E-02 |
| RABE1_HUMAN\|Q15276_T408 | 0.708113039 | 4.28E-04 | 4.17E-03 |
| ZCHC7_HUMAN\|Q8N3Z6_S482 | 0.708099943 | 4.36E-03 | 4.24E-02 |
| PKHA4_HUMAN\|Q9H4M7_S194 | 0.708091663 | 4.22E-03 | 4.36E-02 |
| SEPT9_HUMAN\|Q9UHD8_T42 | 0.708075929 | 1.23E-03 | 1.77E-02 |
| PLCH2_HUMAN\|O75038_S562 | 0.708072877 | 1.53E-04 | 1.25E-03 |
| NHRF1_HUMAN\|O14745_S280 | 0.708064709 | 1.53E-03 | 3.85E-02 |
| GRIN1_HUMAN\|Q7Z2K8_S615 | 0.707991359 | 1.97E-03 | 1.54E-02 |
| SRGP3_HUMAN\|O43295_S858 | 0.707984945 | 3.11E-03 | 2.19E-02 |
| MAP4_HUMAN\|P27816_S928 | 0.707977706 | 2.83E-03 | 3.09E-02 |
| PDLI4_HUMAN\|P50479_S112 | 0.707970773 | 1.95E-03 | 1.60E-02 |
| KI26B_HUMAN\|Q2KJY2_S1006_S1018_S1021 | 0.707963812 | 3.28E-04 | 4.04E-03 |
| ABI2_HUMAN\|Q9NYB9_S368 | 0.707954033 | 2.07E-04 | 5.16E-03 |
| YBOX1_HUMAN\|P67809_S174_S176 | 0.707952487 | 1.36E-03 | 1.86E-02 |
| NBN_HUMAN\|O60934_T337 | 0.707922869 | 5.01E-03 | 2.00E-02 |
| TP53B_HUMAN\|Q12888_S366_T357 | 0.707897335 | 3.17E-03 | 1.93E-02 |
| ASHWN_HUMAN\|Q9BVC5_S193 | 0.707856451 | 2.51E-04 | 3.36E-03 |
| EFHD2_HUMAN\|Q96C19_S74 | 0.707853942 | 2.73E-03 | 1.39E-02 |

| | | | |
|---|---|---|---|
| MAST4_HUMAN\|O15021_S1387_S1394 | 0.707604263 | 8.33E-04 | 1.12E-02 |
| DEN4C_HUMAN\|Q5VZ89_S1184 | 0.707591866 | 1.06E-03 | 5.79E-03 |
| G3BP1_HUMAN\|Q13283_S231 | 0.707591531 | 6.49E-04 | 6.77E-03 |
| SRRM2_HUMAN\|Q9UQ35_S2272 | 0.707545767 | 2.17E-04 | 1.09E-02 |
| TFE3_HUMAN\|P19532_S548_S556 | 0.707509377 | 4.43E-03 | 3.72E-02 |
| SRSF6_HUMAN\|Q13247_S301 | 0.707485549 | 5.01E-03 | 3.98E-02 |
| MK14_HUMAN\|Q16539_Y182 | 0.707258071 | 4.21E-03 | 3.59E-02 |
| GYS1_HUMAN\|P13807_S727 | 0.707128218 | 9.10E-04 | 2.73E-02 |
| DLG5_HUMAN\|Q8TDM6_S1264 | 0.70710675 | 2.24E-03 | 1.70E-02 |
| FA83H_HUMAN\|Q6ZRV2_S914 | 0.707105622 | 2.23E-03 | 2.76E-02 |
| GOGA4_HUMAN\|Q13439_T31 | 0.707080869 | 2.81E-04 | 5.97E-03 |
| PDLI2_HUMAN\|Q96JY6_S74 | 0.70706322 | 1.19E-04 | 1.05E-03 |
| S38A2_HUMAN\|Q96QD8_S19_S21 | 0.706990966 | 9.50E-03 | 3.87E-02 |
| LIPA1_HUMAN\|Q13136_S666_S668 | 0.706989632 | 1.25E-03 | 6.58E-03 |
| SPAST_HUMAN\|Q9UBP0_S270 | 0.706945927 | 6.15E-04 | 4.11E-03 |
| CR025_HUMAN\|Q96B23_S66 | 0.706925806 | 6.52E-04 | 2.20E-02 |
| PAXI_HUMAN\|P49023_S258 | 0.706916318 | 7.54E-04 | 7.59E-03 |
| HEAT6_HUMAN\|Q6AI08_S643 | 0.706849911 | 1.91E-04 | 2.68E-03 |
| KIF1C_HUMAN\|O43896_S817 | 0.706822817 | 1.74E-03 | 1.40E-02 |
| SAFB1_HUMAN\|Q15424_S604 | 0.706669438 | 1.37E-04 | 3.55E-03 |
| IF4B_HUMAN\|P23588_S597 | 0.706539879 | 1.14E-02 | 4.50E-02 |
| NELFE_HUMAN\|P18615_S353 | 0.706526121 | 9.38E-05 | 3.66E-03 |
| FA83H_HUMAN\|Q6ZRV2_S936 | 0.706509784 | 5.71E-03 | 2.20E-02 |
| PHLB2_HUMAN\|Q86SQ0_S402 | 0.706504629 | 1.97E-03 | 2.35E-02 |
| LMO7_HUMAN\|Q8WWI1_S1586 | 0.706404028 | 1.15E-03 | 1.73E-02 |
| LIPB1_HUMAN\|Q86W92_S579 | 0.706353048 | 6.59E-04 | 3.92E-03 |
| TP53B_HUMAN\|Q12888_S359_S366 | 0.706302242 | 1.05E-02 | 3.61E-02 |
| SSH2_HUMAN\|Q76I76_T486 | 0.706299024 | 1.29E-03 | 1.89E-02 |
| UBP8_HUMAN\|P40818_T577 | 0.706211356 | 3.08E-04 | 2.18E-03 |
| ES8L1_HUMAN\|Q8TE68_S182_T187 | 0.706209695 | 3.01E-04 | 3.83E-03 |
| TB182_HUMAN\|Q9C0C2_T875 | 0.706207503 | 1.33E-03 | 1.75E-02 |

| | | | |
|---|---|---|---|
| GGYF2_HUMAN\|Q6Y7W6_S26 | 0.706193716 | 2.92E-04 | 1.35E-02 |
| IASPP_HUMAN\|Q8WUF5_S113 | 0.706143129 | 1.17E-03 | 2.34E-02 |
| SKIV2_HUMAN\|Q15477_S255 | 0.706122188 | 2.37E-03 | 2.32E-02 |
| IF4B_HUMAN\|P23588_S497_S504_T500 | 0.706073788 | 5.77E-03 | 4.84E-02 |
| AFF4_HUMAN\|Q9UHB7_S836 | 0.706057204 | 4.35E-03 | 3.98E-02 |
| PARP4_HUMAN\|Q9UKK3_S1722 | 0.705993611 | 2.05E-03 | 3.47E-02 |
| G3P_HUMAN\|P04406_S210 | 0.705989714 | 4.17E-04 | 2.74E-03 |
| YBOX3_HUMAN\|P16989_S369_S370 | 0.705984796 | 2.99E-03 | 2.18E-02 |
| NIBL1_HUMAN\|Q96TA1_S692 | 0.7059586 | 1.22E-03 | 9.39E-03 |
| OXR1_HUMAN\|Q8N573_S201_S202 | 0.705904324 | 8.53E-04 | 1.30E-02 |
| FOXJ3_HUMAN\|Q9UPW0_S223 | 0.705887141 | 1.50E-03 | 7.66E-03 |
| NDRG1_HUMAN\|Q92597_S336_T328 | 0.705875367 | 7.22E-03 | 3.10E-02 |
| PDLI2_HUMAN\|Q96JY6_S74 | 0.70586649 | 1.13E-04 | 1.06E-03 |
| AHNK_HUMAN\|Q09666_S177 | 0.705821248 | 3.06E-04 | 6.43E-03 |
| CTNB1_HUMAN\|P35222_S552 | 0.705811176 | 3.30E-03 | 4.73E-02 |
| LAD1_HUMAN\|O00515_T358 | 0.705786287 | 3.17E-03 | 1.93E-02 |
| PRSR2_HUMAN\|Q86WR7_S18 | 0.705775039 | 8.76E-05 | 2.63E-03 |
| ZC3H4_HUMAN\|Q9UPT8_S1114_T1106 | 0.705597088 | 2.88E-03 | 1.29E-02 |
| FA83H_HUMAN\|Q6ZRV2_S647 | 0.705587041 | 1.80E-04 | 1.54E-03 |
| UN13B_HUMAN\|O14795_S254 | 0.705582613 | 7.17E-03 | 3.50E-02 |
| PRSR2_HUMAN\|Q86WR7_S18 | 0.705531332 | 8.65E-05 | 2.99E-03 |
| SRSF6_HUMAN\|Q13247_S303 | 0.705528947 | 5.45E-03 | 2.86E-02 |
| ANXA2_HUMAN\|P07355_T19 | 0.705498832 | 1.30E-04 | 3.98E-03 |
| RAI1_HUMAN\|Q7Z5J4_S1108 | 0.705450638 | 9.53E-04 | 1.50E-02 |
| PPR35_HUMAN\|Q8TAP8_S52 | 0.705403259 | 4.19E-03 | 4.13E-02 |
| GRDN_HUMAN\|Q3V6T2_S1702 | 0.705336127 | 2.88E-03 | 1.45E-02 |
| SPIR1_HUMAN\|Q08AE8_T509 | 0.705176979 | 1.97E-03 | 2.35E-02 |
| SAV1_HUMAN\|Q9H4B6_S27 | 0.705168546 | 4.74E-04 | 5.39E-03 |
| ARH40_HUMAN\|Q8TER5_S262 | 0.705068655 | 6.45E-04 | 3.87E-03 |
| AKAP2_HUMAN\|Q9Y2D5_S720 | 0.704895255 | 9.51E-04 | 1.50E-02 |
| TMF1_HUMAN\|P82094_S77 | 0.704882659 | 2.01E-03 | 3.46E-02 |

| | | | |
|---|---|---|---|
| KIZ_HUMAN|Q2M2Z5_S668_T665 | 0.704815425 | 2.71E-04 | 3.57E-03 |
| K1C14_HUMAN|P02533_S432_S435 | 0.704716637 | 1.52E-02 | 4.84E-02 |
| CARL1_HUMAN|Q5VZK9_S968 | 0.704699044 | 1.75E-04 | 1.51E-03 |
| AHNK_HUMAN|Q09666_S511 | 0.704686587 | 6.65E-03 | 3.81E-02 |
| FA83H_HUMAN|Q6ZRV2_S1098 | 0.704682997 | 1.32E-03 | 1.93E-02 |
| LARP4_HUMAN|Q71RC2_S597 | 0.704607389 | 1.96E-03 | 9.46E-03 |
| TCF20_HUMAN|Q9UGU0_S1755_S1757 | 0.704567268 | 2.24E-03 | 2.72E-02 |
| SI1L1_HUMAN|O43166_S1647_S1650 | 0.704508056 | 4.73E-03 | 1.92E-02 |
| K2C6B_HUMAN|P04259_T552 | 0.704470583 | 5.62E-03 | 2.17E-02 |
| SRGP3_HUMAN|O43295_S1073 | 0.704418149 | 1.58E-03 | 8.81E-03 |
| PKP3_HUMAN|Q9Y446_S238 | 0.704389842 | 2.07E-04 | 1.60E-03 |
| DEFI6_HUMAN|Q9H4E7_S597 | 0.704373051 | 1.16E-04 | 3.52E-03 |
| CE170_HUMAN|Q5SW79_S630_S633 | 0.704361661 | 8.05E-04 | 1.83E-02 |
| OSB10_HUMAN|Q9BXB5_S26_S32 | 0.704358075 | 2.92E-04 | 5.25E-03 |
| 2A5D_HUMAN|Q14738_S598 | 0.704357172 | 4.06E-03 | 2.81E-02 |
| LAR4B_HUMAN|Q92615_T732 | 0.704313795 | 1.24E-04 | 1.90E-03 |
| NU214_HUMAN|P35658_S433_T437 | 0.704306122 | 1.08E-02 | 4.81E-02 |
| P4K2A_HUMAN|Q9BTU6_S462_S468 | 0.704281435 | 8.54E-04 | 1.14E-02 |
| ANR17_HUMAN|O75179_S2041_S2045 | 0.704254807 | 3.93E-03 | 1.67E-02 |
| AHNK_HUMAN|Q09666_S212_S216 | 0.704245299 | 7.32E-04 | 1.26E-02 |
| CTU2_HUMAN|Q2VPK5_S419 | 0.704201908 | 2.39E-04 | 5.31E-03 |
| TB182_HUMAN|Q9C0C2_S936 | 0.704165495 | 1.07E-03 | 5.83E-03 |
| ARFG1_HUMAN|Q8N6T3_T135 | 0.704155021 | 1.11E-03 | 5.99E-03 |
| VIME_HUMAN|P08670_T458 | 0.704037457 | 8.81E-04 | 1.43E-02 |
| LARP4_HUMAN|Q71RC2_S597 | 0.704016993 | 1.92E-03 | 2.33E-02 |
| RIOK1_HUMAN|Q9BRS2_S22 | 0.70400889 | 2.64E-05 | 6.06E-04 |
| AKAP2_HUMAN|Q9Y2D5_S152_S155 | 0.703994982 | 1.49E-03 | 1.32E-02 |
| GOGA4_HUMAN|Q13439_T31 | 0.703917013 | 2.42E-04 | 1.92E-03 |
| IF4G1_HUMAN|Q04637_S1231 | 0.703909571 | 2.04E-03 | 4.39E-02 |
| EPN2_HUMAN|O95208_S173 | 0.703909295 | 4.91E-03 | 3.22E-02 |
| ANR54_HUMAN|Q6NXT1_S63 | 0.703875532 | 3.85E-03 | 3.93E-02 |

| | | | |
|---|---|---|---|
| ATX2_HUMAN\|Q99700_T741 | 0.703850455 | 1.56E-05 | 2.07E-04 |
| LMNB2_HUMAN\|Q03252_S424 | 0.703707064 | 6.44E-03 | 3.73E-02 |
| IF4B_HUMAN\|P23588_S497_S498_S504 | 0.703658905 | 6.94E-04 | 1.15E-02 |
| IWS1_HUMAN\|Q96ST2_S720_T725 | 0.703569606 | 7.42E-03 | 4.13E-02 |
| K2C5_HUMAN\|P13647_S75 | 0.703485511 | 9.36E-03 | 3.29E-02 |
| TB10B_HUMAN\|Q4KMP7_S22 | 0.7034841 | 9.70E-04 | 1.51E-02 |
| PTN14_HUMAN\|Q15678_S594 | 0.703467301 | 2.55E-05 | 5.97E-04 |
| ZC3H1_HUMAN\|O60293_S805_S809 | 0.703417102 | 6.46E-03 | 4.75E-02 |
| IRS2_HUMAN\|Q9Y4H2_T350 | 0.703414629 | 5.83E-04 | 2.08E-02 |
| PDLI2_HUMAN\|Q96JY6_S134 | 0.703399463 | 6.34E-05 | 2.03E-03 |
| 1A01_HUMAN\|P30443_S356_S359 | 0.703392546 | 5.26E-03 | 3.38E-02 |
| DLG5_HUMAN\|Q8TDM6_S1264 | 0.703378952 | 1.98E-03 | 1.06E-02 |
| ML12A_HUMAN\|P19105_S19 | 0.703369643 | 1.22E-03 | 1.77E-02 |
| IPP2M_HUMAN\|Q6NXS1_S122 | 0.70336067 | 3.37E-03 | 3.10E-02 |
| ARG28_HUMAN\|Q8N1W1_S476_S486 | 0.703308073 | 4.22E-03 | 1.98E-02 |
| SRPK2_HUMAN\|P78362_S494_S497 | 0.703168593 | 1.42E-03 | 1.83E-02 |
| RAI14_HUMAN\|Q9P0K7_T251 | 0.703052315 | 5.01E-03 | 2.00E-02 |
| AHNK_HUMAN\|Q09666_S5448 | 0.703050624 | 1.22E-03 | 7.16E-03 |
| RL15_HUMAN\|P61313_S34 | 0.702986871 | 2.27E-03 | 1.19E-02 |
| AHNK_HUMAN\|Q09666_S5110 | 0.702965904 | 3.14E-03 | 4.58E-02 |
| RB3GP_HUMAN\|Q15042_S537 | 0.702797424 | 3.60E-04 | 1.07E-02 |
| PML_HUMAN\|P29590_T867 | 0.702670472 | 1.95E-03 | 2.53E-02 |
| ARFG1_HUMAN\|Q8N6T3_T135 | 0.702595052 | 1.05E-03 | 1.59E-02 |
| TFE3_HUMAN\|P19532_S564_S568 | 0.702436146 | 3.84E-05 | 1.58E-03 |
| UBP24_HUMAN\|Q9UPU5_S1143 | 0.702399261 | 1.59E-03 | 2.12E-02 |
| PKHF2_HUMAN\|Q9H8W4_S239 | 0.702389341 | 1.89E-03 | 1.98E-02 |
| ARP19_HUMAN\|P56211_S62 | 0.70236277 | 1.70E-03 | 2.15E-02 |
| KLC4_HUMAN\|Q9NSK0_S611 | 0.702336728 | 2.41E-03 | 3.91E-02 |
| ANR54_HUMAN\|Q6NXT1_S63 | 0.702325367 | 3.66E-03 | 1.56E-02 |
| SPT5H_HUMAN\|O00267_S666 | 0.702305349 | 2.78E-04 | 6.28E-03 |
| MTCL1_HUMAN\|Q9Y4B5_S77 | 0.70229444 | 4.73E-03 | 4.45E-02 |

| | | | |
|---|---|---|---|
| BORG5_HUMAN\|Q00587_S303 | 0.702224998 | 9.90E-04 | 1.45E-02 |
| AHNK_HUMAN\|Q09666_S5752 | 0.702089391 | 2.36E-04 | 3.40E-02 |
| K2C5_HUMAN\|P13647_S18 | 0.702060723 | 2.83E-03 | 3.25E-02 |
| ZC3HD_HUMAN\|Q5T200_S64 | 0.702021428 | 3.09E-03 | 1.36E-02 |
| PITM3_HUMAN\|Q9BZ71_S504 | 0.701948906 | 1.80E-03 | 2.15E-02 |
| UBP2L_HUMAN\|Q14157_S467_S477 | 0.701916617 | 1.43E-03 | 2.03E-02 |
| KC1D_HUMAN\|P48730_S383 | 0.701825034 | 3.01E-03 | 3.24E-02 |
| ZO2_HUMAN\|Q9UDY2_S244 | 0.701813455 | 8.31E-03 | 2.98E-02 |
| EGFR_HUMAN\|P00533_S1205 | 0.701798224 | 6.45E-03 | 2.41E-02 |
| C2CD5_HUMAN\|Q86YS7-2_S855 | 0.70177642 | 2.25E-03 | 2.77E-02 |
| TWF1_HUMAN\|Q12792_T349 | 0.701756 | 3.70E-04 | 7.28E-03 |
| RBM7_HUMAN\|Q9Y580_S137 | 0.701733903 | 4.67E-04 | 2.99E-03 |
| CAMP3_HUMAN\|Q9P1Y5_S756 | 0.701638763 | 1.31E-03 | 1.85E-02 |
| COHA1_HUMAN\|Q9UMD9_S172 | 0.701603163 | 2.17E-04 | 1.66E-03 |
| SI1L1_HUMAN\|O43166_S1255 | 0.701495141 | 6.53E-03 | 3.94E-02 |
| K1671_HUMAN\|Q9BY89_S128 | 0.701493419 | 2.65E-04 | 3.51E-03 |
| PCM1_HUMAN\|Q15154_S65_S69 | 0.701422092 | 3.71E-05 | 1.48E-03 |
| LMNA_HUMAN\|P02545_S407_S414 | 0.701242988 | 2.92E-03 | 2.15E-02 |
| PRC2C_HUMAN\|Q9Y520_T2691 | 0.701157041 | 1.33E-03 | 1.23E-02 |
| F193A_HUMAN\|P78312_S1144 | 0.701147865 | 2.01E-03 | 2.57E-02 |
| MAP1A_HUMAN\|P78559_S2022 | 0.701146911 | 1.01E-03 | 1.57E-02 |
| CRBG1_HUMAN\|Q9Y4K1_S75 | 0.701105427 | 1.18E-03 | 1.76E-02 |
| SMN_HUMAN\|Q16637_S31 | 0.701039197 | 6.65E-03 | 4.85E-02 |
| P33MX_HUMAN\|Q96A73_S183 | 0.700987497 | 5.93E-03 | 2.27E-02 |
| PKP3_HUMAN\|Q9Y446_S331 | 0.700892311 | 3.36E-03 | 1.65E-02 |
| NU214_HUMAN\|P35658_S1023 | 0.70084963 | 4.36E-03 | 3.98E-02 |
| 2A5D_HUMAN\|Q14738_S598 | 0.700848374 | 3.64E-03 | 3.72E-02 |
| SRSF9_HUMAN\|Q13242_S193 | 0.700655911 | 1.10E-03 | 1.36E-02 |
| DEFI6_HUMAN\|Q9H4E7_S597 | 0.70057183 | 9.65E-05 | 2.78E-03 |
| YAP1_HUMAN\|P46937_S109_T110 | 0.700533758 | 1.38E-04 | 1.16E-03 |
| LUZP1_HUMAN\|Q86V48_S575 | 0.70052433 | 1.91E-04 | 6.58E-03 |

| | | | |
|---|---|---|---|
| AHNK_HUMAN\|Q09666_S3412 | 0.700428442 | 2.53E-03 | 1.17E-02 |
| SRPK2_HUMAN\|P78362_S496_T498 | 0.700425935 | 2.09E-03 | 4.42E-02 |
| KKCC2_HUMAN\|Q96RR4-3_S522 | 0.700346069 | 8.38E-03 | 3.96E-02 |
| SLK_HUMAN\|Q9H2G2_S372 | 0.700345461 | 5.26E-05 | 5.56E-04 |
| ALDOA_HUMAN\|P04075_S46 | 0.700311678 | 3.04E-03 | 1.52E-02 |
| MAP7_HUMAN\|Q14244_S202 | 0.700255455 | 5.43E-04 | 5.98E-03 |
| FA83H_HUMAN\|Q6ZRV2_S523 | 0.700179176 | 6.22E-05 | 1.17E-03 |
| AHNK_HUMAN\|Q09666_S212_S216 | 0.700057208 | 6.17E-04 | 1.09E-02 |
| NUP50_HUMAN\|Q9UKX7_S221 | 0.7000058 | 5.52E-04 | 2.02E-02 |
| UBP2L_HUMAN\|Q14157_S467_S477 | 0.69999361 | 1.33E-03 | 1.75E-02 |
| M4K4_HUMAN\|O95819_S842 | 0.699967034 | 8.20E-04 | 5.20E-03 |
| OPTN_HUMAN\|Q96CV9_T525 | 0.699954027 | 4.73E-04 | 8.93E-03 |
| FA83H_HUMAN\|Q6ZRV2_S998 | 0.699926181 | 1.13E-04 | 4.26E-03 |
| NU214_HUMAN\|P35658_T987 | 0.699915934 | 1.18E-03 | 6.94E-03 |
| UT14A_HUMAN\|Q9BVJ6_S434 | 0.699913589 | 2.23E-03 | 4.57E-02 |
| ROCK1_HUMAN\|Q13464_S1341 | 0.699849081 | 4.56E-03 | 4.43E-02 |
| DGC14_HUMAN\|Q96DF8_T386 | 0.699758429 | 6.88E-04 | 7.08E-03 |
| LMNA_HUMAN\|P02545_S404_S406_S407 | 0.699749884 | 4.79E-03 | 2.20E-02 |
| PLEC_HUMAN\|Q15149_Y4615 | 0.699664215 | 2.03E-03 | 2.58E-02 |
| ABLM3_HUMAN\|O94929_S373_S379 | 0.699642643 | 7.62E-03 | 4.39E-02 |
| FARP1_HUMAN\|Q9Y4F1_T24 | 0.699639937 | 5.56E-03 | 2.49E-02 |
| IF4B_HUMAN\|P23588_S495_S497 | 0.69961667 | 5.46E-04 | 1.42E-02 |
| GMIP_HUMAN\|Q9P107_S914_S919 | 0.699538183 | 8.21E-04 | 5.21E-03 |
| ARHG5_HUMAN\|Q12774_S445_S450 | 0.699498513 | 1.58E-04 | 9.83E-03 |
| TLN1_HUMAN\|Q9Y490_T2270 | 0.699431908 | 3.26E-03 | 1.42E-02 |
| TGS1_HUMAN\|Q96RS0_S89 | 0.699431253 | 3.58E-03 | 2.43E-02 |
| K2C5_HUMAN\|P13647_S572 | 0.699314867 | 3.64E-03 | 2.46E-02 |
| MTAP2_HUMAN\|P11137_S1782_S1787_S1790 | 0.69929924 | 1.08E-03 | 1.64E-02 |
| K1671_HUMAN\|Q9BY89_S127 | 0.699288699 | 8.21E-04 | 1.36E-02 |
| N4BP1_HUMAN\|O75113_T242 | 0.699282966 | 8.83E-04 | 8.72E-03 |
| TRAF7_HUMAN\|Q6Q0C0_S108 | 0.699237779 | 6.76E-03 | 4.91E-02 |

| | | | |
|---|---|---|---|
| IRS2_HUMAN\|Q9Y4H2_S620 | 0.699229871 | 4.19E-03 | 4.20E-02 |
| JUNB_HUMAN\|P17275_T255 | 0.699220841 | 1.36E-04 | 4.02E-03 |
| UBP24_HUMAN\|Q9UPU5_S1141 | 0.69916509 | 1.80E-03 | 2.23E-02 |
| ARP19_HUMAN\|P56211_Y59 | 0.699164756 | 1.21E-03 | 1.73E-02 |
| PHF14_HUMAN\|O94880_S290_T287 | 0.699095112 | 4.65E-03 | 4.48E-02 |
| PLEC_HUMAN\|Q15149_S4386 | 0.699065516 | 8.71E-04 | 1.43E-02 |
| PTN3_HUMAN\|P26045_S425 | 0.699025081 | 4.42E-03 | 2.99E-02 |
| LMO7_HUMAN\|Q8WWI1_S1586 | 0.699018095 | 8.60E-04 | 4.86E-03 |
| MED1_HUMAN\|Q15648_Y1431 | 0.699006151 | 3.17E-04 | 6.82E-03 |
| K1C17_HUMAN\|Q04695_S10 | 0.698872136 | 4.69E-03 | 3.12E-02 |
| CWC25_HUMAN\|Q9NXE8_S218 | 0.698856333 | 1.49E-03 | 2.03E-02 |
| PKHA1_HUMAN\|Q9HB21_S332 | 0.698836915 | 1.38E-04 | 5.14E-03 |
| FBP1L_HUMAN\|Q5T0N5_S295 | 0.698766162 | 3.15E-03 | 2.21E-02 |
| AAPK1_HUMAN\|Q13131_T488_T490 | 0.698748377 | 1.53E-03 | 3.71E-02 |
| AFAP1_HUMAN\|Q8N556_S548 | 0.698650822 | 1.33E-04 | 1.13E-03 |
| YAP1_HUMAN\|P46937_S109_T110 | 0.698602612 | 1.26E-04 | 3.87E-03 |
| AFF4_HUMAN\|Q9UHB7_S836 | 0.698595585 | 3.44E-03 | 1.68E-02 |
| GRB10_HUMAN\|Q13322_S104 | 0.698540627 | 2.25E-04 | 5.36E-03 |
| CACO1_HUMAN\|Q9P1Z2_S563 | 0.69851238 | 1.25E-04 | 1.92E-03 |
| K1C16_HUMAN\|P08779_S447 | 0.698486472 | 8.18E-03 | 2.94E-02 |
| BI2L1_HUMAN\|Q9UHR4_S414 | 0.698476584 | 4.54E-03 | 4.60E-02 |
| LIPA1_HUMAN\|Q13136_S666_S668 | 0.698466961 | 8.97E-04 | 2.73E-02 |
| PHF14_HUMAN\|O94880_T287 | 0.698288312 | 2.34E-03 | 3.81E-02 |
| PACN3_HUMAN\|Q9UKS6_S344_S354 | 0.698274769 | 9.90E-04 | 2.85E-02 |
| TBC17_HUMAN\|Q9HA65_T620 | 0.698260475 | 1.97E-03 | 2.55E-02 |
| SEPT9_HUMAN\|Q9UHD8_T38_T42 | 0.698223315 | 6.59E-04 | 2.20E-02 |
| MILK1_HUMAN\|Q8N3F8_S644 | 0.698203265 | 1.65E-03 | 2.04E-02 |
| M3K20_HUMAN\|Q9NYL2_S727 | 0.698176951 | 1.18E-04 | 3.28E-03 |
| PRAG1_HUMAN\|Q86YV5_S907 | 0.698170159 | 8.24E-04 | 1.39E-02 |
| KC1D_HUMAN\|P48730_S383 | 0.698118399 | 2.66E-03 | 1.94E-02 |
| K1C14_HUMAN\|P02533_S435_S438 | 0.69809677 | 8.80E-03 | 4.84E-02 |

| | | | |
|---|---|---|---|
| SI1L3_HUMAN\|O60292_S172 | 0.698084172 | 2.26E-05 | 5.63E-04 |
| PEAK1_HUMAN\|Q9H792_S1035 | 0.698024379 | 7.22E-04 | 7.37E-03 |
| LMNB2_HUMAN\|Q03252_S424 | 0.697902506 | 5.41E-03 | 3.44E-02 |
| N4BP1_HUMAN\|O75113_T242 | 0.697779032 | 8.34E-04 | 4.75E-03 |
| NCOA7_HUMAN\|Q8NI08_S208 | 0.697742212 | 9.89E-04 | 9.73E-03 |
| PRSR2_HUMAN\|Q86WR7_S43 | 0.697732614 | 2.05E-04 | 4.75E-03 |
| TBCD5_HUMAN\|Q92609_S522 | 0.697709131 | 2.33E-03 | 1.54E-02 |
| APBB2_HUMAN\|Q92870_S409_S412 | 0.697677178 | 8.79E-04 | 1.43E-02 |
| AR6P4_HUMAN\|Q66PJ3_S241 | 0.697609675 | 6.86E-03 | 3.90E-02 |
| CAMP2_HUMAN\|Q08AD1_S1319 | 0.697555489 | 6.37E-04 | 1.11E-02 |
| NDRG1_HUMAN\|Q92597_S330_S333 | 0.697546326 | 1.27E-03 | 2.48E-02 |
| LMNA_HUMAN\|P02545_S403_S406 | 0.697524599 | 3.22E-03 | 4.65E-02 |
| CE170_HUMAN\|Q5SW79_S1165 | 0.69750828 | 7.60E-03 | 4.39E-02 |
| PP12C_HUMAN\|Q9BZL4_S453 | 0.697484912 | 8.33E-05 | 3.30E-03 |
| CE170_HUMAN\|Q5SW79_S636 | 0.697430422 | 2.19E-05 | 2.74E-04 |
| IRS2_HUMAN\|Q9Y4H2_S1186_T1202 | 0.697312406 | 2.43E-04 | 5.45E-03 |
| ZC3HD_HUMAN\|Q5T200_S242 | 0.697303213 | 2.22E-04 | 1.81E-03 |
| PSD3_HUMAN\|Q9NYI0_S1012_S1020 | 0.697280503 | 2.58E-04 | 3.43E-03 |
| CCM2_HUMAN\|Q9BSQ5_S384 | 0.697132386 | 2.76E-03 | 1.98E-02 |
| FRYL_HUMAN\|O94915_T1959 | 0.697115528 | 1.19E-03 | 1.08E-02 |
| LIPA3_HUMAN\|O75145_S17 | 0.69708125 | 6.92E-03 | 3.93E-02 |
| AHNK_HUMAN\|Q09666_S511 | 0.697058807 | 5.34E-03 | 2.09E-02 |
| CAAP1_HUMAN\|Q9H8G2_T90 | 0.696984443 | 2.42E-03 | 1.12E-02 |
| LMNA_HUMAN\|P02545_S414 | 0.69697773 | 2.39E-03 | 2.87E-02 |
| REPS1_HUMAN\|Q96D71_S162_S166_T173 | 0.696967642 | 3.84E-04 | 8.07E-03 |
| LARP1_HUMAN\|Q6PKG0_S774 | 0.696939025 | 1.61E-04 | 3.04E-03 |
| AAPK1_HUMAN\|Q13131_S508 | 0.696938678 | 4.15E-05 | 6.60E-04 |
| AFAD_HUMAN\|P55196_S1696 | 0.696927644 | 1.80E-03 | 9.78E-03 |
| NDRG1_HUMAN\|Q92597_T328 | 0.696905677 | 1.85E-03 | 1.00E-02 |
| 1433G_HUMAN\|P61981_T234 | 0.696869852 | 9.29E-04 | 5.18E-03 |
| LASP1_HUMAN\|Q14847_T104 | 0.696852865 | 7.87E-04 | 2.48E-02 |

| | | | |
|---|---|---|---|
| SARG_HUMAN\|Q9BW04_T456 | 0.696784169 | 1.03E-04 | 9.33E-04 |
| SON_HUMAN\|P18583_S94 | 0.696781988 | 2.03E-04 | 4.72E-03 |
| CHSP1_HUMAN\|Q9Y2V2_S41 | 0.696767687 | 3.08E-03 | 1.36E-02 |
| AHNK_HUMAN\|Q09666_S559 | 0.696695674 | 2.53E-04 | 3.37E-03 |
| LAR4B_HUMAN\|Q92615_S731_S736 | 0.696632988 | 2.56E-03 | 4.03E-02 |
| SRSF9_HUMAN\|Q13242_S193 | 0.696573836 | 9.44E-04 | 9.07E-03 |
| K1C17_HUMAN\|Q04695_S28 | 0.696549623 | 6.59E-03 | 2.87E-02 |
| SLK_HUMAN\|Q9H2G2_S372 | 0.696521702 | 4.32E-05 | 1.75E-03 |
| TB182_HUMAN\|Q9C0C2_T875 | 0.696518476 | 9.17E-04 | 8.89E-03 |
| S38A2_HUMAN\|Q96QD8_S17_S21 | 0.696487004 | 1.15E-02 | 4.52E-02 |
| LAD1_HUMAN\|O00515_S301 | 0.696477058 | 5.62E-03 | 3.35E-02 |
| K1522_HUMAN\|Q9P206_S161 | 0.696464251 | 6.62E-04 | 1.60E-02 |
| PRSR2_HUMAN\|Q86WR7_S18 | 0.696440652 | 5.36E-05 | 1.99E-03 |
| GOGA4_HUMAN\|Q13439_T31 | 0.69643903 | 1.70E-04 | 2.45E-03 |
| MAP7_HUMAN\|Q14244_S209 | 0.696419206 | 6.98E-03 | 3.01E-02 |
| RHG23_HUMAN\|Q9P227_S351 | 0.696339037 | 1.55E-03 | 7.87E-03 |
| CARL1_HUMAN\|Q5VZK9_S968 | 0.69632738 | 1.18E-04 | 1.91E-03 |
| ALKB5_HUMAN\|Q6P6C2_S384 | 0.696291649 | 2.10E-04 | 2.91E-03 |
| NIBL1_HUMAN\|Q96TA1_S691 | 0.69629018 | 8.84E-05 | 1.18E-03 |
| K1C14_HUMAN\|P02533_S44_S53_S54 | 0.696288601 | 1.41E-03 | 1.27E-02 |
| PYGO2_HUMAN\|Q9BRQ0_T302 | 0.696284899 | 9.19E-04 | 5.13E-03 |
| PTN3_HUMAN\|P26045_S425 | 0.696230698 | 4.07E-03 | 1.70E-02 |
| SON_HUMAN\|P18583_S94 | 0.696222815 | 1.98E-04 | 1.65E-03 |
| PLXA2_HUMAN\|O75051_S1630_S1633 | 0.696217663 | 4.37E-03 | 4.24E-02 |
| FA53C_HUMAN\|Q9NYF3_S122 | 0.696203775 | 5.99E-04 | 3.63E-03 |
| TB182_HUMAN\|Q9C0C2_S872 | 0.696186454 | 7.67E-04 | 4.45E-03 |
| RAI14_HUMAN\|Q9P0K7_S296 | 0.696181602 | 5.17E-03 | 3.35E-02 |
| TASOR_HUMAN\|Q9UK61_S344 | 0.696146035 | 8.40E-04 | 1.36E-02 |
| SRSF4_HUMAN\|Q08170_S460 | 0.696143261 | 6.48E-03 | 4.86E-02 |
| PEAK1_HUMAN\|Q9H792_S1035 | 0.695923809 | 6.62E-04 | 4.36E-03 |
| TB182_HUMAN\|Q9C0C2_S1711 | 0.695916801 | 9.29E-04 | 1.98E-02 |

| | | | |
|---|---|---|---|
| AFF4_HUMAN\|Q9UHB7_S1043 | 0.695869166 | 3.24E-04 | 9.92E-03 |
| K1671_HUMAN\|Q9BY89_S404 | 0.69585809 | 2.24E-04 | 1.81E-03 |
| RHG32_HUMAN\|A7KAX9_S892 | 0.695812499 | 4.53E-04 | 5.23E-03 |
| PDZD2_HUMAN\|O15018_T1099 | 0.69579222 | 1.05E-02 | 3.60E-02 |
| HDAC4_HUMAN\|P56524_S266 | 0.695778757 | 8.29E-04 | 1.39E-02 |
| PDE1C_HUMAN\|Q14123_S469 | 0.695685102 | 1.01E-03 | 5.58E-03 |
| BNIP2_HUMAN\|Q12982_S114 | 0.695602071 | 7.97E-03 | 4.37E-02 |
| LMNA_HUMAN\|P02545_S12 | 0.695596922 | 4.34E-03 | 1.79E-02 |
| AHNK_HUMAN\|Q09666_S210 | 0.695525133 | 1.84E-04 | 2.14E-03 |
| NUCKS_HUMAN\|Q9H1E3_S54_S58_S61 | 0.695524951 | 5.09E-04 | 3.21E-03 |
| EMAL4_HUMAN\|Q9HC35_S146 | 0.695511621 | 2.05E-03 | 1.09E-02 |
| SUZ12_HUMAN\|Q15022_S541_S546 | 0.695393851 | 1.02E-02 | 4.10E-02 |
| WWC3_HUMAN\|Q9ULE0_T909 | 0.695322266 | 2.54E-03 | 1.86E-02 |
| 4EBP1_HUMAN\|Q13541_S44 | 0.695302852 | 1.72E-03 | 2.10E-02 |
| AHNK_HUMAN\|Q09666_T4766 | 0.695241919 | 2.36E-03 | 1.84E-02 |
| KHDR1_HUMAN\|Q07666_S18 | 0.695208617 | 2.84E-03 | 1.28E-02 |
| PER1_HUMAN\|O15534_S815 | 0.69514842 | 5.43E-03 | 2.45E-02 |
| RABE1_HUMAN\|Q15276_S407_S410 | 0.695080656 | 1.82E-05 | 2.34E-04 |
| K1C14_HUMAN\|P02533_S429_S432 | 0.695049003 | 8.59E-03 | 3.07E-02 |
| ADRM1_HUMAN\|Q16186_S211 | 0.695028204 | 2.67E-03 | 4.13E-02 |
| JIP4_HUMAN\|O60271_S815 | 0.694963528 | 5.48E-04 | 9.93E-03 |
| LARP6_HUMAN\|Q9BRS8_S56_S58 | 0.694950837 | 1.85E-03 | 1.54E-02 |
| FA76B_HUMAN\|Q5HYJ3_S338 | 0.694838994 | 3.73E-03 | 1.59E-02 |
| RTKN_HUMAN\|Q9BST9_S520 | 0.694826804 | 2.75E-04 | 2.10E-03 |
| WWC3_HUMAN\|Q9ULE0_T909 | 0.694704229 | 2.49E-03 | 1.91E-02 |
| GGYF1_HUMAN\|O75420_S24 | 0.694703092 | 8.38E-04 | 5.29E-03 |
| TB182_HUMAN\|Q9C0C2_S1715 | 0.694677863 | 6.03E-03 | 3.54E-02 |
| AFAP1_HUMAN\|Q8N556_S548 | 0.694573085 | 1.09E-04 | 1.03E-03 |
| K1C16_HUMAN\|P08779_S446_S449 | 0.694401858 | 3.61E-03 | 4.87E-02 |
| FA83B_HUMAN\|Q5T0W9_S802 | 0.694334233 | 1.33E-03 | 1.58E-02 |
| IF4E_HUMAN\|P06730_T203 | 0.694284397 | 1.67E-03 | 3.02E-02 |

| | | | |
|---|---|---|---|
| SRCAP_HUMAN\|Q6ZRS2_S2790 | 0.69427596 | 4.24E-03 | 1.76E-02 |
| WDR13_HUMAN\|Q9H1Z4_S70_S74 | 0.694209452 | 1.49E-03 | 2.76E-02 |
| MTMR2_HUMAN\|Q13614_S631 | 0.694185492 | 2.03E-03 | 1.65E-02 |
| YBOX3_HUMAN\|P16989_S203 | 0.694127531 | 1.04E-03 | 5.70E-03 |
| GGYF2_HUMAN\|Q6Y7W6_S23 | 0.694127171 | 2.70E-04 | 5.88E-03 |
| RTKN_HUMAN\|Q9BST9_S543 | 0.694053366 | 5.94E-04 | 3.99E-03 |
| CR025_HUMAN\|Q96B23_S143_S145 | 0.694046558 | 5.76E-04 | 1.44E-02 |
| RHG29_HUMAN\|Q52LW3_S1029 | 0.693965555 | 1.69E-03 | 2.15E-02 |
| RHG32_HUMAN\|A7KAX9_S892 | 0.693935944 | 4.20E-04 | 2.97E-03 |
| ANXA2_HUMAN\|P07355_T19 | 0.693889943 | 7.43E-05 | 5.16E-03 |
| KIF1C_HUMAN\|O43896_S817 | 0.693872038 | 1.10E-03 | 1.68E-02 |
| AHNK_HUMAN\|Q09666_T5729 | 0.693839915 | 3.92E-04 | 8.10E-03 |
| AAK1_HUMAN\|Q2M2I8_T389 | 0.693811805 | 4.50E-03 | 2.88E-02 |
| GRAP1_HUMAN\|Q4V328_S664 | 0.693802712 | 2.22E-03 | 1.77E-02 |
| NONO_HUMAN\|Q15233_T450 | 0.693719535 | 9.07E-04 | 8.84E-03 |
| YETS2_HUMAN\|Q9ULM3_S536 | 0.693598263 | 9.24E-04 | 1.47E-02 |
| NIBL1_HUMAN\|Q96TA1_S665 | 0.693580477 | 3.55E-04 | 3.61E-03 |
| NDRG1_HUMAN\|Q92597_T328 | 0.693512096 | 1.64E-03 | 8.18E-03 |
| SRRM2_HUMAN\|Q9UQ35_S2449 | 0.693409237 | 2.09E-03 | 3.48E-02 |
| PRR12_HUMAN\|Q9ULL5_S44 | 0.693402528 | 1.90E-03 | 2.46E-02 |
| DOCK6_HUMAN\|Q96HP0_S1343 | 0.69340055 | 4.03E-03 | 4.01E-02 |
| OSTF1_HUMAN\|Q92882_S213 | 0.693373958 | 2.27E-03 | 2.78E-02 |
| BAG3_HUMAN\|O95817_S289 | 0.693371889 | 4.13E-03 | 1.72E-02 |
| CLASR_HUMAN\|Q8N2M8_S547 | 0.693365165 | 1.21E-04 | 1.05E-03 |
| ALKB5_HUMAN\|Q6P6C2_S384 | 0.693295921 | 1.83E-04 | 1.57E-03 |
| CCNL2_HUMAN\|Q96S94_S330 | 0.693239109 | 1.71E-04 | 4.17E-03 |
| NDRG1_HUMAN\|Q92597_S326_S330_S333 | 0.693205587 | 2.88E-03 | 3.27E-02 |
| TMF1_HUMAN\|P82094_S77 | 0.693157217 | 1.32E-03 | 9.88E-03 |
| KLC3_HUMAN\|Q6P597_S497_S502 | 0.693103681 | 8.01E-05 | 7.92E-04 |
| CT2NL_HUMAN\|Q9P2B4_S568 | 0.69304739 | 8.32E-04 | 4.75E-03 |
| STX7_HUMAN\|O15400_S131 | 0.69294497 | 4.34E-04 | 3.06E-03 |

| | | | |
|---|---|---|---|
| CR025_HUMAN\|Q96B23_S143_S145_S147 | 0.692926445 | 4.65E-03 | 2.15E-02 |
| SHRM2_HUMAN\|Q13796_S764 | 0.692904831 | 3.30E-03 | 3.69E-02 |
| KLF16_HUMAN\|Q9BXK1_S238 | 0.69283553 | 3.94E-03 | 1.67E-02 |
| AHNK_HUMAN\|Q09666_S5731 | 0.692827406 | 3.23E-05 | 1.42E-03 |
| HMGA2_HUMAN\|P52926_S101_S105_T100 | 0.692747023 | 3.92E-03 | 2.59E-02 |
| PININ_HUMAN\|Q9H307_S66 | 0.692742008 | 3.68E-03 | 2.14E-02 |
| SRP09_HUMAN\|P49458_T85 | 0.692635872 | 1.63E-03 | 2.97E-02 |
| HD_HUMAN\|P42858_S1864 | 0.692584245 | 6.71E-03 | 2.91E-02 |
| LMNA_HUMAN\|P02545_S407_S414 | 0.692578161 | 2.18E-03 | 2.52E-02 |
| KHDR1_HUMAN\|Q07666_S20 | 0.692488369 | 7.02E-04 | 4.13E-03 |
| AHNK_HUMAN\|Q09666_T566 | 0.692478908 | 1.08E-03 | 5.88E-03 |
| SSFA2_HUMAN\|P28290_S668 | 0.692477657 | 5.83E-03 | 4.85E-02 |
| FA83H_HUMAN\|Q6ZRV2_S516 | 0.692477395 | 3.36E-03 | 1.65E-02 |
| ACTN4_HUMAN\|O43707_S909 | 0.692449552 | 5.36E-03 | 2.09E-02 |
| BCAR3_HUMAN\|O75815_S375 | 0.692441945 | 4.00E-03 | 1.69E-02 |
| FA83B_HUMAN\|Q5T0W9_S869 | 0.692438 | 1.11E-05 | 3.24E-04 |
| PLEC_HUMAN\|Q15149_S4384_S4385 | 0.692425494 | 4.18E-03 | 3.58E-02 |
| 4EBP1_HUMAN\|Q13541_T70 | 0.692424733 | 1.71E-03 | 2.15E-02 |
| K2C8_HUMAN\|P05787_S21_S24_S27 | 0.692393314 | 3.48E-03 | 3.81E-02 |
| SRRM1_HUMAN\|Q8IYB3_S738_S740_S743 | 0.69238128 | 7.39E-04 | 7.47E-03 |
| GSK3A_HUMAN\|P49840_T19 | 0.692376148 | 3.38E-04 | 4.16E-03 |
| K2C5_HUMAN\|P13647_S82_T78 | 0.692366289 | 1.33E-03 | 1.83E-02 |
| CR025_HUMAN\|Q96B23_S145_S147 | 0.692353413 | 3.88E-04 | 2.60E-03 |
| ZCRB1_HUMAN\|Q8TBF4_S216 | 0.692287707 | 4.19E-04 | 4.09E-03 |
| CCNL2_HUMAN\|Q96S94_S330 | 0.692190889 | 1.63E-04 | 4.06E-03 |
| PHF14_HUMAN\|O94880_T287 | 0.692126859 | 1.91E-03 | 1.58E-02 |
| PDLI1_HUMAN\|O00151_S130 | 0.692082442 | 1.14E-03 | 6.74E-03 |
| ARG28_HUMAN\|Q8N1W1_S513 | 0.692080861 | 1.49E-03 | 7.64E-03 |
| MA7D1_HUMAN\|Q3KQU3_S366 | 0.692045187 | 1.06E-04 | 3.41E-03 |
| RPGF6_HUMAN\|Q8TEU7_S1590 | 0.691978371 | 2.16E-03 | 1.02E-02 |
| TB182_HUMAN\|Q9C0C2_S936 | 0.691950923 | 6.62E-04 | 7.03E-03 |

| | | | |
|---|---|---|---|
| TB182_HUMAN\|Q9C0C2_S836 | 0.691941089 | 5.73E-04 | 5.29E-03 |
| COIL_HUMAN\|P38432_S566_S571 | 0.691939969 | 1.55E-03 | 2.03E-02 |
| KC1D_HUMAN\|P48730_S383 | 0.691931827 | 2.16E-03 | 1.02E-02 |
| PHLB2_HUMAN\|Q86SQ0_Y567 | 0.691875324 | 4.50E-04 | 3.15E-03 |
| ABI2_HUMAN\|Q9NYB9_S368 | 0.69177941 | 9.57E-05 | 1.66E-03 |
| SRRM2_HUMAN\|Q9UQ35_T1492 | 0.691709889 | 1.06E-03 | 9.89E-03 |
| TISB_HUMAN\|Q07352_S336 | 0.691675505 | 5.70E-03 | 3.58E-02 |
| SOX15_HUMAN\|O60248_S37 | 0.691628445 | 1.25E-04 | 3.70E-03 |
| KAZRN_HUMAN\|Q674X7-3_S8 | 0.691558254 | 8.60E-04 | 5.40E-03 |
| DDX17_HUMAN\|Q92841_S599 | 0.691548495 | 2.47E-04 | 2.73E-03 |
| DGC14_HUMAN\|Q96DF8_T386 | 0.691539671 | 4.90E-04 | 4.69E-02 |
| DBNL_HUMAN\|Q9UJU6_S269 | 0.691529476 | 5.73E-03 | 3.60E-02 |
| JUPI2_HUMAN\|Q9H910_S97 | 0.691520734 | 3.50E-03 | 4.87E-02 |
| PRSR2_HUMAN\|Q86WR7_S43 | 0.691435137 | 1.52E-04 | 5.62E-03 |
| KPCD2_HUMAN\|Q9BZL6_S214 | 0.691401162 | 3.70E-03 | 1.58E-02 |
| DDI2_HUMAN\|Q5TDH0_S106 | 0.691366504 | 3.82E-04 | 2.75E-03 |
| ZCCHV_HUMAN\|Q7Z2W4_S378 | 0.691360963 | 5.16E-05 | 5.08E-03 |
| MGMT_HUMAN\|P16455_S201 | 0.691292789 | 2.80E-05 | 2.92E-03 |
| KLC3_HUMAN\|Q6P597_S497 | 0.691274875 | 1.84E-04 | 1.57E-03 |
| SPIR1_HUMAN\|Q08AE8_T509 | 0.691189563 | 1.20E-03 | 1.08E-02 |
| COHA1_HUMAN\|Q9UMD9_T387 | 0.691185775 | 9.50E-04 | 9.09E-03 |
| K1522_HUMAN\|Q9P206_S929 | 0.691182579 | 3.02E-04 | 4.81E-03 |
| ABLM3_HUMAN\|O94929_S372_S379 | 0.691128343 | 3.88E-03 | 3.87E-02 |
| PSF2_HUMAN\|Q9Y248_S182 | 0.69109576 | 5.17E-03 | 3.35E-02 |
| PKP2_HUMAN\|Q99959_S132_S135 | 0.691064374 | 4.00E-03 | 1.89E-02 |
| SF3B1_HUMAN\|O75533_T248 | 0.691003517 | 4.41E-03 | 1.82E-02 |
| M3K7_HUMAN\|O43318_S439_S455 | 0.69097035 | 8.39E-03 | 3.50E-02 |
| PRC2A_HUMAN\|P48634_S761 | 0.690862903 | 5.73E-04 | 6.23E-03 |
| FA83H_HUMAN\|Q6ZRV2_S974 | 0.69081192 | 5.64E-04 | 1.00E-02 |
| YBOX1_HUMAN\|P67809_S165_S176 | 0.690733232 | 1.10E-03 | 1.63E-02 |
| NKTR_HUMAN\|P30414_T1155 | 0.690728633 | 1.49E-03 | 2.03E-02 |

| | | | |
|---|---|---|---|
| JIP4_HUMAN\|O60271_S815 | 0.690627524 | 4.63E-04 | 5.32E-03 |
| BACH1_HUMAN\|O14867_S445 | 0.69057917 | 1.25E-03 | 1.80E-02 |
| IF4B_HUMAN\|P23588_S597 | 0.690578715 | 7.44E-03 | 3.59E-02 |
| SQSTM_HUMAN\|Q13501_T269 | 0.690353769 | 1.10E-03 | 1.05E-02 |
| RIOK2_HUMAN\|Q9BVS4_S380_S382_S385 | 0.690263804 | 8.54E-04 | 4.85E-03 |
| MA7D1_HUMAN\|Q3KQU3_S313 | 0.690188536 | 7.84E-03 | 4.49E-02 |
| AKP13_HUMAN\|Q12802_S2728 | 0.690176082 | 1.53E-04 | 1.35E-03 |
| P66B_HUMAN\|Q8WXI9_S334 | 0.690175547 | 7.14E-05 | 2.65E-03 |
| TB182_HUMAN\|Q9C0C2_S1715 | 0.690164263 | 5.31E-03 | 2.08E-02 |
| CRBG1_HUMAN\|Q9Y4K1_S75 | 0.690137126 | 7.92E-04 | 1.31E-02 |
| COMT_HUMAN\|P21964_S267 | 0.690105035 | 7.99E-04 | 4.61E-03 |
| DOCK6_HUMAN\|Q96HP0_S1343 | 0.690020376 | 3.63E-03 | 1.75E-02 |
| CSN3_HUMAN\|Q9UNS2_S421 | 0.690011738 | 5.13E-05 | 2.28E-03 |
| FA83H_HUMAN\|Q6ZRV2_S1025 | 0.690001692 | 1.27E-03 | 7.35E-03 |
| AJUBA_HUMAN\|Q96IF1_S69 | 0.689927977 | 4.88E-03 | 3.03E-02 |
| DOCK6_HUMAN\|Q96HP0_S882 | 0.689920338 | 3.62E-03 | 2.45E-02 |
| RHG05_HUMAN\|Q13017_S1218 | 0.68991038 | 6.29E-04 | 6.74E-03 |
| NDRG2_HUMAN\|Q9UN36_S332 | 0.689900675 | 5.08E-04 | 3.21E-03 |
| MAP1B_HUMAN\|P46821_S1792 | 0.689899628 | 3.69E-03 | 2.59E-02 |
| CND1_HUMAN\|Q15021_S1333 | 0.689889784 | 9.93E-04 | 2.92E-02 |
| SMC1A_HUMAN\|Q14683_S956 | 0.689887695 | 4.43E-03 | 4.03E-02 |
| PYGO2_HUMAN\|Q9BRQ0_T302 | 0.689886651 | 7.18E-04 | 1.25E-02 |
| ACLY_HUMAN\|P53396_S455 | 0.689876673 | 5.57E-05 | 1.85E-03 |
| UBP24_HUMAN\|Q9UPU5_S1143 | 0.689859617 | 9.98E-04 | 1.54E-02 |
| ZNRF2_HUMAN\|Q8NHG8_S135 | 0.689857751 | 3.61E-03 | 2.45E-02 |
| MISP_HUMAN\|Q8IVT2_T377 | 0.689849895 | 1.66E-03 | 9.14E-03 |
| UBP31_HUMAN\|Q70CQ4_S1088 | 0.689758396 | 2.04E-05 | 5.19E-04 |
| CAP1_HUMAN\|Q01518_S308 | 0.689727131 | 5.71E-04 | 1.03E-02 |
| FARP1_HUMAN\|Q9Y4F1_T24 | 0.689692924 | 4.10E-03 | 2.69E-02 |
| TB182_HUMAN\|Q9C0C2_S1711 | 0.689660713 | 7.29E-04 | 4.26E-03 |
| PDLI2_HUMAN\|Q96JY6_S117 | 0.689649911 | 3.40E-03 | 2.04E-02 |

| | | | |
|---|---|---|---|
| TIAM1_HUMAN\|Q13009_S358 | 0.689613737 | 5.35E-04 | 3.65E-03 |
| IRS1_HUMAN\|P35568_S1100 | 0.689596126 | 8.43E-04 | 8.64E-03 |
| ARHGG_HUMAN\|Q5VV41_S107 | 0.689582625 | 9.11E-05 | 2.96E-03 |
| C2C4C_HUMAN\|Q8TF44_S156 | 0.689495848 | 4.26E-03 | 2.76E-02 |
| CNOT2_HUMAN\|Q9NZN8_S165 | 0.689493735 | 3.97E-04 | 4.73E-03 |
| MA7D1_HUMAN\|Q3KQU3_S834 | 0.689489891 | 2.79E-03 | 3.22E-02 |
| KLC3_HUMAN\|Q6P597_S497 | 0.689462188 | 1.70E-04 | 1.36E-03 |
| SYNRG_HUMAN\|Q9UMZ2_S935 | 0.689415519 | 1.62E-03 | 8.99E-03 |
| KLC3_HUMAN\|Q6P597_S495 | 0.689382338 | 1.43E-02 | 4.59E-02 |
| I2BP2_HUMAN\|Q7Z5L9_S460 | 0.689353757 | 4.08E-06 | 7.34E-05 |
| MARCS_HUMAN\|P29966_T150 | 0.68935136 | 4.60E-03 | 4.64E-02 |
| MAP7_HUMAN\|Q14244_S315 | 0.689327324 | 8.38E-04 | 8.35E-03 |
| SMC1A_HUMAN\|Q14683_S956 | 0.689317954 | 4.36E-03 | 2.04E-02 |
| MACF1_HUMAN\|Q9UPN3_S7330_S7345 | 0.689308088 | 2.02E-03 | 9.70E-03 |
| ES8L2_HUMAN\|Q9H6S3_S28 | 0.689264304 | 5.25E-05 | 1.80E-03 |
| SRRM2_HUMAN\|Q9UQ35_S1582 | 0.689247648 | 1.45E-04 | 1.20E-03 |
| BL1S3_HUMAN\|Q6QNY0_S65_T63 | 0.689158864 | 8.33E-04 | 1.39E-02 |
| OSBL8_HUMAN\|Q9BZF1_S807_S808_S810 | 0.689145937 | 5.82E-03 | 4.85E-02 |
| ARAF_HUMAN\|P10398_S257_T253 | 0.689134132 | 2.09E-03 | 2.44E-02 |
| DCA10_HUMAN\|Q5QP82_S349 | 0.689063046 | 2.10E-03 | 1.43E-02 |
| P4K2A_HUMAN\|Q9BTU6_S468 | 0.689056922 | 8.19E-03 | 4.60E-02 |
| OSB10_HUMAN\|Q9BXB5_S26_S32 | 0.689056138 | 1.44E-04 | 2.19E-03 |
| PLEC_HUMAN\|Q15149_S4386 | 0.688950166 | 5.83E-04 | 3.93E-03 |
| YBOX3_HUMAN\|P16989_S203 | 0.688899931 | 8.53E-04 | 1.40E-02 |
| IRS2_HUMAN\|Q9Y4H2_T520 | 0.688885852 | 4.61E-03 | 4.45E-02 |
| GSK3A_HUMAN\|P49840_S21 | 0.688868065 | 5.99E-04 | 6.51E-03 |
| YAP1_HUMAN\|P46937_S105_S109 | 0.688804439 | 6.16E-05 | 6.09E-04 |
| TACC1_HUMAN\|O75410_S44 | 0.688625871 | 1.47E-03 | 2.73E-02 |
| PDLI1_HUMAN\|O00151_S130 | 0.688618093 | 9.94E-04 | 2.85E-02 |
| ZCCHV_HUMAN\|Q7Z2W4_S378 | 0.688612716 | 4.52E-05 | 9.20E-04 |
| AHNK_HUMAN\|Q09666_T4564 | 0.688607271 | 2.80E-03 | 2.09E-02 |

| | | | |
|---|---|---|---|
| SPD2A_HUMAN\|Q5TCZ1_S819 | 0.688571153 | 1.31E-02 | 4.29E-02 |
| K1671_HUMAN\|Q9BY89_S404 | 0.688563537 | 1.62E-04 | 4.04E-03 |
| MA7D1_HUMAN\|Q3KQU3_S834 | 0.688504883 | 2.70E-03 | 1.38E-02 |
| CD97_HUMAN\|P48960_S831 | 0.688458825 | 9.79E-04 | 5.96E-03 |
| SRRM1_HUMAN\|Q8IYB3_S738_S740_S743 | 0.688436993 | 6.30E-04 | 1.14E-02 |
| NONO_HUMAN\|Q15233_T450 | 0.688436236 | 7.42E-04 | 1.24E-02 |
| SHRM2_HUMAN\|Q13796_S1039 | 0.688392074 | 7.78E-06 | 5.70E-04 |
| HPS5_HUMAN\|Q9UPZ3_S534 | 0.688355335 | 1.56E-03 | 1.30E-02 |
| YAP1_HUMAN\|P46937_T110 | 0.688281204 | 7.45E-03 | 2.71E-02 |
| MAST4_HUMAN\|O15021_S2555 | 0.68825071 | 3.14E-03 | 4.58E-02 |
| HEAT6_HUMAN\|Q6AI08_S643 | 0.688245417 | 7.58E-05 | 7.20E-04 |
| UBP32_HUMAN\|Q8NFA0_S1393 | 0.688245031 | 4.90E-04 | 7.57E-03 |
| DESP_HUMAN\|P15924_S2821_S2825 | 0.68823981 | 8.77E-04 | 5.49E-03 |
| FGD6_HUMAN\|Q6ZV73_S721 | 0.688130151 | 3.92E-03 | 2.59E-02 |
| K2C5_HUMAN\|P13647_S571 | 0.688093535 | 4.70E-04 | 3.01E-03 |
| K1C14_HUMAN\|P02533_S33 | 0.688084843 | 1.87E-03 | 1.55E-02 |
| YAP1_HUMAN\|P46937_S127 | 0.688079055 | 1.23E-02 | 4.76E-02 |
| PAIRB_HUMAN\|Q8NC51_S392 | 0.687925116 | 2.48E-03 | 4.83E-02 |
| CE170_HUMAN\|Q5SW79_S1165 | 0.687771214 | 5.82E-03 | 2.23E-02 |
| DEFI6_HUMAN\|Q9H4E7_S597 | 0.68774799 | 5.09E-05 | 9.92E-04 |
| SPD2A_HUMAN\|Q5TCZ1_S651 | 0.68765386 | 4.52E-03 | 2.90E-02 |
| PDLI4_HUMAN\|P50479_S116 | 0.687575838 | 2.46E-03 | 1.89E-02 |
| ASPC1_HUMAN\|Q9BZE9_S275 | 0.687425282 | 7.47E-04 | 1.27E-02 |
| FSTL3_HUMAN\|O95633_S255 | 0.687420286 | 3.60E-03 | 3.78E-02 |
| CR025_HUMAN\|Q96B23_S66 | 0.687406847 | 2.79E-04 | 6.02E-03 |
| K2C5_HUMAN\|P13647_S571_S574 | 0.687385961 | 2.51E-03 | 2.96E-02 |
| CD2AP_HUMAN\|Q9Y5K6_S458 | 0.687316236 | 1.60E-03 | 8.04E-03 |
| DEN1A_HUMAN\|Q8TEH3_S536_S546 | 0.687250863 | 3.96E-03 | 2.26E-02 |
| N4BP1_HUMAN\|O75113_T242 | 0.687246399 | 5.55E-04 | 6.09E-03 |
| HPS5_HUMAN\|Q9UPZ3_S534 | 0.687180193 | 1.50E-03 | 1.32E-02 |
| FA83H_HUMAN\|Q6ZRV2_S523 | 0.68716434 | 3.09E-05 | 3.63E-04 |

| | | | |
|---|---|---|---|
| RGL2_HUMAN\|O15211_S761 | 0.687142649 | 1.23E-03 | 6.52E-03 |
| LIPB1_HUMAN\|Q86W92_S1001 | 0.687140704 | 1.19E-02 | 3.94E-02 |
| PDLI4_HUMAN\|P50479_S112 | 0.687019906 | 9.21E-04 | 8.91E-03 |
| IASPP_HUMAN\|Q8WUF5_S134 | 0.686978576 | 3.39E-03 | 3.62E-02 |
| TRM1_HUMAN\|Q9NXH9_S625 | 0.686899328 | 2.65E-03 | 1.21E-02 |
| KPCD1_HUMAN\|Q15139_S742 | 0.686883355 | 1.02E-02 | 4.61E-02 |
| K1C14_HUMAN\|P02533_S33 | 0.686858706 | 1.79E-03 | 9.76E-03 |
| NDRG2_HUMAN\|Q9UN36_S332_T330 | 0.686755115 | 8.06E-04 | 1.34E-02 |
| LMNB1_HUMAN\|P20700_S404 | 0.686703079 | 1.26E-02 | 4.84E-02 |
| AKP13_HUMAN\|Q12802_S2728 | 0.686640409 | 1.30E-04 | 1.11E-03 |
| K1C14_HUMAN\|P02533_Y46 | 0.686617708 | 3.27E-03 | 3.53E-02 |
| FOXO3_HUMAN\|O43524_S425 | 0.686573947 | 3.93E-04 | 3.90E-03 |
| MPLKI_HUMAN\|Q8TAP9_T110 | 0.686539368 | 2.88E-03 | 3.27E-02 |
| LARP1_HUMAN\|Q6PKG0_S774 | 0.686417374 | 9.90E-05 | 6.23E-03 |
| CDK13_HUMAN\|Q14004_T1056 | 0.686355093 | 1.70E-03 | 2.31E-02 |
| FA83B_HUMAN\|Q5T0W9_S388 | 0.686331874 | 3.62E-03 | 1.55E-02 |
| RHG23_HUMAN\|Q9P227_S361_S372 | 0.686091742 | 7.07E-05 | 6.24E-03 |
| BAD_HUMAN\|Q92934_Y76 | 0.68599686 | 2.67E-04 | 8.41E-03 |
| FA83H_HUMAN\|Q6ZRV2_S828 | 0.685969179 | 1.46E-04 | 1.21E-03 |
| AHNK_HUMAN\|Q09666_S216 | 0.685911019 | 1.79E-04 | 4.80E-03 |
| BAIP2_HUMAN\|Q9UQB8_S453 | 0.685893064 | 1.20E-03 | 1.72E-02 |
| RBM7_HUMAN\|Q9Y580_S136 | 0.685887127 | 3.63E-03 | 3.26E-02 |
| SPTN1_HUMAN\|Q13813_S1190 | 0.685832572 | 2.94E-04 | 2.21E-03 |
| TBD2A_HUMAN\|Q9BYX2_S920 | 0.685806298 | 5.94E-04 | 1.08E-02 |
| KI16B_HUMAN\|Q96L93_S662 | 0.685784035 | 4.88E-04 | 3.38E-03 |
| PP12C_HUMAN\|Q9BZL4_S453 | 0.685755537 | 4.55E-05 | 1.95E-03 |
| FCHO2_HUMAN\|Q0JRZ9_S509 | 0.685748492 | 2.41E-04 | 1.91E-03 |
| K1C16_HUMAN\|P08779_S449 | 0.685722137 | 4.11E-03 | 2.83E-02 |
| RTKN_HUMAN\|Q9BST9_S220 | 0.685719104 | 2.72E-03 | 1.96E-02 |
| TBCD5_HUMAN\|Q92609_S554 | 0.685661579 | 2.23E-03 | 4.57E-02 |
| ZCRB1_HUMAN\|Q8TBF4_S216 | 0.68565833 | 3.18E-04 | 1.41E-02 |

| | | | |
|---|---|---|---|
| TP53B_HUMAN\|Q12888_S366_T357 | 0.685637844 | 1.53E-03 | 1.92E-02 |
| ARP19_HUMAN\|P56211_Y59 | 0.685637459 | 7.12E-04 | 1.67E-02 |
| MAP7_HUMAN\|Q14244_S183 | 0.685603128 | 2.98E-04 | 6.21E-03 |
| BCL9L_HUMAN\|Q86UU0_S21 | 0.685469976 | 9.42E-04 | 9.45E-03 |
| ITB4_HUMAN\|P16144_T1455 | 0.68525715 | 4.36E-03 | 2.81E-02 |
| CLAP1_HUMAN\|Q7Z460-4_S691 | 0.685243054 | 5.66E-03 | 3.56E-02 |
| RHG23_HUMAN\|Q9P227_S611 | 0.685225787 | 4.37E-04 | 2.84E-03 |
| OSTF1_HUMAN\|Q92882_S213 | 0.685165631 | 1.74E-03 | 2.34E-02 |
| WDR13_HUMAN\|Q9H1Z4_S79 | 0.685156243 | 1.15E-03 | 1.10E-02 |
| SRC8_HUMAN\|Q14247_T401 | 0.685132809 | 5.49E-04 | 3.42E-03 |
| TACC2_HUMAN\|O95359_S1562 | 0.685091648 | 9.97E-04 | 2.08E-02 |
| SMAG1_HUMAN\|Q9UPU9_S420_T424 | 0.685040875 | 1.70E-04 | 4.17E-03 |
| DLG1_HUMAN\|Q12959_S575 | 0.684835905 | 3.44E-03 | 3.65E-02 |
| K1C14_HUMAN\|P02533_S433_S438 | 0.684822227 | 5.17E-03 | 3.35E-02 |
| NUP98_HUMAN\|P52948_S612 | 0.684821408 | 7.39E-04 | 6.30E-03 |
| ALKB5_HUMAN\|Q6P6C2_S384 | 0.684810899 | 1.25E-04 | 1.08E-03 |
| IF4B_HUMAN\|P23588_S497_S498_S504 | 0.684798072 | 3.11E-04 | 3.27E-03 |
| TBD2A_HUMAN\|Q9BYX2_S920 | 0.684796828 | 5.72E-04 | 3.50E-03 |
| CCNH_HUMAN\|P51946_T315 | 0.684776775 | 3.41E-03 | 4.81E-02 |
| LMNB2_HUMAN\|Q03252_S424 | 0.684726339 | 3.60E-03 | 4.87E-02 |
| DIDO1_HUMAN\|Q9BTC0_S523 | 0.684712197 | 7.59E-05 | 2.74E-03 |
| SF3B1_HUMAN\|O75533_T207_T211 | 0.684697922 | 8.57E-03 | 3.55E-02 |
| CE170_HUMAN\|Q5SW79_S636_T628 | 0.684678658 | 3.19E-03 | 2.29E-02 |
| INT12_HUMAN\|Q96CB8_S389 | 0.684596068 | 9.59E-03 | 4.40E-02 |
| TBCD5_HUMAN\|Q92609_S554 | 0.68459384 | 2.15E-03 | 1.46E-02 |
| ITBP1_HUMAN\|O14713_S10_S14 | 0.684585021 | 7.12E-04 | 4.17E-03 |
| SWAHC_HUMAN\|Q53LP3_S212 | 0.684571806 | 1.31E-04 | 2.04E-03 |
| ZO2_HUMAN\|Q9UDY2_S400 | 0.684555726 | 1.21E-02 | 3.99E-02 |
| PTN14_HUMAN\|Q15678_S593_S594 | 0.684503667 | 1.23E-04 | 1.97E-03 |
| COHA1_HUMAN\|Q9UMD9_S172 | 0.684485038 | 9.48E-05 | 3.15E-03 |
| BORG5_HUMAN\|Q00587_S303 | 0.684467019 | 5.08E-04 | 1.90E-02 |

| | | | |
|---|---|---|---|
| SCAF8_HUMAN\|Q9UPN6_S617 | 0.684353315 | 1.13E-03 | 6.06E-03 |
| ABI2_HUMAN\|Q9NYB9_Y371 | 0.684338988 | 3.02E-04 | 3.84E-03 |
| F120A_HUMAN\|Q9NZB2_S1023 | 0.684218551 | 5.08E-04 | 9.52E-03 |
| PDLI1_HUMAN\|O00151_S130 | 0.684196715 | 8.38E-04 | 1.29E-02 |
| NCOA7_HUMAN\|Q8NI08_S208_T210 | 0.684147684 | 6.04E-04 | 1.49E-02 |
| DPYL2_HUMAN\|Q16555_T509 | 0.684121182 | 3.12E-05 | 6.87E-04 |
| DC1L2_HUMAN\|O43237_S407 | 0.684085258 | 7.25E-04 | 4.71E-03 |
| FA83B_HUMAN\|Q5T0W9_S664 | 0.684061292 | 2.83E-04 | 6.35E-03 |
| PAK6_HUMAN\|Q9NQU5_S347_S351 | 0.683985806 | 6.82E-03 | 4.06E-02 |
| IASPP_HUMAN\|Q8WUF5_Y109 | 0.683975633 | 6.74E-04 | 4.42E-03 |
| MA7D1_HUMAN\|Q3KQU3_S460 | 0.683861004 | 8.96E-04 | 5.57E-03 |
| LAD1_HUMAN\|O00515_S301 | 0.68379375 | 3.86E-03 | 1.84E-02 |
| AHNK_HUMAN\|Q09666_T5729 | 0.683775439 | 2.54E-04 | 1.87E-03 |
| CAMP2_HUMAN\|Q08AD1_S1313 | 0.683742623 | 1.22E-03 | 7.14E-03 |
| PLXA2_HUMAN\|O75051_S1630_S1633 | 0.683660174 | 2.96E-03 | 1.48E-02 |
| SON_HUMAN\|P18583_S94 | 0.683619936 | 1.12E-04 | 3.01E-03 |
| MTUS1_HUMAN\|Q9ULD2_S1264_S1268 | 0.683544681 | 2.56E-03 | 4.90E-02 |
| ZC3HD_HUMAN\|Q5T200_S242 | 0.683505347 | 1.18E-04 | 1.04E-03 |
| MLF2_HUMAN\|Q15773_S238_S243 | 0.683458273 | 2.31E-03 | 1.22E-02 |
| 1A01_HUMAN\|P30443_S352_S359 | 0.683392616 | 4.69E-03 | 2.97E-02 |
| HEAT6_HUMAN\|Q6AI08_S643 | 0.683317236 | 5.91E-05 | 6.18E-04 |
| DOCK6_HUMAN\|Q96HP0_S882 | 0.683283319 | 2.94E-03 | 2.15E-02 |
| MICA3_HUMAN\|Q7RTP6_S685 | 0.683279228 | 2.28E-03 | 1.52E-02 |
| PCSK9_HUMAN\|Q8NBP7_S688 | 0.683279016 | 2.70E-03 | 3.10E-02 |
| LIMD1_HUMAN\|Q9UGP4_S424 | 0.683267342 | 7.21E-04 | 1.18E-02 |
| TRAF7_HUMAN\|Q6Q0C0_S108 | 0.683247952 | 4.27E-03 | 3.63E-02 |
| M3K20_HUMAN\|Q9NYL2_S667 | 0.683244114 | 4.25E-06 | 1.75E-04 |
| TCOF_HUMAN\|Q13428_T249 | 0.68316269 | 1.96E-03 | 4.49E-02 |
| YAP1_HUMAN\|P46937_S163 | 0.68312479 | 1.18E-02 | 4.60E-02 |
| SPTN1_HUMAN\|Q13813_S1190 | 0.68310354 | 2.61E-04 | 1.90E-03 |
| CND1_HUMAN\|Q15021_S1333 | 0.683082331 | 7.67E-04 | 2.47E-02 |

| | | | |
|---|---|---|---|
| UBP24_HUMAN\|Q9UPU5_S1141 | 0.683055093 | 1.01E-03 | 9.84E-03 |
| AGFG1_HUMAN\|P52594_S181 | 0.682973158 | 7.19E-05 | 7.34E-04 |
| RU17_HUMAN\|P08621_S226 | 0.682968063 | 1.36E-03 | 1.90E-02 |
| NFIC_HUMAN\|P08651_S305 | 0.682889912 | 2.14E-03 | 2.67E-02 |
| BD1L1_HUMAN\|Q8NFC6_S2986 | 0.682843995 | 9.59E-03 | 3.34E-02 |
| TB182_HUMAN\|Q9C0C2_S872 | 0.682833575 | 4.48E-04 | 5.19E-03 |
| EPN2_HUMAN\|O95208_S172 | 0.682813019 | 7.54E-03 | 4.37E-02 |
| MAP4_HUMAN\|P27816_S941 | 0.682761698 | 1.77E-03 | 2.20E-02 |
| ZC3H1_HUMAN\|O60293_S805 | 0.682753964 | 3.15E-03 | 2.96E-02 |
| PAIRB_HUMAN\|Q8NC51_S392 | 0.682732951 | 2.09E-03 | 3.48E-02 |
| PRC2C_HUMAN\|Q9Y520_S187_S189 | 0.682668181 | 2.13E-03 | 1.01E-02 |
| TBC17_HUMAN\|Q9HA65_T620 | 0.682605285 | 1.15E-03 | 6.12E-03 |
| PLEC_HUMAN\|Q15149_S4389_S4391 | 0.682569718 | 3.82E-03 | 3.28E-02 |
| OSTF1_HUMAN\|Q92882_S213 | 0.682522074 | 1.59E-03 | 1.32E-02 |
| LAD1_HUMAN\|O00515_S38 | 0.682467803 | 1.06E-03 | 2.17E-02 |
| RPTOR_HUMAN\|Q8N122_S859_T865 | 0.682428414 | 1.08E-02 | 4.30E-02 |
| NUP98_HUMAN\|P52948_S623 | 0.682246652 | 3.14E-03 | 1.92E-02 |
| SPD2A_HUMAN\|Q5TCZ1_S644 | 0.682241433 | 4.55E-03 | 3.71E-02 |
| PDLI4_HUMAN\|P50479_T114 | 0.682217592 | 4.87E-04 | 5.51E-03 |
| CYBR1_HUMAN\|Q53TN4_T285 | 0.682215961 | 4.87E-03 | 3.03E-02 |
| LARP1_HUMAN\|Q6PKG0_S1040 | 0.682208103 | 4.20E-04 | 1.16E-02 |
| IF4B_HUMAN\|P23588_S489_S497_S498 | 0.682161665 | 4.91E-04 | 4.69E-02 |
| KLC1_HUMAN\|Q07866-4_S561 | 0.682147427 | 3.30E-04 | 2.43E-03 |
| 1433Z_HUMAN\|P63104_T232 | 0.682034853 | 6.61E-03 | 2.87E-02 |
| RL22L_HUMAN\|Q6P5R6_S120 | 0.68203426 | 2.34E-03 | 1.09E-02 |
| RBM7_HUMAN\|Q9Y580_S204 | 0.681975657 | 1.82E-03 | 1.27E-02 |
| AGFG1_HUMAN\|P52594_S181 | 0.681973034 | 6.84E-05 | 1.22E-03 |
| NKTR_HUMAN\|P30414_T1155 | 0.681897651 | 1.09E-03 | 5.93E-03 |
| ZN185_HUMAN\|O15231_S465 | 0.681886199 | 1.42E-02 | 4.57E-02 |
| LMNB2_HUMAN\|Q03252_S420_S424 | 0.681854915 | 6.83E-03 | 4.06E-02 |
| DSC3_HUMAN\|Q14574-2_S833 | 0.681815309 | 1.15E-02 | 3.85E-02 |

| | | | |
|---|---|---|---|
| DDX17_HUMAN\|Q92841_S571 | 0.68176221 | 1.43E-03 | 8.09E-03 |
| ADNP_HUMAN\|Q9H2P0_S709 | 0.681745819 | 5.44E-05 | 2.18E-03 |
| TRI47_HUMAN\|Q96LD4_S588 | 0.68167815 | 1.85E-03 | 4.17E-02 |
| AHNK_HUMAN\|Q09666_S5749 | 0.681627529 | 4.32E-05 | 4.73E-04 |
| F193A_HUMAN\|P78312_S1144 | 0.68158666 | 9.72E-04 | 5.39E-03 |
| MED1_HUMAN\|Q15648_Y1431 | 0.681575756 | 1.48E-04 | 4.22E-03 |
| PRC2C_HUMAN\|Q9Y520_S2694_T2691 | 0.681538854 | 5.33E-03 | 3.41E-02 |
| RNF25_HUMAN\|Q96BH1_S450 | 0.681446805 | 2.60E-03 | 4.06E-02 |
| CE170_HUMAN\|Q5SW79_S971 | 0.68143517 | 7.30E-04 | 7.41E-03 |
| IRS1_HUMAN\|P35568_S1101 | 0.681372158 | 2.87E-04 | 2.17E-03 |
| RIOK1_HUMAN\|Q9BRS2_S22 | 0.681359087 | 7.06E-06 | 1.15E-04 |
| BNIP2_HUMAN\|Q12982_S114 | 0.681259305 | 5.39E-03 | 4.58E-02 |
| RBM7_HUMAN\|Q9Y580_S137 | 0.68120317 | 1.90E-04 | 4.84E-03 |
| K1C14_HUMAN\|P02533_S437_S438 | 0.681192571 | 3.45E-03 | 1.68E-02 |
| M3K20_HUMAN\|Q9NYL2_S633 | 0.681186656 | 8.58E-03 | 4.77E-02 |
| IF4B_HUMAN\|P23588_T420 | 0.681181103 | 9.80E-04 | 2.07E-02 |
| TOM1_HUMAN\|O60784_S462 | 0.681137012 | 2.67E-03 | 2.85E-02 |
| BACH1_HUMAN\|O14867_S445 | 0.68110827 | 8.74E-04 | 8.66E-03 |
| KLC3_HUMAN\|Q6P597_S167 | 0.681053921 | 2.05E-03 | 1.67E-02 |
| IF4B_HUMAN\|P23588_T420 | 0.681030721 | 9.74E-04 | 1.43E-02 |
| NDRG1_HUMAN\|Q92597_S336_T328 | 0.680945989 | 3.47E-03 | 1.50E-02 |
| PTN14_HUMAN\|Q15678_S593_S594 | 0.680851618 | 1.05E-04 | 1.00E-03 |
| PDLI2_HUMAN\|Q96JY6_S134_S137_S143 | 0.680828847 | 2.37E-03 | 1.24E-02 |
| MA7D1_HUMAN\|Q3KQU3_S460 | 0.680814115 | 7.99E-04 | 8.02E-03 |
| LIPA1_HUMAN\|Q13136_S1172 | 0.680792425 | 2.04E-04 | 4.79E-03 |
| IP3KB_HUMAN\|P27987_S264 | 0.680763471 | 4.25E-03 | 2.91E-02 |
| NUP98_HUMAN\|P52948_T1000 | 0.680718781 | 5.28E-03 | 4.21E-02 |
| PRSR2_HUMAN\|Q86WR7_S212 | 0.68070451 | 3.26E-04 | 9.92E-03 |
| FUBP3_HUMAN\|Q96I24_S569 | 0.680672296 | 4.91E-03 | 2.25E-02 |
| FA83H_HUMAN\|Q6ZRV2_T938 | 0.680653446 | 6.05E-03 | 2.29E-02 |
| TB182_HUMAN\|Q9C0C2_S435 | 0.680622141 | 5.96E-03 | 3.51E-02 |

| | | | |
|---|---|---|---|
| MGMT_HUMAN\|P16455_S201 | 0.680585008 | 1.55E-05 | 8.15E-04 |
| PKHA4_HUMAN\|Q9H4M7_S194 | 0.680580767 | 1.68E-03 | 9.26E-03 |
| SVIL_HUMAN\|O95425_T679 | 0.6805711 | 6.00E-05 | 6.27E-04 |
| HNRL2_HUMAN\|Q1KMD3_S161_S168 | 0.680498036 | 5.82E-05 | 5.79E-04 |
| EFHD2_HUMAN\|Q96C19_S74 | 0.680423704 | 1.07E-03 | 5.83E-03 |
| Z280C_HUMAN\|Q8ND82_S80 | 0.680342262 | 5.10E-04 | 1.36E-02 |
| ACLY_HUMAN\|P53396_S455 | 0.680295869 | 3.46E-05 | 3.95E-04 |
| TP53B_HUMAN\|Q12888_S366 | 0.680238685 | 9.55E-05 | 2.82E-03 |
| EPN2_HUMAN\|O95208_S173 | 0.680186036 | 2.41E-03 | 1.12E-02 |
| SRGP1_HUMAN\|Q7Z6B7_S906 | 0.680179571 | 1.24E-03 | 6.57E-03 |
| TBCD5_HUMAN\|Q92609_S541 | 0.680151607 | 6.77E-06 | 1.11E-04 |
| LAP2A_HUMAN\|P42166_S159 | 0.68005757 | 9.54E-04 | 5.84E-03 |
| P4K2A_HUMAN\|Q9BTU6_S468 | 0.680042325 | 6.46E-03 | 3.74E-02 |
| RANB3_HUMAN\|Q9H6Z4_T124 | 0.680038115 | 4.47E-03 | 1.83E-02 |
| RIOK1_HUMAN\|Q9BRS2_S21 | 0.679967886 | 3.98E-05 | 8.31E-04 |
| IRS2_HUMAN\|Q9Y4H2_S1185_S1203 | 0.679863707 | 1.86E-03 | 4.33E-02 |
| FA83H_HUMAN\|Q6ZRV2_T938 | 0.679848317 | 5.92E-03 | 2.62E-02 |
| IF4B_HUMAN\|P23588_T420 | 0.67984639 | 9.31E-04 | 7.50E-03 |
| CRBG1_HUMAN\|Q9Y4K1_S22 | 0.679842562 | 1.19E-04 | 1.11E-03 |
| MTCL1_HUMAN\|Q9Y4B5_S77 | 0.679765435 | 2.40E-03 | 1.86E-02 |
| MKL2_HUMAN\|Q9ULH7_S921 | 0.679750066 | 1.08E-03 | 2.94E-02 |
| PP1R7_HUMAN\|Q15435_S24 | 0.679714633 | 7.50E-03 | 4.17E-02 |
| CAVN3_HUMAN\|Q969G5_T205 | 0.679695926 | 3.46E-03 | 3.80E-02 |
| CSKI2_HUMAN\|Q8WXE0_S406 | 0.679674856 | 4.13E-05 | 1.84E-03 |
| SPD2A_HUMAN\|Q5TCZ1_S1043 | 0.679673946 | 3.66E-03 | 3.82E-02 |
| FKBP5_HUMAN\|Q13451_T443 | 0.67967368 | 1.30E-02 | 4.97E-02 |
| ALDOA_HUMAN\|P04075_S46 | 0.679651847 | 1.56E-03 | 7.90E-03 |
| ARP19_HUMAN\|P56211_Y59 | 0.679605634 | 5.59E-04 | 6.15E-03 |
| PRAG1_HUMAN\|Q86YV5_S745 | 0.67960276 | 1.55E-03 | 1.13E-02 |
| VIME_HUMAN\|P08670_S430 | 0.679515586 | 7.89E-04 | 8.18E-03 |
| SH24A_HUMAN\|Q9H788_S316 | 0.679392196 | 2.69E-03 | 4.13E-02 |

| | | | |
|---|---|---|---|
| NU153_HUMAN\|P49790_S334 | 0.679340018 | 9.81E-04 | 2.85E-02 |
| VIME_HUMAN\|P08670_S56 | 0.679315759 | 3.05E-04 | 3.86E-03 |
| PHLB2_HUMAN\|Q86SQ0_S384_S387 | 0.679204321 | 3.55E-03 | 1.53E-02 |
| YAP1_HUMAN\|P46937_S105_S109 | 0.679187997 | 3.74E-05 | 1.48E-03 |
| GSK3A_HUMAN\|P49840_T19 | 0.67918767 | 1.94E-04 | 4.63E-03 |
| PHF8_HUMAN\|Q9UPP1_S804 | 0.679165766 | 2.93E-04 | 4.73E-03 |
| MTMR2_HUMAN\|Q13614_S631 | 0.679158097 | 1.21E-03 | 1.09E-02 |
| AGFG1_HUMAN\|P52594_S181 | 0.679031538 | 5.91E-05 | 1.94E-03 |
| LMNA_HUMAN\|P02545_S404_S406 | 0.678972731 | 1.07E-03 | 9.94E-03 |
| MAP1A_HUMAN\|P78559_S2022 | 0.678967223 | 4.07E-04 | 2.70E-03 |
| K2C5_HUMAN\|P13647_S571 | 0.678966726 | 3.19E-04 | 7.02E-03 |
| CDK16_HUMAN\|Q00536_T135 | 0.678911522 | 2.44E-03 | 1.27E-02 |
| PRAG1_HUMAN\|Q86YV5_S712 | 0.678891985 | 1.42E-04 | 3.78E-03 |
| PLEC_HUMAN\|Q15149_Y4615 | 0.678791076 | 9.67E-04 | 1.51E-02 |
| AF1L2_HUMAN\|Q8N4X5_S344 | 0.678750532 | 8.14E-04 | 5.18E-03 |
| TBCD4_HUMAN\|O60343_S318 | 0.67872944 | 5.49E-04 | 3.42E-03 |
| PEAK1_HUMAN\|Q9H792_S1036 | 0.678633951 | 1.02E-04 | 2.88E-03 |
| PDLI2_HUMAN\|Q96JY6_S137 | 0.678603909 | 3.64E-04 | 1.07E-02 |
| PLXA2_HUMAN\|O75051_S1630_S1633 | 0.678589549 | 2.52E-03 | 1.63E-02 |
| AKAP2_HUMAN\|Q9Y2D5_S720 | 0.678527082 | 3.24E-04 | 2.27E-03 |
| ZC11A_HUMAN\|O75152_S132 | 0.678519312 | 5.78E-05 | 2.07E-03 |
| RBM7_HUMAN\|Q9Y580_S137 | 0.678509085 | 1.69E-04 | 4.16E-03 |
| VIME_HUMAN\|P08670_S325 | 0.678495616 | 1.09E-02 | 4.33E-02 |
| TR150_HUMAN\|Q9Y2W1_T874 | 0.678423578 | 2.38E-03 | 1.85E-02 |
| PPR35_HUMAN\|Q8TAP8_S52 | 0.678368557 | 1.71E-03 | 9.39E-03 |
| CTU2_HUMAN\|Q2VPK5_S419 | 0.67826217 | 6.99E-05 | 2.89E-03 |
| TBCD4_HUMAN\|O60343_T752 | 0.678257209 | 2.99E-03 | 2.85E-02 |
| PDLI5_HUMAN\|Q96HC4_S313 | 0.678171911 | 1.41E-03 | 1.89E-02 |
| MAN1_HUMAN\|Q9Y2U8_S144 | 0.678139691 | 9.44E-03 | 4.91E-02 |
| OPTN_HUMAN\|Q96CV9_S526 | 0.678106718 | 4.56E-04 | 4.37E-03 |
| YAP1_HUMAN\|P46937_S109 | 0.678053383 | 1.18E-03 | 6.94E-03 |

| | | | |
|---|---|---|---|
| 4EBP1_HUMAN\|Q13541_S44 | 0.678029811 | 9.30E-04 | 1.47E-02 |
| RTKN_HUMAN\|Q9BST9_S529 | 0.677972326 | 1.41E-03 | 7.27E-03 |
| SVIL_HUMAN\|O95425_T679 | 0.677963628 | 5.31E-05 | 5.39E-04 |
| MISP_HUMAN\|Q8IVT2_T377 | 0.677906686 | 1.10E-03 | 5.96E-03 |
| MYPT1_HUMAN\|O14974_T443 | 0.677836039 | 7.43E-04 | 7.50E-03 |
| PKHA4_HUMAN\|Q9H4M7_S194 | 0.677825775 | 1.53E-03 | 3.85E-02 |
| ATX2L_HUMAN\|Q8WWM7_S339 | 0.677824965 | 1.19E-05 | 1.69E-04 |
| GSK3A_HUMAN\|P49840_T19 | 0.67774963 | 1.82E-04 | 4.82E-03 |
| K1C16_HUMAN\|P08779_S446_S449 | 0.677734582 | 2.08E-03 | 9.95E-03 |
| OSBL8_HUMAN\|Q9BZF1_S807_S808_S810 | 0.677725611 | 4.16E-03 | 2.35E-02 |
| SAFB1_HUMAN\|Q15424_S604 | 0.677720595 | 3.11E-05 | 1.60E-03 |
| TB182_HUMAN\|Q9C0C2_S1103 | 0.677642516 | 1.56E-03 | 1.95E-02 |
| KLC2_HUMAN\|Q9H0B6_S608_S611 | 0.677580593 | 8.10E-05 | 3.25E-03 |
| RING1_HUMAN\|Q06587_S38 | 0.677579633 | 1.05E-03 | 9.83E-03 |
| UBXN7_HUMAN\|O94888_S285_S288 | 0.677569133 | 7.67E-03 | 2.78E-02 |
| TISB_HUMAN\|Q07352_S334 | 0.677498341 | 4.13E-03 | 4.32E-02 |
| SIPA1_HUMAN\|Q96FS4_S1036 | 0.677484606 | 9.30E-04 | 7.50E-03 |
| K1671_HUMAN\|Q9BY89_S127 | 0.677462535 | 3.49E-04 | 7.44E-03 |
| ZO1_HUMAN\|Q07157_S1617 | 0.677411107 | 3.82E-04 | 3.84E-03 |
| PRAG1_HUMAN\|Q86YV5_S907 | 0.677392706 | 3.65E-04 | 7.77E-03 |
| AHNK_HUMAN\|Q09666_S216_S220 | 0.677162724 | 2.03E-04 | 4.79E-03 |
| SPTB2_HUMAN\|Q01082-3_S14_S8 | 0.677105507 | 4.73E-04 | 8.93E-03 |
| RAB12_HUMAN\|Q6IQ22_S21 | 0.677018736 | 4.78E-03 | 3.17E-02 |
| PKP3_HUMAN\|Q9Y446_S313 | 0.676946359 | 2.48E-05 | 3.03E-04 |
| DLG1_HUMAN\|Q12959_S575 | 0.676943552 | 2.68E-03 | 1.95E-02 |
| AHNK_HUMAN\|Q09666_S3412 | 0.676849504 | 1.13E-03 | 2.28E-02 |
| KIF1B_HUMAN\|O60333_S1057 | 0.676801487 | 4.96E-04 | 3.43E-03 |
| ENSA_HUMAN\|O43768_S109 | 0.676792715 | 1.92E-03 | 3.36E-02 |
| ANR17_HUMAN\|O75179_S1693 | 0.67669416 | 3.31E-04 | 4.05E-03 |
| PARP4_HUMAN\|Q9UKK3_S1722 | 0.676657953 | 7.17E-04 | 2.40E-02 |
| 1B07_HUMAN\|P01889_S352 | 0.676652644 | 2.65E-03 | 3.07E-02 |

| | | | |
|---|---|---|---|
| PRC2A_HUMAN\|P48634_S761_S764 | 0.676637361 | 7.94E-05 | 1.35E-03 |
| CD2AP_HUMAN\|Q9Y5K6_S458 | 0.676540039 | 1.09E-03 | 6.50E-03 |
| PRSR2_HUMAN\|Q86WR7_S212 | 0.676474869 | 2.72E-04 | 5.91E-03 |
| K1522_HUMAN\|Q9P206_S406 | 0.676427852 | 1.76E-03 | 1.49E-02 |
| BL1S3_HUMAN\|Q6QNY0_S65_T63 | 0.676387076 | 5.09E-04 | 3.21E-03 |
| PKHA1_HUMAN\|Q9HB21_S177 | 0.676341722 | 1.07E-02 | 3.65E-02 |
| PDLI5_HUMAN\|Q96HC4_S111 | 0.676295082 | 7.58E-04 | 1.76E-02 |
| PDLI2_HUMAN\|Q96JY6_S137_S143 | 0.676284696 | 7.32E-04 | 1.25E-02 |
| CRTC3_HUMAN\|Q6UUV7_S370 | 0.676283256 | 4.43E-03 | 1.82E-02 |
| SAV1_HUMAN\|Q9H4B6_S27 | 0.676281951 | 1.36E-04 | 3.53E-03 |
| GGYF1_HUMAN\|O75420_S24 | 0.676272284 | 4.19E-04 | 1.16E-02 |
| MILK1_HUMAN\|Q8N3F8_S640_S644 | 0.676208613 | 7.39E-04 | 4.31E-03 |
| PHIP_HUMAN\|Q8WWQ0_S1783 | 0.676032464 | 1.76E-04 | 3.30E-03 |
| P4K2A_HUMAN\|Q9BTU6_S462_S468 | 0.676019834 | 2.68E-04 | 5.81E-03 |
| ZYX_HUMAN\|Q15942_S259_S267 | 0.675976963 | 1.05E-03 | 9.85E-03 |
| STRN3_HUMAN\|Q13033_S202 | 0.675916105 | 1.19E-04 | 1.11E-03 |
| IF4B_HUMAN\|P23588_S497_S498_S504 | 0.675860375 | 2.10E-04 | 7.03E-03 |
| TBC16_HUMAN\|Q8TBP0_S103_T99 | 0.675802629 | 1.64E-03 | 1.17E-02 |
| K1C14_HUMAN\|P02533_S429_S435 | 0.675787662 | 3.87E-03 | 1.84E-02 |
| PDLI2_HUMAN\|Q96JY6_S134 | 0.675726965 | 1.44E-05 | 4.00E-04 |
| CE170_HUMAN\|Q5SW79_S971 | 0.675704406 | 5.87E-04 | 1.05E-02 |
| DGC14_HUMAN\|Q96DF8_T386 | 0.675613664 | 2.49E-04 | 1.84E-03 |
| IF4G1_HUMAN\|Q04637_S1231 | 0.675404229 | 6.98E-04 | 6.06E-03 |
| BAIP2_HUMAN\|Q9UQB8_S454 | 0.675377549 | 2.96E-04 | 3.81E-03 |
| CSKI2_HUMAN\|Q8WXE0_S403 | 0.675281496 | 1.12E-04 | 3.49E-03 |
| HMGA2_HUMAN\|P52926_S14 | 0.675272573 | 5.31E-04 | 3.63E-03 |
| SPD2A_HUMAN\|Q5TCZ1_S1017 | 0.675154594 | 1.43E-02 | 4.61E-02 |
| NCOA7_HUMAN\|Q8NI08_S211 | 0.675088763 | 4.43E-04 | 5.11E-03 |
| PLEC_HUMAN\|Q15149_S4385_S4386 | 0.675009579 | 5.87E-06 | 3.46E-04 |
| M3K20_HUMAN\|Q9NYL2_S667 | 0.674977392 | 2.63E-06 | 5.48E-05 |
| NSD3_HUMAN\|Q9BZ95_S457 | 0.674955643 | 4.50E-04 | 5.20E-03 |

| | | | |
|---|---|---|---|
| PHLB2_HUMAN\|Q86SQ0_S565 | 0.674937868 | 2.23E-04 | 1.81E-03 |
| K1671_HUMAN\|Q9BY89_S128 | 0.674928753 | 7.90E-05 | 2.81E-03 |
| SAV1_HUMAN\|Q9H4B6_S94 | 0.674904354 | 1.01E-04 | 1.73E-03 |
| ARG28_HUMAN\|Q8N1W1_S476_S486 | 0.674884321 | 1.67E-03 | 2.29E-02 |
| DBNL_HUMAN\|Q9UJU6_S269 | 0.674840378 | 3.58E-03 | 4.87E-02 |
| 1B07_HUMAN\|P01889_S352_S356 | 0.674834032 | 1.89E-03 | 1.02E-02 |
| PML_HUMAN\|P29590_S504 | 0.674832562 | 4.05E-04 | 6.63E-03 |
| 1A01_HUMAN\|P30443_S356_S359 | 0.674771486 | 2.15E-03 | 1.02E-02 |
| FA83H_HUMAN\|Q6ZRV2_S974 | 0.674667322 | 2.90E-04 | 1.35E-02 |
| PALLD_HUMAN\|Q8WX93_S893 | 0.674610417 | 1.63E-03 | 3.81E-02 |
| KAZRN_HUMAN\|Q674X7-3_S8 | 0.674578652 | 4.43E-04 | 2.87E-03 |
| NFIC_HUMAN\|P08651_S339 | 0.67452147 | 3.67E-03 | 1.57E-02 |
| BI2L1_HUMAN\|Q9UHR4_S261_T257 | 0.674507807 | 1.94E-04 | 1.62E-03 |
| CLAP2_HUMAN\|O75122_S1029 | 0.674474095 | 1.84E-03 | 1.28E-02 |
| OSTF1_HUMAN\|Q92882_S213 | 0.674442214 | 1.22E-03 | 1.73E-02 |
| AHNK_HUMAN\|Q09666_S5763 | 0.674356424 | 3.19E-05 | 3.85E-03 |
| SARG_HUMAN\|Q9BW04_T456 | 0.674288948 | 3.32E-05 | 7.26E-04 |
| ZYX_HUMAN\|Q15942_S259 | 0.674272113 | 2.98E-04 | 6.21E-03 |
| SRSF9_HUMAN\|Q13242_S193 | 0.674179344 | 4.02E-04 | 6.05E-03 |
| BCAR3_HUMAN\|O75815_S370 | 0.674153263 | 4.06E-03 | 1.70E-02 |
| SPTB2_HUMAN\|Q01082-3_S8 | 0.674151954 | 1.42E-03 | 2.68E-02 |
| IRS2_HUMAN\|Q9Y4H2_T520 | 0.674131411 | 3.01E-03 | 1.34E-02 |
| ARG28_HUMAN\|Q8N1W1_S476_S486 | 0.674088637 | 1.63E-03 | 1.40E-02 |
| PP14A_HUMAN\|Q96A00_S26 | 0.67393538 | 1.50E-04 | 4.23E-03 |
| RHG12_HUMAN\|Q8IWW6_S215 | 0.673924031 | 4.88E-04 | 5.46E-03 |
| MAP7_HUMAN\|Q14244_T184 | 0.673869033 | 4.57E-04 | 8.55E-03 |
| 1433B_HUMAN\|P31946_S47 | 0.673860607 | 6.67E-03 | 2.89E-02 |
| NCK5L_HUMAN\|Q9HCH0_S489_S494 | 0.673788705 | 5.16E-04 | 9.45E-03 |
| ARH40_HUMAN\|Q8TER5_S1474 | 0.673784674 | 4.66E-04 | 4.46E-03 |
| TB182_HUMAN\|Q9C0C2_S983 | 0.673706773 | 1.65E-03 | 9.11E-03 |
| RAI14_HUMAN\|Q9P0K7_S296 | 0.673643918 | 2.64E-03 | 1.35E-02 |

| | | | |
|---|---|---|---|
| ZN185_HUMAN\|O15231_S153_S155_T154 | 0.6736282 | 4.81E-03 | 2.61E-02 |
| 1B07_HUMAN\|P01889_S352_S356 | 0.673624757 | 1.82E-03 | 8.91E-03 |
| CHM2A_HUMAN\|O43633_S203 | 0.673577356 | 2.74E-04 | 1.98E-03 |
| C2C4C_HUMAN\|Q8TF44_S156 | 0.673476419 | 2.57E-03 | 1.33E-02 |
| 1B07_HUMAN\|P01889_S359 | 0.673475558 | 4.30E-04 | 8.69E-03 |
| STK10_HUMAN\|O94804_T952 | 0.673266515 | 3.53E-03 | 4.87E-02 |
| ARMX3_HUMAN\|Q9UH62_S110 | 0.673217757 | 4.56E-04 | 8.74E-03 |
| K1522_HUMAN\|Q9P206_S929 | 0.673196765 | 1.34E-04 | 1.21E-03 |
| FOXO3_HUMAN\|O43524_S253 | 0.673194738 | 4.73E-05 | 1.66E-03 |
| KIF1C_HUMAN\|O43896_S817 | 0.673159762 | 5.19E-04 | 9.97E-03 |
| PRC2A_HUMAN\|P48634_S1219 | 0.673096594 | 2.58E-04 | 5.64E-03 |
| MRP4_HUMAN\|O15439_T646 | 0.673045177 | 4.30E-03 | 2.78E-02 |
| K1C14_HUMAN\|P02533_S429 | 0.673000462 | 1.36E-02 | 4.43E-02 |
| MTAP2_HUMAN\|P11137_S1782_S1787_S1790 | 0.672984227 | 3.81E-04 | 2.74E-03 |
| MTAP2_HUMAN\|P11137_S1782_S1786_S1790 | 0.672973071 | 4.68E-04 | 9.39E-03 |
| MILK1_HUMAN\|Q8N3F8_S391 | 0.672940712 | 1.04E-05 | 5.98E-04 |
| BORG5_HUMAN\|Q00587_S303 | 0.672936138 | 3.26E-04 | 2.41E-03 |
| MISP_HUMAN\|Q8IVT2_S395_S397 | 0.672892103 | 1.71E-03 | 8.46E-03 |
| LARP4_HUMAN\|Q71RC2_S597 | 0.67289124 | 5.92E-04 | 1.08E-02 |
| PSF2_HUMAN\|Q9Y248_S179_S182 | 0.672890128 | 9.79E-04 | 9.66E-03 |
| DESP_HUMAN\|P15924_S165 | 0.672812521 | 5.55E-03 | 2.89E-02 |
| PDLI2_HUMAN\|Q96JY6_S134_S137_S143 | 0.672803434 | 1.84E-03 | 3.25E-02 |
| COHA1_HUMAN\|Q9UMD9_S78 | 0.67273791 | 9.93E-03 | 4.01E-02 |
| K2C5_HUMAN\|P13647_S571_S574 | 0.672728809 | 1.53E-03 | 2.00E-02 |
| MLF2_HUMAN\|Q15773_S238_S243 | 0.672696481 | 1.63E-03 | 1.40E-02 |
| AHNK_HUMAN\|Q09666_T4766 | 0.672686804 | 1.09E-03 | 5.92E-03 |
| RTKN_HUMAN\|Q9BST9_S529 | 0.672600054 | 1.18E-03 | 1.12E-02 |
| ITB4_HUMAN\|P16144_S1457 | 0.672597857 | 2.35E-03 | 1.84E-02 |
| PLCH2_HUMAN\|O75038_S595 | 0.672569587 | 3.28E-04 | 2.29E-03 |
| IF4G2_HUMAN\|P78344_S17 | 0.67244586 | 5.69E-06 | 9.65E-05 |
| K1C14_HUMAN\|P02533_S429_S435 | 0.672385129 | 3.49E-03 | 3.81E-02 |

| | | | |
|---|---|---|---|
| GIT1_HUMAN\|Q9Y2X7_S385_S388_T392 | 0.672380828 | 5.12E-04 | 3.23E-03 |
| FARP1_HUMAN\|Q9Y4F1_T24 | 0.672376941 | 2.36E-03 | 1.84E-02 |
| SUZ12_HUMAN\|Q15022_S541_S546 | 0.672308401 | 5.52E-03 | 2.14E-02 |
| CHSP1_HUMAN\|Q9Y2V2_S41 | 0.672265705 | 1.32E-03 | 7.61E-03 |
| AHNK_HUMAN\|Q09666_S5763 | 0.672261825 | 2.86E-05 | 5.06E-04 |
| EGFR_HUMAN\|P00533_S1205 | 0.672187597 | 2.59E-03 | 2.91E-02 |
| ARAF_HUMAN\|P10398_S257_T253 | 0.672150989 | 1.20E-03 | 1.08E-02 |
| PRC2A_HUMAN\|P48634_S1219 | 0.672024931 | 2.47E-04 | 3.40E-02 |
| COHA1_HUMAN\|Q9UMD9_S45 | 0.671927213 | 7.86E-04 | 4.55E-03 |
| K2C5_HUMAN\|P13647_S82_T78 | 0.671923238 | 6.10E-04 | 1.11E-02 |
| CAMP3_HUMAN\|Q9P1Y5_S756 | 0.671921693 | 4.35E-04 | 2.83E-03 |
| BIG1_HUMAN\|Q9Y6D6_S397 | 0.671799083 | 3.75E-04 | 2.71E-03 |
| HNRL2_HUMAN\|Q1KMD3_S161_S168_T165 | 0.671762521 | 5.39E-04 | 3.37E-03 |
| DBNL_HUMAN\|Q9UJU6-2_S232 | 0.671716174 | 2.28E-03 | 1.52E-02 |
| BCLF1_HUMAN\|Q9NYF8_S397_T402 | 0.671707389 | 3.96E-03 | 2.75E-02 |
| MFAP1_HUMAN\|P55081_S116_S118 | 0.671704331 | 2.56E-04 | 1.88E-03 |
| AKAP2_HUMAN\|Q9Y2D5_T157 | 0.671554219 | 9.58E-04 | 2.82E-02 |
| YAP1_HUMAN\|P46937_S164 | 0.671551954 | 1.60E-03 | 3.92E-02 |
| SPD2A_HUMAN\|Q5TCZ1_S724 | 0.671485916 | 1.78E-04 | 4.33E-03 |
| MISP_HUMAN\|Q8IVT2_S155 | 0.671417069 | 4.75E-03 | 4.54E-02 |
| SPD2A_HUMAN\|Q5TCZ1_S1002 | 0.671394366 | 1.59E-04 | 1.29E-03 |
| DPYL3_HUMAN\|Q14195-2_S102 | 0.671393433 | 6.44E-04 | 1.57E-02 |
| EXOC2_HUMAN\|Q96KP1_S432 | 0.671389291 | 1.09E-04 | 1.81E-03 |
| MTAP2_HUMAN\|P11137_S1782_S1786_S1790 | 0.671379415 | 4.39E-04 | 5.11E-03 |
| RM09_HUMAN\|Q9BYD2_S264 | 0.67135463 | 9.12E-04 | 1.20E-02 |
| HPBP1_HUMAN\|Q9NZL4_S354 | 0.671346137 | 8.90E-04 | 1.11E-02 |
| FA83B_HUMAN\|Q5T0W9_S388 | 0.671182579 | 2.31E-03 | 3.78E-02 |
| CAMP3_HUMAN\|Q9P1Y5_S553 | 0.671169362 | 1.22E-05 | 2.70E-04 |
| TRM1_HUMAN\|Q9NXH9_S625 | 0.671162836 | 1.61E-03 | 1.40E-02 |
| NEK9_HUMAN\|Q8TD19_T333 | 0.671104497 | 6.26E-03 | 2.75E-02 |
| DC1L2_HUMAN\|O43237_S407 | 0.671062666 | 4.46E-04 | 5.12E-03 |

| | | | |
|---|---|---|---|
| ARFG1_HUMAN\|Q8N6T3_T135 | 0.671012956 | 3.07E-04 | 3.87E-03 |
| AFAD_HUMAN\|P55196_T1232 | 0.670976662 | 2.15E-03 | 2.70E-02 |
| PHF14_HUMAN\|O94880_S290_T287 | 0.670963993 | 1.99E-03 | 1.07E-02 |
| RB6I2_HUMAN\|Q8IUD2_S17_S21 | 0.670951766 | 4.21E-04 | 8.57E-03 |
| BNIP2_HUMAN\|Q12982_S114 | 0.670873495 | 4.03E-03 | 2.29E-02 |
| PDLI2_HUMAN\|Q96JY6_S134_T146 | 0.670699787 | 8.31E-04 | 1.39E-02 |
| DDX10_HUMAN\|Q13206_S831 | 0.670686279 | 5.83E-04 | 2.08E-02 |
| K1C14_HUMAN\|P02533_S53 | 0.670651984 | 2.75E-04 | 2.10E-03 |
| T184B_HUMAN\|Q9Y519_S403 | 0.670640867 | 1.44E-03 | 1.23E-02 |
| K2C5_HUMAN\|P13647_S574 | 0.670557229 | 4.81E-04 | 5.45E-03 |
| SF3B1_HUMAN\|O75533_S194 | 0.670476269 | 1.09E-03 | 5.92E-03 |
| SPD2A_HUMAN\|Q5TCZ1_S642 | 0.670439674 | 1.55E-03 | 8.67E-03 |
| FA76A_HUMAN\|Q8TAV0_S306 | 0.670428346 | 2.44E-04 | 3.40E-02 |
| PKN2_HUMAN\|Q16513_S360 | 0.670418977 | 2.12E-03 | 2.67E-02 |
| PDLI2_HUMAN\|Q96JY6_S121_S129 | 0.670408599 | 1.40E-03 | 2.67E-02 |
| K1671_HUMAN\|Q9BY89_S1441 | 0.670320063 | 3.56E-03 | 4.87E-02 |
| FA83H_HUMAN\|Q6ZRV2_S647 | 0.670284943 | 3.05E-05 | 3.53E-04 |
| PTN14_HUMAN\|Q15678_S593_S594 | 0.670213407 | 6.52E-05 | 2.51E-03 |
| OSB10_HUMAN\|Q9BXB5_S32 | 0.670204816 | 6.62E-04 | 6.85E-03 |
| CARL1_HUMAN\|Q5VZK9_S1290_T1278 | 0.670069793 | 3.47E-04 | 3.54E-03 |
| ARHG5_HUMAN\|Q12774_S445_S450 | 0.670033373 | 3.83E-05 | 6.27E-04 |
| GGA2_HUMAN\|Q9UJY4_S177 | 0.670014578 | 2.26E-03 | 1.51E-02 |
| MAP7_HUMAN\|Q14244_S315 | 0.670012299 | 3.91E-04 | 2.80E-03 |
| BI2L1_HUMAN\|Q9UHR4_S414 | 0.670006377 | 1.86E-03 | 2.27E-02 |
| FXR2_HUMAN\|P51116_S453 | 0.669968124 | 7.71E-04 | 4.97E-03 |
| NSD3_HUMAN\|Q9BZ95_S457 | 0.66981112 | 3.64E-04 | 7.31E-03 |
| NUP50_HUMAN\|Q9UKX7_S221 | 0.669795211 | 1.54E-04 | 1.26E-03 |
| IF4G1_HUMAN\|Q04637_S1187 | 0.669774582 | 3.34E-04 | 2.31E-03 |
| OSTF1_HUMAN\|Q92882_S213 | 0.669725339 | 1.04E-03 | 1.25E-02 |
| KI26B_HUMAN\|Q2KJY2_S1006_S1018_S1021 | 0.669655328 | 5.48E-05 | 2.07E-03 |
| RTKN_HUMAN\|Q9BST9_S543 | 0.669653076 | 2.26E-04 | 3.06E-03 |

| | | | |
|---|---|---|---|
| OSTF1_HUMAN\|Q92882_S213 | 0.669587666 | 1.04E-03 | 1.01E-02 |
| EPIPL_HUMAN\|P58107_S2716 | 0.669279491 | 2.09E-05 | 4.00E-04 |
| EPN1_HUMAN\|Q9Y6I3_S454 | 0.669246714 | 1.27E-04 | 7.46E-03 |
| K1C17_HUMAN\|Q04695_S11 | 0.669145613 | 9.73E-04 | 9.62E-03 |
| PTN13_HUMAN\|Q12923_S908 | 0.669072572 | 1.17E-03 | 2.34E-02 |
| SOX15_HUMAN\|O60248_S37 | 0.669005764 | 4.02E-05 | 8.45E-04 |
| LASP1_HUMAN\|Q14847_T104 | 0.668951233 | 2.55E-04 | 5.96E-03 |
| S38A2_HUMAN\|Q96QD8_S17_S21 | 0.668950437 | 5.51E-03 | 3.48E-02 |
| FOXO3_HUMAN\|O43524_S253 | 0.66891395 | 3.80E-05 | 1.60E-03 |
| VIME_HUMAN\|P08670_S56 | 0.668878754 | 1.94E-04 | 9.92E-03 |
| K1671_HUMAN\|Q9BY89_S404 | 0.668733762 | 6.53E-05 | 1.22E-03 |
| TB182_HUMAN\|Q9C0C2_S1711 | 0.668695211 | 3.15E-04 | 3.94E-03 |
| TR150_HUMAN\|Q9Y2W1_T874 | 0.668655436 | 1.74E-03 | 2.34E-02 |
| CTNA1_HUMAN\|P35221_S641_T645 | 0.668654348 | 1.16E-03 | 6.21E-03 |
| DOCK6_HUMAN\|Q96HP0_S1343 | 0.668646236 | 1.83E-03 | 1.45E-02 |
| BZW2_HUMAN\|Q9Y6E2_S412 | 0.668569178 | 6.17E-03 | 4.62E-02 |
| DC1I2_HUMAN\|Q13409-2_S84 | 0.668553389 | 5.86E-04 | 1.03E-02 |
| CO052_HUMAN\|Q6ZUT6_S193 | 0.668544821 | 9.98E-04 | 1.55E-02 |
| LMNB2_HUMAN\|Q03252_S420 | 0.66850086 | 3.63E-04 | 2.63E-03 |
| UBE4B_HUMAN\|O95155_S84_S88 | 0.668373489 | 7.86E-03 | 4.49E-02 |
| ZFP91_HUMAN\|Q96JP5_S83 | 0.668370897 | 6.73E-05 | 6.56E-04 |
| IASPP_HUMAN\|Q8WUF5_S113 | 0.66833619 | 2.69E-04 | 1.95E-03 |
| AR6P4_HUMAN\|Q66PJ3_S241 | 0.668335412 | 2.89E-03 | 1.29E-02 |
| Q659E9_HUMAN\|Q659E9_S168 | 0.668319378 | 5.88E-05 | 5.50E-03 |
| CAMP2_HUMAN\|Q08AD1_S1319 | 0.668295269 | 1.92E-04 | 2.74E-03 |
| CCDC9_HUMAN\|Q9Y3X0_S376_S386_T381 | 0.668167342 | 4.98E-04 | 9.21E-03 |
| IWS1_HUMAN\|Q96ST2_T725 | 0.668152125 | 3.60E-03 | 3.70E-02 |
| IF4B_HUMAN\|P23588_S489_S497_S498 | 0.668142545 | 2.72E-04 | 2.94E-03 |
| PKP3_HUMAN\|Q9Y446_S123 | 0.668108801 | 2.31E-03 | 1.81E-02 |
| AHNK_HUMAN\|Q09666_S210_S216_S220 | 0.668096029 | 2.17E-04 | 1.77E-03 |
| CARL1_HUMAN\|Q5VZK9_S1290_T1278 | 0.668075082 | 3.21E-04 | 6.71E-03 |

| | | | |
|---|---|---|---|
| SP100_HUMAN|P23497_S476 | 0.668036934 | 1.94E-03 | 2.27E-02 |
| K2C75_HUMAN|O95678_S44 | 0.668020946 | 2.35E-03 | 1.09E-02 |
| ACAP2_HUMAN|Q15057_S384 | 0.667971904 | 3.60E-05 | 3.49E-03 |
| EPN2_HUMAN|O95208_S172 | 0.667894905 | 5.13E-03 | 2.03E-02 |
| SQSTM_HUMAN|Q13501_T269 | 0.667871602 | 4.65E-04 | 5.34E-03 |
| GPTC2_HUMAN|Q9NW75_S115_S117 | 0.667816861 | 5.30E-05 | 1.02E-03 |
| K1C14_HUMAN|P02533_S11 | 0.667760305 | 2.26E-04 | 3.12E-03 |
| FA83H_HUMAN|Q6ZRV2_S914 | 0.667757147 | 5.28E-04 | 3.31E-03 |
| BD1L1_HUMAN|Q8NFC6_S659_T658 | 0.667734983 | 2.32E-04 | 5.24E-03 |
| ZCH18_HUMAN|Q86VM9_T796 | 0.667707902 | 3.50E-03 | 3.69E-02 |
| PDLI2_HUMAN|Q96JY6_S117 | 0.667637562 | 1.72E-03 | 1.83E-02 |
| LARP6_HUMAN|Q9BRS8_S56_S58 | 0.667629647 | 6.68E-04 | 1.19E-02 |
| ACD_HUMAN|Q96AP0_S425 | 0.66761544 | 2.12E-04 | 5.19E-03 |
| AKAP2_HUMAN|Q9Y2D5_S152_T157 | 0.667464588 | 3.98E-04 | 2.83E-03 |
| SF3B1_HUMAN|O75533_T235 | 0.667443343 | 7.85E-03 | 4.49E-02 |
| ZCHC8_HUMAN|Q6NZY4_S500_T492_T496 | 0.667435953 | 1.31E-03 | 1.81E-02 |
| YAP1_HUMAN|P46937_S164 | 0.667408995 | 1.40E-03 | 1.89E-02 |
| S38A2_HUMAN|Q96QD8_S17_S22 | 0.667407471 | 6.85E-03 | 2.53E-02 |
| UFD1_HUMAN|Q92890_S299 | 0.667353466 | 9.53E-03 | 3.87E-02 |
| MFAP1_HUMAN|P55081_T267 | 0.667246268 | 2.27E-03 | 1.06E-02 |
| AHNK_HUMAN|Q09666_S511 | 0.667244586 | 2.18E-03 | 2.48E-02 |
| ABI2_HUMAN|Q9NYB9_S368 | 0.667230913 | 2.86E-05 | 6.43E-04 |
| RHG23_HUMAN|Q9P227_S351 | 0.667217518 | 5.16E-04 | 3.53E-03 |
| DDI2_HUMAN|Q5TDH0_S194 | 0.667206778 | 1.16E-03 | 1.61E-02 |
| F122B_HUMAN|Q7Z309_S115_S119 | 0.667194569 | 1.36E-04 | 3.88E-03 |
| XPO4_HUMAN|Q9C0E2_S521 | 0.667170421 | 5.07E-04 | 4.78E-03 |
| RB3GP_HUMAN|Q15042_S537 | 0.667107081 | 7.87E-05 | 2.49E-03 |
| PLEC_HUMAN|Q15149_S4386 | 0.667093256 | 2.37E-04 | 1.79E-03 |
| ITB4_HUMAN|P16144_S1457 | 0.66706868 | 1.96E-03 | 1.54E-02 |
| SKT_HUMAN|Q5T5P2_S1794 | 0.666926587 | 6.04E-04 | 6.52E-03 |
| AHNK_HUMAN|Q09666_T4430 | 0.666903898 | 1.20E-03 | 1.08E-02 |

| | | | |
|---|---|---|---|
| CSKI2_HUMAN\|Q8WXE0_S406 | 0.666903822 | 2.15E-05 | 1.02E-03 |
| KLC4_HUMAN\|Q9NSK0_T612 | 0.666836699 | 1.11E-02 | 3.76E-02 |
| HAUS6_HUMAN\|Q7Z4H7_T584 | 0.666827101 | 2.66E-03 | 1.68E-02 |
| MA7D1_HUMAN\|Q3KQU3_S366 | 0.66681971 | 2.96E-05 | 1.27E-03 |
| PRAG1_HUMAN\|Q86YV5_S907 | 0.666817363 | 2.39E-04 | 5.31E-03 |
| MILK1_HUMAN\|Q8N3F8_S471 | 0.666644135 | 3.91E-05 | 8.21E-04 |
| PTN14_HUMAN\|Q15678_S593_S594 | 0.666637601 | 5.56E-05 | 5.57E-04 |
| MAST4_HUMAN\|O15021_S2555 | 0.66660586 | 1.57E-03 | 2.18E-02 |
| MILK1_HUMAN\|Q8N3F8_S391 | 0.666583702 | 7.37E-06 | 4.50E-04 |
| MET_HUMAN\|P08581_S997 | 0.666532603 | 5.64E-03 | 2.91E-02 |
| NU107_HUMAN\|P57740_S37 | 0.666385034 | 4.46E-04 | 1.78E-02 |
| AFAD_HUMAN\|P55196_T1330 | 0.666239206 | 5.99E-04 | 2.15E-02 |
| EXPH5_HUMAN\|Q8NEV8_S341 | 0.666063709 | 4.50E-04 | 1.23E-02 |
| CHSP1_HUMAN\|Q9Y2V2_S41 | 0.666049278 | 1.06E-03 | 1.02E-02 |
| ACINU_HUMAN\|Q9UKV3_S240_S243 | 0.666008932 | 1.22E-04 | 1.55E-03 |
| DBNL_HUMAN\|Q9UJU6-2_S232 | 0.665982545 | 1.90E-03 | 1.49E-02 |
| KKCC2_HUMAN\|Q96RR4-3_S522 | 0.665961807 | 3.16E-03 | 4.59E-02 |
| FCHO2_HUMAN\|Q0JRZ9_S387 | 0.665898962 | 1.08E-02 | 4.80E-02 |
| BIG1_HUMAN\|Q9Y6D6_S397 | 0.665868508 | 2.94E-04 | 3.78E-03 |
| SVIL_HUMAN\|O95425-2_S263_S270 | 0.665867612 | 8.73E-04 | 7.15E-03 |
| CF132_HUMAN\|Q5T0Z8_T297 | 0.665861687 | 5.34E-03 | 4.94E-02 |
| LARP6_HUMAN\|Q9BRS8_S56_S58 | 0.66583655 | 6.23E-04 | 6.57E-03 |
| LARP1_HUMAN\|Q6PKG0_S627_Y633 | 0.665707375 | 4.59E-03 | 1.88E-02 |
| TMF1_HUMAN\|P82094_S72 | 0.665629 | 1.51E-04 | 1.84E-03 |
| TWF1_HUMAN\|Q12792_T348 | 0.66561813 | 1.74E-05 | 2.27E-04 |
| EF1D_HUMAN\|P29692_S119 | 0.665604768 | 6.10E-04 | 3.69E-03 |
| MKL2_HUMAN\|Q9ULH7_S921 | 0.665600292 | 6.43E-04 | 1.11E-02 |
| YAP1_HUMAN\|P46937_S109_T119 | 0.665384114 | 2.64E-05 | 1.31E-03 |
| TP53B_HUMAN\|Q12888_S366 | 0.665349361 | 4.82E-05 | 2.19E-03 |
| LIPA1_HUMAN\|Q13136_S666_S668 | 0.665328355 | 2.36E-04 | 3.20E-03 |
| K1522_HUMAN\|Q9P206_S327 | 0.665288365 | 5.64E-05 | 1.05E-03 |

| | | | |
|---|---|---|---|
| VIME_HUMAN\|P08670_S459 | 0.665256059 | 1.66E-04 | 1.45E-03 |
| EPHA2_HUMAN\|P29317_S897_S901 | 0.665255941 | 3.02E-03 | 4.45E-02 |
| ADNP_HUMAN\|Q9H2P0_S921 | 0.665099192 | 1.76E-04 | 1.39E-03 |
| EPN2_HUMAN\|O95208_S172 | 0.665089086 | 4.76E-03 | 2.19E-02 |
| ZC3HD_HUMAN\|Q5T200_S1014_S1017 | 0.665063469 | 1.37E-03 | 7.84E-03 |
| DLG1_HUMAN\|Q12959_S573 | 0.665003149 | 9.61E-03 | 3.35E-02 |
| SP100_HUMAN\|P23497_S180 | 0.664997728 | 1.32E-03 | 3.34E-02 |
| SENP3_HUMAN\|Q9H4L4_S169_S181_T176 | 0.664976549 | 9.79E-03 | 3.39E-02 |
| TM1L1_HUMAN\|O75674_S321_S323 | 0.664933216 | 1.75E-03 | 1.48E-02 |
| AT11C_HUMAN\|Q8NB49_T1124 | 0.664931693 | 4.57E-04 | 4.38E-03 |
| K2C5_HUMAN\|P13647_S571 | 0.6649191 | 1.72E-04 | 2.47E-03 |
| FA83B_HUMAN\|Q5T0W9_S664 | 0.664853858 | 1.27E-04 | 1.10E-03 |
| JUNB_HUMAN\|P17275_T255 | 0.664817794 | 2.48E-05 | 1.27E-03 |
| NKAP_HUMAN\|Q8N5F7_S149 | 0.664804454 | 1.33E-04 | 1.13E-03 |
| NFIC_HUMAN\|P08651_S294_S304 | 0.664765871 | 1.51E-03 | 2.11E-02 |
| TB182_HUMAN\|Q9C0C2_S1029 | 0.664697624 | 1.74E-04 | 6.13E-03 |
| C2CD5_HUMAN\|Q86YS7-2_S855 | 0.664666228 | 5.76E-04 | 3.52E-03 |
| PLEC_HUMAN\|Q15149-2_S24 | 0.664653867 | 2.56E-03 | 2.76E-02 |
| E7ET48_HUMAN\|E7ET48_S441 | 0.664519726 | 1.10E-03 | 8.57E-03 |
| SAFB1_HUMAN\|Q15424_S604 | 0.664482009 | 1.54E-05 | 4.23E-04 |
| AHNK2_HUMAN\|Q8IVF2_S1712 | 0.664416443 | 4.47E-04 | 1.69E-02 |
| CT2NL_HUMAN\|Q9P2B4_S568 | 0.66416757 | 2.77E-04 | 6.35E-03 |
| FA83B_HUMAN\|Q5T0W9_S664 | 0.664126667 | 1.24E-04 | 2.39E-02 |
| IRS1_HUMAN\|P35568_S1101 | 0.664089155 | 1.37E-04 | 1.15E-03 |
| ITB4_HUMAN\|P16144_S1515_S1517 | 0.664066011 | 4.70E-03 | 2.57E-02 |
| K2C5_HUMAN\|P13647_S16 | 0.664009494 | 2.46E-03 | 1.89E-02 |
| REPS1_HUMAN\|Q96D71_S162_S166_T173 | 0.663985686 | 9.19E-05 | 2.68E-03 |
| DC1L1_HUMAN\|Q9Y6G9_S516 | 0.663955256 | 1.57E-04 | 1.90E-03 |
| K1C16_HUMAN\|P08779_S468 | 0.663944067 | 5.26E-04 | 5.82E-03 |
| TGS1_HUMAN\|Q96RS0_S154 | 0.663851301 | 7.84E-04 | 5.03E-03 |
| LSR_HUMAN\|Q86X29_T505 | 0.663819782 | 9.32E-03 | 4.87E-02 |

| | | | |
|---|---|---|---|
| CIRBP_HUMAN\|Q14011_S148 | 0.663789899 | 1.96E-04 | 1.53E-03 |
| SRG2B_HUMAN\|P0DMP2_S206 | 0.663735557 | 8.41E-04 | 1.06E-02 |
| SQSTM_HUMAN\|Q13501_S266 | 0.663694037 | 1.08E-04 | 1.80E-03 |
| MA7D1_HUMAN\|Q3KQU3_S313 | 0.663693344 | 3.79E-03 | 3.85E-02 |
| AHNK_HUMAN\|Q09666_S5763 | 0.663678102 | 1.82E-05 | 4.79E-04 |
| NELFE_HUMAN\|P18615_S353 | 0.663654311 | 1.04E-05 | 3.19E-04 |
| NHRF1_HUMAN\|O14745_S280 | 0.663634791 | 2.54E-04 | 8.15E-03 |
| FARP1_HUMAN\|Q9Y4F1_S427 | 0.663631723 | 3.18E-04 | 3.96E-03 |
| NCOA7_HUMAN\|Q8NI08_S384 | 0.663606646 | 8.63E-05 | 2.84E-03 |
| XPC_HUMAN\|Q01831_S883_S884_S891 | 0.663602195 | 4.20E-03 | 4.36E-02 |
| ZC3HD_HUMAN\|Q5T200_S64 | 0.663572936 | 8.47E-04 | 1.40E-02 |
| CNKR1_HUMAN\|Q969H4_S305_S314 | 0.663527883 | 1.72E-03 | 8.51E-03 |
| BASP1_HUMAN\|P80723_T196 | 0.663409574 | 1.05E-03 | 1.59E-02 |
| K1C14_HUMAN\|P02533_S51 | 0.663338097 | 4.59E-03 | 1.88E-02 |
| TB182_HUMAN\|Q9C0C2_S920 | 0.663325932 | 1.03E-03 | 2.86E-02 |
| RB6I2_HUMAN\|Q8IUD2_S17_S21 | 0.663291702 | 3.08E-04 | 2.18E-03 |
| PININ_HUMAN\|Q9H307_S690 | 0.663259385 | 3.44E-03 | 2.05E-02 |
| UBP8_HUMAN\|P40818_T577 | 0.663236762 | 4.22E-05 | 4.63E-04 |
| RTKN_HUMAN\|Q9BST9_S529 | 0.66321677 | 8.59E-04 | 8.54E-03 |
| PKP3_HUMAN\|Q9Y446_S313 | 0.663188413 | 1.21E-05 | 1.73E-04 |
| TB182_HUMAN\|Q9C0C2_T875 | 0.66316833 | 2.40E-04 | 3.40E-02 |
| RIOK1_HUMAN\|Q9BRS2_S21_S22 | 0.66309723 | 2.46E-03 | 1.81E-02 |
| ATRX_HUMAN\|P46100_S784 | 0.662978937 | 4.21E-03 | 3.50E-02 |
| PKP3_HUMAN\|Q9Y446_S331 | 0.662919913 | 9.80E-04 | 5.42E-03 |
| TBC9B_HUMAN\|Q66K14_S1241 | 0.662919885 | 2.75E-05 | 4.90E-04 |
| TB10B_HUMAN\|Q4KMP7_S656 | 0.66291545 | 2.22E-03 | 1.05E-02 |
| HDAC4_HUMAN\|P56524_S266 | 0.662835083 | 2.11E-04 | 4.86E-03 |
| ITSN1_HUMAN\|Q15811_S315 | 0.662721283 | 1.73E-04 | 2.48E-03 |
| UBP2L_HUMAN\|Q14157_S470_S477 | 0.662623307 | 2.55E-03 | 3.01E-02 |
| MICA1_HUMAN\|Q8TDZ2_S960 | 0.662534639 | 8.15E-05 | 1.12E-03 |
| PLXB2_HUMAN\|O15031_S1570 | 0.662464297 | 2.07E-03 | 1.61E-02 |

| | | | |
|---|---|---|---|
| PKP2_HUMAN\|Q99959_S197 | 0.662335425 | 3.01E-04 | 2.14E-03 |
| UT14A_HUMAN\|Q9BVJ6_S437 | 0.66230555 | 1.74E-03 | 4.04E-02 |
| EDC3_HUMAN\|Q96F86_S131 | 0.662290777 | 2.47E-05 | 1.14E-03 |
| FA83H_HUMAN\|Q6ZRV2_Y875 | 0.662263752 | 2.32E-05 | 2.81E-04 |
| SRRM1_HUMAN\|Q8IYB3_T846 | 0.662254354 | 4.18E-03 | 2.87E-02 |
| LAR4B_HUMAN\|Q92615_S731_S736 | 0.662241287 | 8.09E-04 | 4.65E-03 |
| LMNA_HUMAN\|P02545_S619 | 0.662201378 | 2.19E-03 | 1.75E-02 |
| SP100_HUMAN\|P23497_S180 | 0.66219908 | 1.20E-03 | 9.21E-03 |
| WDR13_HUMAN\|Q9H1Z4_S70_S74 | 0.662184936 | 4.50E-04 | 1.69E-02 |
| CCD86_HUMAN\|Q9H6F5_S217 | 0.662181959 | 2.42E-03 | 3.91E-02 |
| AFAD_HUMAN\|P55196_S1799 | 0.662131616 | 1.59E-03 | 1.32E-02 |
| ACAP2_HUMAN\|Q15057_S381_S384 | 0.662112596 | 3.81E-04 | 7.92E-03 |
| SARG_HUMAN\|Q9BW04_S429 | 0.66207942 | 3.11E-03 | 1.55E-02 |
| MA7D1_HUMAN\|Q3KQU3_S446 | 0.661955847 | 9.13E-05 | 2.68E-03 |
| LAP2A_HUMAN\|P42166_T74 | 0.661924451 | 2.94E-03 | 2.15E-02 |
| KKCC2_HUMAN\|Q96RR4-3_S522 | 0.66187798 | 2.80E-03 | 2.93E-02 |
| OSTF1_HUMAN\|Q92882_S213 | 0.66179697 | 7.95E-04 | 1.09E-02 |
| PHF20_HUMAN\|Q9BVI0_S878_S880 | 0.661641225 | 1.89E-04 | 4.51E-03 |
| AFF4_HUMAN\|Q9UHB7_S836 | 0.661626084 | 1.02E-03 | 8.12E-03 |
| AHNK_HUMAN\|Q09666_S5763 | 0.66161667 | 1.63E-05 | 8.42E-04 |
| RHG23_HUMAN\|Q9P227_S361_S372 | 0.661615524 | 1.98E-05 | 9.60E-04 |
| FA83H_HUMAN\|Q6ZRV2_S936 | 0.661586971 | 1.40E-03 | 7.98E-03 |
| SFPQ_HUMAN\|P23246_T687 | 0.661546294 | 2.47E-03 | 1.14E-02 |
| IRS2_HUMAN\|Q9Y4H2_S1186_S1203 | 0.661419595 | 3.62E-04 | 1.53E-02 |
| RUFY3_HUMAN\|Q7L099-3_S618 | 0.661242627 | 1.53E-05 | 4.23E-04 |
| RHG23_HUMAN\|Q9P227_S361_S372 | 0.661148266 | 1.93E-05 | 4.96E-04 |
| SCEL_HUMAN\|O95171_S91 | 0.661092205 | 6.84E-03 | 4.06E-02 |
| GRB10_HUMAN\|Q13322_S104 | 0.661050662 | 3.86E-05 | 1.60E-03 |
| PKP4_HUMAN\|Q99569_S236 | 0.660975758 | 3.80E-04 | 2.56E-03 |
| DC1L1_HUMAN\|Q9Y6G9_S421 | 0.660935437 | 1.04E-02 | 4.18E-02 |
| SMAG1_HUMAN\|Q9UPU9_S420_T424 | 0.660917656 | 5.35E-05 | 1.99E-03 |

| | | | |
|---|---|---|---|
| ZYX_HUMAN\|Q15942_S259 | 0.660894658 | 1.69E-04 | 2.45E-03 |
| TISB_HUMAN\|Q07352_S54 | 0.660884235 | 3.49E-05 | 3.92E-04 |
| AAK1_HUMAN\|Q2M2I8_T389 | 0.660831224 | 1.68E-03 | 9.26E-03 |
| AF1L2_HUMAN\|Q8N4X5_T284 | 0.660821339 | 8.58E-04 | 1.89E-02 |
| ZCH18_HUMAN\|Q86VM9_S532_S534 | 0.660721227 | 7.90E-06 | 1.24E-04 |
| IASPP_HUMAN\|Q8WUF5_S74 | 0.660698192 | 2.58E-05 | 3.11E-04 |
| CF132_HUMAN\|Q5T0Z8_T297 | 0.660696695 | 4.66E-03 | 4.40E-02 |
| TP53B_HUMAN\|Q12888_S366 | 0.660529411 | 3.85E-05 | 6.28E-04 |
| SMTN_HUMAN\|P53814_T791 | 0.660479092 | 7.63E-03 | 2.77E-02 |
| ELAV1_HUMAN\|Q15717_S202 | 0.660446825 | 3.71E-03 | 3.78E-02 |
| EDC3_HUMAN\|Q96F86_S131 | 0.660405133 | 2.25E-05 | 2.79E-04 |
| TRI29_HUMAN\|Q14134_S152 | 0.660391714 | 5.38E-04 | 1.41E-02 |
| EGFR_HUMAN\|P00533_S1205 | 0.660305183 | 1.76E-03 | 9.62E-03 |
| LIPA3_HUMAN\|O75145_S1164 | 0.660289836 | 2.60E-04 | 4.45E-03 |
| PML_HUMAN\|P29590_T867 | 0.66025679 | 4.02E-04 | 8.37E-03 |
| DDI2_HUMAN\|Q5TDH0_S106 | 0.660206003 | 9.55E-05 | 2.82E-03 |
| FSTL3_HUMAN\|O95633_S255 | 0.66019306 | 1.47E-03 | 1.62E-02 |
| SGTA_HUMAN\|O43765_S301_S305 | 0.660178519 | 9.67E-03 | 4.42E-02 |
| PPR18_HUMAN\|Q6NYC8_S209_S213 | 0.660157042 | 1.34E-04 | 2.01E-03 |
| YETS2_HUMAN\|Q9ULM3_S536 | 0.660020924 | 2.47E-04 | 5.91E-03 |
| TB182_HUMAN\|Q9C0C2_S1711 | 0.660001261 | 2.20E-04 | 5.01E-03 |
| ARP19_HUMAN\|P56211_S62 | 0.659963772 | 3.27E-04 | 4.04E-03 |
| BORG4_HUMAN\|Q9H3Q1_S292 | 0.659931361 | 2.01E-03 | 1.64E-02 |
| TGS1_HUMAN\|Q96RS0_S55_T60 | 0.659905943 | 1.94E-05 | 4.96E-04 |
| SPD2A_HUMAN\|Q5TCZ1_S642 | 0.659904068 | 1.10E-03 | 5.95E-03 |
| ACINU_HUMAN\|Q9UKV3_T254 | 0.659893125 | 4.22E-03 | 4.21E-02 |
| NIPA_HUMAN\|Q86WB0_S321 | 0.659754332 | 2.34E-05 | 4.40E-04 |
| PLXA2_HUMAN\|O75051_S1633 | 0.659746405 | 1.19E-03 | 1.12E-02 |
| ZN638_HUMAN\|Q14966_S1401 | 0.659734597 | 1.22E-03 | 1.09E-02 |
| ITB4_HUMAN\|P16144_S1515_S1517 | 0.659579052 | 4.15E-03 | 3.56E-02 |
| PKP3_HUMAN\|Q9Y446_S238 | 0.659575049 | 2.46E-05 | 1.27E-03 |

| | | | |
|---|---|---|---|
| PAXI_HUMAN\|P49023_S106 | 0.659559941 | 8.44E-04 | 1.88E-02 |
| MISP_HUMAN\|Q8IVT2_S471 | 0.659425314 | 9.57E-04 | 5.31E-03 |
| PHLB2_HUMAN\|Q86SQ0_S330_S334 | 0.659409208 | 5.15E-03 | 3.14E-02 |
| PHLB2_HUMAN\|Q86SQ0_S402 | 0.659407523 | 3.60E-04 | 2.46E-03 |
| PRAG1_HUMAN\|Q86YV5_S712 | 0.65939857 | 5.90E-05 | 8.77E-04 |
| UBP24_HUMAN\|Q9UPU5_S1143 | 0.659352834 | 3.02E-04 | 3.83E-03 |
| TWF1_HUMAN\|Q12792_T348 | 0.659338301 | 1.26E-05 | 1.78E-04 |
| K2C8_HUMAN\|P05787_S24 | 0.659184385 | 2.72E-03 | 3.16E-02 |
| CCDC6_HUMAN\|Q16204_S244 | 0.659166625 | 1.57E-03 | 1.13E-02 |
| UBP2L_HUMAN\|Q14157_S467_S477 | 0.659124101 | 2.57E-04 | 5.96E-03 |
| K2C5_HUMAN\|P13647_S21_T24 | 0.659020441 | 3.30E-03 | 1.62E-02 |
| NUMA1_HUMAN\|Q14980_T2000 | 0.659014947 | 1.09E-04 | 4.15E-03 |
| SPD2A_HUMAN\|Q5TCZ1_S724 | 0.659005281 | 1.05E-04 | 9.47E-04 |
| DBNL_HUMAN\|Q9UJU6_T270 | 0.658982952 | 2.62E-03 | 1.98E-02 |
| AHNK_HUMAN\|Q09666_T5729 | 0.658962981 | 8.38E-05 | 1.49E-03 |
| SYNEM_HUMAN\|O15061_S1044_S1049 | 0.658960676 | 4.26E-04 | 4.97E-03 |
| PLXA2_HUMAN\|O75051_S1630_S1633 | 0.658906769 | 1.32E-03 | 1.16E-02 |
| PLEC_HUMAN\|Q15149_S4382_S4396 | 0.658850886 | 4.08E-04 | 2.89E-03 |
| PATL1_HUMAN\|Q86TB9_S179_S184 | 0.658843304 | 2.38E-03 | 1.85E-02 |
| MARCS_HUMAN\|P29966_T150 | 0.658840302 | 1.77E-03 | 2.20E-02 |
| SWAHC_HUMAN\|Q53LP3_S212 | 0.658776371 | 4.09E-05 | 1.53E-03 |
| K1C14_HUMAN\|P02533_S432 | 0.658766951 | 3.29E-03 | 1.62E-02 |
| AKTS1_HUMAN\|Q96B36_T246 | 0.65865969 | 2.51E-03 | 3.00E-02 |
| PHF14_HUMAN\|O94880_S290_T287 | 0.658606314 | 1.35E-03 | 1.17E-02 |
| STAT1_HUMAN\|P42224_S727 | 0.658590835 | 2.49E-04 | 1.27E-02 |
| AKAP2_HUMAN\|Q9Y2D5_S152_S155 | 0.658570123 | 2.54E-04 | 1.98E-03 |
| DKC1_HUMAN\|O60832_S485_S494 | 0.658559324 | 5.52E-05 | 5.55E-04 |
| MPLKI_HUMAN\|Q8TAP9_T110 | 0.658549625 | 1.11E-03 | 5.98E-03 |
| PHLB2_HUMAN\|Q86SQ0_S334 | 0.65850394 | 9.54E-04 | 5.30E-03 |
| GPBP1_HUMAN\|Q86WP2_S381 | 0.658427175 | 8.48E-04 | 2.64E-02 |
| MA7D1_HUMAN\|Q3KQU3_S366 | 0.658418779 | 1.91E-05 | 2.45E-04 |

| | | | |
|---|---|---|---|
| PEAK1_HUMAN\|Q9H792_S1036 | 0.658315542 | 3.75E-05 | 1.72E-03 |
| TKT_HUMAN\|P29401_S295 | 0.658306189 | 3.91E-05 | 4.28E-04 |
| FRM4B_HUMAN\|Q9Y2L6_S628 | 0.658277076 | 8.21E-04 | 4.71E-03 |
| IF4G2_HUMAN\|P78344_S17 | 0.658209248 | 2.44E-06 | 5.20E-05 |
| AHNK_HUMAN\|Q09666_S5448 | 0.6581401 | 2.06E-04 | 2.84E-03 |
| SIK3_HUMAN\|Q9Y2K2_S493 | 0.658084483 | 4.17E-05 | 4.59E-04 |
| STRN3_HUMAN\|Q13033_S202 | 0.658058287 | 5.31E-05 | 5.39E-04 |
| PKHG3_HUMAN\|A1L390_S618 | 0.658039504 | 2.25E-03 | 1.06E-02 |
| RHG21_HUMAN\|Q5T5U3_S615 | 0.658002516 | 9.78E-05 | 8.99E-04 |
| NUP98_HUMAN\|P52948_S612 | 0.657932467 | 2.48E-04 | 4.31E-03 |
| GMIP_HUMAN\|Q9P107_S914_S919 | 0.657915149 | 1.61E-04 | 4.35E-03 |
| ROA3_HUMAN\|P51991_S355 | 0.657898816 | 3.27E-05 | 3.73E-04 |
| RB3GP_HUMAN\|Q15042_S537 | 0.657814827 | 5.24E-05 | 5.08E-03 |
| K2C5_HUMAN\|P13647_S18 | 0.657768424 | 5.72E-04 | 1.44E-02 |
| STX7_HUMAN\|O15400_S131 | 0.657744266 | 9.60E-05 | 7.17E-03 |
| AF1L2_HUMAN\|Q8N4X5_T284 | 0.657610962 | 7.68E-04 | 1.24E-02 |
| AFAD_HUMAN\|P55196_S1172_S1173_S1182 | 0.657587094 | 5.62E-05 | 1.05E-03 |
| AHNK2_HUMAN\|Q8IVF2_S294_S297 | 0.657580604 | 2.25E-05 | 2.75E-04 |
| KC1D_HUMAN\|P48730_S331 | 0.657533214 | 2.25E-03 | 2.59E-02 |
| LMNB1_HUMAN\|P20700_T19 | 0.657508457 | 8.20E-05 | 1.12E-03 |
| YBOX1_HUMAN\|P67809_S165_S176 | 0.657436276 | 3.06E-04 | 3.87E-03 |
| PKP3_HUMAN\|Q9Y446_S238 | 0.657431021 | 2.21E-05 | 1.06E-03 |
| ATN1_HUMAN\|P54259_S77_S79 | 0.657403985 | 2.84E-03 | 2.11E-02 |
| F118A_HUMAN\|Q9NWS6_S349 | 0.657393461 | 3.38E-05 | 5.83E-04 |
| CSKI2_HUMAN\|Q8WXE0_S403 | 0.657388372 | 4.89E-05 | 5.12E-04 |
| REPS1_HUMAN\|Q96D71_S162_S170_S174 | 0.657372851 | 2.58E-04 | 2.00E-03 |
| ITB4_HUMAN\|P16144_S1069 | 0.65735548 | 3.05E-03 | 2.15E-02 |
| OSB11_HUMAN\|Q9BXB4_S181 | 0.657334691 | 1.35E-04 | 3.88E-03 |
| LAD1_HUMAN\|O00515_S485 | 0.657329477 | 1.28E-03 | 6.71E-03 |
| LAP2A_HUMAN\|P42166_T74 | 0.657317095 | 2.58E-03 | 3.02E-02 |
| CTNB1_HUMAN\|P35222_S552 | 0.657217241 | 5.73E-04 | 3.50E-03 |

| | | | |
|---|---|---|---|
| F117A_HUMAN\|Q9C073_S67 | 0.657204363 | 6.99E-04 | 4.58E-03 |
| NBN_HUMAN\|O60934_S343 | 0.657186366 | 4.22E-04 | 4.94E-03 |
| IWS1_HUMAN\|Q96ST2_T725 | 0.656989049 | 2.62E-03 | 3.05E-02 |
| TB10B_HUMAN\|Q4KMP7_S656 | 0.656974906 | 1.86E-03 | 1.54E-02 |
| P66B_HUMAN\|Q8WXI9_S334 | 0.656900654 | 1.31E-05 | 1.83E-04 |
| LAR4B_HUMAN\|Q92615_T732 | 0.656880309 | 1.12E-05 | 6.37E-04 |
| AKA11_HUMAN\|Q9UKA4_S448 | 0.656826634 | 1.38E-03 | 1.81E-02 |
| K1C14_HUMAN\|P02533_S28 | 0.656822327 | 3.98E-04 | 2.83E-03 |
| SYTL1_HUMAN\|Q8IYJ3_S216 | 0.656771619 | 8.30E-03 | 4.52E-02 |
| UBE4B_HUMAN\|O95155_S84_S88 | 0.656767367 | 5.85E-03 | 3.45E-02 |
| P4K2A_HUMAN\|Q9BTU6_S462_S468 | 0.656642234 | 1.17E-04 | 8.10E-03 |
| LARP6_HUMAN\|Q9BRS8_S447_T450 | 0.656638436 | 1.71E-03 | 1.45E-02 |
| WDR13_HUMAN\|Q9H1Z4_S79 | 0.656624171 | 3.94E-04 | 7.67E-03 |
| ITB4_HUMAN\|P16144_S1069 | 0.656542704 | 2.98E-03 | 3.21E-02 |
| JUND_HUMAN\|P17535_S100 | 0.656481471 | 1.21E-03 | 9.32E-03 |
| LMNB2_HUMAN\|Q03252_S424 | 0.656447135 | 1.43E-03 | 2.03E-02 |
| CRBG1_HUMAN\|Q9Y4K1_S141 | 0.656418827 | 9.03E-04 | 1.43E-02 |
| ITA6_HUMAN\|P23229_S1128 | 0.656410122 | 1.84E-03 | 8.99E-03 |
| NFIC_HUMAN\|P08651_S333_S339 | 0.656334747 | 5.33E-04 | 3.64E-03 |
| ADNP_HUMAN\|Q9H2P0_S709 | 0.656314444 | 1.51E-05 | 8.00E-04 |
| ASPC1_HUMAN\|Q9BZE9_S275 | 0.656277852 | 2.21E-04 | 7.22E-03 |
| CTNA1_HUMAN\|P35221_T654_T658 | 0.656192535 | 1.36E-03 | 7.05E-03 |
| LARP4_HUMAN\|Q71RC2_S597 | 0.65615664 | 3.03E-04 | 3.85E-03 |
| NF1_HUMAN\|P21359_S2515 | 0.65611825 | 1.09E-04 | 1.72E-03 |
| PRP31_HUMAN\|Q8WWY3_S498 | 0.656071822 | 7.03E-06 | 1.13E-04 |
| K2C6B_HUMAN\|P04259_S546 | 0.656040031 | 2.63E-04 | 3.51E-03 |
| GRAP1_HUMAN\|Q4V328_S664 | 0.656036743 | 6.14E-04 | 1.51E-02 |
| COHA1_HUMAN\|Q9UMD9_S85_S93_S96 | 0.65599039 | 5.83E-04 | 3.55E-03 |
| BAD_HUMAN\|Q92934_Y76 | 0.655958777 | 7.08E-05 | 2.37E-03 |
| PLCH2_HUMAN\|O75038_S982 | 0.655924498 | 1.39E-05 | 1.89E-04 |
| RFIP5_HUMAN\|Q9BXF6_S296 | 0.655919097 | 1.18E-05 | 3.41E-04 |

| | | | |
|---|---|---|---|
| KLC2_HUMAN\|Q9H0B6_S610 | 0.655891256 | 1.76E-05 | 2.28E-04 |
| SHRM2_HUMAN\|Q13796_S1039 | 0.655754161 | 1.11E-06 | 6.25E-05 |
| BRAF_HUMAN\|P15056_S729 | 0.655708589 | 5.08E-03 | 2.72E-02 |
| ROA1_HUMAN\|P09651_S361_S364 | 0.655704878 | 3.46E-03 | 1.68E-02 |
| CRBG1_HUMAN\|Q9Y4K1_S75 | 0.655567365 | 2.14E-04 | 5.22E-03 |
| LAD1_HUMAN\|O00515_S485 | 0.655534774 | 1.20E-03 | 7.06E-03 |
| AFF4_HUMAN\|Q9UHB7_S1043 | 0.655480599 | 5.17E-05 | 5.34E-04 |
| M3K20_HUMAN\|Q9NYL2_S633 | 0.655458425 | 4.47E-03 | 2.08E-02 |
| RB6I2_HUMAN\|Q8IUD2_S17_S21 | 0.655384935 | 2.22E-04 | 5.04E-03 |
| URFB1_HUMAN\|Q6BDS2_S1106 | 0.655378009 | 1.49E-04 | 2.89E-03 |
| KPCD1_HUMAN\|Q15139_S742 | 0.655322332 | 4.42E-03 | 4.03E-02 |
| PKP2_HUMAN\|Q99959_S81 | 0.655234578 | 5.03E-03 | 2.29E-02 |
| NIBL1_HUMAN\|Q96TA1_S691_S696 | 0.655213659 | 2.88E-04 | 6.15E-03 |
| CAMP2_HUMAN\|Q08AD1_S1319 | 0.655189099 | 1.10E-04 | 9.82E-04 |
| BCAR3_HUMAN\|O75815_S375 | 0.655150291 | 1.22E-03 | 1.73E-02 |
| TRI25_HUMAN\|Q14258_S100 | 0.655123615 | 2.56E-03 | 1.94E-02 |
| PTK6_HUMAN\|Q13882_Y447 | 0.655018711 | 1.28E-04 | 3.47E-03 |
| COHA1_HUMAN\|Q9UMD9_S93_S96 | 0.654988654 | 4.79E-03 | 4.56E-02 |
| DESP_HUMAN\|P15924_S2815 | 0.654921175 | 1.03E-04 | 2.44E-03 |
| CLAP1_HUMAN\|Q7Z460_S1091 | 0.654899418 | 9.95E-05 | 2.84E-03 |
| ACAP2_HUMAN\|Q15057_S384 | 0.654895975 | 1.96E-05 | 9.55E-04 |
| MTMR2_HUMAN\|Q13614_S58 | 0.654851056 | 4.02E-03 | 2.65E-02 |
| AFF4_HUMAN\|Q9UHB7_S1043 | 0.654798255 | 5.01E-05 | 5.33E-04 |
| TB182_HUMAN\|Q9C0C2_S1666 | 0.654790413 | 1.28E-05 | 2.78E-04 |
| DGC14_HUMAN\|Q96DF8_T386 | 0.65477229 | 9.90E-05 | 3.19E-03 |
| PALLD_HUMAN\|Q8WX93_S726 | 0.654717189 | 3.34E-03 | 1.45E-02 |
| AKAP2_HUMAN\|Q9Y2D5_S720 | 0.654685162 | 1.16E-04 | 1.89E-03 |
| PDS5A_HUMAN\|Q29RF7_S1276 | 0.654663574 | 1.55E-03 | 1.94E-02 |
| SMG9_HUMAN\|Q9H0W8_T56 | 0.654451584 | 1.87E-04 | 2.63E-03 |
| K1522_HUMAN\|Q9P206_T162 | 0.6544427 | 2.66E-03 | 1.36E-02 |
| CARL1_HUMAN\|Q5VZK9_S1290_T1278 | 0.654438473 | 1.87E-04 | 9.80E-03 |

| | | | |
|---|---|---|---|
| DC1L1_HUMAN\|Q9Y6G9_S516_T515 | 0.654435333 | 2.04E-04 | 1.17E-02 |
| TB182_HUMAN\|Q9C0C2_S435 | 0.654360428 | 2.85E-03 | 1.44E-02 |
| NCOA7_HUMAN\|Q8NI08_S384 | 0.654351679 | 5.61E-05 | 5.91E-04 |
| NDRG1_HUMAN\|Q92597_S330_T328 | 0.654279092 | 4.69E-04 | 9.18E-03 |
| SPT5H_HUMAN\|O00267_S666 | 0.654239083 | 3.03E-05 | 3.57E-04 |
| AFAD_HUMAN\|P55196_T1330 | 0.65420107 | 3.90E-04 | 4.22E-02 |
| SAFB1_HUMAN\|Q15424_S604 | 0.654177371 | 8.79E-06 | 1.37E-04 |
| UBP24_HUMAN\|Q9UPU5_S1143 | 0.654047476 | 2.44E-04 | 3.28E-03 |
| BASP1_HUMAN\|P80723_T196 | 0.654036468 | 7.43E-04 | 1.73E-02 |
| CCNL2_HUMAN\|Q96S94_S330 | 0.654020027 | 2.53E-05 | 3.07E-04 |
| AHNK_HUMAN\|Q09666_S572 | 0.654012979 | 1.50E-03 | 7.66E-03 |
| MTUS1_HUMAN\|Q9ULD2_S1264_S1268 | 0.653958889 | 9.48E-04 | 1.41E-02 |
| NUMA1_HUMAN\|Q14980_T2000 | 0.653906361 | 8.62E-05 | 5.77E-03 |
| K1C17_HUMAN\|Q04695_S13 | 0.653895838 | 4.52E-04 | 5.16E-03 |
| MTAP2_HUMAN\|P11137_S1782_S1787_S1790 | 0.653754598 | 1.71E-04 | 1.36E-03 |
| TNIK_HUMAN\|Q9UKE5_S707 | 0.653742252 | 9.43E-03 | 3.30E-02 |
| TP53B_HUMAN\|Q12888_S1462_S1481 | 0.653735362 | 4.53E-03 | 2.50E-02 |
| SRPK2_HUMAN\|P78362_S497_T492 | 0.653721295 | 1.53E-04 | 9.73E-03 |
| RGS3_HUMAN\|P49796_S946_T941 | 0.65368542 | 1.15E-04 | 1.02E-03 |
| REQU_HUMAN\|Q92785_S142 | 0.653683622 | 6.11E-04 | 1.11E-02 |
| HPS5_HUMAN\|Q9UPZ3_S534 | 0.653631085 | 4.39E-04 | 2.84E-03 |
| PKHG3_HUMAN\|A1L390_T1042 | 0.653600309 | 3.27E-03 | 3.53E-02 |
| IF4G2_HUMAN\|P78344_T508 | 0.653569506 | 1.15E-05 | 7.88E-04 |
| KC1D_HUMAN\|P48730_S331 | 0.653560082 | 1.99E-03 | 1.56E-02 |
| SL9A1_HUMAN\|P19634_S783 | 0.653550988 | 1.29E-02 | 4.96E-02 |
| AHNK_HUMAN\|Q09666_S212 | 0.653541865 | 1.40E-03 | 2.67E-02 |
| SPR2E_HUMAN\|P22531_S60 | 0.65351177 | 7.98E-03 | 3.36E-02 |
| MYPT1_HUMAN\|O14974_S445 | 0.653492807 | 1.70E-04 | 2.46E-03 |
| ZN827_HUMAN\|Q17R98_S963_S984_T959 | 0.653459986 | 8.44E-04 | 5.30E-03 |
| AF1L2_HUMAN\|Q8N4X5_T284 | 0.653452232 | 6.65E-04 | 2.20E-02 |
| NHRF1_HUMAN\|O14745_S280 | 0.653409373 | 1.63E-04 | 9.95E-03 |

| | | | |
|---|---|---|---|
| M3K20_HUMAN\|Q9NYL2_S727 | 0.653281928 | 1.40E-05 | 8.53E-04 |
| BRD4_HUMAN\|O60885_S1117 | 0.653237896 | 5.24E-03 | 2.37E-02 |
| MILK1_HUMAN\|Q8N3F8_S644 | 0.653222466 | 3.31E-04 | 4.05E-03 |
| SRRM2_HUMAN\|Q9UQ35_S1616_S1621 | 0.653222176 | 1.40E-03 | 7.21E-03 |
| IRS2_HUMAN\|Q9Y4H2_S608 | 0.653203182 | 9.33E-05 | 9.12E-04 |
| ZO2_HUMAN\|Q9UDY2_S400 | 0.652986085 | 5.59E-03 | 2.50E-02 |
| EIF3F_HUMAN\|O00303_S258 | 0.65298304 | 1.24E-02 | 4.08E-02 |
| ZCHC7_HUMAN\|Q8N3Z6_S482 | 0.65294633 | 6.42E-04 | 1.14E-02 |
| PRAG1_HUMAN\|Q86YV5_Y413 | 0.652904352 | 1.19E-04 | 1.11E-03 |
| ASPC1_HUMAN\|Q9BZE9_S275 | 0.652898248 | 1.93E-04 | 2.69E-03 |
| AHNK_HUMAN\|Q09666_S2397 | 0.652856153 | 1.61E-03 | 3.92E-02 |
| IF4E_HUMAN\|P06730_T203 | 0.652810617 | 3.64E-04 | 4.39E-03 |
| RBP56_HUMAN\|Q92804_T222 | 0.652729865 | 1.12E-03 | 6.67E-03 |
| K1C19_HUMAN\|P08727_S35 | 0.652681798 | 1.29E-03 | 6.77E-03 |
| OSB10_HUMAN\|Q9BXB5_S26_S32 | 0.652678225 | 2.48E-05 | 5.87E-04 |
| DAPK3_HUMAN\|O43293_S311_S312 | 0.652647103 | 1.58E-04 | 4.31E-03 |
| PHLB1_HUMAN\|Q86UU1_S430 | 0.652631813 | 6.75E-04 | 5.94E-03 |
| AHNK_HUMAN\|Q09666_S572 | 0.652606712 | 1.44E-03 | 8.13E-03 |
| CLAP1_HUMAN\|Q7Z460_S1091 | 0.652598802 | 8.98E-05 | 3.02E-03 |
| BORG4_HUMAN\|Q9H3Q1_S140_S141 | 0.652578325 | 1.19E-02 | 3.94E-02 |
| RPB1_HUMAN\|P24928_T1877_T1885 | 0.65257419 | 4.74E-05 | 9.51E-04 |
| K1C14_HUMAN\|P02533_S437 | 0.652428417 | 3.61E-03 | 1.55E-02 |
| SP100_HUMAN\|P23497_S476 | 0.652375103 | 1.14E-03 | 3.03E-02 |
| IF2B_HUMAN\|P20042_T111 | 0.652353598 | 2.12E-03 | 2.67E-02 |
| BORG2_HUMAN\|Q9UKI2_S100 | 0.652332653 | 1.32E-03 | 1.74E-02 |
| SMG9_HUMAN\|Q9H0W8_T56 | 0.652317902 | 1.71E-04 | 2.46E-03 |
| HEXI2_HUMAN\|Q96MH2_S76 | 0.652250119 | 1.58E-02 | 4.96E-02 |
| ERF_HUMAN\|P50548_T526 | 0.652139266 | 1.17E-03 | 6.94E-03 |
| AHNK_HUMAN\|Q09666_S5731 | 0.651930749 | 3.17E-06 | 1.40E-04 |
| PEAK1_HUMAN\|Q9H792_S1036 | 0.65190717 | 2.71E-05 | 3.18E-04 |
| 1B07_HUMAN\|P01889_S352_S356 | 0.651874776 | 8.94E-04 | 8.78E-03 |

| | | | |
|---|---|---|---|
| HECD1_HUMAN\|Q9ULT8_S1571 | 0.651864885 | 3.41E-03 | 3.77E-02 |
| LMNA_HUMAN\|P02545_S407_T409 | 0.651864283 | 1.48E-03 | 8.33E-03 |
| BAIP2_HUMAN\|Q9UQB8_S453 | 0.651856408 | 3.31E-04 | 4.10E-03 |
| HBS1L_HUMAN\|Q9Y450-2_S246 | 0.651854648 | 2.17E-04 | 7.22E-03 |
| FGD6_HUMAN\|Q6ZV73_T693 | 0.651770598 | 4.70E-04 | 6.81E-03 |
| UBP2L_HUMAN\|Q14157_S467_S477 | 0.651664063 | 1.87E-04 | 4.49E-03 |
| NCOA7_HUMAN\|Q8NI08_S384 | 0.651626274 | 4.94E-05 | 9.77E-04 |
| SVIL_HUMAN\|O95425_T679 | 0.651608394 | 1.52E-05 | 4.08E-04 |
| KC1A_HUMAN\|P48729_T321 | 0.651580489 | 4.58E-03 | 2.12E-02 |
| CSKI2_HUMAN\|Q8WXE0_S403 | 0.651558847 | 3.72E-05 | 1.72E-03 |
| 4EBP1_HUMAN\|Q13541_T70 | 0.651528623 | 3.89E-04 | 2.60E-03 |
| ZFP91_HUMAN\|Q96JP5_S82 | 0.651479888 | 3.64E-04 | 2.49E-03 |
| STK10_HUMAN\|O94804_S448_S455 | 0.65143134 | 1.15E-04 | 1.80E-03 |
| NAV1_HUMAN\|Q8NEY1_T1006 | 0.651418829 | 3.37E-03 | 4.81E-02 |
| SACS_HUMAN\|Q9NZJ4_S4264 | 0.651407603 | 1.55E-03 | 1.12E-02 |
| BAIP2_HUMAN\|Q9UQB8_T340 | 0.651384015 | 7.33E-03 | 4.27E-02 |
| LAP2A_HUMAN\|P42166_S66_S67_T74 | 0.651373506 | 7.38E-04 | 4.30E-03 |
| P4K2A_HUMAN\|Q9BTU6_S468 | 0.651355527 | 2.93E-03 | 2.80E-02 |
| PLEC_HUMAN\|Q15149-2_S24 | 0.651274746 | 1.67E-03 | 1.36E-02 |
| ZC3HD_HUMAN\|Q5T200_S64 | 0.651246725 | 5.47E-04 | 1.03E-02 |
| SFR19_HUMAN\|Q9H7N4_S719_S725 | 0.651232278 | 1.45E-03 | 1.30E-02 |
| LARP1_HUMAN\|Q6PKG0_S1040 | 0.651231116 | 1.21E-04 | 1.11E-03 |
| ZCH18_HUMAN\|Q86VM9_T796 | 0.651173424 | 2.16E-03 | 1.65E-02 |
| NSD3_HUMAN\|Q9BZ95_S457 | 0.651110054 | 1.66E-04 | 9.98E-03 |
| ABLM1_HUMAN\|O14639_S431_T429 | 0.651096304 | 9.32E-05 | 1.51E-03 |
| NDK8_HUMAN\|O60361_S105 | 0.651088192 | 2.46E-04 | 1.93E-03 |
| ITB4_HUMAN\|P16144_S1515_S1517 | 0.650939627 | 3.24E-03 | 2.92E-02 |
| CLAP2_HUMAN\|O75122_S1029 | 0.650893784 | 8.23E-04 | 1.27E-02 |
| KI26B_HUMAN\|Q2KJY2_S1724 | 0.650866549 | 1.06E-03 | 6.38E-03 |
| KC1D_HUMAN\|P48730_S328 | 0.650854188 | 4.38E-03 | 1.81E-02 |
| K1C14_HUMAN\|P02533_T9 | 0.650766686 | 2.51E-03 | 1.30E-02 |

| | | | |
|---|---|---|---|
| 1B07_HUMAN\|P01889_S352_S356 | 0.650706643 | 8.60E-04 | 1.41E-02 |
| BORG2_HUMAN\|Q9UKI2_S100 | 0.650676442 | 1.25E-03 | 3.22E-02 |
| AHNK_HUMAN\|Q09666_S41 | 0.650609151 | 4.91E-03 | 2.25E-02 |
| CTNB1_HUMAN\|P35222_T551 | 0.650598173 | 6.69E-03 | 2.49E-02 |
| ABI2_HUMAN\|Q9NYB9_Y371 | 0.650570202 | 7.10E-05 | 2.65E-03 |
| RFIP5_HUMAN\|Q9BXF6_S395 | 0.650567679 | 1.57E-04 | 5.64E-03 |
| BCAR3_HUMAN\|O75815_S375 | 0.650294395 | 1.04E-03 | 1.01E-02 |
| RMP_HUMAN\|O94763_S372 | 0.650260408 | 8.87E-04 | 5.00E-03 |
| LMNA_HUMAN\|P02545_S404_S407 | 0.650230069 | 2.94E-04 | 2.21E-03 |
| BAG6_HUMAN\|P46379_S1117 | 0.650221387 | 6.11E-03 | 4.65E-02 |
| FUBP3_HUMAN\|Q96I24_S569 | 0.650203391 | 2.05E-03 | 3.47E-02 |
| TISB_HUMAN\|Q07352_S92 | 0.650147158 | 1.58E-03 | 8.81E-03 |
| STMN1_HUMAN\|P16949_S16 | 0.650136571 | 5.01E-04 | 4.69E-02 |
| SCRIB_HUMAN\|Q14160_S1486 | 0.650059395 | 2.73E-04 | 1.32E-02 |
| GRIN1_HUMAN\|Q7Z2K8_S615 | 0.649972683 | 2.07E-04 | 5.17E-03 |
| LIPA3_HUMAN\|O75145_S17 | 0.649950554 | 1.63E-03 | 1.40E-02 |
| UBP2L_HUMAN\|Q14157_S477 | 0.649898398 | 4.29E-03 | 4.25E-02 |
| LRRF2_HUMAN\|Q9Y608_S320_S323 | 0.649839233 | 1.82E-03 | 1.53E-02 |
| SRRM1_HUMAN\|Q8IYB3_S752_S754_S756 | 0.649793719 | 3.66E-05 | 4.06E-04 |
| CLAP1_HUMAN\|Q7Z460-4_S691 | 0.649721063 | 1.96E-03 | 1.06E-02 |
| LARP1_HUMAN\|Q6PKG0_S517_S521 | 0.649639323 | 1.83E-05 | 2.37E-04 |
| DEFI6_HUMAN\|Q9H4E7_S597 | 0.649596624 | 7.05E-06 | 4.44E-04 |
| TB182_HUMAN\|Q9C0C2_S691 | 0.649539261 | 2.82E-04 | 6.05E-03 |
| SMC6_HUMAN\|Q96SB8_S669 | 0.649480524 | 2.15E-03 | 1.64E-02 |
| MILK1_HUMAN\|Q8N3F8_S640_S644 | 0.649466388 | 2.78E-04 | 6.35E-03 |
| TB182_HUMAN\|Q9C0C2_S983 | 0.649414112 | 7.19E-04 | 7.35E-03 |
| CNOT2_HUMAN\|Q9NZN8_S165 | 0.649298781 | 7.39E-05 | 7.45E-04 |
| YAP1_HUMAN\|P46937_S109_T110 | 0.649279546 | 9.81E-06 | 1.44E-04 |
| YAP1_HUMAN\|P46937_S105_S109 | 0.649263888 | 7.52E-06 | 1.20E-04 |
| BORG4_HUMAN\|Q9H3Q1_S292 | 0.649247123 | 1.44E-03 | 7.40E-03 |
| TR150_HUMAN\|Q9Y2W1_T874 | 0.64921747 | 9.16E-04 | 5.13E-03 |

| | | | |
|---|---|---|---|
| K1C14_HUMAN\|P02533_S28 | 0.649207792 | 2.92E-04 | 3.77E-03 |
| CE170_HUMAN\|Q5SW79_S971 | 0.649175679 | 2.08E-04 | 2.91E-03 |
| CRBG1_HUMAN\|Q9Y4K1_S410 | 0.649143625 | 3.54E-03 | 3.46E-02 |
| PEAK1_HUMAN\|Q9H792_T574 | 0.648969316 | 5.86E-05 | 8.77E-04 |
| PHF20_HUMAN\|Q9BVI0_S878_S880 | 0.648953633 | 1.09E-04 | 3.42E-03 |
| FOSL2_HUMAN\|P15408_S308_S320 | 0.648932711 | 3.53E-03 | 2.08E-02 |
| CHSP1_HUMAN\|Q9Y2V2_S52 | 0.648862655 | 1.87E-03 | 1.47E-02 |
| ALKB5_HUMAN\|Q6P6C2_S374_S384 | 0.648839039 | 6.86E-03 | 2.53E-02 |
| EDC3_HUMAN\|Q96F86_S131 | 0.648828565 | 1.26E-05 | 3.65E-04 |
| AHNK_HUMAN\|Q09666_S177 | 0.648795153 | 1.95E-05 | 4.95E-04 |
| ACAP2_HUMAN\|Q15057_S384_T382 | 0.648742518 | 5.35E-07 | 1.51E-05 |
| IRS2_HUMAN\|Q9Y4H2_S1186_S1203 | 0.648703577 | 2.23E-04 | 1.22E-02 |
| DC1L1_HUMAN\|Q9Y6G9_S510_S516 | 0.648389412 | 1.51E-04 | 1.83E-03 |
| K1C16_HUMAN\|P08779_S448 | 0.648342391 | 7.05E-04 | 4.14E-03 |
| MP2K4_HUMAN\|P45985_S393 | 0.648319223 | 5.84E-03 | 2.99E-02 |
| NUP98_HUMAN\|P52948_T1000 | 0.648259776 | 2.02E-03 | 4.54E-02 |
| RAB12_HUMAN\|Q6IQ22_S21 | 0.648244872 | 2.06E-03 | 1.60E-02 |
| JUND_HUMAN\|P17535_S100 | 0.648238979 | 9.12E-04 | 2.73E-02 |
| RTKN_HUMAN\|Q9BST9_S561 | 0.648200455 | 3.36E-04 | 1.48E-02 |
| ADNP_HUMAN\|Q9H2P0_S709 | 0.648149428 | 9.87E-06 | 3.07E-04 |
| TLN1_HUMAN\|Q9Y490_T2270 | 0.6481294 | 6.03E-04 | 6.52E-03 |
| AHNK_HUMAN\|Q09666_S511 | 0.647973384 | 1.18E-03 | 6.94E-03 |
| PLEC_HUMAN\|Q15149_S4618_S4626_T4628 | 0.647921178 | 3.28E-04 | 5.14E-03 |
| RAI14_HUMAN\|Q9P0K7_T297 | 0.647915226 | 1.91E-03 | 1.03E-02 |
| RB6I2_HUMAN\|Q8IUD2_S17_S21 | 0.647902593 | 1.62E-04 | 1.42E-03 |
| PLEC_HUMAN\|Q15149_S4622_Y4612 | 0.647878465 | 5.47E-04 | 1.03E-02 |
| UBP2L_HUMAN\|Q14157_S470_S477 | 0.647837367 | 1.60E-03 | 2.21E-02 |
| VIME_HUMAN\|P08670_S34 | 0.647781566 | 4.33E-04 | 8.69E-03 |
| HMGA2_HUMAN\|P52926_S101_S105_T100 | 0.647761387 | 8.61E-04 | 8.79E-03 |
| PAK6_HUMAN\|Q9NQU5_S347_S351 | 0.647744898 | 2.36E-03 | 1.23E-02 |
| LAP2A_HUMAN\|P42166_S167_T160 | 0.647728607 | 3.94E-04 | 4.72E-03 |

| | | | |
|---|---|---|---|
| PRC2A_HUMAN\|P48634_S1085_S1089_S1092 | 0.647650984 | 9.30E-05 | 1.51E-03 |
| COHA1_HUMAN\|Q9UMD9_S79 | 0.647607801 | 6.74E-03 | 2.91E-02 |
| MA7D1_HUMAN\|Q3KQU3_S446 | 0.647568998 | 4.65E-05 | 9.41E-04 |
| YAP1_HUMAN\|P46937_S109_T110 | 0.647489691 | 8.91E-06 | 5.31E-04 |
| AFAD_HUMAN\|P55196_T1232 | 0.647480305 | 1.04E-03 | 6.26E-03 |
| K2C5_HUMAN\|P13647_S16_S21 | 0.647478496 | 7.83E-03 | 4.30E-02 |
| KS6A1_HUMAN\|Q15418_S363_S369 | 0.647474616 | 1.26E-02 | 4.15E-02 |
| ANS1A_HUMAN\|Q92625_S649 | 0.6474584 | 1.05E-05 | 1.52E-04 |
| KPCD1_HUMAN\|Q15139_S742 | 0.64734195 | 3.53E-03 | 4.87E-02 |
| FA83H_HUMAN\|Q6ZRV2_S523 | 0.647329268 | 3.31E-06 | 3.32E-04 |
| MET_HUMAN\|P08581_S997_T993 | 0.647313083 | 2.22E-03 | 4.57E-02 |
| GRDN_HUMAN\|Q3V6T2_S1653 | 0.647234124 | 1.88E-05 | 4.90E-04 |
| MAP4_HUMAN\|P27816_S928 | 0.647223249 | 3.02E-04 | 6.72E-03 |
| NDRG1_HUMAN\|Q92597_S333_S336_T328 | 0.647165002 | 4.77E-05 | 1.85E-03 |
| TP53B_HUMAN\|Q12888_S830_S834 | 0.647098999 | 3.12E-03 | 1.37E-02 |
| K2C5_HUMAN\|P13647_Y66 | 0.647081863 | 3.48E-05 | 3.92E-04 |
| K1C14_HUMAN\|P02533_S12 | 0.647070047 | 1.36E-04 | 2.04E-03 |
| ATRX_HUMAN\|P46100_S784 | 0.647039184 | 2.63E-03 | 1.67E-02 |
| K2C5_HUMAN\|P13647_S26 | 0.64703514 | 3.02E-04 | 3.83E-03 |
| GOGA4_HUMAN\|Q13439_T31 | 0.647018981 | 1.42E-05 | 4.02E-04 |
| PER1_HUMAN\|O15534_S815 | 0.646980591 | 1.25E-03 | 6.59E-03 |
| PRAG1_HUMAN\|Q86YV5_S907 | 0.646972965 | 1.05E-04 | 1.77E-03 |
| SMAG2_HUMAN\|Q5PRF9_S271 | 0.646897838 | 2.68E-03 | 4.13E-02 |
| PRAG1_HUMAN\|Q86YV5_S510 | 0.646879743 | 1.81E-03 | 9.84E-03 |
| PAIRB_HUMAN\|Q8NC51_S392 | 0.646862459 | 6.20E-04 | 1.08E-02 |
| LRRF2_HUMAN\|Q9Y608_S320_S324 | 0.646860424 | 4.33E-03 | 1.79E-02 |
| BZW2_HUMAN\|Q9Y6E2_S412 | 0.64682365 | 3.47E-03 | 2.06E-02 |
| ELF1_HUMAN\|P32519_S432 | 0.646823524 | 1.28E-06 | 4.85E-05 |
| CRK_HUMAN\|P46108_S304 | 0.646752864 | 1.05E-05 | 1.51E-03 |
| ANXA1_HUMAN\|P04083_S37 | 0.646749585 | 4.32E-03 | 2.02E-02 |
| AFAD_HUMAN\|P55196_S1799 | 0.646703939 | 9.78E-04 | 1.55E-02 |

| | | | |
|---|---|---|---|
| TB182_HUMAN\|Q9C0C2_S1533 | 0.646689645 | 3.74E-04 | 1.09E-02 |
| TISB_HUMAN\|Q07352_S334 | 0.646611637 | 1.65E-03 | 9.11E-03 |
| NU214_HUMAN\|P35658_S1023 | 0.646495845 | 7.13E-04 | 6.15E-03 |
| NCK5L_HUMAN\|Q9HCH0_S489_S494 | 0.646483368 | 1.73E-04 | 2.48E-03 |
| K1C17_HUMAN\|Q04695_S24 | 0.646465399 | 3.37E-04 | 7.28E-03 |
| TWF1_HUMAN\|Q12792_S143 | 0.646457387 | 2.36E-04 | 4.14E-03 |
| SARG_HUMAN\|Q9BW04_S429 | 0.646417363 | 1.96E-03 | 2.53E-02 |
| SPD2A_HUMAN\|Q5TCZ1_S1002 | 0.64635401 | 5.40E-05 | 5.70E-04 |
| VIME_HUMAN\|P08670_T458 | 0.6462868 | 7.39E-05 | 7.45E-04 |
| ELAV1_HUMAN\|Q15717_S202 | 0.646229021 | 2.47E-03 | 1.28E-02 |
| IF2B_HUMAN\|P20042_T111 | 0.646195022 | 1.75E-03 | 3.11E-02 |
| UBP5_HUMAN\|P45974_S785 | 0.64617538 | 1.29E-02 | 4.95E-02 |
| NSD3_HUMAN\|Q9BZ95_S457 | 0.646147283 | 1.34E-04 | 1.67E-03 |
| UBXN7_HUMAN\|O94888_S280 | 0.646125646 | 3.38E-04 | 3.50E-03 |
| COHA1_HUMAN\|Q9UMD9_S148 | 0.646075265 | 6.22E-03 | 2.34E-02 |
| MAP4_HUMAN\|P27816_S1073 | 0.645917164 | 2.00E-04 | 4.75E-03 |
| RELB_HUMAN\|Q01201_S573 | 0.645906757 | 1.51E-04 | 8.50E-03 |
| OSB11_HUMAN\|Q9BXB4_S181 | 0.645890387 | 8.14E-05 | 2.71E-03 |
| MA7D1_HUMAN\|Q3KQU3_S446 | 0.645858861 | 4.29E-05 | 4.70E-04 |
| 1A01_HUMAN\|P30443_S356_S359 | 0.645783595 | 8.17E-04 | 5.19E-03 |
| K1C17_HUMAN\|Q04695_S12 | 0.645674333 | 1.39E-03 | 1.26E-02 |
| BORG4_HUMAN\|Q9H3Q1_S292 | 0.645525852 | 1.28E-03 | 1.78E-02 |
| NUCKS_HUMAN\|Q9H1E3_S181_T179 | 0.645467432 | 7.41E-03 | 3.16E-02 |
| PHLB2_HUMAN\|Q86SQ0_S330_S334 | 0.645428408 | 3.56E-03 | 4.87E-02 |
| CRBG1_HUMAN\|Q9Y4K1_S75 | 0.645311475 | 1.43E-04 | 2.19E-03 |
| ZCHC8_HUMAN\|Q6NZY4_S500_T492_T496 | 0.645245912 | 6.18E-04 | 1.12E-02 |
| YAP1_HUMAN\|P46937_S164 | 0.645201523 | 6.52E-04 | 1.11E-02 |
| PKN2_HUMAN\|Q16513_S362 | 0.645168943 | 3.55E-04 | 2.44E-03 |
| BNIP2_HUMAN\|Q12982_S114 | 0.645101581 | 1.89E-03 | 2.00E-02 |
| IRS2_HUMAN\|Q9Y4H2_S620 | 0.645062433 | 7.54E-04 | 4.88E-03 |
| NDRG2_HUMAN\|Q9UN36_S332_T330 | 0.645060545 | 1.46E-04 | 4.19E-03 |

| | | | |
|---|---|---|---|
| DESP_HUMAN\|P15924_S32 | 0.645003922 | 7.35E-03 | 4.28E-02 |
| STK10_HUMAN\|O94804_S455 | 0.644919901 | 9.91E-05 | 9.60E-04 |
| K1C14_HUMAN\|P02533_S13 | 0.644848958 | 5.79E-05 | 1.07E-03 |
| PKP2_HUMAN\|Q99959_S197 | 0.644771101 | 1.52E-04 | 1.34E-03 |
| IASPP_HUMAN\|Q8WUF5_S134 | 0.644719073 | 8.97E-04 | 2.73E-02 |
| DC1I2_HUMAN\|Q13409-2_S84 | 0.644712111 | 2.28E-04 | 1.22E-02 |
| PLEC_HUMAN\|Q15149_S4384_S4385 | 0.644696432 | 9.16E-04 | 1.13E-02 |
| CTNA1_HUMAN\|P35221_S641_T645 | 0.644558256 | 5.03E-04 | 3.48E-03 |
| PR38B_HUMAN\|Q5VTL8_S529 | 0.64452702 | 2.02E-04 | 2.87E-03 |
| SPIR1_HUMAN\|Q08AE8_T509 | 0.644436158 | 2.08E-04 | 2.91E-03 |
| CRTC3_HUMAN\|Q6UUV7_S370 | 0.64437398 | 1.76E-03 | 1.41E-02 |
| DCBD2_HUMAN\|Q96PD2_S727 | 0.644366805 | 2.40E-03 | 1.25E-02 |
| TGS1_HUMAN\|Q96RS0_S85_S89 | 0.644324784 | 3.30E-03 | 1.44E-02 |
| K1C16_HUMAN\|P08779_S450 | 0.644134553 | 3.02E-04 | 2.15E-03 |
| VIP2_HUMAN\|O43314_S1006_S1013_S1016 | 0.64403525 | 1.61E-05 | 2.13E-04 |
| EFNB1_HUMAN\|P98172_S281 | 0.643933197 | 5.01E-04 | 1.34E-02 |
| SMC1A_HUMAN\|Q14683_S956 | 0.643879002 | 1.08E-03 | 5.88E-03 |
| AKTS1_HUMAN\|Q96B36_S212 | 0.643798996 | 4.34E-03 | 1.79E-02 |
| FUBP3_HUMAN\|Q96I24_S569 | 0.643711752 | 1.69E-03 | 8.41E-03 |
| LMNB2_HUMAN\|Q03252_S420 | 0.64370155 | 1.30E-04 | 1.11E-03 |
| ABLM1_HUMAN\|O14639_S458 | 0.643523451 | 3.58E-03 | 3.87E-02 |
| GMIP_HUMAN\|Q9P107_S914_S919 | 0.643429655 | 8.88E-05 | 8.26E-04 |
| AHNK_HUMAN\|Q09666_T4430 | 0.643257358 | 5.25E-04 | 1.38E-02 |
| ARP19_HUMAN\|P56211_S62 | 0.643193881 | 1.62E-04 | 5.79E-03 |
| GGA2_HUMAN\|Q9UJY4_S177 | 0.643153239 | 9.28E-04 | 5.70E-03 |
| SCRIB_HUMAN\|Q14160_S1475_S1486 | 0.643137 | 6.95E-03 | 3.41E-02 |
| RHG23_HUMAN\|Q9P227_S611 | 0.643100848 | 6.87E-05 | 7.07E-04 |
| RIN1_HUMAN\|Q13671_S349 | 0.642972127 | 2.52E-03 | 2.51E-02 |
| TOR4A_HUMAN\|Q9NXH8_S63 | 0.642921793 | 2.73E-03 | 1.39E-02 |
| AHNK_HUMAN\|Q09666_S5749 | 0.642896552 | 5.44E-06 | 4.27E-04 |
| RD23B_HUMAN\|P54727_T162 | 0.642884185 | 8.62E-03 | 4.04E-02 |

| | | | |
|---|---|---|---|
| YAP1_HUMAN\|P46937_S164 | 0.642874657 | 6.01E-04 | 1.49E-02 |
| IF4G1_HUMAN\|Q04637_S1187 | 0.642854552 | 1.04E-04 | 9.96E-04 |
| YAP1_HUMAN\|P46937_S109 | 0.642817363 | 3.24E-04 | 2.27E-03 |
| PHF20_HUMAN\|Q9BVI0_S878_S880 | 0.642812852 | 8.31E-05 | 1.49E-03 |
| LMNA_HUMAN\|P02545_S406_S407 | 0.642800921 | 2.73E-04 | 1.97E-03 |
| TAF6_HUMAN\|P49848_S532 | 0.642680509 | 2.56E-05 | 4.68E-04 |
| AHNK2_HUMAN\|Q8IVF2_S5175 | 0.64255578 | 5.46E-04 | 1.98E-02 |
| AHNK_HUMAN\|Q09666_S3426 | 0.642529203 | 1.04E-04 | 9.39E-04 |
| SLK_HUMAN\|Q9H2G2_T820 | 0.642506902 | 1.49E-05 | 2.03E-04 |
| IASPP_HUMAN\|Q8WUF5_S395 | 0.642480558 | 1.83E-03 | 2.40E-02 |
| MRP_HUMAN\|P49006_T178 | 0.642435214 | 3.85E-05 | 4.22E-04 |
| ZYX_HUMAN\|Q15942_S259_S267 | 0.642432185 | 3.03E-04 | 2.27E-03 |
| LMNA_HUMAN\|P02545_S392_S395 | 0.642351364 | 1.21E-02 | 4.71E-02 |
| RPGF6_HUMAN\|Q8TEU7_S1590 | 0.642321741 | 3.43E-04 | 2.52E-03 |
| KC1A_HUMAN\|P48729_T321 | 0.642207027 | 3.51E-03 | 1.52E-02 |
| RBM14_HUMAN\|Q96PK6_S571 | 0.642192194 | 3.64E-03 | 2.56E-02 |
| TENS4_HUMAN\|Q8IZW8_S253 | 0.64218441 | 3.56E-05 | 3.96E-04 |
| FA83G_HUMAN\|A6ND36_S726 | 0.642182406 | 7.72E-04 | 4.97E-03 |
| CBX3_HUMAN\|Q13185_S93 | 0.642042691 | 5.56E-03 | 3.33E-02 |
| CR025_HUMAN\|Q96B23_S66_S69 | 0.642003423 | 6.99E-03 | 3.97E-02 |
| ELAV1_HUMAN\|Q15717_S202 | 0.641989825 | 2.19E-03 | 2.73E-02 |
| PLEC_HUMAN\|Q15149_S4622_Y4612 | 0.641953222 | 4.41E-04 | 5.09E-03 |
| WAC2A_HUMAN\|Q641Q2_S390 | 0.641891078 | 1.31E-03 | 1.81E-02 |
| CDK13_HUMAN\|Q14004_T1056 | 0.641836521 | 3.63E-04 | 2.63E-03 |
| RRP1B_HUMAN\|Q14684_S706 | 0.641775069 | 5.29E-04 | 4.97E-03 |
| IF4G2_HUMAN\|P78344_T508 | 0.641743014 | 6.10E-06 | 2.21E-04 |
| SCRIB_HUMAN\|Q14160_S853 | 0.64163529 | 5.13E-03 | 2.03E-02 |
| AHNK_HUMAN\|Q09666_S210_S216_S220 | 0.641477015 | 6.82E-05 | 6.63E-04 |
| BASP1_HUMAN\|P80723_T196 | 0.641453687 | 4.64E-04 | 2.98E-03 |
| NFKB1_HUMAN\|P19838_S907 | 0.641453372 | 3.47E-06 | 1.02E-04 |
| PAK6_HUMAN\|Q9NQU5_S347_S351 | 0.641422805 | 1.94E-03 | 2.34E-02 |

| | | | |
|---|---|---|---|
| RBP10_HUMAN\|Q6VN20_S365 | 0.641408036 | 1.98E-05 | 1.09E-03 |
| ARHGB_HUMAN\|O15085_T668_T672 | 0.641386515 | 4.57E-05 | 1.63E-03 |
| AKAP2_HUMAN\|Q9Y2D5_T157 | 0.641361734 | 3.13E-04 | 3.28E-03 |
| ACINU_HUMAN\|Q9UKV3_T254 | 0.641319457 | 2.53E-03 | 2.85E-02 |
| YAP1_HUMAN\|P46937_S164 | 0.641221126 | 5.67E-04 | 6.20E-03 |
| NIBL1_HUMAN\|Q96TA1_S641_S646 | 0.641125679 | 2.62E-03 | 2.50E-02 |
| ZN609_HUMAN\|O15014_S358_S361 | 0.641119432 | 8.86E-03 | 4.74E-02 |
| DBNL_HUMAN\|Q9UJU6-2_S232 | 0.641030401 | 8.16E-04 | 1.85E-02 |
| TOM20_HUMAN\|Q15388_S138 | 0.6409092 | 3.92E-03 | 1.86E-02 |
| TR10A_HUMAN\|O00220_S466 | 0.640904966 | 4.75E-03 | 3.81E-02 |
| HEXI1_HUMAN\|O94992_S237_S252 | 0.640843893 | 6.08E-03 | 4.65E-02 |
| SMG9_HUMAN\|Q9H0W8_S53 | 0.640834901 | 2.39E-05 | 2.93E-04 |
| ARAF_HUMAN\|P10398_S157 | 0.64080177 | 5.26E-05 | 5.39E-04 |
| K1671_HUMAN\|Q9BY89_S128 | 0.64074304 | 1.55E-05 | 2.07E-04 |
| RAI14_HUMAN\|Q9P0K7_S296 | 0.640717092 | 9.21E-04 | 1.47E-02 |
| LMNA_HUMAN\|P02545_S404_S406_S414 | 0.64068631 | 5.43E-03 | 3.45E-02 |
| M4K4_HUMAN\|O95819_S842 | 0.640651611 | 6.15E-05 | 6.09E-04 |
| ROA1_HUMAN\|P09651_S361_S364 | 0.640604208 | 2.22E-03 | 1.05E-02 |
| PAXI_HUMAN\|P49023_S103_S106 | 0.640467477 | 2.80E-03 | 4.25E-02 |
| DTD1_HUMAN\|Q8TEA8_S196_S197 | 0.640417198 | 2.49E-05 | 4.60E-04 |
| K1C16_HUMAN\|P08779_S448_S451 | 0.640413522 | 4.94E-04 | 5.50E-03 |
| STIM1_HUMAN\|Q13586_S668 | 0.640378024 | 4.82E-03 | 1.94E-02 |
| LIPA1_HUMAN\|Q13136_S666_S668 | 0.640367536 | 8.10E-05 | 1.46E-03 |
| MAN1_HUMAN\|Q9Y2U8_S144 | 0.640348865 | 3.37E-03 | 2.02E-02 |
| AKAP2_HUMAN\|Q9Y2D5_S748 | 0.640344947 | 2.73E-03 | 2.89E-02 |
| K1C17_HUMAN\|Q04695_S39 | 0.640310628 | 2.68E-03 | 1.22E-02 |
| STX7_HUMAN\|O15400_S131 | 0.64026631 | 4.38E-05 | 1.58E-03 |
| TB182_HUMAN\|Q9C0C2_S1103 | 0.640236499 | 4.16E-04 | 4.07E-03 |
| RRBP1_HUMAN\|Q9P2E9_S615 | 0.640234524 | 2.64E-04 | 5.77E-03 |
| FMNL2_HUMAN\|Q96PY5_S1016 | 0.640212014 | 6.35E-04 | 5.70E-03 |
| GPBP1_HUMAN\|Q86WP2_S381 | 0.640167984 | 4.37E-04 | 2.84E-03 |

| | | | |
|---|---|---|---|
| AHNK2_HUMAN\|Q8IVF2_S5175 | 0.640155707 | 5.00E-04 | 7.71E-03 |
| LYRIC_HUMAN\|Q86UE4_S298 | 0.640143756 | 8.38E-03 | 3.96E-02 |
| COHA1_HUMAN\|Q9UMD9_T182 | 0.640118095 | 1.58E-03 | 2.05E-02 |
| ES8L1_HUMAN\|Q8TE68_S182_T187 | 0.640080773 | 1.18E-05 | 1.69E-04 |
| SQSTM_HUMAN\|Q13501_S266 | 0.63991508 | 3.56E-05 | 7.66E-04 |
| LMO7_HUMAN\|Q8WWI1_S1421 | 0.639867715 | 1.86E-03 | 9.03E-03 |
| LMNA_HUMAN\|P02545_S616 | 0.639775543 | 1.84E-04 | 1.57E-03 |
| NDRG1_HUMAN\|Q92597_S330_S333_S336 | 0.639771172 | 6.85E-03 | 3.90E-02 |
| PKHF2_HUMAN\|Q9H8W4_S239 | 0.639764963 | 1.70E-04 | 4.16E-03 |
| RFIP5_HUMAN\|Q9BXF6_S296 | 0.639729057 | 4.85E-06 | 3.33E-04 |
| ZN654_HUMAN\|Q8IZM8_S576 | 0.639683445 | 8.63E-03 | 3.08E-02 |
| AHNK_HUMAN\|Q09666_S210_S216 | 0.639675106 | 1.31E-04 | 1.64E-03 |
| FRYL_HUMAN\|O94915_T1959 | 0.639674578 | 1.17E-04 | 1.03E-03 |
| MGMT_HUMAN\|P16455_S201 | 0.639644802 | 1.47E-06 | 1.47E-04 |
| AFAD_HUMAN\|P55196_T1232 | 0.639638559 | 8.09E-04 | 8.30E-03 |
| K2C5_HUMAN\|P13647_S50 | 0.639593385 | 6.28E-05 | 1.14E-03 |
| NUFP2_HUMAN\|Q7Z417_S212_S214 | 0.639507842 | 3.14E-04 | 6.39E-03 |
| ADCY6_HUMAN\|O43306_S54 | 0.639394939 | 1.97E-04 | 1.64E-03 |
| MA7D1_HUMAN\|Q3KQU3_S254 | 0.639368607 | 3.16E-05 | 1.33E-03 |
| ZN827_HUMAN\|Q17R98_S963_S984_T959 | 0.639309296 | 5.17E-04 | 3.25E-03 |
| ZCCHV_HUMAN\|Q7Z2W4_S387 | 0.639302607 | 2.51E-03 | 1.16E-02 |
| TB182_HUMAN\|Q9C0C2_S1652 | 0.639295568 | 2.05E-04 | 5.17E-03 |
| FA83H_HUMAN\|Q6ZRV2_S1009 | 0.639222604 | 1.32E-03 | 2.55E-02 |
| CR025_HUMAN\|Q96B23_S143_S145 | 0.639162583 | 5.09E-05 | 9.92E-04 |
| LARP1_HUMAN\|Q6PKG0_S766_S774 | 0.639147531 | 2.16E-03 | 2.68E-02 |
| K1C15_HUMAN\|P19012_S34 | 0.639069963 | 7.18E-03 | 4.21E-02 |
| FBP1L_HUMAN\|Q5T0N5_S295 | 0.639011466 | 4.36E-04 | 2.83E-03 |
| PATL1_HUMAN\|Q86TB9_S179_S184 | 0.63896674 | 1.32E-03 | 2.55E-02 |
| SRS10_HUMAN\|O75494_S106 | 0.638939064 | 2.76E-03 | 4.21E-02 |
| RCC1_HUMAN\|P18754_S11 | 0.638929082 | 4.28E-03 | 2.77E-02 |
| YAP1_HUMAN\|P46937_S164 | 0.638908615 | 5.22E-04 | 3.57E-03 |

| | | | |
|---|---|---|---|
| ARAF_HUMAN\|P10398_S157 | 0.638906014 | 4.85E-05 | 9.57E-04 |
| PHLB1_HUMAN\|Q86UU1_S461 | 0.63890461 | 5.64E-06 | 9.74E-05 |
| TPBG_HUMAN\|Q13641_S415_S418 | 0.638902342 | 5.63E-05 | 8.54E-04 |
| TWF1_HUMAN\|Q12792_T349 | 0.63885937 | 2.20E-05 | 2.70E-04 |
| KDM3B_HUMAN\|Q7LBC6_S293 | 0.638849871 | 6.16E-06 | 1.57E-04 |
| SLAI2_HUMAN\|Q9P270_S349_S353_S357 | 0.638812758 | 4.32E-05 | 1.75E-03 |
| AHNK_HUMAN\|Q09666_T5729 | 0.638797177 | 3.28E-05 | 1.39E-03 |
| Z280C_HUMAN\|Q8ND82_S80 | 0.638749419 | 8.74E-05 | 5.78E-03 |
| KANK1_HUMAN\|Q14678_S144 | 0.638731951 | 1.51E-06 | 3.39E-05 |
| MA7D1_HUMAN\|Q3KQU3_S834 | 0.638728514 | 4.93E-04 | 1.90E-02 |
| KIZ_HUMAN\|Q2M2Z5_S623 | 0.638594702 | 5.90E-05 | 1.09E-03 |
| HELZ_HUMAN\|P42694_T1163 | 0.638580996 | 1.26E-06 | 3.10E-05 |
| CRBG1_HUMAN\|Q9Y4K1_S141 | 0.638506128 | 4.95E-04 | 9.34E-03 |
| CE170_HUMAN\|Q5SW79_S971 | 0.638493229 | 1.35E-04 | 3.88E-03 |
| TISB_HUMAN\|Q07352_S92 | 0.638436672 | 1.09E-03 | 1.05E-02 |
| ML12A_HUMAN\|P19105_T18 | 0.638356789 | 3.82E-04 | 1.10E-02 |
| NP1L1_HUMAN\|P55209_T62 | 0.638340591 | 3.84E-03 | 3.87E-02 |
| SPD2A_HUMAN\|Q5TCZ1_S1038 | 0.638263082 | 1.13E-06 | 1.32E-04 |
| RTKN_HUMAN\|Q9BST9_T518 | 0.63824054 | 2.30E-04 | 1.84E-03 |
| K2C5_HUMAN\|P13647_T19 | 0.638177437 | 4.94E-03 | 3.05E-02 |
| RFIP5_HUMAN\|Q9BXF6_S397 | 0.638098392 | 2.05E-04 | 6.92E-03 |
| SMC1A_HUMAN\|Q14683_S956 | 0.637959482 | 8.93E-04 | 8.78E-03 |
| 3BP1_HUMAN\|Q9Y3L3_T601 | 0.637954352 | 4.74E-03 | 2.99E-02 |
| RHG29_HUMAN\|Q52LW3_S176 | 0.63792046 | 2.96E-03 | 2.74E-02 |
| INADL_HUMAN\|Q8NI35_S645 | 0.637831263 | 4.10E-05 | 8.41E-04 |
| STIM1_HUMAN\|Q13586_S668 | 0.637792131 | 4.49E-03 | 4.32E-02 |
| BIG1_HUMAN\|Q9Y6D6_S394 | 0.637783069 | 1.36E-04 | 3.88E-03 |
| TGS1_HUMAN\|Q96RS0_S154 | 0.637782406 | 3.11E-04 | 2.19E-03 |
| TBC15_HUMAN\|Q8TC07-2_S205 | 0.637771292 | 1.37E-03 | 1.25E-02 |
| MA7D1_HUMAN\|Q3KQU3_S254 | 0.637750888 | 2.92E-05 | 6.54E-04 |
| PKHA1_HUMAN\|Q9HB21_S332 | 0.637734253 | 6.95E-06 | 1.13E-04 |

| | | | |
|---|---|---|---|
| AHNK_HUMAN\|Q09666_S5731 | 0.637710678 | 1.37E-06 | 1.35E-04 |
| PDLI2_HUMAN\|Q96JY6_S137_S143 | 0.637677438 | 1.68E-04 | 2.45E-03 |
| PACN2_HUMAN\|Q9UNF0_S399_S425 | 0.637649475 | 1.14E-04 | 1.01E-03 |
| VIP2_HUMAN\|O43314_S1006_S1013_S1016 | 0.637647167 | 1.17E-05 | 7.73E-04 |
| NUP50_HUMAN\|Q9UKX7_S221 | 0.63763853 | 3.68E-05 | 1.78E-03 |
| TB10B_HUMAN\|Q4KMP7_S22 | 0.637592896 | 6.99E-05 | 2.20E-03 |
| AMOL2_HUMAN\|Q9Y2J4_S759_S762 | 0.637504855 | 8.30E-03 | 2.98E-02 |
| FSTL3_HUMAN\|O95633_S255 | 0.637404559 | 6.57E-04 | 1.11E-02 |
| OSB10_HUMAN\|Q9BXB5_S32 | 0.63738331 | 1.89E-04 | 1.60E-03 |
| PRAG1_HUMAN\|Q86YV5_Y413 | 0.6373786 | 6.15E-05 | 1.13E-03 |
| AFAP1_HUMAN\|Q8N556_S664_S668 | 0.637357071 | 1.19E-02 | 3.95E-02 |
| WAC2A_HUMAN\|Q641Q2_S284_S288 | 0.637354268 | 2.78E-04 | 2.99E-03 |
| ITB4_HUMAN\|P16144_S1454_T1455 | 0.637237588 | 4.79E-03 | 1.94E-02 |
| K1C14_HUMAN\|P02533_S432_S433_S437 | 0.637162425 | 1.73E-03 | 8.57E-03 |
| TB182_HUMAN\|Q9C0C2_S872 | 0.637145279 | 6.35E-05 | 6.55E-04 |
| ARHG5_HUMAN\|Q12774_S450 | 0.637123031 | 1.42E-03 | 3.49E-02 |
| LMNB2_HUMAN\|Q03252_S421 | 0.637114357 | 1.50E-03 | 7.66E-03 |
| STIM2_HUMAN\|Q9P246_S522 | 0.63704432 | 7.02E-03 | 2.58E-02 |
| NP1L1_HUMAN\|P55209_T62 | 0.637005555 | 3.71E-03 | 4.95E-02 |
| AHNK_HUMAN\|Q09666_S5864 | 0.636995968 | 8.84E-03 | 3.14E-02 |
| ARH40_HUMAN\|Q8TER5_S262 | 0.636985816 | 3.70E-05 | 1.48E-03 |
| PAXI_HUMAN\|P49023_S126_S130 | 0.636909768 | 3.05E-03 | 2.79E-02 |
| AHNK2_HUMAN\|Q8IVF2_S294_S297 | 0.636870591 | 7.52E-06 | 1.20E-04 |
| LMO7_HUMAN\|Q8WWI1_S1421 | 0.636868279 | 1.70E-03 | 2.30E-02 |
| NFIX_HUMAN\|Q14938_S280 | 0.636829345 | 2.82E-05 | 5.01E-04 |
| VIME_HUMAN\|P08670_S430 | 0.636688064 | 1.44E-04 | 1.29E-03 |
| MISP_HUMAN\|Q8IVT2_S394_S395 | 0.636621227 | 7.33E-05 | 6.25E-03 |
| K1C14_HUMAN\|P02533_S10 | 0.63661673 | 1.34E-02 | 4.37E-02 |
| VIME_HUMAN\|P08670_S56 | 0.636534384 | 4.50E-05 | 1.77E-03 |
| CAMP1_HUMAN\|Q5T5Y3_S1080 | 0.636523367 | 8.73E-05 | 1.17E-03 |
| EPN1_HUMAN\|Q9Y6I3_S454 | 0.636467067 | 3.08E-05 | 1.36E-03 |

| | | | |
|---|---|---|---|
| SGTB_HUMAN\|Q96EQ0_S297 | 0.636313761 | 1.00E-04 | 1.31E-03 |
| LIPA1_HUMAN\|Q13136_S1172 | 0.636295728 | 2.61E-05 | 6.04E-04 |
| BAD_HUMAN\|Q92934_S74 | 0.636253371 | 3.21E-04 | 9.90E-03 |
| IRS2_HUMAN\|Q9Y4H2_T350 | 0.636167164 | 3.28E-05 | 7.11E-04 |
| CAMP3_HUMAN\|Q9P1Y5_S756 | 0.636116751 | 1.07E-04 | 2.96E-03 |
| PAK1_HUMAN\|Q13153_S223 | 0.636096185 | 6.69E-04 | 5.92E-03 |
| PAK6_HUMAN\|Q9NQU5_S308 | 0.63608416 | 4.94E-03 | 2.26E-02 |
| BAIP2_HUMAN\|Q9UQB8_S366 | 0.636072202 | 3.93E-03 | 1.67E-02 |
| TR10A_HUMAN\|O00220_S466 | 0.636070129 | 4.18E-03 | 2.73E-02 |
| ANXA2_HUMAN\|P07355_T19 | 0.636067683 | 4.06E-06 | 2.98E-04 |
| GRDN_HUMAN\|Q3V6T2_S1653 | 0.636005617 | 1.09E-05 | 1.56E-04 |
| M3K20_HUMAN\|Q9NYL2_S633 | 0.635982034 | 2.66E-03 | 1.21E-02 |
| G3BP1_HUMAN\|Q13283_S230 | 0.635858237 | 1.98E-03 | 2.54E-02 |
| FOXO3_HUMAN\|O43524_S294 | 0.635849309 | 1.17E-03 | 1.62E-02 |
| PPR35_HUMAN\|Q8TAP8_S52 | 0.635799503 | 3.69E-04 | 4.45E-03 |
| MRP_HUMAN\|P49006_T178 | 0.635719189 | 2.79E-05 | 1.50E-03 |
| MICA3_HUMAN\|Q7RTP6_S1636 | 0.635651565 | 3.59E-03 | 1.74E-02 |
| DC1I2_HUMAN\|Q13409-2_S84 | 0.63564604 | 1.57E-04 | 2.66E-02 |
| LMNB2_HUMAN\|Q03252_S424 | 0.635634447 | 6.87E-04 | 1.20E-02 |
| LAD1_HUMAN\|O00515_S38_T40_T41 | 0.635517582 | 8.38E-04 | 2.63E-02 |
| CDK16_HUMAN\|Q00536_T135 | 0.635480078 | 6.10E-04 | 6.52E-03 |
| USO1_HUMAN\|O60763_S952 | 0.635478144 | 2.34E-03 | 2.38E-02 |
| F117A_HUMAN\|Q9C073_S67 | 0.635365067 | 3.01E-04 | 2.14E-03 |
| MEF2D_HUMAN\|Q14814_S231 | 0.63534547 | 4.05E-03 | 2.80E-02 |
| PLEC_HUMAN\|Q15149_S4386_S4396 | 0.635336573 | 1.35E-05 | 1.86E-04 |
| FA83H_HUMAN\|Q6ZRV2_S523 | 0.635296799 | 1.64E-06 | 3.63E-05 |
| MINK1_HUMAN\|Q8N4C8_S701 | 0.635279352 | 5.56E-03 | 2.49E-02 |
| AHNK_HUMAN\|Q09666_S210_S216 | 0.635269675 | 1.09E-04 | 3.11E-03 |
| TR150_HUMAN\|Q9Y2W1_T874 | 0.63521744 | 5.68E-04 | 6.20E-03 |
| ITA6_HUMAN\|P23229_S1128 | 0.635198258 | 9.05E-04 | 1.46E-02 |
| RIOK2_HUMAN\|Q9BVS4_S382_S385 | 0.635176237 | 1.39E-05 | 1.91E-04 |

| | | | |
|---|---|---|---|
| PAK2_HUMAN\|Q13177_S141 | 0.63513871 | 7.21E-04 | 7.57E-03 |
| BZW2_HUMAN\|Q9Y6E2_S412 | 0.635107803 | 2.51E-03 | 2.73E-02 |
| 3BP1_HUMAN\|Q9Y3L3_S596 | 0.634924284 | 1.56E-04 | 4.31E-03 |
| MILK1_HUMAN\|Q8N3F8_S323 | 0.634922837 | 5.76E-03 | 3.41E-02 |
| MFAP1_HUMAN\|P55081_S80 | 0.634852533 | 1.41E-05 | 1.91E-04 |
| MILK1_HUMAN\|Q8N3F8_S391 | 0.63478081 | 1.29E-06 | 6.20E-05 |
| TB182_HUMAN\|Q9C0C2_T875 | 0.634744883 | 7.13E-05 | 2.65E-03 |
| PP14A_HUMAN\|Q96A00_S128_S134_S136 | 0.634607606 | 2.75E-05 | 3.50E-03 |
| RIN1_HUMAN\|Q13671_S333_S337 | 0.634497002 | 1.02E-04 | 2.42E-03 |
| ANR17_HUMAN\|O75179_S2041_S2045 | 0.634433264 | 3.63E-04 | 2.63E-03 |
| PRAG1_HUMAN\|Q86YV5_S510 | 0.634404853 | 1.24E-03 | 1.84E-02 |
| MRGBP_HUMAN\|Q9NV56_S195 | 0.634381247 | 1.63E-03 | 8.16E-03 |
| SDS3_HUMAN\|Q9H7L9_S45_S53_T49 | 0.634376991 | 1.06E-03 | 5.81E-03 |
| MILK1_HUMAN\|Q8N3F8_T318 | 0.634195854 | 4.61E-03 | 2.94E-02 |
| PKP3_HUMAN\|Q9Y446_S313_S314 | 0.634183697 | 4.46E-05 | 4.85E-04 |
| PKHA1_HUMAN\|Q9HB21_S177 | 0.634181656 | 3.76E-03 | 1.80E-02 |
| SAFB1_HUMAN\|Q15424_S247 | 0.63409738 | 5.67E-05 | 2.44E-03 |
| SPTB2_HUMAN\|Q01082-3_S10 | 0.633990662 | 2.12E-04 | 1.63E-03 |
| ERCC5_HUMAN\|P28715_S382_S384 | 0.633967926 | 1.91E-03 | 9.25E-03 |
| WDR13_HUMAN\|Q9H1Z4_S79 | 0.633916775 | 1.60E-04 | 9.83E-03 |
| MARK2_HUMAN\|Q7KZI7-13_S376 | 0.633882314 | 4.40E-03 | 1.81E-02 |
| SENP3_HUMAN\|Q9H4L4_S169_S181_T176 | 0.633781201 | 4.51E-03 | 2.10E-02 |
| S23A2_HUMAN\|Q9UGH3_S78_S81 | 0.633739537 | 3.95E-03 | 2.25E-02 |
| NIBL1_HUMAN\|Q96TA1_S691 | 0.633680026 | 3.97E-06 | 1.04E-03 |
| K1C14_HUMAN\|P02533_T9 | 0.633638759 | 1.47E-03 | 1.31E-02 |
| EIF3A_HUMAN\|Q14152_S882 | 0.633613206 | 3.17E-03 | 2.21E-02 |
| K2C5_HUMAN\|P13647_S82_T78 | 0.633527973 | 1.26E-04 | 2.00E-03 |
| 1433B_HUMAN\|P31946_S47 | 0.633498231 | 2.23E-03 | 1.77E-02 |
| MA7D1_HUMAN\|Q3KQU3_T97 | 0.63348097 | 1.59E-03 | 8.01E-03 |
| COHA1_HUMAN\|Q9UMD9_S61 | 0.63343777 | 1.88E-03 | 1.56E-02 |
| CE170_HUMAN\|Q5SW79_S971 | 0.6331236 | 1.08E-04 | 3.44E-03 |

| | | | |
|---|---|---|---|
| VIME_HUMAN\|P08670_S430 | 0.633037042 | 1.23E-04 | 1.07E-03 |
| ZO2_HUMAN\|Q9UDY2_S398 | 0.633017184 | 2.17E-03 | 2.20E-02 |
| PATL1_HUMAN\|Q86TB9_S179_S184 | 0.632968401 | 1.10E-03 | 1.63E-02 |
| CE170_HUMAN\|Q5SW79_S630_S636 | 0.632927341 | 3.23E-05 | 7.07E-04 |
| PDLI2_HUMAN\|Q96JY6_S134 | 0.632927015 | 1.29E-06 | 1.80E-04 |
| OXR1_HUMAN\|Q8N573_S201_S202 | 0.632823795 | 3.80E-05 | 6.26E-04 |
| CAMP2_HUMAN\|Q08AD1_S1319 | 0.632685832 | 4.10E-05 | 1.84E-03 |
| ZEP1_HUMAN\|P15822_S537 | 0.632664992 | 1.10E-03 | 1.55E-02 |
| PKHA1_HUMAN\|Q9HB21_S332 | 0.632633725 | 5.36E-06 | 9.31E-05 |
| TPD54_HUMAN\|O43399_S96 | 0.632594618 | 3.29E-03 | 3.05E-02 |
| CCNH_HUMAN\|P51946_T315 | 0.632481713 | 6.73E-04 | 4.42E-03 |
| TT39B_HUMAN\|Q5VTQ0_S120 | 0.632432841 | 2.22E-03 | 2.72E-02 |
| AAPK1_HUMAN\|Q13131_T488_T490 | 0.632372632 | 1.33E-04 | 2.07E-03 |
| RFIP5_HUMAN\|Q9BXF6_S397 | 0.632363521 | 1.63E-04 | 1.95E-03 |
| K1C17_HUMAN\|Q04695_T9 | 0.632349085 | 9.12E-04 | 5.65E-03 |
| MAP4_HUMAN\|P27816_S941 | 0.632278545 | 2.84E-04 | 6.44E-03 |
| UBP2L_HUMAN\|Q14157_S470_S477 | 0.632220415 | 9.55E-04 | 9.13E-03 |
| RPRD2_HUMAN\|Q5VT52_T723 | 0.632093003 | 6.37E-06 | 5.16E-04 |
| SARG_HUMAN\|Q9BW04_T456 | 0.6320596 | 3.57E-06 | 3.40E-04 |
| SNTB2_HUMAN\|Q13425_S222 | 0.632045759 | 6.14E-03 | 4.99E-02 |
| PLXB2_HUMAN\|O15031_S1570 | 0.632034497 | 7.58E-04 | 4.90E-03 |
| ML12A_HUMAN\|P19105_T18 | 0.632025966 | 3.03E-04 | 6.64E-03 |
| CHSP1_HUMAN\|Q9Y2V2_S52 | 0.632010767 | 1.07E-03 | 1.03E-02 |
| EGFR_HUMAN\|P00533_S1042 | 0.631943968 | 3.98E-03 | 2.27E-02 |
| PRSR2_HUMAN\|Q86WR7_S212 | 0.631881175 | 3.76E-05 | 6.24E-04 |
| LRRF2_HUMAN\|Q9Y608_S320_S324 | 0.631788672 | 2.85E-03 | 2.11E-02 |
| IF4E_HUMAN\|P06730_T203 | 0.631770701 | 1.60E-04 | 4.01E-03 |
| ZBT21_HUMAN\|Q9ULJ3_S435_T431 | 0.631742339 | 3.75E-05 | 8.01E-04 |
| PP14A_HUMAN\|Q96A00_S26 | 0.631727496 | 2.04E-05 | 2.53E-04 |
| MISP_HUMAN\|Q8IVT2_S400 | 0.63166962 | 1.18E-03 | 3.18E-02 |
| TBC17_HUMAN\|Q9HA65_T620 | 0.631662679 | 1.75E-04 | 4.62E-03 |

| | | | |
|---|---|---|---|
| ZC11A_HUMAN\|O75152_S132 | 0.631637198 | 4.84E-06 | 4.24E-04 |
| AFAD_HUMAN\|P55196_S1799 | 0.63161856 | 6.02E-04 | 6.52E-03 |
| PHF2_HUMAN\|O75151_S1058 | 0.631616011 | 5.22E-04 | 7.34E-03 |
| SENP3_HUMAN\|Q9H4L4_S169_S181_T176 | 0.631522413 | 4.25E-03 | 2.76E-02 |
| TENS4_HUMAN\|Q8IZW8_S242_S248_S253 | 0.631431879 | 7.58E-03 | 3.23E-02 |
| K1C16_HUMAN\|P08779_S446_S449 | 0.631343618 | 3.94E-04 | 2.82E-03 |
| SPT5H_HUMAN\|O00267_S666 | 0.631274786 | 9.97E-06 | 1.49E-04 |
| JIP4_HUMAN\|O60271_S815 | 0.631273002 | 4.15E-05 | 4.57E-04 |
| AFAD_HUMAN\|P55196_S1172_S1173_S1182 | 0.6312709 | 1.75E-05 | 8.76E-04 |
| RHG21_HUMAN\|Q5T5U3_S615 | 0.631223045 | 2.99E-05 | 1.27E-03 |
| HNRH3_HUMAN\|P31942_S298 | 0.631150525 | 1.32E-04 | 1.12E-03 |
| AR6P4_HUMAN\|Q66PJ3_S239 | 0.631129307 | 1.20E-04 | 1.05E-03 |
| NUCKS_HUMAN\|Q9H1E3_S75_S79 | 0.631127261 | 9.04E-03 | 4.79E-02 |
| HIRP3_HUMAN\|Q9BW71_S551 | 0.631080681 | 7.00E-03 | 3.43E-02 |
| PHLB1_HUMAN\|Q86UU1_S430 | 0.631057524 | 3.09E-04 | 3.65E-02 |
| HOME3_HUMAN\|Q9NSC5_S159 | 0.631049456 | 6.67E-03 | 3.30E-02 |
| HSPB1_HUMAN\|P04792_S82 | 0.631017583 | 7.86E-06 | 1.90E-04 |
| ATX2L_HUMAN\|Q8WWM7_S339 | 0.630903968 | 7.78E-07 | 2.02E-05 |
| TB182_HUMAN\|Q9C0C2_S1652 | 0.630883168 | 1.46E-04 | 4.09E-03 |
| ARHGB_HUMAN\|O15085_T668_T672 | 0.630719884 | 2.79E-05 | 3.31E-04 |
| AHNK_HUMAN\|Q09666_S4092 | 0.630679901 | 3.56E-03 | 2.43E-02 |
| RABL6_HUMAN\|Q3YEC7_S425_S427 | 0.630641571 | 9.47E-06 | 1.45E-04 |
| LAD1_HUMAN\|O00515_S485 | 0.630623974 | 5.07E-04 | 1.90E-02 |
| K2C5_HUMAN\|P13647_S77_S82 | 0.630514471 | 6.81E-03 | 3.89E-02 |
| LIMA1_HUMAN\|Q9UHB6_Y229 | 0.630488506 | 2.41E-03 | 1.57E-02 |
| MISP_HUMAN\|Q8IVT2_S376 | 0.63046885 | 1.92E-04 | 1.61E-03 |
| ACINU_HUMAN\|Q9UKV3_S240 | 0.630429663 | 2.56E-04 | 5.64E-03 |
| PEAK1_HUMAN\|Q9H792_S1036 | 0.630406208 | 8.89E-06 | 2.83E-04 |
| PTK6_HUMAN\|Q13882_Y447 | 0.630344856 | 4.19E-05 | 8.66E-04 |
| ERCC5_HUMAN\|P28715_S382_S384 | 0.630329099 | 1.72E-03 | 2.10E-02 |
| PRC2A_HUMAN\|P48634_S1147 | 0.630227442 | 1.64E-05 | 8.32E-04 |

| | | | |
|---|---|---|---|
| VIME_HUMAN\|P08670_S430 | 0.63021303 | 1.10E-04 | 1.81E-03 |
| AKAP2_HUMAN\|Q9Y2D5_S720 | 0.630195209 | 3.86E-05 | 4.31E-04 |
| MAP1B_HUMAN\|P46821_S1792 | 0.630143511 | 5.35E-04 | 9.96E-03 |
| SRGP1_HUMAN\|Q7Z6B7_S906 | 0.630129422 | 2.07E-04 | 2.91E-03 |
| IWS1_HUMAN\|Q96ST2_S720_T725 | 0.630051554 | 8.41E-04 | 5.29E-03 |
| CLD4_HUMAN\|O14493_Y208 | 0.62998966 | 4.73E-03 | 1.92E-02 |
| CAVN3_HUMAN\|Q969G5_T205 | 0.629955267 | 7.26E-04 | 7.38E-03 |
| COHA1_HUMAN\|Q9UMD9_S61 | 0.629864989 | 1.69E-03 | 8.41E-03 |
| KPCD2_HUMAN\|Q9BZL6_S710 | 0.629768268 | 8.66E-03 | 3.58E-02 |
| MISP_HUMAN\|Q8IVT2_S575 | 0.629683041 | 1.87E-04 | 1.12E-02 |
| OSBP1_HUMAN\|P22059_S379_S382_S386 | 0.629678778 | 1.81E-03 | 1.94E-02 |
| IF4G2_HUMAN\|P78344_T508 | 0.629677319 | 3.17E-06 | 1.29E-04 |
| ACINU_HUMAN\|Q9UKV3_S240 | 0.629643721 | 2.49E-04 | 1.27E-02 |
| MET_HUMAN\|P08581_S997_T993 | 0.629633986 | 1.27E-03 | 1.47E-02 |
| AKP13_HUMAN\|Q12802_S2728 | 0.62960645 | 8.28E-06 | 2.69E-04 |
| ARG28_HUMAN\|Q8N1W1_S513 | 0.629396949 | 1.32E-04 | 2.00E-03 |
| PUM1_HUMAN\|Q14671_S709 | 0.629379784 | 8.49E-06 | 1.33E-04 |
| K2C1_HUMAN\|P04264_S66 | 0.629298814 | 5.19E-03 | 2.35E-02 |
| COBL1_HUMAN\|Q53SF7_S294 | 0.629289199 | 1.40E-05 | 1.92E-04 |
| SYNE2_HUMAN\|Q8WXH0_T2876 | 0.629272279 | 2.79E-03 | 2.62E-02 |
| NUP98_HUMAN\|P52948_S623 | 0.629269005 | 5.35E-04 | 7.48E-03 |
| CAMP2_HUMAN\|Q08AD1_S1313 | 0.629188028 | 1.63E-04 | 4.50E-03 |
| RB6I2_HUMAN\|Q8IUD2_S17_S21 | 0.629155582 | 7.22E-05 | 1.27E-03 |
| ARHGB_HUMAN\|O15085_T668_T672 | 0.629150991 | 2.59E-05 | 6.03E-04 |
| STK10_HUMAN\|O94804_S450_S454 | 0.629146685 | 4.86E-04 | 5.46E-03 |
| SSH1_HUMAN\|Q8WYL5_S937 | 0.629021076 | 2.40E-05 | 7.42E-04 |
| CAP1_HUMAN\|Q01518_S308 | 0.628970528 | 4.53E-05 | 1.77E-03 |
| ABLM1_HUMAN\|O14639_S655 | 0.628955168 | 1.20E-04 | 1.11E-03 |
| AHNK_HUMAN\|Q09666_S5752 | 0.628920106 | 6.21E-06 | 4.68E-04 |
| DLG1_HUMAN\|Q12959_S573 | 0.628892716 | 3.83E-03 | 1.83E-02 |
| HNRPC_HUMAN\|P07910_S239 | 0.62885819 | 4.62E-05 | 4.93E-04 |

| | | | |
|---|---|---|---|
| UBXN7_HUMAN\|O94888_S285_S288 | 0.62876423 | 1.96E-03 | 4.29E-02 |
| AHNK_HUMAN\|Q09666_S5110 | 0.628726168 | 2.28E-04 | 2.55E-03 |
| IASPP_HUMAN\|Q8WUF5_S335 | 0.628697752 | 3.52E-05 | 3.94E-04 |
| FRYL_HUMAN\|O94915_T1959 | 0.628639095 | 7.22E-05 | 1.27E-03 |
| DAPK3_HUMAN\|O43293_S311_S312 | 0.628581923 | 5.90E-05 | 2.09E-03 |
| AKAP1_HUMAN\|Q92667_S151 | 0.628581234 | 1.25E-03 | 9.55E-03 |
| STX7_HUMAN\|O15400_S131 | 0.628572428 | 2.56E-05 | 5.99E-04 |
| 4EBP1_HUMAN\|Q13541_T70 | 0.628538436 | 1.60E-04 | 2.38E-03 |
| AR6P4_HUMAN\|Q66PJ3_S239 | 0.628465591 | 1.07E-04 | 3.41E-03 |
| NUFP2_HUMAN\|Q7Z417_S212_S214 | 0.628444289 | 2.07E-04 | 5.17E-03 |
| ARG28_HUMAN\|Q8N1W1_S558 | 0.628407444 | 1.43E-03 | 2.03E-02 |
| SSH2_HUMAN\|Q76I76_S487 | 0.628377558 | 1.72E-06 | 3.74E-05 |
| RBL2_HUMAN\|Q08999_S966 | 0.628335922 | 5.10E-03 | 2.02E-02 |
| DLG1_HUMAN\|Q12959_S571 | 0.628327279 | 3.52E-04 | 2.57E-03 |
| NDE1_HUMAN\|Q9NXR1_S282 | 0.628280138 | 4.17E-03 | 4.34E-02 |
| CING_HUMAN\|Q9P2M7_S129 | 0.62824319 | 1.82E-03 | 8.91E-03 |
| K1C14_HUMAN\|P02533_S435_S438 | 0.628226107 | 1.07E-03 | 6.43E-03 |
| K2C5_HUMAN\|P13647_S21_T24 | 0.628179049 | 1.28E-03 | 1.78E-02 |
| AKAP2_HUMAN\|Q9Y2D5_S720 | 0.628142295 | 3.51E-05 | 1.00E-02 |
| KLC4_HUMAN\|Q9NSK0_S611 | 0.628079008 | 1.72E-04 | 1.36E-03 |
| NDRG1_HUMAN\|Q92597_S330_T328 | 0.628072237 | 1.68E-04 | 2.45E-03 |
| DBNL_HUMAN\|Q9UJU6_T271 | 0.628034944 | 1.29E-04 | 1.17E-03 |
| FOXK2_HUMAN\|Q01167_S398 | 0.627942239 | 2.16E-03 | 2.23E-02 |
| NU214_HUMAN\|P35658_S986_S988 | 0.62783816 | 1.32E-03 | 7.61E-03 |
| CHSP1_HUMAN\|Q9Y2V2_S52 | 0.627710354 | 9.26E-04 | 1.98E-02 |
| KS6A3_HUMAN\|P51812_S715 | 0.62769338 | 2.75E-06 | 2.33E-04 |
| SI1L1_HUMAN\|O43166_S1255 | 0.627671501 | 7.18E-04 | 1.25E-02 |
| MAP7_HUMAN\|Q14244_S209 | 0.627669006 | 8.74E-04 | 1.42E-02 |
| NELFE_HUMAN\|P18615_S353 | 0.627651821 | 1.49E-06 | 1.84E-04 |
| SRSF6_HUMAN\|Q13247_S303 | 0.627541846 | 4.54E-04 | 7.17E-03 |
| IP3KB_HUMAN\|P27987_S43 | 0.627411091 | 4.67E-05 | 7.19E-04 |

| | | | |
|---|---|---|---|
| MA7D1_HUMAN\|Q3KQU3_S254 | 0.627400183 | 1.77E-05 | 8.78E-04 |
| RFIP5_HUMAN\|Q9BXF6_S296 | 0.627360334 | 2.42E-06 | 5.16E-05 |
| PDLI2_HUMAN\|Q96JY6_S137_S143 | 0.627356749 | 1.12E-04 | 3.49E-03 |
| RPRD2_HUMAN\|Q5VT52_T723 | 0.627356691 | 5.00E-06 | 8.88E-05 |
| SEPT9_HUMAN\|Q9UHD8_T38_T42 | 0.627328159 | 3.27E-05 | 1.10E-03 |
| AHNK_HUMAN\|Q09666_S5752 | 0.627294955 | 5.70E-06 | 9.78E-05 |
| LARP4_HUMAN\|Q71RC2_S597 | 0.627146873 | 8.92E-05 | 3.02E-03 |
| ATX2L_HUMAN\|Q8WWM7_S680_S684 | 0.627020839 | 4.41E-06 | 1.63E-04 |
| K1C16_HUMAN\|P08779_S447_S450 | 0.627018328 | 2.44E-04 | 1.82E-03 |
| NIBL1_HUMAN\|Q96TA1_S692 | 0.626987706 | 5.67E-05 | 5.39E-03 |
| P3C2A_HUMAN\|O00443_S259 | 0.626927228 | 5.68E-05 | 8.60E-04 |
| GGA2_HUMAN\|Q9UJY4_S177 | 0.626889333 | 5.27E-04 | 3.31E-03 |
| MA7D1_HUMAN\|Q3KQU3_S366 | 0.626878227 | 3.47E-06 | 1.37E-04 |
| BORG5_HUMAN\|Q00587_S303 | 0.626872875 | 5.14E-05 | 5.34E-04 |
| PP14A_HUMAN\|Q96A00_S26 | 0.626839966 | 1.60E-05 | 9.44E-04 |
| RL15_HUMAN\|P61313_S34 | 0.626749206 | 1.42E-04 | 1.19E-03 |
| SRC8_HUMAN\|Q14247_T399_T411 | 0.626729478 | 6.63E-05 | 9.43E-04 |
| RCC1_HUMAN\|P18754_S11 | 0.626723326 | 3.12E-03 | 1.55E-02 |
| PKN2_HUMAN\|Q16513_S362 | 0.626713804 | 1.77E-04 | 4.64E-03 |
| LAD1_HUMAN\|O00515_S301 | 0.626690207 | 6.08E-04 | 8.96E-03 |
| VIME_HUMAN\|P08670_S430 | 0.626599138 | 9.42E-05 | 3.15E-03 |
| IF6_HUMAN\|P56537_S243 | 0.626595831 | 6.02E-03 | 2.29E-02 |
| AHNK_HUMAN\|Q09666_S2397 | 0.626574206 | 6.87E-04 | 1.14E-02 |
| SRRM1_HUMAN\|Q8IYB3_S740_S743 | 0.626550891 | 2.27E-04 | 1.83E-03 |
| LIPA3_HUMAN\|O75145_S1164 | 0.626485567 | 5.92E-05 | 8.77E-04 |
| ARHGB_HUMAN\|O15085_S1457_T1462 | 0.62641696 | 5.92E-04 | 5.42E-03 |
| ITB4_HUMAN\|P16144_S1457 | 0.626415692 | 4.83E-04 | 3.36E-03 |
| SEPT9_HUMAN\|Q9UHD8_T38_T42 | 0.626386742 | 3.14E-05 | 1.51E-03 |
| MARCS_HUMAN\|P29966_T150 | 0.626383311 | 5.87E-04 | 1.09E-02 |
| AKP13_HUMAN\|Q12802_S2728 | 0.626381067 | 7.04E-06 | 5.61E-04 |
| LARP1_HUMAN\|Q6PKG0_S517_S521 | 0.626356363 | 5.84E-06 | 9.84E-05 |

| | | | |
|---|---|---|---|
| PHLB1_HUMAN\|Q86UU1_S430 | 0.626330738 | 2.60E-04 | 1.90E-03 |
| NDRG1_HUMAN\|Q92597_S330_S336 | 0.626202184 | 1.10E-03 | 1.01E-02 |
| RBM7_HUMAN\|Q9Y580_S137 | 0.626170178 | 1.44E-05 | 1.92E-04 |
| TOM20_HUMAN\|Q15388_S138 | 0.626131517 | 2.56E-03 | 4.03E-02 |
| BASP1_HUMAN\|P80723_T196 | 0.626089566 | 2.55E-04 | 5.64E-03 |
| HELZ_HUMAN\|P42694_T1163 | 0.626068596 | 6.05E-07 | 1.67E-05 |
| SRGP3_HUMAN\|O43295_S858 | 0.626049044 | 1.51E-04 | 2.23E-03 |
| ATX2L_HUMAN\|Q8WWM7_S339 | 0.625993454 | 5.80E-07 | 3.31E-05 |
| ATAT_HUMAN\|Q5SQI0_S315 | 0.625930091 | 4.68E-06 | 4.16E-04 |
| SH24A_HUMAN\|Q9H788_S316 | 0.625910217 | 4.74E-04 | 4.50E-03 |
| MA7D1_HUMAN\|Q3KQU3_S254 | 0.625812747 | 1.63E-05 | 9.52E-04 |
| 4EBP1_HUMAN\|Q13541_T70 | 0.625777002 | 1.44E-04 | 8.29E-03 |
| PRR12_HUMAN\|Q9ULL5_S44 | 0.625747606 | 1.43E-04 | 1.20E-03 |
| F193A_HUMAN\|P78312_S1144 | 0.625747437 | 1.00E-04 | 2.93E-03 |
| KPCD2_HUMAN\|Q9BZL6_S212 | 0.625696503 | 3.38E-04 | 2.34E-03 |
| LMNA_HUMAN\|P02545_S303 | 0.625688366 | 1.78E-03 | 9.71E-03 |
| P4K2A_HUMAN\|Q9BTU6_S468 | 0.625668705 | 1.38E-03 | 1.87E-02 |
| SLK_HUMAN\|Q9H2G2_S779 | 0.625643698 | 5.13E-03 | 3.14E-02 |
| MK01_HUMAN\|P28482_T185_Y187 | 0.625591903 | 7.61E-04 | 1.28E-02 |
| TBCD4_HUMAN\|O60343_S318 | 0.625580056 | 7.02E-05 | 2.20E-03 |
| K1C14_HUMAN\|P02533_S51_S54 | 0.625563693 | 9.28E-04 | 9.35E-03 |
| OPTN_HUMAN\|Q96CV9_S177 | 0.625523389 | 1.04E-03 | 5.70E-03 |
| SF3B1_HUMAN\|O75533_S194 | 0.62535646 | 1.92E-04 | 1.61E-03 |
| SPTB2_HUMAN\|Q01082_S2341 | 0.625349893 | 7.26E-07 | 2.01E-05 |
| TB182_HUMAN\|Q9C0C2_S1715 | 0.625186477 | 7.13E-04 | 4.64E-03 |
| NDRG1_HUMAN\|Q92597_S326 | 0.625108147 | 1.13E-02 | 3.81E-02 |
| RECQ4_HUMAN\|O94761_S251 | 0.625081389 | 3.32E-04 | 2.30E-03 |
| SAFB1_HUMAN\|Q15424_S247 | 0.625069761 | 3.79E-05 | 1.60E-03 |
| AHNK_HUMAN\|Q09666_S212 | 0.625046077 | 5.44E-04 | 5.08E-03 |
| HABP4_HUMAN\|Q5JVS0_S108 | 0.625027269 | 1.26E-05 | 1.78E-04 |
| ARFG1_HUMAN\|Q8N6T3_T135 | 0.624732167 | 4.47E-05 | 1.77E-03 |

| | | | |
|---|---|---|---|
| SON_HUMAN\|P18583_S94 | 0.624683009 | 6.70E-06 | 1.69E-04 |
| ENOA_HUMAN\|P06733_S40 | 0.624673284 | 7.82E-04 | 8.12E-03 |
| SRPK2_HUMAN\|P78362_S494_S497 | 0.624632611 | 7.14E-05 | 6.24E-03 |
| TBC9B_HUMAN\|Q66K14_S411 | 0.624601998 | 9.22E-04 | 1.47E-02 |
| SRRM1_HUMAN\|Q8IYB3_S740_S743 | 0.624508209 | 2.10E-04 | 5.17E-03 |
| SLBP_HUMAN\|Q14493_S20 | 0.624287712 | 2.38E-04 | 1.25E-02 |
| STK10_HUMAN\|O94804_S448_S455 | 0.62427639 | 3.64E-05 | 7.74E-04 |
| PEAK1_HUMAN\|Q9H792_S1036 | 0.62426952 | 6.42E-06 | 1.07E-04 |
| OSB10_HUMAN\|Q9BXB5_S26_S30 | 0.624265985 | 1.19E-03 | 1.46E-02 |
| MED24_HUMAN\|O75448_S873 | 0.62422616 | 3.05E-03 | 1.35E-02 |
| K1C14_HUMAN\|P02533_S33 | 0.624198944 | 1.62E-04 | 1.31E-03 |
| DEFI6_HUMAN\|Q9H4E7_S590_S597_T595 | 0.624189266 | 3.07E-04 | 3.65E-02 |
| SHRM1_HUMAN\|Q2M3G4_S40 | 0.624131564 | 6.38E-03 | 2.79E-02 |
| MICA3_HUMAN\|Q7RTP6_S977 | 0.624088164 | 4.12E-07 | 5.73E-05 |
| SPTB2_HUMAN\|Q01082-3_S10 | 0.624068282 | 1.44E-04 | 1.29E-03 |
| SEPT7_HUMAN\|Q16181_S423 | 0.623993109 | 1.61E-03 | 2.95E-02 |
| CC120_HUMAN\|Q96HB5_S296 | 0.623989961 | 6.00E-03 | 4.53E-02 |
| RIN1_HUMAN\|Q13671_S349 | 0.62388298 | 1.43E-03 | 1.60E-02 |
| SNX17_HUMAN\|Q15036_S336 | 0.623866336 | 4.06E-03 | 2.66E-02 |
| CTNA1_HUMAN\|P35221_S641_S652 | 0.623812837 | 1.70E-04 | 1.47E-03 |
| PKP2_HUMAN\|Q99959_S132 | 0.623806423 | 6.12E-04 | 6.53E-03 |
| LAD1_HUMAN\|O00515_S123 | 0.623800973 | 4.38E-03 | 2.96E-02 |
| PP1R7_HUMAN\|Q15435_S24 | 0.623758833 | 1.62E-03 | 2.14E-02 |
| DC1L2_HUMAN\|O43237_S407 | 0.623716849 | 7.00E-05 | 6.75E-04 |
| PFD4_HUMAN\|Q9NQP4_S125 | 0.623627401 | 7.07E-04 | 4.15E-03 |
| LMNA_HUMAN\|P02545_S406_S407 | 0.623603877 | 1.25E-04 | 1.15E-03 |
| CO052_HUMAN\|Q6ZUT6_S55 | 0.623562646 | 4.55E-03 | 2.11E-02 |
| REPS1_HUMAN\|Q96D71_S162_S170_S174 | 0.62345345 | 6.54E-05 | 2.27E-03 |
| AKAP1_HUMAN\|Q92667_S151 | 0.623429488 | 1.06E-03 | 2.92E-02 |
| MGMT_HUMAN\|P16455_S201 | 0.623428174 | 5.57E-07 | 7.36E-05 |
| INT1_HUMAN\|Q8N201_T83 | 0.623325258 | 2.41E-03 | 1.79E-02 |

| | | | |
|---|---|---|---|
| PHLB2_HUMAN\|Q86SQ0_T404 | 0.623287947 | 2.44E-04 | 1.93E-03 |
| RBP56_HUMAN\|Q92804_S226 | 0.623287831 | 1.05E-02 | 4.71E-02 |
| WAC2A_HUMAN\|Q641Q2_S390 | 0.623266951 | 7.08E-04 | 4.62E-03 |
| UBP24_HUMAN\|Q9UPU5_S1141 | 0.62323115 | 9.47E-05 | 8.76E-04 |
| SH3K1_HUMAN\|Q96B97_S587 | 0.623190234 | 4.41E-05 | 8.89E-04 |
| UBE4B_HUMAN\|O95155_S84_S88 | 0.623141194 | 2.37E-03 | 1.10E-02 |
| MK01_HUMAN\|P28482_T185_Y187 | 0.62312247 | 7.03E-04 | 4.59E-03 |
| ITB4_HUMAN\|P16144_T1455 | 0.623052473 | 5.75E-04 | 3.89E-03 |
| ASPP1_HUMAN\|Q96KQ4_S477 | 0.622964897 | 4.76E-03 | 3.00E-02 |
| IASPP_HUMAN\|Q8WUF5_S134 | 0.622776454 | 4.31E-04 | 2.81E-03 |
| NDRG1_HUMAN\|Q92597_S330_S333_S336 | 0.622733074 | 4.49E-03 | 4.38E-02 |
| K1C14_HUMAN\|P02533_S435_S437 | 0.62270521 | 1.40E-03 | 7.98E-03 |
| HES1_HUMAN\|Q14469_S10 | 0.622693609 | 5.13E-03 | 2.03E-02 |
| PRC2C_HUMAN\|Q9Y520_S2694_T2691 | 0.622668723 | 8.76E-04 | 4.95E-03 |
| SRSF6_HUMAN\|Q13247_S301 | 0.622653599 | 3.11E-04 | 5.49E-03 |
| COHA1_HUMAN\|Q9UMD9_S78 | 0.622585449 | 2.73E-03 | 1.24E-02 |
| VIME_HUMAN\|P08670_S55 | 0.622524764 | 9.94E-04 | 9.42E-03 |
| SLAI2_HUMAN\|Q9P270_S349_S353_S357 | 0.622468494 | 2.10E-05 | 5.22E-04 |
| CLIP1_HUMAN\|P30622_S200 | 0.622436772 | 1.29E-03 | 1.20E-02 |
| LMNA_HUMAN\|P02545_S407_T409 | 0.622393519 | 5.51E-04 | 3.42E-03 |
| DC1L2_HUMAN\|O43237_S383 | 0.622378078 | 1.04E-04 | 2.90E-03 |
| NFIC_HUMAN\|P08651_S473 | 0.62232306 | 2.82E-03 | 3.25E-02 |
| AHNK_HUMAN\|Q09666_S212_S216 | 0.622311447 | 1.96E-05 | 1.09E-03 |
| CING_HUMAN\|Q9P2M7_S129_S134 | 0.622241516 | 3.34E-04 | 2.31E-03 |
| TGS1_HUMAN\|Q96RS0_S89 | 0.621998226 | 2.75E-04 | 2.10E-03 |
| LAR4B_HUMAN\|Q92615_T732 | 0.621995756 | 1.73E-06 | 3.74E-05 |
| HNRPC_HUMAN\|P07910_S240 | 0.621991072 | 1.59E-04 | 1.40E-03 |
| PARD3_HUMAN\|Q8TEW0_S187 | 0.621897534 | 1.14E-02 | 3.82E-02 |
| PP12C_HUMAN\|Q9BZL4_S453 | 0.621804664 | 1.37E-06 | 7.47E-05 |
| THTM_HUMAN\|P25325-2_S15 | 0.621777038 | 1.73E-03 | 1.47E-02 |
| SF3A3_HUMAN\|Q12874_S365_S367_S369 | 0.621669144 | 1.49E-02 | 4.75E-02 |

| | | | |
|---|---|---|---|
| K1C16_HUMAN\|P08779_S468 | 0.62164253 | 9.66E-05 | 9.37E-04 |
| CLAP1_HUMAN\|Q7Z460_Y634 | 0.621592772 | 5.90E-03 | 3.67E-02 |
| SCRIB_HUMAN\|Q14160_S1348 | 0.621475932 | 1.59E-03 | 8.84E-03 |
| MTUS1_HUMAN\|Q9ULD2_S1264_S1268 | 0.621468348 | 2.95E-04 | 3.78E-03 |
| BACD2_HUMAN\|Q13829_S280 | 0.621414824 | 6.40E-04 | 4.25E-03 |
| BAIP2_HUMAN\|Q9UQB8_T340 | 0.62139395 | 3.43E-03 | 1.68E-02 |
| ANK3_HUMAN\|Q12955_S959 | 0.621384643 | 3.07E-03 | 3.30E-02 |
| CBX4_HUMAN\|O00257_S182 | 0.621259096 | 3.78E-04 | 2.55E-03 |
| TM1L1_HUMAN\|O75674_S321 | 0.621250401 | 1.47E-03 | 1.87E-02 |
| LYAR_HUMAN\|Q9NX58_S276 | 0.621235504 | 7.72E-06 | 1.88E-04 |
| GBF1_HUMAN\|Q92538_S1781_S1784 | 0.621142416 | 2.73E-04 | 2.94E-03 |
| 3BP1_HUMAN\|Q9Y3L3_S695 | 0.621113073 | 2.39E-04 | 7.70E-03 |
| TB182_HUMAN\|Q9C0C2_S1715 | 0.621084351 | 6.22E-04 | 1.08E-02 |
| SGTB_HUMAN\|Q96EQ0_S297 | 0.620984841 | 5.07E-05 | 2.27E-03 |
| DC1L2_HUMAN\|O43237_S383 | 0.620957455 | 9.87E-05 | 9.05E-04 |
| DC1L1_HUMAN\|Q9Y6G9_S421 | 0.620906299 | 4.03E-03 | 1.69E-02 |
| ROA3_HUMAN\|P51991_S356 | 0.620851861 | 1.87E-05 | 2.39E-04 |
| TMF1_HUMAN\|P82094_S338_S344 | 0.620843469 | 1.22E-05 | 1.73E-04 |
| IRS2_HUMAN\|Q9Y4H2_S620 | 0.620685845 | 3.29E-04 | 2.29E-03 |
| DESP_HUMAN\|P15924_S2820_S2821 | 0.620659565 | 2.01E-04 | 1.67E-03 |
| K1C17_HUMAN\|Q04695_S11_S12 | 0.620654741 | 5.70E-04 | 6.24E-03 |
| ML12A_HUMAN\|P19105_S19 | 0.620636171 | 4.10E-05 | 1.95E-03 |
| DAPK3_HUMAN\|O43293_S311_S312 | 0.620515231 | 4.22E-05 | 4.53E-04 |
| ABLM1_HUMAN\|O14639_S431_T429 | 0.620352053 | 2.18E-05 | 5.35E-04 |
| PP14A_HUMAN\|Q96A00_S26 | 0.620322004 | 1.16E-05 | 2.03E-03 |
| TENS4_HUMAN\|Q8IZW8_S248_S253 | 0.620139143 | 2.12E-03 | 2.42E-02 |
| NRK_HUMAN\|Q7Z2Y5_S854 | 0.620078246 | 1.87E-06 | 3.95E-05 |
| PLEC_HUMAN\|Q15149_S4382_S4396 | 0.620076404 | 8.64E-05 | 2.62E-03 |
| PININ_HUMAN\|Q9H307_S690 | 0.620076167 | 9.55E-04 | 1.23E-02 |
| K1C17_HUMAN\|Q04695_S17 | 0.620012312 | 2.20E-03 | 1.16E-02 |
| T184B_HUMAN\|Q9Y519_S403 | 0.619992286 | 2.22E-04 | 5.31E-03 |

| | | | |
|---|---|---|---|
| K1143_HUMAN\|Q96AT1_S146 | 0.619964618 | 1.05E-03 | 1.59E-02 |
| FKB15_HUMAN\|Q5T1M5_S344 | 0.619937267 | 7.88E-05 | 1.35E-03 |
| LMNB1_HUMAN\|P20700_T20 | 0.619907709 | 1.38E-03 | 1.03E-02 |
| CLAP2_HUMAN\|O75122_S463 | 0.619825394 | 2.22E-03 | 2.24E-02 |
| VIME_HUMAN\|P08670_S55 | 0.619784729 | 9.11E-04 | 1.44E-02 |
| SPTB2_HUMAN\|Q01082_S2341 | 0.619783589 | 5.39E-07 | 1.04E-04 |
| GPTC2_HUMAN\|Q9NW75_S115_S117 | 0.61972249 | 4.34E-06 | 7.95E-05 |
| S6A15_HUMAN\|Q9H2J7_S699 | 0.619682638 | 2.56E-03 | 4.03E-02 |
| M3K3_HUMAN\|Q99759_S337 | 0.619656877 | 4.80E-04 | 4.55E-03 |
| K1143_HUMAN\|Q96AT1_S146 | 0.619577611 | 1.03E-03 | 1.60E-02 |
| CREB1_HUMAN\|P16220_S271 | 0.619433501 | 8.61E-04 | 1.89E-02 |
| RCC1_HUMAN\|P18754_S11 | 0.619430766 | 2.57E-03 | 4.90E-02 |
| NDRG1_HUMAN\|Q92597_S330_S336 | 0.619409669 | 8.86E-04 | 5.53E-03 |
| CT2NL_HUMAN\|Q9P2B4_Y447 | 0.619351426 | 1.12E-03 | 3.17E-02 |
| PDLI5_HUMAN\|Q96HC4_S111 | 0.61931018 | 7.12E-05 | 1.31E-03 |
| PAK2_HUMAN\|Q13177_Y139 | 0.619301229 | 5.57E-04 | 3.44E-03 |
| MISP_HUMAN\|Q8IVT2_S400 | 0.61926621 | 8.04E-04 | 4.62E-03 |
| PKP4_HUMAN\|Q99569_S233 | 0.619161767 | 3.11E-05 | 3.65E-04 |
| FXR2_HUMAN\|P51116_S453 | 0.619041655 | 1.02E-04 | 9.22E-04 |
| K1C17_HUMAN\|Q04695_S11 | 0.619031627 | 1.39E-04 | 1.25E-03 |
| PAXI_HUMAN\|P49023_S106 | 0.618987544 | 1.82E-04 | 4.38E-03 |
| AKAP2_HUMAN\|Q9Y2D5_S473 | 0.618971904 | 2.04E-03 | 1.09E-02 |
| SPR2B_HUMAN\|P35325_S60 | 0.618848984 | 9.33E-03 | 3.82E-02 |
| RFIP1_HUMAN\|Q6WKZ4_S501 | 0.61876097 | 1.87E-03 | 9.05E-03 |
| LARP6_HUMAN\|Q9BRS8_S56_S58 | 0.618726313 | 9.06E-05 | 2.67E-03 |
| PLXB2_HUMAN\|O15031_S1570 | 0.618683761 | 4.76E-04 | 9.45E-03 |
| RTKN_HUMAN\|Q9BST9_S543 | 0.618668874 | 2.69E-05 | 1.22E-03 |
| AHNK_HUMAN\|Q09666_S212_S216 | 0.61861061 | 1.65E-05 | 4.44E-04 |
| MAP4_HUMAN\|P27816_S1073 | 0.618571152 | 6.31E-05 | 1.14E-03 |
| PCIF1_HUMAN\|Q9H4Z3_T150 | 0.618522914 | 9.93E-04 | 5.49E-03 |
| SRRM2_HUMAN\|Q9UQ35_S2044_S2046 | 0.618403858 | 2.79E-04 | 3.65E-03 |

| | | | |
|---|---|---|---|
| ZDHC5_HUMAN\|Q9C0B5_S621 | 0.618399858 | 1.50E-03 | 1.10E-02 |
| R3HD1_HUMAN\|Q15032_S973 | 0.618389641 | 7.02E-06 | 1.12E-03 |
| UBP2L_HUMAN\|Q14157_S609 | 0.618377687 | 1.52E-04 | 2.91E-03 |
| SPTB2_HUMAN\|Q01082-3_S14_S8 | 0.618325061 | 4.13E-05 | 6.60E-04 |
| RAI14_HUMAN\|Q9P0K7_S281_S290 | 0.618313088 | 3.44E-03 | 3.14E-02 |
| LMNA_HUMAN\|P02545_S404_S407 | 0.618244904 | 7.82E-05 | 7.39E-04 |
| 3BP1_HUMAN\|Q9Y3L3_S695 | 0.618164627 | 2.15E-04 | 1.76E-03 |
| NSRP1_HUMAN\|Q9H0G5_T275 | 0.618145605 | 2.45E-04 | 1.82E-03 |
| UBP2L_HUMAN\|Q14157_S470_S477 | 0.618112893 | 5.90E-04 | 1.05E-02 |
| DOCK6_HUMAN\|Q96HP0_S880 | 0.617979996 | 3.88E-04 | 7.60E-03 |
| NUCKS_HUMAN\|Q9H1E3_S181_T179 | 0.617899978 | 3.70E-03 | 3.30E-02 |
| COIL_HUMAN\|P38432_S566_S571 | 0.617884916 | 1.01E-04 | 1.73E-03 |
| RTKN_HUMAN\|Q9BST9_S529 | 0.6178618 | 1.73E-04 | 4.23E-03 |
| KLC3_HUMAN\|Q6P597_S495_S502 | 0.617823598 | 1.97E-03 | 3.43E-02 |
| IF4B_HUMAN\|P23588_S497_S504_T500 | 0.617746518 | 2.77E-04 | 8.67E-03 |
| HIRP3_HUMAN\|Q9BW71_S196_S199 | 0.617703582 | 2.94E-05 | 3.47E-04 |
| RFIP5_HUMAN\|Q9BXF6_S296 | 0.617693986 | 1.39E-06 | 7.47E-05 |
| SPTB2_HUMAN\|Q01082-3_S8 | 0.617690142 | 1.90E-04 | 1.60E-03 |
| DESP_HUMAN\|P15924_S22 | 0.617664224 | 1.12E-03 | 1.03E-02 |
| F122B_HUMAN\|Q7Z309_S115_S119 | 0.617636206 | 1.46E-05 | 4.12E-04 |
| 1B07_HUMAN\|P01889_S359 | 0.617493213 | 4.47E-05 | 4.85E-04 |
| ABLM1_HUMAN\|O14639_S458 | 0.617479669 | 1.68E-03 | 8.36E-03 |
| EDC3_HUMAN\|Q96F86_S131 | 0.617360186 | 2.48E-06 | 5.02E-05 |
| SPTN1_HUMAN\|Q13813_S1190 | 0.617274467 | 1.14E-05 | 3.39E-04 |
| TFE3_HUMAN\|P19532_S548_S556 | 0.617012114 | 2.03E-04 | 3.71E-03 |
| UBE4B_HUMAN\|O95155_S103 | 0.616997642 | 1.13E-03 | 1.59E-02 |
| EXOC7_HUMAN\|Q9UPT5_S250 | 0.616983245 | 5.04E-05 | 9.87E-04 |
| DC1L2_HUMAN\|O43237_S407 | 0.616947893 | 5.32E-05 | 1.81E-03 |
| SKT_HUMAN\|Q5T5P2_S169 | 0.61691233 | 1.31E-04 | 1.12E-03 |
| ARHG2_HUMAN\|Q92974_S886 | 0.616775595 | 1.68E-03 | 3.03E-02 |
| BICD2_HUMAN\|Q8TD16_S608_S615 | 0.616728001 | 6.17E-03 | 3.77E-02 |

| | | | |
|---|---|---|---|
| TAOK3_HUMAN\|Q9H2K8_S442 | 0.616718528 | 3.42E-04 | 4.16E-03 |
| RABL6_HUMAN\|Q3YEC7_S492 | 0.616681576 | 1.22E-05 | 1.72E-04 |
| SPR2B_HUMAN\|P35325_S60 | 0.616660421 | 8.88E-03 | 3.15E-02 |
| CSN3_HUMAN\|Q9UNS2_S421 | 0.616582288 | 1.14E-06 | 4.50E-05 |
| TP53B_HUMAN\|Q12888_S359_S366 | 0.616432 | 8.38E-04 | 1.88E-02 |
| SRRM1_HUMAN\|Q8IYB3_T846 | 0.616414205 | 1.07E-03 | 5.82E-03 |
| BAD_HUMAN\|Q92934_S74 | 0.616396643 | 1.53E-04 | 3.95E-03 |
| PLEC_HUMAN\|Q15149_S4389_S4391 | 0.616369535 | 4.59E-04 | 7.21E-03 |
| KANK2_HUMAN\|Q63ZY3_S540 | 0.616212568 | 1.74E-03 | 8.57E-03 |
| PACN3_HUMAN\|Q9UKS6_S383 | 0.616212011 | 2.96E-03 | 1.49E-02 |
| 3BP1_HUMAN\|Q9Y3L3_T601 | 0.616138594 | 2.71E-03 | 1.23E-02 |
| DC1L1_HUMAN\|Q9Y6G9_T513_T515 | 0.616134247 | 4.39E-05 | 6.85E-04 |
| MRCKG_HUMAN\|Q6DT37_S1544 | 0.616116267 | 1.02E-03 | 8.12E-03 |
| PP14A_HUMAN\|Q96A00_S128_S134_S136 | 0.616001013 | 1.11E-05 | 3.24E-04 |
| PKP4_HUMAN\|Q99569_S233 | 0.615896507 | 2.70E-05 | 1.48E-03 |
| TP53B_HUMAN\|Q12888_S366_T357 | 0.615767741 | 1.27E-04 | 1.10E-03 |
| RABL6_HUMAN\|Q3YEC7_S492 | 0.615712916 | 1.17E-05 | 1.69E-04 |
| RALY_HUMAN\|Q9UKM9_S288_T286_T298 | 0.615664152 | 3.84E-04 | 2.58E-03 |
| NUCKS_HUMAN\|Q9H1E3_S75_S79 | 0.615651357 | 6.33E-03 | 2.38E-02 |
| PDS5A_HUMAN\|Q29RF7_S1276 | 0.615650744 | 3.90E-04 | 1.12E-02 |
| PTN1_HUMAN\|P18031_S50 | 0.615603324 | 4.80E-03 | 3.01E-02 |
| MAP7_HUMAN\|Q14244_S183 | 0.615600475 | 1.14E-05 | 3.39E-04 |
| AKTS1_HUMAN\|Q96B36_T246 | 0.615564776 | 6.41E-04 | 1.14E-02 |
| MAST4_HUMAN\|O15021_S2555 | 0.615556823 | 2.70E-04 | 5.83E-03 |
| KC1A_HUMAN\|P48729_S312 | 0.615551231 | 1.19E-02 | 4.64E-02 |
| XCT_HUMAN\|Q9UPY5_S26 | 0.615428707 | 1.03E-03 | 2.15E-02 |
| ARG28_HUMAN\|Q8N1W1_S558 | 0.615381263 | 9.50E-04 | 5.28E-03 |
| XRN2_HUMAN\|Q9H0D6_S448 | 0.615302296 | 1.37E-04 | 4.02E-03 |
| AHNK_HUMAN\|Q09666_T4100 | 0.615272573 | 8.30E-04 | 4.75E-03 |
| CCD86_HUMAN\|Q9H6F5_S113 | 0.615043674 | 4.51E-03 | 3.02E-02 |
| MILK1_HUMAN\|Q8N3F8_S578 | 0.614985901 | 8.60E-06 | 5.18E-04 |

| | | | |
|---|---|---|---|
| PHLB2_HUMAN\|Q86SQ0_S334_T328 | 0.61491879 | 1.46E-03 | 2.73E-02 |
| SPTN1_HUMAN\|Q13813_S1190 | 0.614818237 | 1.01E-05 | 7.16E-04 |
| DLG1_HUMAN\|Q12959_S571 | 0.61473702 | 2.09E-04 | 2.85E-03 |
| DDX10_HUMAN\|Q13206_S831 | 0.614603002 | 6.09E-05 | 8.91E-04 |
| SYMPK_HUMAN\|Q92797_S1173 | 0.614590541 | 3.80E-03 | 1.62E-02 |
| SARG_HUMAN\|Q9BW04_S502 | 0.61450173 | 7.08E-06 | 4.43E-04 |
| KLC4_HUMAN\|Q9NSK0_S611 | 0.614498599 | 1.02E-04 | 9.83E-04 |
| ARVC_HUMAN\|O00192_S203 | 0.614447951 | 7.85E-05 | 1.42E-03 |
| KPCD2_HUMAN\|Q9BZL6_S214 | 0.614443676 | 2.98E-04 | 9.29E-03 |
| LMNA_HUMAN\|P02545_S403_S406 | 0.614437696 | 1.63E-04 | 1.31E-03 |
| CLAP2_HUMAN\|O75122_S1029 | 0.614411573 | 2.18E-04 | 7.22E-03 |
| ZCHC8_HUMAN\|Q6NZY4_T479 | 0.61435388 | 2.77E-05 | 1.34E-03 |
| K1C14_HUMAN\|P02533_S11 | 0.614292928 | 1.95E-05 | 2.50E-04 |
| NDRG1_HUMAN\|Q92597_S330_T328 | 0.61417686 | 9.52E-05 | 1.53E-03 |
| PARP4_HUMAN\|Q9UKK3_S1722 | 0.614169965 | 6.29E-05 | 2.16E-03 |
| PLEC_HUMAN\|Q15149_S4386_S4396 | 0.614155796 | 4.54E-06 | 3.15E-04 |
| ABI2_HUMAN\|Q9NYB9_Y371 | 0.614151234 | 1.37E-05 | 3.83E-04 |
| UBXN7_HUMAN\|O94888_S285_S288 | 0.614109797 | 1.26E-03 | 2.47E-02 |
| SCRIB_HUMAN\|Q14160_S1475 | 0.614086642 | 1.50E-03 | 1.90E-02 |
| BASP1_HUMAN\|P80723_T196 | 0.614044446 | 1.58E-04 | 4.31E-03 |
| PHF8_HUMAN\|Q9UPP1_S804 | 0.614043128 | 1.40E-05 | 6.17E-04 |
| TB182_HUMAN\|Q9C0C2_S1029 | 0.614024431 | 1.75E-05 | 8.88E-04 |
| NGAP_HUMAN\|Q9UJF2_T620 | 0.613991341 | 6.46E-03 | 2.42E-02 |
| WAC2A_HUMAN\|Q641Q2_S284_S288 | 0.613909921 | 1.08E-04 | 4.15E-03 |
| CO052_HUMAN\|Q6ZUT6_S193 | 0.613871473 | 1.20E-04 | 1.11E-03 |
| LAP2A_HUMAN\|P42166_T74 | 0.613866133 | 6.96E-04 | 1.21E-02 |
| STMN1_HUMAN\|P16949_S25 | 0.613835382 | 1.53E-05 | 1.77E-03 |
| FSTL3_HUMAN\|O95633_S255 | 0.613815791 | 2.72E-04 | 3.58E-03 |
| 3BP5L_HUMAN\|Q7L8J4_S343 | 0.613810181 | 8.27E-06 | 1.29E-04 |
| MGMT_HUMAN\|P16455_S201 | 0.613771582 | 3.11E-07 | 6.48E-05 |
| BAIP2_HUMAN\|Q9UQB8_T360 | 0.613716219 | 7.35E-03 | 4.11E-02 |

| | | | |
|---|---|---|---|
| COHA1_HUMAN\|Q9UMD9_T387 | 0.613692258 | 3.81E-05 | 4.29E-04 |
| SPTB2_HUMAN\|Q01082-3_S14_S8 | 0.613684545 | 3.37E-05 | 3.37E-03 |
| K2C1_HUMAN\|P04264_S531 | 0.61365245 | 1.05E-02 | 3.60E-02 |
| CRBG1_HUMAN\|Q9Y4K1_S22 | 0.61364446 | 5.72E-06 | 2.78E-04 |
| DESP_HUMAN\|P15924_S166 | 0.613612649 | 3.90E-04 | 3.89E-03 |
| SRGP1_HUMAN\|Q7Z6B7_S906 | 0.613590623 | 1.11E-04 | 3.00E-03 |
| ARH40_HUMAN\|Q8TER5_T371 | 0.613570637 | 1.72E-06 | 3.97E-05 |
| AHNK_HUMAN\|Q09666_S511 | 0.613544717 | 3.70E-04 | 1.08E-02 |
| TCOF_HUMAN\|Q13428_T1270 | 0.613536337 | 3.88E-03 | 3.87E-02 |
| B4DKV4_HUMAN\|B4DKV4_S19 | 0.613505387 | 3.86E-03 | 1.84E-02 |
| LMNA_HUMAN\|P02545_S392_S395 | 0.613490422 | 6.35E-03 | 2.38E-02 |
| CRK_HUMAN\|P46108_S304 | 0.613462772 | 1.70E-06 | 1.51E-04 |
| TPPP_HUMAN\|O94811_S160 | 0.613259927 | 1.09E-03 | 1.66E-02 |
| TANK_HUMAN\|Q92844_S129 | 0.613175104 | 4.91E-04 | 9.12E-03 |
| TMF1_HUMAN\|P82094_S77 | 0.613156516 | 5.25E-05 | 4.25E-03 |
| K2C6B_HUMAN\|P04259_S546 | 0.613144475 | 4.06E-05 | 4.41E-04 |
| RBP10_HUMAN\|Q6VN20_S365 | 0.613058607 | 5.35E-06 | 3.81E-04 |
| PRC2A_HUMAN\|P48634_S761_S764 | 0.612853157 | 3.49E-06 | 2.91E-04 |
| MRP_HUMAN\|P49006_T178 | 0.612840283 | 9.11E-06 | 1.35E-04 |
| REPS1_HUMAN\|Q96D71_S162_S170_S174 | 0.61283688 | 4.20E-05 | 8.53E-04 |
| RFIP5_HUMAN\|Q9BXF6_S395 | 0.612803708 | 2.96E-05 | 5.16E-04 |
| UBXN7_HUMAN\|O94888_S285_S288 | 0.612781991 | 1.20E-03 | 1.73E-02 |
| GSK3A_HUMAN\|P49840_S21 | 0.612642741 | 2.65E-05 | 1.19E-03 |
| CCDC9_HUMAN\|Q9Y3X0_S386_T381 | 0.612624273 | 6.95E-05 | 6.73E-04 |
| DLG1_HUMAN\|Q12959_S571 | 0.61262188 | 1.92E-04 | 1.51E-03 |
| M3K3_HUMAN\|Q99759_S337_S340 | 0.612553381 | 1.21E-05 | 2.70E-04 |
| LMNA_HUMAN\|P02545_S628 | 0.612449054 | 6.62E-03 | 2.88E-02 |
| ARVC_HUMAN\|O00192_S203 | 0.612439851 | 7.19E-05 | 7.34E-04 |
| K1C14_HUMAN\|P02533_S17 | 0.612395978 | 1.98E-03 | 1.62E-02 |
| EIF3F_HUMAN\|O00303_S258 | 0.61230167 | 4.86E-03 | 4.80E-02 |
| M3K20_HUMAN\|Q9NYL2_S727 | 0.612184176 | 1.85E-06 | 1.80E-04 |

| | | | |
|---|---|---|---|
| OSBP1_HUMAN\|P22059_S379_S382_S386 | 0.612044388 | 1.09E-03 | 1.29E-02 |
| UBP2L_HUMAN\|Q14157_S467_S477 | 0.611935206 | 3.16E-05 | 6.97E-04 |
| AHNK_HUMAN\|Q09666_S2397 | 0.611894355 | 4.19E-04 | 1.16E-02 |
| PAR10_HUMAN\|Q53GL7_S431 | 0.611835532 | 9.32E-06 | 1.86E-03 |
| DBNL_HUMAN\|Q9UJU6-2_S232 | 0.611818842 | 2.85E-04 | 1.31E-02 |
| MTCL1_HUMAN\|Q9Y4B5_S196 | 0.611708964 | 2.09E-04 | 4.86E-03 |
| SRS10_HUMAN\|O75494_S106 | 0.61167042 | 1.18E-03 | 1.75E-02 |
| RAB12_HUMAN\|Q6IQ22_S21 | 0.611609334 | 6.48E-04 | 4.29E-03 |
| MTAP2_HUMAN\|P11137_S1782_S1786_S1790 | 0.611555716 | 3.45E-05 | 3.89E-04 |
| K1C14_HUMAN\|P02533_S435 | 0.611542698 | 4.28E-04 | 3.02E-03 |
| COHA1_HUMAN\|Q9UMD9_S130 | 0.611500032 | 7.55E-03 | 4.37E-02 |
| AHNK_HUMAN\|Q09666_T158 | 0.611491858 | 4.08E-03 | 1.92E-02 |
| AHNK_HUMAN\|Q09666_S3412 | 0.611428091 | 9.55E-05 | 9.30E-04 |
| DCBD2_HUMAN\|Q96PD2_S724_S727 | 0.611148027 | 1.76E-03 | 8.66E-03 |
| PDLI2_HUMAN\|Q96JY6_T279 | 0.611145904 | 1.93E-03 | 9.31E-03 |
| NDRG1_HUMAN\|Q92597_S330_S336 | 0.611044314 | 6.75E-04 | 1.19E-02 |
| CDK16_HUMAN\|Q00536_S95 | 0.610944483 | 6.19E-05 | 6.10E-04 |
| EXOC7_HUMAN\|Q9UPT5_S250 | 0.610943915 | 3.93E-05 | 4.29E-04 |
| RIOK1_HUMAN\|Q9BRS2_S21_S22 | 0.610912876 | 4.72E-04 | 9.44E-03 |
| CCM2_HUMAN\|Q9BSQ5_S384 | 0.610911199 | 1.12E-04 | 9.94E-04 |
| AHNK_HUMAN\|Q09666_T4766 | 0.610694142 | 1.08E-04 | 1.02E-03 |
| BORG4_HUMAN\|Q9H3Q1_S64 | 0.610442508 | 3.10E-05 | 6.86E-04 |
| AHNK2_HUMAN\|Q8IVF2_S294_T298 | 0.610287705 | 1.70E-04 | 4.61E-03 |
| ZC11A_HUMAN\|O75152_S132 | 0.610211791 | 1.47E-06 | 7.80E-05 |
| RGS3_HUMAN\|P49796_S1044 | 0.610165183 | 2.08E-05 | 5.26E-04 |
| MTAP2_HUMAN\|P11137_S1782_S1786_S1790 | 0.610107502 | 3.23E-05 | 3.76E-04 |
| NDRG2_HUMAN\|Q9UN36_S332_T330 | 0.610036896 | 3.09E-05 | 6.86E-04 |
| AHNK_HUMAN\|Q09666_T158 | 0.610013517 | 3.94E-03 | 4.00E-02 |
| PLEC_HUMAN\|Q15149_S4385_S4396 | 0.609974309 | 2.11E-03 | 1.12E-02 |
| KAZRN_HUMAN\|Q674X7-3_S8 | 0.60971856 | 2.85E-05 | 6.51E-04 |
| TCOF_HUMAN\|Q13428_T249 | 0.609664788 | 1.35E-04 | 2.02E-03 |

| | | | |
|---|---|---|---|
| K1C16_HUMAN\|P08779_S466 | 0.609642357 | 1.65E-04 | 1.33E-03 |
| VIME_HUMAN\|P08670_S51 | 0.609630803 | 1.33E-04 | 1.21E-03 |
| BCAR3_HUMAN\|O75815_S375 | 0.609463614 | 2.49E-04 | 3.33E-03 |
| RTKN_HUMAN\|Q9BST9_S561 | 0.60941795 | 7.81E-05 | 2.49E-03 |
| PAK2_HUMAN\|Q13177_S141 | 0.609404404 | 3.03E-04 | 2.27E-03 |
| MISP_HUMAN\|Q8IVT2_S376 | 0.609339352 | 8.64E-05 | 1.43E-03 |
| IF4B_HUMAN\|P23588_S597 | 0.60925346 | 6.01E-04 | 2.12E-02 |
| FA83H_HUMAN\|Q6ZRV2_S1098 | 0.609225957 | 2.51E-05 | 3.00E-04 |
| NUMA1_HUMAN\|Q14980_S1969 | 0.609190913 | 4.48E-03 | 2.48E-02 |
| ROA1_HUMAN\|P09651_S364 | 0.609188487 | 4.61E-03 | 1.88E-02 |
| SPR2E_HUMAN\|P22531_S60 | 0.60898052 | 2.42E-03 | 2.88E-02 |
| MILK1_HUMAN\|Q8N3F8_S323 | 0.608789298 | 3.02E-03 | 1.51E-02 |
| PLEC_HUMAN\|Q15149_S4385_S4396 | 0.60877024 | 2.04E-03 | 2.41E-02 |
| PKHA4_HUMAN\|Q9H4M7_S194 | 0.60874335 | 1.10E-04 | 3.00E-03 |
| NUP53_HUMAN\|Q8NFH5_S105 | 0.608739307 | 2.59E-04 | 5.96E-03 |
| RRBP1_HUMAN\|Q9P2E9_S615 | 0.608735882 | 7.04E-05 | 5.16E-03 |
| HCFC1_HUMAN\|P51610_S1507 | 0.608725195 | 1.32E-03 | 9.88E-03 |
| PHLB2_HUMAN\|Q86SQ0_T898 | 0.608625248 | 4.04E-03 | 1.70E-02 |
| LRRF2_HUMAN\|Q9Y608_S324 | 0.608388533 | 5.63E-04 | 3.82E-03 |
| S38A2_HUMAN\|Q96QD8_S17_S22 | 0.608365172 | 1.25E-03 | 1.17E-02 |
| NGAP_HUMAN\|Q9UJF2_S954 | 0.608346836 | 9.57E-05 | 2.82E-03 |
| K2C5_HUMAN\|P13647_S18_S21 | 0.608164233 | 4.21E-03 | 4.21E-02 |
| PAXX_HUMAN\|Q9BUH6_S148 | 0.608104392 | 1.30E-05 | 2.81E-04 |
| EIF3A_HUMAN\|Q14152_S882 | 0.608098823 | 1.57E-03 | 1.38E-02 |
| MAP7_HUMAN\|Q14244_T184 | 0.608086755 | 2.76E-05 | 6.35E-04 |
| LRRF2_HUMAN\|Q9Y608_S324 | 0.607990575 | 5.55E-04 | 3.44E-03 |
| K1C14_HUMAN\|P02533_S53 | 0.607868336 | 1.56E-05 | 2.07E-04 |
| RBM7_HUMAN\|Q9Y580_S137 | 0.607850224 | 5.78E-06 | 2.12E-04 |
| PAK6_HUMAN\|Q9NQU5_S347_S351 | 0.607794762 | 6.47E-04 | 6.77E-03 |
| MICA3_HUMAN\|Q7RTP6_S977 | 0.607741316 | 1.59E-07 | 6.13E-06 |
| K2C5_HUMAN\|P13647_S16 | 0.607725572 | 3.70E-04 | 2.52E-03 |

| | | | |
|---|---|---|---|
| UBXN7_HUMAN\|O94888_S285 | 0.607666748 | 1.55E-03 | 2.85E-02 |
| PKN2_HUMAN\|Q16513_S360 | 0.607504789 | 2.63E-04 | 1.92E-03 |
| MILK1_HUMAN\|Q8N3F8_S323 | 0.607492007 | 2.92E-03 | 1.31E-02 |
| SSFA2_HUMAN\|P28290_S668 | 0.607477193 | 4.51E-04 | 4.33E-03 |
| MAST4_HUMAN\|O15021_S2555 | 0.607376409 | 2.00E-04 | 1.56E-03 |
| ITA6_HUMAN\|P23229_S1128 | 0.607359039 | 3.36E-04 | 2.47E-03 |
| ZN768_HUMAN\|Q9H5H4_S83 | 0.607307927 | 4.84E-04 | 3.09E-03 |
| ZCHC8_HUMAN\|Q6NZY4_T479 | 0.607275244 | 2.02E-05 | 2.53E-04 |
| AHNK_HUMAN\|Q09666_S5864 | 0.607186866 | 4.40E-03 | 4.48E-02 |
| MARCS_HUMAN\|P29966_T150 | 0.606985941 | 2.91E-04 | 3.77E-03 |
| SARG_HUMAN\|Q9BW04_S502 | 0.606864583 | 4.84E-06 | 3.64E-04 |
| REEP4_HUMAN\|Q9H6H4_S253 | 0.606844881 | 9.08E-04 | 8.84E-03 |
| K2C5_HUMAN\|P13647_S16 | 0.606754018 | 3.57E-04 | 2.60E-03 |
| STX5_HUMAN\|Q13190_S9 | 0.60675264 | 2.35E-03 | 1.23E-02 |
| SPTB2_HUMAN\|Q01082-3_S14 | 0.606731398 | 1.88E-04 | 9.80E-03 |
| LAP2A_HUMAN\|P42166_T160 | 0.606612993 | 1.36E-05 | 1.87E-04 |
| TAOK3_HUMAN\|Q9H2K8_S442 | 0.606588904 | 2.41E-04 | 1.81E-03 |
| PEA15_HUMAN\|Q15121_S104 | 0.606584343 | 8.43E-04 | 1.88E-02 |
| ICE1_HUMAN\|Q9Y2F5_S1712 | 0.606558319 | 6.72E-05 | 6.56E-04 |
| TENS4_HUMAN\|Q8IZW8_S248_S253 | 0.606554044 | 1.45E-03 | 1.92E-02 |
| EDC3_HUMAN\|Q96F86_S131 | 0.606538174 | 1.40E-06 | 1.84E-04 |
| NU214_HUMAN\|P35658_S986_T987 | 0.606441391 | 3.08E-03 | 1.89E-02 |
| CBX3_HUMAN\|Q13185_S93 | 0.606391553 | 2.21E-03 | 2.50E-02 |
| WDR44_HUMAN\|Q5JSH3_S561 | 0.606313253 | 2.04E-05 | 7.94E-04 |
| ROCK1_HUMAN\|Q13464_S1341 | 0.60625351 | 2.13E-04 | 1.75E-03 |
| AHNK_HUMAN\|Q09666_S3426 | 0.606112758 | 2.09E-05 | 1.18E-03 |
| MTAP2_HUMAN\|P11137_S1787_S1790_T1780 | 0.606100758 | 6.53E-06 | 1.08E-04 |
| NUP98_HUMAN\|P52948_S623 | 0.605870549 | 2.28E-04 | 4.39E-03 |
| AHNK_HUMAN\|Q09666_S41 | 0.60583245 | 1.44E-03 | 1.23E-02 |
| AHNK_HUMAN\|Q09666_T4100 | 0.605807165 | 6.13E-04 | 4.10E-03 |
| K2C1_HUMAN\|P04264_S608_S612 | 0.605650643 | 5.80E-03 | 2.23E-02 |

| | | | |
|---|---|---|---|
| RNF10_HUMAN\|Q8N5U6_S41 | 0.605643067 | 2.54E-04 | 1.88E-03 |
| BRD4_HUMAN\|O60885_S1117 | 0.60561101 | 1.45E-03 | 2.73E-02 |
| TBCD4_HUMAN\|O60343_S570 | 0.605553105 | 6.45E-04 | 6.87E-03 |
| ACINU_HUMAN\|Q9UKV3_T254 | 0.605540421 | 8.88E-04 | 1.44E-02 |
| K1C15_HUMAN\|P19012_S48 | 0.605471231 | 5.27E-03 | 3.38E-02 |
| CD97_HUMAN\|P48960_S831 | 0.605392778 | 2.83E-05 | 1.51E-03 |
| CCDC9_HUMAN\|Q9Y3X0_S376_S386_S390 | 0.605374809 | 1.71E-03 | 2.31E-02 |
| 2A5D_HUMAN\|Q14738_S598 | 0.605347884 | 1.28E-04 | 1.17E-03 |
| VIME_HUMAN\|P08670_S325 | 0.605290173 | 1.58E-03 | 8.01E-03 |
| TAOK3_HUMAN\|Q9H2K8_S442 | 0.605263302 | 2.30E-04 | 3.15E-03 |
| LMNA_HUMAN\|P02545_S404_S407_T409 | 0.605094491 | 7.90E-04 | 4.57E-03 |
| SPTB2_HUMAN\|Q01082-3_S14_S8 | 0.605044255 | 2.30E-05 | 2.80E-04 |
| SRRM1_HUMAN\|Q8IYB3_S713_S715 | 0.605024959 | 1.32E-03 | 1.22E-02 |
| TLN1_HUMAN\|Q9Y490_T2270 | 0.60501917 | 1.29E-04 | 1.17E-03 |
| DOCK9_HUMAN\|Q9BZ29-5_S37 | 0.605004086 | 3.77E-04 | 3.80E-03 |
| IF4B_HUMAN\|P23588_S497_S498 | 0.604977157 | 1.26E-04 | 1.59E-03 |
| CE170_HUMAN\|Q5SW79_S630_S633 | 0.604888973 | 1.02E-05 | 3.05E-04 |
| PDLI2_HUMAN\|Q96JY6_S134 | 0.604842926 | 2.51E-07 | 8.40E-06 |
| PLCB3_HUMAN\|Q01970_S1105 | 0.604602005 | 3.07E-03 | 1.35E-02 |
| PRC2A_HUMAN\|P48634_S1004 | 0.604554511 | 6.03E-05 | 6.29E-04 |
| PHLB2_HUMAN\|Q86SQ0_S334_T328 | 0.604552458 | 1.08E-03 | 1.00E-02 |
| K1C17_HUMAN\|Q04695_S24 | 0.604525194 | 6.00E-05 | 2.36E-03 |
| TANK_HUMAN\|Q92844_S129 | 0.604518683 | 3.65E-04 | 7.73E-03 |
| OPTN_HUMAN\|Q96CV9_T204 | 0.604480665 | 2.75E-06 | 5.59E-05 |
| ABLM3_HUMAN\|O94929_S373_S379 | 0.604420508 | 3.44E-04 | 2.52E-03 |
| NDRG1_HUMAN\|Q92597_S330_S336 | 0.604416219 | 5.42E-04 | 3.38E-03 |
| NDRG1_HUMAN\|Q92597_S364 | 0.604367525 | 2.06E-05 | 5.16E-04 |
| JUN_HUMAN\|P05412_S63 | 0.604343902 | 3.03E-04 | 3.20E-03 |
| KMT2A_HUMAN\|Q03164_T3028 | 0.604312833 | 5.53E-03 | 2.48E-02 |
| TM1L1_HUMAN\|O75674_S321_S323 | 0.604273339 | 2.19E-04 | 1.22E-02 |
| YAP1_HUMAN\|P46937_S163 | 0.604081311 | 1.48E-03 | 1.88E-02 |

| | | | |
|---|---|---|---|
| Z280C_HUMAN\|Q8ND82_S80 | 0.604030111 | 1.79E-05 | 8.97E-04 |
| TASOR_HUMAN\|Q9UK61_S344 | 0.604013713 | 1.46E-05 | 1.73E-03 |
| PHRF1_HUMAN\|Q9P1Y6_S915_T917 | 0.604007395 | 4.37E-03 | 2.81E-02 |
| K2C6B_HUMAN\|P04259_S37 | 0.603762099 | 1.08E-03 | 2.20E-02 |
| PRC2A_HUMAN\|P48634_S1147 | 0.603721495 | 4.59E-06 | 1.82E-04 |
| LIPA3_HUMAN\|O75145_S1164 | 0.60360904 | 2.05E-05 | 2.92E-03 |
| MILK1_HUMAN\|Q8N3F8_S644 | 0.603599926 | 4.93E-05 | 2.23E-03 |
| RAI14_HUMAN\|Q9P0K7_T251 | 0.603524835 | 1.78E-04 | 9.46E-03 |
| GOGA4_HUMAN\|Q13439_S89_S92 | 0.603482747 | 5.34E-05 | 1.03E-03 |
| RAI14_HUMAN\|Q9P0K7_T297 | 0.603392166 | 4.65E-04 | 9.38E-03 |
| LIPA3_HUMAN\|O75145_S1164 | 0.603289132 | 2.02E-05 | 9.65E-04 |
| SRGP1_HUMAN\|Q7Z6B7_S906 | 0.603278628 | 7.44E-05 | 1.36E-03 |
| FBP1L_HUMAN\|Q5T0N5_S295 | 0.603274247 | 1.21E-04 | 1.11E-03 |
| PSF2_HUMAN\|Q9Y248_S182 | 0.603125919 | 2.59E-04 | 2.00E-03 |
| PSF2_HUMAN\|Q9Y248_S182_T180 | 0.603115803 | 5.99E-04 | 6.51E-03 |
| USO1_HUMAN\|O60763_S952 | 0.602950238 | 9.15E-04 | 1.13E-02 |
| SMTN_HUMAN\|P53814_S301 | 0.602923759 | 4.97E-03 | 1.99E-02 |
| CR025_HUMAN\|Q96B23_S67 | 0.6028397 | 5.78E-04 | 1.44E-02 |
| SPD2A_HUMAN\|Q5TCZ1_S724 | 0.602828073 | 8.84E-06 | 1.34E-04 |
| SOS2_HUMAN\|Q07890_S1267 | 0.602820031 | 6.05E-04 | 2.16E-02 |
| OPTN_HUMAN\|Q96CV9_T525 | 0.602819836 | 4.78E-06 | 1.34E-04 |
| K1C16_HUMAN\|P08779_S449 | 0.602755445 | 2.31E-04 | 1.85E-03 |
| NP1L1_HUMAN\|P55209_T62 | 0.602742027 | 1.45E-03 | 1.30E-02 |
| RGL2_HUMAN\|O15211_S761 | 0.602676001 | 4.73E-05 | 5.08E-04 |
| TENS4_HUMAN\|Q8IZW8_S242_S248_S253 | 0.602671708 | 3.84E-03 | 2.56E-02 |
| TBC16_HUMAN\|Q8TBP0_S103_T99 | 0.602644805 | 1.02E-04 | 2.97E-03 |
| IRS2_HUMAN\|Q9Y4H2_S606 | 0.6026278 | 2.10E-03 | 1.00E-02 |
| COHA1_HUMAN\|Q9UMD9_S130 | 0.602595683 | 6.18E-03 | 2.33E-02 |
| TT39B_HUMAN\|Q5VTQ0_S120 | 0.60254777 | 8.70E-04 | 1.90E-02 |
| SMG9_HUMAN\|Q9H0W8_S53 | 0.602463868 | 3.63E-06 | 6.69E-05 |
| SPD2A_HUMAN\|Q5TCZ1_S644 | 0.602343788 | 4.03E-04 | 2.86E-03 |

| | | | |
|---|---|---|---|
| ARVC_HUMAN\|O00192_T642 | 0.602227468 | 4.08E-05 | 8.41E-04 |
| CARL1_HUMAN\|Q5VZK9_S1290_T1278 | 0.602207758 | 2.15E-05 | 1.20E-03 |
| CRBG1_HUMAN\|Q9Y4K1_S22 | 0.602143992 | 3.31E-06 | 2.57E-04 |
| K1522_HUMAN\|Q9P206_S406 | 0.602119458 | 1.19E-04 | 1.05E-03 |
| NFIC_HUMAN\|P08651_S333_S339 | 0.601949785 | 6.31E-05 | 6.21E-04 |
| AHNK_HUMAN\|Q09666_S5749_S5763 | 0.601883039 | 7.84E-07 | 2.02E-05 |
| K1C14_HUMAN\|P02533_S429_S435 | 0.601815204 | 3.30E-04 | 2.29E-03 |
| KLRG2_HUMAN\|A4D1S0_S215 | 0.601813559 | 2.02E-04 | 4.02E-03 |
| K1C14_HUMAN\|P02533_S433_S438 | 0.601805974 | 3.39E-04 | 2.49E-03 |
| MICA2_HUMAN\|O94851_S699 | 0.601794931 | 8.02E-05 | 2.06E-03 |
| KI26B_HUMAN\|Q2KJY2_S1006_S1018_S1021 | 0.601771999 | 1.76E-06 | 5.97E-04 |
| AHNK_HUMAN\|Q09666_S5752 | 0.60163904 | 1.44E-06 | 1.84E-04 |
| STK10_HUMAN\|O94804_S450_S454 | 0.601543419 | 1.87E-04 | 2.63E-03 |
| K2C5_HUMAN\|P13647_S11_S14 | 0.601335752 | 5.37E-06 | 1.87E-04 |
| AFAD_HUMAN\|P55196_T1330 | 0.601276731 | 5.40E-05 | 1.03E-03 |
| ARHGB_HUMAN\|O15085_T668_T672 | 0.60127581 | 6.85E-06 | 2.32E-04 |
| K1C16_HUMAN\|P08779_S450 | 0.601256121 | 5.16E-05 | 5.49E-04 |
| FOSL2_HUMAN\|P15408_S19 | 0.601208539 | 2.61E-03 | 2.49E-02 |
| MAP4_HUMAN\|P27816_S941 | 0.601115527 | 8.31E-05 | 1.40E-03 |
| PKN2_HUMAN\|Q16513_S362 | 0.601101707 | 6.55E-05 | 6.75E-04 |
| MRP4_HUMAN\|O15439_T646 | 0.601069123 | 4.36E-04 | 3.07E-03 |
| LMNA_HUMAN\|P02545_S404_S406_S414 | 0.600855823 | 1.88E-03 | 9.10E-03 |
| REQU_HUMAN\|Q92785_S142 | 0.600818434 | 8.15E-05 | 3.25E-03 |
| K2C75_HUMAN\|O95678_S535 | 0.60076085 | 1.99E-06 | 8.77E-05 |
| ARAF_HUMAN\|P10398_S257_T253 | 0.600660592 | 9.77E-05 | 8.99E-04 |
| IP3KB_HUMAN\|P27987_S264 | 0.600587574 | 3.54E-04 | 2.58E-03 |
| FA83H_HUMAN\|Q6ZRV2_S1024 | 0.600483125 | 3.87E-04 | 2.60E-03 |
| F193A_HUMAN\|P78312_S1144 | 0.600426498 | 3.24E-05 | 1.54E-03 |
| TAOK3_HUMAN\|Q9H2K8_S324 | 0.600409024 | 9.33E-06 | 2.15E-04 |
| STIM1_HUMAN\|Q13586_S668 | 0.600380044 | 1.53E-03 | 2.07E-02 |
| SRC8_HUMAN\|Q14247_S405_T401 | 0.600334545 | 2.30E-03 | 4.61E-02 |

| | | | |
|---|---|---|---|
| CCDC9_HUMAN\|Q9Y3X0_S386_T381 | 0.600064779 | 4.13E-05 | 8.58E-04 |
| LIMA1_HUMAN\|Q9UHB6_Y229 | 0.600006149 | 1.00E-03 | 1.45E-02 |
| ADRM1_HUMAN\|Q16186_S211 | 0.599992927 | 7.37E-05 | 7.06E-04 |
| K2C1_HUMAN\|P04264_S612 | 0.599954104 | 5.89E-03 | 2.26E-02 |
| MYPT1_HUMAN\|O14974_S507 | 0.599881641 | 5.69E-05 | 5.69E-04 |
| AHNK2_HUMAN\|Q8IVF2_S294_T298 | 0.599808741 | 1.14E-04 | 3.49E-03 |
| PKP3_HUMAN\|Q9Y446_S314 | 0.599790753 | 9.69E-06 | 1.42E-04 |
| ASPC1_HUMAN\|Q9BZE9_S275 | 0.599760454 | 2.05E-05 | 2.60E-04 |
| OSBL8_HUMAN\|Q9BZF1_S807_S808_S810 | 0.599727991 | 3.09E-04 | 5.49E-03 |
| K1C14_HUMAN\|P02533_S433 | 0.599685118 | 2.08E-03 | 9.91E-03 |
| BIG1_HUMAN\|Q9Y6D6_S397 | 0.599684443 | 1.54E-05 | 5.58E-03 |
| UBP2L_HUMAN\|Q14157_S477 | 0.599680911 | 9.83E-04 | 1.55E-02 |
| HNRPC_HUMAN\|P07910_S239 | 0.599651083 | 1.25E-05 | 1.75E-04 |
| ZCHC8_HUMAN\|Q6NZY4_T479 | 0.599618115 | 1.44E-05 | 7.86E-04 |
| CE170_HUMAN\|Q5SW79_S633_S636 | 0.599590364 | 1.94E-03 | 3.37E-02 |
| TWF1_HUMAN\|Q12792_T349 | 0.599451486 | 3.34E-06 | 6.39E-05 |
| AHNK_HUMAN\|Q09666_S216_S220 | 0.59943127 | 5.31E-06 | 9.08E-05 |
| NDRG1_HUMAN\|Q92597_S333_S336_T328 | 0.599405855 | 4.41E-06 | 3.11E-04 |
| DLG1_HUMAN\|Q12959_S571 | 0.599371317 | 1.14E-04 | 3.49E-03 |
| SMG9_HUMAN\|Q9H0W8_T56 | 0.599335825 | 1.78E-05 | 2.30E-04 |
| PLEC_HUMAN\|Q15149_S4613_S4618 | 0.59923895 | 2.01E-05 | 1.09E-03 |
| WDR13_HUMAN\|Q9H1Z4_S70_S74 | 0.599234929 | 3.35E-05 | 1.58E-03 |
| PRKDC_HUMAN\|P78527_S3205 | 0.59922775 | 1.45E-03 | 8.19E-03 |
| ABI2_HUMAN\|Q9NYB9_S227 | 0.599004896 | 5.65E-04 | 1.06E-02 |
| DDI2_HUMAN\|Q5TDH0_S194 | 0.599000938 | 9.16E-05 | 8.49E-04 |
| EMAL4_HUMAN\|Q9HC35_S146 | 0.59895926 | 5.23E-05 | 2.12E-03 |
| PAK6_HUMAN\|Q9NQU5_S308 | 0.598937624 | 1.72E-03 | 8.51E-03 |
| DCAF6_HUMAN\|Q58WW2_S657 | 0.598912641 | 1.62E-04 | 3.38E-03 |
| SPIR1_HUMAN\|Q08AE8_T509 | 0.598854497 | 3.28E-05 | 3.74E-04 |
| GSK3A_HUMAN\|P49840_S21 | 0.598839521 | 1.46E-05 | 7.91E-04 |
| ITB4_HUMAN\|P16144_S1454_T1455 | 0.598822761 | 1.67E-03 | 1.36E-02 |

| | | | |
|---|---|---|---|
| LMNA_HUMAN|P02545_S403_S406 | 0.598662334 | 8.69E-05 | 8.55E-04 |
| MARCS_HUMAN|P29966_T150 | 0.598659253 | 2.14E-04 | 2.91E-03 |
| AHNK_HUMAN|Q09666_T4564 | 0.598645423 | 1.21E-04 | 1.11E-03 |
| PFD4_HUMAN|Q9NQP4_S125 | 0.598601552 | 3.08E-04 | 2.29E-03 |
| KIZ_HUMAN|Q2M2Z5_S623 | 0.598531082 | 8.55E-06 | 1.31E-04 |
| TPPP_HUMAN|O94811_S160 | 0.598530392 | 6.66E-04 | 1.14E-02 |
| RHG29_HUMAN|Q52LW3_S176 | 0.59837632 | 9.39E-04 | 1.50E-02 |
| K2C6B_HUMAN|P04259_S541 | 0.598361763 | 5.31E-05 | 1.02E-03 |
| CC120_HUMAN|Q96HB5_S296 | 0.598269988 | 3.13E-03 | 3.45E-02 |
| PPR18_HUMAN|Q6NYC8_S490 | 0.598248663 | 2.24E-06 | 7.32E-05 |
| ARVC_HUMAN|O00192_S203 | 0.598045838 | 3.80E-05 | 8.06E-04 |
| SPD2A_HUMAN|Q5TCZ1_S644 | 0.597962413 | 3.49E-04 | 6.93E-03 |
| LARP1_HUMAN|Q6PKG0_S766_S774 | 0.597929609 | 6.37E-04 | 1.10E-02 |
| MPLKI_HUMAN|Q8TAP9_S80 | 0.597914729 | 3.74E-03 | 1.79E-02 |
| K1C16_HUMAN|P08779_S448_S451 | 0.597913685 | 9.20E-05 | 9.04E-04 |
| MFF_HUMAN|Q9GZY8_S157 | 0.597890417 | 8.45E-04 | 1.88E-02 |
| ERF_HUMAN|P50548_S24 | 0.597876897 | 1.54E-05 | 3.15E-04 |
| TB182_HUMAN|Q9C0C2_S920 | 0.59781669 | 9.06E-05 | 2.76E-03 |
| SSFA2_HUMAN|P28290_S668 | 0.597740755 | 3.27E-04 | 1.45E-02 |
| DAPK3_HUMAN|O43293_S311_S312 | 0.597634856 | 1.59E-05 | 9.32E-04 |
| ITB4_HUMAN|P16144_S1486_T1487 | 0.597602944 | 7.39E-04 | 2.44E-02 |
| ITB4_HUMAN|P16144_T1455 | 0.597573505 | 2.29E-04 | 3.15E-03 |
| SI1L1_HUMAN|O43166_S1255 | 0.597569357 | 2.67E-04 | 5.81E-03 |
| BORG4_HUMAN|Q9H3Q1_S73 | 0.597542985 | 1.24E-03 | 6.57E-03 |
| MILK1_HUMAN|Q8N3F8_S578 | 0.597522397 | 3.80E-06 | 7.05E-05 |
| NFIX_HUMAN|Q14938_S265 | 0.597489765 | 6.39E-04 | 1.56E-02 |
| TB182_HUMAN|Q9C0C2_S1533 | 0.597411683 | 5.32E-05 | 5.39E-04 |
| DESP_HUMAN|P15924_S176 | 0.597284263 | 3.93E-03 | 1.87E-02 |
| NDRG1_HUMAN|Q92597_T366 | 0.597279272 | 1.14E-04 | 1.87E-03 |
| CCAR2_HUMAN|Q8N163_S678 | 0.597241916 | 6.28E-03 | 4.68E-02 |
| PHLB2_HUMAN|Q86SQ0_S334 | 0.597200488 | 1.10E-04 | 1.04E-03 |

| | | | |
|---|---|---|---|
| TFE3_HUMAN\|P19532_S556 | 0.597143786 | 1.25E-04 | 2.49E-03 |
| MAP7_HUMAN\|Q14244_S209 | 0.597115182 | 3.12E-04 | 6.74E-03 |
| AKTS1_HUMAN\|Q96B36_S187 | 0.597092574 | 1.98E-04 | 5.07E-03 |
| BAIP2_HUMAN\|Q9UQB8_Y491 | 0.597070083 | 8.24E-03 | 4.63E-02 |
| CREB1_HUMAN\|P16220_S271 | 0.59700252 | 3.97E-04 | 2.64E-03 |
| FA83B_HUMAN\|Q5T0W9_S664 | 0.596974345 | 6.57E-06 | 4.56E-04 |
| IRS2_HUMAN\|Q9Y4H2_S1186_T1202 | 0.596946022 | 2.13E-06 | 7.13E-05 |
| GRIN1_HUMAN\|Q7Z2K8_S615 | 0.596887683 | 2.06E-05 | 2.56E-04 |
| CTU2_HUMAN\|Q2VPK5_S419 | 0.596855674 | 1.05E-06 | 2.68E-05 |
| K2C5_HUMAN\|P13647_S572 | 0.596854237 | 8.31E-05 | 1.40E-03 |
| ABI2_HUMAN\|Q9NYB9_S227 | 0.596811447 | 5.27E-04 | 5.83E-03 |
| NCOA7_HUMAN\|Q8NI08_S384 | 0.59671893 | 3.33E-06 | 2.58E-04 |
| ARFG1_HUMAN\|Q8N6T3_T135 | 0.596650057 | 1.30E-05 | 8.62E-04 |
| EPN3_HUMAN\|Q9H201_S188_S191 | 0.596462705 | 1.03E-04 | 9.92E-04 |
| GRSF1_HUMAN\|Q12849_S118 | 0.596459711 | 2.41E-05 | 8.70E-04 |
| ZO1_HUMAN\|Q07157_T868_Y843 | 0.596353662 | 1.99E-03 | 2.08E-02 |
| Z280C_HUMAN\|Q8ND82_S80 | 0.596337513 | 1.24E-05 | 2.71E-04 |
| LMNA_HUMAN\|P02545_S404_S406 | 0.596333171 | 3.88E-05 | 4.33E-04 |
| CCDC9_HUMAN\|Q9Y3X0_S376 | 0.596305744 | 1.55E-05 | 4.23E-04 |
| SARG_HUMAN\|Q9BW04_T151 | 0.59630329 | 3.72E-03 | 4.95E-02 |
| LMNA_HUMAN\|P02545_S414 | 0.596241482 | 5.02E-05 | 5.23E-04 |
| IRS2_HUMAN\|Q9Y4H2_S679 | 0.596162252 | 2.61E-05 | 3.08E-04 |
| SPTB2_HUMAN\|Q01082_T2328 | 0.596157661 | 4.56E-03 | 1.87E-02 |
| T22D3_HUMAN\|Q99576-3_S34 | 0.59598237 | 6.05E-03 | 2.29E-02 |
| CO052_HUMAN\|Q6ZUT6_S55 | 0.595966382 | 2.20E-03 | 1.67E-02 |
| COHA1_HUMAN\|Q9UMD9_S130 | 0.595953671 | 5.30E-03 | 2.40E-02 |
| SMC1A_HUMAN\|Q14683_S956 | 0.595906014 | 2.16E-04 | 5.22E-03 |
| GGA2_HUMAN\|Q9UJY4_S177 | 0.595905916 | 1.69E-04 | 9.08E-03 |
| K2C8_HUMAN\|P05787_S24 | 0.595903595 | 3.17E-04 | 3.97E-03 |
| ITB4_HUMAN\|P16144_S1457_T1455 | 0.595886219 | 5.20E-03 | 3.16E-02 |
| TBCD5_HUMAN\|Q92609_S43 | 0.595857047 | 8.17E-05 | 8.07E-04 |

| | | | |
|---|---|---|---|
| PAK2_HUMAN\|Q13177_Y139 | 0.595811922 | 2.54E-04 | 1.98E-03 |
| HNRPC_HUMAN\|P07910_S240 | 0.59580714 | 5.73E-05 | 5.72E-04 |
| CCNL2_HUMAN\|Q96S94_S330 | 0.595531053 | 1.16E-06 | 1.67E-04 |
| NECP2_HUMAN\|Q9NVZ3_T182 | 0.595492235 | 5.91E-06 | 2.15E-04 |
| OSB10_HUMAN\|Q9BXB5_S26_S32 | 0.595474871 | 1.28E-06 | 1.33E-04 |
| HSPB1_HUMAN\|P04792_S78_S82 | 0.595416004 | 7.01E-04 | 6.07E-03 |
| NUP53_HUMAN\|Q8NFH5_S105 | 0.595229717 | 1.61E-04 | 4.50E-03 |
| SSH2_HUMAN\|Q76I76_T486 | 0.595099938 | 1.07E-05 | 1.58E-04 |
| K1C17_HUMAN\|Q04695_S12 | 0.595064145 | 2.28E-04 | 1.83E-03 |
| HMGN1_HUMAN\|P05114_S7 | 0.595052246 | 1.83E-04 | 4.39E-03 |
| FA83H_HUMAN\|Q6ZRV2_S1098 | 0.595010023 | 1.31E-05 | 6.98E-04 |
| AHNK2_HUMAN\|Q8IVF2_S294_T298 | 0.594876575 | 9.36E-05 | 3.66E-03 |
| PP1R7_HUMAN\|Q15435_S24 | 0.594852199 | 6.82E-04 | 1.14E-02 |
| KAZRN_HUMAN\|Q674X7-3_S8 | 0.59474834 | 1.45E-05 | 8.97E-04 |
| MFAP1_HUMAN\|P55081_S80 | 0.594743292 | 1.71E-06 | 3.72E-05 |
| TBCD5_HUMAN\|Q92609_S43 | 0.594694295 | 7.82E-05 | 1.35E-03 |
| RNF10_HUMAN\|Q8N5U6_S41 | 0.594669421 | 1.67E-04 | 1.45E-03 |
| MTAP2_HUMAN\|P11137_T1602_T1605 | 0.594635599 | 1.56E-03 | 1.73E-02 |
| ARP19_HUMAN\|P56211_S62 | 0.594634082 | 1.85E-05 | 4.81E-04 |
| TCOF_HUMAN\|Q13428_T249 | 0.594633867 | 7.42E-05 | 7.08E-04 |
| JAM1_HUMAN\|Q9Y624_S296 | 0.594550352 | 1.44E-02 | 4.63E-02 |
| PNISR_HUMAN\|Q8TF01_S762 | 0.59445721 | 1.13E-03 | 6.05E-03 |
| TDIF2_HUMAN\|Q5QJE6_T232 | 0.594457077 | 3.19E-03 | 2.23E-02 |
| LRRF2_HUMAN\|Q9Y608_S320_S323 | 0.594397315 | 2.80E-04 | 2.02E-03 |
| PHLB2_HUMAN\|Q86SQ0_T898 | 0.594396388 | 2.85E-03 | 2.04E-02 |
| CR025_HUMAN\|Q96B23_S145 | 0.594361704 | 1.41E-06 | 5.21E-05 |
| FA83B_HUMAN\|Q5T0W9_S388 | 0.594354219 | 1.92E-04 | 1.61E-03 |
| KC1D_HUMAN\|P48730_S331 | 0.594196468 | 2.81E-04 | 2.02E-03 |
| OSB11_HUMAN\|Q9BXB4_S174_S181 | 0.594193171 | 1.89E-03 | 2.46E-02 |
| PTN1_HUMAN\|P18031_S50 | 0.594187381 | 2.73E-03 | 2.89E-02 |
| PHLB1_HUMAN\|Q86UU1_S430 | 0.594159084 | 7.60E-05 | 2.48E-03 |

| | | | |
|---|---|---|---|
| ACAP2_HUMAN\|Q15057_S384 | 0.59413974 | 1.05E-06 | 1.08E-04 |
| FA83H_HUMAN\|Q6ZRV2_S1098 | 0.594116269 | 1.26E-05 | 3.57E-04 |
| NCOA7_HUMAN\|Q8NI08_S208_T210 | 0.594091221 | 1.29E-05 | 2.79E-04 |
| MA7D1_HUMAN\|Q3KQU3_S313 | 0.594090739 | 4.44E-04 | 3.12E-03 |
| DOCK6_HUMAN\|Q96HP0_S880 | 0.594055985 | 1.60E-04 | 1.41E-03 |
| ZO1_HUMAN\|Q07157_T868_Y843 | 0.594029305 | 1.87E-03 | 1.96E-02 |
| LMNA_HUMAN\|P02545_S303 | 0.594007408 | 6.71E-04 | 3.99E-03 |
| COBL1_HUMAN\|Q53SF7_T298 | 0.593930772 | 8.55E-05 | 2.60E-03 |
| K1C17_HUMAN\|Q04695_S10 | 0.593836011 | 1.07E-04 | 1.01E-03 |
| MA7D1_HUMAN\|Q3KQU3_S366 | 0.593769682 | 5.33E-07 | 6.65E-05 |
| TISB_HUMAN\|Q07352_S92 | 0.593758669 | 2.46E-04 | 1.83E-03 |
| BORG4_HUMAN\|Q9H3Q1_S64 | 0.593730908 | 1.47E-05 | 8.97E-04 |
| G3BP1_HUMAN\|Q13283_S230 | 0.593666469 | 5.64E-04 | 1.00E-02 |
| NHRF1_HUMAN\|O14745_S280 | 0.59364865 | 1.02E-05 | 6.28E-04 |
| GRAP1_HUMAN\|Q4V328_S666 | 0.593637495 | 1.32E-04 | 2.00E-03 |
| LIMA1_HUMAN\|Q9UHB6_S617 | 0.593630315 | 3.84E-04 | 2.58E-03 |
| OPTN_HUMAN\|Q96CV9_T204 | 0.593615054 | 1.53E-06 | 3.43E-05 |
| TGS1_HUMAN\|Q96RS0_S85_S89 | 0.593525754 | 7.78E-04 | 8.10E-03 |
| ARMX3_HUMAN\|Q9UH62_S110 | 0.593290712 | 1.38E-05 | 2.91E-04 |
| ATX2L_HUMAN\|Q8WWM7_S306 | 0.593239658 | 4.25E-03 | 4.15E-02 |
| SLK_HUMAN\|Q9H2G2_S655 | 0.593124436 | 3.20E-04 | 2.25E-03 |
| ARH40_HUMAN\|Q8TER5_T371 | 0.593120797 | 5.91E-07 | 7.69E-05 |
| MISP_HUMAN\|Q8IVT2_S575 | 0.593090988 | 4.61E-05 | 7.14E-04 |
| MA7D1_HUMAN\|Q3KQU3_S834 | 0.593076799 | 8.83E-05 | 2.63E-03 |
| UT14A_HUMAN\|Q9BVJ6_S437 | 0.593065209 | 1.79E-04 | 2.10E-03 |
| RBP10_HUMAN\|Q6VN20_S365 | 0.59300894 | 2.08E-06 | 9.96E-05 |
| UBP24_HUMAN\|Q9UPU5_S1143 | 0.59291927 | 1.73E-05 | 2.25E-04 |
| CE170_HUMAN\|Q5SW79_T628 | 0.592838468 | 5.89E-03 | 4.48E-02 |
| M4K4_HUMAN\|O95819_S639 | 0.5928321 | 4.90E-04 | 9.62E-03 |
| ZYX_HUMAN\|Q15942_S259_S267 | 0.592812554 | 4.11E-05 | 1.53E-03 |
| K2C6B_HUMAN\|P04259_S546 | 0.592691366 | 1.58E-05 | 2.09E-04 |

| | | | |
|---|---|---|---|
| FA83G_HUMAN\|A6ND36_S610_S614 | 0.592576115 | 2.84E-04 | 2.04E-03 |
| NDE1_HUMAN\|Q9NXR1_S282 | 0.592572702 | 1.62E-03 | 2.14E-02 |
| UBP24_HUMAN\|Q9UPU5_S1141 | 0.592554049 | 2.53E-05 | 3.07E-04 |
| CAMP2_HUMAN\|Q08AD1_S980 | 0.592505748 | 1.86E-04 | 1.47E-03 |
| NIBL1_HUMAN\|Q96TA1_S692_S696 | 0.592470384 | 5.24E-05 | 5.08E-03 |
| GGA2_HUMAN\|Q9UJY4_S177 | 0.592456027 | 1.48E-04 | 3.85E-03 |
| ARH40_HUMAN\|Q8TER5_S262 | 0.592338144 | 4.96E-06 | 1.91E-04 |
| ARG35_HUMAN\|A5YM69_S450 | 0.592222519 | 4.37E-04 | 8.50E-03 |
| UBE4B_HUMAN\|O95155_S84_S88 | 0.592167114 | 9.68E-04 | 5.91E-03 |
| NDRG1_HUMAN\|Q92597_S364 | 0.592132929 | 1.16E-05 | 1.68E-04 |
| LMNA_HUMAN\|P02545_S407_S414 | 0.592121924 | 4.63E-05 | 4.93E-04 |
| ATRX_HUMAN\|P46100_S784 | 0.592121208 | 4.41E-04 | 7.07E-03 |
| IWS1_HUMAN\|Q96ST2_S398 | 0.591987443 | 2.60E-03 | 1.66E-02 |
| PACN3_HUMAN\|Q9UKS6_S383 | 0.591987108 | 1.55E-03 | 1.37E-02 |
| CTNA1_HUMAN\|P35221_S641_S652 | 0.591974632 | 4.89E-05 | 5.12E-04 |
| PGAM1_HUMAN\|P18669_S14 | 0.591926182 | 7.65E-05 | 7.64E-04 |
| HIRP3_HUMAN\|Q9BW71_S196_S199 | 0.591838028 | 9.02E-06 | 1.40E-04 |
| K2C6A_HUMAN\|P02538_S546 | 0.591835697 | 1.46E-02 | 4.69E-02 |
| SEPT9_HUMAN\|Q9UHD8_T38_T42 | 0.591834836 | 6.46E-06 | 1.07E-04 |
| K1C14_HUMAN\|P02533_S13 | 0.591597658 | 4.32E-06 | 7.94E-05 |
| BORG4_HUMAN\|Q9H3Q1_S292 | 0.591565144 | 2.08E-04 | 2.85E-03 |
| PP1R7_HUMAN\|Q15435_S24 | 0.591557592 | 6.16E-04 | 9.03E-03 |
| PDLI2_HUMAN\|Q96JY6_S137_S143 | 0.591542656 | 2.58E-05 | 1.17E-03 |
| SRC8_HUMAN\|Q14247_S405_T401 | 0.591439042 | 1.81E-03 | 1.26E-02 |
| T22D4_HUMAN\|Q9Y3Q8_T167 | 0.591239344 | 3.80E-05 | 1.48E-03 |
| SRRM1_HUMAN\|Q8IYB3_T846 | 0.591228018 | 4.74E-04 | 9.44E-03 |
| MTAP2_HUMAN\|P11137_S1787_S1790_T1780 | 0.591215526 | 3.04E-06 | 5.91E-05 |
| K2C8_HUMAN\|P05787_S21_S24_S27 | 0.591209632 | 8.34E-05 | 1.40E-03 |
| RAI14_HUMAN\|Q9P0K7_S281_S290 | 0.591195552 | 1.67E-03 | 3.98E-02 |
| SSFA2_HUMAN\|P28290_S92 | 0.591174436 | 4.85E-03 | 3.19E-02 |
| DCBD2_HUMAN\|Q96PD2_S726_S727 | 0.590970044 | 1.23E-03 | 6.53E-03 |

| | | | |
|---|---|---|---|
| LMNA_HUMAN\|P02545_S392_S395 | 0.590734008 | 3.69E-03 | 4.93E-02 |
| MTMRA_HUMAN\|Q9NXD2_T770 | 0.590725965 | 4.90E-03 | 2.24E-02 |
| TENS4_HUMAN\|Q8IZW8_S248_S253 | 0.590697163 | 9.14E-04 | 8.88E-03 |
| NUCKS_HUMAN\|Q9H1E3_S181_T179 | 0.590663735 | 1.77E-03 | 1.41E-02 |
| AHNK_HUMAN\|Q09666_S4092 | 0.590601811 | 1.18E-03 | 2.35E-02 |
| STIM1_HUMAN\|Q13586_S668 | 0.590522966 | 1.13E-03 | 6.71E-03 |
| VIP2_HUMAN\|O43314_S1006_S1013_S1016 | 0.59040635 | 1.01E-06 | 5.83E-05 |
| ML12A_HUMAN\|P19105_T18 | 0.59033152 | 6.10E-05 | 6.33E-04 |
| NDE1_HUMAN\|Q9NXR1_S282 | 0.590280133 | 1.52E-03 | 7.74E-03 |
| PLXB2_HUMAN\|O15031_S1570 | 0.590264988 | 1.69E-04 | 1.36E-03 |
| I2BP2_HUMAN\|Q7Z5L9_S455_S460 | 0.590110347 | 4.87E-05 | 7.48E-04 |
| XCT_HUMAN\|Q9UPY5_S26 | 0.590080364 | 4.37E-04 | 4.22E-03 |
| COHA1_HUMAN\|Q9UMD9_S172_T176_T182 | 0.589993991 | 5.53E-03 | 3.31E-02 |
| AFAD_HUMAN\|P55196_S1172_S1173_S1182 | 0.589969851 | 2.64E-06 | 1.18E-04 |
| ERF_HUMAN\|P50548_T526 | 0.589954076 | 1.28E-04 | 1.10E-03 |
| KIZ_HUMAN\|Q2M2Z5_S623 | 0.589865853 | 5.53E-06 | 3.66E-04 |
| AHNK_HUMAN\|Q09666_S5031 | 0.589857319 | 7.24E-05 | 7.37E-04 |
| FA83H_HUMAN\|Q6ZRV2_S513_S516 | 0.589823886 | 4.27E-04 | 4.97E-03 |
| K1671_HUMAN\|Q9BY89_S128 | 0.589817811 | 1.21E-06 | 6.76E-05 |
| XPC_HUMAN\|Q01831_S883_S884_S891 | 0.589785731 | 4.17E-04 | 4.90E-03 |
| PRAG1_HUMAN\|Q86YV5_S510 | 0.589772331 | 2.98E-04 | 2.13E-03 |
| MICA2_HUMAN\|O94851_S699 | 0.589756508 | 4.78E-05 | 1.24E-03 |
| BORG4_HUMAN\|Q9H3Q1_S64 | 0.589730502 | 1.23E-05 | 7.84E-04 |
| CTNA1_HUMAN\|P35221_S652_S655_T658 | 0.589707407 | 6.23E-04 | 4.15E-03 |
| SAFB1_HUMAN\|Q15424_S604 | 0.589657059 | 2.23E-07 | 7.94E-06 |
| RFIP1_HUMAN\|Q6WKZ4_S501 | 0.589548307 | 8.01E-04 | 5.13E-03 |
| ES8L1_HUMAN\|Q8TE68_S182_T187 | 0.589508987 | 7.80E-07 | 1.32E-04 |
| GRB10_HUMAN\|Q13322_S104 | 0.589400584 | 1.01E-06 | 2.50E-05 |
| ELAV1_HUMAN\|Q15717_S202 | 0.589372767 | 4.32E-04 | 8.69E-03 |
| ZO2_HUMAN\|Q9UDY2_T1027 | 0.589317769 | 1.14E-05 | 6.43E-04 |
| RTKN_HUMAN\|Q9BST9_S561 | 0.589313947 | 3.56E-05 | 6.01E-04 |

| | | | |
|---|---|---|---|
| TENS4_HUMAN\|Q8IZW8_S253 | 0.589152879 | 2.86E-06 | 5.76E-05 |
| TAOK3_HUMAN\|Q9H2K8_S324 | 0.589059876 | 5.57E-06 | 4.13E-04 |
| DP13A_HUMAN\|Q9UKG1_S401 | 0.588969748 | 9.45E-05 | 3.05E-03 |
| ARHGG_HUMAN\|Q5VV41_S107 | 0.588942616 | 5.02E-07 | 1.43E-05 |
| ARP19_HUMAN\|P56211_Y59 | 0.588848587 | 9.76E-06 | 3.01E-04 |
| K1C17_HUMAN\|Q04695_S24 | 0.588813247 | 3.03E-05 | 6.76E-04 |
| YETS2_HUMAN\|Q9ULM3_S536 | 0.588625491 | 1.15E-05 | 1.68E-04 |
| COHA1_HUMAN\|Q9UMD9_S79 | 0.588581121 | 1.42E-03 | 7.31E-03 |
| CSCL2_HUMAN\|Q5T3F8_T110 | 0.588511819 | 6.90E-04 | 9.17E-03 |
| AHNK_HUMAN\|Q09666_S210_S216 | 0.588425758 | 1.43E-05 | 8.53E-04 |
| UBP2L_HUMAN\|Q14157_S477 | 0.588339275 | 6.87E-04 | 4.06E-03 |
| FOSL1_HUMAN\|P15407_S252_T267 | 0.588235481 | 6.07E-03 | 3.08E-02 |
| NIBL1_HUMAN\|Q96TA1_S641 | 0.588068345 | 9.00E-04 | 7.31E-03 |
| G3BP1_HUMAN\|Q13283_S230 | 0.588018035 | 4.73E-04 | 5.39E-03 |
| ADAM9_HUMAN\|Q13443_S758 | 0.587912088 | 7.65E-04 | 9.92E-03 |
| LRRF2_HUMAN\|Q9Y608_S320_S323 | 0.587858863 | 2.21E-04 | 5.30E-03 |
| DNJC1_HUMAN\|Q96KC8_S381 | 0.587843864 | 2.95E-03 | 2.73E-02 |
| REPS1_HUMAN\|Q96D71_S162_S166_T173 | 0.587767829 | 2.57E-06 | 2.21E-04 |
| ITA6_HUMAN\|P23229_S1128 | 0.587732259 | 1.60E-04 | 2.38E-03 |
| NEK9_HUMAN\|Q8TD19_S332 | 0.587725201 | 6.09E-03 | 2.30E-02 |
| ABCF1_HUMAN\|Q8NE71_S109_T108 | 0.587721358 | 8.37E-07 | 2.13E-05 |
| AHNK_HUMAN\|Q09666_S5752_S5763 | 0.587548453 | 2.93E-04 | 3.65E-02 |
| ABCF1_HUMAN\|Q8NE71_S109_T108 | 0.587532965 | 8.29E-07 | 2.24E-05 |
| AHNK_HUMAN\|Q09666_S5031 | 0.587354396 | 6.57E-05 | 1.22E-03 |
| EPHA2_HUMAN\|P29317_S897_S901 | 0.587271033 | 2.05E-04 | 2.34E-03 |
| AHNK_HUMAN\|Q09666_S5749 | 0.587111578 | 2.26E-07 | 6.11E-05 |
| AHNK_HUMAN\|Q09666_S5784 | 0.587074435 | 1.15E-02 | 3.85E-02 |
| EFNB1_HUMAN\|P98172_T284 | 0.587024034 | 1.53E-03 | 2.82E-02 |
| KANK1_HUMAN\|Q14678_S144 | 0.586959992 | 7.79E-08 | 3.58E-06 |
| B4DKV4_HUMAN\|B4DKV4_S19 | 0.586602743 | 1.84E-03 | 8.99E-03 |
| IASPP_HUMAN\|Q8WUF5_T341 | 0.586427257 | 4.46E-03 | 3.00E-02 |

| | | | |
|---|---|---|---|
| MA7D1_HUMAN\|Q3KQU3_S366 | 0.586289071 | 3.46E-07 | 1.10E-05 |
| RAI14_HUMAN\|Q9P0K7_S296 | 0.586181383 | 1.38E-04 | 4.02E-03 |
| FCHO2_HUMAN\|Q0JRZ9_S387 | 0.586068506 | 1.39E-03 | 1.20E-02 |
| ES8L1_HUMAN\|Q8TE68_S182_T187 | 0.585977881 | 6.41E-07 | 1.84E-05 |
| ASPP2_HUMAN\|Q13625_S558 | 0.585811152 | 1.77E-06 | 6.22E-05 |
| OSB11_HUMAN\|Q9BXB4_S181 | 0.585767818 | 4.87E-06 | 1.90E-04 |
| ML12A_HUMAN\|P19105_S19 | 0.585691982 | 8.42E-06 | 1.30E-04 |
| TGS1_HUMAN\|Q96RS0_S55_T60 | 0.585579415 | 4.12E-07 | 1.25E-05 |
| ZCCHV_HUMAN\|Q7Z2W4_S378 | 0.585572283 | 2.41E-07 | 3.80E-05 |
| CAMP2_HUMAN\|Q08AD1_S1313 | 0.585534487 | 2.85E-05 | 3.32E-04 |
| F193A_HUMAN\|P78312_S1144 | 0.5855114 | 1.62E-05 | 8.30E-04 |
| LAD1_HUMAN\|O00515_S485 | 0.585510142 | 9.42E-05 | 2.82E-03 |
| DBNL_HUMAN\|Q9UJU6_S275 | 0.585436194 | 3.84E-04 | 2.75E-03 |
| PDLI2_HUMAN\|Q96JY6_S134 | 0.585272685 | 7.81E-08 | 1.46E-05 |
| PRAG1_HUMAN\|Q86YV5_S907 | 0.585233308 | 7.05E-06 | 2.29E-04 |
| DC1L2_HUMAN\|O43237_S383 | 0.585166898 | 2.38E-05 | 5.67E-04 |
| FA83G_HUMAN\|A6ND36_S726 | 0.584954164 | 1.01E-04 | 9.21E-04 |
| MEF2D_HUMAN\|Q14814_S231 | 0.584929501 | 9.82E-04 | 1.55E-02 |
| S38A2_HUMAN\|Q96QD8_S17_S22 | 0.58480932 | 5.85E-04 | 3.94E-03 |
| PDLI1_HUMAN\|O00151_S130 | 0.584693733 | 1.09E-05 | 6.16E-04 |
| SAFB1_HUMAN\|Q15424_S247 | 0.584634184 | 5.83E-06 | 9.97E-05 |
| SLK_HUMAN\|Q9H2G2_S779 | 0.584600298 | 1.82E-03 | 1.53E-02 |
| VIP2_HUMAN\|O43314_S1006_S1013_S1016 | 0.584550204 | 7.40E-07 | 8.87E-05 |
| CRTC3_HUMAN\|Q6UUV7_S370 | 0.584350704 | 2.64E-04 | 2.03E-03 |
| DDI2_HUMAN\|Q5TDH0_S106 | 0.584251067 | 2.30E-06 | 2.12E-04 |
| FLNB_HUMAN\|O75369-8_S1474 | 0.584196175 | 5.10E-03 | 4.45E-02 |
| DC1L2_HUMAN\|O43237_S446 | 0.584101 | 2.93E-03 | 1.31E-02 |
| PDLI2_HUMAN\|Q96JY6_S137 | 0.584098617 | 6.03E-06 | 4.36E-04 |
| R3HD1_HUMAN\|Q15032_S973 | 0.584082436 | 1.17E-06 | 4.59E-05 |
| GPBP1_HUMAN\|Q86WP2_S381 | 0.584001158 | 4.84E-05 | 1.69E-03 |
| MTAP2_HUMAN\|P11137_T1592_T1605 | 0.583956572 | 5.73E-03 | 2.55E-02 |

| | | | |
|---|---|---|---|
| PKN2_HUMAN\|Q16513_S362 | 0.583907116 | 3.29E-05 | 7.21E-04 |
| K2C6A_HUMAN\|P02538_S541 | 0.583756468 | 8.37E-03 | 3.49E-02 |
| EXOC7_HUMAN\|Q9UPT5_S250 | 0.583660424 | 1.24E-05 | 6.74E-04 |
| SHRM2_HUMAN\|Q13796_S1039 | 0.583530538 | 1.23E-08 | 5.53E-06 |
| CCAR2_HUMAN\|Q8N163_S678 | 0.583497783 | 4.62E-03 | 3.83E-02 |
| TT39B_HUMAN\|Q5VTQ0_S120 | 0.583414789 | 4.59E-04 | 3.21E-03 |
| LIPA1_HUMAN\|Q13136_S1172 | 0.583339454 | 1.86E-06 | 3.95E-05 |
| CAMP2_HUMAN\|Q08AD1_S691 | 0.58318092 | 2.68E-04 | 2.06E-03 |
| K2C6A_HUMAN\|P02538_S58 | 0.583111556 | 8.17E-03 | 4.60E-02 |
| AHNK2_HUMAN\|Q8IVF2_S1712 | 0.583069862 | 1.42E-05 | 2.95E-04 |
| K1143_HUMAN\|Q96AT1_S146 | 0.583051951 | 3.15E-04 | 3.94E-03 |
| ARH40_HUMAN\|Q8TER5_S262 | 0.582975516 | 3.22E-06 | 6.26E-05 |
| CO052_HUMAN\|Q6ZUT6_S55 | 0.582867536 | 1.53E-03 | 1.35E-02 |
| PAXI_HUMAN\|P49023_S106 | 0.582760212 | 4.13E-05 | 4.57E-03 |
| PARD3_HUMAN\|Q8TEW0_S187 | 0.582730411 | 4.74E-03 | 4.54E-02 |
| AHNK_HUMAN\|Q09666_S5031 | 0.582698651 | 5.46E-05 | 1.04E-03 |
| K1143_HUMAN\|Q96AT1_S146 | 0.582687987 | 3.11E-04 | 6.74E-03 |
| ASPC1_HUMAN\|Q9BZE9_S275 | 0.582571044 | 9.55E-06 | 1.44E-03 |
| ATAT_HUMAN\|Q5SQI0_S315 | 0.582569167 | 4.11E-07 | 1.25E-05 |
| RRBP1_HUMAN\|Q9P2E9_S615 | 0.582530719 | 2.19E-05 | 8.24E-04 |
| TBC15_HUMAN\|Q8TC07_S70 | 0.582410138 | 2.56E-03 | 4.03E-02 |
| ETV3_HUMAN\|P41162_S139 | 0.582372478 | 3.24E-03 | 2.31E-02 |
| K1671_HUMAN\|Q9BY89_S404 | 0.582261883 | 9.40E-07 | 4.80E-05 |
| SON_HUMAN\|P18583_S94 | 0.582248392 | 7.87E-07 | 4.96E-05 |
| ROA1_HUMAN\|P09651_S364 | 0.582216138 | 2.32E-03 | 1.22E-02 |
| TPR_HUMAN\|P12270_S2155 | 0.582125032 | 2.88E-05 | 3.36E-04 |
| NDRG1_HUMAN\|Q92597_T366 | 0.582066404 | 6.27E-05 | 2.02E-03 |
| NEK9_HUMAN\|Q8TD19_T333 | 0.581978558 | 4.83E-04 | 9.21E-03 |
| LYRIC_HUMAN\|Q86UE4_S298 | 0.581855036 | 1.77E-03 | 1.90E-02 |
| LDB1_HUMAN\|Q86U70_S302 | 0.581806993 | 2.02E-03 | 2.09E-02 |
| IRS2_HUMAN\|Q9Y4H2_S606 | 0.581800032 | 1.19E-03 | 1.72E-02 |

| | | | |
|---|---|---|---|
| SWAHC_HUMAN\|Q53LP3_S212 | 0.581790095 | 1.05E-06 | 2.58E-05 |
| PDLI1_HUMAN\|O00151_S130 | 0.58176694 | 9.47E-06 | 4.62E-04 |
| VIME_HUMAN\|P08670_S56 | 0.581700388 | 3.12E-06 | 2.48E-04 |
| SRRM1_HUMAN\|Q8IYB3_S738_S740_S743 | 0.58168476 | 4.97E-06 | 8.86E-05 |
| PKN2_HUMAN\|Q16513_S360 | 0.581535408 | 1.04E-04 | 1.76E-03 |
| SCRIB_HUMAN\|Q14160_S1475_S1486 | 0.58152877 | 1.36E-03 | 3.40E-02 |
| K1C17_HUMAN\|Q04695_S11_S13 | 0.581402802 | 7.14E-05 | 1.26E-03 |
| PKN2_HUMAN\|Q16513_S360 | 0.581390658 | 1.03E-04 | 1.63E-03 |
| NIPA_HUMAN\|Q86WB0_S408_S409 | 0.581324772 | 2.91E-07 | 2.02E-05 |
| ZCHC8_HUMAN\|Q6NZY4_S500_T492_T496 | 0.581319177 | 5.95E-05 | 2.17E-03 |
| PALLD_HUMAN\|Q8WX93_S893 | 0.581315357 | 6.03E-05 | 8.86E-04 |
| LIPA3_HUMAN\|O75145_S1164 | 0.581259638 | 6.98E-06 | 2.35E-04 |
| NGAP_HUMAN\|Q9UJF2_S954 | 0.581010761 | 3.25E-05 | 1.63E-03 |
| PAXI_HUMAN\|P49023_S250 | 0.581008985 | 1.14E-03 | 3.03E-02 |
| TCOF_HUMAN\|Q13428_T249 | 0.581007294 | 4.26E-05 | 1.88E-03 |
| WDR7_HUMAN\|Q9Y4E6_S1153 | 0.580996163 | 1.76E-03 | 1.41E-02 |
| SCRIB_HUMAN\|Q14160_S853 | 0.580945693 | 9.40E-04 | 5.77E-03 |
| IRS2_HUMAN\|Q9Y4H2_S606 | 0.58092327 | 1.16E-03 | 1.10E-02 |
| CDK13_HUMAN\|Q14004_T1056 | 0.580905917 | 3.62E-05 | 1.69E-03 |
| FCHO2_HUMAN\|Q0JRZ9_S387 | 0.580716598 | 1.19E-03 | 7.01E-03 |
| PDLI2_HUMAN\|Q96JY6_S134_S137_S143 | 0.580641523 | 7.38E-05 | 6.25E-03 |
| COHA1_HUMAN\|Q9UMD9_S76_S79 | 0.580498263 | 2.70E-03 | 2.98E-02 |
| SRP09_HUMAN\|P49458_T85 | 0.580460021 | 2.08E-05 | 2.57E-04 |
| NDRG1_HUMAN\|Q92597_S326_S330_S333 | 0.58045242 | 4.38E-05 | 1.58E-03 |
| STIM2_HUMAN\|Q9P246_S522 | 0.580443079 | 1.51E-03 | 1.99E-02 |
| ITB4_HUMAN\|P16144_S1069 | 0.580340558 | 2.47E-04 | 3.31E-03 |
| TGS1_HUMAN\|Q96RS0_S55_T60 | 0.580322155 | 3.11E-07 | 1.05E-05 |
| ABLM3_HUMAN\|O94929_S372_S379 | 0.580315193 | 6.68E-05 | 1.24E-03 |
| TR150_HUMAN\|Q9Y2W1_T874 | 0.580269937 | 7.64E-05 | 2.38E-03 |
| AFAD_HUMAN\|P55196_S1696 | 0.580221187 | 2.53E-05 | 3.02E-04 |
| FA83B_HUMAN\|Q5T0W9_S664 | 0.580212183 | 3.06E-06 | 2.68E-04 |

| | | | |
|---|---|---|---|
| KS6A1_HUMAN\|Q15418_S363_S369 | 0.580094036 | 2.72E-03 | 3.10E-02 |
| AHNK_HUMAN\|Q09666_S210 | 0.58000525 | 4.77E-07 | 6.47E-05 |
| SARG_HUMAN\|Q9BW04_S429 | 0.579990576 | 2.37E-04 | 7.70E-03 |
| ARG28_HUMAN\|Q8N1W1_S476_S486 | 0.579887072 | 4.83E-05 | 5.09E-04 |
| PKP2_HUMAN\|Q99959_S82 | 0.579832255 | 6.95E-03 | 4.11E-02 |
| ACAP2_HUMAN\|Q15057_S381_S384 | 0.579830321 | 1.60E-05 | 2.11E-04 |
| COHA1_HUMAN\|Q9UMD9_T182 | 0.579718169 | 2.15E-04 | 5.22E-03 |
| DDX17_HUMAN\|Q92841_S571 | 0.579712256 | 3.49E-05 | 1.70E-03 |
| K2C5_HUMAN\|P13647_S62 | 0.579702093 | 2.16E-05 | 5.46E-04 |
| EGFR_HUMAN\|P00533_S1204 | 0.579679167 | 3.96E-04 | 4.73E-03 |
| K2C5_HUMAN\|P13647_S18_S21 | 0.57960863 | 2.00E-03 | 3.45E-02 |
| MTCL1_HUMAN\|Q9Y4B5_S1679_T1675 | 0.579598213 | 1.89E-06 | 2.09E-04 |
| LMNA_HUMAN\|P02545_S616 | 0.579553308 | 1.34E-05 | 1.86E-04 |
| CREB1_HUMAN\|P16220_S271 | 0.579388003 | 2.10E-04 | 4.83E-03 |
| TIF1B_HUMAN\|Q13263_S596_S601 | 0.579363148 | 5.18E-03 | 2.05E-02 |
| WWC3_HUMAN\|Q9ULE0_T909 | 0.579283372 | 3.29E-05 | 3.81E-04 |
| LARP1_HUMAN\|Q6PKG0_S774 | 0.579281319 | 5.00E-07 | 3.52E-05 |
| ITB4_HUMAN\|P16144_S1454_S1457 | 0.579156244 | 3.12E-03 | 2.19E-02 |
| ARH40_HUMAN\|Q8TER5_S1513 | 0.57911542 | 2.83E-07 | 1.72E-05 |
| PDLI5_HUMAN\|Q96HC4_S111 | 0.578965485 | 1.11E-05 | 3.24E-04 |
| SPTB2_HUMAN\|Q01082_S2341 | 0.578690173 | 5.84E-08 | 5.97E-06 |
| EMAL4_HUMAN\|Q9HC35_S146 | 0.578596532 | 2.21E-05 | 5.38E-04 |
| LRRF2_HUMAN\|Q9Y608_S320_S324 | 0.578542285 | 5.75E-04 | 6.24E-03 |
| TAOK3_HUMAN\|Q9H2K8_S324 | 0.578521917 | 3.43E-06 | 2.91E-04 |
| CAMP3_HUMAN\|Q9P1Y5_S756 | 0.578433867 | 9.79E-06 | 3.07E-04 |
| MICA3_HUMAN\|Q7RTP6_S977 | 0.578402501 | 2.81E-08 | 6.93E-06 |
| SRRM1_HUMAN\|Q8IYB3_S738_S740_S743 | 0.578371078 | 4.21E-06 | 7.53E-05 |
| ARVC_HUMAN\|O00192_T642 | 0.578362648 | 1.36E-05 | 1.87E-04 |
| ITB4_HUMAN\|P16144_S1454_T1455 | 0.578180586 | 9.11E-04 | 9.26E-03 |
| LASP1_HUMAN\|Q14847_T104 | 0.578122624 | 4.36E-06 | 3.29E-04 |
| NSD3_HUMAN\|Q9BZ95_S457 | 0.577792432 | 5.64E-06 | 2.78E-04 |

| | | | |
|---|---|---|---|
| CO052_HUMAN\|Q6ZUT6_S193 | 0.577658995 | 2.56E-05 | 3.04E-04 |
| YAP1_HUMAN\|P46937_S109_T119 | 0.577616286 | 2.06E-07 | 7.26E-06 |
| REPS1_HUMAN\|Q96D71_S162_S170_S174 | 0.577590863 | 9.23E-06 | 1.42E-04 |
| K1C10_HUMAN\|P13645_S56 | 0.577340688 | 9.75E-03 | 3.38E-02 |
| TENS4_HUMAN\|Q8IZW8_S198 | 0.577248604 | 2.04E-03 | 1.59E-02 |
| WDR7_HUMAN\|Q9Y4E6_S1153 | 0.577208018 | 1.58E-03 | 8.00E-03 |
| SLK_HUMAN\|Q9H2G2_S372 | 0.577182898 | 5.85E-08 | 2.83E-06 |
| STK10_HUMAN\|O94804_S450_S454 | 0.576943355 | 7.70E-05 | 7.67E-04 |
| MGMT_HUMAN\|P16455_S201 | 0.576850499 | 3.17E-08 | 5.37E-06 |
| K1522_HUMAN\|Q9P206_T162 | 0.576776952 | 2.18E-04 | 1.67E-03 |
| TPR_HUMAN\|P12270_S1847_T1839 | 0.576745527 | 1.07E-03 | 2.19E-02 |
| I3L2J0_HUMAN\|I3L2J0_T472 | 0.57673669 | 1.47E-02 | 4.70E-02 |
| LMNA_HUMAN\|P02545_S22_T19 | 0.576720079 | 4.49E-03 | 3.67E-02 |
| ELAV1_HUMAN\|Q15717_S202 | 0.576685014 | 2.85E-04 | 2.05E-03 |
| HMGN1_HUMAN\|P05114_S7 | 0.576678001 | 8.85E-05 | 5.78E-03 |
| ITB4_HUMAN\|P16144_S1454_S1457 | 0.576601769 | 2.92E-03 | 1.31E-02 |
| DAPK3_HUMAN\|O43293_S311_S312 | 0.576577735 | 6.37E-06 | 2.26E-04 |
| TOP2B_HUMAN\|Q02880_S1550 | 0.576386636 | 1.12E-03 | 8.66E-03 |
| CCDC9_HUMAN\|Q9Y3X0_S376 | 0.57630851 | 6.20E-06 | 2.10E-04 |
| TBC16_HUMAN\|Q8TBP0_S103_T99 | 0.576192532 | 3.33E-05 | 1.64E-03 |
| PHLB2_HUMAN\|Q86SQ0_S402 | 0.576065737 | 1.28E-05 | 1.78E-04 |
| ROA1_HUMAN\|P09651_S363 | 0.576065418 | 5.92E-03 | 2.27E-02 |
| CCDC9_HUMAN\|Q9Y3X0_S386_T381 | 0.576059181 | 1.49E-05 | 4.01E-04 |
| K2C6B_HUMAN\|P04259_S541 | 0.576001559 | 1.95E-05 | 2.50E-04 |
| IWS1_HUMAN\|Q96ST2_S398 | 0.575851405 | 1.71E-03 | 1.83E-02 |
| RPB1_HUMAN\|P24928_S1847_T1856 | 0.575797811 | 2.87E-04 | 5.22E-03 |
| S23A2_HUMAN\|Q9UGH3_S78_S81 | 0.575681194 | 7.25E-04 | 1.18E-02 |
| OSB11_HUMAN\|Q9BXB4_S181 | 0.575569504 | 2.95E-06 | 2.42E-04 |
| NCK5L_HUMAN\|Q9HCH0_S489_S494 | 0.575531491 | 7.87E-06 | 2.47E-04 |
| TBCD5_HUMAN\|Q92609_S541 | 0.575493814 | 1.48E-08 | 5.28E-06 |
| LPP_HUMAN\|Q93052_S609 | 0.575363908 | 2.25E-04 | 2.53E-03 |

| | | | |
|---|---|---|---|
| PKP3_HUMAN\|Q9Y446_S183 | 0.575333079 | 6.49E-05 | 1.17E-03 |
| K2C5_HUMAN\|P13647_S62 | 0.575324241 | 1.77E-05 | 2.30E-04 |
| CR025_HUMAN\|Q96B23_S66 | 0.575293502 | 1.07E-06 | 4.28E-05 |
| EGFR_HUMAN\|P00533_S991 | 0.575013762 | 1.06E-05 | 2.43E-04 |
| NUCKS_HUMAN\|Q9H1E3_S75_S79 | 0.574911403 | 2.31E-03 | 2.81E-02 |
| 1B07_HUMAN\|P01889_S352 | 0.574891038 | 8.78E-05 | 8.19E-04 |
| AR6P4_HUMAN\|Q66PJ3_S241 | 0.574831268 | 1.16E-04 | 3.52E-03 |
| K2C5_HUMAN\|P13647_S18_S21 | 0.574676447 | 1.75E-03 | 2.34E-02 |
| LDLR_HUMAN\|P01130_S854 | 0.574636802 | 1.68E-03 | 1.19E-02 |
| CCDC9_HUMAN\|Q9Y3X0_S376_S386_T381 | 0.574585671 | 1.03E-05 | 5.98E-04 |
| TB10B_HUMAN\|Q4KMP7_S22 | 0.574392565 | 4.48E-06 | 1.80E-04 |
| GOGA4_HUMAN\|Q13439_S89_S92 | 0.574206125 | 1.51E-05 | 2.04E-04 |
| SIK3_HUMAN\|Q9Y2K2_S493 | 0.574100693 | 6.38E-07 | 1.72E-05 |
| NFIC_HUMAN\|P08651_S284_S304_T293 | 0.574049415 | 4.03E-05 | 6.48E-04 |
| SARG_HUMAN\|Q9BW04_S502 | 0.573852239 | 9.04E-07 | 1.02E-04 |
| JMY_HUMAN\|Q8N9B5_S115 | 0.573801291 | 2.07E-04 | 5.17E-03 |
| CR025_HUMAN\|Q96B23_S143_S145 | 0.573799936 | 2.01E-06 | 1.35E-04 |
| PDLI2_HUMAN\|Q96JY6_S134_S137_S143 | 0.573797089 | 5.70E-05 | 8.61E-04 |
| SPTB2_HUMAN\|Q01082-3_S8 | 0.5737683 | 3.46E-05 | 3.89E-04 |
| RBM27_HUMAN\|Q9P2N5_T447 | 0.573711232 | 1.71E-05 | 2.23E-04 |
| PP14A_HUMAN\|Q96A00_S128_S134_S136 | 0.573388291 | 1.25E-06 | 3.08E-05 |
| SRGP1_HUMAN\|Q7Z6B7_S906 | 0.573343194 | 2.28E-05 | 2.81E-04 |
| TENS4_HUMAN\|Q8IZW8_S248_S253 | 0.573337419 | 5.45E-04 | 3.40E-03 |
| K1522_HUMAN\|Q9P206_S929 | 0.573312333 | 8.93E-07 | 2.06E-04 |
| BIG1_HUMAN\|Q9Y6D6_S397 | 0.573299607 | 4.32E-06 | 7.69E-05 |
| TBD2A_HUMAN\|Q9BYX2_S920 | 0.573219908 | 5.11E-06 | 1.79E-04 |
| RAI14_HUMAN\|Q9P0K7_T251 | 0.573201372 | 5.64E-05 | 2.07E-03 |
| K1C16_HUMAN\|P08779_S449 | 0.573140381 | 7.12E-05 | 6.86E-04 |
| EGFR_HUMAN\|P00533_S695 | 0.5731092 | 7.87E-04 | 6.67E-03 |
| NDRG2_HUMAN\|Q9UN36_S332_T330 | 0.573000075 | 5.36E-06 | 3.63E-04 |
| DC1L1_HUMAN\|Q9Y6G9_S421 | 0.572979495 | 1.16E-03 | 1.68E-02 |

| | | | |
|---|---|---|---|
| GYS1_HUMAN\|P13807_T703 | 0.572884855 | 2.48E-05 | 1.27E-03 |
| K2C75_HUMAN\|O95678_S40 | 0.572847482 | 1.47E-03 | 1.31E-02 |
| M3K20_HUMAN\|Q9NYL2_S727 | 0.572843691 | 2.53E-07 | 3.92E-05 |
| DDX3X_HUMAN\|O00571_S612 | 0.57269914 | 3.35E-03 | 1.64E-02 |
| COHA1_HUMAN\|Q9UMD9_S172_T176_T182 | 0.572682624 | 3.70E-03 | 1.58E-02 |
| COHA1_HUMAN\|Q9UMD9_S61_S62_S70 | 0.572671078 | 2.30E-03 | 4.61E-02 |
| MTUS1_HUMAN\|Q9ULD2_T1204 | 0.572557803 | 8.35E-06 | 1.99E-04 |
| ERCC5_HUMAN\|P28715_S382_S384 | 0.572423998 | 2.75E-04 | 5.89E-03 |
| YBOX3_HUMAN\|P16989_S369_T366 | 0.572351365 | 3.71E-06 | 6.94E-05 |
| LIPA1_HUMAN\|Q13136_S666 | 0.572338267 | 6.58E-06 | 1.08E-04 |
| K2C5_HUMAN\|P13647_S11_S8 | 0.572321543 | 2.85E-06 | 1.19E-04 |
| MAP1B_HUMAN\|P46821_S1427 | 0.572279864 | 6.21E-07 | 3.50E-05 |
| YBOX3_HUMAN\|P16989_S369_T366 | 0.572207394 | 3.69E-06 | 6.77E-05 |
| DNJC1_HUMAN\|Q96KC8_S381 | 0.572192452 | 1.91E-03 | 2.02E-02 |
| SRRM1_HUMAN\|Q8IYB3_T846 | 0.572122074 | 2.48E-04 | 1.94E-03 |
| DNJC1_HUMAN\|Q96KC8_S381 | 0.572107526 | 1.91E-03 | 1.32E-02 |
| YAP1_HUMAN\|P46937_S163 | 0.572083065 | 5.74E-04 | 6.23E-03 |
| PRKDC_HUMAN\|P78527_S3205 | 0.572062281 | 6.16E-04 | 3.71E-03 |
| T2FA_HUMAN\|P35269_T389 | 0.572032481 | 2.16E-04 | 4.23E-03 |
| RAI14_HUMAN\|Q9P0K7_S281_S290 | 0.572030928 | 9.76E-04 | 7.79E-03 |
| NDRG1_HUMAN\|Q92597_S333_S336_T328 | 0.571810649 | 1.03E-06 | 5.16E-05 |
| SRP09_HUMAN\|P49458_T85 | 0.571802718 | 1.43E-05 | 1.92E-04 |
| YBOX1_HUMAN\|P67809_S165_S176 | 0.571800182 | 7.88E-06 | 1.24E-04 |
| CND1_HUMAN\|Q15021_S1330_S1333 | 0.571793249 | 1.82E-03 | 8.91E-03 |
| PATL1_HUMAN\|Q86TB9_S179_S184 | 0.571780762 | 1.57E-04 | 4.31E-03 |
| CNKR1_HUMAN\|Q969H4_S305_S314 | 0.571757463 | 6.90E-05 | 2.37E-03 |
| ARH40_HUMAN\|Q8TER5_T371 | 0.571684323 | 1.90E-07 | 1.81E-05 |
| RENT1_HUMAN\|Q92900_S1127 | 0.571456677 | 8.02E-03 | 4.39E-02 |
| NUFP2_HUMAN\|Q7Z417_S212_S214 | 0.571436547 | 2.20E-05 | 5.36E-04 |
| K1522_HUMAN\|Q9P206_S406 | 0.571413142 | 3.48E-05 | 3.98E-04 |
| NIBL1_HUMAN\|Q96TA1_S691_S696 | 0.571308794 | 1.03E-05 | 1.86E-03 |

| | | | |
|---|---|---|---|
| KPCD2_HUMAN\|Q9BZL6_S212 | 0.571255491 | 4.48E-05 | 2.07E-03 |
| ACINU_HUMAN\|Q9UKV3_T563 | 0.571207131 | 1.90E-07 | 1.29E-05 |
| K1C14_HUMAN\|P02533_Y46 | 0.571206166 | 4.47E-05 | 8.96E-04 |
| MAP4_HUMAN\|P27816_S1073 | 0.571176755 | 7.43E-06 | 4.63E-04 |
| PHLB1_HUMAN\|Q86UU1_S489 | 0.571098531 | 2.90E-04 | 6.08E-03 |
| TBCD4_HUMAN\|O60343_S318 | 0.570969852 | 7.36E-06 | 2.36E-04 |
| MILK1_HUMAN\|Q8N3F8_S578 | 0.570963234 | 1.07E-06 | 5.33E-05 |
| PHLB1_HUMAN\|Q86UU1_S489 | 0.570933661 | 2.88E-04 | 3.72E-03 |
| VIME_HUMAN\|P08670_S459 | 0.570901708 | 1.81E-06 | 1.72E-04 |
| AHNK_HUMAN\|Q09666_S5099 | 0.570772962 | 4.23E-04 | 8.27E-03 |
| PLXA2_HUMAN\|O75051_S1630_S1633 | 0.570733363 | 4.99E-05 | 5.08E-03 |
| DBNL_HUMAN\|Q9UJU6_S275 | 0.570707744 | 2.39E-04 | 5.31E-03 |
| LRRF2_HUMAN\|Q9Y608_S320_S324 | 0.570643814 | 4.46E-04 | 3.13E-03 |
| RHG29_HUMAN\|Q52LW3_S176 | 0.570578372 | 3.95E-04 | 6.54E-03 |
| PKN2_HUMAN\|Q16513_S362 | 0.570531989 | 1.90E-05 | 9.29E-04 |
| KS6A1_HUMAN\|Q15418_S363_S369 | 0.570388072 | 2.14E-03 | 2.67E-02 |
| SEPT7_HUMAN\|Q16181_T426 | 0.57029655 | 7.04E-04 | 7.42E-03 |
| K1671_HUMAN\|Q9BY89_S127 | 0.569957042 | 3.32E-06 | 6.38E-05 |
| SEC62_HUMAN\|Q99442_T375 | 0.569916612 | 1.57E-03 | 1.70E-02 |
| BNIP2_HUMAN\|Q12982_S114 | 0.569907309 | 1.63E-04 | 3.08E-03 |
| IF4G2_HUMAN\|P78344_T508 | 0.569863704 | 1.08E-07 | 4.51E-06 |
| TPPP_HUMAN\|O94811_S160 | 0.5698401 | 2.44E-04 | 1.82E-03 |
| MRE11_HUMAN\|P49959_S649 | 0.569822233 | 2.12E-04 | 1.63E-03 |
| PAK2_HUMAN\|Q13177_S141 | 0.569587909 | 7.39E-05 | 7.06E-04 |
| PRC2A_HUMAN\|P48634_S1325_S1328_T1323 | 0.569571479 | 4.47E-03 | 4.36E-02 |
| GPBP1_HUMAN\|Q86WP2_S381 | 0.569502185 | 2.64E-05 | 1.46E-03 |
| PTN13_HUMAN\|Q12923_S908 | 0.569451369 | 2.72E-05 | 3.18E-04 |
| DCBD2_HUMAN\|Q96PD2_S726_S727 | 0.569380233 | 6.22E-04 | 6.68E-03 |
| 3BP1_HUMAN\|Q9Y3L3_S596 | 0.56936104 | 1.13E-05 | 3.39E-04 |
| KI26B_HUMAN\|Q2KJY2_S1006_S1018_S1021 | 0.569209639 | 3.07E-07 | 1.83E-05 |
| F193A_HUMAN\|P78312_S1144 | 0.569176317 | 7.45E-06 | 1.19E-04 |

| | | | |
|---|---|---|---|
| AHNK_HUMAN\|Q09666_S5749_S5752 | 0.569171216 | 2.33E-06 | 2.14E-04 |
| PCSK9_HUMAN\|Q8NBP7_S688 | 0.568876945 | 4.02E-05 | 1.64E-03 |
| LIPA1_HUMAN\|Q13136_S666 | 0.568738284 | 5.57E-06 | 2.08E-04 |
| YAP1_HUMAN\|P46937_S127 | 0.568550975 | 4.53E-04 | 2.92E-03 |
| CE170_HUMAN\|Q5SW79_S636 | 0.568442497 | 7.35E-09 | 5.95E-07 |
| PLEC_HUMAN\|Q15149_T4030 | 0.568439975 | 3.44E-03 | 2.37E-02 |
| SCRIB_HUMAN\|Q14160_S1475 | 0.568420498 | 3.55E-04 | 3.61E-03 |
| BIG1_HUMAN\|Q9Y6D6_S397 | 0.568238576 | 3.37E-06 | 3.32E-04 |
| BRD4_HUMAN\|O60885_S470 | 0.568234336 | 6.16E-05 | 6.09E-04 |
| NDK8_HUMAN\|O60361_S105 | 0.568231881 | 7.68E-06 | 1.21E-04 |
| SOS2_HUMAN\|Q07890_S1267 | 0.568223008 | 1.81E-04 | 4.38E-03 |
| SPF30_HUMAN\|O75940_S201 | 0.568191839 | 1.06E-04 | 2.48E-03 |
| T22D4_HUMAN\|Q9Y3Q8_T167 | 0.5681339 | 1.42E-05 | 4.02E-04 |
| DC1L2_HUMAN\|O43237_S383 | 0.568102235 | 1.18E-05 | 3.41E-04 |
| LAP2A_HUMAN\|P42166_S159 | 0.568040589 | 1.28E-05 | 1.80E-04 |
| 4EBP1_HUMAN\|Q13541_T70 | 0.568039933 | 1.32E-05 | 7.29E-04 |
| IRS2_HUMAN\|Q9Y4H2_S679 | 0.567953393 | 7.92E-06 | 1.24E-04 |
| ABLM3_HUMAN\|O94929_S373_S379 | 0.56789604 | 8.49E-05 | 1.50E-03 |
| Q659E9_HUMAN\|Q659E9_S168 | 0.567856843 | 2.64E-07 | 1.67E-05 |
| BORG4_HUMAN\|Q9H3Q1_S64 | 0.567724346 | 4.46E-06 | 1.80E-04 |
| SI1L1_HUMAN\|O43166_S1255 | 0.567681193 | 9.55E-05 | 8.80E-04 |
| DLG1_HUMAN\|Q12959_S573 | 0.56756549 | 6.55E-04 | 1.11E-02 |
| STK10_HUMAN\|O94804_S448_S455 | 0.567492085 | 2.90E-06 | 5.80E-05 |
| TRI29_HUMAN\|Q14134_S152 | 0.567435801 | 1.18E-05 | 1.69E-04 |
| SMTN_HUMAN\|P53814_S357 | 0.567392225 | 3.86E-03 | 4.09E-02 |
| K1C14_HUMAN\|P02533_S44_S53_S54 | 0.567377461 | 5.06E-06 | 8.96E-05 |
| AHNK_HUMAN\|Q09666_T4564 | 0.567339492 | 3.56E-05 | 3.96E-04 |
| CDC23_HUMAN\|Q9UJX2_T596 | 0.567330984 | 4.70E-03 | 2.57E-02 |
| WAC2A_HUMAN\|Q641Q2_S390 | 0.567188664 | 9.45E-05 | 1.52E-03 |
| OSB11_HUMAN\|Q9BXB4_S174_S181 | 0.566828748 | 8.80E-04 | 5.51E-03 |
| KIZ_HUMAN\|Q2M2Z5_S623 | 0.56682005 | 1.69E-06 | 8.71E-05 |

| | | | |
|---|---|---|---|
| RBP10_HUMAN\|Q6VN20_S365 | 0.56679869 | 5.96E-07 | 7.69E-05 |
| ZDHC5_HUMAN\|Q9C0B5_S704 | 0.5667683 | 6.21E-04 | 4.15E-03 |
| NDRG1_HUMAN\|Q92597_S330_T328 | 0.5666125 | 1.22E-05 | 6.71E-04 |
| KI16B_HUMAN\|Q96L93_S662 | 0.566397386 | 3.23E-06 | 1.41E-04 |
| PML_HUMAN\|P29590_T867 | 0.566371968 | 7.43E-06 | 1.19E-04 |
| UBXN7_HUMAN\|O94888_S280_S288 | 0.566205418 | 6.22E-04 | 4.15E-03 |
| BAG3_HUMAN\|O95817_S289_T285 | 0.566081916 | 4.65E-05 | 5.01E-04 |
| BAIP2_HUMAN\|Q9UQB8_T360 | 0.566073395 | 2.36E-03 | 1.10E-02 |
| CR025_HUMAN\|Q96B23_S143_S145_S147 | 0.566066614 | 5.59E-05 | 1.08E-03 |
| PKP3_HUMAN\|Q9Y446_S280 | 0.566040263 | 3.16E-04 | 9.79E-03 |
| RAI14_HUMAN\|Q9P0K7_S290 | 0.565997683 | 2.44E-04 | 5.86E-03 |
| COHA1_HUMAN\|Q9UMD9_S85_S93_S96 | 0.565912409 | 1.55E-05 | 4.13E-04 |
| RGL2_HUMAN\|O15211_S761 | 0.565900439 | 9.94E-06 | 3.03E-04 |
| K1C16_HUMAN\|P08779_S468 | 0.565848653 | 8.41E-06 | 1.32E-04 |
| PAXI_HUMAN\|P49023_S103_S106 | 0.565785219 | 2.70E-04 | 1.32E-02 |
| NIPA_HUMAN\|Q86WB0_S408_S409 | 0.565771485 | 1.30E-07 | 5.22E-06 |
| NDRG1_HUMAN\|Q92597_S330_S333 | 0.565699501 | 4.19E-06 | 7.52E-05 |
| SMAG2_HUMAN\|Q5PRF9_S271 | 0.565664131 | 2.11E-04 | 2.40E-03 |
| WRIP1_HUMAN\|Q96S55_T116 | 0.565643255 | 1.86E-03 | 1.01E-02 |
| BORG5_HUMAN\|Q00587_S303 | 0.56563852 | 3.81E-06 | 1.62E-04 |
| AHNK2_HUMAN\|Q8IVF2_S294_T298 | 0.565638495 | 2.87E-05 | 1.25E-03 |
| GYS1_HUMAN\|P13807_T703 | 0.565548296 | 1.84E-05 | 7.43E-04 |
| JMY_HUMAN\|Q8N9B5_S115 | 0.565346616 | 1.52E-04 | 1.25E-03 |
| SEPT9_HUMAN\|Q9UHD8_T42 | 0.565289784 | 2.80E-06 | 2.28E-04 |
| PEA15_HUMAN\|Q15121_S116 | 0.565222612 | 1.76E-04 | 6.14E-03 |
| RAE1_HUMAN\|P24386_S651 | 0.565219659 | 1.13E-06 | 1.13E-04 |
| ANK3_HUMAN\|Q12955_S959 | 0.565145829 | 6.50E-04 | 3.88E-03 |
| VIME_HUMAN\|P08670_S34 | 0.565001733 | 1.48E-05 | 4.01E-04 |
| PDLI1_HUMAN\|O00151_S130 | 0.564992849 | 4.16E-06 | 1.73E-04 |
| RNF25_HUMAN\|Q96BH1_S450 | 0.564984439 | 5.28E-05 | 5.39E-04 |
| MYPT1_HUMAN\|O14974_S554 | 0.564866872 | 1.69E-03 | 2.15E-02 |

| | | | |
|---|---|---|---|
| K1C17_HUMAN\|Q04695_S17 | 0.564772743 | 3.79E-04 | 2.56E-03 |
| MILK1_HUMAN\|Q8N3F8_S644 | 0.564713819 | 1.02E-05 | 1.49E-04 |
| I2BPL_HUMAN\|Q9H1B7_S659 | 0.56471232 | 3.65E-03 | 1.56E-02 |
| PTN13_HUMAN\|Q12923_S908 | 0.56431505 | 2.20E-05 | 2.74E-04 |
| CRBG1_HUMAN\|Q9Y4K1_S22 | 0.564280589 | 5.30E-07 | 7.18E-05 |
| AHNK_HUMAN\|Q09666_T4430 | 0.564218284 | 2.53E-05 | 5.96E-04 |
| SLAI2_HUMAN\|Q9P270_S349_S353_S357 | 0.56420458 | 1.45E-06 | 3.48E-05 |
| AAK1_HUMAN\|Q2M2I8_S21 | 0.564118062 | 3.90E-05 | 4.27E-04 |
| IRS2_HUMAN\|Q9Y4H2_S1186_S1203 | 0.56403569 | 6.93E-06 | 5.15E-04 |
| KC1D_HUMAN\|P48730_S328 | 0.563999825 | 3.32E-04 | 2.44E-03 |
| K2C6B_HUMAN\|P04259_S34 | 0.56393339 | 8.42E-04 | 4.79E-03 |
| PAXI_HUMAN\|P49023_S250 | 0.563891242 | 6.85E-04 | 7.24E-03 |
| ERCC5_HUMAN\|P28715_S382_S384 | 0.563796244 | 2.05E-04 | 6.92E-03 |
| K2C5_HUMAN\|P13647_S26 | 0.563766071 | 7.80E-06 | 1.23E-04 |
| COBL1_HUMAN\|Q53SF7_T298 | 0.563535197 | 2.68E-05 | 6.24E-04 |
| TR150_HUMAN\|Q9Y2W1_T874 | 0.563401357 | 3.98E-05 | 4.41E-04 |
| CREB1_HUMAN\|P16220_S271 | 0.563232114 | 1.15E-04 | 1.07E-03 |
| AAK1_HUMAN\|Q2M2I8_S21 | 0.562917411 | 3.73E-05 | 4.23E-04 |
| EIF3H_HUMAN\|O15372_S183 | 0.562861257 | 3.39E-04 | 2.34E-03 |
| MAP4_HUMAN\|P27816_S928 | 0.562840303 | 8.28E-06 | 2.58E-04 |
| OSB11_HUMAN\|Q9BXB4_S174_S181 | 0.56283131 | 7.84E-04 | 1.33E-02 |
| MTCL1_HUMAN\|Q9Y4B5_S1679_T1675 | 0.562529949 | 8.37E-07 | 9.86E-05 |
| DC1L2_HUMAN\|O43237_S407 | 0.562504149 | 5.41E-06 | 3.81E-04 |
| SRRM1_HUMAN\|Q8IYB3_S738_S743 | 0.562464103 | 1.74E-03 | 3.10E-02 |
| ADRM1_HUMAN\|Q16186_S211 | 0.561994329 | 1.42E-05 | 1.91E-04 |
| PKP1_HUMAN\|Q13835_S191 | 0.561943353 | 1.86E-03 | 9.04E-03 |
| K1C14_HUMAN\|P02533_S12 | 0.561774826 | 2.34E-06 | 5.01E-05 |
| AHNK_HUMAN\|Q09666_S5752_S5763 | 0.561689561 | 1.19E-04 | 1.05E-03 |
| NELFE_HUMAN\|P18615_S353 | 0.561687246 | 3.72E-08 | 2.22E-06 |
| RGS3_HUMAN\|P49796_S1044 | 0.561681755 | 1.90E-06 | 9.51E-05 |
| NUFP2_HUMAN\|Q7Z417_S212_S214 | 0.561550634 | 1.46E-05 | 8.97E-04 |

| | | | |
|---|---|---|---|
| PPR35_HUMAN\|Q8TAP8_S52 | 0.561534635 | 1.77E-05 | 4.71E-04 |
| XCT_HUMAN\|Q9UPY5_S26 | 0.561534014 | 1.55E-04 | 3.96E-03 |
| RHG23_HUMAN\|Q9P227_S361_S372 | 0.561491239 | 6.54E-08 | 8.25E-06 |
| CCM2_HUMAN\|Q9BSQ5_S384 | 0.561385612 | 1.34E-05 | 1.85E-04 |
| OSBL1_HUMAN\|Q9BXW6_S472_S481 | 0.561289121 | 8.78E-03 | 4.10E-02 |
| IRS2_HUMAN\|Q9Y4H2_S1186_S1203 | 0.56127375 | 6.16E-06 | 4.39E-04 |
| RHG21_HUMAN\|Q5T5U3_S615 | 0.561203664 | 1.11E-06 | 5.45E-05 |
| R3HD2_HUMAN\|Q9Y2K5_S349 | 0.561140447 | 3.39E-03 | 3.34E-02 |
| COHA1_HUMAN\|Q9UMD9_S85_S93_S96 | 0.56110356 | 1.25E-05 | 1.75E-04 |
| K2C5_HUMAN\|P13647_S16_S26 | 0.561043261 | 1.08E-03 | 1.04E-02 |
| MAST4_HUMAN\|O15021_S2555 | 0.560863196 | 3.38E-05 | 3.89E-04 |
| OSB11_HUMAN\|Q9BXB4_S181 | 0.5607817 | 1.40E-06 | 3.21E-05 |
| ARHG5_HUMAN\|Q12774_S445_S450 | 0.560581675 | 1.24E-07 | 1.12E-04 |
| PHLB2_HUMAN\|Q86SQ0_S330_S334 | 0.560481586 | 3.01E-04 | 3.83E-03 |
| CLIP1_HUMAN\|P30622_S200 | 0.560451708 | 1.83E-04 | 1.44E-03 |
| TBCD4_HUMAN\|O60343_S570 | 0.560450104 | 1.53E-04 | 2.29E-03 |
| K2C5_HUMAN\|P13647_S573 | 0.560165006 | 3.62E-03 | 4.87E-02 |
| DPYL3_HUMAN\|Q14195_S568 | 0.559987645 | 5.15E-04 | 3.53E-03 |
| K1671_HUMAN\|Q9BY89_S128 | 0.559943038 | 2.56E-07 | 8.84E-06 |
| PKN2_HUMAN\|Q16513_S362 | 0.559878881 | 1.22E-05 | 3.50E-04 |
| DCA10_HUMAN\|Q5QP82_S350 | 0.559837745 | 1.97E-05 | 3.82E-04 |
| SRS10_HUMAN\|O75494_S106 | 0.559764732 | 1.99E-04 | 4.72E-03 |
| IF4G2_HUMAN\|P78344_T508 | 0.559551888 | 5.92E-08 | 2.94E-06 |
| 3BP1_HUMAN\|Q9Y3L3_S596 | 0.559480015 | 7.46E-06 | 1.19E-04 |
| LMNA_HUMAN\|P02545_S407_S414 | 0.559422985 | 1.12E-05 | 1.59E-04 |
| K1522_HUMAN\|Q9P206_S404_S409 | 0.559370962 | 1.34E-06 | 3.27E-05 |
| DPYL2_HUMAN\|Q16555_S570 | 0.559337007 | 5.77E-04 | 1.44E-02 |
| TM230_HUMAN\|Q96A57_T25 | 0.559325765 | 2.16E-04 | 4.23E-03 |
| TBCD4_HUMAN\|O60343_S318 | 0.559268758 | 4.47E-06 | 8.14E-05 |
| IF2B_HUMAN\|P20042_T111 | 0.559255123 | 9.36E-05 | 1.64E-03 |
| OSBL8_HUMAN\|Q9BZF1_S807_S808_S810 | 0.559190505 | 6.55E-05 | 1.58E-03 |

| | | | |
|---|---|---|---|
| ENSA_HUMAN|O43768_S109 | 0.559052459 | 2.55E-05 | 3.03E-04 |
| TB182_HUMAN|Q9C0C2_S1715 | 0.558954817 | 6.89E-05 | 2.87E-03 |
| M4K4_HUMAN|O95819_S639 | 0.558855357 | 1.50E-04 | 4.23E-03 |
| MAP4_HUMAN|P27816_S941 | 0.558826818 | 1.40E-05 | 1.90E-04 |
| NOLC1_HUMAN|Q14978_S688_S698 | 0.558723309 | 1.05E-02 | 4.20E-02 |
| K2C6B_HUMAN|P04259_S37 | 0.558709029 | 2.41E-04 | 3.25E-03 |
| M4K4_HUMAN|O95819_S639 | 0.558579106 | 1.48E-04 | 1.22E-03 |
| FKB15_HUMAN|Q5T1M5_S344 | 0.558555866 | 6.64E-06 | 4.23E-04 |
| CREB1_HUMAN|P16220_S271 | 0.558407153 | 9.52E-05 | 3.15E-03 |
| FRYL_HUMAN|O94915_T1959 | 0.558396244 | 2.73E-06 | 5.59E-05 |
| MARCS_HUMAN|P29966_S118 | 0.558267185 | 2.70E-04 | 2.93E-03 |
| IRS2_HUMAN|Q9Y4H2_S1185_S1203 | 0.558174473 | 2.39E-05 | 1.12E-03 |
| NDRG1_HUMAN|Q92597_S333_S336 | 0.558168752 | 1.72E-05 | 9.93E-04 |
| PAK2_HUMAN|Q13177_S192 | 0.55814919 | 4.88E-03 | 1.96E-02 |
| GRAP1_HUMAN|Q4V328_S666_S669 | 0.558135374 | 3.93E-03 | 2.60E-02 |
| RHG29_HUMAN|Q52LW3_S176 | 0.557960467 | 2.63E-04 | 8.34E-03 |
| ZO2_HUMAN|Q9UDY2_S1159 | 0.557942546 | 7.09E-04 | 7.46E-03 |
| ARH40_HUMAN|Q8TER5_T371 | 0.55762092 | 8.96E-08 | 8.23E-06 |
| TANC2_HUMAN|Q9HCD6_S1827 | 0.557569495 | 7.20E-07 | 3.28E-05 |
| UBP5_HUMAN|P45974_S779 | 0.557533481 | 1.06E-04 | 9.51E-04 |
| SPD2A_HUMAN|Q5TCZ1_S1043 | 0.557521587 | 7.05E-05 | 2.40E-03 |
| CE170_HUMAN|Q5SW79_S1165 | 0.557521175 | 9.12E-05 | 1.61E-03 |
| SPF30_HUMAN|O75940_S201 | 0.557457321 | 7.17E-05 | 5.16E-03 |
| AFAD_HUMAN|P55196_S1799 | 0.557415774 | 4.71E-05 | 4.97E-04 |
| N4BP1_HUMAN|O75113_T242 | 0.557355475 | 2.03E-06 | 4.48E-05 |
| CND1_HUMAN|Q15021_S1333 | 0.557243891 | 3.39E-06 | 2.91E-04 |
| T2FA_HUMAN|P35269_T389 | 0.557223265 | 1.32E-04 | 4.01E-03 |
| MK03_HUMAN|P27361_T198_T202 | 0.557104624 | 3.12E-05 | 1.32E-03 |
| COHA1_HUMAN|Q9UMD9_S76_S79 | 0.557016872 | 1.47E-03 | 2.07E-02 |
| RGS3_HUMAN|P49796_S1044 | 0.556982297 | 1.49E-06 | 1.84E-04 |
| TM1L1_HUMAN|O75674_S321 | 0.556979412 | 1.91E-04 | 1.12E-02 |

| | | | |
|---|---|---|---|
| SPR2B_HUMAN\|P35325_S60 | 0.556967435 | 1.98E-03 | 2.35E-02 |
| PCSK9_HUMAN\|Q8NBP7_S688 | 0.556859948 | 2.44E-05 | 7.35E-03 |
| PKP1_HUMAN\|Q13835_S142 | 0.556846881 | 1.31E-03 | 1.21E-02 |
| ZYX_HUMAN\|Q15942_S259 | 0.55669694 | 1.25E-06 | 2.94E-05 |
| PACN3_HUMAN\|Q9UKS6_S344_S354 | 0.556680007 | 2.73E-06 | 8.54E-05 |
| MISP_HUMAN\|Q8IVT2_S394_S395 | 0.556588597 | 2.26E-06 | 7.35E-05 |
| MAP4_HUMAN\|P27816_S928 | 0.556566083 | 6.21E-06 | 1.05E-04 |
| DLG5_HUMAN\|Q8TDM6_S1264 | 0.556512391 | 6.83E-06 | 2.24E-04 |
| PKP4_HUMAN\|Q99569_S447 | 0.556357975 | 5.03E-04 | 3.48E-03 |
| IF4B_HUMAN\|P23588_S495_S497 | 0.556180723 | 5.08E-07 | 2.58E-05 |
| TPPP_HUMAN\|O94811_S160 | 0.556153442 | 1.47E-04 | 1.31E-03 |
| AHNK2_HUMAN\|Q8IVF2_S294_T298 | 0.555926182 | 1.92E-05 | 4.93E-04 |
| MISP_HUMAN\|Q8IVT2_S155 | 0.55588459 | 1.44E-04 | 1.29E-03 |
| EGFR_HUMAN\|P00533_T693 | 0.555834717 | 7.45E-05 | 1.04E-03 |
| K2C5_HUMAN\|P13647_S36 | 0.555829142 | 1.62E-05 | 4.29E-04 |
| YBOX3_HUMAN\|P16989_S369_S370 | 0.555612837 | 9.69E-06 | 1.47E-04 |
| S23A2_HUMAN\|Q9UGH3_S78_S81 | 0.55543855 | 3.79E-04 | 5.80E-03 |
| PP12C_HUMAN\|Q9BZL4_S453 | 0.555432123 | 2.70E-08 | 1.61E-06 |
| SLTM_HUMAN\|Q9NWH9_S371 | 0.555293475 | 1.07E-02 | 4.26E-02 |
| MISP_HUMAN\|Q8IVT2_S400 | 0.555246258 | 9.88E-05 | 1.29E-03 |
| PTN1_HUMAN\|P18031_S50 | 0.555117393 | 8.94E-04 | 1.45E-02 |
| IASPP_HUMAN\|Q8WUF5_S134 | 0.555079976 | 3.81E-05 | 1.48E-03 |
| DESP_HUMAN\|P15924_S53 | 0.555045098 | 1.56E-03 | 7.92E-03 |
| SPR2B_HUMAN\|P35325_S60 | 0.554997717 | 1.88E-03 | 2.47E-02 |
| AR6P4_HUMAN\|Q66PJ3-7_S44 | 0.554708823 | 2.95E-03 | 1.48E-02 |
| SPR2E_HUMAN\|P22531_S60 | 0.554636088 | 4.57E-04 | 2.94E-03 |
| GYS1_HUMAN\|P13807_T703 | 0.554631748 | 1.17E-05 | 3.44E-04 |
| AFF4_HUMAN\|Q9UHB7_S836 | 0.554460324 | 1.80E-05 | 1.02E-03 |
| NDRG2_HUMAN\|Q9UN36_S332_T330 | 0.554392473 | 2.14E-06 | 9.26E-05 |
| LAD1_HUMAN\|O00515_S512 | 0.554239643 | 2.47E-06 | 7.84E-05 |
| K2C5_HUMAN\|P13647_S21_T19 | 0.554224588 | 4.96E-04 | 5.57E-03 |

| | | | |
|---|---|---|---|
| KLC1_HUMAN|Q07866-4_S631 | 0.554167861 | 1.70E-07 | 6.37E-06 |
| CAMP3_HUMAN|Q9P1Y5_S756 | 0.553934993 | 3.38E-06 | 6.44E-05 |
| ARAF_HUMAN|P10398_S157 | 0.553913033 | 1.11E-06 | 2.77E-05 |
| K1C19_HUMAN|P08727_S22 | 0.553904779 | 2.29E-05 | 5.69E-04 |
| RTKN_HUMAN|Q9BST9_S543 | 0.553897299 | 1.51E-06 | 1.49E-04 |
| MA7D1_HUMAN|Q3KQU3_S313 | 0.553874927 | 1.12E-04 | 3.49E-03 |
| EPN3_HUMAN|Q9H201_S188_S191 | 0.55368608 | 1.95E-05 | 2.46E-04 |
| STIM1_HUMAN|Q13586_S668 | 0.55363893 | 3.37E-04 | 4.12E-03 |
| OSB11_HUMAN|Q9BXB4_S177_S181 | 0.553622864 | 1.93E-03 | 2.03E-02 |
| PAXX_HUMAN|Q9BUH6_S152 | 0.553459816 | 3.21E-04 | 4.01E-03 |
| DOCK6_HUMAN|Q96HP0_S880 | 0.553406539 | 3.25E-05 | 7.07E-04 |
| AHNK_HUMAN|Q09666_S5752_S5763 | 0.553394143 | 8.89E-05 | 3.02E-03 |
| TARA_HUMAN|Q9H2D6_S1955 | 0.55338739 | 4.24E-04 | 2.78E-03 |
| ES8L1_HUMAN|Q8TE68_S182_T187 | 0.553342173 | 1.01E-07 | 1.71E-05 |
| JUPI2_HUMAN|Q9H910_S97 | 0.553278031 | 2.49E-05 | 1.14E-03 |
| RFIP3_HUMAN|O75154_S52 | 0.553209214 | 9.15E-06 | 2.12E-04 |
| NUFP2_HUMAN|Q7Z417_S212_S214 | 0.553124539 | 1.03E-05 | 3.05E-04 |
| RABL6_HUMAN|Q3YEC7_S425_S427 | 0.553095614 | 1.33E-07 | 5.36E-06 |
| OSB11_HUMAN|Q9BXB4_S177_S181 | 0.553058156 | 1.90E-03 | 1.98E-02 |
| COBL1_HUMAN|Q53SF7_S294_T298 | 0.553029118 | 1.33E-05 | 6.98E-04 |
| SARG_HUMAN|Q9BW04_S502 | 0.553018056 | 3.04E-07 | 4.58E-05 |
| KHDR1_HUMAN|Q07666_S18 | 0.552993992 | 9.73E-06 | 6.60E-04 |
| AHNK_HUMAN|Q09666_S5749_S5752 | 0.552930768 | 1.04E-06 | 1.56E-04 |
| PACS1_HUMAN|Q6VY07_T504 | 0.55285154 | 4.37E-05 | 9.00E-04 |
| STK10_HUMAN|O94804_T952 | 0.552824209 | 7.39E-05 | 7.45E-04 |
| ZBT21_HUMAN|Q9ULJ3_S422 | 0.55281854 | 7.30E-04 | 1.03E-02 |
| MILK1_HUMAN|Q8N3F8_S640_S644 | 0.552817209 | 5.93E-06 | 1.01E-04 |
| AHNK_HUMAN|Q09666_S212_S216 | 0.552710774 | 6.20E-07 | 1.70E-05 |
| FAT1_HUMAN|Q14517_S4473 | 0.552690073 | 2.87E-04 | 3.74E-03 |
| SLAI2_HUMAN|Q9P270_S349_S353_S357 | 0.552655402 | 8.37E-07 | 1.38E-04 |
| SMCR8_HUMAN|Q8TEV9_S417 | 0.552643345 | 7.45E-03 | 4.14E-02 |

| | | | |
|---|---|---|---|
| WDR13_HUMAN\|Q9H1Z4_S79 | 0.552534317 | 4.59E-06 | 8.27E-05 |
| FKB15_HUMAN\|Q5T1M5_S344 | 0.552526821 | 5.17E-06 | 9.09E-05 |
| VIME_HUMAN\|P08670_T458 | 0.552517506 | 7.35E-07 | 9.05E-05 |
| HPS5_HUMAN\|Q9UPZ3_S534 | 0.552478036 | 6.56E-06 | 1.08E-04 |
| HIRP3_HUMAN\|Q9BW71_S551_S555 | 0.552325574 | 2.18E-03 | 1.47E-02 |
| UBP24_HUMAN\|Q9UPU5_S1143 | 0.552103872 | 2.54E-06 | 5.31E-05 |
| CT2NL_HUMAN\|Q9P2B4_S443 | 0.551993406 | 9.36E-03 | 3.29E-02 |
| TB182_HUMAN\|Q9C0C2_T1282 | 0.551818561 | 2.53E-06 | 2.28E-04 |
| NDRG1_HUMAN\|Q92597_T366 | 0.551800391 | 1.80E-05 | 2.32E-04 |
| IASPP_HUMAN\|Q8WUF5_S395 | 0.551784499 | 9.93E-05 | 2.84E-03 |
| NGAP_HUMAN\|Q9UJF2_T620 | 0.551723323 | 1.42E-03 | 1.96E-02 |
| TBCD5_HUMAN\|Q92609_S43 | 0.551584708 | 1.50E-05 | 7.77E-04 |
| ITB4_HUMAN\|P16144_S1486_T1487 | 0.551581003 | 1.56E-04 | 1.37E-03 |
| PKP3_HUMAN\|Q9Y446_S238 | 0.551526268 | 9.13E-08 | 4.05E-06 |
| GRB10_HUMAN\|Q13322_S104 | 0.551503176 | 1.30E-07 | 1.35E-05 |
| MAP4_HUMAN\|P27816_S941 | 0.551493761 | 1.02E-05 | 3.15E-04 |
| SP100_HUMAN\|P23497_S407_S410 | 0.551460043 | 1.03E-02 | 4.65E-02 |
| BORG5_HUMAN\|Q00587_S303 | 0.551334562 | 2.03E-06 | 1.88E-04 |
| ITB4_HUMAN\|P16144_S1457_T1455 | 0.551246331 | 1.71E-03 | 8.46E-03 |
| ROCK1_HUMAN\|Q13464_S1333_S1336_S1341 | 0.551240877 | 4.54E-04 | 2.92E-03 |
| K1C14_HUMAN\|P02533_T9 | 0.551104426 | 8.01E-05 | 7.52E-04 |
| SP100_HUMAN\|P23497_S476 | 0.55110369 | 2.21E-05 | 4.20E-04 |
| HMGN1_HUMAN\|P05114_S7 | 0.550986561 | 3.05E-05 | 3.76E-03 |
| PRC2A_HUMAN\|P48634_S1325_S1328_T1323 | 0.550635468 | 2.89E-03 | 1.46E-02 |
| PTN1_HUMAN\|P18031_S50 | 0.550619847 | 7.80E-04 | 1.00E-02 |
| GYS1_HUMAN\|P13807_T703 | 0.550543821 | 9.85E-06 | 6.18E-04 |
| K1C16_HUMAN\|P08779_S448_S451 | 0.550531532 | 1.17E-05 | 1.69E-04 |
| RAI14_HUMAN\|Q9P0K7_T297 | 0.550514829 | 7.41E-05 | 2.49E-03 |
| SPD2A_HUMAN\|Q5TCZ1_S644 | 0.550465759 | 6.96E-05 | 6.73E-04 |
| PRAG1_HUMAN\|Q86YV5_S907 | 0.550441023 | 1.41E-06 | 3.21E-05 |
| BASP1_HUMAN\|P80723_T196 | 0.550265546 | 9.86E-06 | 1.44E-04 |

| | | | |
|---|---|---|---|
| SPF30_HUMAN\|O75940_S201 | 0.55023771 | 5.49E-05 | 1.84E-03 |
| COF1_HUMAN\|P23528_S3 | 0.550106077 | 8.53E-03 | 3.54E-02 |
| M3K20_HUMAN\|Q9NYL2_S727 | 0.550049364 | 7.84E-08 | 2.66E-05 |
| MRE11_HUMAN\|P49959_S649 | 0.550003193 | 1.11E-04 | 1.04E-03 |
| KHDR1_HUMAN\|Q07666_S20 | 0.549841708 | 8.11E-07 | 9.77E-05 |
| PGAM1_HUMAN\|P18669_S14 | 0.549798044 | 1.50E-05 | 2.03E-04 |
| KLF16_HUMAN\|Q9BXK1_S109 | 0.549515931 | 1.13E-03 | 1.03E-02 |
| KC1D_HUMAN\|P48730_S383 | 0.549442204 | 7.85E-06 | 2.47E-04 |
| AHNK_HUMAN\|Q09666_S212_S216 | 0.549423847 | 5.23E-07 | 1.62E-05 |
| NCK5L_HUMAN\|Q9HCH0_S489_S494 | 0.549260837 | 2.31E-06 | 4.72E-05 |
| RS6_HUMAN\|P62753_S236 | 0.549252112 | 2.48E-03 | 3.97E-02 |
| AFAD_HUMAN\|P55196_S1799 | 0.549228547 | 3.50E-05 | 1.42E-03 |
| AKTS1_HUMAN\|Q96B36_S203 | 0.549159217 | 1.06E-03 | 1.33E-02 |
| JMY_HUMAN\|Q8N9B5_S115 | 0.549001565 | 8.27E-05 | 2.87E-03 |
| CCD86_HUMAN\|Q9H6F5_S217 | 0.548907999 | 5.82E-05 | 8.77E-04 |
| 4EBP1_HUMAN\|Q13541_T70 | 0.54883319 | 5.70E-06 | 3.91E-04 |
| SLAI2_HUMAN\|Q9P270_S349_S353_S357 | 0.548783305 | 6.96E-07 | 3.81E-05 |
| SRRM1_HUMAN\|Q8IYB3_T846 | 0.548745628 | 1.09E-04 | 1.81E-03 |
| NDRG1_HUMAN\|Q92597_S364 | 0.54865844 | 1.37E-06 | 1.46E-04 |
| EFNB1_HUMAN\|P98172_S281 | 0.548610052 | 9.53E-06 | 1.45E-04 |
| UBXN7_HUMAN\|O94888_S280_S288 | 0.548603741 | 3.58E-04 | 1.07E-02 |
| RTKN_HUMAN\|Q9BST9_S529 | 0.548533825 | 1.23E-05 | 8.24E-04 |
| MAP4_HUMAN\|P27816_S1073 | 0.54847886 | 2.52E-06 | 1.15E-04 |
| FA83B_HUMAN\|Q5T0W9_S664 | 0.548331935 | 6.98E-07 | 4.56E-05 |
| LAR4B_HUMAN\|Q92615_T732 | 0.548327666 | 2.77E-08 | 1.36E-05 |
| DAPK3_HUMAN\|O43293_S311_S312 | 0.548190998 | 1.80E-06 | 4.09E-05 |
| K2C7_HUMAN\|P08729_S38 | 0.548051798 | 9.82E-04 | 2.07E-02 |
| TERA_HUMAN\|P55072_S794 | 0.547837328 | 1.49E-04 | 1.32E-03 |
| K1C14_HUMAN\|P02533_S11 | 0.5476793 | 6.59E-07 | 1.77E-05 |
| TENS4_HUMAN\|Q8IZW8_S242_S248_S253 | 0.547651522 | 9.27E-04 | 9.35E-03 |
| UFD1_HUMAN\|Q92890_S299 | 0.547411692 | 3.72E-04 | 2.53E-03 |

| | | | |
|---|---|---|---|
| DP13A_HUMAN\|Q9UKG1_S401 | 0.547348574 | 1.99E-05 | 1.09E-03 |
| DBNL_HUMAN\|Q9UJU6_S275 | 0.54716647 | 1.09E-04 | 1.72E-03 |
| SI1L1_HUMAN\|O43166_S1255 | 0.547151685 | 4.60E-05 | 1.95E-03 |
| ERBB2_HUMAN\|P04626_S998 | 0.547137295 | 2.84E-04 | 1.31E-02 |
| ADCY6_HUMAN\|O43306_S54 | 0.546991093 | 3.63E-06 | 6.69E-05 |
| SEPT9_HUMAN\|Q9UHD8_T42 | 0.546930677 | 1.18E-06 | 2.82E-05 |
| AFAD_HUMAN\|P55196_T1330 | 0.546911163 | 6.28E-06 | 1.05E-04 |
| MISP_HUMAN\|Q8IVT2_S471 | 0.546874878 | 1.60E-05 | 9.32E-04 |
| AMOL2_HUMAN\|Q9Y2J4_S759 | 0.546799206 | 4.32E-07 | 1.30E-05 |
| K2C5_HUMAN\|P13647_S21_T19 | 0.546789936 | 3.92E-04 | 8.21E-03 |
| UBP32_HUMAN\|Q8NFA0_S1393 | 0.546484237 | 5.21E-07 | 5.43E-05 |
| SPD2A_HUMAN\|Q5TCZ1_S644 | 0.546461738 | 6.05E-05 | 2.19E-03 |
| K2C75_HUMAN\|O95678_S535 | 0.546246357 | 1.01E-07 | 4.37E-06 |
| ATX2L_HUMAN\|Q8WWM7_S306 | 0.546177168 | 1.31E-03 | 3.47E-02 |
| 2A5E_HUMAN\|Q16537_S32_S34 | 0.546134342 | 2.23E-04 | 5.04E-03 |
| ITB4_HUMAN\|P16144_S1486_T1487 | 0.546005685 | 1.28E-04 | 2.01E-03 |
| REPS1_HUMAN\|Q96D71_S162_S166_T173 | 0.545901177 | 3.17E-07 | 2.11E-05 |
| AHNK_HUMAN\|Q09666_S5745_S5749 | 0.545876974 | 1.48E-07 | 4.22E-05 |
| TMF1_HUMAN\|P82094_S72_S77 | 0.545702273 | 1.62E-04 | 8.88E-03 |
| CR025_HUMAN\|Q96B23_S143_S145 | 0.545612521 | 4.50E-07 | 1.22E-04 |
| T22D4_HUMAN\|Q9Y3Q8_T167 | 0.545545344 | 5.22E-06 | 3.81E-04 |
| BAIP2_HUMAN\|Q9UQB8_S366 | 0.54517104 | 2.75E-04 | 3.61E-03 |
| SGTA_HUMAN\|O43765_S305 | 0.545093675 | 2.57E-06 | 8.09E-05 |
| SRRM1_HUMAN\|Q8IYB3_S738_S743 | 0.545058883 | 1.09E-03 | 1.65E-02 |
| LMNB2_HUMAN\|Q03252_S421_S424 | 0.544948359 | 7.90E-06 | 1.24E-04 |
| BAD_HUMAN\|Q92934_S74 | 0.544943199 | 8.72E-06 | 2.07E-04 |
| NDRG1_HUMAN\|Q92597_S326_S330_S333 | 0.544793099 | 9.58E-06 | 6.60E-04 |
| GPBP1_HUMAN\|Q86WP2_S381 | 0.544617186 | 9.04E-06 | 6.62E-04 |
| IF2B_HUMAN\|P20042_T111 | 0.544595373 | 5.44E-05 | 5.50E-04 |
| AHNK_HUMAN\|Q09666_T5729 | 0.544524031 | 2.83E-07 | 6.29E-05 |
| IASPP_HUMAN\|Q8WUF5_Y109 | 0.544490566 | 1.96E-06 | 4.14E-05 |

| | | | |
|---|---|---|---|
| RAI14_HUMAN\|Q9P0K7_T251 | 0.544473184 | 1.81E-05 | 3.55E-04 |
| PRC2A_HUMAN\|P48634_S1004 | 0.544286964 | 4.95E-06 | 8.60E-05 |
| ZO1_HUMAN\|Q07157_T868_Y843 | 0.544195119 | 4.47E-04 | 4.30E-03 |
| ANK3_HUMAN\|Q12955_S959 | 0.54419358 | 3.49E-04 | 4.24E-03 |
| K2C1_HUMAN\|P04264_S66 | 0.544053742 | 4.16E-04 | 2.74E-03 |
| K1C17_HUMAN\|Q04695_T9 | 0.543967605 | 3.03E-05 | 3.52E-04 |
| TPPP_HUMAN\|O94811_S160 | 0.543865961 | 9.25E-05 | 1.51E-03 |
| TISB_HUMAN\|Q07352_S336 | 0.543747793 | 4.39E-05 | 1.76E-03 |
| CCNL2_HUMAN\|Q96S94_S330 | 0.543570748 | 6.20E-08 | 8.20E-06 |
| IRS2_HUMAN\|Q9Y4H2_S1185_S1203 | 0.543567483 | 1.35E-05 | 8.41E-04 |
| K2C5_HUMAN\|P13647_S50 | 0.543469471 | 4.87E-07 | 1.41E-05 |
| OSB10_HUMAN\|Q9BXB5_S26_S32 | 0.543406982 | 7.24E-08 | 1.78E-05 |
| LPP_HUMAN\|Q93052_S609 | 0.54319547 | 7.50E-05 | 2.46E-03 |
| IRS2_HUMAN\|Q9Y4H2_T350 | 0.543191615 | 4.09E-07 | 6.00E-05 |
| SPTB2_HUMAN\|Q01082_S2341 | 0.543164961 | 8.39E-09 | 3.45E-06 |
| K1C14_HUMAN\|P02533_S39 | 0.54304061 | 1.05E-03 | 1.02E-02 |
| RBP10_HUMAN\|Q6VN20_S365 | 0.542868678 | 1.87E-07 | 6.72E-06 |
| CAMP3_HUMAN\|Q9P1Y5_S756 | 0.542692428 | 2.06E-06 | 9.01E-05 |
| CAMP2_HUMAN\|Q08AD1_S1319 | 0.542683462 | 5.97E-07 | 1.75E-05 |
| ITB4_HUMAN\|P16144_S1457_T1455 | 0.542596813 | 1.36E-03 | 1.24E-02 |
| SARG_HUMAN\|Q9BW04_S429 | 0.542580215 | 6.47E-05 | 2.26E-03 |
| K1C18_HUMAN\|P05783_S7 | 0.542343444 | 1.05E-04 | 9.99E-04 |
| IWS1_HUMAN\|Q96ST2_T725 | 0.54224177 | 6.57E-05 | 6.46E-04 |
| TANK_HUMAN\|Q92844_S129 | 0.54210232 | 3.70E-05 | 7.93E-04 |
| WDR13_HUMAN\|Q9H1Z4_S79 | 0.541974616 | 2.80E-06 | 1.25E-04 |
| GGYF2_HUMAN\|Q6Y7W6_S23 | 0.541896172 | 2.26E-07 | 1.47E-05 |
| AKTS1_HUMAN\|Q96B36_S187 | 0.541887935 | 2.38E-05 | 1.27E-03 |
| DCBD2_HUMAN\|Q96PD2_S726_S727 | 0.541816303 | 2.46E-04 | 1.93E-03 |
| TISB_HUMAN\|Q07352_S334 | 0.541733891 | 4.69E-05 | 4.96E-04 |
| ITB4_HUMAN\|P16144_S1454_S1457 | 0.541717018 | 1.17E-03 | 1.11E-02 |
| DBNL_HUMAN\|Q9UJU6-2_S232 | 0.541647833 | 1.73E-05 | 8.85E-04 |

| | | | |
|---|---|---|---|
| FCHO2_HUMAN\|Q0JRZ9_S387 | 0.541476834 | 3.73E-04 | 1.56E-02 |
| DEFI6_HUMAN\|Q9H4E7_S590_S597_T595 | 0.541121168 | 1.34E-05 | 8.41E-04 |
| YBOX3_HUMAN\|P16989_S369_S370 | 0.540853334 | 5.13E-06 | 9.05E-05 |
| DOCK6_HUMAN\|Q96HP0_S880 | 0.540774378 | 1.93E-05 | 4.94E-04 |
| FA83H_HUMAN\|Q6ZRV2_S513_S516 | 0.540765764 | 8.00E-05 | 7.52E-04 |
| MTCL1_HUMAN\|Q9Y4B5_S196 | 0.540742825 | 1.50E-05 | 9.05E-04 |
| SI1L1_HUMAN\|O43166_S1255 | 0.54045609 | 3.61E-05 | 4.11E-04 |
| GGYF2_HUMAN\|Q6Y7W6_S26 | 0.540427567 | 9.67E-08 | 7.49E-06 |
| NIPA_HUMAN\|Q86WB0_S408_S409 | 0.540380207 | 3.45E-08 | 1.56E-05 |
| LMNA_HUMAN\|P02545_S428 | 0.540310072 | 1.03E-07 | 4.45E-06 |
| ITB4_HUMAN\|P16144_S1486_T1487 | 0.540271722 | 1.04E-04 | 2.90E-03 |
| TACC1_HUMAN\|O75410_S44 | 0.540244972 | 3.02E-06 | 6.00E-05 |
| TBCD4_HUMAN\|O60343_S570 | 0.540207705 | 7.75E-05 | 7.71E-04 |
| SPTB2_HUMAN\|Q01082_T2328 | 0.540156364 | 1.16E-03 | 1.68E-02 |
| PKP4_HUMAN\|Q99569_S233 | 0.540150022 | 8.07E-07 | 2.06E-05 |
| RELB_HUMAN\|Q01201_S573 | 0.540138772 | 1.14E-06 | 1.32E-04 |
| CE170_HUMAN\|Q5SW79_S1165 | 0.540096259 | 4.85E-05 | 5.19E-04 |
| LAP2A_HUMAN\|P42166_S167_T160 | 0.540081928 | 6.01E-06 | 1.02E-04 |
| RELB_HUMAN\|Q01201_S573 | 0.540068529 | 1.14E-06 | 1.19E-04 |
| TB182_HUMAN\|Q9C0C2_S1652 | 0.540055359 | 2.83E-06 | 5.56E-05 |
| K2C6A_HUMAN\|P02538_T548 | 0.540030878 | 3.48E-03 | 1.69E-02 |
| LARP1_HUMAN\|Q6PKG0_S774 | 0.539766139 | 6.29E-08 | 1.48E-05 |
| SARG_HUMAN\|Q9BW04_S131 | 0.539732542 | 3.53E-04 | 6.98E-03 |
| ES8L1_HUMAN\|Q8TE68_S182_T187 | 0.539680681 | 4.58E-08 | 6.89E-06 |
| NIBL1_HUMAN\|Q96TA1_S641_S646 | 0.539603483 | 1.22E-04 | 8.36E-03 |
| REEP4_HUMAN\|Q9H6H4_S253 | 0.53942598 | 8.40E-05 | 8.28E-04 |
| DC1L1_HUMAN\|Q9Y6G9_S516_T515 | 0.539365474 | 1.66E-06 | 5.91E-05 |
| 3BP1_HUMAN\|Q9Y3L3_T601 | 0.539272841 | 3.11E-04 | 6.36E-03 |
| AHNK_HUMAN\|Q09666_S5745_S5749 | 0.539232775 | 1.02E-07 | 8.94E-06 |
| MYPT1_HUMAN\|O14974_S554 | 0.539198649 | 8.58E-04 | 4.86E-03 |
| K2C6A_HUMAN\|P02538_S31 | 0.539155564 | 3.59E-03 | 1.55E-02 |

| | | | |
|---|---|---|---|
| PARD3_HUMAN\|Q8TEW0_S225 | 0.539109689 | 4.47E-06 | 8.14E-05 |
| PSF2_HUMAN\|Q9Y248_S182 | 0.538918607 | 1.87E-05 | 2.39E-04 |
| AHNK_HUMAN\|Q09666_S5731 | 0.538864121 | 2.84E-09 | 1.92E-06 |
| CCNH_HUMAN\|P51946_T315 | 0.53879163 | 2.59E-05 | 3.07E-04 |
| IF4B_HUMAN\|P23588_S597 | 0.538780893 | 4.42E-05 | 2.07E-03 |
| AKTS1_HUMAN\|Q96B36_T246 | 0.538704297 | 4.14E-05 | 1.95E-03 |
| K2C5_HUMAN\|P13647_S26 | 0.538651289 | 2.31E-06 | 4.72E-05 |
| TB182_HUMAN\|Q9C0C2_T875 | 0.538613999 | 7.64E-07 | 9.57E-05 |
| K1C14_HUMAN\|P02533_S32 | 0.538562755 | 1.22E-06 | 3.01E-05 |
| DEFI6_HUMAN\|Q9H4E7_S590_S597_T595 | 0.53854053 | 1.21E-05 | 1.70E-04 |
| COHA1_HUMAN\|Q9UMD9_S85_S93_T97 | 0.538426917 | 1.38E-04 | 1.16E-03 |
| DPYL2_HUMAN\|Q16555_S570 | 0.538350393 | 3.12E-04 | 2.32E-03 |
| ZCH18_HUMAN\|Q86VM9_T796 | 0.538346873 | 5.12E-05 | 5.32E-04 |
| DC1L1_HUMAN\|Q9Y6G9_S421 | 0.538277353 | 4.36E-04 | 8.72E-03 |
| UBXN7_HUMAN\|O94888_S285_S288 | 0.538229938 | 9.86E-05 | 1.70E-03 |
| K2C5_HUMAN\|P13647_S11_S14 | 0.538171398 | 1.75E-07 | 6.42E-06 |
| RAI14_HUMAN\|Q9P0K7_S281 | 0.538090839 | 2.98E-07 | 4.25E-05 |
| SEC62_HUMAN\|Q99442_T375 | 0.538050189 | 6.19E-04 | 1.08E-02 |
| CCDC9_HUMAN\|Q9Y3X0_S376 | 0.537954848 | 1.01E-06 | 1.09E-04 |
| SRRM1_HUMAN\|Q8IYB3_S740_S743 | 0.537933756 | 5.29E-06 | 9.07E-05 |
| PP14A_HUMAN\|Q96A00_S128_S134_S136 | 0.537922427 | 1.86E-07 | 3.27E-05 |
| DEFI6_HUMAN\|Q9H4E7_S590_S597_T595 | 0.537868368 | 1.18E-05 | 6.90E-04 |
| PDS5A_HUMAN\|Q29RF7_S1276 | 0.537867925 | 1.66E-05 | 1.88E-03 |
| AFAD_HUMAN\|P55196_T1232 | 0.537714504 | 2.42E-05 | 2.91E-04 |
| KLC1_HUMAN\|Q07866-4_S631 | 0.537653292 | 6.74E-08 | 3.21E-06 |
| SEPT9_HUMAN\|Q9UHD8_T42 | 0.537642368 | 7.54E-07 | 4.86E-05 |
| DCBD2_HUMAN\|Q96PD2_S724_S727 | 0.537605206 | 1.51E-04 | 1.34E-03 |
| MA7D3_HUMAN\|Q8IWC1_S524 | 0.537443976 | 8.31E-04 | 4.75E-03 |
| YAP1_HUMAN\|P46937_S109_T119 | 0.537440237 | 1.89E-08 | 5.40E-06 |
| PLEC_HUMAN\|Q15149_S4406 | 0.537288409 | 2.71E-03 | 2.88E-02 |
| K1C14_HUMAN\|P02533_S437_S438 | 0.537009018 | 1.55E-05 | 2.07E-04 |

| | | | |
|---|---|---|---|
| K2C5_HUMAN\|P13647_S26_T19 | 0.536921036 | 7.45E-03 | 4.33E-02 |
| AHNK_HUMAN\|Q09666_S5746_S5752 | 0.536856375 | 2.81E-07 | 6.29E-05 |
| K1C16_HUMAN\|P08779_S449_S452 | 0.536666714 | 4.31E-04 | 2.81E-03 |
| 3BP1_HUMAN\|Q9Y3L3_S596 | 0.53650258 | 2.81E-06 | 2.28E-04 |
| SGTA_HUMAN\|O43765_S301_S305 | 0.536415472 | 3.36E-04 | 1.01E-02 |
| YTHD2_HUMAN\|Q9Y5A9_S39 | 0.536329184 | 2.78E-03 | 1.74E-02 |
| CR025_HUMAN\|Q96B23_S145_S147 | 0.536199184 | 1.48E-07 | 5.85E-06 |
| NDRG1_HUMAN\|Q92597_S333_S336_T328 | 0.536143743 | 1.44E-07 | 2.52E-05 |
| AHNK_HUMAN\|Q09666_S5749_S5763 | 0.536128692 | 1.66E-08 | 1.22E-06 |
| TERA_HUMAN\|P55072_S794 | 0.536113658 | 1.01E-04 | 9.21E-04 |
| K1522_HUMAN\|Q9P206_S161 | 0.535824044 | 3.22E-07 | 1.08E-05 |
| CCDC9_HUMAN\|Q9Y3X0_S376_S386_S390 | 0.535546059 | 2.11E-04 | 1.62E-03 |
| EGFR_HUMAN\|P00533_S1039 | 0.535540552 | 8.47E-03 | 4.59E-02 |
| K2C5_HUMAN\|P13647_S11_S8 | 0.535440233 | 4.04E-07 | 1.25E-05 |
| TBCD5_HUMAN\|Q92609_S522 | 0.53535634 | 5.85E-06 | 4.51E-04 |
| DEFI6_HUMAN\|Q9H4E7_S590_S597_T595 | 0.535303243 | 1.06E-05 | 3.25E-04 |
| JUPI2_HUMAN\|Q9H910_S97 | 0.53522064 | 1.19E-05 | 8.08E-04 |
| SPTB2_HUMAN\|Q01082_T2328 | 0.535102385 | 1.02E-03 | 2.86E-02 |
| OSB10_HUMAN\|Q9BXB5_S32 | 0.535021811 | 2.42E-06 | 2.17E-04 |
| K1C17_HUMAN\|Q04695_S12 | 0.534953147 | 1.97E-05 | 2.47E-04 |
| AHNK_HUMAN\|Q09666_S5752_S5763 | 0.534719487 | 4.50E-05 | 4.86E-04 |
| K2C75_HUMAN\|O95678_S535 | 0.534602166 | 5.22E-08 | 2.61E-06 |
| AF1L2_HUMAN\|Q8N4X5_S344 | 0.534229421 | 2.25E-06 | 2.10E-04 |
| MAP7_HUMAN\|Q14244_S209 | 0.534192227 | 3.09E-05 | 3.57E-04 |
| F10A1_HUMAN\|P50502_S76_S79 | 0.534143088 | 5.01E-03 | 2.00E-02 |
| RHG21_HUMAN\|Q5T5U3_S615 | 0.533959798 | 2.91E-07 | 2.35E-05 |
| K1522_HUMAN\|Q9P206_S339 | 0.533836202 | 2.20E-03 | 1.04E-02 |
| RENT1_HUMAN\|Q92900_S1107_T1111 | 0.533790373 | 3.49E-04 | 4.24E-03 |
| ENAH_HUMAN\|Q8N8S7_T489 | 0.533771228 | 2.63E-03 | 2.93E-02 |
| K1522_HUMAN\|Q9P206_S929 | 0.533669843 | 1.00E-07 | 2.12E-05 |
| BICD2_HUMAN\|Q8TD16_S582 | 0.53347791 | 3.46E-03 | 2.39E-02 |

| | | | |
|---|---|---|---|
| MARCS_HUMAN\|P29966_T150 | 0.533309789 | 1.56E-05 | 2.07E-04 |
| K1C14_HUMAN\|P02533_S54_Y46 | 0.53324611 | 2.36E-05 | 5.70E-04 |
| K2C6A_HUMAN\|P02538_S547 | 0.533234113 | 6.79E-03 | 4.05E-02 |
| PKP4_HUMAN\|Q99569_S447 | 0.533132504 | 2.49E-04 | 1.84E-03 |
| APBB2_HUMAN\|Q92870_S409_S412 | 0.533098666 | 5.81E-07 | 1.74E-05 |
| FA83B_HUMAN\|Q5T0W9_S664 | 0.532935581 | 3.38E-07 | 1.11E-05 |
| SCRIB_HUMAN\|Q14160_S1285 | 0.532595001 | 4.76E-05 | 5.02E-04 |
| EMAL3_HUMAN\|Q32P44_S889 | 0.53248854 | 6.30E-04 | 5.68E-03 |
| ASPC1_HUMAN\|Q9BZE9_S275 | 0.532401136 | 9.47E-07 | 1.09E-04 |
| ZO2_HUMAN\|Q9UDY2_T1027 | 0.532274262 | 8.44E-07 | 2.14E-05 |
| MAP7_HUMAN\|Q14244_S202 | 0.532153704 | 1.35E-07 | 5.37E-06 |
| AHNK_HUMAN\|Q09666_S5746_S5752 | 0.532133 | 2.19E-07 | 3.49E-05 |
| RHG29_HUMAN\|Q52LW3_S176 | 0.532040062 | 1.10E-04 | 1.42E-03 |
| GOGA4_HUMAN\|Q13439_S89_S92 | 0.532023052 | 2.21E-06 | 4.76E-05 |
| NDRG1_HUMAN\|Q92597_S330_T328 | 0.532018291 | 2.49E-06 | 5.02E-05 |
| EXOC7_HUMAN\|Q9UPT5_S250 | 0.532016147 | 1.29E-06 | 6.20E-05 |
| PRC2A_HUMAN\|P48634_S1147 | 0.531983469 | 1.22E-07 | 1.00E-05 |
| ROCK1_HUMAN\|Q13464_S1341 | 0.531883472 | 1.30E-05 | 1.82E-04 |
| LARP1_HUMAN\|Q6PKG0_S631_Y633 | 0.531872066 | 9.18E-04 | 5.13E-03 |
| NGAP_HUMAN\|Q9UJF2_T620 | 0.531858897 | 8.35E-04 | 1.39E-02 |
| GRIN1_HUMAN\|Q7Z2K8_S615 | 0.531803388 | 9.18E-07 | 1.44E-04 |
| LMNB1_HUMAN\|P20700_S23_T20 | 0.531738948 | 2.39E-03 | 1.57E-02 |
| KC1D_HUMAN\|P48730_S331 | 0.531661139 | 2.77E-05 | 6.25E-04 |
| PKN2_HUMAN\|Q16513_S362 | 0.53156271 | 3.67E-06 | 2.75E-04 |
| E41L2_HUMAN\|O43491_S87 | 0.531549108 | 1.70E-06 | 3.71E-05 |
| TAOK3_HUMAN\|Q9H2K8_S324 | 0.531522823 | 3.74E-07 | 1.13E-04 |
| STAT1_HUMAN\|P42224_S727 | 0.53145673 | 8.28E-07 | 3.65E-05 |
| TB182_HUMAN\|Q9C0C2_T875 | 0.531224322 | 5.26E-07 | 1.62E-05 |
| NELFE_HUMAN\|P18615_S353 | 0.531216353 | 6.44E-09 | 5.29E-07 |
| MILK1_HUMAN\|Q8N3F8_S640_S644 | 0.53095478 | 2.35E-06 | 2.06E-04 |
| BAIP2_HUMAN\|Q9UQB8_Y491 | 0.530920074 | 1.79E-03 | 8.79E-03 |

| | | | |
|---|---|---|---|
| SPTB2_HUMAN\|Q01082-3_S14 | 0.530909909 | 9.91E-06 | 6.18E-04 |
| RBP10_HUMAN\|Q6VN20_S365 | 0.530904943 | 1.04E-07 | 8.98E-06 |
| PDLI5_HUMAN\|Q96HC4_S111 | 0.530897674 | 1.02E-06 | 2.50E-05 |
| K1C14_HUMAN\|P02533_S28 | 0.53081412 | 1.24E-06 | 2.93E-05 |
| LRRF2_HUMAN\|Q9Y608_S320_S323 | 0.530509821 | 2.43E-05 | 2.96E-04 |
| TOP2B_HUMAN\|Q02880_S1550_S1552 | 0.530491405 | 1.29E-04 | 8.64E-03 |
| K2C6A_HUMAN\|P02538_S546 | 0.530437912 | 3.94E-03 | 1.87E-02 |
| 1433Z_HUMAN\|P63104_T232 | 0.530302104 | 6.65E-05 | 6.51E-04 |
| LMNA_HUMAN\|P02545_S618 | 0.530013992 | 2.72E-03 | 2.03E-02 |
| ACINU_HUMAN\|Q9UKV3_T254 | 0.529831249 | 7.71E-05 | 3.14E-03 |
| CRTC3_HUMAN\|Q6UUV7_S370 | 0.529821527 | 3.97E-05 | 8.31E-04 |
| NDRG1_HUMAN\|Q92597_T366 | 0.529624902 | 6.94E-06 | 2.26E-04 |
| BORG4_HUMAN\|Q9H3Q1_S292 | 0.529591763 | 2.09E-05 | 2.65E-04 |
| PACS1_HUMAN\|Q6VY07_T504 | 0.52944742 | 1.87E-05 | 9.29E-04 |
| ZCCHV_HUMAN\|Q7Z2W4_T380 | 0.52933994 | 4.17E-06 | 3.43E-04 |
| INADL_HUMAN\|Q8NI35_S645 | 0.529242119 | 2.58E-07 | 8.59E-06 |
| SSFA2_HUMAN\|P28290_S668 | 0.529151392 | 3.03E-05 | 1.58E-03 |
| K1C14_HUMAN\|P02533_S435_S438 | 0.528965804 | 2.69E-05 | 3.17E-04 |
| K2C5_HUMAN\|P13647_S21_T19 | 0.528884585 | 2.20E-04 | 7.22E-03 |
| K2C6A_HUMAN\|P02538_S547 | 0.528755975 | 6.16E-03 | 2.32E-02 |
| AF1L2_HUMAN\|Q8N4X5_S344 | 0.528730815 | 1.76E-06 | 3.76E-05 |
| K2C6A_HUMAN\|P02538_S58 | 0.528535208 | 2.36E-03 | 1.23E-02 |
| OXR1_HUMAN\|Q8N573_S201_S202 | 0.52852844 | 2.45E-07 | 3.80E-05 |
| COBL1_HUMAN\|Q53SF7_T322 | 0.52848336 | 2.48E-07 | 6.12E-05 |
| LAD1_HUMAN\|O00515_S38_S39 | 0.52845482 | 1.69E-04 | 4.13E-03 |
| K2C5_HUMAN\|P13647_S62 | 0.528359193 | 1.92E-06 | 4.32E-05 |
| ERBB2_HUMAN\|P04626_S998 | 0.528353769 | 1.48E-04 | 3.85E-03 |
| ZCCHV_HUMAN\|Q7Z2W4_S378 | 0.528313377 | 1.17E-08 | 9.60E-07 |
| K2C5_HUMAN\|P13647_S11_S14 | 0.527934492 | 9.78E-08 | 4.28E-06 |
| LAP2A_HUMAN\|P42166_S156_T160 | 0.527930335 | 9.54E-05 | 1.53E-03 |
| COBL1_HUMAN\|Q53SF7_T322 | 0.527911595 | 2.42E-07 | 2.08E-05 |

| | | | |
|---|---|---|---|
| SCRIB_HUMAN\|Q14160_S1348 | 0.527764161 | 6.87E-05 | 6.66E-04 |
| NDRG1_HUMAN\|Q92597_S326_S330_S333 | 0.527752983 | 4.50E-06 | 4.11E-04 |
| FA83H_HUMAN\|Q6ZRV2_S974 | 0.527647687 | 2.77E-07 | 1.71E-05 |
| IF6_HUMAN\|P56537_S243 | 0.527626541 | 4.31E-04 | 3.04E-03 |
| MAN1_HUMAN\|Q9Y2U8_S144 | 0.527511514 | 7.67E-05 | 2.48E-03 |
| AHNK_HUMAN\|Q09666_S4092 | 0.527511136 | 1.72E-04 | 1.36E-03 |
| SPD2A_HUMAN\|Q5TCZ1_S651 | 0.52743202 | 2.31E-05 | 5.69E-04 |
| TB182_HUMAN\|Q9C0C2_S691 | 0.527206904 | 1.36E-06 | 5.26E-04 |
| GSK3A_HUMAN\|P49840_T19 | 0.527118631 | 1.61E-07 | 6.13E-06 |
| PKP1_HUMAN\|Q13835_S205 | 0.527112411 | 3.39E-03 | 1.47E-02 |
| PKN2_HUMAN\|Q16513_S360 | 0.526962329 | 1.31E-05 | 1.82E-04 |
| BORG5_HUMAN\|Q00587_S303 | 0.526919905 | 6.84E-07 | 1.24E-04 |
| AHNK_HUMAN\|Q09666_S5752_S5763 | 0.526822381 | 3.36E-05 | 7.28E-04 |
| CREB1_HUMAN\|P16220_S271 | 0.526819955 | 2.70E-05 | 6.26E-04 |
| ANK3_HUMAN\|Q12955_S959 | 0.526780909 | 2.05E-04 | 2.83E-03 |
| BASP1_HUMAN\|P80723_T196 | 0.526752764 | 3.25E-06 | 1.41E-04 |
| ZCCHV_HUMAN\|Q7Z2W4_T380 | 0.526388684 | 3.70E-06 | 6.94E-05 |
| MTCL1_HUMAN\|Q9Y4B5_S196 | 0.526329067 | 8.57E-06 | 1.33E-04 |
| ZCH18_HUMAN\|Q86VM9_S532_S534 | 0.526219026 | 2.57E-09 | 2.79E-07 |
| NDE1_HUMAN\|Q9NXR1_S282 | 0.526199824 | 2.30E-04 | 3.15E-03 |
| STXB4_HUMAN\|Q6ZWJ1_S463 | 0.526156026 | 7.43E-04 | 4.33E-03 |
| GSK3A_HUMAN\|P49840_T19 | 0.526002566 | 1.52E-07 | 5.89E-06 |
| ABI2_HUMAN\|Q9NYB9_S368 | 0.525768805 | 1.37E-08 | 9.32E-07 |
| PRAG1_HUMAN\|Q86YV5_S907 | 0.525722313 | 4.36E-07 | 1.40E-05 |
| ANXA2_HUMAN\|P07355_S127 | 0.525643958 | 5.19E-04 | 3.26E-03 |
| K2C6A_HUMAN\|P02538_Y62 | 0.525407847 | 2.73E-03 | 1.39E-02 |
| CLAP1_HUMAN\|Q7Z460_S1091 | 0.525225567 | 1.79E-07 | 6.60E-06 |
| ZCH18_HUMAN\|Q86VM9_T796 | 0.525015768 | 3.14E-05 | 3.67E-04 |
| SEPT7_HUMAN\|Q16181_T426 | 0.524897946 | 1.80E-04 | 1.42E-03 |
| MET_HUMAN\|P08581_S997_T993 | 0.524880347 | 2.94E-05 | 5.14E-04 |
| LRRF2_HUMAN\|Q9Y608_S320_S323 | 0.52467414 | 1.91E-05 | 4.93E-04 |

| | | | |
|---|---|---|---|
| NHRF1_HUMAN\|O14745_S280 | 0.524638075 | 2.85E-07 | 1.72E-05 |
| K1671_HUMAN\|Q9BY89_S127 | 0.524573874 | 3.92E-07 | 1.22E-05 |
| NBEL2_HUMAN\|Q6ZNJ1_T1867 | 0.52453353 | 3.26E-05 | 5.65E-04 |
| PEA15_HUMAN\|Q15121_S116 | 0.524495374 | 3.81E-05 | 6.27E-04 |
| K2C5_HUMAN\|P13647_S571_S574 | 0.524411371 | 3.49E-06 | 1.37E-04 |
| GSK3A_HUMAN\|P49840_S21 | 0.52431976 | 4.99E-07 | 1.56E-05 |
| PDLI5_HUMAN\|Q96HC4_S111 | 0.524317432 | 7.22E-07 | 9.05E-05 |
| T22D4_HUMAN\|Q9Y3Q8_T167 | 0.524225607 | 1.98E-06 | 2.15E-04 |
| MICA3_HUMAN\|Q7RTP6_S685_S687 | 0.524223272 | 1.08E-04 | 1.39E-03 |
| SGTA_HUMAN\|O43765_S301_S305 | 0.52416934 | 2.30E-04 | 1.84E-03 |
| IRS2_HUMAN\|Q9Y4H2_S606 | 0.524071929 | 2.21E-04 | 1.80E-03 |
| WAC2A_HUMAN\|Q641Q2_S375 | 0.5240602 | 4.01E-05 | 4.36E-04 |
| CAMP2_HUMAN\|Q08AD1_S1319 | 0.524044338 | 2.37E-07 | 2.08E-05 |
| MFF_HUMAN\|Q9GZY8_S157 | 0.523899843 | 6.19E-05 | 6.42E-04 |
| K2C5_HUMAN\|P13647_S574 | 0.523830083 | 4.92E-07 | 1.41E-05 |
| SMTN_HUMAN\|P53814_S514 | 0.523785555 | 1.16E-03 | 3.17E-02 |
| AHNK_HUMAN\|Q09666_S5749_S5752 | 0.523534981 | 2.32E-07 | 8.17E-06 |
| CE170_HUMAN\|Q5SW79_S636_T628 | 0.523396425 | 9.89E-06 | 1.49E-04 |
| DOCK4_HUMAN\|Q8N1I0_S1620 | 0.523349828 | 2.65E-04 | 3.50E-03 |
| PSF2_HUMAN\|Q9Y248_S182_T180 | 0.523252412 | 2.90E-05 | 3.43E-04 |
| BAD_HUMAN\|Q92934_Y76 | 0.523094812 | 1.00E-07 | 7.54E-06 |
| TB182_HUMAN\|Q9C0C2_S1029 | 0.523018768 | 1.81E-07 | 1.24E-05 |
| TB182_HUMAN\|Q9C0C2_S1158 | 0.52300529 | 4.77E-06 | 8.56E-05 |
| PA24A_HUMAN\|P47712_S729 | 0.522990333 | 3.17E-04 | 5.51E-03 |
| K1522_HUMAN\|Q9P206_S612 | 0.522885096 | 1.75E-05 | 1.94E-03 |
| NDRG1_HUMAN\|Q92597_S330_S336_T328 | 0.522831947 | 1.95E-03 | 2.34E-02 |
| K2C5_HUMAN\|P13647_S50 | 0.522822388 | 1.56E-07 | 6.08E-06 |
| TBCD4_HUMAN\|O60343_S570 | 0.522541976 | 4.23E-05 | 1.55E-03 |
| AHNK2_HUMAN\|Q8IVF2_S294_T298 | 0.522492022 | 4.55E-06 | 1.65E-04 |
| PKP2_HUMAN\|Q99959_S132 | 0.522157004 | 1.75E-05 | 4.57E-04 |
| ANS1A_HUMAN\|Q92625_S649 | 0.522154728 | 1.11E-08 | 2.07E-06 |

| | | | |
|---|---|---|---|
| NBN_HUMAN\|O60934_S343 | 0.522073406 | 1.45E-06 | 3.48E-05 |
| SON_HUMAN\|P18583_S94 | 0.522000689 | 3.32E-08 | 3.60E-05 |
| 2A5E_HUMAN\|Q16537_S32_S34 | 0.521952443 | 9.88E-05 | 9.05E-04 |
| GRB10_HUMAN\|Q13322_S104 | 0.521904561 | 2.52E-08 | 7.19E-06 |
| ZBT21_HUMAN\|Q9ULJ3_S422 | 0.521863054 | 2.94E-04 | 4.73E-03 |
| EIF3A_HUMAN\|Q14152_S882 | 0.5217276 | 1.14E-04 | 1.01E-03 |
| CE170_HUMAN\|Q5SW79_S636_T628 | 0.521563252 | 9.18E-06 | 1.35E-04 |
| PDLI2_HUMAN\|Q96JY6_S121_S124_S127 | 0.521462046 | 8.37E-03 | 4.55E-02 |
| ML12A_HUMAN\|P19105_T18 | 0.521322011 | 3.41E-06 | 6.44E-05 |
| ATAT_HUMAN\|Q5SQI0_S315 | 0.521225393 | 1.06E-08 | 3.38E-06 |
| RTKN_HUMAN\|Q9BST9_S543 | 0.521200948 | 3.33E-07 | 2.17E-05 |
| IASPP_HUMAN\|Q8WUF5_S395 | 0.521198211 | 3.38E-05 | 7.30E-04 |
| AR6P4_HUMAN\|Q66PJ3-7_S44 | 0.521196262 | 1.27E-03 | 1.12E-02 |
| HMGA1_HUMAN\|P17096_S49 | 0.521111615 | 6.50E-04 | 4.30E-03 |
| AKAP2_HUMAN\|Q9Y2D5_S748 | 0.521106337 | 6.01E-05 | 8.86E-04 |
| SRPK2_HUMAN\|P78362_S496_T498 | 0.521073066 | 2.17E-06 | 7.16E-05 |
| AFAD_HUMAN\|P55196_S1172_S1173_S1182 | 0.521029736 | 1.01E-07 | 3.08E-05 |
| SI1L1_HUMAN\|O43166_S1255 | 0.520911146 | 1.76E-05 | 4.70E-04 |
| F120A_HUMAN\|Q9NZB2_S1023 | 0.520863543 | 5.34E-07 | 1.51E-05 |
| MED24_HUMAN\|O75448_S862_S873 | 0.520834952 | 2.11E-04 | 2.40E-03 |
| LMNA_HUMAN\|P02545_S404_S406_S414 | 0.520719012 | 1.63E-04 | 1.43E-03 |
| VIME_HUMAN\|P08670_S51 | 0.520626655 | 3.08E-06 | 5.92E-05 |
| LMNB2_HUMAN\|Q03252_S421_S424 | 0.520565257 | 2.47E-06 | 5.01E-05 |
| GRB10_HUMAN\|Q13322_S104 | 0.52055152 | 2.34E-08 | 3.16E-06 |
| TNIK_HUMAN\|Q9UKE5_S707 | 0.520507711 | 2.47E-04 | 1.94E-03 |
| GRSF1_HUMAN\|Q12849_S118 | 0.520506826 | 7.12E-07 | 3.28E-05 |
| JMY_HUMAN\|Q8N9B5_S115 | 0.520446359 | 2.72E-05 | 1.20E-03 |
| G3BP1_HUMAN\|Q13283_S250 | 0.52041781 | 1.69E-03 | 8.39E-03 |
| CDK13_HUMAN\|Q14004_T1056 | 0.520399185 | 3.08E-06 | 5.92E-05 |
| CRBG1_HUMAN\|Q9Y4K1_S22 | 0.520368183 | 5.98E-08 | 6.01E-06 |
| CRBG1_HUMAN\|Q9Y4K1_S141 | 0.520099037 | 6.28E-06 | 2.25E-04 |

| | | | |
|---|---|---|---|
| CR025_HUMAN\|Q96B23_S66_S69 | 0.520019011 | 2.28E-04 | 1.73E-03 |
| WRIP1_HUMAN\|Q96S55_T116 | 0.519932636 | 5.56E-04 | 3.44E-03 |
| SYTL1_HUMAN\|Q8IYJ3_S216 | 0.519833187 | 1.77E-04 | 2.53E-03 |
| PHLB2_HUMAN\|Q86SQ0_Y567 | 0.519777593 | 1.93E-07 | 6.87E-06 |
| PKP3_HUMAN\|Q9Y446_S313_S314 | 0.519773989 | 2.24E-07 | 7.74E-06 |
| AFAD_HUMAN\|P55196_T1330 | 0.519615171 | 2.05E-06 | 9.96E-05 |
| UBP20_HUMAN\|Q9Y2K6_S132_S134 | 0.519602684 | 4.23E-04 | 6.23E-03 |
| UTP18_HUMAN\|Q9Y5J1_S121 | 0.519342536 | 3.83E-03 | 2.20E-02 |
| BAIP2_HUMAN\|Q9UQB8_Y491 | 0.519322087 | 1.33E-03 | 7.66E-03 |
| PHLB1_HUMAN\|Q86UU1_S489 | 0.519224076 | 4.90E-05 | 5.12E-04 |
| RENT1_HUMAN\|Q92900_S1127 | 0.51921263 | 2.60E-03 | 1.97E-02 |
| K2C6B_HUMAN\|P04259_S541 | 0.519020625 | 1.25E-06 | 2.94E-05 |
| AHNK_HUMAN\|Q09666_S216_S220 | 0.519019554 | 8.06E-08 | 7.67E-06 |
| JMY_HUMAN\|Q8N9B5_S115 | 0.518998468 | 2.56E-05 | 5.99E-04 |
| IWS1_HUMAN\|Q96ST2_T725 | 0.518888714 | 2.86E-05 | 6.51E-04 |
| K2C5_HUMAN\|P13647_S11_S8 | 0.518792146 | 1.62E-07 | 6.13E-06 |
| COBL1_HUMAN\|Q53SF7_T322 | 0.518722024 | 1.54E-07 | 2.61E-05 |
| TANK_HUMAN\|Q92844_S129 | 0.518647176 | 1.47E-05 | 1.96E-04 |
| FA83B_HUMAN\|Q5T0W9_T687 | 0.518405501 | 4.84E-03 | 4.59E-02 |
| NDRG1_HUMAN\|Q92597_S330_S333 | 0.518178422 | 3.87E-07 | 1.26E-05 |
| TB182_HUMAN\|Q9C0C2_S1711 | 0.518174296 | 2.91E-07 | 9.86E-06 |
| SPIR1_HUMAN\|Q08AE8_T509 | 0.518107961 | 9.48E-07 | 2.48E-05 |
| R3HD2_HUMAN\|Q9Y2K5_T856 | 0.518102585 | 9.15E-05 | 1.21E-03 |
| K1C14_HUMAN\|P02533_S13 | 0.517795713 | 8.39E-08 | 3.76E-06 |
| LMNA_HUMAN\|P02545_S404_S407_T409 | 0.517792658 | 3.80E-05 | 4.28E-04 |
| TR10A_HUMAN\|O00220_S466 | 0.517632033 | 1.11E-04 | 3.13E-03 |
| NDRG1_HUMAN\|Q92597_S326_S330_S333 | 0.517510243 | 2.83E-06 | 1.19E-04 |
| MAP4_HUMAN\|P27816_S941 | 0.517507365 | 2.20E-06 | 4.54E-05 |
| IASPP_HUMAN\|Q8WUF5_S74 | 0.517256033 | 1.28E-08 | 8.87E-07 |
| ASPP2_HUMAN\|Q13625_S556 | 0.517139698 | 2.09E-06 | 7.04E-05 |
| TCOF_HUMAN\|Q13428_T249 | 0.517127445 | 2.68E-06 | 5.52E-05 |

| | | | |
|---|---|---|---|
| GRAP1_HUMAN\|Q4V328_S664 | 0.517055227 | 2.65E-06 | 5.29E-05 |
| ARHG5_HUMAN\|Q12774_S450 | 0.516982068 | 2.52E-05 | 4.64E-04 |
| AHNK_HUMAN\|Q09666_S5749_S5752 | 0.516836272 | 1.64E-07 | 1.25E-05 |
| TB182_HUMAN\|Q9C0C2_S1158 | 0.51669939 | 3.62E-06 | 6.69E-05 |
| ABI2_HUMAN\|Q9NYB9_Y371 | 0.516658219 | 1.20E-07 | 4.95E-06 |
| CAVN3_HUMAN\|Q969G5_T205 | 0.51663763 | 1.18E-05 | 6.50E-04 |
| PKP3_HUMAN\|Q9Y446_S238 | 0.516436984 | 1.34E-08 | 3.14E-06 |
| RGS3_HUMAN\|P49796_S1044 | 0.516427913 | 1.73E-07 | 6.42E-06 |
| TAOK3_HUMAN\|Q9H2K8_S442 | 0.51633886 | 8.60E-06 | 1.32E-04 |
| L7UQD0_HUMAN\|L7UQD0_S268 | 0.516337722 | 3.48E-06 | 6.53E-05 |
| CRBG1_HUMAN\|Q9Y4K1_S410 | 0.516320793 | 6.39E-05 | 9.22E-04 |
| RS6_HUMAN\|P62753_S236 | 0.516209335 | 1.09E-03 | 8.53E-03 |
| F122A_HUMAN\|Q96E09_S37 | 0.516032232 | 7.98E-04 | 4.61E-03 |
| SARG_HUMAN\|Q9BW04_S131 | 0.516000827 | 1.69E-04 | 2.45E-03 |
| BICD2_HUMAN\|Q8TD16_S582 | 0.515830232 | 2.31E-03 | 4.61E-02 |
| DDX3X_HUMAN\|O00571_S612 | 0.515793277 | 8.03E-04 | 4.62E-03 |
| AHNK_HUMAN\|Q09666_S5099 | 0.515661553 | 6.83E-05 | 2.87E-03 |
| YAP1_HUMAN\|P46937_T110_T119 | 0.515558139 | 1.63E-05 | 2.15E-04 |
| NDRG1_HUMAN\|Q92597_S333_S336_T328 | 0.515556508 | 4.47E-08 | 1.73E-05 |
| K1C14_HUMAN\|P02533_S432 | 0.515530543 | 2.03E-05 | 2.53E-04 |
| ML12A_HUMAN\|P19105_S19 | 0.515479482 | 2.81E-07 | 9.01E-06 |
| OSB11_HUMAN\|Q9BXB4_S181 | 0.515398419 | 1.33E-07 | 1.08E-05 |
| PRC2A_HUMAN\|P48634_S1085_S1089_S1092 | 0.515363882 | 2.08E-07 | 7.29E-06 |
| AHNK_HUMAN\|Q09666_S5784 | 0.51528641 | 2.49E-03 | 2.94E-02 |
| STIM2_HUMAN\|Q9P246_S522 | 0.51525024 | 1.90E-04 | 2.67E-03 |
| APBB2_HUMAN\|Q92870_S409_S412 | 0.5151877 | 2.38E-07 | 8.16E-06 |
| K1C17_HUMAN\|Q04695_S24 | 0.515111579 | 9.53E-07 | 2.48E-05 |
| SF3A3_HUMAN\|Q12874_S365_S367_S369 | 0.515078993 | 1.41E-03 | 8.01E-03 |
| COHA1_HUMAN\|Q9UMD9_S93_S96 | 0.515009359 | 6.13E-05 | 1.13E-03 |
| K2C5_HUMAN\|P13647_S574 | 0.514912039 | 3.06E-07 | 2.07E-05 |
| K2C6A_HUMAN\|P02538_S85 | 0.514845919 | 2.63E-03 | 1.98E-02 |

| | | | |
|---|---|---|---|
| ERCC5_HUMAN\|P28715_S382_S384 | 0.514838135 | 3.63E-05 | 7.74E-04 |
| M3K20_HUMAN\|Q9NYL2_S727 | 0.514701817 | 1.25E-08 | 3.08E-06 |
| M3K20_HUMAN\|Q9NYL2_S727 | 0.514679421 | 1.25E-08 | 8.79E-07 |
| MISP_HUMAN\|Q8IVT2_S400 | 0.514549434 | 2.35E-05 | 5.64E-04 |
| FA83H_HUMAN\|Q6ZRV2_S1098 | 0.51440939 | 2.62E-07 | 8.99E-06 |
| OSB10_HUMAN\|Q9BXB5_S32 | 0.514265112 | 9.30E-07 | 4.80E-05 |
| CF132_HUMAN\|Q5T0Z8_T297 | 0.514262845 | 4.86E-05 | 9.65E-04 |
| GYS1_HUMAN\|P13807_S727 | 0.514030842 | 1.32E-07 | 9.57E-06 |
| AHNK_HUMAN\|Q09666_S5749_S5763 | 0.5139996 | 4.31E-09 | 2.60E-06 |
| IRS2_HUMAN\|Q9Y4H2_S679 | 0.513941785 | 7.32E-07 | 9.57E-05 |
| MISP_HUMAN\|Q8IVT2_S395 | 0.513902541 | 3.59E-05 | 6.04E-04 |
| ABI2_HUMAN\|Q9NYB9_S368 | 0.513756567 | 6.91E-09 | 6.14E-07 |
| K2C8_HUMAN\|P05787_S477 | 0.513667634 | 1.42E-05 | 1.91E-04 |
| LAP2A_HUMAN\|P42166_T160 | 0.51364871 | 1.81E-07 | 6.64E-06 |
| BORG5_HUMAN\|Q00587_S303 | 0.513595069 | 3.75E-07 | 5.34E-05 |
| JUNB_HUMAN\|P17275_T255 | 0.513570103 | 5.94E-09 | 4.95E-07 |
| PKP1_HUMAN\|Q13835_S205 | 0.513271964 | 2.42E-03 | 1.87E-02 |
| DLG1_HUMAN\|Q12959_S573 | 0.513121528 | 1.10E-04 | 3.43E-03 |
| NDRG1_HUMAN\|Q92597_S364 | 0.513046152 | 2.17E-07 | 1.59E-05 |
| KMT2A_HUMAN\|Q03164_T3028 | 0.512991828 | 4.73E-04 | 9.21E-03 |
| INADL_HUMAN\|Q8NI35_S645 | 0.512912817 | 1.16E-07 | 4.83E-06 |
| EFNB1_HUMAN\|P98172_S281 | 0.512830784 | 1.79E-06 | 4.09E-05 |
| JMY_HUMAN\|Q8N9B5_S115 | 0.512777843 | 1.99E-05 | 2.54E-04 |
| TANK_HUMAN\|Q92844_S129 | 0.512603047 | 1.15E-05 | 3.41E-04 |
| REST_HUMAN\|Q13127_S864 | 0.512576792 | 6.17E-05 | 1.17E-03 |
| GSK3A_HUMAN\|P49840_S21 | 0.512506512 | 2.86E-07 | 9.12E-06 |
| COHA1_HUMAN\|Q9UMD9_S61 | 0.512473233 | 3.26E-05 | 3.79E-04 |
| RBM7_HUMAN\|Q9Y580_S136 | 0.512260982 | 7.12E-06 | 1.76E-04 |
| CAMP2_HUMAN\|Q08AD1_S691 | 0.5122273 | 2.20E-05 | 2.70E-04 |
| K1C19_HUMAN\|P08727_S13 | 0.512141758 | 3.79E-05 | 8.06E-04 |
| RGS3_HUMAN\|P49796_S1044 | 0.512107083 | 1.37E-07 | 5.41E-06 |

| | | | |
|---|---|---|---|
| CLAP1_HUMAN\|Q7Z460_S1091 | 0.511870941 | 8.83E-08 | 1.02E-05 |
| CR025_HUMAN\|Q96B23_S67 | 0.511853166 | 2.17E-05 | 2.35E-03 |
| NDE1_HUMAN\|Q9NXR1_S282 | 0.511830818 | 1.46E-04 | 3.73E-03 |
| E7ETA6_HUMAN\|E7ETA6_T1557 | 0.511616834 | 1.36E-04 | 3.88E-03 |
| K2C6A_HUMAN\|P02538_S34 | 0.511487122 | 1.07E-02 | 3.65E-02 |
| FGD6_HUMAN\|Q6ZV73_S721 | 0.511408281 | 9.40E-06 | 6.60E-04 |
| PHLB2_HUMAN\|Q86SQ0_S565 | 0.511376753 | 1.26E-07 | 5.08E-06 |
| TANK_HUMAN\|Q92844_S129 | 0.511325241 | 1.09E-05 | 1.62E-04 |
| PHRF1_HUMAN\|Q9P1Y6_S915_T917 | 0.51109036 | 3.84E-04 | 4.60E-03 |
| SPTB2_HUMAN\|Q01082_S2341 | 0.5110826 | 1.44E-09 | 5.56E-07 |
| RTKN_HUMAN\|Q9BST9_S529 | 0.511021602 | 2.71E-06 | 2.26E-04 |
| PAXI_HUMAN\|P49023_S250 | 0.51095922 | 1.27E-04 | 2.00E-03 |
| LAP2A_HUMAN\|P42166_T74 | 0.510792928 | 2.11E-05 | 2.66E-04 |
| MAP4_HUMAN\|P27816_S941 | 0.510716511 | 1.60E-06 | 1.85E-04 |
| NDRG1_HUMAN\|Q92597_S330_S333_S336 | 0.510685661 | 1.81E-04 | 1.55E-03 |
| FOSL2_HUMAN\|P15408_S19 | 0.51057363 | 1.50E-04 | 1.83E-03 |
| MFF_HUMAN\|Q9GZY8_S157 | 0.510571572 | 3.69E-05 | 4.08E-04 |
| SMG9_HUMAN\|Q9H0W8_T56 | 0.510558816 | 2.83E-07 | 9.63E-06 |
| SYTL1_HUMAN\|Q8IYJ3_S216 | 0.510530065 | 1.31E-04 | 1.12E-03 |
| PPR18_HUMAN\|Q6NYC8_S209_S213 | 0.510524295 | 1.24E-07 | 5.06E-06 |
| SEPT7_HUMAN\|Q16181_T426 | 0.510410353 | 1.14E-04 | 1.87E-03 |
| AHNK_HUMAN\|Q09666_S5745_S5749 | 0.510348329 | 2.00E-08 | 5.43E-06 |
| ZDHC5_HUMAN\|Q9C0B5_S704 | 0.510266226 | 8.66E-05 | 8.10E-04 |
| K2C5_HUMAN\|P13647_S64 | 0.510016525 | 2.48E-05 | 5.87E-04 |
| ATX2L_HUMAN\|Q8WWM7_S306 | 0.509990983 | 4.92E-04 | 9.32E-03 |
| NCK2_HUMAN\|O43639_S90_T95 | 0.50989891 | 5.46E-03 | 2.12E-02 |
| K2C8_HUMAN\|P05787_S477 | 0.509832083 | 1.20E-05 | 7.76E-04 |
| PLEC_HUMAN\|Q15149_S4406 | 0.509824113 | 1.39E-03 | 3.44E-02 |
| ITA2_HUMAN\|P17301_S1181 | 0.509736371 | 1.46E-05 | 5.58E-03 |
| MARCS_HUMAN\|P29966_T150 | 0.509705214 | 5.60E-06 | 9.70E-05 |
| REPS1_HUMAN\|Q96D71_S162_S166_T173 | 0.509465706 | 4.83E-08 | 2.60E-06 |

| | | | |
|---|---|---|---|
| MYPT1_HUMAN\|O14974_S554 | 0.509343779 | 3.75E-04 | 7.94E-03 |
| S6A15_HUMAN\|Q9H2J7_S699 | 0.509333516 | 7.84E-05 | 7.40E-04 |
| YAP1_HUMAN\|P46937_T110_T119 | 0.509304241 | 1.28E-05 | 7.07E-04 |
| STK10_HUMAN\|O94804_T952 | 0.509210586 | 1.52E-05 | 2.05E-04 |
| DOCK4_HUMAN\|Q8N1I0_S1620 | 0.50919336 | 1.67E-04 | 4.46E-03 |
| CTSL2_HUMAN\|Q05D32_S9 | 0.509184252 | 8.09E-05 | 1.12E-03 |
| DCBD2_HUMAN\|Q96PD2_S724_S727 | 0.509141643 | 5.15E-05 | 1.00E-03 |
| G3BP1_HUMAN\|Q13283_S250 | 0.509106681 | 1.28E-03 | 2.49E-02 |
| DDI2_HUMAN\|Q5TDH0_S194 | 0.50897572 | 1.96E-06 | 2.00E-04 |
| SWAHC_HUMAN\|Q53LP3_S212 | 0.508942344 | 2.73E-08 | 1.74E-06 |
| CF132_HUMAN\|Q5T0Z8_T297 | 0.508873004 | 4.02E-05 | 8.34E-04 |
| MAP4_HUMAN\|P27816_S928 | 0.508804875 | 6.36E-07 | 1.72E-05 |
| NIPA_HUMAN\|Q86WB0_S408_S409 | 0.508732876 | 6.49E-09 | 3.20E-06 |
| SRPK2_HUMAN\|P78362_S497_T492 | 0.508674767 | 2.75E-07 | 1.71E-05 |
| SYTL1_HUMAN\|Q8IYJ3_S216 | 0.508431491 | 1.22E-04 | 3.22E-03 |
| NAV1_HUMAN\|Q8NEY1_S648 | 0.508110526 | 7.12E-04 | 1.67E-02 |
| AHNK_HUMAN\|Q09666_S5749_S5763 | 0.508093682 | 3.00E-09 | 8.12E-06 |
| MK03_HUMAN\|P27361_T198_T202 | 0.507861927 | 4.04E-06 | 3.77E-04 |
| PPR18_HUMAN\|Q6NYC8_S209_S213 | 0.507709721 | 1.08E-07 | 4.53E-06 |
| PKP3_HUMAN\|Q9Y446_S314 | 0.507700201 | 1.25E-07 | 2.25E-05 |
| PKP2_HUMAN\|Q99959_S132 | 0.507686219 | 1.02E-05 | 1.52E-04 |
| PAK2_HUMAN\|Q13177_S192 | 0.507503967 | 1.52E-03 | 8.50E-03 |
| TB10B_HUMAN\|Q4KMP7_S656 | 0.506997545 | 1.09E-05 | 1.56E-04 |
| MA7D1_HUMAN\|Q3KQU3_S452 | 0.506950348 | 3.53E-06 | 6.60E-05 |
| SAFB1_HUMAN\|Q15424_S247 | 0.506850219 | 1.21E-07 | 4.98E-06 |
| TGS1_HUMAN\|Q96RS0_S85_S89 | 0.506619723 | 4.83E-05 | 9.65E-04 |
| ES8L1_HUMAN\|Q8TE68_S182_T187 | 0.50657087 | 6.45E-09 | 2.50E-06 |
| MISP_HUMAN\|Q8IVT2_S395_S400 | 0.506449379 | 9.80E-05 | 2.12E-02 |
| R3HD2_HUMAN\|Q9Y2K5_T856 | 0.506440221 | 6.15E-05 | 4.83E-03 |
| PRC2A_HUMAN\|P48634_S1085_S1089_S1092 | 0.506423871 | 1.34E-07 | 5.39E-06 |
| PKN2_HUMAN\|Q16513_S360 | 0.506342959 | 5.77E-06 | 3.91E-04 |

| | | | |
|---|---|---|---|
| AHNK_HUMAN\|Q09666_S216_S220 | 0.506138755 | 3.99E-08 | 2.30E-06 |
| CRBG1_HUMAN\|Q9Y4K1_S141 | 0.506106266 | 3.60E-06 | 6.81E-05 |
| PKP3_HUMAN\|Q9Y446_S180 | 0.506081939 | 2.95E-06 | 1.21E-04 |
| CRBG1_HUMAN\|Q9Y4K1_S141 | 0.505906303 | 3.58E-06 | 3.40E-04 |
| K2C5_HUMAN\|P13647_S571_S574 | 0.505672765 | 1.43E-06 | 3.24E-05 |
| 3BP1_HUMAN\|Q9Y3L3_S596 | 0.505443497 | 7.29E-07 | 3.95E-05 |
| K1C17_HUMAN\|Q04695_S13 | 0.505441953 | 5.91E-07 | 1.75E-05 |
| TANK_HUMAN\|Q92844_S129 | 0.505366439 | 8.59E-06 | 6.38E-04 |
| TCOF_HUMAN\|Q13428_T249 | 0.505277002 | 1.56E-06 | 8.19E-05 |
| SRS10_HUMAN\|O75494_S106 | 0.505274037 | 2.36E-05 | 5.70E-04 |
| GPBP1_HUMAN\|Q86WP2_S381 | 0.505127898 | 1.52E-06 | 3.61E-05 |
| SVIL_HUMAN\|O95425_T852 | 0.50501549 | 2.26E-04 | 1.82E-03 |
| CE170_HUMAN\|Q5SW79_S971 | 0.504991526 | 3.29E-07 | 1.09E-05 |
| CD97_HUMAN\|P48960_S831 | 0.504953979 | 1.74E-07 | 6.56E-06 |
| PKP1_HUMAN\|Q13835_S121 | 0.50490147 | 2.56E-04 | 1.98E-03 |
| TIF1B_HUMAN\|Q13263_S596_S598_S601 | 0.504870352 | 1.33E-03 | 6.91E-03 |
| K1C15_HUMAN\|P19012_S34 | 0.504696988 | 1.18E-04 | 1.10E-03 |
| DP13A_HUMAN\|Q9UKG1_S401 | 0.504665714 | 3.72E-06 | 6.81E-05 |
| 1B07_HUMAN\|P01889_S359 | 0.504422867 | 2.81E-07 | 9.01E-06 |
| KC1D_HUMAN\|P48730_S383 | 0.504385286 | 9.91E-07 | 2.56E-05 |
| MILK1_HUMAN\|Q8N3F8_T318 | 0.504376115 | 1.18E-04 | 3.14E-03 |
| PHLB2_HUMAN\|Q86SQ0_Y567 | 0.504217946 | 8.97E-08 | 1.68E-05 |
| AKTS1_HUMAN\|Q96B36_S187 | 0.503709947 | 4.93E-06 | 1.74E-04 |
| PP12C_HUMAN\|Q9BZL4_S453 | 0.503634701 | 1.07E-09 | 1.32E-07 |
| K1C17_HUMAN\|Q04695_S11 | 0.503271049 | 6.89E-07 | 1.83E-05 |
| DC1I2_HUMAN\|Q13409-2_S84 | 0.5029232 | 3.24E-07 | 1.91E-05 |
| NU214_HUMAN\|P35658_S1023 | 0.502794535 | 2.00E-06 | 2.00E-04 |
| K1C15_HUMAN\|P19012_S34 | 0.50278675 | 1.10E-04 | 9.82E-04 |
| PHLB2_HUMAN\|Q86SQ0_S334_T328 | 0.50266615 | 4.23E-05 | 8.55E-04 |
| AHNK_HUMAN\|Q09666_S5739 | 0.502587279 | 9.20E-06 | 5.42E-04 |
| CR025_HUMAN\|Q96B23_S143_S145_S147 | 0.502500335 | 3.96E-06 | 7.40E-04 |

| | | | |
|---|---|---|---|
| STIM2_HUMAN\|Q9P246_S522 | 0.50235928 | 1.21E-04 | 1.87E-03 |
| MICA3_HUMAN\|Q7RTP6_S685_S687 | 0.502354165 | 5.25E-05 | 5.08E-03 |
| COHA1_HUMAN\|Q9UMD9_S61_S62_S70 | 0.502235849 | 3.30E-04 | 6.64E-03 |
| WAC2A_HUMAN\|Q641Q2_S375 | 0.501948334 | 1.70E-05 | 2.22E-04 |
| RBP10_HUMAN\|Q6VN20_S365 | 0.501905123 | 2.50E-08 | 1.65E-06 |
| PATL1_HUMAN\|Q86TB9_S179_S184 | 0.501891795 | 1.37E-05 | 3.96E-04 |
| SPD2A_HUMAN\|Q5TCZ1_S644 | 0.501812869 | 1.20E-05 | 6.90E-04 |
| PAR10_HUMAN\|Q53GL7_S431 | 0.501785867 | 5.36E-08 | 4.99E-06 |
| 3BP1_HUMAN\|Q9Y3L3_S596 | 0.50170726 | 6.19E-07 | 1.78E-05 |
| PRC2A_HUMAN\|P48634_S1147 | 0.50165833 | 2.48E-08 | 1.49E-06 |
| COBL1_HUMAN\|Q53SF7_S294_T298 | 0.501508467 | 1.58E-06 | 1.85E-04 |
| F193A_HUMAN\|P78312_S1144 | 0.501401454 | 2.32E-07 | 2.08E-05 |
| RHG23_HUMAN\|Q9P227_S361_S372 | 0.501374277 | 1.54E-09 | 1.74E-07 |
| STMN1_HUMAN\|P16949_S25 | 0.501275947 | 5.06E-08 | 1.19E-05 |
| SARG_HUMAN\|Q9BW04_T151 | 0.501074489 | 2.87E-04 | 2.17E-03 |
| CRK_HUMAN\|P46108_S304 | 0.50093299 | 2.17E-09 | 4.36E-07 |
| SLAI2_HUMAN\|Q9P270_S349_S353_S357 | 0.50092609 | 6.84E-08 | 3.21E-06 |
| MAP7_HUMAN\|Q14244_S202 | 0.500712434 | 2.29E-08 | 3.16E-06 |
| ITPR3_HUMAN\|Q14573_S934 | 0.50058246 | 1.05E-02 | 3.61E-02 |
| IRS2_HUMAN\|Q9Y4H2_T350 | 0.500543967 | 4.88E-08 | 1.19E-05 |
| NEK9_HUMAN\|Q8TD19_T333 | 0.500475199 | 3.13E-05 | 3.61E-04 |
| PAIRB_HUMAN\|Q8NC51_S392 | 0.500433098 | 1.84E-06 | 6.40E-05 |
| T22D3_HUMAN\|Q99576-3_S34 | 0.500410423 | 4.81E-04 | 9.21E-03 |
| E7ETA6_HUMAN\|E7ETA6_T1557 | 0.500365916 | 9.02E-05 | 2.67E-03 |
| PCSK9_HUMAN\|Q8NBP7_S688 | 0.500311088 | 2.03E-06 | 1.35E-04 |
| K1522_HUMAN\|Q9P206_S929 | 0.499865802 | 1.43E-08 | 5.97E-06 |
| OSB11_HUMAN\|Q9BXB4_S181 | 0.499726969 | 5.74E-08 | 2.94E-06 |
| JAM1_HUMAN\|Q9Y624_S284 | 0.499632936 | 1.32E-03 | 6.88E-03 |
| NIBL1_HUMAN\|Q96TA1_S641_S646 | 0.499590951 | 3.18E-05 | 1.38E-03 |
| CAP1_HUMAN\|Q01518_S34 | 0.499358868 | 1.23E-03 | 1.83E-02 |
| CDC20_HUMAN\|Q12834_S41 | 0.498971601 | 2.53E-04 | 6.00E-03 |

| | | | |
|---|---|---|---|
| GTR1_HUMAN\|P11166_S490 | 0.49890604 | 2.26E-03 | 3.73E-02 |
| IASPP_HUMAN\|Q8WUF5_S395 | 0.49867648 | 1.49E-05 | 4.01E-04 |
| AHNK_HUMAN\|Q09666_S5746_S5752 | 0.498579861 | 3.62E-08 | 4.09E-06 |
| LMNB1_HUMAN\|P20700_S23_T20 | 0.498560977 | 1.04E-03 | 9.79E-03 |
| K1C14_HUMAN\|P02533_S51_S54 | 0.498538708 | 6.80E-06 | 1.12E-04 |
| PAK2_HUMAN\|Q13177_S197 | 0.498306778 | 8.91E-04 | 5.55E-03 |
| 4EBP1_HUMAN\|Q13541_T70 | 0.498195311 | 5.68E-07 | 1.06E-04 |
| BAIP2_HUMAN\|Q9UQB8_S366 | 0.498068566 | 5.69E-05 | 5.98E-04 |
| SPD2A_HUMAN\|Q5TCZ1_S1043 | 0.498039704 | 7.91E-06 | 2.61E-04 |
| TRI47_HUMAN\|Q96LD4_S588 | 0.498009094 | 2.31E-06 | 7.45E-05 |
| JMY_HUMAN\|Q8N9B5_S115 | 0.497952637 | 1.08E-05 | 3.30E-04 |
| K1C14_HUMAN\|P02533_S12 | 0.497727437 | 7.50E-08 | 3.42E-06 |
| PKP3_HUMAN\|Q9Y446_S183 | 0.497702137 | 3.17E-06 | 6.07E-05 |
| GRAP1_HUMAN\|Q4V328_S666 | 0.497569352 | 3.24E-06 | 6.18E-05 |
| PKP1_HUMAN\|Q13835_S191 | 0.497463539 | 2.64E-04 | 2.03E-03 |
| KLC3_HUMAN\|Q6P597_S167_S173 | 0.497274303 | 3.25E-06 | 6.19E-05 |
| NDRG1_HUMAN\|Q92597_S330_S333_S336 | 0.49708531 | 1.16E-04 | 1.03E-03 |
| SENP3_HUMAN\|Q9H4L4_S169_S181_T176 | 0.497070496 | 7.30E-05 | 1.27E-03 |
| AHNK2_HUMAN\|Q8IVF2_S1710 | 0.497063261 | 1.69E-06 | 5.97E-04 |
| PHLB1_HUMAN\|Q86UU1_S489 | 0.496901949 | 2.18E-05 | 5.47E-04 |
| SEC62_HUMAN\|Q99442_T375 | 0.496698807 | 1.67E-04 | 1.99E-03 |
| CR025_HUMAN\|Q96B23_S145_S147 | 0.496576568 | 1.51E-08 | 5.28E-06 |
| GAB1_HUMAN\|Q13480_S683 | 0.496555513 | 3.24E-08 | 3.74E-06 |
| COBL1_HUMAN\|Q53SF7_T322 | 0.496533202 | 5.11E-08 | 5.44E-06 |
| NBN_HUMAN\|O60934_S343 | 0.496521805 | 4.43E-07 | 1.31E-05 |
| AFAD_HUMAN\|P55196_S1172_S1173_S1182 | 0.496366456 | 3.05E-08 | 1.88E-06 |
| IF4B_HUMAN\|P23588_S348 | 0.496244549 | 8.31E-03 | 2.98E-02 |
| CE170_HUMAN\|Q5SW79_S633_S636 | 0.496042158 | 8.62E-05 | 1.43E-03 |
| TENS4_HUMAN\|Q8IZW8_S248_S253 | 0.495978562 | 4.50E-05 | 2.07E-03 |
| K1C14_HUMAN\|P02533_S433_S438 | 0.495963068 | 4.60E-06 | 8.08E-05 |
| LMNB2_HUMAN\|Q03252_S420_S424 | 0.495875786 | 9.05E-06 | 1.40E-04 |

| | | | |
|---|---|---|---|
| MA7D1_HUMAN\|Q3KQU3_S313 | 0.495789158 | 1.28E-05 | 1.80E-04 |
| HIRP3_HUMAN\|Q9BW71_S551 | 0.49565276 | 1.80E-04 | 3.64E-03 |
| IMA1_HUMAN\|P52292_S62 | 0.495590518 | 3.86E-03 | 3.87E-02 |
| ARAF_HUMAN\|P10398_S257_T253 | 0.49556941 | 1.61E-06 | 3.74E-05 |
| RU17_HUMAN\|P08621_S320 | 0.495194377 | 1.43E-04 | 2.13E-03 |
| TP53B_HUMAN\|Q12888_S366_T357 | 0.495065284 | 9.95E-07 | 2.56E-05 |
| BAIP2_HUMAN\|Q9UQB8_T360 | 0.494957773 | 3.42E-04 | 4.20E-03 |
| PARP4_HUMAN\|Q9UKK3_S1722 | 0.494894533 | 3.32E-07 | 1.93E-05 |
| K2C5_HUMAN\|P13647_S571_S574 | 0.49489029 | 8.44E-07 | 2.25E-05 |
| LAR4B_HUMAN\|Q92615_T732 | 0.494860294 | 1.21E-09 | 8.18E-07 |
| BAIP2_HUMAN\|Q9UQB8_T340 | 0.494789197 | 7.67E-05 | 7.27E-04 |
| KC1A_HUMAN\|P48729-3_T321 | 0.49460944 | 5.61E-04 | 3.81E-03 |
| CTNA1_HUMAN\|P35221_S652_S655_T658 | 0.494523474 | 2.60E-05 | 3.07E-04 |
| K1C17_HUMAN\|Q04695_S10 | 0.494439336 | 1.20E-06 | 2.85E-05 |
| LAD1_HUMAN\|O00515_S485 | 0.494404865 | 2.05E-06 | 2.02E-04 |
| K2C6A_HUMAN\|P02538_S546 | 0.494326648 | 1.62E-03 | 1.40E-02 |
| SRRM2_HUMAN\|Q9UQ35_S2692 | 0.494316651 | 1.05E-04 | 9.47E-04 |
| LAR4B_HUMAN\|Q92615_S731_S736 | 0.493927017 | 1.08E-06 | 2.72E-05 |
| LIPA1_HUMAN\|Q13136_S666_S668 | 0.493808899 | 6.63E-08 | 3.21E-06 |
| EGFR_HUMAN\|P00533_S1204 | 0.493656056 | 1.56E-05 | 2.07E-04 |
| PRC2A_HUMAN\|P48634_S1325_S1328_T1323 | 0.493522532 | 6.97E-04 | 1.66E-02 |
| PRC2A_HUMAN\|P48634_S761_S764 | 0.493317078 | 5.78E-09 | 4.90E-07 |
| BICD2_HUMAN\|Q8TD16_S582 | 0.493269502 | 1.35E-03 | 7.74E-03 |
| PACS1_HUMAN\|Q6VY07_T504 | 0.493216832 | 4.82E-06 | 1.71E-04 |
| AHNK_HUMAN\|Q09666_T158 | 0.492923484 | 1.46E-04 | 1.21E-03 |
| PDLI2_HUMAN\|Q96JY6_S137_S143 | 0.49270885 | 3.19E-07 | 2.11E-05 |
| MTCL1_HUMAN\|Q9Y4B5_S196 | 0.492649193 | 2.23E-06 | 9.53E-05 |
| ABLM1_HUMAN\|O14639_S458 | 0.492614807 | 2.24E-05 | 5.42E-04 |
| K1C14_HUMAN\|P02533_S435_S437 | 0.492416939 | 1.21E-05 | 1.73E-04 |
| STMN1_HUMAN\|P16949_S25 | 0.492361631 | 3.12E-08 | 8.46E-06 |
| CRBG1_HUMAN\|Q9Y4K1_S141 | 0.492295374 | 2.07E-06 | 4.30E-05 |

| | | | |
|---|---|---|---|
| HIRP3_HUMAN\|Q9BW71_S555 | 0.492258496 | 5.39E-03 | 2.84E-02 |
| ZYX_HUMAN\|Q15942_S259 | 0.492069923 | 4.20E-08 | 2.35E-06 |
| SCRIB_HUMAN\|Q14160_S1285 | 0.491976284 | 9.47E-06 | 6.60E-04 |
| LARP1_HUMAN\|Q6PKG0_S766 | 0.491723224 | 4.62E-04 | 3.23E-03 |
| LIPA1_HUMAN\|Q13136_S1172 | 0.491577882 | 1.27E-08 | 1.01E-06 |
| K2C75_HUMAN\|O95678_S61 | 0.491488474 | 5.05E-05 | 9.89E-04 |
| CE170_HUMAN\|Q5SW79_S627 | 0.491348731 | 3.11E-03 | 3.17E-02 |
| AHNK_HUMAN\|Q09666_S212_S216 | 0.491335582 | 2.34E-08 | 3.16E-06 |
| ABI2_HUMAN\|Q9NYB9_Y371 | 0.491163661 | 3.27E-08 | 1.81E-06 |
| MA7D1_HUMAN\|Q3KQU3_S452 | 0.490742529 | 1.73E-06 | 1.85E-04 |
| PDLI2_HUMAN\|Q96JY6_S121_S124_S127 | 0.490660701 | 4.57E-03 | 4.43E-02 |
| TP53B_HUMAN\|Q12888_S356 | 0.490311107 | 1.33E-04 | 4.96E-03 |
| WDR7_HUMAN\|Q9Y4E6_S1153 | 0.490124916 | 1.02E-04 | 2.88E-03 |
| HMGA1_HUMAN\|P17096_S49 | 0.490044613 | 2.55E-04 | 1.88E-03 |
| FGD6_HUMAN\|Q6ZV73_T693 | 0.489982518 | 7.63E-07 | 9.57E-05 |
| MB12A_HUMAN\|Q96EY5_S188 | 0.48994854 | 1.17E-05 | 1.66E-04 |
| FLNB_HUMAN\|O75369-8_S1474 | 0.489678876 | 4.01E-04 | 3.96E-03 |
| K2C5_HUMAN\|P13647_T19 | 0.4894309 | 4.84E-05 | 9.56E-04 |
| ATN1_HUMAN\|P54259_S77_S79 | 0.488938734 | 5.33E-06 | 9.28E-05 |
| CBX3_HUMAN\|Q13185_S93 | 0.488866615 | 6.40E-05 | 5.88E-03 |
| MILK1_HUMAN\|Q8N3F8_T318 | 0.488696969 | 7.18E-05 | 6.89E-04 |
| GAB1_HUMAN\|Q13480_S683 | 0.48864486 | 2.14E-08 | 7.74E-06 |
| IRS2_HUMAN\|Q9Y4H2_S608 | 0.48824027 | 4.20E-08 | 2.23E-06 |
| VIME_HUMAN\|P08670_S34 | 0.488226718 | 4.31E-07 | 1.39E-05 |
| SRS10_HUMAN\|O75494_S106 | 0.488071362 | 1.14E-05 | 1.68E-04 |
| FOXO3_HUMAN\|O43524_S294 | 0.487746545 | 5.20E-06 | 1.39E-04 |
| PDLI1_HUMAN\|O00151_S130 | 0.487578899 | 7.01E-08 | 3.16E-05 |
| RELB_HUMAN\|Q01201_S573 | 0.487492612 | 7.64E-08 | 6.37E-06 |
| REQU_HUMAN\|Q92785_S142 | 0.487449057 | 5.94E-07 | 1.75E-05 |
| K2C5_HUMAN\|P13647_S21 | 0.486652081 | 1.44E-03 | 1.29E-02 |
| PCSK9_HUMAN\|Q8NBP7_S688 | 0.486355349 | 1.06E-06 | 6.07E-05 |

| | | | |
|---|---|---|---|
| K1143_HUMAN\|Q96AT1_S146 | 0.486204694 | 8.91E-06 | 2.76E-04 |
| WDR13_HUMAN\|Q9H1Z4_S70_S74 | 0.486035794 | 1.57E-07 | 3.15E-05 |
| UBXN7_HUMAN\|O94888_S280_S288 | 0.485979643 | 4.22E-05 | 4.53E-04 |
| MTCL1_HUMAN\|Q9Y4B5_S1679_T1675 | 0.485956993 | 1.90E-08 | 3.82E-06 |
| NGAP_HUMAN\|Q9UJF2_S946_S954 | 0.485821104 | 3.43E-03 | 3.37E-02 |
| K1C14_HUMAN\|P02533_S435 | 0.485737783 | 2.40E-06 | 4.89E-05 |
| CR025_HUMAN\|Q96B23_S143_S145_S147 | 0.485695464 | 1.88E-06 | 1.96E-04 |
| DC1L2_HUMAN\|O43237_S383 | 0.485692483 | 3.41E-07 | 1.11E-05 |
| CR025_HUMAN\|Q96B23_S67 | 0.485641331 | 7.61E-06 | 5.07E-04 |
| DEN1A_HUMAN\|Q8TEH3_T532 | 0.485630573 | 1.94E-04 | 2.24E-03 |
| KC1A_HUMAN\|P48729_S312 | 0.485628412 | 4.10E-04 | 2.70E-03 |
| CAVN3_HUMAN\|Q969G5_T205 | 0.485594163 | 3.36E-06 | 1.34E-04 |
| PHLB2_HUMAN\|Q86SQ0_T898 | 0.485414759 | 1.37E-04 | 2.05E-03 |
| DLG1_HUMAN\|Q12959_S573 | 0.485258441 | 4.09E-05 | 8.55E-04 |
| K2C5_HUMAN\|P13647_S82_T78 | 0.485152362 | 8.58E-08 | 8.01E-06 |
| LARP1_HUMAN\|Q6PKG0_S1040 | 0.485068401 | 6.87E-08 | 3.21E-06 |
| MAP4_HUMAN\|P27816_S1073 | 0.485013238 | 1.02E-07 | 3.08E-05 |
| TENS4_HUMAN\|Q8IZW8_S242_S248_S253 | 0.484813404 | 1.51E-04 | 3.82E-03 |
| PEA15_HUMAN\|Q15121_S116 | 0.484628272 | 7.61E-06 | 1.18E-03 |
| UBP5_HUMAN\|P45974_S785 | 0.484577976 | 2.25E-04 | 1.71E-03 |
| PACS1_HUMAN\|Q6VY07_T504 | 0.48453587 | 3.46E-06 | 6.50E-05 |
| GPBP1_HUMAN\|Q86WP2_S381 | 0.484498889 | 5.74E-07 | 7.41E-05 |
| BCAR3_HUMAN\|O75815_S375 | 0.48437709 | 1.59E-06 | 3.74E-05 |
| HMGA2_HUMAN\|P52926_S101_S105_T100 | 0.484219807 | 7.57E-07 | 1.98E-05 |
| TB182_HUMAN\|Q9C0C2_T875 | 0.484110518 | 4.54E-08 | 2.35E-06 |
| WFS1_HUMAN\|O76024_S235 | 0.484102627 | 1.02E-04 | 1.33E-03 |
| BAIP2_HUMAN\|Q9UQB8_T360 | 0.483861565 | 2.47E-04 | 1.94E-03 |
| EGFR_HUMAN\|P00533_S1039 | 0.483528826 | 2.82E-03 | 2.73E-02 |
| PHF14_HUMAN\|O94880_T287 | 0.483304096 | 5.98E-07 | 1.75E-05 |
| K2C6A_HUMAN\|P02538_S541 | 0.483166478 | 7.45E-04 | 4.33E-03 |
| COBL1_HUMAN\|Q53SF7_T298 | 0.482836995 | 1.01E-06 | 2.50E-05 |

| | | | |
|---|---|---|---|
| MAN1_HUMAN|Q9Y2U8_S144 | 0.482778574 | 1.27E-05 | 4.52E-04 |
| NDE1_HUMAN|Q9NXR1_S282 | 0.482741618 | 5.61E-05 | 1.08E-03 |
| K2C8_HUMAN|P05787_S477 | 0.482610014 | 3.54E-06 | 6.60E-05 |
| LMNA_HUMAN|P02545_S616_S619 | 0.482575943 | 3.58E-03 | 2.43E-02 |
| EPN1_HUMAN|Q9Y6I3_S454 | 0.48252369 | 2.39E-08 | 3.15E-06 |
| NUP53_HUMAN|Q8NFH5_S105 | 0.482486651 | 2.11E-06 | 9.96E-05 |
| EXPH5_HUMAN|Q8NEV8_S341 | 0.481990733 | 1.43E-07 | 5.58E-06 |
| AHNK_HUMAN|Q09666_S5752 | 0.481957359 | 1.40E-09 | 1.85E-07 |
| PKP1_HUMAN|Q13835_S185 | 0.48178399 | 1.70E-04 | 1.47E-03 |
| M3K20_HUMAN|Q9NYL2_S727 | 0.481604835 | 2.19E-09 | 2.34E-07 |
| CR025_HUMAN|Q96B23_S69 | 0.48158936 | 1.59E-04 | 4.00E-03 |
| OSB10_HUMAN|Q9BXB5_S32 | 0.48157972 | 1.98E-07 | 3.35E-05 |
| GGYF2_HUMAN|Q6Y7W6_S30 | 0.481573509 | 2.02E-04 | 1.57E-03 |
| K1C14_HUMAN|P02533_S44 | 0.481444786 | 3.24E-06 | 1.31E-04 |
| SPTB2_HUMAN|Q01082-3_S14 | 0.481397198 | 1.23E-06 | 1.39E-04 |
| CBPC1_HUMAN|Q9UPW5_T1170 | 0.48135788 | 1.05E-03 | 1.48E-02 |
| FA83H_HUMAN|Q6ZRV2_S513_S516 | 0.481338884 | 8.75E-06 | 1.33E-04 |
| 3BP1_HUMAN|Q9Y3L3_S596 | 0.48110251 | 2.47E-07 | 3.92E-05 |
| K2C5_HUMAN|P13647_S21_S26 | 0.481034765 | 1.96E-06 | 8.75E-05 |
| ZDHC5_HUMAN|Q9C0B5_S704 | 0.481017844 | 2.80E-05 | 1.50E-03 |
| STX5_HUMAN|Q13190_S9 | 0.48096397 | 4.62E-05 | 4.93E-04 |
| NDE1_HUMAN|Q9NXR1_S282 | 0.480873968 | 5.26E-05 | 5.56E-04 |
| K2C6A_HUMAN|P02538_S37 | 0.480806719 | 1.62E-03 | 1.40E-02 |
| RTKN_HUMAN|Q9BST9_S520_S529 | 0.480794102 | 9.32E-07 | 1.44E-04 |
| FA83H_HUMAN|Q6ZRV2_S1025 | 0.480703548 | 1.41E-07 | 5.53E-06 |
| GYS1_HUMAN|P13807_S727_S731 | 0.48059347 | 3.04E-06 | 9.31E-05 |
| TMF1_HUMAN|P82094_S72_S77 | 0.480055256 | 1.52E-05 | 3.12E-04 |
| K1C14_HUMAN|P02533_S32 | 0.480014616 | 5.15E-08 | 2.61E-06 |
| EXPH5_HUMAN|Q8NEV8_S341 | 0.479816701 | 1.29E-07 | 5.25E-06 |
| PKP3_HUMAN|Q9Y446_S123 | 0.479750992 | 2.95E-06 | 5.89E-05 |
| PATL1_HUMAN|Q86TB9_S179_S184 | 0.479600726 | 6.07E-06 | 1.01E-04 |

| | | | |
|---|---|---|---|
| FOSL1_HUMAN\|P15407_S252_S265 | 0.479506727 | 1.15E-04 | 1.48E-03 |
| K1522_HUMAN\|Q9P206_S612 | 0.479421249 | 2.93E-06 | 2.61E-04 |
| CCDC9_HUMAN\|Q9Y3X0_S376_S386_T381 | 0.479351169 | 1.17E-07 | 1.27E-05 |
| LAP2A_HUMAN\|P42166_S167_T160 | 0.479299526 | 4.54E-07 | 1.33E-05 |
| SOS2_HUMAN\|Q07890_S1267 | 0.479200864 | 5.37E-06 | 2.02E-04 |
| K2C5_HUMAN\|P13647_S572 | 0.47899913 | 3.66E-07 | 1.15E-05 |
| NGAP_HUMAN\|Q9UJF2_T620 | 0.478991598 | 1.86E-04 | 1.58E-03 |
| TCOF_HUMAN\|Q13428_T249 | 0.478924776 | 4.53E-07 | 6.14E-05 |
| TP53B_HUMAN\|Q12888_S356 | 0.478878785 | 9.32E-05 | 9.12E-04 |
| SEPT7_HUMAN\|Q16181_S423 | 0.478841166 | 1.48E-05 | 2.01E-04 |
| PKP3_HUMAN\|Q9Y446_S280 | 0.478735112 | 1.68E-05 | 2.21E-04 |
| TRI22_HUMAN\|Q8IYM9_S383 | 0.478582737 | 4.31E-05 | 3.70E-03 |
| NDRG2_HUMAN\|Q9UN36_S332_T330 | 0.478247843 | 3.84E-08 | 2.08E-06 |
| LAD1_HUMAN\|O00515_S512 | 0.478024736 | 4.76E-08 | 2.87E-05 |
| K2C5_HUMAN\|P13647_S18 | 0.477890264 | 1.49E-07 | 5.85E-06 |
| AHNK_HUMAN\|Q09666_S5749_S5763 | 0.477883683 | 4.54E-10 | 4.66E-07 |
| LMNB2_HUMAN\|Q03252_S424 | 0.477872228 | 7.69E-07 | 2.10E-05 |
| PHLB2_HUMAN\|Q86SQ0_S565 | 0.477857291 | 2.37E-08 | 7.14E-06 |
| NDRG1_HUMAN\|Q92597_S333_S336_T328 | 0.477509737 | 4.78E-09 | 1.36E-06 |
| MISP_HUMAN\|Q8IVT2_S395_S400 | 0.477499379 | 3.80E-05 | 3.51E-03 |
| LPP_HUMAN\|Q93052_S609 | 0.477468813 | 6.78E-06 | 1.11E-04 |
| SRRM2_HUMAN\|Q9UQ35_S1152 | 0.477425428 | 5.90E-06 | 4.51E-04 |
| PKP3_HUMAN\|Q9Y446_S180_S183 | 0.477408927 | 1.75E-06 | 7.93E-05 |
| CND1_HUMAN\|Q15021_S1333 | 0.477399102 | 6.36E-08 | 5.65E-06 |
| UTP18_HUMAN\|Q9Y5J1_S121 | 0.477218933 | 1.47E-03 | 1.67E-02 |
| CCDC9_HUMAN\|Q9Y3X0_S376 | 0.477159083 | 4.85E-08 | 2.60E-06 |
| LAD1_HUMAN\|O00515_S38_T40_T41 | 0.476718494 | 1.63E-06 | 1.51E-04 |
| SRPK2_HUMAN\|P78362_S494_S497 | 0.476697933 | 1.17E-07 | 8.59E-06 |
| DOCK4_HUMAN\|Q8N1I0_S1620 | 0.476637329 | 5.54E-05 | 2.07E-03 |
| ERBB2_HUMAN\|P04626_S998 | 0.475583433 | 2.03E-05 | 3.89E-04 |
| K1522_HUMAN\|Q9P206_S161 | 0.475581192 | 1.16E-08 | 8.40E-07 |

| | | | |
|---|---|---|---|
| COHA1_HUMAN\|Q9UMD9_S93_T97 | 0.47540656 | 1.01E-05 | 3.03E-04 |
| ARP19_HUMAN\|P56211_Y59 | 0.475229343 | 2.38E-08 | 1.45E-06 |
| SPD2A_HUMAN\|Q5TCZ1_S1043 | 0.475087531 | 3.27E-06 | 6.19E-05 |
| MISP_HUMAN\|Q8IVT2_S155 | 0.475024714 | 8.47E-06 | 6.12E-04 |
| CR025_HUMAN\|Q96B23_S66_S67_T74 | 0.47499565 | 3.50E-03 | 3.18E-02 |
| HMGA2_HUMAN\|P52926_S101_S105_T100 | 0.474765348 | 4.70E-07 | 1.48E-05 |
| TRI22_HUMAN\|Q8IYM9_S384 | 0.474443274 | 6.66E-07 | 1.72E-04 |
| DOCK4_HUMAN\|Q8N1I0_S1620 | 0.474342395 | 5.11E-05 | 1.26E-02 |
| TB182_HUMAN\|Q9C0C2_S920 | 0.47416155 | 4.41E-07 | 2.30E-05 |
| DESP_HUMAN\|P15924_S22 | 0.474041212 | 6.81E-06 | 2.24E-04 |
| EGFR_HUMAN\|P00533_S1204 | 0.473867315 | 6.83E-06 | 1.12E-04 |
| PR38B_HUMAN\|Q5VTL8_S529 | 0.473584071 | 4.48E-08 | 2.34E-06 |
| OSB11_HUMAN\|Q9BXB4_S174_S181 | 0.473545396 | 4.67E-05 | 4.96E-04 |
| AFAD_HUMAN\|P55196_S1799 | 0.473398471 | 2.00E-06 | 4.45E-05 |
| AKTS1_HUMAN\|Q96B36_S187 | 0.473325229 | 1.32E-06 | 1.46E-04 |
| CE170_HUMAN\|Q5SW79_S971 | 0.473168519 | 6.82E-08 | 3.21E-06 |
| DBNL_HUMAN\|Q9UJU6_T271 | 0.473158655 | 1.79E-07 | 6.60E-06 |
| IRS2_HUMAN\|Q9Y4H2_T350 | 0.472917086 | 1.19E-08 | 4.30E-06 |
| GAB1_HUMAN\|Q13480_S683 | 0.47287583 | 9.32E-09 | 4.21E-06 |
| LMNA_HUMAN\|P02545_S619 | 0.472819588 | 1.61E-06 | 3.74E-05 |
| SRRM2_HUMAN\|Q9UQ35_S2692 | 0.472802636 | 5.21E-05 | 5.52E-04 |
| RPRD2_HUMAN\|Q5VT52_T723 | 0.472757736 | 1.02E-09 | 1.50E-07 |
| NCK5L_HUMAN\|Q9HCH0_S489_S494 | 0.47268506 | 5.19E-08 | 2.73E-06 |
| TACC1_HUMAN\|O75410_S44 | 0.472613808 | 1.08E-07 | 4.51E-06 |
| K1522_HUMAN\|Q9P206_S404_S409 | 0.472606359 | 1.25E-08 | 8.79E-07 |
| NAV1_HUMAN\|Q8NEY1_T1006 | 0.472588475 | 8.06E-06 | 2.63E-04 |
| MYPT1_HUMAN\|O14974_S554 | 0.472512861 | 1.27E-04 | 1.94E-03 |
| RHG21_HUMAN\|Q5T5U3_S615 | 0.472474437 | 1.26E-08 | 1.01E-06 |
| TENS4_HUMAN\|Q8IZW8_S248_S253 | 0.47243128 | 1.98E-05 | 2.53E-04 |
| K2C8_HUMAN\|P05787_S21_S24_S27 | 0.472395623 | 3.31E-07 | 1.09E-05 |
| AHNK_HUMAN\|Q09666_S5749 | 0.472125247 | 1.83E-10 | 4.32E-08 |

| | | | |
|---|---|---|---|
| DPYL3_HUMAN\|Q14195_S568 | 0.472006524 | 3.12E-05 | 1.60E-03 |
| NUP53_HUMAN\|Q8NFH5_S105 | 0.471778953 | 1.36E-06 | 1.46E-04 |
| MTCL1_HUMAN\|Q9Y4B5_S1679_T1675 | 0.471646317 | 9.21E-09 | 3.33E-06 |
| MTCL1_HUMAN\|Q9Y4B5_S1679_T1675 | 0.471238547 | 9.02E-09 | 6.79E-07 |
| SPD2A_HUMAN\|Q5TCZ1_S651 | 0.471220575 | 2.72E-06 | 5.40E-05 |
| TB182_HUMAN\|Q9C0C2_S1652 | 0.471189067 | 1.02E-07 | 2.12E-05 |
| K2C5_HUMAN\|P13647_S18_S21 | 0.470774037 | 7.25E-05 | 1.27E-03 |
| TB182_HUMAN\|Q9C0C2_S1652 | 0.470750799 | 9.94E-08 | 8.83E-06 |
| TARA_HUMAN\|Q9H2D6_S1955 | 0.470710273 | 3.10E-05 | 6.87E-04 |
| PKP3_HUMAN\|Q9Y446_S123 | 0.470691842 | 2.06E-06 | 4.29E-05 |
| K2C6A_HUMAN\|P02538_S22 | 0.470649977 | 5.08E-04 | 3.50E-03 |
| DFFA_HUMAN\|O00273-2_S257 | 0.47052814 | 2.48E-04 | 1.94E-03 |
| MEF2D_HUMAN\|Q14814_S231 | 0.470509737 | 2.18E-05 | 2.68E-04 |
| K1C19_HUMAN\|P08727_S22 | 0.469826531 | 4.33E-07 | 1.30E-05 |
| K2C5_HUMAN\|P13647_S574 | 0.469713179 | 2.51E-08 | 1.65E-06 |
| DP13A_HUMAN\|Q9UKG1_S401 | 0.469676642 | 8.91E-07 | 1.05E-04 |
| AHNK2_HUMAN\|Q8IVF2_S457 | 0.469658067 | 4.86E-05 | 5.11E-04 |
| BORG2_HUMAN\|Q9UKI2_S100 | 0.469601183 | 8.43E-07 | 3.65E-05 |
| K1671_HUMAN\|Q9BY89_S1441 | 0.469567767 | 3.84E-06 | 6.97E-05 |
| MAP4_HUMAN\|P27816_S1073 | 0.469303321 | 4.46E-08 | 2.47E-06 |
| CLIP1_HUMAN\|P30622_S200 | 0.469219938 | 7.30E-06 | 1.17E-04 |
| RSF1_HUMAN\|Q96T23_S1277_T1278 | 0.469136218 | 6.30E-03 | 4.69E-02 |
| TB182_HUMAN\|Q9C0C2_T1282 | 0.469049519 | 3.99E-08 | 1.66E-05 |
| NAV1_HUMAN\|Q8NEY1_S648 | 0.46888229 | 2.34E-04 | 1.77E-03 |
| SARG_HUMAN\|Q9BW04_S502 | 0.468829818 | 3.04E-09 | 9.69E-07 |
| PP1R7_HUMAN\|Q15435_S24 | 0.468821225 | 9.27E-06 | 3.78E-04 |
| MET_HUMAN\|P08581_S990_S997 | 0.468591444 | 2.13E-03 | 4.47E-02 |
| IF4B_HUMAN\|P23588_S497_S504_T500 | 0.468533256 | 3.56E-07 | 1.93E-05 |
| TB182_HUMAN\|Q9C0C2_S429 | 0.468220036 | 2.50E-05 | 3.00E-04 |
| AHNK_HUMAN\|Q09666_S5739 | 0.46809524 | 2.26E-06 | 2.01E-04 |
| SEPT9_HUMAN\|Q9UHD8_T38_T42 | 0.468047235 | 1.39E-08 | 3.14E-06 |

| | | | |
|---|---|---|---|
| AHNK2_HUMAN\|Q8IVF2_S1583 | 0.468035994 | 4.48E-06 | 7.89E-05 |
| AHNK_HUMAN\|Q09666_S4092 | 0.468005015 | 2.27E-05 | 2.81E-04 |
| PACN3_HUMAN\|Q9UKS6_S341_S354 | 0.468004977 | 2.17E-03 | 1.47E-02 |
| NP1L1_HUMAN\|P55209_T62 | 0.467631253 | 1.90E-05 | 1.08E-03 |
| MPLKI_HUMAN\|Q8TAP9_S80 | 0.467525883 | 7.47E-05 | 7.11E-04 |
| KC1A_HUMAN\|P48729-3_T321 | 0.467433 | 2.41E-04 | 1.81E-03 |
| ANXA2_HUMAN\|P07355_S127 | 0.467397366 | 9.35E-05 | 9.12E-04 |
| MK01_HUMAN\|P28482_T185_Y187 | 0.467248725 | 2.41E-06 | 1.02E-04 |
| SPTB2_HUMAN\|Q01082_T2328 | 0.467105143 | 1.55E-04 | 5.62E-03 |
| CAVN3_HUMAN\|Q969G5_T205 | 0.466951003 | 1.55E-06 | 3.44E-05 |
| B4DKV4_HUMAN\|B4DKV4_S22 | 0.466931357 | 1.29E-04 | 1.96E-03 |
| IF2B_HUMAN\|P20042_T111 | 0.466875256 | 2.49E-06 | 5.26E-05 |
| PKP1_HUMAN\|Q13835_S121 | 0.466827117 | 6.75E-05 | 6.57E-04 |
| DBNL_HUMAN\|Q9UJU6_S269 | 0.466672013 | 2.82E-06 | 5.72E-05 |
| DC1I2_HUMAN\|Q13409-2_S81 | 0.466544227 | 1.60E-05 | 3.25E-04 |
| TARA_HUMAN\|Q9H2D6_S1955 | 0.466346934 | 2.68E-05 | 1.19E-03 |
| LASP1_HUMAN\|Q14847_T104 | 0.466160583 | 1.49E-08 | 3.23E-06 |
| ETV3_HUMAN\|P41162_S139 | 0.46612477 | 1.06E-04 | 1.01E-03 |
| PKP3_HUMAN\|Q9Y446_S313_S314 | 0.465991169 | 1.56E-08 | 5.28E-06 |
| NGAP_HUMAN\|Q9UJF2_S946_S954 | 0.465981879 | 2.23E-03 | 4.57E-02 |
| ITB4_HUMAN\|P16144_S1486_T1487 | 0.465952587 | 5.77E-06 | 1.98E-04 |
| CE170_HUMAN\|Q5SW79_S630_S636 | 0.465858143 | 6.31E-09 | 5.70E-07 |
| IASPP_HUMAN\|Q8WUF5_S395 | 0.465555347 | 4.25E-06 | 7.58E-05 |
| HS90A_HUMAN\|P07900_S726 | 0.46540807 | 8.81E-04 | 5.51E-03 |
| PKP1_HUMAN\|Q13835_S205 | 0.465377469 | 6.83E-04 | 4.48E-03 |
| GRB10_HUMAN\|Q13322_S104 | 0.465336276 | 9.84E-10 | 1.26E-07 |
| K1522_HUMAN\|Q9P206_S929 | 0.465301876 | 1.83E-09 | 6.61E-07 |
| SLK_HUMAN\|Q9H2G2_S779 | 0.465290157 | 5.56E-05 | 5.57E-04 |
| COHA1_HUMAN\|Q9UMD9_T182 | 0.465255947 | 2.63E-06 | 1.18E-04 |
| IASPP_HUMAN\|Q8WUF5_T341 | 0.465127156 | 1.86E-04 | 1.58E-03 |
| PARD3_HUMAN\|Q8TEW0_S225 | 0.4651104 | 1.07E-07 | 4.51E-06 |

| | | | |
|---|---|---|---|
| TOM1_HUMAN\|O60784_S462 | 0.465053954 | 1.12E-06 | 4.45E-05 |
| TB182_HUMAN\|Q9C0C2_S1652 | 0.464988756 | 7.44E-08 | 9.01E-06 |
| PAK2_HUMAN\|Q13177_S197 | 0.464825522 | 3.66E-04 | 2.50E-03 |
| ZO2_HUMAN\|Q9UDY2_T1027 | 0.464680579 | 3.22E-08 | 3.88E-06 |
| LMNB2_HUMAN\|Q03252_S420_S424 | 0.464243583 | 2.19E-06 | 4.75E-05 |
| LAD1_HUMAN\|O00515_S301 | 0.464177078 | 8.99E-07 | 1.02E-04 |
| EXOC7_HUMAN\|Q9UPT5_S250 | 0.463850195 | 5.51E-08 | 2.87E-06 |
| CCDC9_HUMAN\|Q9Y3X0_S386_T381 | 0.463408825 | 8.04E-08 | 3.66E-06 |
| K1C13_HUMAN\|P13646_S11 | 0.463335919 | 3.05E-04 | 2.28E-03 |
| BL1S3_HUMAN\|Q6QNY0_S65 | 0.463042063 | 4.03E-04 | 6.63E-03 |
| NDRG1_HUMAN\|Q92597_S330_S336_T328 | 0.462867534 | 4.06E-04 | 8.43E-03 |
| K1C14_HUMAN\|P02533_S44_S53_S54 | 0.462457314 | 1.82E-08 | 1.18E-06 |
| COHA1_HUMAN\|Q9UMD9_S85_S93_T97 | 0.462338278 | 8.74E-06 | 1.33E-04 |
| CR025_HUMAN\|Q96B23_S66_S69 | 0.462300563 | 3.28E-05 | 1.63E-03 |
| R3HD2_HUMAN\|Q9Y2K5_S349 | 0.462033489 | 2.56E-04 | 8.16E-03 |
| CBX3_HUMAN\|Q13185_S93 | 0.461720988 | 2.55E-05 | 4.67E-04 |
| SPAS2_HUMAN\|Q86XZ4_S538 | 0.461648249 | 9.56E-05 | 3.70E-03 |
| K1C18_HUMAN\|P05783_S7 | 0.461378918 | 4.45E-06 | 1.63E-04 |
| K2C5_HUMAN\|P13647_S16_S21 | 0.461315901 | 2.69E-05 | 6.13E-04 |
| K1C17_HUMAN\|Q04695_S28 | 0.461201184 | 5.85E-07 | 1.63E-05 |
| AHNK_HUMAN\|Q09666_T5824 | 0.460826906 | 4.00E-04 | 4.75E-03 |
| S20A2_HUMAN\|Q08357_Y423 | 0.460768392 | 1.15E-05 | 1.68E-04 |
| NDRG1_HUMAN\|Q92597_S330_T328 | 0.460733374 | 7.31E-08 | 3.48E-06 |
| NBEL1_HUMAN\|Q6ZS30_S733 | 0.460666043 | 7.09E-03 | 4.17E-02 |
| ERCC5_HUMAN\|P28715_S382_S384 | 0.460495647 | 4.49E-06 | 8.14E-05 |
| LAD1_HUMAN\|O00515_S38_S39 | 0.460439941 | 1.53E-05 | 3.15E-04 |
| JUPI2_HUMAN\|Q9H910_S97 | 0.460438049 | 4.65E-07 | 2.71E-05 |
| RAI14_HUMAN\|Q9P0K7-4_S261 | 0.460412827 | 6.35E-08 | 1.43E-05 |
| ATN1_HUMAN\|P54259_S77_S79 | 0.460297057 | 1.46E-06 | 3.30E-05 |
| FGD6_HUMAN\|Q6ZV73_S721 | 0.460177732 | 1.25E-06 | 4.79E-05 |
| DBNL_HUMAN\|Q9UJU6_T270 | 0.460109731 | 2.88E-06 | 5.78E-05 |

| | | | |
|---|---|---|---|
| FOSL2_HUMAN\|P15408_S19 | 0.460019765 | 2.26E-05 | 8.34E-04 |
| R3HD1_HUMAN\|Q15032_S973 | 0.460019391 | 9.99E-10 | 9.34E-07 |
| FAT1_HUMAN\|Q14517_S4473 | 0.459780687 | 9.16E-06 | 1.35E-04 |
| SSFA2_HUMAN\|P28290_S92 | 0.459458631 | 1.54E-04 | 1.36E-03 |
| K1C14_HUMAN\|P02533_S17 | 0.459400076 | 9.73E-06 | 1.47E-04 |
| RAI14_HUMAN\|Q9P0K7_S290 | 0.459379802 | 5.15E-06 | 8.86E-05 |
| MISP_HUMAN\|Q8IVT2_S395_S400 | 0.459270065 | 2.05E-05 | 2.60E-04 |
| R3HD1_HUMAN\|Q15032_S973 | 0.459024682 | 9.41E-10 | 8.50E-07 |
| 2A5E_HUMAN\|Q16537_S32_S34 | 0.45899413 | 1.01E-05 | 3.03E-04 |
| RBM15_HUMAN\|Q96T37_S656 | 0.458989409 | 4.77E-04 | 8.95E-03 |
| PP14A_HUMAN\|Q96A00_S128_S134_S136 | 0.458678793 | 2.00E-09 | 6.77E-07 |
| LMNB1_HUMAN\|P20700_S23_T20 | 0.458672507 | 3.54E-04 | 7.20E-03 |
| K1C14_HUMAN\|P02533_S32 | 0.458571839 | 1.51E-08 | 2.41E-06 |
| HIRP3_HUMAN\|Q9BW71_S555 | 0.458359872 | 2.66E-03 | 2.60E-02 |
| SRC8_HUMAN\|Q14247_Y446 | 0.458250837 | 1.38E-03 | 7.13E-03 |
| PKP3_HUMAN\|Q9Y446_S183 | 0.458113596 | 6.15E-07 | 1.78E-05 |
| UBXN7_HUMAN\|O94888_S285 | 0.457931286 | 8.72E-06 | 1.33E-04 |
| ATAT_HUMAN\|Q5SQI0_S315 | 0.457900421 | 1.92E-10 | 2.08E-07 |
| UBXN7_HUMAN\|O94888_S288 | 0.457813626 | 3.63E-04 | 1.07E-02 |
| AHNK_HUMAN\|Q09666_S5752_S5763 | 0.457670219 | 2.22E-06 | 2.10E-04 |
| FA83B_HUMAN\|Q5T0W9_T687 | 0.457643001 | 1.26E-03 | 1.11E-02 |
| SRRM1_HUMAN\|Q8IYB3_S738_S743 | 0.457632854 | 7.82E-05 | 1.42E-03 |
| K2C6A_HUMAN\|P02538_S547 | 0.457540601 | 1.08E-03 | 6.49E-03 |
| SARG_HUMAN\|Q9BW04_T151 | 0.457417095 | 7.57E-05 | 2.74E-03 |
| ERBB2_HUMAN\|P04626_S998 | 0.457381777 | 9.68E-06 | 6.60E-04 |
| CHSP1_HUMAN\|Q9Y2V2_S52 | 0.457112062 | 7.91E-07 | 2.03E-05 |
| AHNK_HUMAN\|Q09666_S5739 | 0.457059937 | 1.42E-06 | 1.84E-04 |
| COHA1_HUMAN\|Q9UMD9_S85_S93_T97 | 0.457059165 | 7.12E-06 | 2.30E-04 |
| NFIB_HUMAN\|O00712_S276_S295 | 0.4570131 | 5.73E-07 | 2.83E-05 |
| NUFP2_HUMAN\|Q7Z417_S212_S214 | 0.456729439 | 1.45E-07 | 5.79E-06 |
| K1C13_HUMAN\|P13646_S11 | 0.456450586 | 2.45E-04 | 1.82E-03 |

| | | | |
|---|---|---|---|
| SRRM1_HUMAN\|Q8IYB3_S738_S743 | 0.456284343 | 7.49E-05 | 1.30E-03 |
| K2C5_HUMAN\|P13647_S77_S82 | 0.455971394 | 3.82E-05 | 8.06E-04 |
| IWS1_HUMAN\|Q96ST2_S398 | 0.455792601 | 5.13E-05 | 1.48E-03 |
| K1C14_HUMAN\|P02533_S17 | 0.455587233 | 8.33E-06 | 1.29E-04 |
| K1C14_HUMAN\|P02533_Y46 | 0.455463316 | 1.93E-07 | 6.94E-06 |
| GRAP1_HUMAN\|Q4V328_S664 | 0.455160071 | 1.75E-07 | 6.42E-06 |
| FAT1_HUMAN\|Q14517_S4473 | 0.455154101 | 7.59E-06 | 1.20E-04 |
| K2C5_HUMAN\|P13647_S572 | 0.455121076 | 1.07E-07 | 4.53E-06 |
| LARP1_HUMAN\|Q6PKG0_S766_S774 | 0.455043903 | 4.87E-06 | 3.94E-04 |
| MAP7_HUMAN\|Q14244_S315 | 0.45476045 | 1.22E-08 | 2.14E-06 |
| DFFA_HUMAN\|O00273-2_S257 | 0.454753955 | 1.51E-04 | 1.24E-03 |
| TPR_HUMAN\|P12270_S1847_T1839 | 0.454583477 | 1.76E-05 | 2.29E-04 |
| ARH40_HUMAN\|Q8TER5_S262 | 0.454509992 | 6.29E-09 | 1.27E-06 |
| TOP2B_HUMAN\|Q02880_S1550_S1552 | 0.454355398 | 7.62E-06 | 1.87E-04 |
| LARP1_HUMAN\|Q6PKG0_S766_S774 | 0.454224995 | 4.73E-06 | 1.86E-04 |
| CR025_HUMAN\|Q96B23_S67 | 0.454181499 | 2.05E-06 | 6.96E-05 |
| BCR_HUMAN\|P11274_T385 | 0.454043331 | 3.58E-03 | 2.43E-02 |
| CRBG1_HUMAN\|Q9Y4K1_S141 | 0.453977843 | 4.29E-07 | 2.52E-05 |
| MB12A_HUMAN\|Q96EY5_S188 | 0.453793277 | 2.60E-06 | 5.21E-05 |
| PDLI2_HUMAN\|Q96JY6_S137_S143 | 0.453716 | 5.00E-08 | 2.56E-06 |
| SVIL_HUMAN\|O95425_T852 | 0.453616628 | 4.67E-05 | 4.96E-04 |
| DESP_HUMAN\|P15924_T13 | 0.453484419 | 6.11E-04 | 3.70E-03 |
| RTKN_HUMAN\|Q9BST9_S529_T518 | 0.453247973 | 2.66E-10 | 3.61E-07 |
| COHA1_HUMAN\|Q9UMD9_S172_T176_T182 | 0.453245193 | 1.47E-04 | 2.18E-03 |
| ARAF_HUMAN\|P10398_S257_T253 | 0.45322252 | 2.74E-07 | 1.98E-05 |
| NDRG1_HUMAN\|Q92597_S333_T328 | 0.452833245 | 1.51E-04 | 1.24E-03 |
| CE170_HUMAN\|Q5SW79_S630_S636 | 0.452811454 | 3.05E-09 | 3.07E-07 |
| AKTS1_HUMAN\|Q96B36_T246 | 0.452436795 | 1.24E-06 | 6.86E-05 |
| AHNK2_HUMAN\|Q8IVF2_S5175 | 0.452339709 | 1.67E-07 | 1.16E-05 |
| K1C19_HUMAN\|P08727_S13 | 0.451510144 | 3.05E-06 | 6.04E-05 |
| SNUT1_HUMAN\|O43290_T430 | 0.451400407 | 3.15E-05 | 6.91E-04 |

| | | | |
|---|---|---|---|
| CCDC9_HUMAN\|Q9Y3X0_S376 | 0.451235273 | 1.26E-08 | 6.81E-06 |
| NAV1_HUMAN\|Q8NEY1_T1006 | 0.451070971 | 3.43E-06 | 2.62E-04 |
| REPS1_HUMAN\|Q96D71_S162_S166_T173 | 0.450984951 | 2.12E-09 | 2.55E-07 |
| DP13A_HUMAN\|Q9UKG1_S401 | 0.450942024 | 4.07E-07 | 2.48E-05 |
| AHNK_HUMAN\|Q09666_S5752_S5763 | 0.45091103 | 1.68E-06 | 1.51E-04 |
| CND1_HUMAN\|Q15021_S1330_S1333 | 0.450395251 | 4.91E-05 | 5.25E-04 |
| FKBP5_HUMAN\|Q13451_T443 | 0.450380499 | 1.60E-05 | 2.12E-04 |
| LMNA_HUMAN\|P02545_S22_T19 | 0.450237759 | 1.51E-04 | 2.27E-03 |
| RL1D1_HUMAN\|O76021_T465 | 0.450149959 | 5.52E-06 | 3.84E-04 |
| PDS5A_HUMAN\|Q29RF7_S1276 | 0.450131315 | 2.51E-07 | 1.62E-05 |
| DESP_HUMAN\|P15924_S176 | 0.449891158 | 5.66E-05 | 1.05E-03 |
| SOS2_HUMAN\|Q07890_S1267 | 0.449672328 | 1.47E-06 | 3.51E-05 |
| SUN2_HUMAN\|Q9UH99_S112_S113_T107 | 0.449640109 | 1.97E-03 | 1.55E-02 |
| NBEL1_HUMAN\|Q6ZS30_S733 | 0.449604599 | 5.74E-03 | 3.40E-02 |
| DPYL2_HUMAN\|Q16555_S570 | 0.449447297 | 1.83E-05 | 2.35E-04 |
| NDRG1_HUMAN\|Q92597_S333_T328 | 0.449274172 | 1.36E-04 | 1.22E-03 |
| HDGF_HUMAN\|P51858_S103 | 0.448986023 | 1.43E-04 | 1.20E-03 |
| NUFP2_HUMAN\|Q7Z417_S212_S214 | 0.448827972 | 1.00E-07 | 4.36E-06 |
| MAP7_HUMAN\|Q14244_S183 | 0.448643599 | 1.16E-09 | 1.59E-07 |
| ENAH_HUMAN\|Q8N8S7_T489 | 0.448399197 | 2.87E-04 | 2.17E-03 |
| MISP_HUMAN\|Q8IVT2_S400 | 0.448153579 | 1.94E-06 | 4.34E-05 |
| K2C5_HUMAN\|P13647_S36 | 0.448028764 | 8.87E-08 | 3.94E-06 |
| SPTB2_HUMAN\|Q01082_S2341 | 0.447901761 | 4.31E-11 | 7.78E-08 |
| IRS2_HUMAN\|Q9Y4H2_T350 | 0.44748929 | 3.17E-09 | 1.45E-06 |
| PKP3_HUMAN\|Q9Y446_S280 | 0.446910006 | 5.29E-06 | 9.25E-05 |
| GTR1_HUMAN\|P11166_S490 | 0.44685697 | 5.68E-04 | 3.49E-03 |
| 4EBP1_HUMAN\|Q13541_S112 | 0.446778628 | 2.57E-04 | 1.89E-03 |
| GAB1_HUMAN\|Q13480_S683 | 0.44643052 | 2.26E-09 | 6.12E-06 |
| CF132_HUMAN\|Q5T0Z8_T297 | 0.446279065 | 3.94E-06 | 7.11E-05 |
| TGS1_HUMAN\|Q96RS0_S85_S89 | 0.446214812 | 5.67E-06 | 9.75E-05 |
| RAI14_HUMAN\|Q9P0K7-4_S261 | 0.446058805 | 2.88E-08 | 2.23E-05 |

| | | | |
|---|---|---|---|
| AHNK_HUMAN\|Q09666_S2397 | 0.446050779 | 6.48E-07 | 3.06E-05 |
| MKL2_HUMAN\|Q9ULH7_S921 | 0.445995933 | 2.88E-08 | 3.39E-06 |
| LAD1_HUMAN\|O00515_S512 | 0.445846323 | 8.01E-09 | 4.29E-06 |
| K2C8_HUMAN\|P05787_S24 | 0.445733233 | 4.67E-07 | 1.48E-05 |
| PKP1_HUMAN\|Q13835_S142 | 0.445582582 | 3.09E-05 | 3.57E-04 |
| PSN2_HUMAN\|P49810_S7_T3 | 0.445495208 | 9.62E-04 | 2.82E-02 |
| CRBG1_HUMAN\|Q9Y4K1_S22 | 0.445174639 | 1.27E-09 | 9.85E-07 |
| K2C6A_HUMAN\|P02538_Y62 | 0.445164324 | 3.15E-04 | 2.22E-03 |
| 4EBP1_HUMAN\|Q13541_S112 | 0.445143624 | 2.45E-04 | 1.93E-03 |
| MISP_HUMAN\|Q8IVT2_S395 | 0.444656269 | 2.85E-06 | 2.37E-04 |
| PHRF1_HUMAN\|Q9P1Y6_S915_T917 | 0.444589532 | 5.16E-05 | 5.34E-04 |
| B4DKV4_HUMAN\|B4DKV4_S22 | 0.444546682 | 5.78E-05 | 5.76E-04 |
| MAP7_HUMAN\|Q14244_S202 | 0.444540799 | 8.41E-10 | 1.14E-07 |
| LMNA_HUMAN\|P02545_S619 | 0.44420619 | 4.39E-07 | 1.31E-05 |
| TBC15_HUMAN\|Q8TC07_S70 | 0.444204822 | 5.29E-05 | 5.39E-04 |
| K1C17_HUMAN\|Q04695_S13 | 0.444079585 | 2.36E-08 | 1.45E-06 |
| K2C5_HUMAN\|P13647_S21 | 0.444005198 | 4.74E-04 | 3.03E-03 |
| KMT2A_HUMAN\|Q03164_T3028 | 0.443924638 | 5.08E-05 | 2.09E-03 |
| MISP_HUMAN\|Q8IVT2_S395_S400 | 0.443920814 | 1.20E-05 | 3.51E-04 |
| LAD1_HUMAN\|O00515_S301 | 0.443872045 | 3.53E-07 | 1.13E-04 |
| KC1D_HUMAN\|P48730_S331 | 0.443619916 | 7.04E-07 | 1.99E-05 |
| ABLM1_HUMAN\|O14639_S458 | 0.443405805 | 3.00E-06 | 2.42E-04 |
| NUCKS_HUMAN\|Q9H1E3_S181_T179 | 0.443059113 | 1.45E-05 | 1.73E-03 |
| CND1_HUMAN\|Q15021_S1330_S1333 | 0.442968933 | 3.83E-05 | 1.84E-03 |
| IASPP_HUMAN\|Q8WUF5_S134 | 0.442908191 | 4.21E-07 | 3.12E-05 |
| K2C6A_HUMAN\|P02538_S58 | 0.442507537 | 2.33E-04 | 1.77E-03 |
| K2C5_HUMAN\|P13647_S573 | 0.442460472 | 1.46E-04 | 1.21E-03 |
| SEPT9_HUMAN\|Q9UHD8_T38_T42 | 0.442229441 | 3.56E-09 | 3.45E-07 |
| ACINU_HUMAN\|Q9UKV3_S240 | 0.442078786 | 9.91E-08 | 7.54E-06 |
| NDRG1_HUMAN\|Q92597_S326_S330_S333 | 0.441911464 | 7.48E-08 | 9.01E-06 |
| GRAP1_HUMAN\|Q4V328_S666_S669 | 0.441670247 | 2.01E-04 | 2.85E-03 |

| | | | |
|---|---|---|---|
| LMNB1_HUMAN\|P20700_S23_T20 | 0.441291641 | 2.15E-04 | 4.92E-03 |
| LAP2A_HUMAN\|P42166_S156_T160 | 0.441284044 | 4.57E-06 | 8.25E-05 |
| ARM10_HUMAN\|Q8N2F6-2_S45 | 0.440953236 | 4.94E-03 | 4.02E-02 |
| TB182_HUMAN\|Q9C0C2_T1282 | 0.440861739 | 8.92E-09 | 3.45E-06 |
| NIPA4_HUMAN\|Q0D2K0_S427 | 0.440785631 | 4.73E-04 | 6.81E-03 |
| NDRG1_HUMAN\|Q92597_S364 | 0.440754997 | 3.99E-09 | 3.73E-07 |
| SNTB2_HUMAN\|Q13425_S222 | 0.440380306 | 2.84E-05 | 5.02E-04 |
| NDRG1_HUMAN\|Q92597_S330_S336_T328 | 0.440098323 | 2.12E-04 | 1.63E-03 |
| PAXI_HUMAN\|P49023_S106 | 0.439899027 | 4.65E-08 | 4.67E-06 |
| RYBP_HUMAN\|Q8N488_T186 | 0.43982059 | 2.67E-03 | 2.85E-02 |
| SARG_HUMAN\|Q9BW04_S131 | 0.439717646 | 1.31E-05 | 1.83E-04 |
| K1C14_HUMAN\|P02533_S51_S54 | 0.439693426 | 4.50E-07 | 1.33E-05 |
| RL29_HUMAN\|P47914_S142 | 0.439631852 | 1.38E-02 | 4.47E-02 |
| ARP19_HUMAN\|P56211_S62 | 0.439625764 | 5.26E-09 | 4.53E-07 |
| TV23B_HUMAN\|Q9NYZ1_T199 | 0.439484033 | 1.30E-04 | 1.63E-03 |
| PKN2_HUMAN\|Q16513_S360 | 0.439212447 | 3.48E-07 | 5.10E-05 |
| SEPT9_HUMAN\|Q9UHD8_T42 | 0.439130527 | 5.39E-09 | 1.17E-06 |
| MAP7_HUMAN\|Q14244_T184 | 0.438838503 | 5.09E-09 | 4.53E-07 |
| AFAD_HUMAN\|P55196_S1172_S1173_S1182 | 0.438364237 | 1.73E-09 | 2.23E-07 |
| JUPI2_HUMAN\|Q9H910_S97 | 0.438349656 | 1.68E-07 | 3.20E-05 |
| K1C17_HUMAN\|Q04695_S11_S13 | 0.438282739 | 7.67E-08 | 3.58E-06 |
| NP1L1_HUMAN\|P55209_T62 | 0.43790029 | 6.42E-06 | 2.26E-04 |
| K1C19_HUMAN\|P08727_S22 | 0.437823158 | 8.18E-08 | 3.67E-06 |
| SARG_HUMAN\|Q9BW04_S502 | 0.437816573 | 5.16E-10 | 4.66E-07 |
| PAXI_HUMAN\|P49023_S250 | 0.437507367 | 9.05E-06 | 1.35E-04 |
| SSFA2_HUMAN\|P28290_S92 | 0.437427668 | 7.96E-05 | 7.51E-04 |
| NDRG1_HUMAN\|Q92597_S333_S336 | 0.43712771 | 5.92E-08 | 2.94E-06 |
| K1C14_HUMAN\|P02533_S39 | 0.437073636 | 3.64E-05 | 4.14E-04 |
| TPR_HUMAN\|P12270_S1847_T1839 | 0.437063527 | 8.99E-06 | 1.34E-04 |
| PKP3_HUMAN\|Q9Y446_S314 | 0.436925532 | 3.55E-09 | 3.37E-07 |
| COHA1_HUMAN\|Q9UMD9_S61_S62_S70 | 0.43685502 | 4.22E-05 | 4.53E-04 |

| | | | |
|---|---|---|---|
| DP13A_HUMAN|Q9UKG1_S401 | 0.436485644 | 2.20E-07 | 2.06E-05 |
| MK01_HUMAN|P28482_T185_Y187 | 0.436119955 | 6.75E-07 | 8.13E-05 |
| RBM15_HUMAN|Q96T37_S656 | 0.435915908 | 2.57E-04 | 3.40E-02 |
| TP53B_HUMAN|Q12888_S366_T357 | 0.435900343 | 7.51E-08 | 1.51E-05 |
| K2C75_HUMAN|O95678_S40 | 0.435438736 | 2.15E-05 | 2.65E-04 |
| ATAT_HUMAN|Q5SQI0_S315 | 0.435303695 | 4.35E-11 | 3.37E-08 |
| CLAP1_HUMAN|Q7Z460_S1091 | 0.435206709 | 1.28E-09 | 3.48E-07 |
| AHNK_HUMAN|Q09666_S5745_S5749 | 0.43492212 | 2.33E-10 | 4.07E-08 |
| LAP2A_HUMAN|P42166_T74 | 0.434886688 | 1.18E-06 | 2.82E-05 |
| AHNK_HUMAN|Q09666_S5746_S5752 | 0.434579206 | 9.96E-10 | 1.26E-07 |
| FGD6_HUMAN|Q6ZV73_S721 | 0.43407086 | 4.26E-07 | 1.22E-04 |
| PAWR_HUMAN|Q96IZ0_S231 | 0.433826951 | 2.88E-09 | 2.94E-07 |
| AHNK2_HUMAN|Q8IVF2_S1710 | 0.433710454 | 8.56E-08 | 6.82E-06 |
| RENT1_HUMAN|Q92900_S1127 | 0.433351846 | 3.04E-04 | 2.16E-03 |
| K1C18_HUMAN|P05783_S7 | 0.433163404 | 1.33E-06 | 3.07E-05 |
| MILK1_HUMAN|Q8N3F8_S640_S644 | 0.433055952 | 3.05E-08 | 3.75E-06 |
| MINK1_HUMAN|Q8N4C8_S701 | 0.432860419 | 1.36E-05 | 1.89E-04 |
| B4DKV4_HUMAN|B4DKV4_S19_S22 | 0.432723176 | 2.63E-05 | 6.06E-04 |
| UBXN7_HUMAN|O94888_S285 | 0.432201618 | 3.07E-06 | 5.92E-05 |
| PSF2_HUMAN|Q9Y248_S182_T180 | 0.432150607 | 4.65E-07 | 1.35E-05 |
| K2C5_HUMAN|P13647_S21_T19 | 0.431672101 | 6.45E-06 | 2.16E-04 |
| LMNB2_HUMAN|Q03252_S424 | 0.431645588 | 6.96E-08 | 3.22E-06 |
| K2C75_HUMAN|O95678_S40 | 0.431346548 | 1.87E-05 | 2.41E-04 |
| S20A2_HUMAN|Q08357_Y423 | 0.430961485 | 3.40E-06 | 6.44E-05 |
| 2A5E_HUMAN|Q16537_S32_S34 | 0.430868634 | 3.35E-06 | 6.40E-05 |
| HS90A_HUMAN|P07900_S726 | 0.4305129 | 3.59E-04 | 2.46E-03 |
| WDR13_HUMAN|Q9H1Z4_S70_S74 | 0.430447918 | 8.48E-09 | 6.57E-07 |
| NGAP_HUMAN|Q9UJF2_T620 | 0.430414598 | 4.12E-05 | 8.58E-04 |
| K1522_HUMAN|Q9P206_S612 | 0.430361045 | 3.40E-07 | 5.12E-05 |
| CDC20_HUMAN|Q12834_S41 | 0.430146407 | 2.27E-05 | 1.08E-03 |
| PKP1_HUMAN|Q13835_S142 | 0.430141165 | 1.69E-05 | 2.22E-04 |

| | | | |
|---|---|---|---|
| LMNB1_HUMAN\|P20700_S23_T20 | 0.430140958 | 1.55E-04 | 2.94E-03 |
| TARA_HUMAN\|Q9H2D6_S1955 | 0.429873503 | 7.60E-06 | 2.41E-04 |
| PEA15_HUMAN\|Q15121_S116 | 0.429871104 | 6.88E-07 | 8.67E-05 |
| GPBP1_HUMAN\|Q86WP2_S381 | 0.429733041 | 3.82E-08 | 6.06E-06 |
| AKTS1_HUMAN\|Q96B36_S187 | 0.429563592 | 1.80E-07 | 1.75E-05 |
| SEPT9_HUMAN\|Q9UHD8_T42 | 0.429224785 | 3.21E-09 | 1.45E-06 |
| PHLB2_HUMAN\|Q86SQ0_T898 | 0.429202275 | 2.27E-05 | 2.81E-04 |
| ODPA_HUMAN\|P08559_S232 | 0.429166248 | 1.68E-03 | 9.26E-03 |
| CBPC1_HUMAN\|Q9UPW5_T1170 | 0.428931624 | 2.45E-04 | 4.28E-03 |
| RB3GP_HUMAN\|Q15042_S537 | 0.428875054 | 9.55E-10 | 2.07E-07 |
| NUCKS_HUMAN\|Q9H1E3_S181_T179 | 0.428781858 | 8.42E-06 | 5.19E-04 |
| K2C5_HUMAN\|P13647_S16_S26 | 0.428632465 | 1.38E-05 | 1.89E-04 |
| LAD1_HUMAN\|O00515_S301 | 0.428526847 | 1.71E-07 | 3.20E-05 |
| FGD6_HUMAN\|Q6ZV73_T693 | 0.428491401 | 5.01E-08 | 4.85E-06 |
| EGFR_HUMAN\|P00533_S1039 | 0.428479159 | 7.43E-04 | 2.37E-02 |
| OPTN_HUMAN\|Q96CV9_T204 | 0.428398893 | 8.61E-11 | 4.67E-07 |
| ELMO3_HUMAN\|Q96BJ8-3_S5 | 0.428304428 | 9.27E-03 | 4.86E-02 |
| IWS1_HUMAN\|Q96ST2_T725 | 0.428261529 | 8.95E-07 | 2.37E-05 |
| RTKN_HUMAN\|Q9BST9_S529_T518 | 0.427904395 | 6.32E-11 | 8.57E-08 |
| GTR1_HUMAN\|P11166_S490 | 0.427847776 | 3.26E-04 | 2.41E-03 |
| PDLI2_HUMAN\|Q96JY6_S137_S143 | 0.427814585 | 1.41E-08 | 1.07E-06 |
| PKP1_HUMAN\|Q13835_S185 | 0.427441749 | 2.37E-05 | 2.87E-04 |
| SARG_HUMAN\|Q9BW04_S502 | 0.427230987 | 2.79E-10 | 4.73E-08 |
| EDC4_HUMAN\|Q6P2E9_S723 | 0.427059149 | 2.34E-04 | 3.20E-03 |
| VIME_HUMAN\|P08670_S34 | 0.426959509 | 1.97E-08 | 1.24E-06 |
| K2C7_HUMAN\|P08729_S38 | 0.426457386 | 1.88E-05 | 2.43E-04 |
| ATAT_HUMAN\|Q5SQI0_S315 | 0.42617965 | 2.37E-11 | 7.74E-09 |
| CE170_HUMAN\|Q5SW79_S630_S633 | 0.426060602 | 8.82E-10 | 1.17E-07 |
| UBXN7_HUMAN\|O94888_S285_S288 | 0.426031234 | 1.10E-06 | 2.76E-05 |
| LMNA_HUMAN\|P02545_S616_S619 | 0.425957788 | 9.78E-04 | 5.41E-03 |
| K2C6A_HUMAN\|P02538_T548 | 0.425899162 | 1.57E-04 | 1.28E-03 |

| | | | |
|---|---|---|---|
| AHNK_HUMAN\|Q09666_S5739 | 0.425692392 | 3.69E-07 | 2.82E-05 |
| AHNK_HUMAN\|Q09666_S5749_S5763 | 0.425675982 | 1.57E-11 | 3.61E-08 |
| RRBP1_HUMAN\|Q9P2E9_T225 | 0.425557892 | 1.42E-06 | 1.32E-04 |
| NDRG1_HUMAN\|Q92597_S330_S336_T328 | 0.425488497 | 1.37E-04 | 2.13E-03 |
| ABLM1_HUMAN\|O14639_S458 | 0.425460904 | 1.38E-06 | 3.33E-05 |
| PHLB2_HUMAN\|Q86SQ0_S330_S334 | 0.425430489 | 2.85E-06 | 5.57E-05 |
| RAI14_HUMAN\|Q9P0K7_S296 | 0.425210838 | 1.83E-07 | 6.67E-06 |
| MK03_HUMAN\|P27361_T198_T202 | 0.425031592 | 9.45E-08 | 4.13E-06 |
| NDRG2_HUMAN\|Q9UN36_S332_T330 | 0.424822216 | 1.81E-09 | 2.01E-07 |
| ACINU_HUMAN\|Q9UKV3_T254 | 0.424796489 | 1.61E-06 | 3.74E-05 |
| TRI22_HUMAN\|Q8IYM9_S384 | 0.424697895 | 5.77E-08 | 5.22E-06 |
| LARP1_HUMAN\|Q6PKG0_S766_S774 | 0.424685455 | 1.54E-06 | 1.85E-04 |
| B4DKV4_HUMAN\|B4DKV4_S19_S22 | 0.424618655 | 1.91E-05 | 2.42E-04 |
| EFNB1_HUMAN\|P98172_T284 | 0.42414369 | 6.13E-06 | 1.02E-04 |
| JUPI2_HUMAN\|Q9H910_S97 | 0.424043193 | 8.59E-08 | 1.01E-05 |
| AKTS1_HUMAN\|Q96B36_T246 | 0.423875308 | 3.53E-07 | 2.26E-05 |
| PP14A_HUMAN\|Q96A00_S26 | 0.423534407 | 1.78E-10 | 6.90E-08 |
| CAP1_HUMAN\|Q01518_S34 | 0.423406359 | 1.60E-04 | 3.37E-03 |
| MTCL1_HUMAN\|Q9Y4B5_S1679_T1675 | 0.423394228 | 7.60E-10 | 2.29E-07 |
| DESP_HUMAN\|P15924_T13 | 0.422919303 | 2.66E-04 | 3.51E-03 |
| AKTS1_HUMAN\|Q96B36_T246 | 0.422834673 | 3.37E-07 | 5.07E-05 |
| K1C14_HUMAN\|P02533_S39 | 0.422745679 | 2.18E-05 | 2.68E-04 |
| SPTB2_HUMAN\|Q01082-3_S14 | 0.422511788 | 8.76E-08 | 6.89E-06 |
| CE170_HUMAN\|Q5SW79_S630_S633 | 0.422334704 | 7.14E-10 | 1.17E-07 |
| SRPK2_HUMAN\|P78362_S494_T498 | 0.422201415 | 3.95E-06 | 1.14E-04 |
| WAC2A_HUMAN\|Q641Q2_S284 | 0.422077654 | 4.11E-07 | 2.21E-05 |
| NIBL1_HUMAN\|Q96TA1_S691_S696 | 0.421972827 | 1.40E-08 | 1.95E-06 |
| ITA2_HUMAN\|P17301_S1180 | 0.421891231 | 1.06E-07 | 1.12E-04 |
| KC1A_HUMAN\|P48729-3_T323 | 0.421868416 | 4.66E-03 | 1.89E-02 |
| TBC9B_HUMAN\|Q66K14_S411 | 0.421746418 | 6.14E-07 | 1.69E-05 |
| K1C17_HUMAN\|Q04695_S11_S12 | 0.421553219 | 7.75E-08 | 3.58E-06 |

| | | | |
|---|---|---|---|
| GYS1_HUMAN\|P13807_S710 | 0.421461754 | 4.81E-04 | 3.07E-03 |
| ANK3_HUMAN\|Q12955_S959 | 0.421109839 | 5.87E-06 | 1.00E-04 |
| RAI14_HUMAN\|Q9P0K7-4_S261 | 0.42101633 | 7.00E-09 | 4.21E-06 |
| PATL1_HUMAN\|Q86TB9_S179_S184 | 0.420978958 | 6.45E-07 | 1.84E-05 |
| PSN2_HUMAN\|P49810_S7_T3 | 0.420733594 | 4.75E-04 | 5.38E-03 |
| EIF3A_HUMAN\|Q14152_S882 | 0.420542043 | 3.27E-06 | 6.33E-05 |
| MA7D1_HUMAN\|Q3KQU3_S254 | 0.420262965 | 1.64E-10 | 6.90E-08 |
| NDRG1_HUMAN\|Q92597_T366 | 0.420226854 | 3.96E-08 | 2.30E-06 |
| GGA2_HUMAN\|Q9UJY4_S177 | 0.420024991 | 7.30E-08 | 1.51E-05 |
| K2C5_HUMAN\|P13647_S36 | 0.419368015 | 1.92E-08 | 1.38E-06 |
| COHA1_HUMAN\|Q9UMD9_T182 | 0.418999433 | 3.55E-07 | 2.26E-05 |
| ITB4_HUMAN\|P16144_S1454_T1455 | 0.418714242 | 3.15E-06 | 6.19E-05 |
| PP14A_HUMAN\|Q96A00_S26 | 0.418056945 | 1.27E-10 | 1.15E-07 |
| RBP56_HUMAN\|Q92804_S226 | 0.41797894 | 3.22E-05 | 1.63E-03 |
| CRBG1_HUMAN\|Q9Y4K1_S22 | 0.417181622 | 2.92E-10 | 2.38E-07 |
| TCOF_HUMAN\|Q13428_T249 | 0.416500236 | 2.06E-08 | 6.19E-06 |
| CBPC1_HUMAN\|Q9UPW5_T1170 | 0.416226799 | 1.67E-04 | 1.99E-03 |
| WDR7_HUMAN\|Q9Y4E6_S1153 | 0.415908612 | 6.25E-06 | 2.10E-04 |
| ELMO3_HUMAN\|Q96BJ8-3_S5 | 0.415436347 | 7.40E-03 | 3.58E-02 |
| PRC2A_HUMAN\|P48634_S1147 | 0.415303052 | 2.18E-10 | 4.07E-08 |
| SEPT9_HUMAN\|Q9UHD8_T42 | 0.415284706 | 1.53E-09 | 1.74E-07 |
| MA7D1_HUMAN\|Q3KQU3_S254 | 0.415218348 | 1.20E-10 | 2.60E-08 |
| AHNK_HUMAN\|Q09666_T4430 | 0.41500526 | 3.10E-08 | 1.73E-06 |
| MAP4_HUMAN\|P27816_S1073 | 0.414999265 | 2.23E-09 | 2.63E-07 |
| NGAP_HUMAN\|Q9UJF2_T620 | 0.414917811 | 2.48E-05 | 1.15E-03 |
| PKP3_HUMAN\|Q9Y446_S180 | 0.414809389 | 5.75E-08 | 2.81E-06 |
| ENAH_HUMAN\|Q8N8S7_T489 | 0.414727516 | 1.07E-04 | 1.79E-03 |
| PRC2A_HUMAN\|P48634_S761_S764 | 0.414679342 | 6.37E-11 | 1.73E-08 |
| UBXN7_HUMAN\|O94888_S288 | 0.414254148 | 9.91E-05 | 1.57E-03 |
| GRAP1_HUMAN\|Q4V328_S666 | 0.41407276 | 9.18E-08 | 4.05E-06 |
| K2C8_HUMAN\|P05787_S21_S24_S27 | 0.414057299 | 1.43E-08 | 9.55E-07 |

| | | | |
|---|---|---|---|
| K2C7_HUMAN\|P08729_S38 | 0.413837003 | 1.18E-05 | 1.69E-04 |
| STIM2_HUMAN\|Q9P246_S522 | 0.413517535 | 3.63E-06 | 6.84E-05 |
| RTKN_HUMAN\|Q9BST9_S520_S529 | 0.413501581 | 5.13E-08 | 7.31E-06 |
| PHRF1_HUMAN\|Q9P1Y6_S915_T917 | 0.412773997 | 1.82E-05 | 2.37E-04 |
| FGD6_HUMAN\|Q6ZV73_T693 | 0.412605332 | 2.42E-08 | 2.19E-05 |
| CAP1_HUMAN\|Q01518_S34 | 0.412435637 | 1.16E-04 | 6.97E-03 |
| AKTS1_HUMAN\|Q96B36_S187 | 0.412357018 | 7.99E-08 | 1.55E-05 |
| K2C5_HUMAN\|P13647_T581 | 0.412296619 | 4.17E-04 | 2.74E-03 |
| CCDC9_HUMAN\|Q9Y3X0_S376_S386_T381 | 0.41221457 | 3.81E-09 | 2.29E-06 |
| K2C8_HUMAN\|P05787_Y25 | 0.412024999 | 2.96E-04 | 2.22E-03 |
| CLAP1_HUMAN\|Q7Z460_S1091 | 0.411964754 | 3.36E-10 | 5.35E-08 |
| MISP_HUMAN\|Q8IVT2_S395 | 0.411222899 | 7.77E-07 | 9.57E-05 |
| NIBL1_HUMAN\|Q96TA1_S692_S696 | 0.411195211 | 2.67E-08 | 3.36E-06 |
| SPD2A_HUMAN\|Q5TCZ1_S644 | 0.410943889 | 3.41E-07 | 2.64E-05 |
| WDR13_HUMAN\|Q9H1Z4_S70_S74 | 0.410590962 | 2.86E-09 | 6.74E-07 |
| C2CD2_HUMAN\|Q9Y426_S473 | 0.410284425 | 2.18E-03 | 1.15E-02 |
| NP1L1_HUMAN\|P55209_T62 | 0.410018063 | 2.23E-06 | 4.59E-05 |
| ABLM1_HUMAN\|O14639_S458 | 0.409986035 | 6.89E-07 | 3.81E-05 |
| PAXI_HUMAN\|P49023_S103_S106 | 0.409605557 | 6.05E-07 | 2.95E-05 |
| K2C5_HUMAN\|P13647_S18 | 0.409476876 | 3.33E-09 | 3.22E-07 |
| K2C5_HUMAN\|P13647_T581 | 0.409273511 | 3.84E-04 | 2.75E-03 |
| K1C19_HUMAN\|P08727_S13 | 0.409064363 | 4.39E-07 | 1.31E-05 |
| PHLB1_HUMAN\|Q86UU1_S430 | 0.408763365 | 2.04E-08 | 6.49E-06 |
| SPD2A_HUMAN\|Q5TCZ1_S1043 | 0.408531014 | 2.22E-07 | 3.64E-05 |
| AHNK_HUMAN\|Q09666_S5749_S5763 | 0.408105905 | 4.91E-12 | 1.33E-08 |
| C2CD2_HUMAN\|Q9Y426_S473 | 0.408032319 | 2.07E-03 | 9.91E-03 |
| LMBD1_HUMAN\|Q9NUN5_S528 | 0.407959026 | 1.23E-03 | 1.73E-02 |
| TRI22_HUMAN\|Q8IYM9_S384 | 0.407792544 | 2.42E-08 | 8.12E-06 |
| MTCL1_HUMAN\|Q9Y4B5_S196 | 0.407787201 | 6.20E-08 | 8.20E-06 |
| 3BP1_HUMAN\|Q9Y3L3_T601 | 0.40776082 | 3.94E-06 | 1.50E-04 |
| MA7D1_HUMAN\|Q3KQU3_S254 | 0.407445821 | 7.36E-11 | 2.22E-08 |

| | | | |
|---|---|---|---|
| PRC2A_HUMAN\|P48634_S1325_S1328_T1323 | 0.406975 | 5.84E-05 | 5.80E-04 |
| MICA3_HUMAN\|Q7RTP6_S977 | 0.406940044 | 7.51E-13 | 5.09E-10 |
| COBL1_HUMAN\|Q53SF7_T298 | 0.406624435 | 3.57E-08 | 4.09E-06 |
| PHLB1_HUMAN\|Q86UU1_S489 | 0.406052543 | 6.12E-07 | 1.78E-05 |
| LMNB1_HUMAN\|P20700_S23_T20 | 0.40598513 | 7.37E-05 | 5.16E-03 |
| K1522_HUMAN\|Q9P206_S929 | 0.40583547 | 4.53E-11 | 1.75E-08 |
| LAP2A_HUMAN\|P42166_S156_T160 | 0.405615734 | 1.18E-06 | 2.82E-05 |
| LAD1_HUMAN\|O00515_S38_S39 | 0.404796483 | 1.74E-06 | 1.85E-04 |
| SRS10_HUMAN\|O75494_S106 | 0.404595061 | 2.27E-07 | 2.08E-05 |
| PKP2_HUMAN\|Q99959_S132 | 0.404588908 | 1.64E-07 | 6.16E-06 |
| NDRG1_HUMAN\|Q92597_S330_S336_T328 | 0.404558023 | 7.21E-05 | 7.35E-04 |
| MAP7_HUMAN\|Q14244_S183 | 0.404541075 | 7.73E-11 | 4.66E-08 |
| COHA1_HUMAN\|Q9UMD9_T182 | 0.404469693 | 1.84E-07 | 6.67E-06 |
| COHA1_HUMAN\|Q9UMD9_S93_T97 | 0.404058424 | 5.45E-07 | 1.64E-05 |
| ARP19_HUMAN\|P56211_Y59 | 0.403736649 | 3.35E-10 | 5.35E-08 |
| COHA1_HUMAN\|Q9UMD9_S93_S96 | 0.403111493 | 8.05E-07 | 2.18E-05 |
| ROA1_HUMAN\|P09651_S360_Y366 | 0.40309704 | 4.87E-04 | 3.38E-03 |
| LASP1_HUMAN\|Q14847_T104 | 0.402866407 | 4.55E-10 | 2.85E-07 |
| K2C6A_HUMAN\|P02538_S22 | 0.402833099 | 6.67E-05 | 6.53E-04 |
| PDLI1_HUMAN\|O00151_S130 | 0.402630684 | 4.73E-10 | 2.85E-07 |
| DP13A_HUMAN\|Q9UKG1_S401 | 0.402448001 | 5.01E-08 | 2.66E-06 |
| K2C5_HUMAN\|P13647_S16_S26 | 0.40242222 | 5.05E-06 | 8.75E-05 |
| K2C5_HUMAN\|P13647_S77_S82 | 0.402000441 | 5.10E-06 | 8.81E-05 |
| JUPI2_HUMAN\|Q9H910_S97 | 0.401882567 | 2.96E-08 | 1.70E-06 |
| IASPP_HUMAN\|Q8WUF5_S134 | 0.401515858 | 6.93E-08 | 3.32E-06 |
| ACINU_HUMAN\|Q9UKV3_T254 | 0.40109721 | 6.26E-07 | 1.70E-05 |
| PHLB2_HUMAN\|Q86SQ0_S334_T328 | 0.400864499 | 1.01E-06 | 2.50E-05 |
| PARD3_HUMAN\|Q8TEW0_S220 | 0.400723633 | 1.37E-05 | 1.89E-04 |
| RSF1_HUMAN\|Q96T23_S1277_T1278 | 0.400719176 | 1.53E-03 | 1.70E-02 |
| K1522_HUMAN\|Q9P206_S612 | 0.4004385 | 8.52E-08 | 6.82E-06 |
| RENT1_HUMAN\|Q92900_S1107_T1111 | 0.400261317 | 4.35E-06 | 7.71E-05 |

| | | | |
|---|---|---|---|
| B4DKV4_HUMAN\|B4DKV4_S22 | 0.399805304 | 1.00E-05 | 1.46E-04 |
| FGD6_HUMAN\|Q6ZV73_S721 | 0.399677956 | 9.95E-08 | 1.93E-05 |
| K1143_HUMAN\|Q96AT1_S146 | 0.399382655 | 2.26E-07 | 8.01E-06 |
| K1143_HUMAN\|Q96AT1_S146 | 0.399133345 | 2.24E-07 | 7.74E-06 |
| PARD3_HUMAN\|Q8TEW0_S220 | 0.399039589 | 1.29E-05 | 1.79E-04 |
| SARG_HUMAN\|Q9BW04_S502 | 0.398969518 | 5.27E-11 | 1.68E-08 |
| NUCKS_HUMAN\|Q9H1E3_S181_T179 | 0.398834312 | 2.59E-06 | 2.23E-04 |
| ZO2_HUMAN\|Q9UDY2_T1027 | 0.398801308 | 1.12E-09 | 1.35E-07 |
| K1C17_HUMAN\|Q04695_S24 | 0.398708534 | 1.83E-09 | 2.31E-07 |
| ITA2_HUMAN\|P17301_S1181 | 0.398703165 | 6.93E-08 | 1.50E-05 |
| CR025_HUMAN\|Q96B23_S143_S145 | 0.39824387 | 7.65E-11 | 1.38E-07 |
| XPC_HUMAN\|Q01831_S883_S884_S891 | 0.398069021 | 1.60E-07 | 6.13E-06 |
| TB182_HUMAN\|Q9C0C2_S1652 | 0.398044649 | 2.23E-09 | 2.34E-07 |
| SEPT9_HUMAN\|Q9UHD8_T38_T42 | 0.397327008 | 3.12E-10 | 1.06E-07 |
| LAD1_HUMAN\|O00515_S512 | 0.397273776 | 4.80E-10 | 6.50E-07 |
| MTCL1_HUMAN\|Q9Y4B5_S196 | 0.396917438 | 3.84E-08 | 2.26E-06 |
| WDR13_HUMAN\|Q9H1Z4_S70_S74 | 0.396804343 | 1.33E-09 | 5.53E-07 |
| PKP3_HUMAN\|Q9Y446_S223 | 0.396792806 | 2.14E-04 | 2.95E-03 |
| CR025_HUMAN\|Q96B23_S143_S145_S147 | 0.396775139 | 2.70E-08 | 8.12E-06 |
| MK03_HUMAN\|P27361_T198_T202 | 0.39673859 | 2.39E-08 | 3.16E-06 |
| CHSP1_HUMAN\|Q9Y2V2_S52 | 0.396719702 | 3.54E-08 | 2.16E-06 |
| SYTL1_HUMAN\|Q8IYJ3_S216 | 0.396380849 | 2.19E-06 | 4.75E-05 |
| SARG_HUMAN\|Q9BW04_S429 | 0.396254558 | 2.00E-07 | 7.09E-06 |
| COF1_HUMAN\|P23528_S3 | 0.395576915 | 2.33E-04 | 1.77E-03 |
| WDR7_HUMAN\|Q9Y4E6_S1153 | 0.395384022 | 2.68E-06 | 5.52E-05 |
| MTCL1_HUMAN\|Q9Y4B5_S1679_T1675 | 0.395104164 | 1.70E-10 | 3.41E-08 |
| K2C5_HUMAN\|P13647_S21_S26 | 0.394925015 | 2.89E-08 | 1.80E-06 |
| MISP_HUMAN\|Q8IVT2_S400 | 0.394787378 | 2.27E-07 | 3.84E-05 |
| LASP1_HUMAN\|Q14847_T104 | 0.394784674 | 2.87E-10 | 5.57E-08 |
| TB182_HUMAN\|Q9C0C2_T1282 | 0.394422141 | 6.86E-10 | 2.19E-07 |
| EGFR_HUMAN\|P00533_S1039 | 0.394240231 | 2.94E-04 | 4.73E-03 |

| | | | |
|---|---|---|---|
| REQU_HUMAN\|Q92785_T149 | 0.394193003 | 6.76E-09 | 3.33E-06 |
| ROA1_HUMAN\|P09651_S360_Y366 | 0.393560656 | 3.77E-04 | 2.55E-03 |
| GYS1_HUMAN\|P13807_T703 | 0.393447054 | 7.70E-09 | 3.48E-06 |
| RBM15_HUMAN\|Q96T37_S656 | 0.392662552 | 7.47E-05 | 1.36E-03 |
| PP14A_HUMAN\|Q96A00_S26 | 0.391874467 | 2.43E-11 | 7.74E-09 |
| PDLI5_HUMAN\|Q96HC4_S111 | 0.39155151 | 3.51E-10 | 5.43E-08 |
| TARA_HUMAN\|Q9H2D6_S1955 | 0.390914036 | 1.85E-06 | 4.19E-05 |
| TRI22_HUMAN\|Q8IYM9_S383 | 0.390868275 | 1.50E-06 | 5.49E-05 |
| MICA3_HUMAN\|Q7RTP6_S1636 | 0.389776321 | 1.33E-06 | 3.07E-05 |
| AHNK_HUMAN\|Q09666_S5397 | 0.389751399 | 3.79E-05 | 4.17E-04 |
| RBM15_HUMAN\|Q96T37_S656 | 0.388961426 | 6.69E-05 | 2.54E-03 |
| PP14A_HUMAN\|Q96A00_S26 | 0.388842624 | 2.00E-11 | 3.61E-08 |
| TT39B_HUMAN\|Q5VTQ0_S120 | 0.388587296 | 1.21E-07 | 4.98E-06 |
| NOLC1_HUMAN\|Q14978_S688_S698 | 0.388435752 | 2.00E-04 | 1.56E-03 |
| 3BP1_HUMAN\|Q9Y3L3_T601 | 0.387422883 | 1.86E-06 | 4.21E-05 |
| AHNK_HUMAN\|Q09666_S5749_S5752 | 0.387037153 | 1.22E-10 | 3.01E-08 |
| RTKN_HUMAN\|Q9BST9_S529_T518 | 0.386359575 | 5.75E-12 | 1.04E-08 |
| SLK_HUMAN\|Q9H2G2_S779 | 0.386343372 | 3.76E-06 | 7.00E-05 |
| COHA1_HUMAN\|Q9UMD9_S61_S62_S70 | 0.386191268 | 7.19E-06 | 2.40E-04 |
| TB182_HUMAN\|Q9C0C2_T1282 | 0.386014086 | 4.25E-10 | 2.88E-07 |
| ARG35_HUMAN\|A5YM69_S450 | 0.385342276 | 1.16E-07 | 8.58E-06 |
| K2C5_HUMAN\|P13647_T19 | 0.385222493 | 7.62E-07 | 1.99E-05 |
| K2C6A_HUMAN\|P02538_S85 | 0.385051398 | 5.88E-05 | 6.17E-04 |
| ITB4_HUMAN\|P16144_S1454_S1457 | 0.384076769 | 7.02E-06 | 1.14E-04 |
| MK03_HUMAN\|P27361_T198_T202 | 0.384075814 | 1.27E-08 | 4.32E-06 |
| K2C5_HUMAN\|P13647_S573 | 0.383989243 | 2.14E-05 | 2.69E-04 |
| NP1L1_HUMAN\|P55209_T62 | 0.383950019 | 7.96E-07 | 2.17E-05 |
| MAP7_HUMAN\|Q14244_T184 | 0.383948438 | 2.16E-10 | 7.81E-08 |
| ITA2_HUMAN\|P17301_S1181 | 0.383544788 | 3.05E-08 | 8.46E-06 |
| K2C5_HUMAN\|P13647_S21 | 0.382985942 | 7.91E-05 | 7.83E-04 |
| RAI14_HUMAN\|Q9P0K7_T251 | 0.382793133 | 1.28E-08 | 5.28E-06 |

| | | | |
|---|---|---|---|
| ZC3HD_HUMAN|Q5T200_S1014_S1017 | 0.38268623 | 1.28E-08 | 2.16E-06 |
| K2C5_HUMAN|P13647_S82_T78 | 0.382509872 | 2.05E-10 | 4.64E-08 |
| ARP19_HUMAN|P56211_S62 | 0.382465003 | 1.67E-10 | 3.41E-08 |
| ADNP_HUMAN|Q9H2P0_S409 | 0.382326042 | 7.76E-04 | 4.50E-03 |
| CAP1_HUMAN|Q01518_S34 | 0.38220046 | 4.61E-05 | 1.63E-03 |
| NUP53_HUMAN|Q8NFH5_S105 | 0.380907098 | 2.60E-08 | 3.36E-06 |
| K2C5_HUMAN|P13647_S64 | 0.380760453 | 8.20E-08 | 3.67E-06 |
| LMNB1_HUMAN|P20700_S23_T20 | 0.38073098 | 3.26E-05 | 1.10E-03 |
| KPCD2_HUMAN|Q9BZL6_S710 | 0.380179196 | 6.48E-06 | 1.07E-04 |
| NGAP_HUMAN|Q9UJF2_T620 | 0.380098667 | 7.55E-06 | 2.41E-04 |
| TM1L1_HUMAN|O75674_S321 | 0.379775574 | 1.99E-07 | 1.33E-05 |
| PDLI2_HUMAN|Q96JY6_S137 | 0.379318497 | 1.86E-10 | 5.30E-08 |
| FA83B_HUMAN|Q5T0W9_T687 | 0.379089803 | 1.66E-04 | 1.34E-03 |
| K1C17_HUMAN|Q04695_S11_S13 | 0.378859347 | 2.66E-09 | 2.83E-07 |
| AHNK_HUMAN|Q09666_T5824 | 0.378807851 | 3.45E-05 | 3.89E-04 |
| MTAP2_HUMAN|P11137_T1602_T1605 | 0.378671287 | 8.80E-07 | 3.72E-05 |
| AHNK_HUMAN|Q09666_T5824 | 0.377900383 | 3.35E-05 | 3.87E-04 |
| PHF14_HUMAN|O94880_T287 | 0.37786924 | 5.11E-09 | 4.53E-07 |
| K1522_HUMAN|Q9P206_S929 | 0.377773404 | 7.34E-12 | 1.99E-08 |
| IASPP_HUMAN|Q8WUF5_S395 | 0.377671528 | 1.20E-07 | 4.95E-06 |
| K1C14_HUMAN|P02533_S44 | 0.377645102 | 1.94E-08 | 1.38E-06 |
| SRC8_HUMAN|Q14247_S405_S418_T401 | 0.377500742 | 7.18E-07 | 3.28E-05 |
| R3HD2_HUMAN|Q9Y2K5_T856 | 0.377428029 | 4.64E-07 | 6.44E-05 |
| IRS2_HUMAN|Q9Y4H2_S1186_S1203 | 0.377247067 | 1.12E-09 | 2.34E-07 |
| BORG4_HUMAN|Q9H3Q1_S64 | 0.376487445 | 2.29E-10 | 4.97E-08 |
| SPD2A_HUMAN|Q5TCZ1_S1043 | 0.376419742 | 5.65E-08 | 5.88E-06 |
| NDRG1_HUMAN|Q92597_S333_S336 | 0.37640033 | 2.41E-09 | 2.72E-07 |
| NAV1_HUMAN|Q8NEY1_T1006 | 0.375816967 | 1.39E-07 | 1.11E-05 |
| SPD2A_HUMAN|Q5TCZ1_S644 | 0.375550548 | 7.72E-08 | 2.66E-05 |
| MET_HUMAN|P08581_S990_S997 | 0.375282628 | 1.90E-04 | 2.21E-03 |
| CRBG1_HUMAN|Q9Y4K1_S410 | 0.375114439 | 3.71E-07 | 5.29E-05 |

| | | | |
|---|---|---|---|
| K1C14_HUMAN\|P02533_S54_Y46 | 0.374145367 | 1.09E-08 | 9.13E-07 |
| SPTB2_HUMAN\|Q01082-3_S14_S8 | 0.374114026 | 2.04E-10 | 5.54E-07 |
| K2C6A_HUMAN\|P02538_S60 | 0.373738605 | 3.07E-05 | 3.62E-04 |
| IWS1_HUMAN\|Q96ST2_T725 | 0.373676782 | 9.25E-08 | 8.35E-06 |
| SPD2A_HUMAN\|Q5TCZ1_S651 | 0.373322136 | 4.75E-08 | 2.60E-06 |
| CR025_HUMAN\|Q96B23_S66_S69 | 0.373298393 | 1.10E-06 | 2.76E-05 |
| COHA1_HUMAN\|Q9UMD9_S172_T176_T182 | 0.373297693 | 1.06E-05 | 1.53E-04 |
| AHNK_HUMAN\|Q09666_S5397 | 0.373143074 | 2.20E-05 | 2.74E-04 |
| NBEL1_HUMAN\|Q6ZS30_S733 | 0.372927294 | 1.10E-03 | 6.57E-03 |
| IRS2_HUMAN\|Q9Y4H2_S1185_S1203 | 0.372864231 | 8.48E-09 | 1.31E-06 |
| REQU_HUMAN\|Q92785_S142 | 0.372724089 | 1.95E-09 | 2.40E-07 |
| NUP53_HUMAN\|Q8NFH5_S105 | 0.372562241 | 1.78E-08 | 1.16E-06 |
| SLTM_HUMAN\|Q9NWH9_S371 | 0.371402547 | 1.38E-04 | 1.16E-03 |
| SEPT9_HUMAN\|Q9UHD8_T38_T42 | 0.371274658 | 7.28E-11 | 9.86E-08 |
| K2C5_HUMAN\|P13647_S21_S26 | 0.37117659 | 8.18E-09 | 6.43E-07 |
| NGAP_HUMAN\|Q9UJF2_S954 | 0.370616382 | 2.43E-09 | 4.71E-07 |
| SMCR8_HUMAN\|Q8TEV9_S417 | 0.37041759 | 8.04E-05 | 3.25E-03 |
| ADNP_HUMAN\|Q9H2P0_S409 | 0.370069727 | 5.73E-04 | 1.44E-02 |
| LMNA_HUMAN\|P02545_S616_S619 | 0.369616326 | 2.22E-04 | 1.81E-03 |
| MILK1_HUMAN\|Q8N3F8_T318 | 0.369473128 | 1.12E-06 | 5.45E-05 |
| R3HD2_HUMAN\|Q9Y2K5_T856 | 0.369286109 | 3.30E-07 | 4.84E-05 |
| K1C14_HUMAN\|P02533_S54_Y46 | 0.368634406 | 8.08E-09 | 6.43E-07 |
| DPYL3_HUMAN\|Q14195_S568 | 0.368505275 | 7.05E-07 | 1.86E-05 |
| COBL1_HUMAN\|Q53SF7_S294_T298 | 0.368403718 | 3.95E-09 | 1.19E-06 |
| CR025_HUMAN\|Q96B23_S66_S67_T74 | 0.368339495 | 2.60E-04 | 2.85E-03 |
| RETR3_HUMAN\|Q86VR2_S435_T440 | 0.368112955 | 8.51E-03 | 3.53E-02 |
| SEPT7_HUMAN\|Q16181_T426 | 0.366575789 | 7.23E-07 | 2.01E-05 |
| PKP3_HUMAN\|Q9Y446_S180 | 0.366048729 | 6.07E-09 | 5.57E-07 |
| ITB4_HUMAN\|P16144_S1457_T1455 | 0.365282319 | 4.22E-06 | 7.78E-05 |
| IRS2_HUMAN\|Q9Y4H2_T350 | 0.36519655 | 3.73E-11 | 2.02E-07 |
| CR025_HUMAN\|Q96B23_S143_S145_S147 | 0.364834585 | 5.15E-09 | 6.98E-06 |

| | | | |
|---|---|---|---|
| XPC_HUMAN\|Q01831_S883_S884_S891 | 0.364641594 | 3.08E-08 | 1.73E-06 |
| E7ETA6_HUMAN\|E7ETA6_T1557 | 0.364448026 | 2.63E-07 | 2.20E-05 |
| COHA1_HUMAN\|Q9UMD9_S93_T97 | 0.363984435 | 9.24E-08 | 4.07E-06 |
| NAV1_HUMAN\|Q8NEY1_T1006 | 0.363147671 | 7.82E-08 | 3.53E-06 |
| AHNK_HUMAN\|Q09666_T4430 | 0.362937563 | 2.25E-09 | 2.34E-07 |
| TB182_HUMAN\|Q9C0C2_T1282 | 0.362816365 | 1.11E-10 | 1.15E-07 |
| SOS2_HUMAN\|Q07890_S1267 | 0.362468782 | 2.16E-08 | 3.16E-06 |
| RAI14_HUMAN\|Q9P0K7_T297 | 0.362264294 | 2.99E-08 | 1.70E-06 |
| IASPP_HUMAN\|Q8WUF5_S134 | 0.362223217 | 1.16E-08 | 2.10E-06 |
| PHLB2_HUMAN\|Q86SQ0_S330_S334 | 0.361750737 | 2.34E-07 | 8.18E-06 |
| REQU_HUMAN\|Q92785_T149 | 0.361587696 | 1.22E-09 | 1.43E-07 |
| CCDC9_HUMAN\|Q9Y3X0_S376 | 0.361074853 | 8.86E-11 | 2.53E-08 |
| MAP7_HUMAN\|Q14244_S183 | 0.360780612 | 4.69E-12 | 2.31E-09 |
| K1C14_HUMAN\|P02533_Y46 | 0.360709373 | 9.86E-10 | 1.48E-07 |
| K2C5_HUMAN\|P13647_S82_T78 | 0.360652363 | 5.10E-11 | 1.46E-08 |
| ITA2_HUMAN\|P17301_S1181 | 0.360438095 | 8.35E-09 | 1.31E-06 |
| RTKN_HUMAN\|Q9BST9_S529_T518 | 0.360413093 | 1.25E-12 | 6.76E-09 |
| RBM15_HUMAN\|Q96T37_S656 | 0.360005643 | 2.75E-05 | 8.23E-04 |
| BORG4_HUMAN\|Q9H3Q1_S64 | 0.359996567 | 8.94E-11 | 2.20E-08 |
| PSN2_HUMAN\|P49810_S7_T3 | 0.359410049 | 6.46E-05 | 2.06E-03 |
| RTKN_HUMAN\|Q9BST9_S529_T518 | 0.358843729 | 1.14E-12 | 6.84E-10 |
| K2C5_HUMAN\|P13647_S77_S82 | 0.35858902 | 8.67E-07 | 2.30E-05 |
| ITA2_HUMAN\|P17301_S1180 | 0.35786947 | 2.94E-09 | 1.99E-06 |
| LARP1_HUMAN\|Q6PKG0_S766 | 0.356119965 | 7.03E-06 | 1.13E-04 |
| RHG29_HUMAN\|Q52LW3_S176 | 0.355934398 | 1.09E-07 | 2.04E-05 |
| LAD1_HUMAN\|O00515_S38_T40_T41 | 0.355931 | 4.71E-09 | 8.81E-07 |
| SARG_HUMAN\|Q9BW04_S131 | 0.355402876 | 5.39E-07 | 1.64E-05 |
| RL1D1_HUMAN\|O76021_T465 | 0.35520141 | 9.31E-08 | 2.12E-05 |
| CAVN3_HUMAN\|Q969G5_T205 | 0.354929977 | 1.01E-08 | 8.65E-07 |
| MISP_HUMAN\|Q8IVT2_S395_S400 | 0.354151352 | 4.26E-07 | 2.24E-05 |
| RENT1_HUMAN\|Q92900_S1107_T1111 | 0.353653724 | 7.62E-07 | 2.10E-05 |

| | | | |
|---|---|---|---|
| K2C5_HUMAN\|P13647_T19 | 0.35346599 | 1.83E-07 | 6.67E-06 |
| LMNA_HUMAN\|P02545_S618 | 0.352425796 | 1.65E-05 | 2.17E-04 |
| CCDC9_HUMAN\|Q9Y3X0_S376_S386_T381 | 0.351876867 | 1.43E-10 | 6.90E-08 |
| MILK1_HUMAN\|Q8N3F8_T318 | 0.350723943 | 5.43E-07 | 1.64E-05 |
| TB182_HUMAN\|Q9C0C2_S691 | 0.349747658 | 1.23E-10 | 3.71E-08 |
| MAP7_HUMAN\|Q14244_T184 | 0.349428886 | 2.69E-11 | 8.09E-09 |
| EGFR_HUMAN\|P00533_S1039 | 0.349356054 | 7.69E-05 | 2.48E-03 |
| COBL1_HUMAN\|Q53SF7_T298 | 0.349240111 | 2.45E-09 | 2.50E-07 |
| KPCD2_HUMAN\|Q9BZL6_S710 | 0.349229566 | 2.09E-06 | 2.03E-04 |
| DESP_HUMAN\|P15924_T13 | 0.348347783 | 2.73E-05 | 3.28E-04 |
| SPD2A_HUMAN\|Q5TCZ1_S1043 | 0.348126414 | 1.62E-08 | 1.21E-06 |
| RTKN_HUMAN\|Q9BST9_S520_S529 | 0.34807188 | 2.57E-09 | 1.39E-06 |
| DOCK4_HUMAN\|Q8N1I0_S1620 | 0.347488717 | 3.37E-07 | 4.16E-05 |
| RRBP1_HUMAN\|Q9P2E9_T225 | 0.347414776 | 3.41E-08 | 2.31E-05 |
| ETV3_HUMAN\|P41162_S139 | 0.34725753 | 1.15E-06 | 2.78E-05 |
| LASP1_HUMAN\|Q14847_T104 | 0.346677697 | 1.72E-11 | 2.33E-08 |
| K2C5_HUMAN\|P13647_S16_S21 | 0.346019692 | 2.42E-07 | 8.26E-06 |
| LARP1_HUMAN\|Q6PKG0_S766_S774 | 0.345680012 | 6.21E-08 | 3.03E-06 |
| IRS2_HUMAN\|Q9Y4H2_T350 | 0.345560669 | 1.24E-11 | 3.61E-08 |
| CR025_HUMAN\|Q96B23_S143_S145 | 0.345151639 | 2.35E-12 | 1.27E-08 |
| COHA1_HUMAN\|Q9UMD9_S76_S79 | 0.34489969 | 1.40E-06 | 6.58E-05 |
| CF132_HUMAN\|Q5T0Z8_T297 | 0.344673295 | 5.81E-08 | 2.94E-06 |
| PKP3_HUMAN\|Q9Y446_S180_S183 | 0.34448467 | 5.20E-09 | 4.53E-07 |
| K1C17_HUMAN\|Q04695_S11_S12 | 0.344105665 | 9.17E-10 | 1.42E-07 |
| SMCR8_HUMAN\|Q8TEV9_S417 | 0.344026268 | 3.56E-05 | 3.96E-04 |
| MISP_HUMAN\|Q8IVT2_S400 | 0.343845246 | 2.58E-08 | 7.37E-06 |
| RTKN_HUMAN\|Q9BST9_S520_S529 | 0.343838239 | 2.10E-09 | 5.18E-07 |
| ODPA_HUMAN\|P08559_S232 | 0.343570056 | 1.40E-04 | 3.75E-03 |
| PLEC_HUMAN\|Q15149_S4406 | 0.343184604 | 9.73E-06 | 2.24E-04 |
| HMGN1_HUMAN\|P05114_S7 | 0.342997659 | 6.74E-10 | 1.59E-07 |
| MISP_HUMAN\|Q8IVT2_S155 | 0.342637839 | 4.32E-08 | 2.27E-06 |

| | | | |
|---|---|---|---|
| COHA1_HUMAN\|Q9UMD9_S93_S96 | 0.341170025 | 5.24E-08 | 2.61E-06 |
| PACS1_HUMAN\|Q6VY07_T504 | 0.340739477 | 9.41E-09 | 8.23E-07 |
| B4DKV4_HUMAN\|B4DKV4_S19_S22 | 0.340648028 | 4.67E-07 | 1.48E-05 |
| RTKN_HUMAN\|Q9BST9_S529_T518 | 0.340260422 | 3.73E-13 | 1.01E-09 |
| CR025_HUMAN\|Q96B23_S69 | 0.339153635 | 9.49E-07 | 3.93E-05 |
| BCR_HUMAN\|P11274_T385 | 0.338906851 | 2.35E-04 | 1.87E-03 |
| E7ETA6_HUMAN\|E7ETA6_T1557 | 0.338824518 | 7.12E-08 | 3.27E-06 |
| RTKN_HUMAN\|Q9BST9_S529_T518 | 0.338778809 | 3.41E-13 | 3.70E-10 |
| K2C8_HUMAN\|P05787_S24 | 0.337429599 | 1.17E-09 | 1.59E-07 |
| K2C6B_HUMAN\|P04259_S37 | 0.337327336 | 1.37E-08 | 1.07E-06 |
| SRRM2_HUMAN\|Q9UQ35_S1152 | 0.337277073 | 1.47E-08 | 1.11E-06 |
| K2C5_HUMAN\|P13647_S64 | 0.337043723 | 8.89E-09 | 6.78E-07 |
| LAD1_HUMAN\|O00515_S38_T40_T41 | 0.336969254 | 1.72E-09 | 1.33E-06 |
| NDRG1_HUMAN\|Q92597_S326_S330_S333 | 0.336702217 | 2.56E-10 | 5.34E-08 |
| IRS2_HUMAN\|Q9Y4H2_T350 | 0.336523843 | 7.39E-12 | 2.67E-09 |
| TRI22_HUMAN\|Q8IYM9_S383 | 0.336125083 | 1.41E-07 | 2.54E-05 |
| AHNK2_HUMAN\|Q8IVF2_S294_T298 | 0.335352999 | 4.06E-10 | 7.10E-08 |
| TRI22_HUMAN\|Q8IYM9_S384 | 0.335189793 | 4.54E-10 | 6.50E-07 |
| FA83B_HUMAN\|Q5T0W9_T687 | 0.334656547 | 4.50E-05 | 4.86E-04 |
| DBNL_HUMAN\|Q9UJU6_T270 | 0.334554403 | 1.50E-08 | 9.93E-07 |
| PKP3_HUMAN\|Q9Y446_S180_S183 | 0.332568136 | 2.96E-09 | 2.91E-07 |
| AKTS1_HUMAN\|Q96B36_T246 | 0.332423507 | 4.44E-09 | 4.08E-07 |
| K2C75_HUMAN\|O95678_S61 | 0.332340192 | 1.06E-07 | 4.53E-06 |
| EGFR_HUMAN\|P00533_S1039 | 0.33212288 | 4.41E-05 | 6.86E-04 |
| LMNA_HUMAN\|P02545_S616_S619 | 0.332105733 | 7.38E-05 | 1.28E-03 |
| NDRG1_HUMAN\|Q92597_S333_S336_T328 | 0.330999641 | 3.69E-13 | 2.86E-10 |
| K2C6A_HUMAN\|P02538_S60 | 0.32985479 | 5.99E-06 | 1.00E-04 |
| MYPT1_HUMAN\|O14974_S554 | 0.32976019 | 9.95E-07 | 2.56E-05 |
| K2C6A_HUMAN\|P02538_S85 | 0.329017829 | 7.29E-06 | 1.17E-04 |
| K1C14_HUMAN\|P02533_S44 | 0.327129975 | 1.15E-09 | 1.59E-07 |
| RAI14_HUMAN\|Q9P0K7-4_S261 | 0.326532129 | 2.10E-11 | 9.80E-09 |

| | | | |
|---|---|---|---|
| RTKN_HUMAN\|Q9BST9_S520_S529 | 0.325085217 | 8.56E-10 | 1.36E-07 |
| COHA1_HUMAN\|Q9UMD9_T182 | 0.324609336 | 3.94E-09 | 3.73E-07 |
| LARP1_HUMAN\|Q6PKG0_S766_S774 | 0.32319946 | 2.35E-08 | 1.45E-06 |
| SRS10_HUMAN\|O75494_S106 | 0.32283932 | 2.49E-09 | 2.50E-07 |
| RRBP1_HUMAN\|Q9P2E9_T225 | 0.321090542 | 8.61E-09 | 3.45E-06 |
| FGD6_HUMAN\|Q6ZV73_T693 | 0.319356199 | 2.70E-10 | 2.93E-07 |
| PSN2_HUMAN\|P49810_S7_T3 | 0.319103914 | 1.40E-05 | 1.90E-04 |
| CE170_HUMAN\|Q5SW79_S633_S636 | 0.319091771 | 1.07E-07 | 4.51E-06 |
| SPTB2_HUMAN\|Q01082_T2328 | 0.319005387 | 1.18E-06 | 2.94E-05 |
| ITA2_HUMAN\|P17301_S1180 | 0.318700342 | 2.66E-10 | 2.93E-07 |
| IRS2_HUMAN\|Q9Y4H2_T350 | 0.318429636 | 2.60E-12 | 1.57E-09 |
| K2C75_HUMAN\|O95678_S61 | 0.318421786 | 5.75E-08 | 2.81E-06 |
| JUPI2_HUMAN\|Q9H910_S97 | 0.318402457 | 4.10E-10 | 6.01E-08 |
| RHG29_HUMAN\|Q52LW3_S176 | 0.318360175 | 1.95E-08 | 2.11E-05 |
| RL1D1_HUMAN\|O76021_T465 | 0.316853524 | 1.49E-08 | 2.02E-05 |
| RENT1_HUMAN\|Q92900_S1127 | 0.315708339 | 8.28E-06 | 1.29E-04 |
| DBNL_HUMAN\|Q9UJU6_S269 | 0.315515088 | 4.37E-09 | 4.08E-07 |
| K2C5_HUMAN\|P13647_S16_S21 | 0.315296954 | 5.79E-08 | 2.94E-06 |
| TP53B_HUMAN\|Q12888_S356 | 0.313760975 | 2.70E-07 | 8.83E-06 |
| MK01_HUMAN\|P28482_T185_Y187 | 0.313454894 | 2.77E-09 | 2.89E-07 |
| RETR3_HUMAN\|Q86VR2_S435_T440 | 0.312918768 | 2.93E-03 | 2.15E-02 |
| STMN1_HUMAN\|P16949_S25 | 0.312578178 | 7.57E-13 | 6.47E-10 |
| SARG_HUMAN\|Q9BW04_T151 | 0.312061849 | 4.11E-07 | 1.25E-05 |
| TRI22_HUMAN\|Q8IYM9_S383 | 0.311595383 | 4.51E-08 | 1.16E-05 |
| MYPT1_HUMAN\|O14974_S554 | 0.31150171 | 4.91E-07 | 3.51E-05 |
| COHA1_HUMAN\|Q9UMD9_S61_S62_S70 | 0.31120422 | 3.83E-07 | 2.41E-05 |
| IASPP_HUMAN\|Q8WUF5_T341 | 0.311105485 | 1.06E-06 | 2.58E-05 |
| KMT2A_HUMAN\|Q03164_T3028 | 0.310943841 | 2.50E-07 | 8.40E-06 |
| AHNK2_HUMAN\|Q8IVF2_S294_T298 | 0.310818265 | 1.00E-10 | 2.27E-08 |
| K2C5_HUMAN\|P13647_S26_T19 | 0.310739309 | 1.71E-05 | 2.24E-04 |
| RU17_HUMAN\|P08621_S320 | 0.309149054 | 1.85E-07 | 6.67E-06 |

| | | | |
|---|---|---|---|
| PHLB2_HUMAN|Q86SQ0_S334_T328 | 0.309098861 | 2.25E-08 | 1.52E-06 |
| RRBP1_HUMAN|Q9P2E9_T225 | 0.308575841 | 4.37E-09 | 1.82E-06 |
| NDRG1_HUMAN|Q92597_S333_S336_T328 | 0.306572701 | 6.51E-14 | 1.77E-10 |
| SRRM1_HUMAN|Q8IYB3_S738_S743 | 0.305672664 | 2.48E-07 | 8.40E-06 |
| RYBP_HUMAN|Q8N488_T186 | 0.304825173 | 2.51E-04 | 2.77E-03 |
| COBL1_HUMAN|Q53SF7_S294_T298 | 0.304609909 | 1.67E-10 | 6.90E-08 |
| ELMO3_HUMAN|Q96BJ8-3_S5 | 0.30378997 | 7.03E-04 | 2.37E-02 |
| ZC3HD_HUMAN|Q5T200_S1014_S1017 | 0.302386572 | 2.18E-10 | 1.97E-07 |
| GRAP1_HUMAN|Q4V328_S666_S669 | 0.300696741 | 2.00E-06 | 4.20E-05 |
| RTKN_HUMAN|Q9BST9_S520_S529 | 0.299355321 | 2.41E-10 | 1.87E-07 |
| ZC3HD_HUMAN|Q5T200_S1014_S1017 | 0.298574958 | 1.78E-10 | 3.21E-07 |
| CE170_HUMAN|Q5SW79_S633_S636 | 0.297422098 | 4.13E-08 | 2.35E-06 |
| NUP53_HUMAN|Q8NFH5_S105 | 0.295292757 | 4.27E-10 | 6.09E-08 |
| K2C6A_HUMAN|P02538_S37 | 0.294681889 | 2.74E-06 | 5.59E-05 |
| COBL1_HUMAN|Q53SF7_S294_T298 | 0.293778882 | 9.52E-11 | 2.57E-08 |
| MK03_HUMAN|P27361_T198_T202 | 0.293022796 | 9.94E-11 | 2.57E-08 |
| SMCR8_HUMAN|Q8TEV9_S417 | 0.292536395 | 6.32E-06 | 1.06E-04 |
| K2C8_HUMAN|P05787_Y25 | 0.291143371 | 4.82E-06 | 8.40E-05 |
| LMNA_HUMAN|P02545_S618 | 0.290007154 | 1.65E-06 | 3.63E-05 |
| CCDC9_HUMAN|Q9Y3X0_S376_S386_T381 | 0.287934879 | 3.43E-12 | 1.86E-09 |
| SUN2_HUMAN|Q9UH99_S112_S113_T107 | 0.287602818 | 1.46E-05 | 6.29E-04 |
| WDR13_HUMAN|Q9H1Z4_S70_S74 | 0.285036127 | 1.59E-12 | 1.08E-09 |
| K2C6B_HUMAN|P04259_S37 | 0.285019905 | 7.01E-10 | 1.17E-07 |
| T22D4_HUMAN|Q9Y3Q8_T167 | 0.284383719 | 5.94E-12 | 2.68E-09 |
| E7ETA6_HUMAN|E7ETA6_T1557 | 0.283673196 | 3.26E-09 | 7.37E-07 |
| NDRG1_HUMAN|Q92597_S326_S330_S333 | 0.281936009 | 9.49E-12 | 3.83E-09 |
| SNUT1_HUMAN|O43290_T430 | 0.281572457 | 4.35E-08 | 1.16E-05 |
| RL1D1_HUMAN|O76021_T465 | 0.281411393 | 2.45E-09 | 1.90E-06 |
| PKP3_HUMAN|Q9Y446_S223 | 0.281406209 | 5.70E-06 | 9.65E-05 |
| NIBL1_HUMAN|Q96TA1_S641_S646 | 0.281250873 | 5.73E-09 | 9.71E-07 |
| LMNA_HUMAN|P02545_S22_T19 | 0.281158089 | 3.29E-07 | 4.81E-05 |

| | | | |
|---|---|---|---|
| RTKN_HUMAN|Q9BST9_S520_S529 | 0.279585898 | 8.84E-11 | 2.20E-08 |
| ENAH_HUMAN|Q8N8S7_T489 | 0.278544945 | 9.01E-07 | 4.71E-05 |
| SEPT9_HUMAN|Q9UHD8_T38_T42 | 0.277422981 | 2.77E-13 | 1.01E-09 |
| AHNK_HUMAN|Q09666_S5739 | 0.276666547 | 2.67E-10 | 5.37E-08 |
| LASP1_HUMAN|Q14847_T104 | 0.275106831 | 1.95E-13 | 2.64E-10 |
| LPP_HUMAN|Q93052_S609 | 0.274718322 | 1.03E-09 | 9.34E-07 |
| T22D4_HUMAN|Q9Y3Q8_T167 | 0.274423114 | 3.21E-12 | 8.70E-09 |
| GRAP1_HUMAN|Q4V328_S666_S669 | 0.274099107 | 7.20E-07 | 2.01E-05 |
| RU17_HUMAN|P08621_S320 | 0.273794987 | 4.10E-08 | 2.20E-06 |
| K2C5_HUMAN|P13647_S18_S21 | 0.272864694 | 2.01E-08 | 1.41E-06 |
| COHA1_HUMAN|Q9UMD9_S76_S79 | 0.271172164 | 6.14E-08 | 3.03E-06 |
| K2C5_HUMAN|P13647_S26_T19 | 0.270071336 | 3.84E-06 | 6.97E-05 |
| AHNK_HUMAN|Q09666_S5739 | 0.268209254 | 1.69E-10 | 6.90E-08 |
| AHNK_HUMAN|Q09666_S5746_S5752 | 0.267143532 | 2.65E-14 | 7.19E-11 |
| REQU_HUMAN|Q92785_T149 | 0.266593303 | 5.30E-12 | 2.10E-09 |
| TB182_HUMAN|Q9C0C2_T1282 | 0.264468001 | 2.42E-13 | 2.47E-10 |
| T22D4_HUMAN|Q9Y3Q8_T167 | 0.262405091 | 1.51E-12 | 8.16E-10 |
| SRRM2_HUMAN|Q9UQ35_S1152 | 0.26218144 | 3.58E-10 | 6.47E-08 |
| AKTS1_HUMAN|Q96B36_S187 | 0.2620209 | 2.69E-11 | 9.13E-09 |
| K2C6A_HUMAN|P02538_S37 | 0.260791023 | 5.61E-07 | 1.57E-05 |
| ODPA_HUMAN|P08559_S232 | 0.26062459 | 5.74E-06 | 2.12E-04 |
| COHA1_HUMAN|Q9UMD9_S76_S79 | 0.26020314 | 3.70E-08 | 2.03E-06 |
| AHNK_HUMAN|Q09666_S5745_S5749 | 0.259927924 | 2.10E-15 | 1.14E-11 |
| MYPT1_HUMAN|O14974_T696 | 0.25866973 | 1.28E-04 | 1.10E-03 |
| COBL1_HUMAN|Q53SF7_T322 | 0.257841666 | 1.05E-13 | 1.89E-10 |
| AHNK_HUMAN|Q09666_S5739 | 0.257679212 | 9.42E-11 | 2.22E-08 |
| COBL1_HUMAN|Q53SF7_T322 | 0.257562708 | 1.03E-13 | 1.86E-10 |
| SRRM1_HUMAN|Q8IYB3_S738_S743 | 0.256643563 | 2.62E-08 | 1.69E-06 |
| CR025_HUMAN|Q96B23_S143_S145_S147 | 0.255724838 | 9.56E-12 | 5.18E-09 |
| NGAP_HUMAN|Q9UJF2_S946_S954 | 0.251209578 | 4.55E-06 | 1.28E-04 |
| MISP_HUMAN|Q8IVT2_S395_S400 | 0.250190886 | 4.90E-09 | 2.66E-06 |

| | | | |
|---|---|---|---|
| COHA1_HUMAN\|Q9UMD9_S61_S62_S70 | 0.250174235 | 2.52E-08 | 1.65E-06 |
| CCDC9_HUMAN\|Q9Y3X0_S376_S386_T381 | 0.247607516 | 2.73E-13 | 2.47E-10 |
| PSN2_HUMAN\|P49810_S7_T3 | 0.245189703 | 4.74E-07 | 3.42E-05 |
| NAV1_HUMAN\|Q8NEY1_T1006 | 0.244814249 | 2.01E-10 | 3.89E-08 |
| LMNA_HUMAN\|P02545_S22_T19 | 0.242604998 | 5.83E-08 | 3.16E-05 |
| MISP_HUMAN\|Q8IVT2_S395_S400 | 0.241829264 | 3.30E-09 | 2.35E-06 |
| E7ETA6_HUMAN\|E7ETA6_T1557 | 0.241360172 | 2.29E-10 | 4.07E-08 |
| K2C5_HUMAN\|P13647_S18_S21 | 0.23224792 | 2.38E-09 | 2.72E-07 |
| ITA2_HUMAN\|P17301_S1180 | 0.231230462 | 6.07E-13 | 6.47E-10 |
| PKP3_HUMAN\|Q9Y446_S223 | 0.230791466 | 8.34E-07 | 2.24E-05 |
| AKTS1_HUMAN\|Q96B36_T246 | 0.22834345 | 1.35E-11 | 4.87E-09 |
| K2C5_HUMAN\|P13647_S21_T19 | 0.22830472 | 3.61E-10 | 5.43E-08 |
| K2C5_HUMAN\|P13647_S21_T19 | 0.225242124 | 3.01E-10 | 5.62E-08 |
| ZC3HD_HUMAN\|Q5T200_S1014_S1017 | 0.220889879 | 2.08E-12 | 1.02E-09 |
| TB182_HUMAN\|Q9C0C2_T1282 | 0.217649459 | 9.23E-15 | 5.01E-11 |
| COBL1_HUMAN\|Q53SF7_S294_T298 | 0.215807388 | 1.25E-12 | 9.69E-10 |
| CR025_HUMAN\|Q96B23_S143_S145 | 0.215054085 | 1.46E-16 | 7.92E-13 |
| AKTS1_HUMAN\|Q96B36_S187 | 0.207708332 | 8.75E-13 | 7.91E-10 |
| LMNA_HUMAN\|P02545_S22_T19 | 0.207504625 | 1.03E-08 | 5.40E-06 |
| UBXN7_HUMAN\|O94888_S288 | 0.194320698 | 1.37E-08 | 9.32E-07 |
| UBXN7_HUMAN\|O94888_S288 | 0.189651194 | 1.07E-08 | 9.07E-07 |
| SUN2_HUMAN\|Q9UH99_S112_S113_T107 | 0.178499971 | 9.91E-08 | 2.24E-05 |
| LMNA_HUMAN\|P02545_S22_T19 | 0.161996124 | 8.24E-10 | 4.47E-07 |
| SNUT1_HUMAN\|O43290_T430 | 0.127101922 | 9.89E-12 | 3.83E-09 |
| SNUT1_HUMAN\|O43290_T430 | 0.112566625 | 3.46E-12 | 1.56E-09 |

**Supplementary Table 5. Patient characteristics for survival analysis (TriNetX)**

| | Baseline | | | Propensity Score Matched | | |
|---|---|---|---|---|---|---|
| **Demographic** | *H1-Antihistamine n=1629* | *H2-Antihistamine n=5729* | *P-Value* | *H1-Antihistamine n=1619* | *H2-Antihistamine n=1619* | *P-Value* |
| Age at Index (Mean +/- SD) | 64.2 +/- 13.8 | 64.5 +/- 12.2 | | 64.3 +/- 13.8 | 63.7 +/- 12.8 | |
| Male | 958 (58.81%) | 3901 (68.09%) | 2.93E-12 | 958 (59.17%) | 965 (59.61%) | 0.802 |
| Female | 651(39.96%) | 1703 (29.73%) | 5.44E-15 | 641 (39.59%) | 629 (38.85%) | 0.666 |
| White | 962 (59.06%) | 2980 (52.02%) | 5.00E-07 | 952 (58.80%) | 941 (58.12%) | 0.695 |
| Black or African American | 62 (3.81%) | 231 (4.03%) | 0.680 | 62 (3.85%) | 51 (3.25%) | 0.292 |
| **Diagnosis** | | | | | | |
| Hypertensive diseases | 643 (39.47%) | 2076 (36.24%) | 0.017 | 635 (39.22%) | 617 (38.11%) | 0.516 |
| Diseases of the skin and subcutaneous tissue | 376 (23.08%) | 916 (15.99%) | 3.15E-11 | 366 (22.61%) | 373 (23.04%) | 0.769 |
| Diseases of esophagus, stomach and duodenum | 325 (19.95%) | 1226 (21.40%) | 0.206 | 323 (19.95%) | 301 (18.59%) | 0.327 |
| Diabetes mellitus | 319 (19.58%) | 992 (17.32%) | 0.035 | 314 (19.40) | 332 (20.51) | 0.429 |
| Ischemic heart diseases | 231 (14.18%) | 815 (14.23%) | 0.963 | 229 (14.15%) | 252 (15.57%) | 0.256 |
| Acute kidney failure and chronic kidney disease | 225 (13.81%) | 838 (14.63) | 0.409 | 225 (13.90%) | 241 (14.89%) | 0.423 |
| Influenza and pneumonia | 110 (6.75%) | 416 (7.26%) | 0.482 | 110 (6.79%) | 117 (7.23%) | 0.630 |
| Cerebrovascular diseases | 100 (6.14%) | 481 (8.40%) | 0.003 | 100 (6.18%) | 99 (6.12%) | 0.942 |
| Pulmonary heart disease and diseases of pulmonary circulation | 55 (3.38%) | 197 (3.44%) | 0.903 | 55 (3.40%) | 58 (3.85%) | 0.774 |
| Chronic rheumatic heart diseases | 30 (1.84%) | 115 (2.01%) | 0.671 | 30 (1.85%) | 26 (1.61%) | 0.590 |
| **Medication** | | | | | | |
| Dermatological Agents | 1142 (70.10%) | 3915 (68.34%) | 0.174 | 1133 (69.98%) | 1138 (70.29%) | 0.848 |
| Antilipemic Agents | 561 (34.44%) | 1459 (25.47%) | 8.11E-13 | 551 (34.03%) | 553 (34.16%) | 0.941 |
| Beta Blockers/Related | 429 (26.34%) | 1393 (24.32%) | 0.096 | 427 (26.37%) | 419 (25.88%) | 0.749 |
| Antianginals | 101 (6.20%) | 485 (8.47%) | 0.003 | 101 (6.24%) | 120 (7.41%) | 0.185 |

**Supplementary Table 6. Patient characteristics for immunotherapy for survival analysis (TriNetX)**

| | Baseline | | | Propensity Score Matched | | |
|---|---|---|---|---|---|---|
| **Demographic** | *H1-Antihistamine n=389* | *H2-Antihistamine n=4464* | *P-Value* | *H1-Antihistamine n=386* | *H2-Antihistamine n=386* | *P-Value* |
| Age at Index (Mean +/- SD) | 67.37 +/- 12.6 | 66 +/- 12.1 | | 67.38 +/- 12.6 | 67.96 +/- 12.2 | |
| Male | 269 (69.15%) | 3157 (70.72%) | 0.515 | 269 (69.69%) | 260 (67.36%) | 0.486 |
| Female | 123 (31.62%) | 1159 (25.96%) | 0.015 | 122 (31.61%) | 129 (33.42%) | 0.591 |
| White | 257 (66.07%) | 2778 (62.23%) | 0.134 | 255 (66.06%) | 246 (63.73%) | 0.497 |
| Black or African American | 14 (3.60%) | 239 (5.35%) | 0.135 | 14 (3.63%) | 15 (3.89%) | 0.850 |
| **Diagnosis** | | | | | | |
| Malignant neoplasms of ill-defined, other secondary and unspecified sites | 274 (70.44%) | 3587 (80.35%) | 3.29182E-06 | 272 (70.47%) | 272 (70.47%) | 1.000 |
| Hypertensive diseases | 221 (56.81%) | 2495 (55.89%) | 0.726 | 219 (56.74%) | 220 (57.00%) | 0.942 |
| Diseases of the skin and subcutaneous tissue | 188 (48.33%) | 1945 (43.57%) | 0.070 | 186 (48.19%) | 181 (46.89%) | 0.719 |
| Diseases of esophagus, stomach and duodenum | 162 (41.65%) | 1759 (39.40%) | 0.386 | 161 (41.71%) | 166 (43.01%) | 0.716 |
| Malignant neoplasms of lip, oral cavity and pharynx | 160 (41.13%) | 2327 (52.13%) | 3.15868E-05 | 159 (41.19%) | 162 (41.97%) | 0.827 |
| Melanoma and other malignant neoplasms of skin | 150 (38.56%) | 1912 (42.83%) | 0.102 | 150 (38.86%) | 146 (37.82%) | 0.767 |
| Malignant neoplasms of respiratory and intrathoracic organs | 128 (32.91%) | 1548 (34.68%) | 0.481 | 126 (32.64%) | 136 (35.23%) | 0.447 |
| Acute kidney failure and chronic kidney disease | 112 (28.79%) | 942 (21.10%) | 0.0004 | 110 (28.50%) | 106 (27.46%) | 0.748 |
| Ischemic heart diseases | 109 (28.02%) | 1158 (25.94%) | 0.370 | 109 (28.24%) | 117 (30.31%) | 0.527 |
| Diabetes mellitus | 95 (24.42%) | 915 (20.50%) | 0.067 | 95 (24.61%) | 89 (23.06%) | 0.612 |
| Chronic rheumatic heart diseases | 77 (19.79%) | 720 (16.13%) | 0.061 | 76 (19.69%) | 80 (20.73%) | 0.720 |
| Influenza and pneumonia | 56 (14.40%) | 626 (14.02%) | 0.839 | 56 (14.51%) | 68 (17.62%) | 0.240 |
| Cerebrovascular diseases | 42 (10.80%) | 372 (8.33%) | 0.095 | 42 (10.88%) | 44 (11.40%) | 0.819 |
| Pulmonary heart disease and diseases of pulmonary circulation | 25 (6.43%) | 325 (7.28%) | 0.532 | 25 (6.48%) | 21 (5.44%) | 0.543 |
| Malignant neoplasms of lymphoid, hematopoietic and related tissue | 11 (2.83%) | 126 (2.82%) | 0.995 | 11 (2.85%) | 13 (3.37%) | 0.678 |
| **Medication/Procedure** | | | | | | |

| | | | | | | |
|---|---|---|---|---|---|---|
| Dermatological Agents | 373 (95.89%) | 4197 (94.02%) | 0.132 | 370 (95.86%) | 372 (96.37%) | 0.710 |
| Antilipemic Agents | 161 (41.39%) | 1458 (32.66%) | 0.000 | 160 (41.45%) | 167 (43.26%) | 0.610 |
| Beta Blockers/Related | 147 (37.79%) | 1853 (41.51%) | 0.153 | 147 (38.08%) | 140 (36.27%) | 0.602 |
| Antianginals | 38 (9.77%) | 346 (7.75%) | 0.157 | 38 (9.85%) | 34 (8.81%) | 0.621 |
| Surgery | 304 (78.15%) | 4076 (91.31%) | 4.75E-17 | 304 (78.76%) | 310 (80.31%) | 0.592 |

**Supplementary Table 7. Patient characteristics for survival analysis (Atropos)**

| | Baseline | | Propensity Score Matched | |
|---|---|---|---|---|
| **Demographic** | *H1-Antihistamine* | *H2-Antihistamine* | *H1-Antihistamine* | *H2-Antihistamine* |
| N | 3493 | 4173 | 2449 | 2449 |
| Female | 1839 (52.6%) | 1853 (44.4%) | 1168 (47.7%) | 1203 (49.1%) |
| Male | 1654 (47.4%) | 2320 (55.6%) | 1281 (52.3%) | 1246 (50.9%) |
| Mean age (sd) | 62.6 (11.2) | 63.1 (10.5) | 63.1 (11.2) | 63.1 (10.6) |
| White | 1871 (53.6%) | 2242 (53.7%) | 1351 (55.2%) | 1317 (53.8%) |
| Other | 1125 (32.2%) | 1419 (34%) | 793 (32.4%) | 831 (33.9%) |
| Black | 415 (11.9%) | 407 (9.8%) | 243 (9.9%) | 243 (9.9%) |
| Asian | 82 (2.3%) | 105 (2.5%) | 62 (2.5%) | 58 (2.4%) |
| Hispanic | 212 (6.1%) | 194 (4.6%) | 139 (5.7%) | 115 (4.7%) |
| **Index Year** | | | | |
| Mean pre-index days (sd) | 5384.2 (4216.3) | 5683.9 (4845.1) | 5695.8 (4400.7) | 5851.1 (4813.1) |
| Mean followup days (sd) | 1093.9 (1079) | 1011.8 (932.6) | 1069.3 (1043.9) | 1040.2 (973.9) |
| Lost to followup | 0 (0%) | 0 (0%) | 0 (0%) | 0 (0%) |
| Number of encounters (sd) | 4.2 (4) | 5.5 (4.8) | 4.3 (4.2) | 4.1 (4.1) |
| **Catagory/Medication** | | | | |
| CPT (sd) | 50.3 (282.3) | 97.1 (364.4) | 58.9 (332.6) | 44.1 (205.9) |
| ICD9 (sd) | 5.1 (23.2) | 3.4 (13.6) | 3.8 (22.6) | 3.8 (14.6) |
| ICD10 (sd) | 17.8 (48.7) | 33.2 (79.1) | 20.9 (53.8) | 17.6 (56.5) |
| RX (sd) | 189.8 (772.6) | 412.5 (1211.7) | 203.9 (901) | 180.7 (789.8) |
| **Diagnosis** | | | | |
| Comorbidity score (sd) | 5.5 (2.6) | 6 (2.8) | 5.7 (2.7) | 5.7 (2.6) |
| Malignancy | 3493 (100%) | 4173 (100%) | 2449 (100%) | 2449 (100%) |
| Diabetes | 953 (27.28%) | 1203 (28.83%) | 708 (28.91%) | 644 (26.3%) |
| Diabetes with Complications | 390 (11.17%) | 581 (13.92%) | 309 (12.62%) | 290 (11.84%) |

| | | | | |
|---|---|---|---|---|
| Congestive Heart Failure | 290 (8.3%) | 551 (13.2%) | 226 (9.23%) | 271 (11.07%) |
| Myocardial Infarction | 160 (4.58%) | 406 (9.73%) | 138 (5.63%) | 185 (7.55%) |
| Peripheral Vascular Disease | 398 (11.39%) | 815 (19.53%) | 313 (12.78%) | 399 (16.29%) |
| Chronic Pulmonary Disease | 1140 (32.64%) | 1284 (30.77%) | 848 (34.63%) | 708 (28.91%) |
| Cerebrovascular Disease | 383 (10.96%) | 634 (15.19%) | 307 (12.54%) | 317 (12.94%) |
| Dementia | 44 (1.26%) | 63 (1.51%) | 38 (1.55%) | 35 (1.43%) |
| Hemiparaplegia | 41 (1.17%) | 94 (2.25%) | 34 (1.39%) | 37 (1.51%) |
| Mild Liver Disease | 432 (12.37%) | 789 (18.91%) | 344 (14.05%) | 367 (14.99%) |
| Severe Liver Disease | 47 (1.35%) | 113 (2.71%) | 41 (1.67%) | 39 (1.59%) |
| Renal Disease | 379 (10.85%) | 636 (15.24%) | 296 (12.09%) | 307 (12.54%) |
| Peptic Ulcer Disease | 81 (2.32%) | 220 (5.27%) | 67 (2.74%) | 110 (4.49%) |
| Rheumatic Disease | 135 (3.86%) | 194 (4.65%) | 98 (4%) | 103 (4.21%) |
| HIV | 46 (1.32%) | 54 (1.29%) | 36 (1.47%) | 22 (0.9%) |
| **Outcomes** | | | | |
| Death | 444 (12.71%) | 665 (15.94%) | 316 (12.9%) | 364 (14.86%) |
